# Illuminating host-mycobacterial interactions with functional genomic screening to inhibit mycobacterial pathogenesis

**DOI:** 10.1101/2020.03.30.016139

**Authors:** Yong Lai, Gregory H. Babunovic, Liang Cui, Peter C. Dedon, John G. Doench, Sarah M. Fortune, Timothy K. Lu

**Affiliations:** Antimicrobial Resistance Interdisciplinary Research Group, Singapore-MIT Alliance for Research and Technology, 138602, Singapore; Synthetic Biology Group, MIT Synthetic Biology Center, Massachusetts Institute of Technology (MIT), Cambridge, MA 02139, USA; Research Laboratory of Electronics, MIT, Cambridge, MA 02139, USA; Department of Immunology and Infectious Diseases, Harvard T.H. Chan School of Public Health, Boston, MA 02115, USA; Ragon Institute of MGH, MIT, and Harvard, Cambridge, MA 02139, USA; Broad Institute, Cambridge, MA 02139, USA; Department of Electrical Engineering and Computer Science, MIT, Cambridge, MA 02139, USA; Harvard-MIT Division of Health Sciences and Technology, Cambridge, MA 02139, USA; Department of Biological Engineering, MIT, Cambridge, MA 02139, USA.

**Keywords:** Host-pathogen interaction, tuberculosis, mycobacteria, CRISPR screen, type I interferon signaling, AHR signaling, oxylipins, host-directed therapy, cerdulatinib, CH223191

## Abstract

Existing antibiotics are inadequate to defeat tuberculosis (TB), a leading cause of death worldwide. We sought potential targets for host-directed therapies (HDTs) by investigating the host immune response to mycobacterial infection. We used CRISPR/Cas9-mediated high-throughput genetic screens to identify perturbations that improve the survival of human phagocytic cells infected with *Mycobacterium bovis* BCG (Bacillus Calmette-Guérin), as a proxy for *Mycobacterium tuberculosis* (Mtb). Many of these perturbations constrained the growth of intracellular mycobacteria. We identified over 100 genes associated with diverse biological pathways as potential HDT targets. We validated key components of the type I interferon and aryl hydrocarbon receptor signaling pathways that respond to the small-molecule inhibitors cerdulatinib and CH223191, respectively; these inhibitors enhanced human macrophage survival and limited the intracellular growth of Mtb. Thus, high-throughput functional genomic screens can elucidate highly complex host-pathogen interactions and serve to identify HDTs with the potential to improve TB treatment.

## INTRODUCTION

In 2018, about 10 million people contracted tuberculosis (TB); 1.5 million were killed by the disease globally (World Health Organization, 2019). Multidrug resistance contributes significantly to this epidemic, and current treatments are insufficient (World Health Organization, 2019). Understanding the dynamic interactions between the host immune response and the intracellular pathogen *Mycobacterium tuberculosis* (Mtb) is crucial for controlling TB.

As sentinels of the host innate immune system, macrophages recognize, ingest, and kill invading pathogens. In latent TB, macrophages limit pathogen replication by triggering granuloma formation (Pagan and Ramakrishnan, 2018). Mtb evades immune surveillance, persisting and replicating inside macrophages and manipulating host-pathogen interactions (Guler and Brombacher, 2015).

Surprisingly, only 5 to 10% of individuals infected with Mtb develop active TB disease (World Health Organization, 2019), so a healthy host immune system can be sufficient to control Mtb infection. Moreover, Mtb infection and pathogenesis can be controlled by drugs targeting host signaling pathways (Hawn et al., 2013); e.g., the Bcr-Abl tyrosine kinase inhibitor Gleevec (imatinib) (Napier et al., 2011), the AMPK (AMP-activated protein kinase)-activating anti-diabetic drug metformin (Singhal et al., 2014), and the EGFR (epidermal growth factor receptor) inhibitors gefitinib and lapatinib (Stanley et al., 2014). The identification of therapeutic targets in distinct signaling pathways suggests the importance of comprehensively capturing host responses as well as the potential benefit of host-directed therapies (HDTs) for TB treatment.

High-throughput unbiased genome-wide screening can be used to systematically sort through complex host-pathogen interactions to identify host perturbations that block bacterial pathogenesis. RNA interference screens have been utilized to study mycobacterial infection (Kumar et al., 2010; Li et al., 2016; Philips et al., 2005). CRISPR-Cas9-mediated genome-wide screens, displaying high efficiency and minimal off-target effects (Shalem et al., 2014; Wang et al., 2014), revealed host cell responses to non-mycobacterial virulence factors (Blondel et al., 2016; Chang et al., 2019; Pacheco et al., 2018; Parnas et al., 2015; Tao et al., 2016), and a CRISPR interference (CRISPRi)-mediated repression library was used to elucidate the function of essential genes and long noncoding RNAs in host cells responding to cholera-diphtheria toxin (Gilbert et al., 2014).

## RESULTS

### High-throughput strategy to identify host determinants

Mycobacterial virulence can induce necrotic host cell death (Lee et al., 2011; Mahamed et al., 2017); therefore, we used CRISPR-Cas9 knockout and CRISPRi screens to identify host genes whose knockout or knockdown improved host cell survival post-infection with mycobacteria (**Figure 1A**). In order to reduce false positive screen hits, we maximized host-pathogen interactions (i.e., maximized the number of infected host cells) for subsequent high-throughput genetic screening. We first infected human monocytic THP-1 cells with *Mycobacterium bovis* Bacillus Calmette-Guérin (BCG), assessed THP-1-mediated phagocytosis of *M. bovis* BCG with induced green fluorescence (*map24*::GFP), and measured host cell viability post-infection (**Figures 1B** and **S1**). Three rounds of *M. bovis* BCG infection significantly induced host cell death, indicating a selective pressure strong enough for genetic screens (**Figure 1C**).

**Figure 1.**
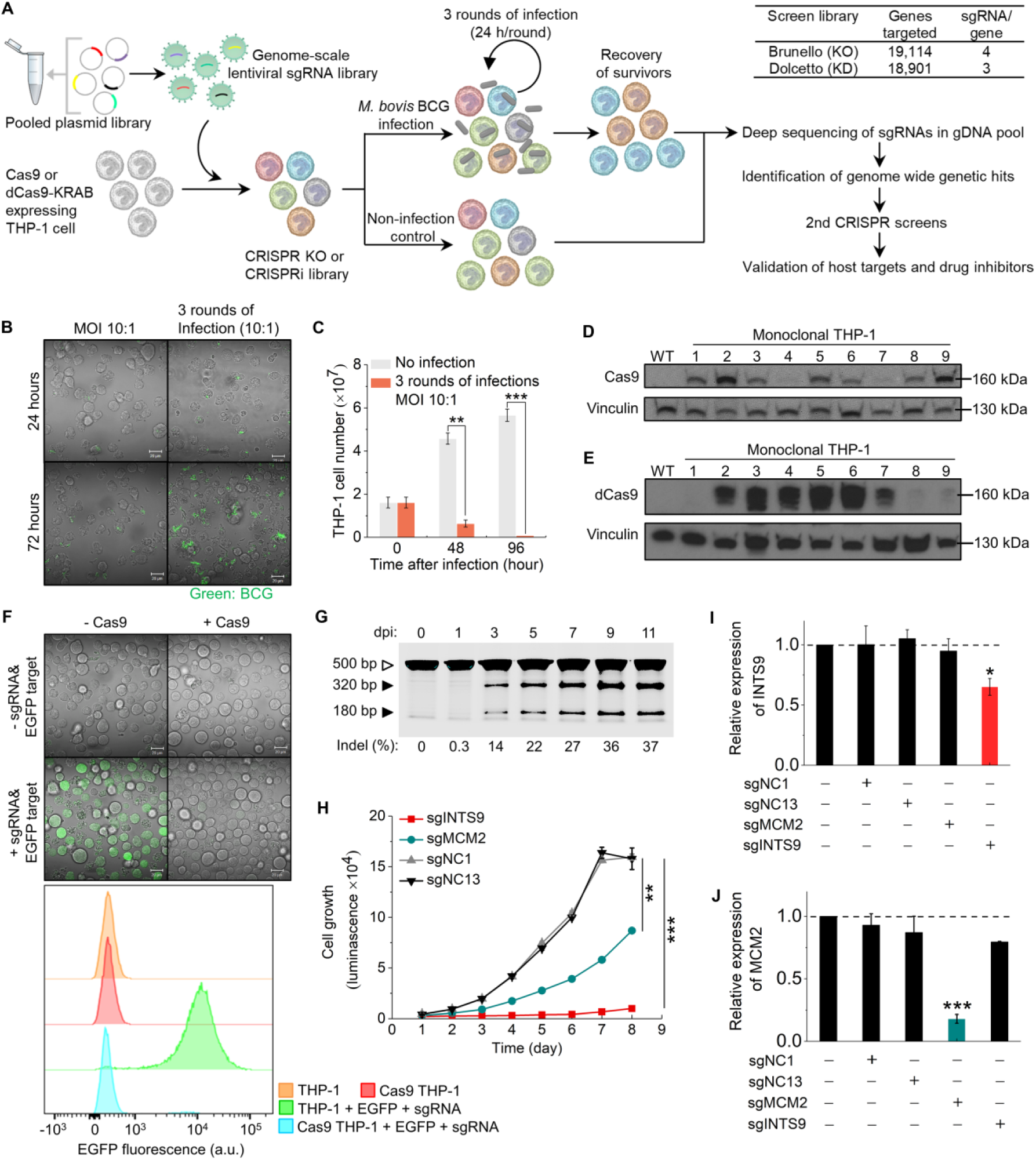
A pooled approach for CRISPR knockout and CRISPRi screening in human THP-1 cells. (A) Strategy for preparing CRISPR libraries and performing genetic screens. (B) THP-1-mediated phagocytosis of *M. bovis* BCG after 3 rounds of infection (MOI 10:1) with induced green fluorescence (*map24*::GFP). (C) Viability of host cells after three rounds of *M. bovis* BCG infection. (D) and (E) Expression of Cas9 and dCas9-KRAB in 9 randomly selected monoclonal THP-1 cells. Wild-type THP-1 cells were used as negative control. Vinculin was used as a loading control. (F) An sgRNA for EGFP was introduced in both wild-type and Cas9-expressing THP-1 cells using a lentivirus (pXPR-011) that also contains EGFP as a target. (G) Cas9-expressing THP-1 cells were transduced with an sgRNA targeting AAVS1 at a low MOI. Mutations at the AAVS1 locus were detected by SURVEYOR assay. The size of the AAVS1 amplicon is 500 bp. The cleaved product sizes are 320 and 180 bp. (H) Growth measurement associated with sgRNAs targeting INTS9, MCM2, and non-targeting negative controls sgNC1 and sgNC13. (I) and (J) RT-qPCR analysis of INTS9 and MCM2 expression in dCas9-KRAB-expressing THP-1 cells. The values are normalized to GAPDH. Data represent the mean ± SD (n=3) (two-tailed unpaired Student’s *t*-test, * P<0.05 ** P<0.01 *** P<0.001). See also Figure S1 and Table S8.

Next, we prepared CRISPR knockout and CRISPRi THP-1 screening libraries. Starting with heterogeneous polyclonal THP-1 cell populations, we isolated monoclonal blasticidin-resistant cells and cells positive for blue fluorescent protein (BFP), corresponding to Cas9-expressing and dCas9-KRAB-expressing THP-1 cells, respectively. The resulting monoclonal cell lines exhibited different levels of Cas9 and dCas9-KRAB protein expression (**Figures 1D** and **1E**), revealing the importance of isolating monoclonal cell lines from the initial heterogeneous cell populations, so that the cells used for CRISPR screening would have consistent levels of protein expression. To determine the functionality of Cas9 in THP-1 cells, we first tested single-guide RNA (sgRNA) targeting enhanced green fluorescent protein (EGFP). EGFP fluorescence was abolished in 91% of the Cas9-expressing THP-1 cells, indicating EGFP cleavage by Cas9 (**Figure 1F**). Transduction of Cas9-expressing THP-1 cells with lentivirus expressing an sgRNA targeting AAVS1 (adeno-associated virus integration site 1) mediated genomic cleavage at the AAVS1 locus, demonstrating knockout by Cas9 of this THP-1 genomic site (**Figure 1G**). Transduction with sgRNAs targeting genes that affect cell proliferation, such as INTS9 (integrator complex subunit 9) and MCM2 (mini-chromosome maintenance complex component 2), inhibited the growth of dCas9-KRAB-expressing THP-1 cells and consistently repressed the transcription of INTS9 and MCM2, suggesting that dCas9-KRAB can knock down protein expression in THP-1 cells (**Figures 1H-1J**). To conduct genetic screens, we transduced Cas9 and dCas9-KRAB-expressing THP-1 cell lines with lentiviruses encoding sgRNA libraries (see below) and infected pooled libraries with *M. bovis* BCG. Surviving host cells with specific gRNA barcodes were analyzed by next-generation sequencing (**Figure 1A**).

### Genome-wide primary screens for host-mycobacterial interactions

We introduced two genome-wide screen libraries into THP-1 cells expressing Cas9 or dCas9-KRAB. The Brunello CRISPR-Cas9 knockout library contains 76,441 sgRNAs systematically targeting 19,114 distinct human genes with 1,000 non-targeting controls (Doench et al., 2016). The Dolcetto CRISPRi library (Set A) contains 57,050 sgRNAs targeting 18,901 human genes with 500 non-targeting controls (Sanson et al., 2018). Independent biological triplicates of each screen library in uninfected THP-1 cells had uniform and comprehensive sgRNA representation (**Figures S2A-S2C** and **S2G-S2I**). The sgRNA distribution significantly differed between mycobacteria-infected and uninfected cells (**Figures S2D** and **S2J**). Comparing the log2-fold change of sgRNA abundance in infected to uninfected THP-1 cells, we observed a stronger sgRNA-level correlation of biological replicates in CRISPRi screens (Pearson R=0.60 and 0.62; **Figures S2K** and **S2L**) than in CRISPR knockout screens (Pearson R=0.39; **Figures S2E** and **S2F**). Volcano plots of genome-wide screen results (**Figures 2A** and **2B**) showed 141 and 157 genes (FDR<0.1; log2-fold change>1) enriched over the uninfected controls in CRISPR-Cas9 knockout and CRISPRi screens, respectively, with 48 of the same genes enriched in both screens (p-value<2.087E-66) (**Figure 2C**; **Tables S2** and **S3**).

**Figure 2.**
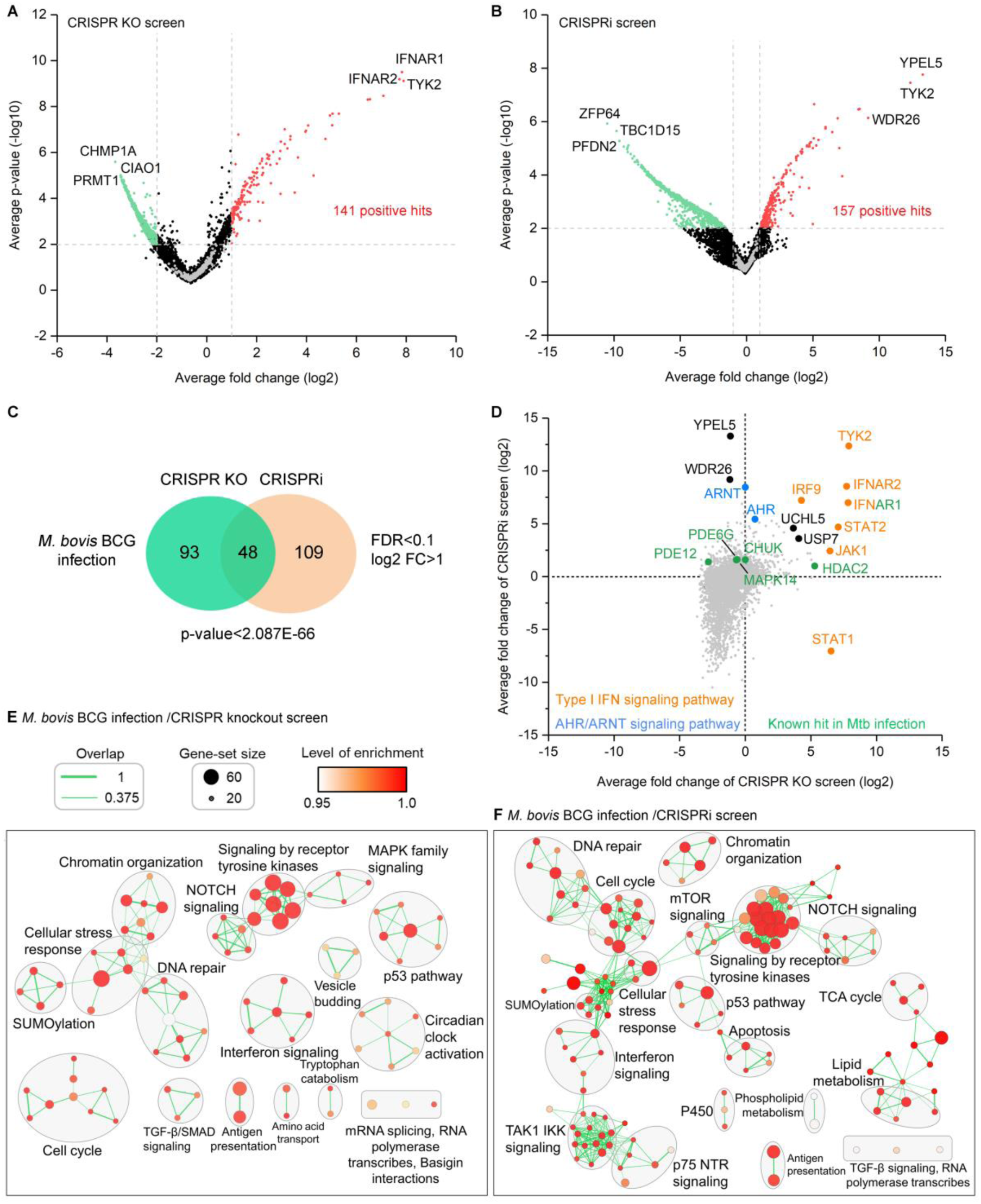
Genome-wide pooled CRISPR knockout and CRISPRi screens to dissect biological pathways in mycobacterial infection. (A) and (B) Volcano plots from CRISPR knockout (A) and CRISPRi (B) screens. For each sgRNA-targeted gene, the x axis shows its enrichment or depletion post-infection, and the y axis shows statistical significance measured by p-value. Positive and negative screen hits are labeled as red and green dots, respectively. Gray dots represent non-targeting controls. For each screen, experiments were carried out in triplicate. (C) Enriched genes in the Venn diagram were filtered with a cut-off of FDR <0.1 and log2-fold change >1 in *M. bovis* BCG infection. The degree of significance of the overlap is given. (D) Gene-centric visualization of average fold change of CRISPR knockout and CRISPRi screens in infected versus non-infected host cells. Selected type I IFN, AHR/ARNT pathway components, and known genetic hits in Mtb infection are highlighted in orange, blue, and green. (E) and (F) Candidate genes identified by CRISPR knockout and CRISPRi screens were functionally categorized to understand the changes in biological functions involved in *M. bovis* BCG infection. Color gradient of nodes represents the enrichment scores of gene-sets. Node size represents the number of genes in the gene set. Edge width represents mutual overlap of genes. See also Figure S2 and Tables S1-S3.

The identification of genes known to contribute to the outcome of Mtb infection supported the reliability of our genome-wide screens (**Figure 2D**). Deletion or inhibition of those genes contributed to host protection against Mtb infection. Deletion of IFNAR1 (interferon alpha and beta receptor subunit 1) in the type I interferon signaling pathway can increase resistance to Mtb in mice (Mayer-Barber et al., 2014), and a proline deletion in IFNAR1 increases resistance to TB in some human populations (Zhang et al., 2018). HDAC (histone deacetylase) regulates the pro-inflammatory response of host cells, and HDAC inhibition restricts intracellular Mtb growth (Seshadri et al., 2017). PDE (phosphodiesterase) inhibitors improve the clearance of Mtb in rabbit lungs, restrict Mtb growth, and increase mouse survival (Maiga et al., 2012; Subbian et al., 2011). Repression of CHUK (conserved helix-loop-helix ubiquitous kinase), identified by a human kinome siRNA screen, decreases intracellular Mtb load (Korbee et al., 2018). Inhibition of p38 MAPK (mitogen activated protein kinase) can restrict Mtb growth in murine macrophages (Stanley et al., 2014). All of these genes were identified by our CRISPR screens as positive genetic hits post-infection.

Many novel host genes were also identified in our screens, including: YPEL5 (yippee-like 5), which may interact with TBK1 (TANK binding kinase 1) (Moreira-Teixeira et al., 2018); WDR26 (WD repeat domain 26), which regulates apoptosis, ubiquitination, signal transduction, and other cellular processes; UCHL5 (ubiquitin C-terminal hydrolase L5), which mediates ubiquitination and protein degradation (Kummari et al., 2015); AHR (aryl hydrocarbon receptor) and ARNT (AHR nuclear translocator), sensors that regulate metabolism and immune functions (Rothhammer and Quintana, 2019); plus many hits without known functions (**Tables S2** and **S3**). Thus, our genome-wide CRISPR-Cas9 knockout and CRISPRi screens identified known and novel host genes that alter macrophage survival after *M. bovis* BCG infection.

We next analyzed biological pathways to identify which knockouts or knockdowns would improve host cell survival in mycobacterial infection. Both CRISPR-Cas9 knockout and CRISPRi screens identified the same pathways, including interferon signaling, DNA repair, chromatin organization, cellular stress response, the p53 pathway, and signaling by receptor tyrosine kinases (**Figures 2E** and **2F**; **Table S1**). The CRISPRi screen also uniquely identified pathways necessary for host cell survival, such as the tricarboxylic acid cycle, mitochondrial biogenesis, and lipid metabolism pathways, supporting the capability of this screen to identify essential host genes that may be missed by CRISPR knockout screens (**Figure 2F**; **Table S1B**).

### Secondary screens for host-mycobacterial interactions

Customized secondary CRISPR knockout and CRISPRi screen libraries were prepared to validate genome-wide screen hits, test known host targets that did not score in the primary screens with either CRISPR knockout or CRISPRi, and allow a clear comparison between CRISPR-Cas9 knockout and CRISPRi screens by making the number of sgRNAs per gene uniform. These libraries each contained 4,740 sgRNAs that target 372 human genes with 10 sgRNAs per gene (**Tables S4** and **S5**). As in the primary screen, pooled secondary screens were based on host cell survival following 3 rounds of *M. bovis* BCG infection. As visualized in volcano plots (**Figures S3A** and **S3B**), CRISPR-Cas9 knockout and CRISPRi screens identified 25 and 26 positively selected genes, respectively, with 8 genes enriched in both screens (FDR<0.05; log2-fold change>0.5; p-value of overlap<9.224E-05; **Figure 3A**). To evaluate screen quality, we calculated a validation rate for genes based on an FDR threshold of <5% in secondary screens; these genes were grouped by their p-value in primary genome-wide screens (Sanson et al., 2018) (**Figure 3B**). Validation rates of genes in secondary screens decreased with increasing p-values in primary CRISPR-Cas9 knockout and CRISPRi screens.

**Figure 3.**
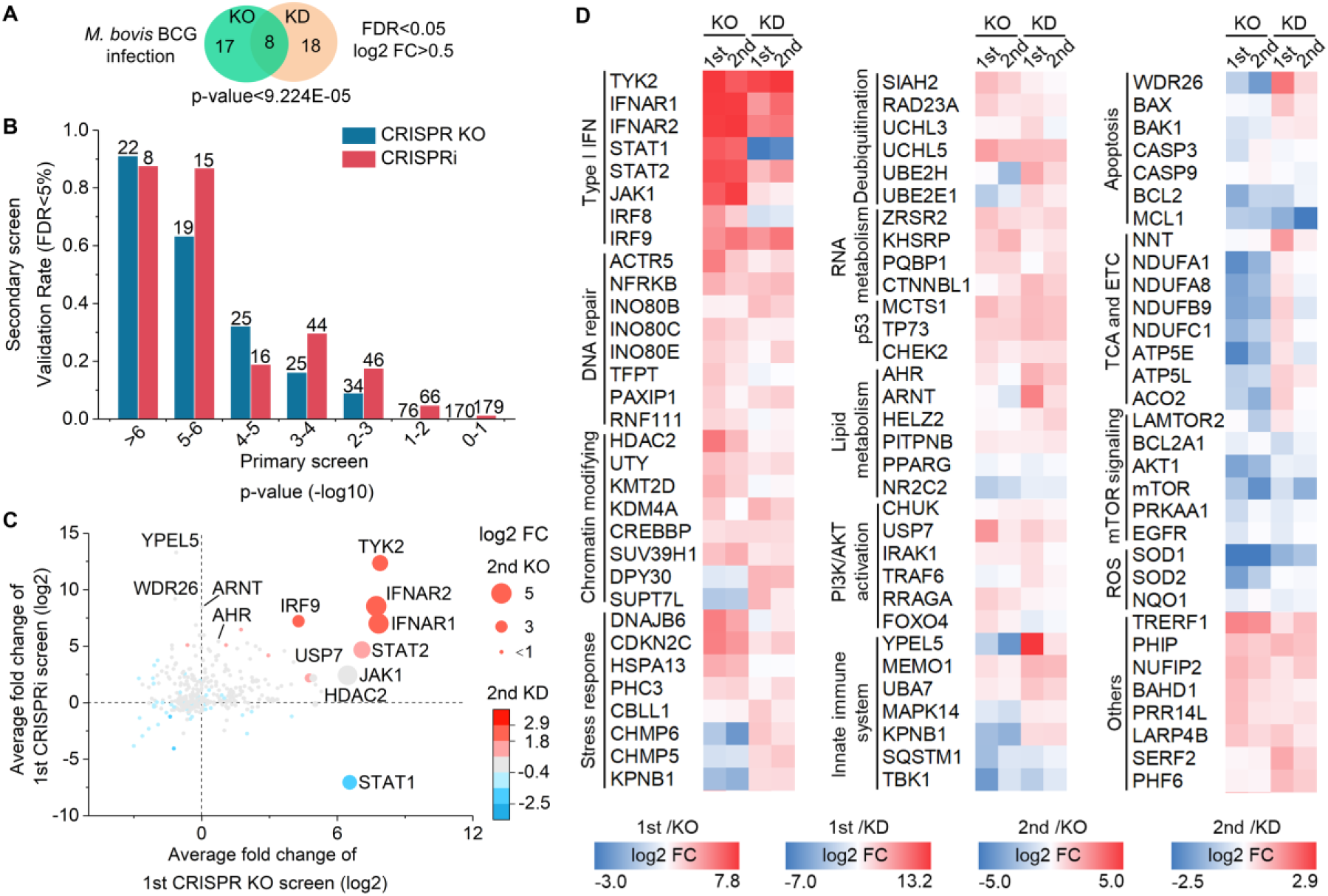
Secondary CRISPR knockout and CRISPRi screens identify host genetic hits in mycobacterial infection. (A) Enriched genes were filtered with a cut-off of FDR <0.05 and log2-fold change >0.5 in *M. bovis* BCG infection. The degree of significance of the overlap is given. (B) Validation rate of genetic hits in secondary screens grouped by their p-value in primary genome-wide screens in *M. bovis* BCG infection. Number of genes per category is indicated. (C) Genetic hits from both primary and secondary screens were ranked by their differential sgRNA abundance between *M. bovis* BCG-infected versus uninfected populations (log2 fold change). (D) Heatmap of screen hits (log2 FC) clustered in different biological pathways in *M. bovis* BCG infection. See also Figure S3 and Tables S4-S7.

The top positively selected genetic hits identified by primary genome-wide screens (i.e., hits that increased the survival of THP-1 cells) were also scored by secondary screens; these included genes involved in type I IFN signaling (TYK2, IFNAR2, IFNAR1, STAT2, JAK1, IRF9), AHR signaling (AHR, ARNT), chromatin modification (HDAC2), PI3K/AKT activation (CHUK, USP7), innate immune functions (YPEL5) and apoptosis (WDR26) (**Figures 3C** and **S3C**; **Tables S6** and **S7**). SOD1 and SOD2, which destroy free superoxide radicals, were identified by both genome-wide and secondary CRISPR screens as negative hits in *M. bovis* BCG infection; i.e., knockdown of these genes reduced host cell survival post-infection (**Figure 3D**). Decreased expression of SOD1 and SOD2 may allow reactive oxygen species to accumulate, which would lead to increased macrophage necrosis (He et al., 2011).

To compare CRISPR knockout to CRISPRi, we examined sgRNA-level correlations of replicates. The correlations of biological triplicates for the CRISPR knockout screens (Pearson R=0.64 and 0.65; **Figures 4A** and **4B**) were not as high as those for the CRISPRi screens (Pearson R=0.88 and 0.78; **Figures 4C** and **4D**). To evaluate screen efficiency, we assessed the separation between positive screen hits (signal) and non-targeting controls (noise) by calculating the signal-to-noise ratio (**Figures 4E** and **4F**). Although CRISPR knockout and CRISPRi screens gave comparable results (S/N=1.1 versus 1.4), hits in the CRISPR knockout screen displayed higher log2-fold changes than those in the CRISPRi screen. Marked changes in gene expression, indicating which genes are most likely to play significant roles in the phenomena being studied, are a critical advantage in genome-wide screens as they can reduce the cost and time spent in confirmatory experiments. In addition, both CRISPR knockout and CRISPRi screens displayed comparable p-values and FDRs (**Figures 4G** and **4H**). Given that specific and important genetic hits, such as AHR and ARNT, were identified by the CRISPRi screens, we consider CRISPRi screens to be a useful complement to genome-wide screens.

**Figure 4.**
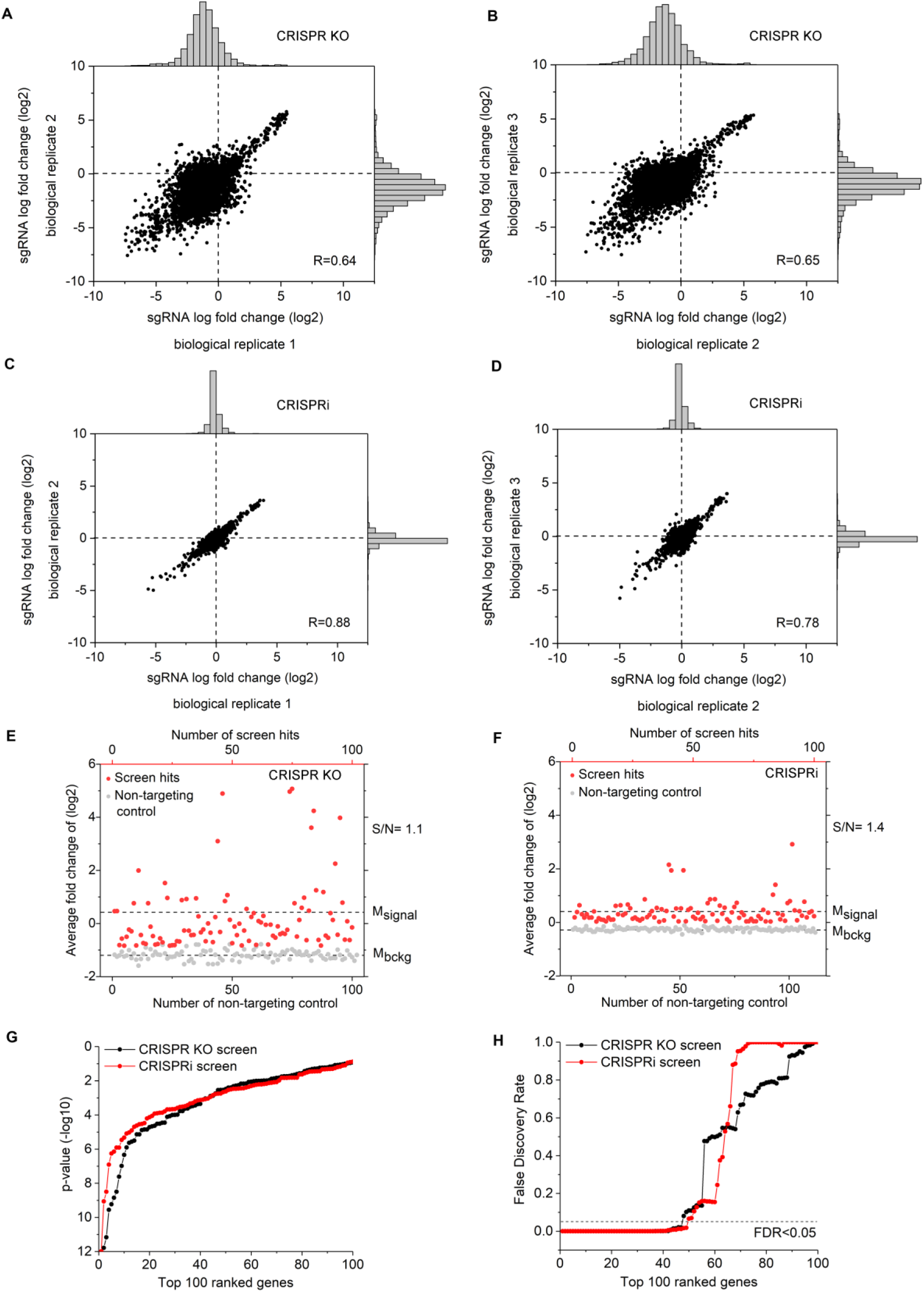
Comparison of secondary CRISPR knockout and CRISPRi screens in mycobacterial infection. (A)-(D) sgRNA-level correlation of replicates in CRISPR knockout screens (A and B) and CRISPRi screens (C and D). Pearson correlation of log2 fold change values between replicates is indicated. (E) and (F) Top 100 positive genetic hits (red) and non-targeting controls (gray) in *M. bovis* BCG infection identified by CRISPR knockout (E) and CRISPRi (F). Broken lines show the means of the hits and non-targeting controls. (G) and (H) p-value (G) and FDR (H) of top 100 positive genetic hits identified by CRISPR knockout and CRISPRi screens in mycobacterial infection.

### Validating hits for host-mycobacterial interactions

Positively selected genetic hits from the aforementioned screens associated with distinct signaling pathways were confirmed by constructing THP-1 cell lines with individual gene knockdowns and testing their phenotypes post-infection. These genes included: TYK2, IFNAR1, JAK1, AHR, CREBBP, UCHL5, TP73, YPEL5, WDR26, TRERF1, and PHIP. The positive correlation between the phenotypes observed in the genome-wide screens and the individual gene validation experiments suggests that repressing each of these screen hits enhanced host cell survival (Pearson R=0.70; **Figure 5A**).

**Figure 5.**
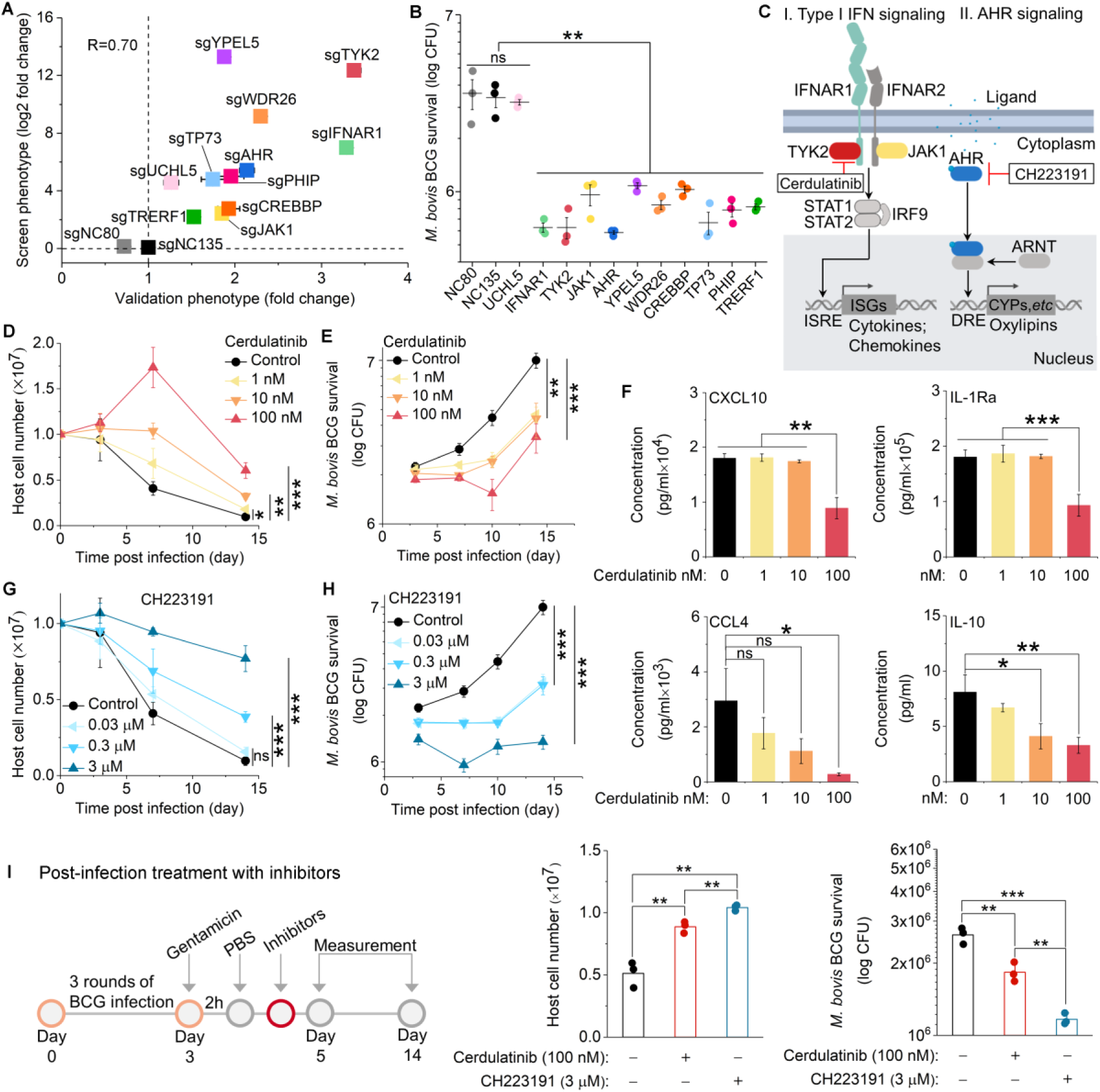
Validation of top positive genetic screen hits and corresponding inhibitors in mycobacterial infection. (A) Correlation of hits between pooled screen and individual validation data. For each hit, the log2 fold change obtained from the genome-wide screening data (Screen phenotype) was plotted against the fold change of cell viability of hits compared to sgNC80 and sgNC135, the non-targeting controls (Validation phenotype). Pearson correlation is indicated. (B) Intracellular *M. bovis* BCG level after infection of THP-1 cells having individual gene knockdowns (day 14). (C) Schematic of biological pathways and corresponding inhibitors associated with top positive genetic hits. (D)-(F) The growth of THP-1 cells (D), intracellular mycobacterial growth (E), and production of infection-induced cytokine and chemokines (F) post-infection in the presence or absence of different concentrations of the TYK2 drug inhibitor cerdulatinib. (G) and (H) The growth of THP-1 cells (G) and intracellular mycobacterial level (H) post-infection in the presence or absence of different concentrations of the AHR inhibitor CH223191. (I) The growth of THP-1 cells and intracellular mycobacterial growth after post-infection treatment with cerdulatinib or CH223191. Data represent the mean ± SD (n = 3). Two-tailed unpaired Student’s *t*-test, * P<0.05 ** P<0.01 *** P<0.001; ns represents not significant. See also Figure S4 and Table S8.

To test whether improved host cell survival was linked to the control of intracellular mycobacterial growth in these individual cell lines, we measured the survival of intracellular bacteria by counting colony-forming units (CFU) post-infection. Repression of gene expression of the same subset of positive screen hits, except for UCHL5, inhibited intracellular *M. bovis* BCG growth (**Figure 5B**).

The type I IFN signaling pathway suppresses the production of IL-1α and IL-1β, two cytokines critical for host resistance to Mtb (Mayer-Barber et al., 2011). Type I IFN signaling was the most highly positively enriched functional pathway in our screens (**Figure 5C**). Six of the top 10 positive hits identified by the genome-wide CRISPR knockout screen and 4 of the top 10 positive hits identified by the CRISPRi screen are key components of this pathway. To characterize the role of these genes in mycobacterial infection, we profiled cytokine and chemokine production in mycobacteria-infected THP-1 cell cultures using a 45-plex Luminex assay. *M. bovis* BCG infection induced the production of many cytokines and chemokines in THP-1 cells (**Figure S4A**). IFNAR1 and TYK2 gene knockdown in THP-1 cells abolished the production of infection-induced cytokines and chemokines, including IL-1Ra, which inhibits the pro-inflammatory function of IL-1α and IL-1β by binding to their receptor, as well as CXCL10 and CCL4, important biomarkers for monitoring TB treatment and disease progression (Sutherland et al., 2016) (**Figure S4C**).

Inhibiting these hits might represent potential HDTs for TB. Small molecule inhibitors that block JAK family members (JAK1, JAK2, JAK3, TYK2) have been developed for immune and inflammatory diseases (Schwartz et al., 2018). Therefore, we tested the function of cerdulatinib, a selective inhibitor of TYK2 (IC_50_=0.5 nM) and JAK1 (IC_50_=12 nM) currently in a phase 2a clinical trial for B-cell malignancies (Coffey et al., 2014). In line with the TYK2 knockdown phenotype, pre-treatment with cerdulatinib enhanced host cell survival and inhibited intracellular *M. bovis* BCG growth dose-dependently (**Figures 5D** and **5E**). Similar results were observed with post-infection treatment (**Figure 5I**). Cerdulatinib (100 nM) also abrogated the production of infection-induced cytokines and chemokines, such as CXCL10, IL-1Ra, CCL4, and IL-10 (**Figure 5F**).

Another intriguing pair of positive genetic hits, AHR and ARNT, which were identified only by the CRISPRi screens, have multiple functions in metabolism and immune responses (Rothhammer and Quintana, 2019). Knockdown of AHR protected host cells and inhibited *M. bovis* BCG growth (**Figures 5A** and **5B**), and the selective AHR inhibitor CH223191 significantly enhanced host cell survival and controlled intracellular *M. bovis* BCG growth dose-dependently (**Figures 5G** and **5H**). Post-infection treatment with CH223191 gave similar results (**Figure 5I**). As a cytosolic transcriptional factor, the AHR/ARNT complex detoxifies xenobiotics by regulating mammalian cytochrome P450 enzymes (CYPs) (**Figure 5C**). To investigate how CH223191 protected host cells, we profiled oxylipin production by LC-MS/MS (**Figures 6A** and **S5**). Oxylipins, which are involved in many immune functions, undergo cytochrome P450-dependent oxidation (Nebert and Karp, 2008). Knockdown of AHR gene expression in THP-1 cells decreased CYP-regulated oxylipin production post-infection (**Figure 6B**). In THP-1 cells without CRISPRi knockdown, CH223191 abrogated the infection-induced and CYP-regulated oxylipin production (**Figures 6D-6F** and **6N**). *M. bovis* BCG infection induced LXA4 (lipoxin A4) production and inhibited the production of PGE2 (prostaglandin E2), a compound critical for blocking Mtb replication and protecting host cells (Chen et al., 2008) (**Figures 6F** and **6G**). Strikingly, CH223191 (3 μM) abolished infection-induced LXA4 production, stimulated PGE2 production, and restored the PGE2/LXA4 ratio to pre-infection levels without affecting arachidonic acid production (**Figures 6C** and **6F-6H**). CH223191 also induced the production of non-CYP-regulated oxylipins 5-HETE, 5-OxoETE, and 11-, 12-, 15-HETE (**Figures 6I-6L**).

**Figure 6.**
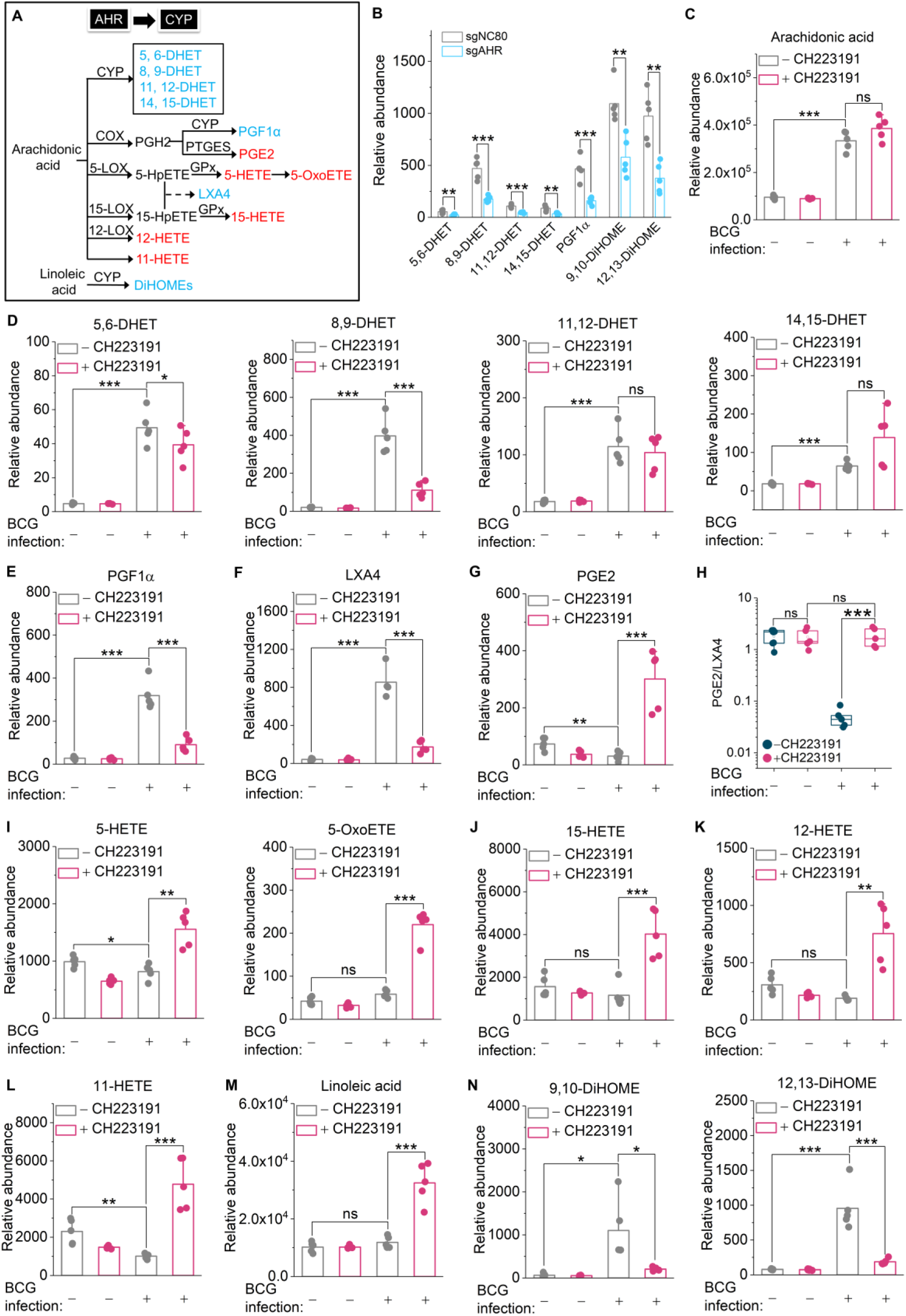
AHR inhibitor regulates host oxylipin metabolism in mycobacterial infection. (A) Diagram of eicosanoids and linoleic acid metabolism in human THP-1 cell line. AHR regulates lipid metabolism by cytochrome P450 (CYP) activation. Metabolites marked in blue represent decreased production in AHR knockdown THP-1 cells or in the presence of CH223191. Metabolites marked in red represent increased production in the presence of CH223191. (B) CYP-regulated oxylipin production in AHR knockdown and non-targeting control THP-1 cells post-infection. (C) Arachidonic acid production. (D) CYP-regulated eicosanoid production. (E) PGF1α production. (F) LXA4 production. (G) PGE2 production. (H) PGE2/LXA4 ratio. (I)-(L) Increased eicosanoid production in the presence of CH223191 post-infection. (M) and (N) Linoleic acid and CYP-regulated metabolites with or without *M. bovis* BCG infection and in the presence or absence of CH223191. Data represent the mean ± SD (n = 5) (two-tailed unpaired Student’s *t*-test, * P<0.05 ** P<0.01 *** P<0.001; ns represents not significant). See also Figure S5.

Host immune responses to *M. bovis* BCG and Mtb are not identical (van der Wel et al., 2007). Our screens with *M. bovis* BCG identified several host targets potentially involved in Mtb infection but required validation with Mtb infection models to establish their clinical relevance. To test the function of TYK2 and AHR inhibitors in Mtb infection, we infected THP-1 cell-Mtb and primary human macrophage-Mtb infection models with Mtb H37Rv-lux, a strain of Mtb expressing a bacterial luciferase (lux) system (Andreu et al., 2010), then added inhibitors and measured host cell and pathogen survival (**Figure 7A**).

**Figure 7.**
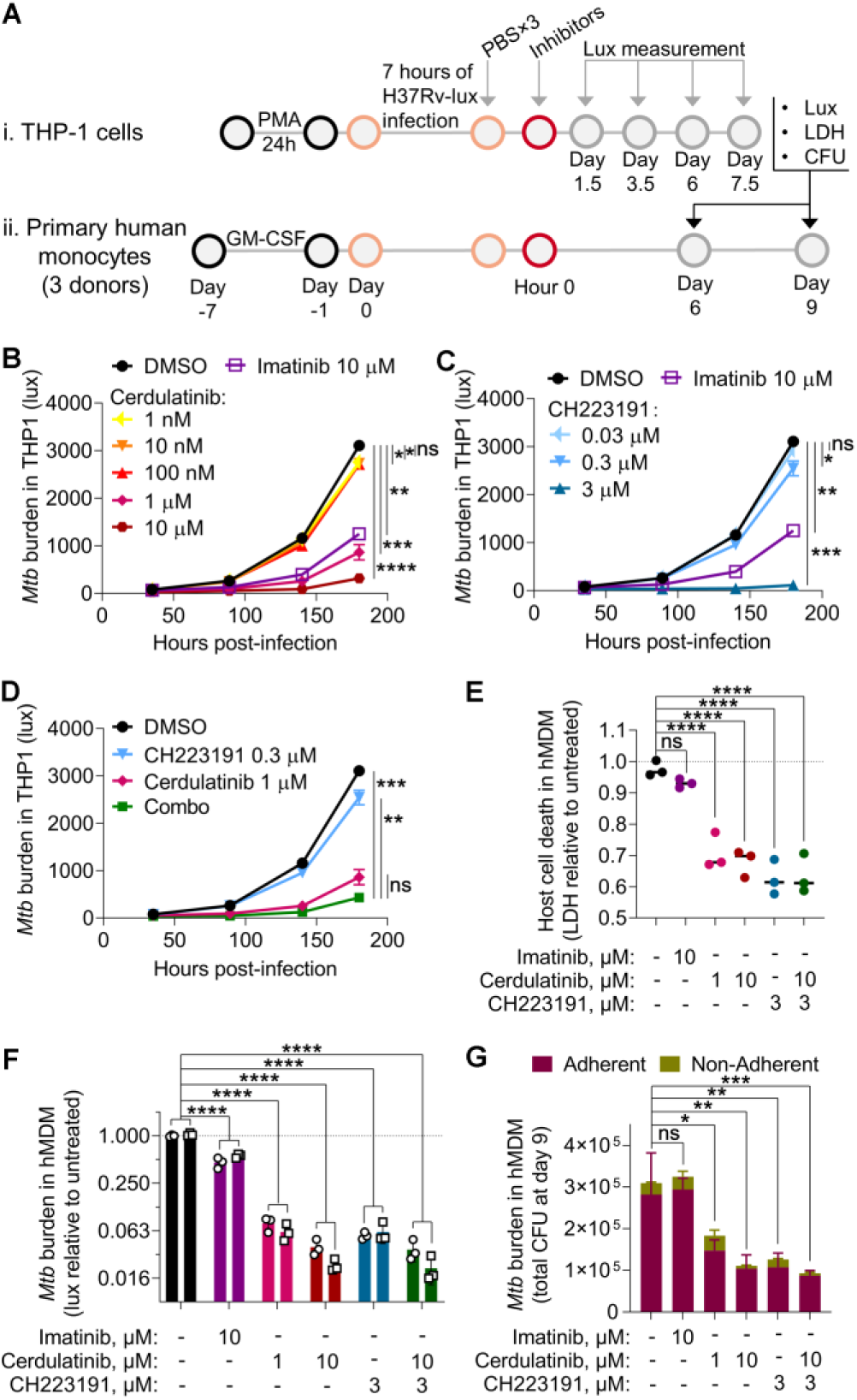
Effects of TYK2 and AHR inhibitors on host cells and *M. tuberculosis* post-infection. (A) Schematic of inhibitor validation in Mtb infection cell models. (B)-(D) Effects of TYK2 inhibitor (B), AHR inhibitor (C), and both inhibitors (“Combo”; D) on growth of *M. tuberculosis* in THP-1 cells, as measured by the luminescence of the Mtb H37Rv-lux strain. (E) Effects of TYK2 and AHR inhibitors on the survival of primary human monocyte-derived macrophages (hMDM) infected with Mtb H37Rv-lux, with death measured at day 9 by lactate dehydrogenase (LDH) release relative to the untreated condition. (F) Effects of TYK2 and AHR inhibitors on the growth of *M. tuberculosis* in hMDM cells, as measured by the luminescence of the H37Rv-lux strain relative to the untreated condition. Circles represent data obtained at day 6; squares represent data obtained at day 9. (G) Effects of TYK2 and AHR inhibitors on the growth of *M. tuberculosis* in hMDM, as measured by total colony-forming units (CFU) of Mtb H37Rv-lux. CFU in the culture supernatant or released after one PBS wash were considered “non-adherent” and marked in deep yellow color, while CFU not released during this wash were considered “adherent” and marked in deep red color. All data represent the mean ± SD (n = 3); imatinib is used at 10 μM as a non-TYK2/AHR inhibitor for comparison in all experiments. Statistics were performed with a two-way results matched ANOVA on time courses (B-D, F) and an ordinary one-way ANOVA for other data (E and G), all followed by Dunnett’s multiple comparisons test. Statistical results are shown only for the final timepoint of growth curves (B-D). Statistics in (G) performed on the sum of both CFU fractions. * P<0.05 ** P<0.01 *** P<0.001 **** P<0.0001

Consistent with results in *M. bovis* BCG infection, both cerdulatinib and CH223191 reduced Mtb growth in THP-1 cells in a dose-dependent fashion (**Figures 7B** and **7C**). While CH223191 was effective against both species at identical concentrations, higher concentrations of cerdulatinib were needed to restrict Mtb growth than for *M. bovis* BCG infection. The drugs were effective in combination, though adding 0.3 μM of CH223191 did not significantly increase the efficacy of 1 μM cerdulatinib (**Figure 7D**). Imatinib, an Mtb HDT, also restricted Mtb growth in this system (**Figures 7B** and **7C**).

In primary human macrophages, cerdulatinib, CH223191, and their combination each significantly reduced host cell death by Mtb as measured by lactate dehydrogenase (LDH) release; these inhibitors outperformed imatinib (**Figure 7E**). The experimental compounds also restricted Mtb growth as measured by bacterial luminescence (**Figure 7F**). Bacterial load was also assessed: cerdulatinib and CH223191 reduced Mtb CFUs (**Figure 7G**). Thus, inhibiting the type I IFN signaling or AHR/ARNT pathway restricts the mycobacterial burden and preserves host cell survival during *M. bovis* BCG and Mtb infection of human monocytic phagocytes, a major cell type infected by Mtb.

## DISCUSSION

Systematic genetic perturbation strategies provide direct causal evidence on how changes in gene expression patterns in macrophages affect intracellular pathogens. To dissect cellular responses to mycobacterial infection, we conducted both genome-wide and focused secondary CRISPR loss-of-function and CRISPRi screens. Unlike other kinds of host genetic screens (Korbee et al., 2018; Kumar et al., 2010; Li et al., 2016; Philips et al., 2005), our high-throughput CRISPR screens require host cell survival and, therefore, directly reveal processes targeted by intracellular mycobacteria in live host cells.

We identified type I IFN signaling as a major driver of cell death in response to mycobacterial infection. This result aligns with those of a number of other studies. Type I IFN signaling correlates with impaired control of Mtb, as assessed by the high transcription levels of IFN-inducible genes in the whole-blood of active TB patients (Berry et al., 2010). Furthermore, deletion of nucleotides TCC in human IFNAR1 (human SNP rs72552343) increases resistance to Mtb (Zhang et al., 2018), as does IFNAR1 deletion in mice (*Ifnar1*−/−) (Mayer-Barber et al., 2014). Type I IFN can exert antimicrobial activity in murine macrophages; however, this effect depends on the production of nitric oxide, which is not consistently associated with a reduced mycobacterial burden in human cells (Banks et al., 2019; Thoma-Uszynski et al., 2001). In our study, perturbations of type I IFN signaling improved host cell survival, decreased bacterial load in both *M. bovis* BCG and Mtb infections, and interfered with the induction of anti-inflammatory cytokines, e.g., IL-10 and IL-1Ra. We infer that the intact type I IFN pathway plays a deleterious role for the host in mycobacterial infection by enhancing pathogen survival. Indeed, type I IFN-driven susceptibility to Mtb can be reversed by repressing IL-1Ra (Ji et al., 2019). Here, more TYK2 inhibitor was required to protect host cells from Mtb than from *M. bovis* BCG, aligning with the magnified type I IFN response to Mtb infection (Novikov et al., 2011; Wassermann et al., 2015); such augmented signaling unsurprisingly mandates more signal transduction inhibition to achieve a similar phenotype.

We observed that AHR and ARNT played a role in host cell death during mycobacterial infection; knockdown or inhibition of AHR also reduced the mycobacterial burden. Decreased PGE2/LXA4 ratios correlate with higher sputum Mtb levels in TB patients (Mayer-Barber et al., 2014), and inhibiting tryptophan metabolism can augment Mtb control (Gautam et al., 2018; Zhang et al., 2013). By inhibiting the activity of AHR, a cytoplasmic sensor that responds to the tryptophan metabolite kynurenine, we could restore pre-infection PGE2/LXA4 ratios and enhance the control of intracellular mycobacterial growth (**Figure 6**). Given the importance of PGE2/LXA4 ratios in host cell defense against Mtb infection (Chen et al., 2008), this rebalancing of oxylipin metabolism by AHR inhibition may contribute to its protective effect. Yet, AHR-deficient mice display increased susceptibility to Mtb infection (Moura-Alves et al., 2014). We did not observe a protective effect of AHR knockout, suggesting that the knockdown or inhibition of AHR sidesteps the deleterious effects of complete knockout. Interestingly, a recent study showed that AHR can bind to rifabutin (RFB), a first-line anti-TB drug, and increase its metabolism, which is likely to decrease the effectiveness of this drug (Puyskens et al., 2019). Inhibition of AHR increased the efficacy of RFB treatment in a zebrafish-*Mycobacterium marinum* infection model (Puyskens et al., 2019). The response of cell networks to slow-growing intracellular pathogens such as Mtb appears, therefore, to be highly complex.

HDTs that boost host immune defense could complement antibiotic treatments and help to achieve the World Health Organization goal of ending the TB epidemic by 2030 (World Health Organization, 2014). We have shown that genome-wide pooled CRISPR-Cas9 knockout and CRISPRi screens can identify key therapeutic targets in immune cell-*Mycobacterium* interactions and that specific small molecule inhibitors and their combinations can re-equilibrate innate immune responses post-infection, potentially preventing the most harmful effects of bacterial pathogenesis. Our top positive genetic hits (TYK2, AHR) indicate potential targets for developing HDTs, and their identification here further illuminates host-pathogen interactions in TB.

## ACKNOWLEDGEMENTS

We thank Karen Pepper for editing the manuscript. This work was supported by the National Institutes of Health (NIH-R01-AI123286-04 to S.M.F.), the United States Defense Threat Reduction Agency (HDTRA1-15-1-0050 to T.K.L.), and the National Research Foundation of Singapore through the Singapore-MIT Alliance for Research and Technology Antimicrobial Resistance IRG (Y.L., L.C., P.C.D., T.K.L.)

## AUTHOR CONTRIBUTIONS

Y.L., G.H.B., L.C., P.C.D., J.G.D., S.M.F., and T.K.L. conceived and designed the research; Y.L. and J.G.D. performed and analyzed genome-wide CRISPR screen experiments; Y.L. and J.G.D. designed, performed and analyzed secondary CRISPR screen experiments; Y.L. conducted validation experiments of genetic hits and small molecule inhibitors in *M. bovis* BCG infection model; G.H.B. conducted validation experiments of small molecule inhibitors in Mtb infection models; Y.L. and L.C. designed and performed oxylipin profile experiments; P.C.D. supervised oxylipin profile experiments. S.M.F. supervised Mtb infection experiments. Y.L. and T.K.L. coordinated the overall research. T.K.L. supervised the overall research. Y.L., G.H.B., L.C., P.C.D., J.G.D., S.M.F., and T.K.L. analyzed the data and wrote the manuscript. All authors discussed the results and reviewed the paper.

## DECLARATION OF INTERESTS

T.K.L. is a co-founder of Senti Biosciences, Synlogic, Engine Biosciences, Tango Therapeutics, Corvium, BiomX, and Eligo Biosciences. T.K.L. also holds financial interests in nest.bio, Ampliphi, IndieBio, MedicusTek, Quark Biosciences, and Personal Genomics.

Y.L. and T.K.L. are co-inventors on a US provisional patent application (no. 62/909727), which is based on discoveries described in this paper.

---

**Figure S1.**
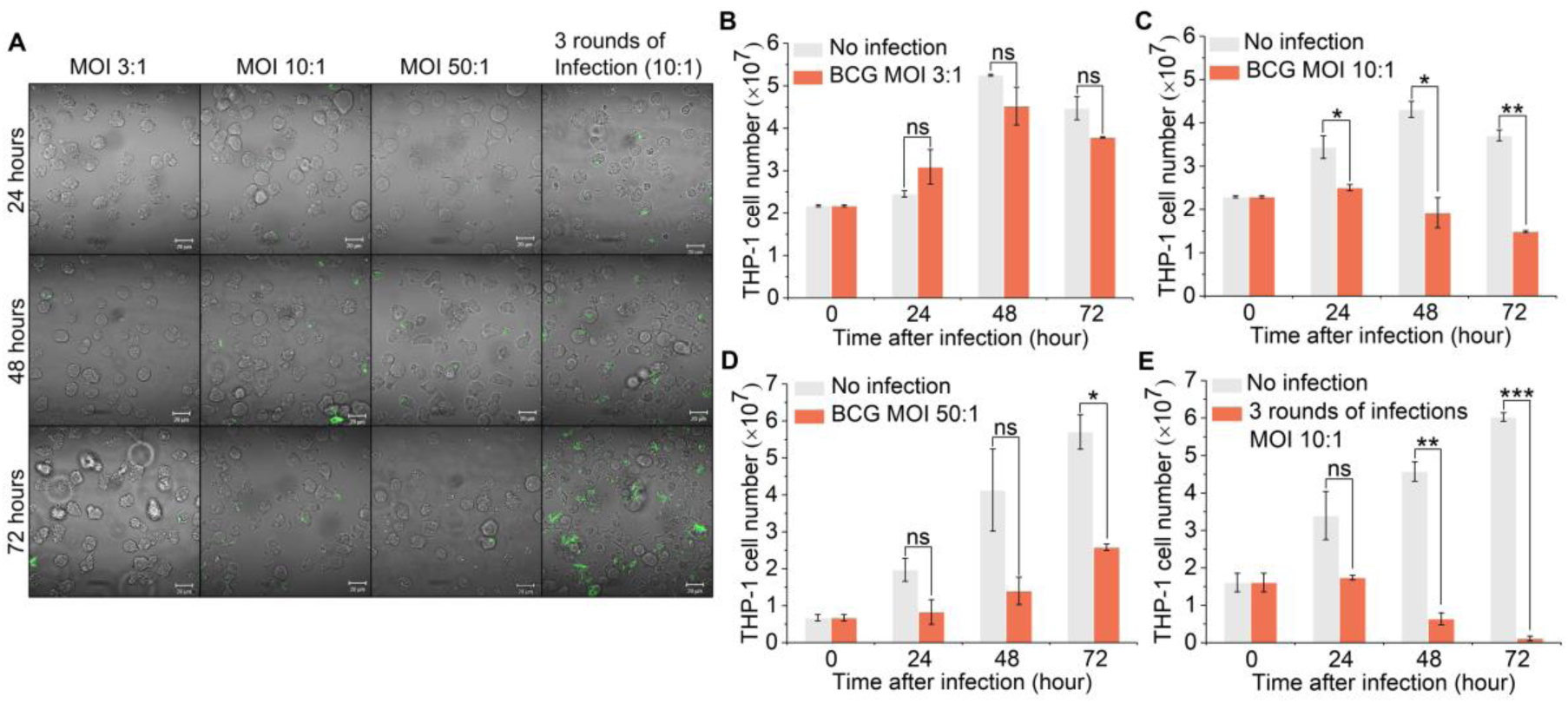
Optimization of conditions for mycobacterial infection of THP-1 cells. Related to Figure 1. (A) Left to right, single round of *M. bovis* BCG infection at MOI (multiplicity of infection, or number of bacterial cells per host cell) of 3:1, 10:1, or 50:1. Far right, three rounds of *M. bovis* BCG infection (0, 18, and 42 h) at an MOI of 10:1. After three rounds of infection over 72 hours, most of the THP-1 cells were infected (right column). (B)-(E) The number of *M. bovis* BCG-infected THP-1 cells after a single round (B, C, and D) or three rounds of infection (E). Data represent mean ± SD (n = 3) (two-tailed unpaired Student’s *t*-test, * P<0.05 ** P<0.01 *** P<0.001; ns represents not significant).

**Figure S2.**
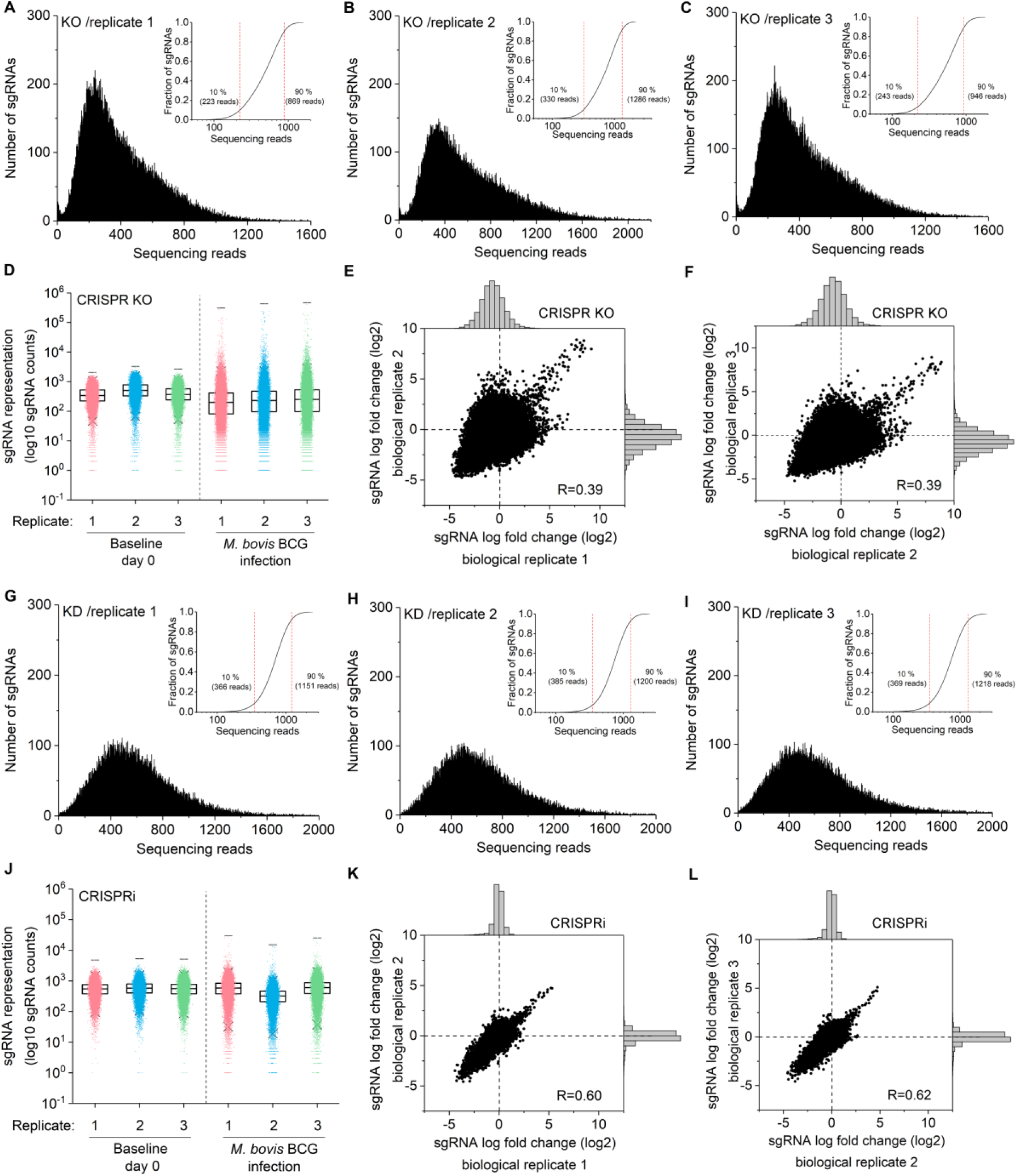
High quality of CRISPR knockout and CRISPRi screening libraries. Related to Figure 2. (A)-(C) Coverage of reads in THP-1 CRISPR knockout libraries. Independent triplicates have <0.0007% undetected guides. The percentage of perfectly matching guides of all triplicates is >86%. The skew ratio of top 10% to bottom 10% of all triplicates is <8.9. (D) Distribution of individual sgRNA in control and *M. bovis* BCG-infected samples after CRISPR knockout screens. (E) and (F) sgRNA-level correlation of replicates in CRISPR knockout screens. R is the Pearson correlation coefficient. (G)-(I) Coverage of reads in CRISPRi libraries. Independent triplicates have <0.00006% undetected guides. The percentage of perfectly matching guides of all triplicates is >80%. The skew ratio of top 10% to bottom 10% of all triplicates is <8.9. (J) Distribution of individual sgRNA in control and *M. bovis* BCG-infected samples after CRISPRi screens. Each point represents individual sgRNAs. Boxes, 25th to 75th percentile; Whiskers, 1st to 99th percentile. (K) and (L) sgRNA-level correlation of replicates in CRISPRi screens. Pearson correlation of log2 fold change values between replicates is indicated.

**Figure S3.**
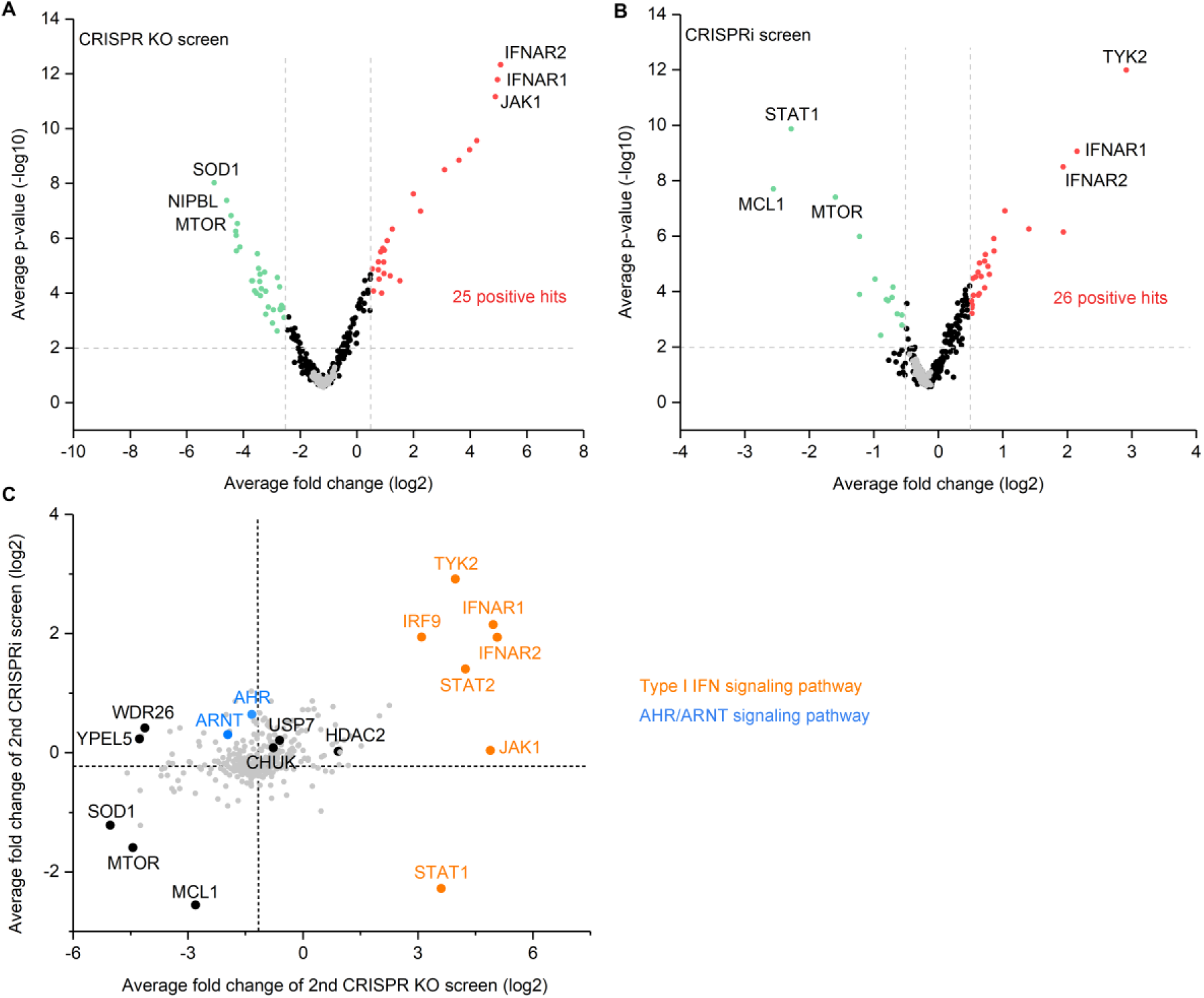
Top genetic hits identified by secondary CRISPR screens in mycobacterial infection. Related to Figure 3. (A) and (B) Volcano plots from secondary CRISPR knockout (A) and CRISPRi (B) screens. For each sgRNA-targeted gene, the x axis shows its enrichment or depletion post-infection, and the y axis shows statistical significance measured by p-value. Positive and negative screen hits are labeled as red and green dots, respectively. Gray dots represent non-targeting controls. For each screen, experiments were carried out in triplicate. (C) Gene-centric visualization of average fold change of secondary screens in infected versus non-infected host cells. Selected type I IFN and AHR/ARNT pathway components are highlighted in orange and blue.

**Figure S4.**
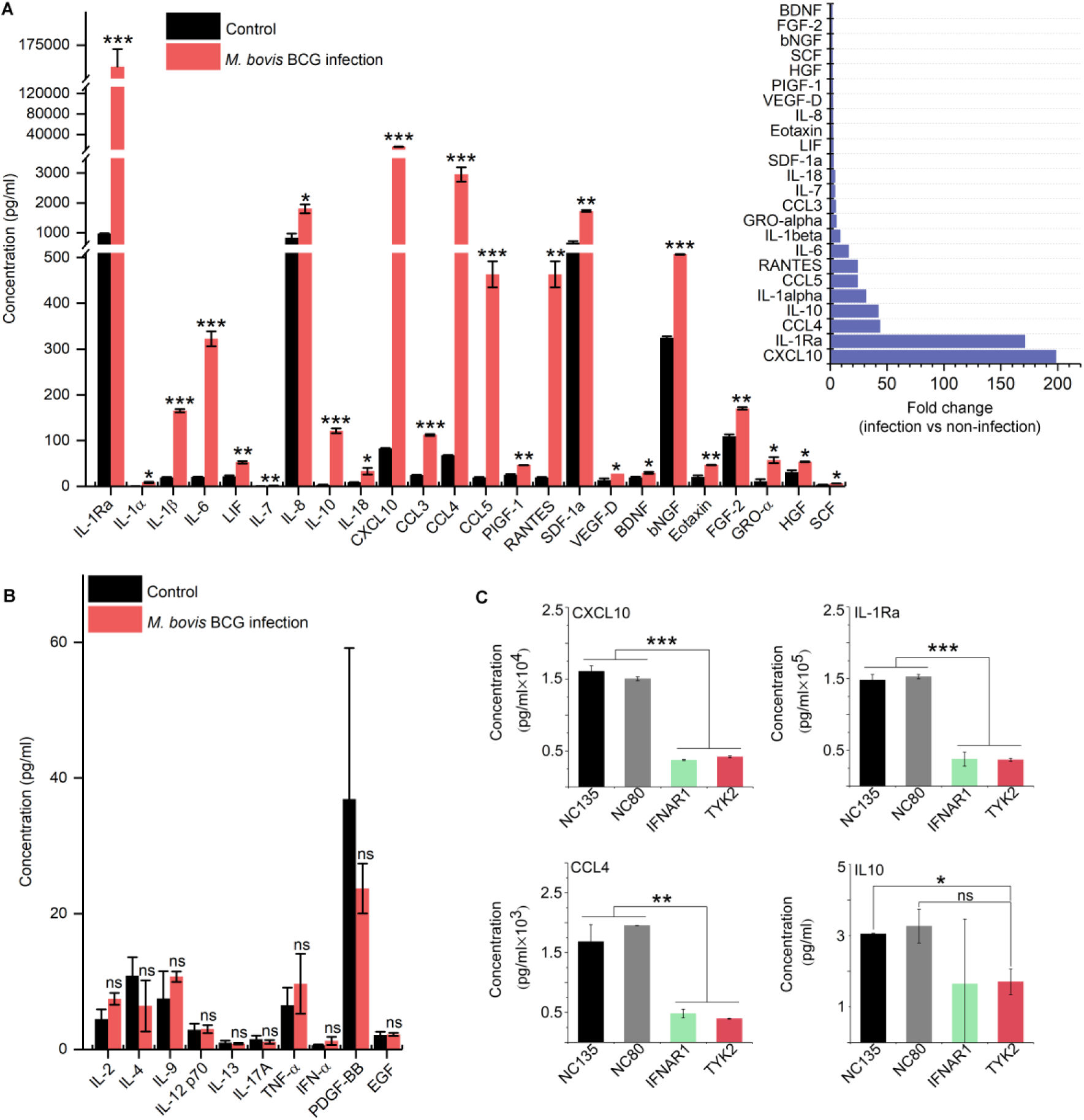
Cytokine and chemokine profile and post-infection treatment of inhibitors in THP-1 cells. Related to Figure 5. (A) and (B) Cytokine and chemokine production of THP-1 cells in the presence or absence of *M. bovis* BCG infection. (C) Infection-induced cytokine and chemokine production in non-targeting control (NC135 and NC80), IFNAR1 and TYK2 gene knockdown THP-1 cells. Data represent the mean ± SD (n = 3) (two-tailed unpaired Student’s *t*-test, * P<0.05 ** P<0.01 *** P<0.001; ns represents not significant).

**Figure S5.**
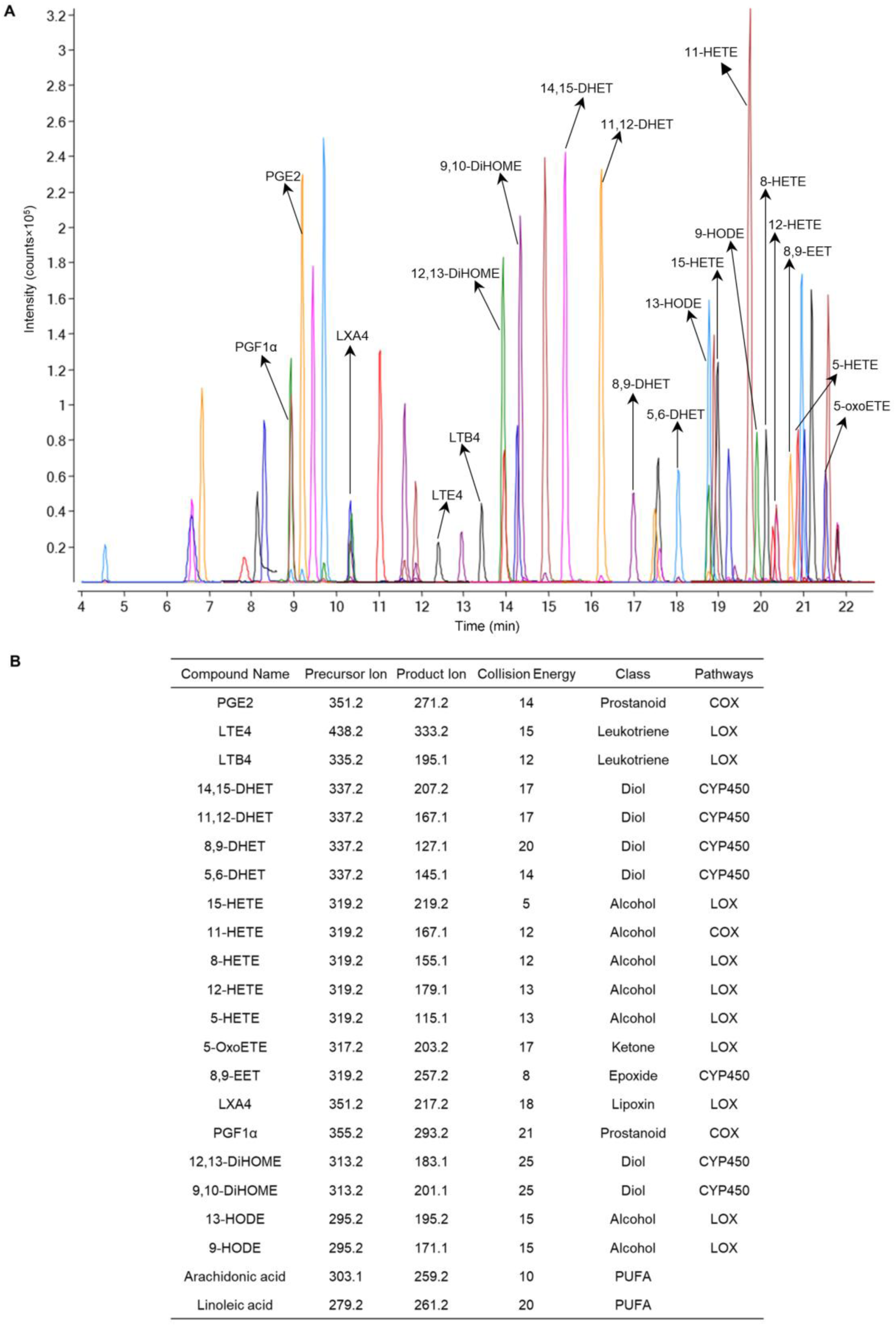
Oxylipin standards used in LC-MS/MS analysis. Related to Figure 6. (A) A representative UPLC-MS/MS chromatogram of a mix of standards. (B) Compounds were quantified in multiple reaction monitoring (MRM) mode with defined time segments. PGE2, prostaglandin E2; LTE4, leukotriene E4; LTB4, leukotriene B4; DHET, dihydroxyeicosatrienoic acid; HETE, hydroxyeicosatetraenoic acid; 5-OxoETE, 5-Oxo-eicosatetraenoic acid; EET, epoxyeicosatrienoic acids; LXA4, lipoxin A4; PGF1α, prostaglandin F1 alpha; 9,10-DiHOME, 9(10)-dihydroxy-12Z-octadecenoic acid; 12,13-DiHOME, 12(13)-dihydroxy-9Z-octadecenoic acid; HODE, hydroxyoctadecaenoic acids.

**Table S1.** Enrichment of biological pathways identified by genome-wide CRISPR knockout and CRISPRi screens in mycobacterial infection. Related to **Figure 2**. (A) and (B) Gene enrichment analysis of genetic hits identified by CRISPR knockout screen (A) and CRISPRi screen (B) in *M. bovis* BCG infection (FDR<0.1 and log2 FC>1).

| **A** CRISPR KO screen |  | **B** CRISPRi screen |  |
| --- | --- | --- | --- |
| Biological pathways | p-value<br>(-log10) | Biological pathways | p-value<br>(-log10) |
| DNA repair | 10.8 | TCA cycle and respiratory electron transport | 13.6 |
| Interferon alpha/beta signaling | 9.1 | Interferon signaling | 11.5 |
| Chromatin organization | 7.2 | DNA repair | 9.3 |
| Cellular responses to stress | 5.0 | Mitochondrial biogenesis | 6.3 |
| SUMOylation | 4.7 | Cellular responses to stress | 6.2 |
| Transcriptional regulation by TP53 | 4.1 | Chromatin organization | 5.6 |
| Signaling by SCF-KIT | 3.6 | Apoptosis | 3.8 |
| Signaling by FGFR | 3.4 | Transcriptional regulation by TP53 | 3.8 |
| PI3K/AKT activation | 3.2 | Activated TAK1 mediates p38 MAPK activation | 3.5 |
| Amino acid transport across the plasma membrane | 3.1 | Regulation of lipid metabolism by PPARalpha | 3.5 |
| NGF-TRKA signaling | 2.9 | Signaling by SCF-KIT | 3.5 |
| MAPK1 (ERK2) activation | 2.9 | Signaling by FGFR | 3.4 |
| Tryptophan catabolism | 2.6 | PI3K/AKT activation | 3.1 |
| Signaling by NOTCH1 | 2.4 | Cell cycle checkpoints | 3.0 |
| Antigen processing: Ubiquitination & Proteasome degradation | 2.4 | Cytochrome P450 - arranged by substrate type | 1.5 |
| Circadian clock | 1.4 | Phospholipid metabolism | 1.3 |

## LEAD CONTACT AND MATERIAL AVAILABILITY

Further information and requests for resources and reagents should be directed to and will be fulfilled by the Lead Contact, Timothy K. Lu. Materials generated in this study are available from the Lead Contact without restrictions.

## EXPERIMENTAL MODEL AND SUBJECT DETAILS

### Mammalian cell cultures

The human monocytic THP-1 cell line was a gift from Jianzhu Chen (Singapore-MIT Alliance for Research and Technology) or purchased from ATCC (TIB-202) for *M. bovis* BCG or Mtb experiments, respectively. The HEK293FT cell line was a gift from Asha Shekaran (Engine Biosciences). HEK293FT cells were cultured in Dulbecco’s modified Eagle’s medium (DMEM; HyClone) supplemented with 10% FBS (Gibco) and penicillin-streptomycin (Pen/Strep; Gibco) at 37°C with 5% CO_2_. THP-1 cells were cultured in RPMI1640 (HyClone or Gibco) with 10% FBS and Pen/Strep at 37°C with 5% CO_2_. Cell growth was measured by CellTiter-Glo luminescent cell viability assay kit (Promega). Prior to Mtb infection, THP-1 cells were adhered to white-bottom 96-well plates (Corning) by 24h of treatment with 50 ng ml^-1^ phorbol 12-myristate 13-acetate (PMA; Calbiochem) followed by 24h of incubation without PMA in antibiotic-free media.

Primary human monocytes were isolated from frozen peripheral blood mononuclear cells, obtained by Ficoll gradient centrifugation of healthy donor leukaphereses (Research Blood Components), by CD14 positive selection (Stemcell Technologies). Monocytes were allowed to mature into macrophages on tissue culture treated dishes using 50 ng ml^-1^ GM-CSF (BioLegend) for 6 days in RPMI1640 with 10% FBS, 25 mM HEPES, and 1× GlutaMAX (Gibco). Matured macrophages were dissociated with accutase (Innovative Cell Technologies), counted, distributed in white-bottom 96-well plates, and allowed to adhere overnight in the same media without GM-CSF. All incubations were performed at 37°C with 5% CO_2_.

### Bacterial strains and growth conditions

*M. bovis* BCG and Mtb H37Rv-lux were grown in Middlebrook 7H9 medium supplemented with 10% albumin-dextrose-catalase (ADC) or oleic acid-albumin-dextrose-catalase (OADC) for *M. bovis* BCG or Mtb, respectively, 0.5% or 0.2% glycerol, and 0.05% Tween-80 at 37°C. A green fluorescent protein (GFP)-reporter *M. bovis* BCG strain was generated by transformation with plasmid P*map24*::GFP, a gift from Jennifer A. Philips (Washington University School of Medicine in St. Louis), to assess mycobacterial infection (Philips et al., 2005). H37Rv-lux was a gift from Bryan D. Bryson (Massachusetts Institute of Technology); it was generated by transformation with a published mycobacterial reporter plasmid (Addgene #26161) that was modified for insertion at the Tweety (mycobacteriophage) site with a zeocin resistance marker (Andreu et al., 2010). When necessary, 25 μg ml^-1^ kanamycin, 100 μg ml^-1^ ampicillin, or 20 μg ml^-1^ zeocin were added to the growth medium.

## METHOD DETAILS

### Reagents and antibodies

Cerdulatinib and imatinib mesylate were purchased from Cayman Chemical, and CH223191 was purchased from Cayman Chemical or Sigma for the *M. bovis* BCG or Mtb experiments, respectively. Compounds were used at the following concentrations: 1 nM to 10 μM cerdulatinib, 0.03-3 μM CH223191, and 10 μM imatinib. The antibodies used were: anti-CRISPR-Cas9 monoclonal antibody 7A9-3A3 (Abcam), vinculin monoclonal antibody (Enzo life sciences), and goat anti-mouse IgG HRP conjugate (Thermo Scientific). Antibiotics in the media were used at the following concentrations: 100 μg ml^-1^ ampicillin, 25 μg ml^-1^ kanamycin, 100 μg ml^-1^ gentamicin, 10 μg ml^-1^ blasticidin, 2 μg ml^-1^ puromycin, 100 U ml^-1^ Pen/Strep, and 20 μg ml^-1^ zeocin.

### *In vitro* bacterial infection

Frozen *M. bovis* BCG with P*map24*::GFP or Mtb H37Rv-lux cultures were thawed and grown in 7H9 medium with appropriate antibiotics to an absorbance at a wavelength of 600 nm of 0.2-0.3 to prevent bacterial clumping (0.3 OD corresponds to ∼ 5×10^7^ bacterial cells for *M. bovis* BCG, and ∼1×10^8^ bacterial cells for Mtb). THP-1 cells were infected with *M. bovis* BCG strains at indicated MOIs (3, 10 and 50) and incubated at 37°C with 5% CO_2_. After 24 hours, *M. bovis* BCG-infected host cells were washed twice with 1×PBS and resuspended in complete RPMI medium containing 100 μg ml^-1^ gentamicin for 2 hours to kill extracellular bacteria. THP-1 cells were then centrifuged, washed with 1×PBS, and maintained in RPMI1640 medium. For 3 rounds of *M. bovis* BCG infection, THP-1 cells were infected once a day for 3 days at an MOI of 10 and extracellular bacteria were removed 24 hours after the third infection. Subsequently, host cells were washed and maintained for the rest of the experiments. The number of viable THP-1 cells were counted in haemocytometer by using trypan blue (Gibco).

Prior to infection, bacterial clumps were removed from Mtb H37Rv-lux cultures by “soft spin” centrifugation of PBS-washed bacteria in cell culture media at 121×*g* with no brake, after which the top half of the supernatant was isolated and used (Saito et al., 2017). Primary macrophages and adherent THP-1 cells were infected with Mtb H37Rv-lux at an MOI of 2 for 7 hours, followed by 3 washes with 1×PBS and incubation in RPMI with 10% FBS, 25 mM HEPES, 1× GlutaMAX, and indicated compounds in 0.1% DMSO. Infection was carried out for 7.5-9 days; all incubations were at 37°C with 5% CO_2_. Luminescence was measured on a BioTek Synergy H1 microplate reader. For cell death measurement by LDH assay (Takara), culture supernatants were clarified with a 5 minute spin at 400×*g*, added at a 5-fold dilution in 1×PBS to the working solution, and incubated for approximately 1 hour before absorbance measurement at 492nm and 620nm on the same plate reader.

### Enumeration of bacteria in infected cells

THP-1 cells (1 ml) infected with *M. bovis* BCG were centrifuged and washed twice with 1×PBS and then lysed with 50 μL of 1×PBS with 1% Triton 100. 10-fold serial dilutions were performed followed by plating on 7H10 agar plates and incubation at 37°C for 3-4 weeks. The number of viable intracellular bacteria was calculated from manually enumerated colony forming units (CFU) on the agar plates.

After Mtb infection, non-adherent bacteria were isolated by combining supernatant and a single 1×PBS wash, while adherent bacteria were those remaining in the well following this wash. Both samples were lysed with 0.1% Triton 100 for at least 5 minutes and diluted 10-fold in 7H9 media followed by 2-fold dilution in 1×PBS with 0.05% Tween-80. Dilutions were plated on 7H11 agar plates and incubated at 37°C for approximately 2 weeks.

### Generation of Cas9-expressing and Cas9-KRAB-expressing THP-1 cell lines

The plasmid lentiCas9-Blast (Addgene #52962) with human codon-optimized sequence Cas9 (SpCas9) was transduced into THP-1 cells to construct the Cas9-expressing THP-1 cell line, which was subsequently expanded in the presence of 10 μg ml^-1^ blasticidin (Sanjana et al., 2014). The plasmid pHR-SFFV-dCas9-BFP-KRAB (Addgene #46911) was transduced into THP-1 cells; dCas9-KRAB-expressing THP-1 cells, which produce blue fluorescent protein (BFP), were subsequently collected using a BD FACS Aria II cell sorter (Gilbert et al., 2013). Monoclonal Cas9 and dCas9-KRAB expressing THP-1 cells, obtained by limiting dilution, served as the parental cell line harboring the human genome-wide CRISPR-Cas9 knockout and CRISPRi libraries, respectively.

### Pooled Genome-wide and secondary CRISPR Screens

Human CRISPR knockout pooled library (Brunello) was obtained from Addgene (#73178). Human CRISPRi pooled library (Dolcetto) was a gift from John Doench (Broad Institute, also available on Addgene #92385). For the secondary screens, we designed CRISPR knockout and CRISPRi libraries, with 10 sgRNAs per gene, targeting 251 genes (169 of which were identified in mycobacterial infection) scored in primary genome-wide screens, 121 of genes from published literature (86 of genes involved in mycobacterial infection), and 100 non-targeting sgRNAs.

### Lentiviral library packaging

Well-dissociated HEK293FT cells were seeded in T175 tissue culture flasks 24 h before transfection in a total volume of 35 ml of DMEM medium at a density of 1.4×10^7^ cells per flask. Cells were transfected at 80-90% confluency using 210 μL of Lipofectamine 2000, 231 μL of PLUS reagent, 7 ml of Opti-MEM, and a DNA mixture of 11.9 μg of pMD2.G (Addgene #12259), 18.2 μg of psPAX2 (Addgene #12260), and 23.8 μg of library plasmid. Flasks were incubated 37°C with 5% CO_2_ for 4 hours, and the media was then replaced with 35 ml DMEM medium with 1% BSA and 10% FBS. Lentivirus was harvested 2 days after the start of transfection and filtered through a 0.45 µm polyethersulfone membrane.

### Lentivirus transduction

Cas9-expressing and dCas9-KRAB-expressing THP-1 cells were transduced with the pooled lentiviral CRISPR knockout and CRISPRi libraries in three biological replicates at an MOI of 0.3 to ensure that only one gene was targeted in each cell. To ensure that each perturbation was fully represented and to reduce spurious effects due to random genome integration in the transduced cell population, screening libraries were prepared with coverage of >500 cells per sgRNA. Lentiviral spinfection was performed by centrifuging 12-well plates at 1,000×*g* for 2 hours at 33°C with THP-1 cells grown in RPMI1640 medium with 10% FBS and 8 μg ml^-1^ of polybrene. Twenty-four hours after lentiviral transduction, cell culture medium was replaced by RPMI1640 with 10% FBS and 2 μg ml^-1^ of puromycin for selection. Following antibiotic selection, a library coverage of > 3000× was maintained for subsequent screens.

### CRISPR screens

After puromycin selection, each CRISPR library replicate was split, one for mycobacterial infection, and one used as control to verify library representation. 100 μg ml^-1^ of gentamicin was added to the cell culture to kill extracellular *M. bovis* BCG post infection. Surviving host cells were harvested and pelleted by centrifugation with coverage of >500 cells per sgRNA. The pooled screens were performed as three independent replicates.

### Genomic DNA extraction, barcode amplification, next generation sequencing and analysis

Genomic DNA (gDNA) from live cells was isolated using a homemade modified salt precipitation method as described previously (Chen et al., 2015). gDNA concentrations were quantitated by Nanodrop. The sgRNA cassette was amplified and prepared for Illumina sequencing (HiSeq2000) as described previously (Sanson et al., 2018). Sequencing reads were deconvoluted to generate a matrix of read counts, which were then normalized under each condition by the following formula: log2 (Reads per sgRNA/(total reads per condition*10^6^+1)). The log2 fold change of each sgRNA was determined by comparing infected and uninfected samples for each biological replicate. To evaluate the rank and statistical significance of genes, a CRISPR screen analysis tool developed by Genetic Perturbation Platform (GPP) at Broad institute was used (https://portals.broadinstitute.org/gpp/public/analysis-tools/crispr-gene-scoring). The significance of the overlap of screen hits between CRISPR knockout and CRISPRi screens was calculated using the hypergeometric distribution. Tests were carried out in R statistical package using the function phyper (q, m, n, k, lower. tail=FALSE), where q is the number of overlap genetic hits, m is the number of genetic hits identified by CRISPR knockout screen, n is the total number of genes in the library, and k is the number of genetic hits identified by CRISPRi screen. The signal-to-noise ratio of secondary screens was calculated by the formula: 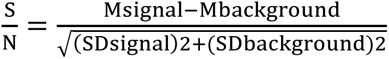. Genetic hits identified by CRISPR screens were used to perform gene-set enrichment using the g:Profiler tool (Raudvere et al., 2019). The KEGG, Reactome and the Gene Ontology (Biological Process) were used as the pathway databases to identify gene sets. Enrichment Map was used for the interpretation of the biological processes (Merico et al., 2010).

### Validation of individual sgRNAs

For each sgRNA, spacer-encoding sense and antisense oligonucleotides with BsmBI-compatible overhangs were annealed, cloned into the lentiGuide-Puro vector (Addgene #52963), and verified by sequencing (**Table S8**). Lentivirus was generated in HEK293FT cells using Lipofectamine 2000 and PLUS reagents following the manufacturer’s instructions. Lentiviral transduction was performed in dCas9-KRAB-expressing THP-1 cells to generate individual knockdown THP-1 cells. After 11 days of puromycin selection, each knockdown THP-1 cell line was infected by *M. bovis* BCG to validate its phenotype.

### RNA extraction and RT-qPCR analysis

Total RNA was extracted using the RNeasy Plus kit (Qiagen) according to the manufacturer’s instructions. cDNA synthesis was performed with iScript cDNA synthesis kit (Bio-Rad). qPCR was performed using iTaq universal SYBR green supermix (Bio-Rad) with a 20 μL reaction consisting of 5 ng of cDNA, 10 μL of supermix, and 0.5 μM of primers. The qPCR reactions were run on CFX Connect real-time PCR detection system (Bio-Rad). Relative quantification of mRNA was performed using GAPDH mRNA as internal control.

### Western blotting

THP-1 cells were lysed in M-PER mammalian protein extraction reagent (Thermo Scientific) supplemented with Pierce protease inhibitor (Thermo Scientific) followed by shaking at 4°C for 10 minutes. Protein concentration was measured by Pierce BCA protein assay (Thermo Scientific). Protein samples were mixed with 4× Laemmli buffer (Bio-Rad), denatured at 70°C for 10 minutes, and loaded onto ExpressPlus PAGE gel (Genscript). Proteins were transferred onto polyvinylidene difluoride (PVDF) membranes using iBlot blotting system (Invitrogen). After blocking with 5% non-fat milk solution in phosphate buffered saline with Tween 20 (PBST), the membrane was incubated with the following primary antibodies: anti-CRISPR-Cas9 monoclonal antibody 7A9-3A3 (Abcam, 1:1,000) and vinculin monoclonal antibody (Enzo, 1:20,000). Secondary incubation was performed with goat anti-mouse IgG HRP conjugate (Thermo Scientific, 1:10,000). Protein bands were visualized with Pierce ECL Western blotting substrate (Thermo Scientific) by autoradiography.

### Surveyor assay

Genomic DNA was extracted from cell cultures using the DNeasy Blood and Tissue kit (Qiagen). The AAVSI target locus was amplified by PCR with high fidelity KOD-Plus-Neo DNA polymerase (TOYOBO) (Wang et al., 2014). 200-400 ng of the PCR amplicons were denatured, reannealed, and incubated with 1 μL of Surveyor Nuclease S and 1 μL of Enhancer S (IDT) at 42°C for 1 h. After incubation, 6 μL of digested product was loaded onto a polyacrylamide gel (4-12%) and run at 120 V for 2.5 h. Gels were stained with ethidium bromide and imaged with a Gel Doc imaging system (Bio-Rad). Quantification was based on band intensity. The ratios of the uncleaved to cleaved DNA bands were used to calculate the percentage of insertion-deletion mutations (indel) in the starting cell population.

### Cytokine and chemokine quantification

Supernatants were collected at indicated times post-bacterial infection. Cytokine and chemokine levels in mycobacterial-infected supernatants were determined using human cytokine and chemokine and growth factor 45-plex panel 1 (Thermo Scientific) according to the manufacturer’s instructions. The results were measured by a Bio-Plex 200 system (Bio-Rad).

### Oxylipin Analysis

Oxylipin extraction and liquid chromatography-mass spectrometry (LC-MS) analysis followed the published reports with modifications (Fang et al., 2017; Lundstrom et al., 2013). Briefly, cell cultures were harvested at given time points, rapidly quenched, and spun down. Cell pellets were resuspended in 1 mL acetonitrile/methanol (1:1), containing 10 μL antioxidant solution (0.2 mg ml^-1^ BHT/EDTA) and 10 μL of internal standard solution, and lysed mechanically with 0.1-mm silica beads by using Qiagen Tissuelyser II. The lysates were collected and evaporated to dryness in a vacuum evaporator, and the dry extracts were redissolved in 100 µL of acetonitrile/methanol (1:1) for LC-MS analysis. The LC-MS/MS analysis was performed with Agilent 1290 ultra pressure liquid chromatography (UPLC, Waldbronn, Germany) coupled to an electrospray ionization with iFunnel technology on a triple quadrupole mass spectrometer. Chromatographic separation was achieved using Acquity UPLC BEH C18 (2.0 × 150 mm, 1.7 μm; Milford, MA, USA) with a flow rate of 0.30 ml min^-1^ at 40°C during a 35 min gradient. (0-3.5 min from 15% B to 33% B, 3.5-5.5 min B to 38%, 5.5-7 min to 42% B, 7-9 min to 48% B, 9-16 min to 60% B, 16-20 min to 75% B, 20-24 min to 85 % B, 24-25 min to 100% B which was held for 2 min, then returned to initial condition over 0.1 min), while using the solvents A, 0.1% acetic acid, and B, 90:10 v/v acetonitrile/isopropanol. The auto-sampler was cooled at 4°C and 5 μL of the extract was injected. Electrospray ionization was performed in negative ion mode with the following source parameters: drying gas (N2) temperature 200°C with a flow of 14 L min^-1^, nebulizer gas pressure 30 psi, sheath gas temperature 400°C with a flow of 11 L min^-1^, capillary voltage 3,000 V, and nozzle voltage 800 V. Data acquisition and processing were performed using MassHunter software (Agilent Technologies, US).

### Flow cytometry

Cas9 activity was tested by transducing pXPR_011 (Addgene #59702), which expresses both GFP and sgRNA against GFP, into Cas9-expressing THP-1 cells. The extent of GFP loss was quantitated by flow cytometry. Ten days post-infection, cells were washed and resuspended in 1×PBS supplemented with 2% FBS. After passing through EASYstrainer cell sieves (VWR) to remove any clumps of cells, samples were measured by LSRII Fortessa flow cytometer (Becton Dickinson). At least 30,000 cells were recorded per sample. BD FACS Aria II Cell Sorter was used to collect dCas9-KRAB-expressing THP-1 cells, which express both BFP and dCas9-KRAB proteins. 110,000 BFP positive cells were collected. Forward and side scatter were used to identify appropriate THP-1 cell populations.

### Imaging

To visualize intracellular GFP-reporter *M. bovis* BCG (P*map24*::GFP), infected THP-1 cells were directly observed under a confocal fluorescence microscope (Zeiss LSM 700).

## QUANTIFICATION AND STATISTICAL ANALYSIS

GraphPad Prism 8 software and Excel were used for all statistical analysis. All of the statistical details of experiments can be found in the corresponding figure legends.

## DATA AND CODE AVAILABILITY

Positive genetic screen datasets are available in Tables S2, S3, S6, and S7. sgRNA target sequences of secondary CRISPR screen libraries are available in Tables S4 and S5. The NGS dataset supporting the current study is available from the corresponding author upon request.

## SUPPLEMENTAL INFORMATION

**Table S2**: Positive genetic hits from the genome-wide CRISPR-Cas9 knockout screen in *M. bovis* BCG infection. Related to Figure 2.

**Table S3**: Positive genetic hits from the genome-wide CRISPRi screen in *M. bovis* BCG infection. Related to Figure 2.

**Table S4**: Secondary CRISPR-Cas9 knockout target genes and sgRNA target sequences. Related to Figure 3.

**Table S5**: Secondary CRISPRi target genes and sgRNA target sequences. Related to Figure 3.

**Table S6**: Positive genetic hits from 2nd CRISPR-Cas9 knockout screen in *M. bovis* BCG infection. Related to Figure 3.

**Table S7**: Positive genetic hits from 2nd CRISPRi screen in *M. bovis* BCG infection. Related to Figure 3.

**Table S8**: Oligonucleotides used for THP-1 gene knockout or knockdown cell construction. Related to Figures 1 and 5.

