## Supplementary material for "Illuminating host-mycobacterial interactions with functional genomic screening to inhibit mycobacterial pathogenesis": Table S2

**Table S2. Positive genetic hits from the genome-wide CRISPR-Cas9 knockout screen in *M. bovis* BCG infection. Related to Figure 2.**

| Gene Symbol | Average LFC | Average -log(p-values) | Number of perturbations | Perturbations | Individual LFCs |
|---|---|---|---|---|---|
| TYK2 | 7.896 | 9.115 | 4 | TTGGGCCTGAGCATCGAAGA;TGAATGACGTGGCATCACTG;CAGGCGGCCCTCATACACGT;AATACCTAGCCACACTCGAG | 7.8;7.83;7.94;8.02 |
| IFNAR1 | 7.834 | 9.496 | 4 | CTCCGCGTACAAGCATCTGA;GTACATTGTATAAAGACCAC;ATAATTGGATAAAATTGTCT;TAGATGACAACTTTATCCTG | 6.76;8.11;8.15;8.31 |
| IFNAR2 | 7.731 | 9.184 | 4 | CGTCATTGAAGAACAGTCAG;ATATCCATGGCTTCCAACGG;TGAGTGGAGAAGCACACACG;TGAAAGTGATAGCGATACTG | 6.59;8.01;8.04;8.28 |
| STAT2 | 7.090 | 8.462 | 4 | ACCTACCCTTGTAATTGAGG;AGCAACATGAGATTGAATCC;ATCATCTCAGCCAACTGGGT;ACGGTCAAATATACCTACCA | 4.95;7.57;7.83;8.01 |
| STAT1 | 6.547 | 8.316 | 4 | TCCCATTACAGGCTCAGTCG;AGAACACGAGACCAATGGTG;CCTGATTAATGATGAACTAG;GACGTTGGAGATCACCACAA | 5.99;6.2;6.66;7.34 |
| JAK1 | 6.460 | 8.298 | 4 | TGGTTTCATTCGAATGACGG;CACACTTACTCTCCACGTCG;CCGGAAGTAGCCATCTACCA;GCCTAGACAGCACCGTAATG | 2.06;6.82;8.46;8.49 |
| HDAC2 | 5.302 | 7.685 | 4 | TCAAAGAGTCCATCAAACAC;CCTCCTCCAAGCATCAGTAA;TACAACAGATCGTGTAATGA;GATGTATCAACCTAGTGCTG | 4.66;4.81;5.45;6.29 |
| ACTR5 | 5.044 | 7.688 | 4 | TCATTTGCCGTGAATACAGT;CGGGACACGCCCAGCCAGCG;TGAGCACAGCTACATCGCTG;TGGGTAAAACATGCGTACAC | 4.83;4.9;5.01;5.44 |
| DNAJB6 | 5.032 | 7.179 | 4 | GACTTCTTTGGGAATCGAAG;TCCTACAGAATTGTCGAGAA;GCCACTACCACCAAATGACG;GCATATGAAGTGCTGTCGGA | 2.24;5.32;5.66;6.91 |

| | | | | | |
|---|---|---|---|---|---|
| CDKN2C | 4.944 | 7.610 | 4 | ACAAACTAGTAAGTTGCTCT;GCAAAGTCTGTAAAGTGTCC;GAATGACAGCGAAACCAGTT;ATGCACAAAATGGATTTGGA | 4.7;4.77;5.09;5.21 |
| TRERF1 | 4.761 | 6.585 | 4 | CGGGCATGACCATAGGCGTG;ATGGATACCTCGAGGCAGGG;GGATCCACCTCAACAACATG;GCAACGTGACGGTCACCCCA | 1.29;5.48;5.96;6.31 |
| IRF9 | 4.292 | 4.994 | 4 | CTTCATCTACAACGGGCGCG;AACTGAGGCCCCCTTTCAAG;ACAATTCCACAGGCCAGCCA;AATTTAAGGAGGTTCCTGAG | 0.45;5.24;5.32;7.06 |
| USP7 | 4.074 | 7.175 | 4 | GGCAACCTTTCAGTTCACTG;TTGATGACGACGTGGTGTCA;AGATGTATGATCCCAAAACG;GGCAGTAGAACAGCTCGATG | 3.73;3.79;4.09;4.69 |
| IRF8 | 3.986 | 6.944 | 4 | GCGTAACCTCGTCTTCCAAG;CTTCTGTGGACGATTACATG;ATGGCTCGGAAATGTCCAGT;TGAATGGTGCGCGTCGTAGG | 3.02;3.53;4.02;5.37 |
| DNTTIP1 | 3.963 | 6.901 | 4 | TCGAGATCCCAACACAAAGG;GGTCCCTACTCACACCATGC;TTGTTGTCATCTGTGAGCGG;GAACGTGCGAGACAATGTTG | 2.7;3.78;4.64;4.74 |
| UCHL5 | 3.666 | 5.780 | 4 | TTTATTATTTCAGGTTGCCG;GAATTAGATGGATTAAGAGA;GTTACTGAACTGTACCCACC;ATGATTGGATCAGTGCAGTA | 0.84;4.27;4.66;4.88 |
| SPEN | 3.362 | 6.701 | 4 | GATATTACCCGGGAGGTACG;ACACATCGCAAAGCTCGCTG;GTTTGCCTACCTCGTGAACG;AAGGAGCGTCTATGCAACCA | 3.07;3.12;3.36;3.9 |
| SEC62 | 3.356 | 6.563 | 4 | TACCAATAAAATAATCAACC;AGAAAAACCTGATCATCATG;GGTGTTTATTACCTCAGTGT;ATCATTTGGCTCATAACTGG | 2.25;3.58;3.77;3.83 |
| C16orf72 | 3.241 | 6.171 | 4 | CCCTAACCGAGCTACTTCAA;GCAATAAGGATGTGTTGGCT;TTTGTAGAGATTGGTGACGG;GCCGTGGCCCAGCTCTACAA | 1.54;3.54;3.93;3.96 |
| HSPA13 | 3.131 | 6.367 | 4 | ATACCAATCACTTTAGGAGT;ACTGACAATGATGTATATGT;TCGAGAGCCAACATATTCTG;TCTTTACTGAATAAACAAGG | 2.02;3.41;3.54;3.55 |

| | | | | | |
|---|---|---|---|---|---|
| KMT2D | 2.980 | 6.465 | 4 | GGTGGAAATTCCCGCCAACG;CCCCTATCCTAGTCCTACGG;CCTTCGAAGCGAAAGTACTG;TTCGGGGTAGACCTCCATAG | 2.76;2.85;3.01;3.3 |
| FOSL2 | 2.959 | 4.194 | 4 | AGTGTGCAAGATTAGCCCCG;CTGAGCCAGGCATATCTACC;GATCACGCCAGGTCTTGGGA;AGGAGAAGCGTCGCATCCGG | -2.73;3.52;4.64;6.4 |
| NFRKB | 2.953 | 6.061 | 4 | TCTCAGATAAAGTAGAACTG;AACCAGCTAAGTCTAGCTCG;CCCTGTGTGAAATACGACAT;CGGACACTTTAACCCCGAGG | 1.64;3.14;3.45;3.59 |
| TMEM30A | 2.928 | 5.717 | 4 | GAGATTTCACGTAACGACGA;GTAGCACCGTGCCAGCCGTA;CATTTATTACAGGGACTGGA;TTGTACCATTAACTTCACAC | 1.45;1.97;4.04;4.26 |
| NUFIP2 | 2.819 | 5.509 | 4 | GAGGACCCTGGAAATAGGGT;GGGCCTGGAGGAACAAGTCG;CAGCGCTGACATAGGGACCT;AGTGTCCCAACCTTAAAACA | 1.33;1.76;3.9;4.28 |
| JUNB | 2.672 | 6.022 | 4 | CACAGCTACGGGATACGGCC;GGGTAAAAGTACTGTCCCGG;CCGGAGTCTCAAAGCGCCTG;CTGAGGTTGGTGTAAACGGG | 1.91;2.58;2.82;3.38 |
| MCTS1 | 2.576 | 4.649 | 4 | CAATTGATAGAGCAATTTCC;TTGTGAAGTAATCTTAGGGT;GGAGCAAATATCATGTGTCC;ATCCTGTCAAAATAGTCCGA | 0.06;3.14;3.5;3.61 |
| SLC3A2 | 2.551 | 5.452 | 4 | CAGGCCCGTGAACTTAGCCG;GACCTTACTCCCAACTACCG;TGAGTGGCAAAATATCACCA;GCGCAGAAGTGGTGGCACAC | 1.15;2.62;2.92;3.51 |
| SIAH2 | 2.516 | 5.857 | 4 | CTTGTGGGCGTGCATGAGAT;GACCGGACACTCGAAGAGCG;TGCAGGAAGCACCAGGACAT;TCTTTGCAGTATGCCACCAC | 1.87;2.09;2.88;3.22 |
| BAHD1 | 2.495 | 5.715 | 4 | ACAGGCCATTGTAGTCGGCG;CAATGGCAAGAACTATCCCA;CAGTAGCACGATCACGGTCA;CCAGCGTGGGTAACAGACAG | 1.76;2.15;2.24;3.83 |
| MBNL1 | 2.490 | 5.842 | 4 | CCCACATTTAAAAACGCAGT;GTATGTCGAGAGTACCAACG;GTATGTAGAGAGTTCCAGAG;CTTGGTGCAACTGAAAACAT | 1.72;2.56;2.67;3.01 |

| | | | | | |
|---|---|---|---|---|---|
| PRR14L | 2.436 | 5.736 | 4 | GGACTATCATGTGTAGAACT;ACAACCAGACATCTTGACAA;GAGGTATTAAAACCCAAGCA;ACTGGGTTACTTAACTGTTG | 1.68;2.23;2.85;2.99 |
| UTY | 2.428 | 5.908 | 4 | TGACTGTCAGCATCCCCCTG;TCAGAACTTCTGTTGAACTG;AGTAGAAGTAGACCAAACCA;AGGATTCATAGAGAGTACCT | 2.06;2.38;2.44;2.84 |
| PHIP | 2.390 | 5.508 | 4 | GTTCATTCACCACCAAACGT;AGCATTGAAACAGACTATCA;TAGGCTGCAAGCATGTTGTG;TCAACATCAACCAAAAAAGG | 1.29;2.59;2.68;2.99 |
| SUV39H1 | 2.266 | 5.107 | 4 | GCCGGGGGGTCTTTGACCGG;ACAGGAACAGGAATATTACC;ATGTACACGAAGGCCCGCGG;GTTCCTCTTAGAGATACCGA | 0.99;2.14;2.66;3.28 |
| C11orf57 | 2.256 | 3.814 | 4 | TTCTTCTGAGAACTCACTCG;CAGATGGAAGATGCTTACCG;CACAGATGCTCACAATAAGG;GATCAGCAAGATATTACCAA | -0.59;2.08;2.43;5.11 |
| FBXW7 | 2.245 | 5.334 | 4 | GTTGGAGTAGAACCTAGACC;AAGAGCGGACCTCAGAACCA;ACATTAGTGGGACATACAGG;ACAGAATTGATACTAACTGG | 1.21;2.49;2.49;2.79 |
| ZRSR2 | 2.227 | 5.720 | 4 | TTCCCTAGGACTCTCACAGA;ACAGTTCCTAGACTTCTATG;AGTTGGAAAATGGTACCACA;TGATAGGCAAAGATTACATG | 1.96;2.0;2.44;2.51 |
| TNRC18 | 2.224 | 5.013 | 4 | TGGTGCCCAAACCTCCAAAG;GTACTGACCATGTGCAACGG;GCTCGGGCAACGCATCCATG;GGATAAGAAAGAGACACAAG | 0.79;2.52;2.77;2.82 |
| SHOC2 | 2.214 | 5.628 | 4 | TCATACCTATAGTATCTGGG;TAACATTTCTACTTTACCAG;GAGCTACATCCAGCGTAATG;TAGTTATACGATTAAAGCGA | 1.76;1.98;2.54;2.58 |
| LARP4B | 2.142 | 5.504 | 4 | CCAGTATGTGCCAATCACAA;AATATCTGAATCTACCCCCG;GGAACTTGCCTGCTTCGAGG;TGCGTTGCTGACCACGTCTG | 1.62;2.09;2.32;2.53 |
| INO80C | 2.128 | 3.018 | 4 | TGCCTTCAATTCAAGCACTC;CTTCCACTCCCGGAATAGTC;TGAGTTTAGCACAGGACCTG;TTAGGATCGTTCAGTTGCCA | -0.67;0.47;4.13;4.57 |

| | | | | | |
|---|---|---|---|---|---|
| RBM48 | 2.081 | 4.609 | 4 | AAAATGTATGTGTTCATCCG;GAAACCATCATAAAACAATG;TACAGAGCATAAAGACACAG;CACACGGGCCAAATATCGAG | 1.01;1.15;2.75;3.41 |
| TFPT | 2.067 | 3.854 | 4 | TTCTGCGATTTAATTCCCGC;GAGAGTGCTGGACTCCTACG;GAAGTGGAGTTTGTGTCAGG;CCACCCCATAGGTGAACGAG | -0.81;2.45;3.0;3.63 |
| SATB1 | 2.023 | 5.302 | 4 | TAGGTGTTGATACGAGCCCA;TATTCATAGATCTACTGACA;ATGCTAAGTACCTGTGAAAG;CATTGAATATGATTGCAAGG | 1.56;1.77;2.22;2.54 |
| KDM4A | 2.021 | 4.887 | 4 | GTGGTATTTCAAGACAAACT;TGTGCACAGTTATGCCAAAG;ATATTTCTTCAGCATTAACG;ATCATACAGACAGCGCAGTG | 1.3;1.33;2.29;3.16 |
| SATB2 | 2.021 | 5.134 | 4 | AACAGACCTCCCTATTAAGG;CAGATTGAGGAAATTCTGCA;AGAGCCAACCAACTCTTCCG;GTGCATCTGTCACATAACTG | 1.35;1.75;2.09;2.89 |
| BAP1 | 2.010 | 4.569 | 4 | TCTACCCCATTGACCATGGT;CACGGACGTATCATCCACCA;GTCCATCAAGACTAGCAGCG;GAACCGTCAGACAGTACTAG | 0.92;1.28;2.53;3.32 |
| DENR | 1.993 | 5.096 | 4 | TGGGTAATCGGCATCTAACT;AGTCTGTTCATTACCAACAG;AAAGAAGACCGTACCACAAA;ATGTTGCTAAATGTAGACAA | 1.21;1.94;2.17;2.65 |
| TNFRSF1B | 1.918 | 5.115 | 4 | AGGAACTGAAACATCAGACG;CTGCGTGTGTTGGGATCGTG;CACCGTGTGTGACTCCTGTG;GGAAACTCAAGCCTGCACTC | 1.51;1.54;2.17;2.46 |
| EWSR1 | 1.908 | 4.962 | 4 | ATGAGTGGCCCTGATAACCG;AAAAAGTAGACTGACCTGGT;GGTCTGCCCATAGGTTGCAG;TGGGTCTTCATAGGACACTG | 1.07;1.99;2.2;2.37 |
| DNAJA1 | 1.895 | 4.585 | 4 | TTTACCGAGTAAAAACAGCA;GGAGAACAGGCAATTAAAGA;TAGGACTGATCCGCTCCCCA;GAGTGCTGTCCCAATTGCCG | 0.77;1.63;2.39;2.79 |
| RB1 | 1.887 | 4.803 | 4 | TGAACTACTTACGAACTGCT;GGTTCTTTGAGCAACATGGG;AACATCTAATGGACTTCCAG;AAACAATCAAAGGACCGAGA | 0.93;1.85;2.32;2.45 |

| | | | | | |
|---|---|---|---|---|---|
| TNFRSF1A | 1.852 | 5.144 | 4 | AAGACCAAAGAAAATGACCA;GTGGACCGGGACACCGTGTG;AGAGGTGCACGGTCCCATTG;GACCAGTCCAATAACCCCTG | 1.48;1.72;2.05;2.17 |
| ARID3A | 1.810 | 4.705 | 4 | AACTCACATGTCCTCGTCGG;ATAGACAGCAACCGACGGGA;ACAGCTCTACGAACTCGACG;GCTGTGGCGTGAGATCACCA | 0.99;1.72;1.81;2.72 |
| KMT2C | 1.802 | 4.890 | 4 | CTGTTGCCAATACACAATCA;ATTCAGACATTAGGCCATCG;CCCATGCGACGACCTCCCCA;GCACACGGTCTAGTTCTCAG | 1.24;1.7;1.83;2.44 |
| NOSIP | 1.784 | 4.558 | 4 | CCACAGATGATGTCCAACCT;GTCAGGTCCGACATGCGCAG;CTCACGTGACAACAGGATCG;GCCCCGCACATGGTCCTGCG | 0.9;1.54;1.92;2.78 |
| RAD23A | 1.764 | 4.223 | 4 | GGGTACCTCAGCACCCCCAG;ATACTCAGAGCCCGTCACTG;GAGTGACGATGTCCCTATCA;CACCGTGAGCAGATACTCCA | 0.22;2.1;2.16;2.57 |
| STAG2 | 1.745 | 4.858 | 4 | AATACTAACCTTGAACCGAC;ATACCTTGTGGATAGCATGT;TTGGAAAACGAGCCAATGAG;TGGAGATTATCCACTTACCA | 1.43;1.54;1.55;2.47 |
| RRAGA | 1.730 | 4.474 | 4 | TTTGATTTACGTGTTTGACG;GTTCCCTAGGAATCGGACGT;GCTGAACGTTGGGAATCAGC;TTATTACCAGTCGTGTCTGG | 0.85;1.47;1.96;2.64 |
| MIER1 | 1.727 | 3.852 | 4 | TCAAAATATTTACATCGACG;GCTGATAATGATGACAACAG;AAATGATGATCAGCTCCTGT;TGATGAACGAACATTAGAAG | -0.18;1.8;2.55;2.74 |
| IKZF5 | 1.712 | 4.900 | 4 | ATCAGCTCTCGACTCTAGCA;GCAAAAACCACACCAACTGG;GAAGCAGAGGCTCTTCAGGG;AGTTACTTCGATCACTGCAG | 1.21;1.84;1.85;1.95 |
| KHSRP | 1.710 | 4.897 | 4 | CGTCGGCGAAAGCGTCCTTG;TTCCACGACAACGCCAACGG;GTGCGGATACAGTTCAAGCA;CAAACCTCTCCGCATCATTG | 1.4;1.49;1.88;2.07 |
| PAXIP1 | 1.708 | 4.266 | 4 | GTGCGTGCATCGACTCGTGA;TCTTCATAAATAATCAGACG;GCACACAAGTTAATCAACCC;GTGATTCTGTCCGTTCAGTG | 0.4;1.8;2.28;2.34 |

| | | | | | |
|---|---|---|---|---|---|
| INO80E | 1.690 | 3.024 | 4 | AAGAGAAGCCCTCCGCTGGG;ATCATCAGATAACAGCGAGA;AGGAGCTGAGGAAAGCGCAA;CGTTCTCGTACTGCAGAAGT | -0.37;0.51;3.15;3.46 |
| PIH1D1 | 1.637 | 4.309 | 4 | TCCTCCCGCCGACGTGACCG;TTTCGCATCCCCATGAGTCT;TAAAGACCAACTCCTCGGAA;GTTGACAGCTACGTCGTAGG | 0.59;1.71;1.89;2.36 |
| PLAGL2 | 1.631 | 3.839 | 4 | AGCTGCCCTCAGCTGCACTG;ATACTCGTAAGGATGTACGG;GGGCAGCGTGTTGTGCACCA;GGTCCCCATTACGTCCCGAG | 0.29;1.18;2.47;2.58 |
| AP3D1 | 1.625 | 4.646 | 4 | TTCACATACTGATCTGACAT;CCAGCGGAACGGGATCTGTG;CATCAGCTTGAAAAAGAGCG;GACCACCAATCAGATCCGTA | 1.07;1.6;1.85;1.98 |
| TP73 | 1.575 | 4.580 | 4 | GGGCGGAACGGATTCCAGCA;GCTGGAAAGTGACCTCAAAG;CATGCCTGTTTACAAGAAAG;AGAGATTATTGCCTTCCACG | 1.27;1.35;1.53;2.15 |
| SPPL3 | 1.555 | 4.590 | 4 | GAAACGTCCACAGCAACCAA;GCTTCCGGGATAGAACGTCA;CACCATCCATGAGAAGCCAA;AGACAGATGCTCCAATTGGA | 1.26;1.39;1.6;1.98 |
| SETD1B | 1.540 | 4.726 | 4 | TTATGTACACAATTCTCCCG;CTACAGGGGAAACCCGAATG;CCAGCAGGCACGAGGCGATG;GGAACTCTCACTGGACGACG | 1.4;1.45;1.65;1.66 |
| RANBP1 | 1.533 | 4.511 | 4 | TTCTGGGAGATCGTTCTCAG;CATCCGCCTCCTCATGCGGA;ACTCGTCGGCGAAGTCAGCG;GTTCATGCCACCTACTGTAG | 1.21;1.3;1.66;1.96 |
| PPP1R2 | 1.532 | 3.728 | 4 | TGGCGCCAGACATCTTAGCC;TCCTCGTCGACATTCCCGCG;CTTGTCATACTAGTATGATG;GACCTCTCACCTGAAGAACG | -0.12;1.74;1.93;2.58 |
| BCORL1 | 1.520 | 3.845 | 4 | TCCCGCATCTGACAGCGCCG;CTGGAACAAGCCCTGAATGG;TTCCCATTGACTGAACACCG;GGAGGCGGGATATATACCAG | 0.32;1.29;2.09;2.38 |
| POU2F1 | 1.497 | 4.131 | 4 | AGCTGGAGGACAGATAACTG;CATAGAGACCAACATCCGTG;ATCATCTCACAGACGCCCCA;GTGATGTTGGGCTCGCTATG | 0.84;1.25;1.52;2.38 |

| | | | | | |
|---|---|---|---|---|---|
| EZH2 | 1.494 | 4.368 | 4 | CTTCTGTGAGCTCATTGCGC;TTATCAGAAGGAAATTTCCG;TTATGATGGGAAAGTACACG;ATGTTGGGGGTACATTCAGG | 0.99;1.4;1.62;1.97 |
| RIC8A | 1.473 | 4.522 | 4 | GCTGCTGGCGCACATCGGTG;GTTTGATGATGCCCAACAGG;AACTCCGTGGAGTCTCCATG;CAGTACAACATCCATGTCTG | 1.22;1.42;1.61;1.64 |
| METTL5 | 1.442 | 3.857 | 4 | TGACATAGATGAAGACGCAT;AATTACTGTATCGAATGACT;AGAGAGTCGCCTGCAACAAG;GAACTGCAATGTTAGGAGCA | 0.68;1.09;1.44;2.56 |
| RNF111 | 1.441 | 4.031 | 4 | AACGAACTGCCATGTAAACA;AGATGGCTATGGATCAAGCA;TATGAGGATGTCCTAATGCA;TAACAGTAGAAATCCTACTG | 0.4;1.69;1.78;1.9 |
| CHD1 | 1.434 | 4.356 | 4 | TTAATTCGCCTAAGAGAACG;TTCCGATGACTCATCAAGTG;ATGCCCAATTTAGACCTCCA;AAGCAGCCATCCTATATTGG | 0.99;1.48;1.59;1.68 |
| KLF13 | 1.410 | 4.264 | 4 | CCGACCTCGAGTCCCCGCAG;GAGTTCTCAGGTGCGCCTTG;CGTGGTGGCGCGGATCCTAG;CGTGTCCATGTCGAGCCGCG | 1.0;1.33;1.51;1.8 |
| MXI1 | 1.405 | 3.943 | 4 | GGTGTTGAAAATGTCTGAGA;CGGCATGGACGGGAATGAAG;GCAAACCAAGTGTTGTGTGC;GCCCGGCTCAACCTCCGTGG | 0.69;1.03;1.9;2.0 |
| BRD1 | 1.399 | 4.139 | 4 | TGTTCTATAGAGCCGCGGTG;GAGGCCTCTGAATATTTACG;AAGACAGATGACGACCGCTG;GCAGCAGTCTCTGATCGACG | 0.82;1.3;1.64;1.83 |
| PRR36 | 1.396 | 3.119 | 4 | AGAAGGTAGGTTCTCCAGAG;AGAAGCTTGGGTTTGCCTAG;GAAGGGACTCCGGATCTCAG;CAGGTAAGGCTCGACGAGTG | 0.26;0.75;1.0;3.57 |
| PQBP1 | 1.396 | 4.072 | 4 | AGCCATGAGAAACTAGACAG;GGGCCACGACAAGTCTGACA;TCGAACACCTTGTACCAGCT;CCCACATGACCCCAACTCCG | 0.94;1.23;1.25;2.16 |
| DYRK1A | 1.392 | 3.978 | 4 | TGAGAAACACCAATTTCCGA;TTACAGGAGTACAAACCACC;TTCAACCAAAATACACCCGA;TCAGCAACCTCTAACTAACC | 0.75;1.05;1.78;1.99 |

| | | | | | |
|---|---|---|---|---|---|
| PIGL | 1.373 | 4.007 | 4 | GGTACACCCAGTGCCTTAGG;AGGGATTTCCCAGATGACCC;AGGTGGTGACTTTCGATGCA;ATAATCATTACACTGGAGAG | 0.85;1.17;1.38;2.1 |
| KLF16 | 1.359 | 3.401 | 4 | GGCCATCTCTTCGGGCGCCG;CGCGCGCACATCCAGGCCGG;CGCAGTCCGGGAAGGGACAG;CAGGTGCGACTTTAGGTGCG | 0.26;0.73;2.04;2.41 |
| XPO6 | 1.355 | 3.799 | 4 | CCTGGCCGAGTACTTTATCG;CGGACAGCACATTACCAGTG;CACCTTATCGACAAGTCGCA;CAACATGATCAGCCCAAGGG | 0.7;0.97;1.54;2.21 |
| PHC3 | 1.339 | 3.818 | 4 | TTTCACACCCGCTACCACTG;TGGGCCAATGTGTACAACAG;ACCAGTTAATAGCACCAGGT;GTCTCCAGCAGCTATAACAG | 0.47;1.26;1.78;1.86 |
| MKL1 | 1.335 | 3.723 | 4 | GAGCAGTGAGCGCTCCGAGG;AGACAGCTCTTCCTTCGATG;TGGCGTAGGATGAGTCCATG;TCCCGGCTTGCCAGTCAGTG | 0.15;1.63;1.65;1.91 |
| PAGR1 | 1.326 | 3.063 | 4 | CTGGCCGTGGAGGATACCGG;AGACACTGCGGCCAGTACGG;GTGCGTGCCCTGCAGCGACG;TTGCAGCTCCAGAGTACCGT | 0.22;0.24;2.01;2.84 |
| SNX12 | 1.297 | 3.170 | 4 | CCACCTATGAGGTTCGCATG;ACTTGGCGGCCCGTAAGCGT;TAAAGGAGTCCTGCGTACGG;GCGGCTTCGAGTTAAGGCGC | 0.34;0.65;1.24;2.97 |
| HMGN2 | 1.295 | 3.883 | 4 | TTGTTTCTTATAGGCTGAAG;TGCTAAGGGAGATAAAGCAA;GTTAGCACTTACCTTTGCAG;ACTTACAGCAGACAACCTCG | 0.64;1.23;1.62;1.68 |
| AP3B1 | 1.282 | 3.957 | 4 | GGAATTATCTAATTGGCCAG;GCAGACAGACCAAGCCATTG;TGCTATGAAGCGGATTGTTG;ACATGCTAACTCGATATGCT | 0.77;1.31;1.34;1.7 |
| PFKP | 1.266 | 3.556 | 4 | CGTCCCCGCCGATCACACAC;GCCGGATGATCAGATCCCAA;GAAGTACGCCTACCTCAACG;GATGTGTGTCAAACTCTCGG | 0.5;0.77;1.8;1.99 |
| PPP1R15B | 1.263 | 3.581 | 4 | AGGTGGACATCCCTGCCACG;ACGCATAGACAATTTCAGTG;CCCTCCTCTAGCCAATCAAG;GCACCTTCTGAAGCAATCCG | 0.58;0.91;1.38;2.18 |

| | | | | | |
|---|---|---|---|---|---|
| CHEK2 | 1.259 | 3.811 | 4 | GGGCCCATAATCGAGCCCAG;AGAGCTGTTTGACAAAGTGG;AGGTAAAGCTGGCTTTCGAG;GCATACATAGAAGATCACAG | 0.69;1.12;1.5;1.73 |
| HNRNPF | 1.259 | 3.619 | 4 | ACTCCCATTTGGATGCACAA;CCATTTCATCTACACTAGAG;TGAAGCCGTAGCCATCACTG;GATAGGGCACAGGTACATTG | 0.39;1.23;1.45;1.97 |
| SPCS1 | 1.258 | 3.246 | 4 | ACCGAGACTTAAGGACCGTG;TGGATTTATCTACGGGTACG;CTAGCTGAACAGATGTTTCA;GCTTCACCCCAGGATTACAA | -1.06;1.76;2.03;2.29 |
| SLC7A3 | 1.255 | 3.745 | 4 | ACCTGGTAGAAAGAGCACAG;GTACTTGCTCGGATCCACAC;CCCATCGGGATGGAACGCTG;AGATATGCCGAACCAGAACG | 0.53;1.31;1.36;1.82 |
| FAM155B | 1.250 | 3.825 | 4 | CAGATGGTCAGCGAGCAGCA;ACTGTATCGAGGCGTACCAG;AGGCGGAAGAGTACTCAATC;CAGCAAGTCCCAGACCGTGT | 0.75;1.09;1.42;1.74 |
| INTS12 | 1.249 | 3.304 | 4 | AACTTACCTACAAACAACGC;CTTGATGAATCTTTGGCTCG;CCGAGATTGTCATAAACCCC;GTTTCCTCGTCAGTAACTAG | 0.42;0.47;1.84;2.27 |
| ZNF699 | 1.227 | 3.787 | 4 | TTTGTGTTCTGTGAGCGATG;CAAAACTCACTGTACATGGG;AAGGGATGAATGATCCACGA;GTGAAGAGAGAACTTATCCA | 0.72;1.04;1.55;1.59 |
| NKTR | 1.223 | 3.534 | 4 | TGCTAAAAGGGAAAAACCTG;AGAGCATATAGACCACCTAG;GATGTGCGAGTTATTGACTG;AAACTGTAAGTGCTTTGGAT | 0.11;1.48;1.53;1.77 |
| GRIPAP1 | 1.220 | 3.288 | 4 | AGGAGGAACTTGCTAAGGTG;TGGCTGAGAACAATGCCTTG;GTTGACCCAGGAATTACAGG;GCTCAGGTACATTCCATGGA | 0.19;1.03;1.34;2.33 |
| SMYD5 | 1.209 | 3.750 | 4 | GTGGGTAGTGAATACTCCTG;CCATCTGGAGTGAACCACTG;AATGCACTTTATCGCTACCG;AGCCACTGAGCAATACCACC | 0.59;1.31;1.33;1.6 |
| ERP44 | 1.204 | 3.616 | 4 | AATGTTATTTCTCGGACAAG;GTGCTGATCACAATCAACTC;TCAGCGATCAGTGAAAGCAT;GCTCCGGATATGGTGTACTT | 0.54;1.13;1.32;1.83 |

| | | | | | |
|---|---|---|---|---|---|
| RAD21 | 1.187 | 2.359 | 4 | GTGTAATTTAGAGAGCAGCG;AAGTGTTGTTTGATCAGTCA;TCTGTTCAGACTCTAATAGG;ACATACTCTAAGTCAGGCAG | -0.59;-0.04;2.2;3.18 |
| OR7C2 | 1.187 | 3.485 | 4 | TGGAGAGGAACAGCCCATGG;GAAGGATCACAAAAAAAGTG;TGTGAAACAGATGTCAGCAA;GAGGTGGGAGTCTGAACTGA | 0.33;1.08;1.61;1.72 |
| MLF2 | 1.167 | 3.614 | 4 | CATTCCCAGCATCCCAAAGG;ATGAGCCGTATGTTGTCAGG;ACCATCACCCGTATTGGAGT;GTCCCGGATGTGATGCCCAA | 0.74;0.87;1.27;1.79 |
| YIPF6 | 1.167 | 3.631 | 4 | TGGAGGGCCCCAATTTGCAG;GTGAGATGGATATATCTGAA;ACAATGCGAGTGTCACACAA;GATGCGAGATCTCATAGGAA | 0.64;1.02;1.35;1.66 |
| AWAT1 | 1.154 | 3.587 | 4 | CATGAGGTACTCCCTAACAA;GAAGTTGCAGAAGGCGCCAA;AAGCACTGGTAGCGGCCACA;CAGATGGCAAGGTAGCTCAA | 0.52;1.17;1.33;1.6 |
| FOXO4 | 1.153 | 3.849 | 4 | GTAACAGGTCCTCGGAAGGG;CCTTGAAGTAGGGTACAGTA;GTTCATCAAGGTTCACAACG;TCAGTCAGCAGTTATGCAGG | 1.01;1.06;1.09;1.45 |
| SBNO2 | 1.117 | 3.512 | 4 | CGAGTGCGTCTACAACCGCG;GGGCTCCCCAGTGACGACCG;CAACGACCTCAAGTACGATG;GCGCCGGTTCGCCTCCATCG | 0.58;1.02;1.25;1.61 |
| RLIM | 1.111 | 3.616 | 4 | GTCACCTATGAAAGTGAACG;TCCCACCTACCAGAGGTCAG;CCCGGCACCATGTGACATTG;TCTTCAGACACAGCTGCCAG | 0.62;1.14;1.32;1.35 |
| CDKN2D | 1.110 | 3.282 | 4 | CAGATGGATTGGAAGTGCCC;GCAGCGCCGTCTTGCCGAAG;ATGGACTGGACTGGTACCGG;GGTGTCCAGGAATCCAGTGC | 0.2;1.08;1.34;1.82 |
| BOD1 | 1.110 | 3.347 | 4 | GGCAGCGACTGGCGCTACTG;ACCAGTTGCGAAATGGTCTG;GGAAGCTGTCAAAAAGGCCC;GTCAACACATCTGGACAAGC | 0.18;1.25;1.34;1.67 |
| INO80 | 1.105 | 2.519 | 4 | TTGCACCAGACTATATCCAA;GTTTCTCACCAATAGCCGAA;CAGAGACAAATCGATATAGG;GACCCCAATTCAGAACACCA | -1.91;0.49;2.84;3.01 |

| | | | | | |
|---|---|---|---|---|---|
| ARMCX6 | 1.105 | 3.430 | 4 | AGAGGAGTGGGACGATGACC;GAGCACACCCAATAAAACAG;TGCGTTTACAAACTGACCAT;TGGGGATTGGACTGAACCTG | 0.53;0.98;1.19;1.72 |
| CEBPD | 1.089 | 3.400 | 4 | CTTCTACGAACCGGGCCGGG;AACAGCAATCACAAGGCGGG;CACCACGGTCTGTGCGCACG;GGCGGCCATGGAGTCGATGT | 0.36;1.18;1.27;1.56 |
| UBE2L6 | 1.087 | 3.339 | 4 | AGCGTGCCACACCAGGACAT;GTTGTGAATTTGATCATGGG;CAAGATCTACCACCCCAACG;CCCCATGGATACTCACGGGT | 0.69;0.78;0.83;2.05 |
| MEMO1 | 1.085 | 3.088 | 4 | ACTCTTCAGTAAATATCTAG;GAAAGTGCACATCGAGAGAG;CTTTAGTTTACGGAGAACTG;AAATCTAGAACTTACCCCAA | 0.14;0.7;1.62;1.88 |
| RXRA | 1.082 | 3.171 | 4 | AGGAAGCCATGTTTCCTGAG;CCTACGTGGAGGCAAACATG;AGGACTGCCTGATTGACAAG;CAAGGACCGGAACGAGAATG | 0.4;0.88;0.9;2.15 |
| HSH2D | 1.081 | 3.354 | 4 | TGGGACTTTCATGATCCCCG;TCATATACATACCTCCTCAG;ACTCTCTTGAGATTGCACCA;ACGACAAGTTCACAGTGGCG | 0.34;1.0;1.48;1.5 |
| GLOD5 | 1.077 | 3.602 | 4 | TCTTCACCGTCATCACGATG;CCAGAAATTTAACCTCCACG;AAGTCTACGGATAAGACATG;TGTGATCAGACATATGTCCA | 0.78;1.04;1.05;1.44 |
| PCGF6 | 1.057 | 3.566 | 4 | GGGAGGCCGGCAGGACTCGG;GGTCTAGAAGTACCTAAACC;CCTCACCCCTGCACCCGCAG;GGTATGAAGACATTCTGTGA | 0.8;0.9;1.18;1.35 |
| VSIG4 | 1.056 | 3.374 | 4 | ATCCAGCAGGCAAAGTACCA;GTGAAGTGGCTGGTACAACG;AGGCTAATCCTCATTCCCTG;ACTACACGTGTGAAGTCACC | 0.53;0.93;1.22;1.55 |
| EDA | 1.050 | 3.114 | 4 | CCAGCTGTGGTGCATCTACA;ACAACTGTTATGGGACCACC;GCAACTCCGAGCGCAACTCT;GCTTAGGTGACGGCTGCCCA | 0.31;0.78;1.16;1.94 |
| DAND5 | 1.048 | 2.984 | 4 | GCTGCAGCGTGGGCAAGACG;CAGATGATTTCGGAGGCGTA;AACCCTCAGGAAGTGATCCA;GGCAGAAGCTGGCACCAGTG | 0.02;0.67;1.72;1.78 |

| | | | | | |
|---|---|---|---|---|---|
| DHX29 | 1.041 | 2.924 | 4 | CCTTCAATATTTCTGAATTG;AATTCTACAAATGATTCTAG;ACCAATACACTCTTATATGG;GATTCTAATGAGTGCCACTG | -1.17;1.36;1.84;2.13 |
| NLGN3 | 1.038 | 3.213 | 4 | TCTTCGGCAAAAGAGTGCCA;GGACTTAGCGGATAATGACG;AGTGATGTGACAACGTGAGG;GACCCGCCGTAAAACACTGG | 0.29;1.02;1.16;1.68 |
| LMO4 | 1.024 | 3.475 | 4 | TTCAGGTTATTTGGAAATAG;CTCATGACGAGTTCACTCGC;GCTGTGCCAATAGCTGTCCA;ACGTCCTGTTACACCAAAAG | 0.64;1.05;1.13;1.28 |
| TCP11X2 | 1.022 | 3.213 | 4 | AAACACACATCCCTACCTCT;TCATTAGGCCCAGAACACGG;ATGGCGTGTTCACTTCCAAG;ACATAGTGTAGTGATGTCTG | 0.46;0.69;1.38;1.56 |
| CREBBP | 1.022 | 3.416 | 4 | CTTAGCCCACTGATGAACGA;CAGGACGGTACTTACGTCTG;CCGCAAATGACTGGTCACGC;ATTGCCCCCCTCCAAACACG | 0.57;0.97;1.27;1.28 |
| NHSL2 | 1.014 | 3.540 | 4 | ACCAGTGAGACACTGCGTGT;GGTTATCACGAGCGACAAGG;AGGAGTTCTCAGTACTGGCG;TCCAGTCAGTAAGATACCAG | 0.81;0.96;1.07;1.21 |
| ERAS | 1.008 | 3.249 | 4 | TTGTTGCCCACGAGGACAAG;CTGGAAGGAAGGGCTCCATG;GAACCACCAGTGCTTCGTGG;AGGAGTTGACCCTGGACAGT | 0.63;0.72;0.96;1.72 |
| ASF1B | 1.007 | 3.285 | 4 | ATCCCAGAGACTGATGCCGT;ATTTGATCAGATCCTAGACT;GAGTGGAAGATCATTTATGT;TGGACAGGAGTTCATCCGAG | 0.48;0.89;1.21;1.45 |
| NO_CURRENT_31 | 1.006 | 2.040 | 4 | ACAGACAACCATAATAGATG;ACACTGAGGACAGAATAACG;ACAGCCCTCACGAGCCCGAA;ACAGCCAGAAAGAGTACTCG | -0.72;0.13;1.12;3.49 |
| PDAP1 | 1.006 | 3.395 | 4 | ACTGGATCTGGACGGGCCAA;AAAAGCGCAAAGGCGTTGAA;CATCGAGAACCCCAACCGGG;GAGGCAGTATACAAGCCCTG | 0.55;1.09;1.09;1.29 |
| HTR7 | 1.004 | 2.903 | 4 | TGAGCCCATCCAAAGAGTGG;GTCAATGCTGATCACGCACA;GGGGATATAAAATGCCACTG;CAGCCGGAGGCATTGTCCGG | 0.06;0.61;1.47;1.87 |
