## Supplementary material for "Illuminating host-mycobacterial interactions with functional genomic screening to inhibit mycobacterial pathogenesis": Table S3

**Table S3. Positive genetic hits from the genome-wide CRISPRi screen in *M. bovis* BCG infection. Related to Figure 2.**

| Gene Symbol | Average LFC | Average -log(p-values) | Number of perturbations | Perturbations | Individual LFCs |
|---|---|---|---|---|---|
| YPEL5 | 13.294 | 7.760 | 3 | GCGGGACTCACCCTGAGCGG;CTGAGCGGCGGTGAGATCCG;CCCGCGGGACTCACCCTGAG | 11.99;13.18;14.71 |
| TYK2 | 12.352 | 7.454 | 3 | AGGTCCTCAGGAAGAAGCCG;GTCCTCAGGAAGAAGCCGCG;TCAAGCGCAGCCAGTCCCCG | 10.53;12.62;13.91 |
| WDR26 | 9.178 | 6.131 | 3 | CGGAGCGGATCCGAGGACAG;AGCGGATCCGAGGACAGAGG;CAACACACCAGGGAGCAGAG | 4.12;11.34;12.08 |
| IFNAR2 | 8.537 | 6.473 | 3 | CCGGCGGGTAGGAATCCCGC;GACGCTTCTTCCCGGCGGGT;CTAAAGACGCTTCTTCCCGG | 7.34;8.06;10.21 |
| ARNT | 8.452 | 6.455 | 3 | GCGGCGACTACTGCCAACCC;GGTGGCATCTGCGGCCATGG;GACTACTGCCAACCCCGGTG | 7.48;7.49;10.39 |
| IRF9 | 7.215 | 3.950 | 3 | CCAAGGATGGGAGTCAGCCT;AACTCAGGTAAGATCAGCCA;AGGTAAGATCAGCCAAGGAT | 0.36;8.62;12.67 |
| IFNAR1 | 6.994 | 5.008 | 3 | TGCGCGAACATGTAACTGGT;GTGCGCGAACATGTAACTGG;GTAACTGGTGGGATCTGCGG | 2.48;3.4;15.1 |
| NNT | 6.875 | 6.119 | 3 | TGTGCCCTGAGGGCCGCGAG;ACGTCCTCTCGCGGCCCTCA;GCCAGCGCGACGTCCTCTCG | 6.6;6.98;7.04 |
| STAG2 | 6.461 | 5.605 | 3 | CCATCATCTCCAGCGCGTGG;ACGCGCTGGAGATGATGGGG;GCCCCATCATCTCCAGCGCG | 3.86;7.37;8.15 |
| SERF2 | 5.966 | 5.737 | 3 | GTGCTCACGGGTCATGGCGA;GGGACGTAACGGAGGCAGGT;AGAAAAGGGCAAGGAGAAGG | 5.29;5.91;6.69 |
| UBE2H | 5.966 | 5.793 | 3 | GGAGCCGGTCAGAGGGTGAG;CGCTTATAGTGGAGGCTGCT;AGCCGGTCAGAGGGTGAGTG | 5.5;5.88;6.52 |
| AHR | 5.423 | 5.314 | 3 | CTGCCCCTGTTCTGAGCTGG;CGGACCGCCAGCTCAGAACA;TCTGTTCCGAGAGCGTGCCC | 3.61;6.21;6.44 |
| KDM4A | 5.270 | 5.099 | 3 | AAAGCAGTAGGATCGGCCAG;CAGGAGCTGAGCCTAAGCCC;TCGCCTGCTGGAAAAGCAGT | 3.8;3.96;8.04 |
| SUPT7L | 5.078 | 5.372 | 3 | GAATGCCCAAGTCCCTCCAG;TGACCGAGGAAACCCCTGGA;GTGACCGAGGAAACCCCTGG | 4.45;5.25;5.53 |

| | | | | | |
|---|---|---|---|---|---|
| SCTR | 5.010 | 2.156 | 3 | GTCCTGATCAATGGGGCGGG;CTGCTCCTCCTCGGACCAGG;TCCGAGGAGGAGCAGTCCCG | 0.1;0.7;14.23 |
| PHIP | 5.000 | 5.008 | 3 | CACACTGACAGCTATAGGGC;CGCCGCCGTTGCTTGAATGG;TGAATGGTGGAGCCGAAGCT | 3.2;4.92;6.87 |
| CTNNBL1 | 4.856 | 5.224 | 3 | GGGCCGGGTACCATGGACGT;CAGAAGTTCGCCCACGTCCA;AGGGAAGTGGAGTATTTGCT | 4.08;4.98;5.51 |
| TP73 | 4.792 | 5.201 | 3 | AAGGGGACGCAGCGAAACCG;CCGCCCGCGCACCCGCCCGG;CGGGCGCGCGAGCCTCCGGG | 4.12;4.93;5.33 |
| TMEM70 | 4.754 | 5.071 | 3 | GAAGCCGTGTCTCGCAGTCG;GAGTCCACGACTGCGAGACA;GTCGTGGACTCGTGCAGCTG | 3.27;5.35;5.64 |
| PHF6 | 4.750 | 5.161 | 3 | TCCAGCAGTGCCTGAGAGCG;GAGCGGGGCTCTGTCGCCGG;GAATGAAAGGAAACAACCTC | 4.01;4.83;5.4 |
| MCTS1 | 4.717 | 3.506 | 3 | GAAAGAAATTGAATTTGCAG;GGAAGGCCGGCACTGACAGT;CCGGCAGATCCCCTCACACG | 0.38;6.28;7.49 |
| STAT2 | 4.675 | 5.069 | 3 | CTGCAACCCTAATCAGGTAC;AATCAGGTACGGGCCCTGAG;ACTGCAACCCTAATCAGGTA | 3.44;5.24;5.35 |
| UCHL5 | 4.571 | 5.099 | 3 | TGAGAGCGAGAGGTGGATCG;GAGCGAGAGGTGGATCGGGG;GGGCAGCTGAGAGCGAGAGG | 4.03;4.49;5.19 |
| INO80B | 4.555 | 5.071 | 3 | GCGGCGTGGGAGCACCTCTG;GCCGCCACAGCTTACTCATG;AGGACCTCATGAGTAAGCTG | 3.92;4.33;5.41 |
| NONO | 4.210 | 4.856 | 3 | TACTCCGAGGAGATACCAGT;CTGGTATCTCCTCGGAGTAA;TCTACCGACTGGTATCTCCT | 3.43;4.07;5.12 |
| GNAI2 | 4.192 | 4.962 | 3 | CTCCCACCACCAGCCCTCTG;GACCCGAGTGCTTCCCGCAG;GCCCTCTGCGGGAAGCACTC | 3.97;4.19;4.41 |
| NFRKB | 4.184 | 3.905 | 3 | CGCGGCCGGAGAAGGGCTGC;GCCCGCCCTCACCTGAACCG;GCCCGCGGTTCAGGTGAGGG | 1.18;4.43;6.94 |
| BAX | 4.149 | 4.800 | 3 | GGCGGTGATGGACGGGTCCG;GGCGGCGGGAGCGGCGGTGA;GGGTCCGGGGAGCAGCCCAG | 3.38;3.72;5.35 |
| UBA7 | 4.090 | 4.876 | 3 | TGTTTGCACTGGCTACAGCA;CTGCTGTAGCCAGTGCAAAC;GCCAGTGCAAACAGGAACCA | 3.62;4.04;4.6 |
| CHMP6 | 4.019 | 3.549 | 3 | TGACGCGGCTCTGCTTCTTG;GCGGCCGAACAGGTTACCCA;GCCGAACAGGTTACCCATGG | 1.41;1.83;8.82 |

| | | | | | |
|---|---|---|---|---|---|
| LARP4B | 3.868 | 4.719 | 3 | GTTGGGGATCTGGAGTCCCG;GCCCGAAAATGTCTCAACCC;GCGAACGGGAAGGAGCGTTG | 3.41;3.58;4.62 |
| FAM214B | 3.764 | 4.550 | 3 | GGAACTACAGGTGTGGGGCA;CCCACGGAACTACAGGTGTG;GACCCACGGAACTACAGGTG | 3.05;3.26;4.98 |
| NIPBL | 3.745 | 4.625 | 3 | GGCGGGGGCGTCCCTCCCTG;GACACCACCAAGCCGCAGGG;CCCGACACCACCAAGCCGCA | 3.15;3.64;4.44 |
| CBLL1 | 3.741 | 4.681 | 3 | CATGGATCACACTGGTAAGG;GGATCACACTGGTAAGGAGG;AGCCGAATCATGGATCACAC | 3.37;3.61;4.24 |
| USP7 | 3.586 | 4.034 | 3 | GAGCCCCGACGACGACGCCG;CCTCCTCGGCGTCGTCGTCG;CCCCGACGACGACGCCGAGG | 2.02;2.56;6.19 |
| ACO2 | 3.497 | 3.363 | 3 | AATGGCGCCCTACAGCCTAC;CCTACAGCCTACTGGTGACT;CCTGCAGCCGAGTCACCAGT | 0.67;4.32;5.49 |
| ATP5L | 3.381 | 3.224 | 3 | GTTACGGACAAATTGGGCCA;GACATTCAGCCGGCGGTTCG;GACTCTCCATTCCAGAACCA | 0.88;2.46;6.8 |
| C14orf2 | 3.359 | 3.084 | 3 | CCTGCGCCAAGGTGAGTACG;TCACCTTGGCGCAGGACAGA;CCAGGTCCCCGTACTCACCT | 0.44;3.83;5.8 |
| PCGF6 | 3.331 | 4.425 | 3 | GTCGGGAGAGACACCAGGCG;GGTGTCTCTCCCGACCATGG;GAGAGACACCAGGCGAGGCG | 2.99;3.21;3.79 |
| RBM6 | 3.328 | 2.566 | 3 | CCGTGGAGGCTTCGCCGCCT;GTCCCGGGCCCTTACCTAGG;CCTAGGCGGCGAAGCCTCCA | 0.58;1.26;8.15 |
| NDUFB9 | 3.310 | 4.171 | 3 | GGGGAAGGTCAGCGCCGTAA;GCTGACCTTCCCCGGCCGCG;CAGTTTCCCGGCTCTCCGCG | 2.18;3.37;4.38 |
| PCGF1 | 3.194 | 3.340 | 3 | TCTTGTACACTGACTGGAGC;GCCAGATTGCGATCGCGATG;GGTCCATCTTGTACACTGAC | 0.76;4.1;4.72 |
| OXSM | 3.177 | 4.361 | 3 | CGACTGTGGAGAAGTGTCCG;GTAGCCAGCGATACCTGTAA;AGCCAGCGATACCTGTAACG | 2.75;3.38;3.4 |
| UBE2L6 | 3.018 | 4.099 | 3 | GGCGAGCATGCGAGTGGTGA;CATGATGGCGAGCATGCGAG;GAAAGGGGTCATCCTATACG | 2.23;3.28;3.55 |
| PINK1 | 3.012 | 1.613 | 3 | GCACCGCCATGGTGGCGCCG;GGGCCGCGGCGCCACCATGG;CGCCTGTCGCACCGCCATGG | -0.74;0.73;9.05 |
| HSH2D | 3.011 | 4.076 | 3 | CGAGGGCCTCGGCCATCCAA;CCCAAGAGCCTCAGCCCTCG;GGCTCTTGGGTCACATACCT | 2.38;2.61;4.04 |

| | | | | | |
|---|---|---|---|---|---|
| ATP5J | 3.004 | 4.173 | 3 | CTGAGTGCAAGGTGACTGGG;GCAGCTCGGGACTGAGTGCA;TGAGTGCAAGGTGACTGGGA | 2.46;3.21;3.33 |
| UBE2E1 | 2.982 | 4.031 | 3 | CTGGCTCCTCCGTGCCGCCG;GCTCCTCCGTGCCGCCGCGG;AGCCGCAGCAGCAGCCGCCG | 2.04;3.36;3.54 |
| SLC7A11 | 2.980 | 4.185 | 3 | GCAGTGGTGGAACGAGGAGG;GCAGCAGTGGTGGAACGAGG;GCAGCAGCAGTGGTGGAACG | 2.55;3.02;3.37 |
| UCHL3 | 2.972 | 1.293 | 3 | CGGCCATGGAGGGTCAACGC;GCTGGAGGCCAATCCCGAGG;GCCGCTGGAGGCCAATCCCG | -0.56;-0.31;9.79 |
| SMC1A | 2.891 | 1.412 | 3 | GCGGCCTGTCCTACTGCCGC;TCCTACTGCCGCCGGCGCCG;CAGTTTCAGGAACCCCATGA | -0.39;0.16;8.9 |
| RAD21 | 2.878 | 3.039 | 3 | TTACCGAACAAGCGATGATG;TACCGAACAAGCGATGATGC;GATGCGGGCCCAAGCTGCAT | 1.21;1.34;6.08 |
| MCMBP | 2.820 | 4.018 | 3 | TGTCGCCAGCAGCTCCTCGG;AGCTGCTGGCGACAGTCAGG;GCTGCTGGCGACAGTCAGGC | 2.36;2.61;3.49 |
| NDUFC1 | 2.786 | 3.565 | 3 | AGGGGACGCAGCAAGGCGGA;GGAAAGGGGACGCAGCAAGG;ACGGCCGCTCGGGAGCCTGG | 1.57;2.26;4.53 |
| MAEA | 2.786 | 2.982 | 3 | GTCCATGACCCTGAAGGTCC;CAAGATGGCGGTGCAGGAGT;GATGGCGGTGCAGGAGTCGG | 0.68;2.6;5.08 |
| MAF1 | 2.775 | 1.351 | 3 | CGCTCGGCTCGCTGACTCGC;TGACTCGCCGGAGCGCTCTG;TGTCCGGCCGGGGGAGGCGG | -0.32;-0.18;8.82 |
| PAXIP1 | 2.763 | 3.701 | 3 | GGAGGTCAAGTATTACGCGG;CAGGGAGGTCAAGTATTACG;TCCTGAGGAGATGTTCAGGG | 1.71;2.55;4.03 |
| CREBBP | 2.762 | 3.942 | 3 | ATCGCGCTCGAAGCCCCGGT;GTAGATCGCGCTCGAAGCCC;TCTTGTCGCGGACCCGACCG | 2.22;2.62;3.45 |
| CHMP5 | 2.742 | 3.662 | 3 | CTTCGGGAAAGCGAAACCCA;CAGTCAGTCAGGCTGGGCGG;GTTTCGCTTTCCCGAAGAGT | 1.61;2.62;4.0 |
| MAP2K6 | 2.677 | 3.876 | 3 | TTCCCTAACGTTGCAACTGG;CTTCCCTAACGTTGCAACTG;CCCAGTTGCAACGTTAGGGA | 2.27;2.35;3.41 |
| KPNB1 | 2.637 | 3.641 | 3 | CAAATGGGCTGCTGGCGGCG;CCCTCCAAATGGGCTGCTGG;GCGGCGGGGGAGGGACCCTG | 1.57;2.76;3.58 |
| SPINT1 | 2.624 | 3.735 | 3 | CCAGCACCCTCGGAACCCAG;GCCATCGCCTTCCTCCCCAG;CATCGCCTTCCTCCCCAGGG | 1.78;2.74;3.35 |

| | | | | | |
|---|---|---|---|---|---|
| PLAGL2 | 2.614 | 2.705 | 3 | GCCGCCACTACCGGGGCCTG;CCCGACCGCCGCCACTACCG;TCAGGCCCCGGTAGTGGCGG | 0.25;3.26;4.33 |
| IDH2 | 2.567 | 2.948 | 3 | GGTAGCCGGCCATCCCAAGC;AAGCTGGAGAGCGAACGAGC;TGGAGAGCGAACGAGCAGGG | 0.54;3.58;3.59 |
| PIGY | 2.469 | 3.061 | 3 | GGCTGCCAGACCATGCTGAG;GCAGCGTGCTCCACTCAGCA;CGGGTGGCGAGGCTGCGGTG | 0.84;2.68;3.89 |
| PYURF | 2.469 | 3.061 | 3 | GGCTGCCAGACCATGCTGAG;GCAGCGTGCTCCACTCAGCA;CGGGTGGCGAGGCTGCGGTG | 0.84;2.68;3.89 |
| CHEK2 | 2.418 | 3.227 | 3 | GAGAGTGTGCGGCTCCAGGT;CGGCTCCAGGTGGGCTCACG;GGAGAGTGTGCGGCTCCAGG | 1.05;2.87;3.34 |
| JAK1 | 2.418 | 3.690 | 3 | CCTGGAGCTGCAGACAGTGC;GGACAGCCGGGACTGGGCGC;TGCAGCTCCAGGATACTCCG | 1.87;2.61;2.78 |
| IRAK1 | 2.410 | 3.535 | 3 | GCGCCCGCCGCGGCCCTGAG;CCCGGCAGGTCCCGGCCCGG;GGACCTGCCGGGGCCTCTCA | 1.79;2.11;3.33 |
| PTCD1 | 2.408 | 3.658 | 3 | AGTCCAAACAGCCTTACCTG;GTAAGGCTGTTTGGACTCCG;AGGCTGTTTGGACTCCGTGG | 2.01;2.23;2.98 |
| ATP5J2 | 2.408 | 3.658 | 3 | AGTCCAAACAGCCTTACCTG;GTAAGGCTGTTTGGACTCCG;AGGCTGTTTGGACTCCGTGG | 2.01;2.23;2.98 |
| ATP5J2-PTCD1 | 2.408 | 3.658 | 3 | AGTCCAAACAGCCTTACCTG;GTAAGGCTGTTTGGACTCCG;AGGCTGTTTGGACTCCGTGG | 2.01;2.23;2.98 |
| NUP153 | 2.402 | 3.503 | 3 | AGCAGGGGTAATAGGCGCAA;CAACGGCGGCGGAGGCTGCA;TGCAGGGACGACGACGACGA | 1.68;2.21;3.31 |
| TRAF6 | 2.385 | 3.307 | 3 | GCGGCCAGCGAAGGTGGCGA;GGAGCCTTCGCCACCTTCGC;AGCAGTGGCGTCCGCAGCTG | 1.3;2.37;3.48 |
| MTPN | 2.371 | 1.553 | 3 | AACCTCTCTGCTGGCCCGG;CCCAGCAGAGAGGTTCCGCC;GGCCACCGGGCCCAGCAGAG | 0.13;0.14;6.84 |
| GID8 | 2.353 | 1.765 | 3 | CTGCGCGGTCTGGGCGGCTG;CCCGCGCTGCGCGGTCTGGG;ACCTCTCCCCAGCCCAGGAG | -0.54;1.21;6.39 |
| SEMA3G | 2.328 | 1.229 | 3 | GGGCCATGCTGGGGAACTGA;CCGGCGGCCAGCGGGACCAG;CGGCCAGCGGGACCAGAGGG | -0.82;0.07;7.74 |
| NDUFS4 | 2.290 | 3.204 | 3 | GGCGGCGGTGTCAATGTCAG;TGGTACTGAGGCAGACGTTG;TACTGAGGCAGACGTTGTGG | 1.09;2.7;3.08 |

| | | | | | |
|---|---|---|---|---|---|
| CCNC | 2.272 | 3.445 | 3 | GCTCCTCGATCAAATCAGCT;GCTGATTTGATCGAGGAGCG;GAGCCGAGCTGATTTGATCG | 1.67;2.15;3.0 |
| PLD6 | 2.250 | 3.649 | 3 | GCCCGGGTTCAGGCTCTCAG;TCCCCTGAGAGCCTGAACCC;TCAGGGGAGCTGGCAGAACC | 2.17;2.19;2.39 |
| FLCN | 2.250 | 3.649 | 3 | GCCCGGGTTCAGGCTCTCAG;TCCCCTGAGAGCCTGAACCC;TCAGGGGAGCTGGCAGAACC | 2.17;2.19;2.39 |
| ZRSR2 | 2.237 | 3.057 | 3 | TGACGTTTCCCGAGAAACCA;AAACCAAGGTAAGCGCCGTA;ACCAAGGTAAGCGCCGTACG | 1.03;2.32;3.36 |
| NENF | 2.227 | 1.477 | 3 | CCATGGTGGGCCCCGCGCCG;ACCAGGGCCAGCGCTGCCAG;GCGCTGCGCGCTCACCATGG | -0.95;0.75;6.89 |
| CSDE1 | 2.208 | 3.412 | 3 | TTATGGCGGCGCTGGAGAGG;CTACCCCCAGTTGGACCCTG;TTGGGGTGAGTACGACCTCA | 1.78;1.86;2.98 |
| PAXBP1 | 2.191 | 2.864 | 3 | GCGAGTATTCGACCGCCGTG;GAATACTCGCTTCCACACCG;GGCCTTTCGGAACATCCCCG | 1.11;1.45;4.01 |
| TRERF1 | 2.186 | 2.643 | 3 | TTGGAGTTGCTTGCGGCGGA;GAGTTTGGAGTTGCTTGCGG;GGAGAGTTTGGAGTTGCTTG | 0.47;2.54;3.55 |
| CDKN2C | 2.184 | 3.412 | 3 | CCTGGTTAGGAGCAAAGGAA;CTCCTAACCAGGCTGCATGA;ACCAGGCTGCATGATGGCAT | 1.54;2.5;2.52 |
| BIK | 2.178 | 2.203 | 3 | CGGAGCAGCGGGCGGCAGCA;CGGGGGCACTCACCTGGCGG;GCAGCGGGCGGCAGCAGGGT | 0.52;1.05;4.96 |
| NCKAP1L | 2.173 | 3.015 | 3 | AGCAGATGTCAAAGACATGA;GTCTGGTGCTCCTCTCTCAG;GACATGATGGCCACTGAGAG | 0.94;2.53;3.05 |
| NDUFA8 | 2.171 | 3.389 | 3 | GACGGGGGCGACGCGGCTGA;GCGGTCGAAAGGGGAGTTCA;TGGCACTCGGCGGTCGAAAG | 1.65;2.05;2.81 |
| DNAJC30 | 2.165 | 2.314 | 3 | TGGAGGTTGCTGCAGGCCCG;GGCAGCGGCTGTTACCTTGG;CGGGCCTGCAGCAACCTCCA | 0.16;2.06;4.27 |
| RAD23A | 2.161 | 2.843 | 3 | CGTGAGTTGCATGTTGTGTG;GCGGGGCCCGAGCGACGCGG;CATGTTGTGTGAGGATCCCG | 0.68;2.69;3.1 |
| ARIH2 | 2.156 | 3.369 | 3 | GCTGCCGCTTCGCCCCAATC;GGGACCGGATTGGGGCGAAG;AGCGCGGGCTTGACCGGCGT | 1.51;2.37;2.58 |
| ATP5E | 2.149 | 2.340 | 3 | GGGCCGAATCGCCAAGACGC;CCACCATGCTGTAGCGAAAG;GTAGCGAAAGCGGAGCTCGT | -0.04;2.64;3.84 |

| | | | | | |
|---|---|---|---|---|---|
| RBM42 | 2.119 | 3.403 | 3 | GGGAGCCCACCCGGACGAAG;AGAGTAGACAGCAGAACCAG;GTAGACAGCAGAACCAGCGG | 1.66;2.26;2.44 |
| KAT2A | 2.113 | 3.143 | 3 | GGCGCCGCTCTCCGCTGCGG;GCCTCCCCCGCAGCGGAGAG;CGCCATGGCCTCCCCCGCAG | 1.13;2.55;2.65 |
| MXI1 | 2.106 | 3.110 | 3 | ACACACAACTGCAGGCAGCG;ACACTCACACACACAACTGC;CTGGAGCAGAGAAGGCGCCG | 1.27;2.01;3.04 |
| BCL2L11 | 2.101 | 3.329 | 3 | GCTCCAACAAACTGCAGACC;CTGCAGACCAGGCGGCTGCG;CCGCCTGGTCTGCAGTTTGT | 1.49;2.36;2.46 |
| LAMTOR 5 | 2.081 | 2.507 | 3 | ATCCCACCCACCGACCACTC;GGGTTCTGAGCCGGAGTGGT;CACTCCGGCTCAGAACCCAG | 0.91;1.07;4.27 |
| NUFIP2 | 2.048 | 3.123 | 3 | GTCCCAGCCGCTTTCAATGG;CTCCTCCATTGAAAGCGGCT;GGCTTCTCCTCCATTGAAAG | 1.19;2.39;2.57 |
| DENR | 2.045 | 3.073 | 3 | GGTTCAGCTGGCGATAGCGG;GTTGGTTCAGCTGGCGATAG;GTTGCATGTGTTGGTTCAGC | 1.44;1.53;3.16 |
| ATP5C1 | 2.034 | 2.722 | 3 | GGCTACCATGTTCTCTCGCG;GACACCCGCGCGAGAGAACA;GCTACCATGTTCTCTCGCGC | 0.98;1.45;3.68 |
| SSTR5 | 2.014 | 2.714 | 3 | GGCTGGGAACAGGGGCTCCA;TGGGAACAGGGGCTCCATGG;GCAGAGCCTGACGCACCCCA | 0.79;1.96;3.3 |
| ATAD1 | 2.010 | 2.963 | 3 | CGGGAACCAGCTACTCACCC;GGGAACCAGCTACTCACCCA;GGAACCAGCTACTCACCCAG | 1.11;1.95;2.97 |
| UTY | 2.001 | 2.681 | 3 | CAAAGTTGTAATTGTGTCAC;ACACAATTACAACTTTGTGC;GAGGTCGCGGCGGCTGCGTG | 0.67;2.25;3.08 |
| A4GALT | 1.983 | 3.248 | 3 | CTAGCTCCAGCGGCGGCGGG;CGTACCTCTAGCTCCAGCGG;CCTCGTACCTCTAGCTCCAG | 1.67;1.77;2.51 |
| MIPEP | 1.981 | 3.175 | 3 | AGCAGGGCAGGGATCTGCGT;CGCTCCCAGCGAAAGCAGCA;GTTGGAGGAAGGGACTGCTC | 1.42;2.05;2.48 |
| TRPC4AP | 1.978 | 2.842 | 3 | CCGACGAGGAGACATGGCGG;AGGAGACATGGCGGCGGCGC;TGTCCGACGAGGAGACATGG | 0.94;2.02;2.97 |
| TADA1 | 1.974 | 2.098 | 3 | GACCTTTGTGAGCGAGCTGG;CGCCTCCAGCTCGCTCACAA;GGCCAAGAAGAACTTAAGCG | 0.66;0.69;4.58 |
| KPNA6 | 1.968 | 3.306 | 3 | CGCCGCCAAGAGTGAGCGAG;CTTGGCGGCGGAGGCAACGG;TGAGCGAGCGGACCCGCGAT | 1.66;2.12;2.12 |

| | | | | | |
|---|---|---|---|---|---|
| HSPA4 | 1.966 | 2.824 | 3 | CGGTTGCAGTACCCACTGGA;AGCGCCTAAGTCCTTCCAGT;GAGCGCCTAAGTCCTTCCAG | 1.01;1.78;3.12 |
| FSTL3 | 1.960 | 2.215 | 3 | CCATGCGTCCCGGGGCGCCA;TGCGTTCGCCATGCGTCCCG;GCCCAAGCCAGGGCCCCCCA | 0.15;1.91;3.82 |
| EP400 | 1.960 | 3.209 | 3 | GGCCGCGTCAGGAGGGCGGG;ATCCTCCCGCCCTCCTGACG;GAGCGCAGCCCTGAGGCCCA | 1.51;1.96;2.41 |
| BAHD1 | 1.943 | 2.161 | 3 | GCGTAGCGGATCCCGGAGCC;GGGAACCGCGAGCGGCGTAG;GCGAGCGGCGTAGCGGATCC | -0.86;3.18;3.51 |
| DMAC2 | 1.937 | 3.228 | 3 | ATTTGGGGCTCTAGGAACTC;ACTCCGGTCACTTACCGCCC;CTCCGGTCACTTACCGCCCA | 1.68;1.73;2.4 |
| SLC15A1 | 1.936 | 1.322 | 3 | GCCGAGCGTACCCATGGCGG;CTGCCAGGAGCACGTCCCGC;CCCGCCGAGCGTACCCATGG | -0.26;0.23;5.84 |
| SNTA1 | 1.930 | 1.193 | 3 | GGCCAGCCGCCACCCTACCC;GACCAAGCGCCCAGGGCAGA;CCCCAGCGGCCGGGGTAGGG | -1.92;0.16;7.55 |
| ARF1 | 1.927 | 3.108 | 3 | AGCCTCTGAGGTGAGGGCGA;GAGCAGCAGCCTCTGAGGTG;GCCCCTCGCCCTCACCTCAG | 1.34;2.14;2.3 |
| KPNA1 | 1.924 | 3.332 | 3 | GAGTCTCGCTGCTCGGTGCG;GTGCGAGGCGGCGGAGAGCG;CTGAGCTGAGTCTCGCTGCT | 1.86;1.9;2.01 |
| HELZ2 | 1.913 | 3.018 | 3 | TGAGAGCTCCTGGGCAGGCT;AGCTCCTGGGCAGGCTCGGC;GCAGGGCAGGCAGCTCCAGG | 1.26;1.91;2.57 |
| MGA | 1.910 | 3.235 | 3 | CCGCCCCCAGCGGGATGTGA;AGTCCTTCACATCCCGCTGG;ACAGAATAACCCGCCCCCAG | 1.65;1.9;2.18 |
| SUV39H1 | 1.902 | 3.194 | 3 | AAGCAGTGGAGGCAGAGGCG;AGCAGTGGAGGCAGAGGCGC;TTTAAAAGGTGAAGCAGTGG | 1.66;1.7;2.35 |
| FAM47B | 1.891 | 2.058 | 3 | GGAGAGGTGGCACAACAGAG;AACAGAGAGGGCCACCATGG;TGGCCTCCGGTCCCCCATGG | 0.53;0.87;4.27 |
| PRR14L | 1.882 | 2.967 | 3 | ACGGAGCCGACAGAGGCCGG;TCCGTAGGCTCAGGTAGACC;TGAGCCTACGGAGCCGACAG | 1.14;2.15;2.35 |
| ZFC3H1 | 1.879 | 2.891 | 3 | GGGCGCTAGGAAAGGGAACT;GAAAGGGAACTGGGTTGCGA;GGTCCGGCGAGAGAGAGCTG | 1.05;2.05;2.54 |
| RGS14 | 1.869 | 2.736 | 3 | GCCGGCCAGAGCTTGCACGG;CGCGGGGGCTGCCCACCATG;GCTGCCCACAGTCCCCATGG | 0.82;2.15;2.64 |

| | | | | | |
|---|---|---|---|---|---|
| LAMTOR2 | 1.865 | 2.988 | 3 | AGCCCAGGGTTAGGAACCGT;GATCTGGGTGCAAAAGCCCA;TGCCTACGGTTCCTAACCCT | 1.36;1.61;2.63 |
| BABAM2 | 1.853 | 2.953 | 3 | CCTGTACCCCACGTAGTCGC;CGGTAACCGGCGACTACGTG;TCGGTAACCGGCGACTACGT | 1.18;2.0;2.38 |
| APEH | 1.852 | 3.235 | 3 | GCCTCGCCCCGGCGGCAGAG;AGACTATGGAACGTCAGGTG;CCCGGGCTGCAGGGACCACG | 1.75;1.84;1.97 |
| OTUB1 | 1.852 | 2.304 | 3 | CCGCTGGGCAGCGACTCCGA;CCTCCTTGGATCTGTACCTT;CCTTCGGAGTCGCTGCCCAG | 0.77;0.99;3.79 |
| PRR16 | 1.838 | 1.420 | 3 | GCGCTCCGCGCTCGGCCGGG;GCTCCCTGGGCAGCGGCGAG;CAGCCACTCGCCGCTGCCCA | 0.03;0.19;5.3 |
| ATP5O | 1.820 | 2.606 | 3 | GGGTTTGACCTACAGCCGCC;GGCAGCCATCTTCTCCCGGG;GGAGAGCCCGGACACTGCTG | 0.75;1.88;2.84 |
| RPS6KA4 | 1.819 | 2.235 | 3 | CAGCGACCCGCCCGCCATGG;GTCGTCCTCGTCCCCCATGG;ATCGTCGTCCTCGTCCCCCA | -0.06;2.68;2.85 |
| GPR162 | 1.815 | 1.139 | 3 | GGGGTCGTCTTACCTCTCTC;CGTCGGGAGTGCGGTCTCCA;TGCTGGGCGCAGCCGGAGAG | -0.58;-0.54;6.56 |
| INTS10 | 1.806 | 2.317 | 3 | GGCGGCGGCGGTGGCTGCCG;GGCGGCTGAGAGTCCAGAGC;CTCTGGACTCTCAGCCGCCA | 0.12;2.63;2.66 |
| IMMP2L | 1.799 | 3.093 | 3 | AGCTGCGCAGCGGGTGCGGA;GAGCTGCGCAGCGGGTGCGG;GGCGTCCGGCGGCGGGTGGG | 1.51;1.79;2.1 |
| ATP5F1 | 1.798 | 1.483 | 3 | GGCGGCGGAAAGTACCACCC;GTTGACCATGCTGTCCCGGG;AAGTACCACCCGGGACAGCA | -0.3;0.73;4.97 |
| ITGB2 | 1.789 | 3.138 | 3 | CGGTGTGCTGGAGTCCTCGG;CCCAGCCAGGAGCACCGCCG;CCGAGGACTCCAGCACACCG | 1.66;1.7;2.01 |
| TRIM25 | 1.789 | 2.541 | 3 | GCAGTTGTGTCCCGACCCCT;CTCTGCCATGGCGCTCCCAG;GCTCTGCCATGGCGCTCCCA | 0.72;1.77;2.88 |
| MAP3K4 | 1.787 | 2.037 | 3 | CGCCGTTTCCAAGATGGCCG;CTCCTGCGGCGGGGTAGAGG;CGGCGGGGTAGAGGCGGAGG | -0.41;2.33;3.43 |
| BABAM1 | 1.786 | 3.082 | 3 | ATGGCGGCTTCCTAGTGAGT;TCGGCGGCTGATTTAGAAGG;GCGGCTTCCTAGTGAGTCGG | 1.46;1.9;2.0 |
| PTGER3 | 1.786 | 1.965 | 3 | GGAAGGCGTGGCTCCCTCCC;CAGAGGTTTCCCAGAGAGGA;GTTTCCCAGAGAGGAAGGCG | 0.11;1.39;3.86 |

| | | | | | |
|---|---|---|---|---|---|
| IFIT1 | 1.780 | 2.540 | 3 | CTAAGCAAAACCCTGCAGAA;AATTAGGCAGCCGTTCTGCA;ACAGAATAGCCAGATCTCAG | 0.77;1.63;2.94 |
| GIGYF2 | 1.767 | 2.674 | 3 | ACGGAACGGCTGGTTACCTG;GTTACCTGCGGTCCGGGACA;GTTGAGGCTGAGGACTGACT | 0.92;1.7;2.68 |
| RHBDD2 | 1.756 | 3.005 | 3 | GCTGCGACACCCGGGCCCCG;GGGAACGACGGCGGCCATGG;AAGCACCAGCTGCGACACCC | 1.39;1.84;2.03 |
| MAU2 | 1.739 | 2.492 | 3 | CAAAATGGCGGCTCAGGCGG;GGCCCAGGCTGCGCAGGCCG;GGCCAAAATGGCGGCTCAGG | 0.77;1.51;2.93 |
| MTIF2 | 1.733 | 1.859 | 3 | GCTGTAGGTAGGAGTTAAGT;TAACTCCTACCTACAGCCCC;CGCCCGTGCTCAGAATCCAG | -0.13;1.45;3.88 |
| GMEB1 | 1.731 | 2.409 | 3 | GGGCGGGCGGCGAGCGGCCA;GTCTCCGTCGGGCGGGCGGG;GCCCGCCCGCCCGACGGAGA | 0.48;2.09;2.62 |
| SLFN5 | 1.729 | 2.695 | 3 | ATTACCCGCTGGGTTCTCCT;GCTCCGTTGCCTGCTCCCGG;CGCTGCCGAGGAGAACCCAG | 0.97;1.68;2.53 |
| NDUFA12 | 1.726 | 2.861 | 3 | TTAGTGCAGGTCCTGAAACG;CAGCAGATCACCGGCCACGG;GATAGCCTCGGAGACCGCCG | 1.29;1.55;2.33 |
| SLC35C2 | 1.720 | 1.345 | 3 | GCGCACTCCCAGAGGCTGGG;GCCTCTGGGAGTGCGCCGGG;GCCTCCCGGCGCACTCCCAG | -0.09;0.25;5.0 |
| ACACA | 1.717 | 2.236 | 3 | TGCGTGGAGGAACTGCTGGG;CCAGCAGTTCCTCCACGCAG;GCGATTCCGGAGCCCCTGCG | 0.12;2.41;2.62 |
| UQCRB | 1.704 | 1.997 | 3 | ACGGTGAAGCGCGACGATGC;AGCAGGCCGGTAAGTAACTG;GCAGGCCGGTAAGTAACTGG | -0.42;2.3;3.23 |
| NDUFAF4 | 1.695 | 2.452 | 3 | TGCGCCGGTGTTCCCACGTG;CTCAGGTTCAGGCCGCACGT;GCATAATGTTATGAGGAGAT | 0.56;2.1;2.42 |
| NFIL3 | 1.695 | 2.895 | 3 | CGCCGGGTCCGCGGGCGCCG;CCGCGCGGCCACGGCGCCCG;GGCGGGCGCGCCGGGTCCGC | 1.35;1.6;2.13 |
| ZDHHC17 | 1.685 | 2.693 | 3 | CACGGTTCCTCACCTCTGGG;GGATGAGTACGATACCGAAG;TGTGCCCCTTCTCCACCCAG | 0.9;2.05;2.11 |
| ZC3H18 | 1.685 | 2.918 | 3 | TTGGCCCTTCCGCTCTTCAC;TCGACACGGGCGGAACAAAC;GGCTCCAGGCTCGACACGGG | 1.46;1.46;2.14 |
| WBP1L | 1.683 | 2.862 | 3 | GAGCGTCAAACAGGAAAAGA;AGCGTCAAACAGGAAAAGAA;GGGAGGTGGCGGCGCCGTGG | 1.34;1.53;2.18 |

| | | | | | |
|---|---|---|---|---|---|
| NPY1R | 1.681 | 1.637 | 3 | CCTGCTTATTAAAGAAGGAA;CTGAGCAGGAGAAATACCAG;GCTTATTAAAGAAGGAAGGG | -0.65;1.27;4.42 |
| TRMT61A | 1.681 | 2.868 | 3 | GCGTCGCAGGTGAGACTCCG;CGGAGGCACGTGTGGAGCCC;GCGCGTCGCGGAGGCACGTG | 1.29;1.67;2.08 |
| RASA4B | 1.676 | 1.555 | 3 | GAAGGAGAGTAAGACAGAGG;GAGAGGGAGGAGAGTAGGAG;GAGAGTAGGAGGGGGAGAGA | 0.05;0.62;4.36 |
| NAA15 | 1.673 | 2.661 | 3 | CCGACGGAGACCCGTAGTGG;GCCGACGGAGACCCGTAGTG;GTTACCACGTGAACCGCCGA | 0.9;1.93;2.2 |
| S1PR3 | 1.667 | 1.283 | 3 | ACAGGGACTCAGGGACCAGA;GGTTCCGGGGACGGGCAACA;GGGACTCAGGGACCAGAAGG | -0.64;0.52;5.12 |
| NDUFB6 | 1.662 | 2.451 | 3 | AGTAACTAGTCCGTAGTTCG;GGTACCAACGCAAAAGGACA;GTAACTAGTCCGTAGTTCGA | 0.58;2.1;2.31 |
| METTL3 | 1.661 | 2.949 | 3 | TATATCCTGGAGCGAGTGCT;GGATATAGCCAATTCTCACG;ATCCTGGAGCGAGTGCTGGG | 1.48;1.55;1.96 |
| GNA13 | 1.659 | 1.708 | 3 | CGCCCTGCGAGTCAGTTCGC;GGCCCGAGCGCGCCCAGGGA;CCGAGCGCGCCCAGGGAGGG | -0.62;1.44;4.16 |
| KLHDC10 | 1.659 | 2.165 | 3 | CCTGTCTCCTGGGTCTCTGG;GCCTGACCCAGCGGAACCAG;CCCGTCAGCGCCTGACCCAG | 0.53;1.26;3.18 |
| LOXL4 | 1.648 | 2.498 | 3 | GGGGCTCACCAGATGGAGCG;GATGGAGCGCGGACAGTCCG;GCGGACAGTCCGGGGCTGGG | 1.04;1.09;2.82 |
| SSB | 1.647 | 2.351 | 3 | CTGTTTGTGAGCCTGTGGCG;GCCCACAGAAGCCGCGCCAC;GGCTGGCCGGCGGCGCTGGG | 0.5;1.93;2.51 |
| FUNDC1 | 1.640 | 1.359 | 3 | CTGGCGGTATCATGGCGACC;ACCCCCCTCCCCAAGGTGAG;TGAGCGGCCAGGGCCCGAAG | -0.19;0.45;4.66 |
| FNIP1 | 1.635 | 2.483 | 3 | GGGGGTGGCGGGGCGGCTGT;GCGGGGCGGCTGTAGGAGCA;GGGGCCTAGCAAGCGCCCAG | 0.72;1.79;2.4 |
| ZBTB14 | 1.628 | 2.029 | 3 | TGGAGTTTGGGGAGGGGGCG;GAAATCGTGCACGTCCCTGA;GGAGGAGGGCTGCTACCATC | -0.16;2.21;2.83 |
| EP300 | 1.625 | 1.521 | 3 | TGTTTGTGTGCTAGGCTGGG;AGCCTAGCACACAAACAAGA;CTAGCACACAAACAAGATGG | -0.22;0.84;4.26 |
| DNTTIP1 | 1.625 | 2.635 | 3 | AGAGTCCAGCGGAGTTGTGG;CAGAGTCCAGCGGAGTTGTG;GGCGTCGCCAGTGGCTCCCA | 0.94;1.78;2.16 |

| | | | | | |
|---|---|---|---|---|---|
| RPAP3 | 1.621 | 2.507 | 3 | TTACGGCCTGGTCAGGTGAG;CAACGCCCGCTCACCTGACC;CAGTGCGGCGGGTTACGGCC | 0.74;1.86;2.26 |
| UQCRQ | 1.619 | 1.872 | 3 | GTCTGGTTGTGCAGTGTTCG;CTGAGCCCTGTGCGTGAGTG;CCAGACCCCACTCACGCACA | -0.12;1.55;3.42 |
| STT3B | 1.618 | 2.187 | 3 | GAGAGCTAGACCCGCCGCCG;TCCGCCATGTTGTGCCCCGG;GCCGCCGGGGCACAACATGG | 0.5;1.4;2.95 |
| HRCT1 | 1.613 | 2.089 | 3 | CTGTCCGGCTCCCAGATGCT;CTGGGCAGAGAGGGACTGTC;GAGGCCCAGCATCTGGGAGC | 0.47;1.22;3.15 |
| MAPK14 | 1.611 | 1.880 | 3 | AGTCGGCCTTGTAGGGGCGA;GCCCGGAGTCGGCCTTGTAG;ACTGCGACGCAGCCCGGAGT | -0.46;1.96;3.33 |
| IQGAP2 | 1.606 | 1.420 | 3 | TTTACTTCAGTGTCAGCGCG;GGCGACCGGCCAGGGAGCGA;GACCGGCCAGGGAGCGAGGG | -0.43;0.76;4.5 |
| CDK3 | 1.606 | 1.319 | 3 | GGCTGGGCAGTGACCCAGGT;AAGACCACCCTTTCCCACCT;GTGACCCAGGTGGGAAAGGG | -0.19;0.37;4.64 |
| BOD1 | 1.604 | 2.760 | 3 | AACCTCTGCTGTATCAGAGG;GGCCCCCTCTGATACAGCAG;CAACCTCTGCTGTATCAGAG | 1.32;1.38;2.11 |
| RAB11A | 1.604 | 2.458 | 3 | GGAGCAGCAGTGGTATCTGT;AGTGGTATCTGTGGGACCAG;GTGGTATCTGTGGGACCAGG | 0.69;1.84;2.28 |
| PARD3B | 1.603 | 1.345 | 3 | CAGGGCCGGCCCCAAGCCGG;CCACTCCAGCCCCCGGCTTG;CGGCCCTGCCAACGCCGCCA | -0.17;0.42;4.56 |
| NDUFA1 | 1.603 | 2.105 | 3 | ACATCTCTGCCCCGTTACCT;TAGGTAACGGGGCAGAGATG;CTTTCAAGGACCCAGAAGTA | -0.04;2.42;2.43 |
| BAK1 | 1.603 | 2.806 | 3 | ACTTGTCCCTGTGCAGCCCG;CTTGTCCCTGTGCAGCCCGA;AGTGATCATAGAGGTGCCAG | 1.35;1.45;2.01 |
| SNTG2 | 1.603 | 1.308 | 3 | GGCCGGCCGAGCTCAACGCC;GTCCTGGCGTTGAGCTCGGC;GACCCCGTCCGAGCCTCCCA | -0.03;0.12;4.72 |
| FOSL2 | 1.601 | 2.630 | 3 | CGAACGAGCGGCGCTCGGCG;AGCGAACGAGCGGCGCTCGG;TCATCTCGGGCAGAGCGCTA | 1.14;1.32;2.34 |
| NDUFV1 | 1.598 | 1.850 | 3 | CCGTGTTGCCAGCATCGCGG;CCGCGATGCTGGCAACACGG;CCGCCGTGTTGCCAGCATCG | -0.53;1.88;3.44 |
| SMAD4 | 1.598 | 2.643 | 3 | CGCAGCGGCGACGACGACCA;AAACCGCTCCGTTACCGCAG;ACAAGTTGGCAGCAACAACA | 0.93;1.91;1.96 |

| | | | | | |
|---|---|---|---|---|---|
| KRBOX4 | 1.586 | 2.482 | 3 | GCGGTGAAGTAAGAGGGACG;GTGAAGTAAGAGGGACGCGG;CCAAGAGCCTGAAACCTCAG | 0.81;1.61;2.34 |
| ACTR5 | 1.586 | 2.631 | 3 | GCACTGGGTCCGGTGCGGCA;CTCCAGCACTGGGTCCGGTG;CGCACCGGACCCAGTGCTGG | 1.07;1.47;2.22 |
| CHUK | 1.583 | 2.816 | 3 | AACCGGCCTTGGAACAACTG;TGGAACAACTGTGGAACCTG;CAGGTTCCACAGTTGTTCCA | 1.27;1.73;1.75 |
| ZC3H10 | 1.581 | 2.486 | 3 | AGGGAGGCGGGAGACTTAGG;GGCGGCAGCAGCAACTACGG;GCAGGCGGCAGCAGCAACTA | 0.94;1.33;2.47 |
| PAQR8 | 1.574 | 1.719 | 3 | CAGCCCGCGCTCCCGCCTCG;AGCTCGTGGCCGGGACCCCG;TCGTGGCCGGGACCCCGAGG | 0.2;0.83;3.69 |
| PDE6G | 1.570 | 1.300 | 3 | ACTACTCACCAAGTGCAGGG;GGCTGCCAGGAAAGACAGCG;GTGGCTGCCAGGAAAGACAG | -0.06;0.17;4.6 |
| NF1 | 1.569 | 2.754 | 3 | GTCTCGGACTGTGATGGCTG;CTGTGATGGCTGTGGGGAGA;CTCGGACTGTGATGGCTGTG | 1.22;1.58;1.91 |
| MMP19 | 1.565 | 1.728 | 3 | TCCCAGAAATCTCAGGTCAG;GAAATCTCAGGTCAGAGGCA;TGCCTAGCACTGCTCCCCCA | 0.01;1.08;3.6 |
| CCDC6 | 1.563 | 2.683 | 3 | TGGGCGCCGGGCGAGCACAG;AGGCCGGGCTGCGAATGAGT;AAGGCCGGGCTGCGAATGAG | 1.21;1.41;2.08 |
| TNPO3 | 1.562 | 2.459 | 3 | CTTCCTCACTGTCTGGGCCA;CAGACAGTGAGGAAGCGCGA;CGGCCGTGGCCCAGACAGTG | 0.97;1.21;2.5 |
| ASH2L | 1.552 | 2.710 | 3 | AGGGGTGGCCGTGATGGCGG;GCGGCGGCAGGAGCAGGACC;AGCAGGACCTGGCCAGGAAG | 1.12;1.72;1.82 |
| SPPL3 | 1.550 | 2.423 | 3 | CGGCGACTGAGGGGGGCGCT;GAGGGGGGCGCTAGGCGATG;GGGCGCTAGGCGATGAGGCG | 0.93;1.21;2.51 |
| CAMK2N1 | 1.550 | 2.594 | 3 | GGAAGGGAGAAAAACAGAGG;GGAAGAGGGTCGGAGAGGGA;AGGGTCGGAGAGGGAAGGAA | 1.13;1.31;2.21 |
| FBN2 | 1.547 | 2.281 | 3 | CCAGCAAGAGAGAGGGCGGG;GCGTGCGGCCAGCAAGAGAG;CGGCCAGCAAGAGAGAGGGC | 0.51;1.77;2.35 |
| ANP32B | 1.541 | 2.708 | 3 | CCCCGGAGCCAAGTTACTCA;AGCCAAGTTACTCACGGGAG;CTCCGCTCCCGTGAGTAACT | 1.17;1.59;1.86 |
| SGK3 | 1.537 | 1.210 | 3 | CTGCTGCGCCACACTCGGCC;GGCCGAGTGTGGCGCAGCAG;GCCGCTGGGAGCTGGTGGCG | -0.32;0.23;4.7 |

| | | | | | |
|---|---|---|---|---|---|
| MYO5A | 1.535 | 2.225 | 3 | TACGCCCCCCGCCTGTGCGG;GAGCTCCGACGCAGCCATGG;GCCTACGCCCCCCGCCTGTG | 0.4;1.91;2.3 |
| RALGAPB | 1.534 | 2.401 | 3 | CGAAGGCCACTCGAGCCAGC;AGGGCGCCAAGGGACGACGA;ACCCGTCAGCGACTCTCCCA | 0.73;1.63;2.25 |
| PROSER1 | 1.533 | 1.976 | 3 | TGTAGGTCCCCATGAGCTGG;GTAGGTCCCCATGAGCTGGT;ACAGCCCCCACCAGCTCATG | 0.58;0.83;3.19 |
| ELOC | 1.533 | 1.444 | 3 | AGGGGCTGTGGCAGTACGCG;TAGGGGCTGTGGCAGTACGC;CTGCCACAGCCCCTATCCCA | 0.13;0.33;4.14 |
| RPRD1B | 1.532 | 2.740 | 3 | TACCGACTGACAGACTGCCA;ACCGACTGACAGACTGCCAG;ACTGCCAGGGGTGCGAGAAG | 1.22;1.63;1.75 |
| CCAR2 | 1.530 | 2.764 | 3 | GCCAGGGCAGCGACACCGGG;TCCGGGTGTCTTTGTCCCCC;ACCGGGGGGACAAAGACACC | 1.35;1.43;1.8 |
| UHMK1 | 1.529 | 1.534 | 3 | AGTCCCGGGAGTCGGTGAGG;TGAAGTCCCGGGAGTCGGTG;CGGCTGCAGGTCCCTCCCTG | 0.15;0.53;3.91 |
| BRCC3 | 1.525 | 2.511 | 3 | GGCCAAGATGGCGGTGCAGG;GAGCGTGGTTGAGACAAACG;GGTGGTGCAGGCGGTGCAGG | 0.92;1.51;2.14 |
| MIER3 | 1.524 | 2.781 | 3 | CAATATGGCGGAGGTGAGCA;CGGAGGTGAGCAGGGAAACC;TAAAGGTACCAATATGGCGG | 1.37;1.46;1.74 |
| RHBDF2 | 1.516 | 2.199 | 3 | CCCCGGGACGCAACTCCGTG;CCGCACGGAGTTGCGTCCCG;GCCCCGCGCTCACCTCCCTG | 0.6;1.32;2.62 |
| TRMT6 | 1.515 | 2.422 | 3 | GGCGGTGGCGACAACCGAGG;GCTGGCGGTGGCGACAACCG;TGGCGACAACCGAGGAGGAG | 0.7;1.85;1.99 |
| LRPPRC | 1.514 | 2.055 | 3 | GGAGCGTGCTTCCCGCTGCG;TTGCTCGAACGTCCCCGCAG;CCAACGCGCGGATCTCAGCA | 0.57;1.04;2.94 |
| DNAJA2 | 1.512 | 2.506 | 3 | GTCGGGCCCACAAGCGGCGT;GGGCCCACAAGCGGCGTCGG;AGCGGAGTCGGGCCCACAAG | 0.84;1.76;1.94 |
| GRK7 | 1.509 | 1.517 | 3 | GAAAGCACAAGAGGGCTGGA;ACTCCCAGGGAAAGCACAAG;GAGCACGGGGCGCACTCCCA | 0.18;0.47;3.88 |
| ZNF717 | 1.508 | 1.169 | 3 | GAAGAGGAACCCGTGGGCCC;CCGGGATCCCCCGGGCCCAC;CCCGGGATCCCCCGGGCCCA | -0.9;0.39;5.03 |
| CIPC | 1.507 | 1.079 | 3 | CGGCCCTAGGTGAGAAAGGG;CCCCGGCCCTAGGTGAGAAA;GGCCCTAGGTGAGAAAGGGA | -0.84;0.09;5.28 |

| | | | | | |
|---|---|---|---|---|---|
| PITPNB | 1.504 | 2.800 | 3 | TATCGGCGGCAGCTGTGAGG;GTATCGGCGGCAGCTGTGAG;GGCGGCGGCGGTGGTATCGG | 1.41;1.47;1.63 |
| BLOC1S5 | 1.500 | 2.331 | 3 | GGAGGGACAGAGACCCCTGT;GGCGGCCTCACAACCCACAG;CGGAGGGACAGAGACCCCTG | 0.58;1.89;2.03 |
| PIAS1 | 1.493 | 2.614 | 3 | TCGAATTCACTTCTAATATT;GGCGGACAGTGCGGAACTAA;TCACTTCTAATATTCGGCCG | 1.08;1.54;1.86 |
| ATF4 | 1.490 | 2.430 | 3 | GAGAAAACTACATCTGTGGG;CGGCCACCATGGCGTATTAG;CTGCCCCTAATACGCCATGG | 0.73;1.83;1.91 |
| MBD4 | 1.484 | 1.664 | 3 | GCAACGCCCAGGGTGTGGGG;GCCCCACACCCTGGGCGTTG;GCCCAGCGCCGCAACGCCCA | -0.22;1.22;3.45 |
| NO_CURRENT_7 | 1.484 | 2.027 | 3 | AAAGCAGAACCCAGTCTCTG;AAAGATATAGCAAATTATGG;AAAGCGACGTAGGCATACTT | 0.11;1.89;2.45 |
| OGFR | 1.483 | 2.683 | 3 | CCCCGACTGCGACTCCACCT;CTCCTCGTCCTCCTCCCAGG;CGACTGCGACTCCACCTGGG | 1.17;1.64;1.64 |
| ABCF2 | 1.479 | 2.420 | 3 | GGCCCAGTGAGTTAGGGGAA;AGCGAGGCCCAGTGAGTTAG;AGGCCCAGTGAGTTAGGGGA | 0.82;1.54;2.08 |
| SLC7A5 | 1.475 | 1.847 | 3 | CCTGGGACACCCGGGAGCCG;GCACACTGCTCGCTGGGCCG;ACCCGCCATGCTCTGCGCAC | -1.15;2.12;3.46 |
| LENG8 | 1.472 | 2.426 | 3 | CGTGTTGCTGATCGCCTGGG;GAAGACCAGAGCCAGCCGGG;GAGCCAGCCGGGTGGCACAG | 1.03;1.13;2.26 |
| DNAJA1 | 1.471 | 2.240 | 3 | CGGAGGAGCGGTAACTACCC;GAACGCTCGGTGAGAGGCGG;CCAGAACGCTCGGTGAGAGG | 0.52;1.75;2.15 |
| RBM26 | 1.467 | 2.216 | 3 | GGCAAAAGTAAAGAAAGCGA;CTCAATGGGAACGATTCAGG;GCTCAATGGGAACGATTCAG | 0.62;1.41;2.38 |
| NDUFS7 | 1.462 | 2.164 | 3 | CCCACGGCCCCAGACCCGCG;GCGGTGCTGTCAGGTGAGCG;TCAGGTGAGCGCGGCACCGG | 0.65;1.18;2.55 |
| NDUFAF2 | 1.451 | 2.072 | 3 | GGAAACTGGGTGGACGGCAT;TGGCAGCGCTGGAAACTGGG;ACTGGGTGGACGGCATGGGT | 0.2;1.98;2.18 |
| SALL2 | 1.447 | 1.785 | 3 | ATGGATATTGGGATTGAGGG;AGAGGCTGCCGCAGACCCAG;GCGGCAGCCTCTGCACCCAG | -0.2;1.55;2.99 |
| SRRM1 | 1.439 | 1.870 | 3 | AGTAGAAGCGCCGGGCGCCG;AGAAGTCCCTGGCCTCACCC;CGAGGCGGCAAGATGGACGC | -0.46;2.17;2.61 |

| | | | | | |
|---|---|---|---|---|---|
| CWC15 | 1.435 | 2.495 | 3 | GAGCCTGCTGCATCGGACCT;CGGACCTCGGCCAGTGTGAG;CGGCCAGTGTGAGCGGCAAG | 0.9;1.7;1.7 |
| EMC7 | 1.432 | 1.570 | 3 | GTCATGGCGGCCGCTCTGTG;GCTGCTGCTGCTGCTATCGG;GGGAAAGAAGCCCCACAGAG | 0.24;0.54;3.52 |
| UQCC1 | 1.432 | 2.008 | 3 | ACGGTAGCTCTCTGGGTCCC;AAATTACTCCTTCACTCACA;GTGCGAGTCCTTGTGAGTGA | 0.31;1.46;2.53 |
| LGALS9B | 1.431 | 1.801 | 3 | CTCCTGGGTGGCAGGAGACA;GCGGTGGAGATGGCCTTCAG;AAAGGCAGCGGTGGCCACAG | -0.3;1.72;2.88 |
| ECSIT | 1.429 | 1.528 | 3 | GGCTGGCCGCGTGAGTAGGT;GCGCGACCTACCTACTCACG;CAGAGCTGGCCTGGAGTCCG | -0.59;1.15;3.73 |
| UBE2E2 | 1.423 | 2.336 | 3 | GCTTCACTTTCCAGGACTCA;AAAGTGAAGCCACCACCTCC;CTTCACTTTCCAGGACTCAG | 0.83;1.35;2.09 |
| RNF31 | 1.421 | 2.151 | 3 | CGGGGGCTGGAGAGTGACCG;GCCCCAGGTCACTCAGACCA;GTGGTCTGAGTGACCTGGGG | 0.64;1.24;2.38 |
| EBI3 | 1.418 | 1.859 | 3 | CATACCTTTCCTTCCACTGC;AGGGCGGGCAGCTGGCCCAG;GCAGCTGGCCCAGAGGACAA | 0.58;0.65;3.02 |
| RBM48 | 1.414 | 1.623 | 3 | ATTTGATCACCACGTCCAGA;GAGGGCGGTATGCGACACAC;TGATCACCACGTCCAGAGGG | -0.18;1.15;3.27 |
| LUC7L2 | 1.409 | 2.556 | 3 | CTTCTGTCCCAAGAACCGGA;TTCTTGGGACAGAAGCGACA;AAGAACCGGACGGAGAGTGA | 1.2;1.28;1.75 |
| TBC1D24 | 1.409 | 2.680 | 3 | AGAAAGCGGCGCGCGGAGGT;CGCGGAGGTGGGTGCGCTCG;GAGGAGAAAGCGGCGCGCGG | 1.36;1.4;1.47 |
| NDUFA13 | 1.408 | 1.717 | 3 | ATGGGCCCATAGCCCCCCGG;CAGGACATGCCTCCGCCGGG;TCGATGGGCCCATAGCCCCC | -0.13;1.34;3.01 |
| NDUFB4 | 1.404 | 1.939 | 3 | GGGTCTCAGGCAGAGTGCGC;GGCTCGACGGCTTATACTTT;GGCAGAGTGCGCAGGCTCGA | 0.66;0.68;2.87 |
| RRAGC | 1.403 | 2.275 | 3 | CTGGCCTGTCAGGGCGCGGG;CGGCCTGGCCTGGCCTGTCA;GGCCTGGCCTGTCAGGGCGC | 0.63;1.67;1.91 |
| SIAH3 | 1.403 | 1.067 | 3 | TGCGCTCCTTCCAGGGCCCA;GTGAGTCCTTGGGCCCTGGA;GCTGGAGCGGAGTGTCTCAG | -0.84;0.16;4.89 |
| BBC3 | 1.399 | 2.242 | 3 | GTGGCCGCTGCTGGGATCGC;ACATGCGAGCGGGCGCCTGG;GGCACCAGCGATCCCAGCAG | 0.62;1.58;1.99 |

| | | | | | |
|---|---|---|---|---|---|
| FAM193B | 1.395 | 1.691 | 3 | CAGGACAACAGCCAATATGG;GAGGGGAGCGGTAGCGGCGG;TGTGAGGGGAGCGGTAGCGG | -0.61;1.6;3.19 |
| NFE2 | 1.395 | 1.782 | 3 | CAGCGCTCAGCAATGGCCCG;CCATTGCTGAGCGCTGAGTG;CCACACTCAGCGCTCAGCAA | 0.27;0.98;2.93 |
| SNRNP25 | 1.394 | 2.036 | 3 | GCGGGCAGAGCCCGGCTGAG;GGGCAGAGCCCGGCTGAGAG;GCTGAGAGGGGCGGCCCTGG | 0.61;1.01;2.56 |
| ZNF628 | 1.392 | 2.570 | 3 | GAGAGGCCGCCGATAAAGGT;AGGGGCGCACTCCCTCCCAA;AGAGGCCGCCGATAAAGGTT | 1.11;1.52;1.54 |
| LMO4 | 1.391 | 2.095 | 3 | GCAGGAGCCCTCGGCCCGCG;CGAGCAGCTGCAGGAGCCCT;CGGCTTCAGGCGCGGCGCAG | 0.33;1.83;2.01 |
| PDE12 | 1.389 | 2.286 | 3 | ATCAGCGGCCTACTGTCAGG;CCTACTGTCAGGTGGAGCTG;CTAGCGGTCGACAAGACCCG | 0.89;1.12;2.15 |
| TMEM255B | 1.388 | 1.892 | 3 | GGCCCCGGGAGAGCCGGGTG;GCGGGGTCCAGCAGGCCCAG;GCACCGGCGGCTGCATCCCG | 0.08;1.59;2.5 |
| B3GLCT | 1.385 | 1.762 | 3 | GGTCAGCCGCGGCGGCAGGG;CCCTACCTGGGCGCGGGGAA;GCCCTACCTGGGCGCGGGGA | 0.23;1.0;2.92 |
| SIAH2 | 1.376 | 2.034 | 3 | GCCGGGAGGAGGGCACCGCG;TCCCGGGCCCCGAACCCCAG;GCGCGGTGCCCTCCTCCCGG | 0.2;1.96;1.97 |
| ZBTB24 | 1.366 | 2.166 | 3 | GTGCGGCGCAGAAGCCCCAG;CTCCGCCCCGCCCGCCTCTG;CGCAGGAGCAGAAACCGGTG | 0.8;1.0;2.29 |
| NDRG1 | 1.362 | 1.695 | 3 | ATCCGCCCTCGGCTTCCGCG;GTCAGTTCACCATCCGCCCT;CGCCCCGCGGAAGCCGAGGG | 0.1;1.06;2.93 |
| KIF2B | 1.361 | 2.436 | 3 | GGAGCGCTCCAGGGCCCTTG;TCCTGCTGCTCCAACCCCAA;GTCCTGCTGCTCCAACCCCA | 1.04;1.32;1.73 |
| RALY | 1.359 | 2.050 | 3 | GGAACCCAGAGAAGCTGAGG;GCGGAACCCAGAGAAGCTGA;CGGAACCCAGAGAAGCTGAG | 0.73;0.87;2.48 |
| KHSRP | 1.359 | 2.529 | 3 | GGCGGCTCAACGCGGGAACA;GGAGGCTGAAGCTGAGGAGG;TGGCGCGGAGGCTGAAGCTG | 1.17;1.35;1.56 |
| TAX1BP1 | 1.358 | 2.324 | 3 | AGAGGTTCGGCGGCTGATGG;ATCAAGGGAGAAAGTTACGC;TGGAAAGACTCCCGGATCAA | 0.79;1.49;1.79 |
| ZFR | 1.355 | 2.397 | 3 | ACAGGGCATATGGGAATCAT;CACCTATGGTGAGTCTAATG;GGCCACATTAGACTCACCAT | 1.04;1.2;1.83 |

| | | | | | |
|---|---|---|---|---|---|
| DIAPH3 | 1.353 | 1.380 | 3 | TGTGCTGACTGTTTGGGTGG;GGCTGTGCTGACTGTTTGGG;GGCTGCGGCCCCGACTTCAG | 0.07;0.38;3.61 |
| ZNF598 | 1.353 | 1.116 | 3 | CGGCGCCACGACCGGCCGAG;CGGCCGGTCGTGGCGCCGAG;GGCCGGATCCCGGACCATGG | -0.46;0.2;4.33 |
| ABHD3 | 1.352 | 1.934 | 3 | GAGGAGAGCCGGCTGGCGAG;TCCTGCGGCGGGAGGAGAGC;CGGGCGAGAGCGGGCGAGAG | 0.18;1.59;2.28 |
| ST3GAL4 | 1.351 | 2.264 | 3 | GCGGCGCAGCGCTGGCGCGA;CGCGACGGCTCGACTCGGCC;GCTGGCGCGACGGCTCGACT | 0.92;1.07;2.07 |
| COX16 | 1.351 | 1.828 | 3 | ATCGGCTCAGAACTCCAAGC;CATCGGCTCAGAACTCCAAG;GAGATTTGGGAGTCTGCGCT | -0.09;1.68;2.46 |
| AHSA1 | 1.350 | 2.128 | 3 | CGGTCCTGAGGCTGTGGCTA;GCTGGCACTAAGCGGTCCTG;GCCGGGCGGCTGGCACTAAG | 0.83;0.88;2.34 |
| UQCRC2 | 1.350 | 1.264 | 3 | ATGAAGCTACTAACCAGAGC;TTCAAGATTGTTCTGACACA;TTCTGACACACGGTCACTGC | -0.19;0.4;3.84 |
| ANKH | 1.349 | 1.476 | 3 | AAAAAAAAGAGGAGGGACGG;GATCTGCCGGGAAAAAAAAG;AAAAAGAGGAGGGACGGCGG | -0.11;0.81;3.35 |
| TTLL12 | 1.345 | 1.828 | 3 | TGGCGGCGCTGGAGTCGGCG;CTGGCGCCATGGAGGCCGAG;CGCGGGTGCTGGCGCCATGG | 0.41;0.92;2.71 |
| MED13 | 1.344 | 1.941 | 3 | TCCCTCACAGCAGCCGCCGC;GAGGCGGCGGTAATGGCGGA;ATGGTGGGTTGTGGCGCCGG | 0.1;1.85;2.08 |
| TBRG1 | 1.342 | 1.925 | 3 | CAGAGACACCCGCAACCGAA;CGTTCCCAGGGCTGACCGCA;AGCTCTCCGGGACGTTCCCA | 0.23;1.48;2.31 |
| WWTR1 | 1.341 | 1.917 | 3 | CGCAGGAGAAGCAGACAGCG;GCTGGGCGAAAAGGAGGCGC;GCTAGTGCTGGGCGAAAAGG | 0.41;1.15;2.47 |
| MPC1L | 1.339 | 2.093 | 3 | GGCAGCCACGCGAAGGCTCG;AGCGTGTGCCCAGTCAGGGT;ACCCGCAGCCGACCCTGACT | 0.47;1.55;1.99 |
| RNPEPL1 | 1.334 | 1.162 | 3 | CGGCACTAGGTGAAATCCAT;TAGGTGAAATCCATGGGCGG;CACTAGGTGAAATCCATGGG | -0.44;0.34;4.11 |
| BCAR1 | 1.329 | 1.616 | 3 | GGACCGGCTCTGGGTGGCCG;CCGCCTCGGCCACCCAGAGC;GCGGCTGGACCGGCTCTGGG | -0.35;1.34;3.0 |
| RBM10 | 1.328 | 2.255 | 3 | CACTGCCGCCTACTCCCAGC;TGTGGCCGGCTGGGAGTAGG;AGGCGGCAGTGAGTTTCCCT | 0.81;1.31;1.87 |

| | | | | | |
|---|---|---|---|---|---|
| BCORL1 | 1.326 | 2.375 | 3 | GGAGAGACAGAGCCACACAC;CCGCGGCTGCTGCCAGTGTG;GGGAGGGCTCAGCAAAGCGG | 1.0;1.25;1.72 |
| HTATSF1 | 1.325 | 2.060 | 3 | CGGCTGCGTCGGCCTGAGCA;TTCTAAACTAAGCCCTGCTC;GCCGACAATGGCGGCTGCGT | 0.37;1.69;1.91 |
| AHCTF1 | 1.323 | 2.409 | 3 | AGGGGAAGGACCCAGAAGCG;CGAGCAGGCAGCGTTGCAAG;CAGGCAGCGTTGCAAGGGGA | 1.07;1.22;1.68 |
| PRMT3 | 1.322 | 1.536 | 3 | TAGCGTCAGGCGCTACCGGT;GCCTGACGCTAACGAGCACA;TGAGGGGCCAGGGTACCCAC | 0.27;0.5;3.19 |
| PSME2 | 1.319 | 2.353 | 3 | AGCAGCATGGCCAAGCCGTG;GTGTGGGGTGCGCCTGAGCG;CAGCATGGCCAAGCCGTGTG | 0.93;1.36;1.67 |
| PPP2R5E | 1.316 | 1.587 | 3 | GTGCCATGGACTCAGCCGCC;CTATTGTCAATATCACCGGG;TGACAATAGGAGAGAGAAAG | -0.2;1.15;3.0 |
| DNAH6 | 1.315 | 1.238 | 3 | AATACAGCAACCACCGCCGA;AAATACAGCAACCACCGCCG;CTCTCTCTGGAGACCCTCGG | -0.45;0.55;3.85 |
| ATP5D | 1.312 | 2.161 | 3 | AGCTGGCGGACAGCGGACTC;GCGGACAGCGGACTCCGGCG;AGCGGGTAGCTGGCGGACAG | 0.77;1.13;2.04 |
| SLC39A8 | 1.304 | 1.776 | 3 | TCAGAGATACAGAGGTTGTG;CAGAGATACAGAGGTTGTGG;CGGACCTGTCAGAGATACAG | 0.29;1.02;2.6 |
| R3HDM1 | 1.304 | 2.248 | 3 | GCAAGTAAGAGTCTTACGAA;AATTAACCCCATCACCAGCC;CAAGTAAGAGTCTTACGAAA | 0.9;1.13;1.89 |
| GTPBP3 | 1.304 | 1.970 | 3 | CAGGTTGTAAATCCATGTGG;TCGCAGGTTGTAAATCCATG;GCGGGGGCTTTGGACCCTGG | 0.44;1.29;2.18 |
| CC2D1A | 1.303 | 1.814 | 3 | GTGGCCGCGCTCCGGTGCGG;CCAGTGGCCGCGCTCCGGTG;GGCAGACCCGGCGAGCCCAG | 0.35;1.03;2.53 |
| GMEB2 | 1.302 | 1.890 | 3 | GGCGGCGGCGTCGGGAGCTG;CCAACAAGGAGCGAAGCCCG;GGCGGCGTCGGGAGCTGCGG | 0.64;0.72;2.55 |
| C4orf32 | 1.292 | 1.035 | 3 | CCTGTCAGCGGCGGGTGCGG;ATCCCAGGGCAGCCTTCGGG;GGCGGGTGCGGCGGATCCCA | -0.64;-0.58;5.1 |
| UQCR10 | 1.292 | 1.747 | 3 | CAATTTCGAAGTCAACGTCG;GGGCGAAGGTGGAGGTCCTG;GGAGGTCCTGCGGAACAGCA | -0.14;1.56;2.46 |
| SH3BP4 | 1.287 | 1.370 | 3 | CGGCTCCGCGGGACTCACGG;GGCGGTAGTGGCGAGGGGCC;CGCCGCCGCCGTGAGTCCCG | -0.09;0.6;3.34 |

| | | | | | |
|---|---|---|---|---|---|
| RBM7 | 1.283 | 1.856 | 3 | GCGTCGCCCCCTCCCTAACC;AGGGGGCGACGCTGAGATGG;GGGCGACGCTGAGATGGGGG | 0.43;1.02;2.4 |
| ZAR1 | 1.283 | 1.627 | 3 | CTGGGGGACGAGGTGCTGGA;CGGGAGCAGTGCGCCCATGG;GTCCAGCACCTCGTCCCCCA | -0.35;1.39;2.81 |
| INO80C | 1.283 | 1.895 | 3 | GAAGTTCCAAGGCCCGCGCT;GGCCCGCGCTGGGAAAAAGG;ACAGCGGAAAGGAAGTTCCA | 0.64;0.75;2.46 |
| DCAF12 | 1.282 | 2.462 | 3 | GAGAATGAGCGGCTTTCCGA;GAAGGGGAAGCGAGAATGAG;ACTTGAGCCGGGAAAGGGAA | 1.2;1.24;1.41 |
| SPEN | 1.282 | 2.265 | 3 | AGGCCCGAGAGTCAGAACCT;GCCCGAGAGTCAGAACCTGG;CAGAACCTGGGGGAGAGGGA | 0.86;1.29;1.7 |
| CDK12 | 1.280 | 2.020 | 3 | TGTGTGGTGTGGAGGTGAAA;GTGAAACGGAGGCAAGAAAG;GTGGTGTGGAGGTGAAACGG | 0.78;0.8;2.27 |
| SPTAN1 | 1.280 | 1.696 | 3 | CGAGGGGTGCTGAAGGACCG;CCGCTGCGGAGTGAACGGTG;GAAGGACCGAGGAGCCTCCG | -0.21;1.47;2.58 |
| R3HCC1L | 1.280 | 1.990 | 3 | AGAGGGAGGGGATACGGGGC;AGCGGAGAGCAACAGCGCGC;GTACAGAGGGAGGGGATACG | 0.47;1.34;2.03 |
| ZKSCAN3 | 1.273 | 2.419 | 3 | TGGGGTTAAATCTCATCCCG;GATGAGATTTAACCCCACAG;GGCGTCGGCCTTCCACTGTG | 1.09;1.32;1.41 |
| ERP44 | 1.271 | 2.041 | 3 | GGAGAATCCTCCGCTGCCGT;ACGGCAGCGGAGGATTCTCC;GAGCCCGGGTCGAGAGGACG | 0.73;0.94;2.14 |
| RSRC1 | 1.269 | 1.961 | 3 | TTAAACTGAAGCAAGTTCGG;GAAGCAAGTTCGGTGGACGC;GCAAGTTCGGTGGACGCCGG | 0.28;1.63;1.89 |
| ARHGAP9 | 1.267 | 1.757 | 3 | TAAAAAGCAGCTGGGGCCTG;GAGGTGACCCAGGGTACTGG;CTCTGGGCTGAGGTGACCCA | 0.45;0.77;2.59 |
| TLN1 | 1.265 | 1.654 | 3 | GGGCGACCCGAGAAGCGGCG;GTGGCCGAGAGAGTGTCGAA;CCGAGAGAGTGTCGAAGGGA | 0.46;0.5;2.83 |
| NO_CURRENT_220 | 1.264 | 1.789 | 3 | GGATCTAGCTACCTCAAAAG;GGATTAATTCGCTAAATGAT;GGATGTGGAAAAGAACCCAG | 0.35;1.0;2.44 |
| SNRNP70 | 1.263 | 1.815 | 3 | TGCGCTCGCTTAGCGGGCGA;CCGCGCGGGTGGCTGAGCAG;TAGCGGGCGACGGAATCAGA | -1.05;2.37;2.47 |
| KRTAP22-1 | 1.262 | 1.407 | 3 | GGTGGCCAGGGCTATGCCAA;GGCCAGGGCTATGCCAAAGG;TGGCATAGCCCTGGCCACCA | 0.02;0.6;3.17 |

| | | | | | |
|---|---|---|---|---|---|
| MIF4GD | 1.258 | 1.118 | 3 | GGGAGCCGGGACCCACCTGC;GCTGCGGAGCCCACGGCCCA;GCCCACGGCCCAGGGTCCCG | -0.43;0.26;3.94 |
| ANKRD17 | 1.252 | 2.084 | 3 | GCTACGCTCTACCGCGACTT;CCGCGACTTCGGCCGCACTG;TTGCTGTGTCGGCCCCAGTG | 0.52;1.55;1.69 |
| PKD2L2 | 1.250 | 1.386 | 3 | GAGGCGTCACGGTGGCACCG;CCATGGCTGAGGCGTCACGG;CCGTGACGCCTCAGCCATGG | -0.35;0.86;3.24 |
| HMBS | 1.248 | 1.989 | 3 | CCGGGGACACGTGGGACCCG;GGCGAGTACCGGGGACACGT;GAGACCAGGAGTCAGACTGT | 0.55;1.19;2.0 |
| SHH | 1.247 | 1.893 | 3 | TAACGGAACACATCGGAGTT;ATAACGGAACACATCGGAGT;CCGCTGATAACGGAACACAT | 0.37;1.28;2.09 |
| SBNO2 | 1.247 | 2.314 | 3 | CGAAACCCGGAAGTGAGCGG;GCTCGGAGAAACAGGCGCCG;AGTGAGCGGCGGCAGCTGCG | 1.03;1.14;1.56 |
| CENPB | 1.245 | 1.614 | 3 | CCGGCGGGCGCGTCGCCGGG;CAGGCGACGGGCGCAGACGA;GCGACGGGCGCAGACGAGGG | 0.06;0.99;2.68 |
| SLC25A15 | 1.244 | 2.269 | 3 | ACGCGGCCACGCCGGCGCGG;GACCCGGAGCCGAGAGCGGG;GGCGGAGCCTGAGCTGGACG | 0.85;1.41;1.47 |
| PROSER3 | 1.243 | 1.496 | 3 | AGAGCGATCGGCCTCACCTG;GTAGTCCCTGGAAGAGCGAT;CTCACCTGCGGTCCATCCCG | -0.36;1.11;2.98 |
| TTLL2 | 1.241 | 1.425 | 3 | AGACCCACAGAGGCCACCCT;GGAACCAGCGCCCAATGAGA;TGCTGCACAGAGACCCACAG | -0.42;0.98;3.16 |
| GLRX5 | 1.234 | 1.998 | 3 | CGTGGGCTCCGGCTTGCGTG;GTCCCTCGGCCGAGCTGCGG;GGCTTGCGTGCGGAGATGAG | 0.64;1.06;2.0 |
| ZNF638 | 1.232 | 2.128 | 3 | GTGCTAAACTGTGTGGGGCG;GGCTGAGTGCTAAACTGTGT;GCTGAGTGCTAAACTGTGTG | 0.67;1.37;1.65 |
| FBXO38 | 1.232 | 2.300 | 3 | CAACCACAATAACAGGCGGA;TCAACCACAATAACAGGCGG;CTCTCAACCACAATAACAGG | 0.93;1.36;1.4 |
| HM13 | 1.229 | 2.117 | 3 | GGTTGCTGCAGCAGGGACGC;TGCTGCAGCAGGGACGCAGG;AGGGACGCAGGCGGAGACAC | 0.64;1.41;1.63 |
| PMAIP1 | 1.227 | 2.039 | 3 | GGATCGGGTGTCCGGACGCG;GCGGAGATGCCAACTACACA;ACACGAACAGTCCTGCAGGC | 0.49;1.53;1.67 |
| STRN | 1.226 | 1.971 | 3 | ATGGCGGCCGCAGATACCCG;GGCAGCTCCCCGGGTATCTG;ACCCGCCTGCTCGTCCATGG | 0.35;1.6;1.73 |

| | | | | | |
|---|---|---|---|---|---|
| PKMYT1 | 1.226 | 1.701 | 3 | GACTGTTCCGGACGCCCGGG;CGGGCGTCCGGAACAGTCGA;GAACAGTCGACGGCAGACTC | -0.24;1.6;2.32 |
| ZSWIM8 | 1.217 | 2.049 | 3 | CAGTCTCCGAGACTTAGGGC;GGCCGGGACGAAGCGCTCGG;TAGGGCCGGGACGAAGCGCT | 0.59;1.35;1.71 |
| MMP13 | 1.216 | 1.933 | 3 | AATGAGTCCAGCTCAAGAAG;ACCATTCAAGATGCATCCAG;AGTCCAGCTCAAGAAGAGGA | 0.62;0.95;2.07 |
| MXD1 | 1.215 | 1.894 | 3 | GGGTTCTGTCACTGTGTCGG;GCAACCACGCTCGACAAGAG;TCCGGGTTCTGTCACTGTGT | 0.59;0.91;2.14 |
| ZNF543 | 1.212 | 1.577 | 3 | GGATTGGCCCTGGAACAGTG;CCAGGCCCCACACTGTTCCA;CGCTGGACGCGCCTACCCAG | -0.2;1.21;2.62 |
| IFNGR2 | 1.211 | 0.966 | 3 | CGGCGACGTGAGCGGCTCCG;CGTCGCCGCCCAAACCGCCA;GCGCCCTGCGCTCGCCATGG | -0.93;-0.34;4.9 |
| CLPX | 1.203 | 1.812 | 3 | CCGTGGAGAGTTCACCTGCC;TTCGCGGGCCCTAGACCCCG;GGGCAGGTGAACTCTCCACG | 0.32;1.19;2.1 |
| MAX | 1.203 | 1.954 | 3 | GTTGATCCGGTGCTGCAGTG;GATCCGGTGCTGCAGTGAGG;CGATAACGATGACATCGAGG | 0.48;1.33;1.8 |
| SERP2 | 1.200 | 1.917 | 3 | AAGCCATGGTGGCCAAACAG;TCAGAGCGCACAAGCCATGG;ACAGCAAAAACATCACCCAG | 0.36;1.43;1.8 |
| ALAS1 | 1.199 | 1.945 | 3 | AGGGCAACGAGCGTTTCGTT;GACTGCAGAGGCGGCGGCGA;TGCAGAGGCGGCGGCGAAGG | 0.7;0.84;2.05 |
| TTLL9 | 1.197 | 1.851 | 3 | GCATAAGCACGCGAGGCGCG;CGACATAACGCGATTCCCCA;GACATAACGCGATTCCCCAG | 0.18;1.57;1.84 |
| CGGBP1 | 1.196 | 1.895 | 3 | TCGGGCAACGGCGGCGACGG;GAGGAGCAAGGGAATAAGGG;GAGCAAGGGAATAAGGGAGG | 0.24;1.62;1.72 |
| SLC43A1 | 1.192 | 2.038 | 3 | CTGTCTCCTCGGAAGAAGCG;TGTCTCCTCGGAAGAAGCGG;GGGAACCCGCCGGGCGCCAG | 0.81;0.89;1.88 |
| THRA | 1.191 | 1.781 | 3 | CCATGGACGCCCCCAGCACG;GTCTCAGCGCCCCGTGCTGG;ACCCGGGCGCAGGAGGCGGG | 0.46;0.9;2.21 |
| RAP1A | 1.190 | 1.800 | 3 | GAGCAGGAGCCACGGCCGAG;AGCCACGGCCGAGAGGAGGG;CAGGAGCCACGGCCGAGAGG | 0.61;0.68;2.29 |
| FKBP1A | 1.190 | 2.028 | 3 | GTGGACCAACAGCGACCTGG;GCGACCTGGCGGCGGTTCCA;GGCGTGGACCAACAGCGACC | 0.57;1.37;1.63 |

| | | | | | |
|---|---|---|---|---|---|
| VIRMA | 1.189 | 2.115 | 3 | CATGGCGGTGGACTCGGCGA;GGCAAACATGGCGGTGGACT;CCCGCCCGCGGCAAACATGG | 0.84;1.04;1.68 |
| CLPP | 1.188 | 2.065 | 3 | CATCGGACGGAAGCCGACCG;ATGTGGCCCGGAATATTGGT;TGTGGCCCGGAATATTGGTA | 0.86;0.87;1.83 |
| TADA3 | 1.178 | 1.690 | 3 | GCGGCCCCTGGGGCCGTAGG;GCGGCCTCCTACGGCCCCAG;GCCCCCTCCCGCGGCCCCTG | 0.16;1.15;2.22 |
| ZFAT | 1.174 | 1.819 | 3 | CAGATTCGCCATAACCTCGC;GAAAAGAGCCGGCGAGGTTA;ATCTGCGGCATCCAACATGG | 0.44;1.04;2.04 |
| BRK1 | 1.173 | 1.569 | 3 | GACAGGAGGATCCGGTGCAG;ACTCCCGGTTAGCCCAGTCC;GGTGCAGCGGGAGATTCACC | -0.04;1.1;2.46 |
| ATXN7L2 | 1.173 | 1.776 | 3 | CCGAGTCTCGATGACTTCGC;CCCGCGAAGTCATCGAGACT;CGGCACCCGCCGCTCCAGAG | 0.24;1.29;1.99 |
| ZNF764 | 1.172 | 1.157 | 3 | GAGGTTACTAGGGCCCCCGA;CTAGAGGGCGTGGAAACTAA;GCCCTCTAGTGCGCCCTCGG | -0.74;0.54;3.71 |
| COX18 | 1.170 | 1.725 | 3 | ACAGCATTTCTGCACCACGG;TGTATGTCCGCTGGATTGTG;TCCCGGGCCTCACAATCCAG | 0.53;0.66;2.32 |
| KLF11 | 1.169 | 2.142 | 3 | GCGGCAACCCGGAGCTCCTG;CAACCCGGAGCTCCTGCGGC;GGCCGCACGTGGGCTGCGGG | 0.89;1.06;1.56 |
| VPS36 | 1.168 | 1.916 | 3 | GGTCCAAACGAAGCGGTCCA;GGCCGCTGGTCCAAACGAAG;GGGTCTCGTTGATCTCCAGG | 0.71;0.8;1.99 |
| FAM160A2 | 1.165 | 1.660 | 3 | GCCGGGCCCGGTTGCTGCTG;ACGGAACGCCGAACCTGGCC;CTAACACCGCAGCAGCAACC | -0.06;1.39;2.16 |
| DDA1 | 1.164 | 2.128 | 3 | GGCTAAGAAGGCGGCTCTGG;TAAGAAGGCGGCTCTGGTGG;GGCTGAGGCGGCGGCCGAGG | 0.91;0.98;1.6 |
| DRD2 | 1.163 | 1.753 | 3 | GGCAGGAGGGAGCGCGGGGA;GGGACGGGGCGGAGGCGGCG;GGCGGGCAGGAGGGAGCGCG | -1.65;2.02;3.13 |
| HMGN2 | 1.163 | 1.550 | 3 | TCTTCACACTGCTCGGGCTC;TTCACACTGCTCGGGCTCCG;GCCCGAGCAGTGTGAAGAAG | -0.12;1.14;2.47 |
| TNR | 1.162 | 1.666 | 3 | ACCGACTGTGCTAAGGCTGT;GGCTCAACACCCAGAGGAGA;GCTGTTGGCTCAACACCCAG | -0.01;1.36;2.14 |
| ATP5G3 | 1.160 | 2.120 | 3 | CCTCCACTTACCTTCCCAGG;CTCCTGGGAAGGTAAGTGGA;CCTCCTGGGAAGGTAAGTGG | 0.74;1.36;1.38 |

| | | | | | |
|---|---|---|---|---|---|
| LAPTM4A | 1.160 | 1.809 | 3 | AGCCGTTTGAGTTTGGCTGC;CGGGTGGAGAACGTTTGTCA;GGGTGGAGAACGTTTGTCAG | 0.46;1.01;2.02 |
| RBL1 | 1.156 | 1.817 | 3 | GAACATCCCTTCAGGCCCCG;TGGGAGGGAGAAAGAAGTCG;GGGAGGGAGAAAGAAGTCGG | 0.31;1.3;1.86 |
| CTSG | 1.154 | 1.608 | 3 | CTGCCTCAGCCCCAGTGGGT;GGCCTTTCTCCTACCCACTG;CTCACCTGCCTCAGCCCCAG | -0.26;1.43;2.29 |
| YWHAE | 1.151 | 2.053 | 3 | ATCCGTCGCGCAGACCCTGC;GTCTGCGCGACGGATGGAAG;GAGCGAGAGGCTGAGAGAGT | 0.81;0.98;1.66 |
| DPPA4 | 1.150 | 1.587 | 3 | CTCCGCTTCTTCTACAAGTA;CTCCATACTTGTAGAAGAAG;TTCTTCTACAAGTATGGAGA | -0.27;1.39;2.33 |
| VPS25 | 1.149 | 1.692 | 3 | GAGCCTCACGTAAAGAAGGG;CGATGGCGATGAGTTTCGAG;GGTGGGAAGCGATACTGCCA | 0.29;1.0;2.16 |
| OPN1SW | 1.148 | 1.675 | 3 | TATCTCTTCAGTGGGGCCGT;TGGTACTGAGGCCCATCCCA;TCAAAAATATCTCTTCAGTG | -0.44;1.84;2.04 |
| FAM133B | 1.148 | 1.573 | 3 | GGCCGGAGAGACTGCCGAAG;GCCGGAGAGACTGCCGAAGA;CACGCCGAGGGAAACCGGGC | 0.12;0.94;2.38 |
| RFX7 | 1.148 | 1.820 | 3 | GACCCGGCAGTAGAAAGCCC;CCCGGCAGTAGAAAGCCCCG;GCCCCGGGGACAAGGAAGGA | 0.38;1.19;1.87 |
| PFKFB2 | 1.147 | 0.991 | 3 | GCGTTACAGGGCAGGCGCCG;GTTCGGTCGCGTTACAGGGC;CAGGCGCCGGGGCCAAGGCA | -0.59;-0.41;4.45 |
| ARL8A | 1.145 | 1.383 | 3 | TGCGGGCGGAGGCTCGAGCG;GTCCCGCGGCTCGGTGCGGG;CCGGTACGGCCCGGGTCCCG | 0.01;0.62;2.8 |
| CSNK1D | 1.143 | 1.836 | 3 | GTTCCCGACTCTCAGCTCCA;CCCGACTCTCAGCTCCATGG;GACTCTCAGCTCCATGGCGG | 0.37;1.28;1.78 |
| GPI | 1.141 | 1.983 | 3 | TGTACCTTCTAGTCCCGCCA;TGCCGGCGCTCCTTCCTCCT;CCGGGTGAGAGCGGCCATGG | 0.55;1.37;1.5 |
| ZMAT5 | 1.140 | 2.163 | 3 | GAAGGTACCTGGCTCCCCGG;GCCCGGAGCTGCAGCCACCG;GCGGAAGGTACCTGGCTCCC | 0.91;1.14;1.36 |
| DDX1 | 1.139 | 1.467 | 3 | GCGAGCAGGCGAAGCCGCGG;CGCCGTGTCAGTCGGGAGGG;GCCGTGTCAGTCGGGAGGGA | -0.37;1.13;2.67 |
| INPP5A | 1.138 | 1.233 | 3 | GCGCAGCCATTAGATCCGCT;GGCTGCGGGAACTTTCCCAG;GCTGCGCGCGGGCCGCTGTG | -0.38;0.63;3.17 |

| | | | | | |
|---|---|---|---|---|---|
| LDLRAD2 | 1.137 | 2.073 | 3 | GCTGGTGGGACACAAGGAGA;TGGTGGGACACAAGGAGAAG;GGTGGGACACAAGGAGAAGG | 0.78;1.14;1.49 |
| RIOX1 | 1.136 | 1.797 | 3 | CCACACCAGTCTTCCCCGCC;GGCCGGGGGCGAAGCTAAAG;CCGGGCGGGGAAGACTGGTG | 0.27;1.35;1.79 |
| LYSMD2 | 1.136 | 2.094 | 3 | GCGGGCGAGGAATCCGCCAT;GCCTTCCCGCAGGGACAGTG;CCTCGCCCGCACTGTCCCTG | 0.87;1.02;1.52 |
| TFPI2 | 1.133 | 1.583 | 3 | CGTCCGAGAAAGCGCCTGGC;TTCTCGGACGCCTTGCCCAG;GCGTCCGAGAAAGCGCCTGG | 0.02;1.11;2.27 |
| KCNMB4 | 1.129 | 2.240 | 3 | GTCAGCGGAGTAGCGGCCAG;GCAGCTGCCGGGCGAGTCAG;CGGGCGAGTCAGCGGAGTAG | 1.05;1.14;1.2 |
| RADIL | 1.128 | 1.795 | 3 | GGCCCCAGCCAACGGCGGGT;TGGGCGCCGGCCCCAGCCAA;GCGCCGGCCCCAGCCAACGG | 0.17;1.55;1.67 |
| PHF10 | 1.125 | 1.366 | 3 | CGCCGCTCCCCTCAGCCCCG;CCGGGCCGTGGGGAGCCAGG;CGGGGCTGAGGGGAGCGGCG | -0.12;0.72;2.77 |
| TARBP1 | 1.125 | 1.918 | 3 | CCACCGGCCCGGGCTCCCAA;GGGCTCCCAAAGGAAGGCGC;CGGCCCGGGCTCCCAAAGGA | 0.5;1.3;1.58 |
| ARRDC3 | 1.121 | 1.006 | 3 | TGAGATTTCTTAAAAAGTCA;TTGAGATTTCTTAAAAAGTC;TTAAAAAGTCAGGGCAGCAG | -0.41;-0.22;3.99 |
| TBC1D14 | 1.119 | 2.099 | 3 | GACCGGCAACTCGCGCGACG;CGCGAGTTGCCGGTCCCGCG;GAGGACGCCGGGCACGCGGT | 0.78;1.28;1.3 |
| C15orf53 | 1.118 | 1.745 | 3 | GGAGCTACAAGGGGCCCAAG;TTCTGATATGGAGCTACAAG;CAAGGGGCCCAAGAGGACCT | 0.28;1.2;1.87 |
| KDM3B | 1.116 | 1.394 | 3 | CGCCGCGTCCGCCATCGCCG;TTGGCGGCGGAGGTGGTGGG;CGGAGGTGGTGGGAGGCGGC | 0.23;0.36;2.76 |
| ASCC2 | 1.112 | 1.629 | 3 | CGCTCAGGGAGCTGTCACCG;GGAGCTGTCACCGTGGTCGG;CGGCGGCGGCACAGAGCCGG | 0.47;0.61;2.27 |
| LAMTOR 3 | 1.109 | 1.813 | 3 | TCAGGGACAGCTTTAAAGAC;AGCTGTCCCTGAAGTGACAG;AAGTGACAGCGGAGAGAACC | 0.42;1.17;1.74 |
| MYO1G | 1.107 | 2.033 | 3 | GTTTCCAGCCGGCAGGATGG;GGTGTTTCCAGCCGGCAGGA;CCAGCAGGCACAGGAAGGTG | 0.78;1.06;1.48 |
| IL3 | 1.106 | 1.191 | 3 | CTCATGTTTGGATCGGCAGG;CAGGCGGCTCATGTTTGGAT;GAGTTGGAGCAGGAGCAGGA | 0.02;0.08;3.22 |

| | | | | | |
|---|---|---|---|---|---|
| PLA2G7 | 1.103 | 1.207 | 3 | CTGCTCGGCCCGCAGCCAGG;GGCCGAGCAGCTCTGGCAGG;GCCCGCAGCCAGGGGGACAG | -0.15;0.4;3.06 |
| TNRC18 | 1.102 | 1.469 | 3 | AGATTTGGGATTCTTTAAAC;TAATAAAATCCAGGTAGCGC;TTAAAAAGCGGGGAGCTCGG | 0.36;0.36;2.58 |
| ATXN7 | 1.102 | 2.036 | 3 | ACATGGAGCATATTTGGAGT;GGAGCATATTTGGAGTTGGT;TGGAGCATATTTGGAGTTGG | 0.78;1.09;1.43 |
| RHOG | 1.101 | 1.578 | 3 | CCAGCTCCCCCGCCTCGGGG;CCCCCAGCTCCCCCGCCTCG;CTCACCTGGTGCCCTCCCCG | 0.15;0.97;2.18 |
| CERS6 | 1.101 | 1.982 | 3 | GCGGCGGCGGCACAGGCTCG;GAGAGCAGCGGCCGCGGAGG;CGCCGCCGCCGCCTCCTCCG | 0.64;1.26;1.41 |
| TTLL1 | 1.100 | 1.753 | 3 | TACTCCTGCGAGCGCCGGAG;CTCCTGCGAGCGCCGGAGAG;ACTCGGAGGAGCGCCCTGCA | 0.54;0.8;1.96 |
| EIF1 | 1.098 | 1.948 | 3 | GGCAGGGGGGAACGGAACGT;GGAAGCGAGGGGGCTCAAGG;CGGAACGTCGGGAAGCGAGG | 0.66;1.09;1.54 |
| NCOA6 | 1.096 | 2.074 | 3 | CGGCTCCGTGAGGCCCTGCC;CTGCCGGGTCGGGCTGCGGG;CCCGACCCGGCAGGGCCTCA | 0.87;1.04;1.37 |
| VAT1 | 1.095 | 1.505 | 3 | GACGAGAGCGCACAGCTGGA;AGCCATGTCCGACGAGAGAG;CTCGGCTACCTCTCTCTCGT | 0.34;0.52;2.43 |
| FASN | 1.094 | 1.126 | 3 | GAGGCTGAAGCGCGGCGGAG;AGCGGGAGGCTGAAGCGCGG;CTGAAGCGCGGCGGAGAGGG | -0.16;0.18;3.27 |
| VAV1 | 1.091 | 2.022 | 3 | GAGCAGGGCAGGCGTGCGGG;AGGCTGCGAGGGTGCACGGC;GGCGAGCAGGGCAGGCGTGC | 0.75;1.15;1.37 |
| E2F6 | 1.090 | 1.756 | 3 | GACGCAGACGGAAAAAGAGG;AAAAAGAGGAGGGAGACCCG;ACGCAGACGGAAAAAGAGGA | 0.16;1.49;1.62 |
| TXN2 | 1.086 | 2.038 | 3 | CACCTCGAGCCACCCCCACA;CTACCTCCCTGCAATGCGAG;CCTCCGCTCGCATTGCAGGG | 0.85;1.0;1.41 |
| CCDC175 | 1.084 | 1.784 | 3 | TGGCGAGAAGCTGGTGCAGG;CGCCCAGCCCTGGGGTCCAG;CTCGCCAGCGCCCAGCCCTG | 0.21;1.51;1.53 |
| NOTCH3 | 1.084 | 1.307 | 3 | ACGCGCCCGGAGCCCAGGGA;GCCCGGAGCCCAGGGAAGGA;CGGAGCCCAGGGAAGGAGGG | 0.03;0.46;2.76 |
| DMD | 1.083 | 2.129 | 3 | TCCACTTTCGGGGAGCCCGG;GCGGCGGCGCTCCACTTTCG;GCCGCCGGGCTCCCCGAAAG | 0.94;1.11;1.2 |

| | | | | | |
|---|---|---|---|---|---|
| CHKB | 1.080 | 1.647 | 3 | GCGGAGCGCAGCCTGGCCTG;CGGTCGAGCCCGCGCCATGG;CCGAGCCCGTCCGAAGGGAG | 0.13;1.21;1.9 |
| RRAGA | 1.080 | 2.173 | 3 | AGTAAGAGCCAGCCCGTCCG;TACTCAGATGGGAGCGCCCG;GGGCTGGCTCTTACTCAGAT | 1.01;1.1;1.12 |
| BCL10 | 1.080 | 1.934 | 3 | CGTCCTTCTTCACTTCAGTG;CGCACCGTCCCTCACCGAGG;CACCGCACCGTCCCTCACCG | 0.8;0.81;1.63 |
| DET1 | 1.077 | 1.902 | 3 | GCTGATGCCCAGAGGCCCCG;TGCATGACGCTGATGCCCAG;GGCATCAGCGTCATGCAGGG | 0.56;1.21;1.46 |
| BAHCC1 | 1.076 | 1.659 | 3 | GGACTGGGCTTCTCGGCGGG;GGGAGGACTGGGCTTCTCGG;GGGGCCGGTCAGCACGCGGG | -0.15;1.64;1.74 |
| RASGRP2 | 1.073 | 1.318 | 3 | GTCAGCGCCTGTGTGGCCGC;TGAGGATGTCAGCGCCTGTG;CAGGCCAGCCTTGAGTCCCG | -0.02;0.57;2.68 |
| AK4 | 1.073 | 1.797 | 3 | CGCCGGGACCGAGTAGCAGG;GTCCGCCTGCTACTCGGTCC;TGCTACTCGGTCCCGGCGCT | 0.66;0.72;1.84 |
| FGFR1 | 1.073 | 1.397 | 3 | CTAGCTGCCGCCCGCCGCCG;GCGGCGTCCTCGGCGGCGGG;CGCCGAGGACGCCGCGCCTG | -0.01;0.75;2.47 |
| CYBA | 1.070 | 1.080 | 3 | GCCGGGTTCGTGTCGCCATG;CGCCATGGGGCAGATCGAGT;GGCCCACTCGATCTGCCCCA | -0.17;-0.04;3.42 |
| GRSF1 | 1.070 | 1.942 | 3 | TGCGCCGCCTCCGGAGTCCA;GTGCCGGCCATGGACTCCGG;TGGACTCCGGAGGCGGCGCA | 0.65;1.16;1.4 |
| RNF146 | 1.068 | 2.012 | 3 | GAGAGGTGGGAGTCGGAGCG;AGAGAGAAAAGACTGCGAGG;GCCAGAGAGAAAAGACTGCG | 0.88;0.89;1.43 |
| FBXO43 | 1.067 | 1.720 | 3 | CCCACGCGAAGCTGGCGTGG;GCCATAGCCCGCATCCCAGG;TGCCATAGCCCGCATCCCAG | 0.51;0.81;1.88 |
| ASIC3 | 1.066 | 1.477 | 3 | CCGGGCCAGGAAGCGCGCGG;CCAAGGAGCTCTCCATGGTG;GCGGCTCGGGATCCGCACCA | -1.26;1.34;3.12 |
| BBX | 1.064 | 1.945 | 3 | AGCAGTGTACAGTGAAGCGG;ACAGTGAAGCGGAGGCAGAG;TGGAGCAGTGTACAGTGAAG | 0.7;1.06;1.43 |
| ENG | 1.064 | 2.037 | 3 | GGGTGCTGGGCTCCAATGGA;ACAGCAGCAGTCCTGGCCCC;CAGCAGCAGTCCTGGCCCCA | 0.83;1.07;1.28 |
| NDUFAF6 | 1.063 | 1.053 | 3 | CCCCCAGACAGAGCCGTGCG;CCAGACAGAGCCGTGCGCGG;CCGCGCACGGCTCTGTCTGG | -0.56;0.28;3.47 |

| | | | | | |
|---|---|---|---|---|---|
| NR2F2 | 1.061 | 1.282 | 3 | TGATCAAATATGCTAAAAAG;ATCAAATATGCTAAAAAGGG;AATGCGATTTATAGGCGCCG | -0.22;0.67;2.73 |
| AMIGO1 | 1.059 | 1.233 | 3 | GGCGCAGCGATCCCAGCGGA;AGGCGCAGCGATCCCAGCGG;CGGAGGCGCAGCGATCCCAG | -0.11;0.47;2.82 |
| IFITM1 | 1.059 | 1.568 | 3 | CGTGGAGCGAAGGGCCGCTG;GAGTGAAGGTTTGAGAAGTG;CCAGCATCCGGACACCACAG | 0.45;0.54;2.19 |
| IFIT2 | 1.059 | 1.899 | 3 | GCAATTCTCAGCTGTTCGGC;CAGTGCAATTCTCAGCTGTT;CAATTCTCAGCTGTTCGGCA | 0.59;1.19;1.4 |
| TMEM120A | 1.058 | 1.551 | 3 | GTAGATCCTCCCAGTCCCGC;CGACTGCCTGCGGGACTGGG;TCCCGCAGGCAGTCGCCCAG | 0.44;0.52;2.22 |
| RAB44 | 1.056 | 1.443 | 3 | GCAGCAGTCACCCTACCACC;CCCAGTGGAACACACAGCCC;CCACCAGGTCCCAGAGCCCA | 0.27;0.52;2.38 |
| PIAS4 | 1.055 | 1.962 | 3 | AGGCCAAAGTGAGTGAGCGG;TGGAGGCCAAAGTGAGTGAG;GCCCTCTCGCCCACTTGCCC | 0.75;1.02;1.39 |
| SMIM24 | 1.053 | 1.368 | 3 | CAGCACCAGAAGGGCCCCCA;CGACCGTCATGGAGACCCTG;GACCGTCATGGAGACCCTGG | -0.98;1.09;3.05 |
| MOB4 | 1.052 | 1.744 | 3 | AGACGCTGGCACTATGGTCA;GCACTATGGTCATGGCGGAG;CGCTGGCACTATGGTCATGG | 0.47;0.96;1.73 |
| ZBTB7A | 1.050 | 2.010 | 3 | CTGGTCGCTCCCTCGCGCGG;GCCCTGGTCGCTCCCTCGCG;GTACTGTACAGTTACTGTAG | 0.8;1.09;1.26 |
| MUM1L1 | 1.050 | 1.525 | 3 | GCATTACACACAGGCTTCGG;GGAGGAGCTGCATTACACAC;ATTACCTCCAAGCTTCTCCA | 0.32;0.66;2.17 |
| ARMCX4 | 1.050 | 1.307 | 3 | TTACCTGCTCCGCTTTCCCC;GGGCCCGGGGAAAGCGGAGC;GGCATCGGGCCCGGGGAAAG | -0.43;0.86;2.72 |
| SS18L1 | 1.048 | 1.985 | 3 | CCGCGCCGCCACCATGTCCG;GTATCCACCTCGATGACCAC;CGCGAAGGCCACGGACATGG | 0.82;0.97;1.35 |
| PRR14 | 1.046 | 1.698 | 3 | TGCTTGACAGTGGATCCCTG;AGAGGGTATCTGCTTGACAG;GAACTCAGCGTAGATCCCCA | 0.31;1.13;1.7 |
| LCTL | 1.045 | 1.563 | 3 | CCCTCTCCAACTGACCCCCA;GGGGACAGCGTAGGCCCTGG;CCTCTCTCAGGGGACAGCGT | 0.38;0.66;2.09 |
| ZNF292 | 1.045 | 1.808 | 3 | GAGCAGGAGAGGTTGAGTTG;TGTGAAGATGGCGGACGAAG;GCAACTCAACCTCTCCTGCT | 0.37;1.38;1.38 |

| | | | | | |
|---|---|---|---|---|---|
| HERC5 | 1.043 | 1.636 | 3 | ACCAGCTGCGCGCAGCTGCG;GCTCTGCAGCGCAACGCCTG;TCAGGCGTTGCGCTGCAGAG | -0.0;1.43;1.71 |
| EED | 1.043 | 0.932 | 3 | GCGGCTGAAACGTCTTTGGA;GCTGAAACGTCTTTGGAAGG;CGTCTTTGGAAGGAGGAAGG | -0.82;-0.29;4.24 |
| STAG1 | 1.041 | 1.322 | 3 | GGAAAATGCCACCAGATGGC;TTGCAGCTCCGTTGAAGGCG;TAGGATTGCAGCTCCGTTGA | -0.05;0.65;2.52 |
| TIGD2 | 1.040 | 1.806 | 3 | CAGACCACACTAAGCGCAAA;GACTCCATTTGCGCTTAGTG;AGCGCAAATGGAGTCTCCTG | 0.44;1.25;1.43 |
| MIEF1 | 1.040 | 1.655 | 3 | CAGAGGCGAGTCCTGCGGAG;CCGGCCAGAGGCGAGTCCTG;GGGGAGAGGTACCCGGCCAG | 0.03;1.45;1.64 |
| NDUFAF5 | 1.038 | 1.789 | 3 | CTTATGTCGGCGACCTTGGG;CAGGGCTCTGGCGCTTATGT;GCAGCTGGAGATGCTGCGGC | 0.55;0.95;1.61 |
| STAT3 | 1.034 | 1.142 | 3 | AAGGGCCTCTCCGAGCCGAG;AGGGCCTCTCCGAGCCGAGG;TTGGCGCTGTCTCTCCCCCT | -0.12;0.24;2.98 |
| SPCS1 | 1.033 | 1.737 | 3 | CGAGACTTAAGGACCGTGAG;CCGAGACTTAAGGACCGTGA;ACCGAGACTTAAGGACCGTG | 0.35;1.21;1.54 |
| FSTL5 | 1.032 | 1.565 | 3 | GGTACTGTCCACTGTGCTGT;GTTCAGCACCGACAGCACAG;CAGTGACTGAGACAAACTGG | -0.35;1.55;1.89 |
| ARMC5 | 1.032 | 1.830 | 3 | CTGCCTCGCGCAGCTCGCGG;GAGAGCGAGTCCGTGAGGGT;GAACGAGAGCGAGTCCGTGA | 0.49;1.24;1.37 |
| PSTPIP1 | 1.032 | 1.662 | 3 | GGCCAGTGGAAGAGGCAGGC;CAGTGGCCAGTGGAAGAGGC;GAAGAGGCAGGCTGGAGCTC | 0.29;1.08;1.73 |
| CCDC42 | 1.030 | 1.778 | 3 | GGAGTTTGAGACTCCACAGA;GTGGCTGGAGACAGGTTGGG;GTTTGAGACTCCACAGAAGG | 0.59;0.86;1.63 |
| C2orf49 | 1.030 | 1.248 | 3 | ACCGCCCACATCCCCCGCCA;TGTGGGCGGTCGCAGCTGCA;ACCATGGCGGGGGATGTGGG | 0.12;0.18;2.78 |
| G6PD | 1.030 | 1.905 | 3 | AGCGCAGGTAACCGGCCGGG;CGAAGCGCAGGTAACCGGCC;ACGAAGCGCAGGTAACCGGC | 0.61;1.23;1.25 |
| ATP5A1 | 1.029 | 1.234 | 3 | CATCTTTGCAGTTACTCCGC;CTTTGCAGTTACTCCGCAGG;GGCTGCAGAAGTACCGCCTG | -0.02;0.39;2.72 |
| CXXC1 | 1.028 | 1.751 | 3 | AACCTGCACAGACCACTCGG;CCACTCGGCGGCGTCCCAGG;GCGAACCTGCACAGACCACT | 0.29;1.4;1.4 |

| | | | | | |
|---|---|---|---|---|---|
| C10orf76 | 1.025 | 1.985 | 3 | TCGGTGGCGTCGAGAGCGAG;CGGTGGCGTCGAGAGCGAGC;AACGGCGCGGGTGGTAGAGG | 0.78;1.11;1.19 |
| HIVEP2 | 1.024 | 1.832 | 3 | GCACGGCCCGGCCACCGCTG;CCTCCCCCGCTACCTGACAA;GCGACGTGCACCGGCGCGCA | 0.57;1.07;1.43 |
| U2AF2 | 1.024 | 1.556 | 3 | CGTCGAAGTCCGACATGCTG;AAGCTGCACAGGGCCCTACG;ACGCGGCCGCCTCAGCATGT | -0.57;1.64;2.0 |
| TNXB | 1.021 | 1.602 | 3 | CTGTTGCAGGGGACAAGTGA;CCTGTTGCAGGGGACAAGTG;TGGCTCAGCCCCTGTTGCAG | 0.1;1.23;1.74 |
| PGLS | 1.019 | 1.941 | 3 | AGAACACCGAGATGAGGCCC;AACTCGAGAACACCGAGATG;GAACACCGAGATGAGGCCCG | 0.83;0.88;1.34 |
| SAG | 1.019 | 1.292 | 3 | TTCAGGCTCATCTGGCAAGA;GTACCAGCTTGCTCAGAACA;TACCAGCTTGCTCAGAACAG | 0.17;0.28;2.61 |
| ITGB6 | 1.018 | 1.672 | 3 | AAATGAAAACAGAGGCTACC;AGCAAGAACTGAAACGAATG;TAAGACTGAAATGAAAACAG | 0.49;0.8;1.77 |
| MID1IP1 | 1.018 | 1.382 | 3 | CAGTGGCTGGACTGTGACTG;TGCAAGCGGCTGCTACGCAA;GCTACGCAAGGGCGCTGCTG | 0.29;0.34;2.42 |
| PCDHAC1 | 1.016 | 1.141 | 3 | TGTAGCGTGTTGGTGGAACG;CCTGGAGAGCACGAGCCGCG;CAGGAGCGTGCTCTTCCCCG | -0.07;0.17;2.94 |
| TGM2 | 1.015 | 1.592 | 3 | TTGGAGGGTCTCGCCGCCAG;GCGGTGGCTCCTTCCACTGG;AGGGTCTCGCCGCCAGTGGA | 0.16;1.11;1.77 |
| PYCR3 | 1.014 | 1.280 | 3 | AGGCAACAAGATGGCAGCTG;CGGCGTCCGAGGCAACAAGA;AAGCCCACGCGCCGCGGAGA | 0.04;0.46;2.54 |
| MRPL3 | 1.014 | 1.477 | 3 | ACGTGGACGCAGTAGCCGTG;CCACGGCTACTGCGTCCACG;CCATTGCGAAAACTTCCCCA | 0.33;0.56;2.15 |
| TRIM49 | 1.013 | 1.631 | 3 | CAGGAAAAGAAATCCCCCGC;CCACTGGGATCCAGCCAGCG;CCCCGCTGGCTGGATCCCAG | 0.42;0.81;1.81 |
| RABGAP1 | 1.010 | 1.523 | 3 | AGGCGGCGGAGCCTCCGGGA;CCCGCCGCTCGCCGTCCCGG;GGAGCCTCCGGGACGGCGAG | -0.81;1.56;2.28 |
| MARK1 | 1.010 | 1.197 | 3 | CGGCCCGAGCCGGCTGCGCG;GGCGGCCGCCAAGTGCCTCT;GCGCGGCACGCAGCGCCCAG | -0.02;0.29;2.76 |
| RPS25 | 1.010 | 1.548 | 3 | TTCGCAATGGTAAGCTTCAG;CCGACATCTTGACGAGGCTG;CTATTCTCCGAGCTTCGCAA | 0.26;0.86;1.91 |

| | | | | | |
|---|---|---|---|---|---|
| NDUFAF7 | 1.009 | 1.399 | 3 | ATGAGTGTACTGCTGAGGTC;ACACTCATGCTGAAATTCGC;TCAGCATGAGTGTACTGCTG | 0.25;0.49;2.29 |
| GBX2 | 1.006 | 1.652 | 3 | GAAGCCGGCGTACTTATCTC;CCCGCCGGAGATAAGTACGC;CGGCTTCGCGCGCTCCCCAG | 0.21;1.24;1.57 |
| AIFM1 | 1.002 | 1.603 | 3 | GTCGCCGAAATGTTCCGGTG;AAGGAGGAGGTCCCGAATAG;CGTGCGTGAGAGGAAAGGGA | 0.43;0.74;1.84 |
| SLC35A4 | 1.002 | 1.341 | 3 | TGACAAGGTGAGTGGCCGTG;GACAAGGTGAGTGGCCGTGG;CCGGCTGCAGATCACCCCCA | 0.27;0.28;2.46 |
| RBM5 | 1.000 | 1.429 | 3 | GCTGCTTCGGTCTCTCCTTG;AGAGACCGAAGCAGCAGTAG;TAGCCGCCGAACCTTGTTGG | 0.24;0.61;2.15 |
