## Supplementary material for "Illuminating host-mycobacterial interactions with functional genomic screening to inhibit mycobacterial pathogenesis": Table S4

**Table S4 Secondary CRISPR-Cas9 KO target genes and sgRNA target sequences. Related to Figure 3.**

| Barcode Sequence | Gene Symbol | Gene ID |
|---|---|---|
| AAAAAACCCGTAGATAGCCT | NDUFA12 | 55967 |
| AAAAACCTAACAAATCCAAG | LRP2 | 4036 |
| AAAAACTGCAATAGCAAACT | YPEL5 | 51646 |
| AAAAATACCGGAATCTGAAG | INO80E | 283899 |
| AAAAATACTAGCCAATGGCC | MAP2K5 | 5607 |
| AAAAATCAGAAATAATCCAT | NO_CURRENT_1 | NO_CURRENT_1 |
| AAAAATTGTGAAAGTAAAGA | ATP5E | 514 |
| AAAACAACTACATACAGGTG | ABI1 | 10006 |
| AAAACACATCAGTATAACAT | RHOA | 387 |
| AAAACAGCTACTATAATCCC | NO_CURRENT_2 | NO_CURRENT_2 |
| AAAACAGGACGATGTGCGGC | NO_CURRENT_3 | NO_CURRENT_3 |
| AAAACATCGACCGAAAGCGT | NO_CURRENT_4 | NO_CURRENT_4 |
| AAAACGGCAAGAAGAACAAG | ZRSR2 | 8233 |
| AAAACTACGTACCATTCTGT | TBXAS1 | 6916 |
| AAAACTGCACATAACTCCGA | PHF6 | 84295 |
| AAAACTGTACGAGTGACGGA | VIPR2 | 7434 |
| AAAAGAAAGGTATTGCTTGG | TMEM30A | 55754 |
| AAAAGACATGTCCCCTCGCT | NO_CURRENT_5 | NO_CURRENT_5 |
| AAAAGAGAATTGTACACTGC | KPNA1 | 3836 |
| AAAAGAGATGAGCCGAAGTG | NLRP3 | 114548 |
| AAAAGCGTTCGGGGTGACTG | IKBKE | 9641 |
| AAAAGCTCTGTGGTGCAGTG | STRAP | 11171 |
| AAAAGGAACGAATTCAGGTA | APIP | 51074 |
| AAAAGGGAATTATATGACAA | DNAJA1 | 3301 |
| AAAAGGTCCATTGGATACAG | MSL2 | 55167 |
| AAAAGTATAATGACAGATGT | TIFA | 92610 |
| AAAATAGCAGTAAACTCAAC | NO_CURRENT_6 | NO_CURRENT_6 |
| AAAATATTTGGATCCACAGG | RBM6 | 10180 |
| AAAATCACTTAAATCACTCT | NO_CURRENT_7 | NO_CURRENT_7 |
| AAAATCCACTGTTACCAATG | KMT2C | 58508 |
| AAAATCCTTTCGCCAGATTA | NO_CURRENT_8 | NO_CURRENT_8 |
| AAAATGCCAGAACATGTCCT | MAP2K5 | 5607 |
| AAAATGGATTATCCTGAGAT | CASP3 | 836 |
| AAAATGGGATGGTGCTAACA | NO_CURRENT_9 | NO_CURRENT_9 |
| AAAATTCAGAGGAGTTACAT | MMP13 | 4322 |
| AAAATTGCAAATCCTGAGAG | TLR2 | 7097 |
| AAACAAAAATCCCTAGATTG | NO_CURRENT_10 | NO_CURRENT_10 |
| AAACAATAAACTTGGAGGAC | HSPA13 | 6782 |
| AAACAATCAAAGGACCGAGA | RB1 | 5925 |
| AAACACAAAAGAATTCACAG | ZNF699 | 374879 |
| AAACACAGGACATGAGGGTA | IRGM | 345611 |
| AAACAGGTCTTCTTGATCCA | NR2C2 | 7182 |
| AAACCAGCTAGTTATTTACA | NO_CURRENT_11 | NO_CURRENT_11 |
| AAACCCATACCATTGACTCT | NO_CURRENT_12 | NO_CURRENT_12 |
| AAACCGACAACAGACTGATA | HDAC2 | 3066 |
| AAACCTCTGGGTTTCACAAG | NO_CURRENT_13 | NO_CURRENT_13 |
| AAACCTGACTGGCTGAAAGT | NDUFC1 | 4717 |
| AAACCTGGCAAATAGAGCCG | NO_CURRENT_14 | NO_CURRENT_14 |

| | | |
|---|---|---|
| AAACGAGATCGAGAAAGGTA | NO_CURRENT_15 | NO_CURRENT_15 |
| AAACTCAAAATCAAGCTTGG | INO80B | 83444 |
| AAACTCACCCGACCTTGACA | GBP5 | 115362 |
| AAACTCTATTTCCAAAACCA | NO_CURRENT_16 | NO_CURRENT_16 |
| AAACTCTTTGCATATAGGCA | TLR1 | 7096 |
| AAACTGCACATACCATATGC | BABAM2 | 9577 |
| AAACTGGATCATCTCAGACA | IL10 | 3586 |
| AAACTGTAGTGCAGGGTCAG | NO_CURRENT_17 | NO_CURRENT_17 |
| AAACTTACAATTAGAGACAG | ANKRD50 | 57182 |
| AAAGAAAAGAGGTTTACACT | NO_CURRENT_18 | NO_CURRENT_18 |
| AAAGAAAGAGGAATAGTAGC | NO_CURRENT_19 | NO_CURRENT_19 |
| AAAGAACCTACAGAGAACTG | MAPK14 | 1432 |
| AAAGAATATAGTATTGGTAC | EIF2AK2 | 5610 |
| AAAGAATCTGAAGTCATGCT | BCL2A1 | 597 |
| AAAGATCATGTACCTAGGCT | NNT | 23530 |
| AAAGATTGCCCCACACCCAT | PTGS1 | 5742 |
| AAAGCCAAAATAGAAAGGCA | PDHB | 5162 |
| AAAGCCCCCATACCTGAATG | ARHGAP33 | 115703 |
| AAAGCGACGTAGGCATACTT | NO_CURRENT_20 | NO_CURRENT_20 |
| AAAGCGTGACCACAGATGAG | RTP1 | 132112 |
| AAAGCTCCAATAATCAACAG | CHUK | 1147 |
| AAAGCTTATATATCAACTCA | NLRC4 | 58484 |
| AAAGGAAATGTAATCCACCA | NO_CURRENT_21 | NO_CURRENT_21 |
| AAAGGACTTCGGCTACGGCG | RRAGC | 64121 |
| AAAGGAGTATATGACACTAA | NO_CURRENT_22 | NO_CURRENT_22 |
| AAAGGATTATTATACACTAA | BMI1 | 648 |
| AAAGGATTATTATACACTAA | COMMD3-BMI1 | 100532731 |
| AAAGGCAATACGTACCACTG | EIF2AK2 | 5610 |
| AAAGGTTGGGCAAAGTCAAG | NO_CURRENT_23 | NO_CURRENT_23 |
| AAAGGTTTACCATCAGCAGA | BMI1 | 648 |
| AAAGGTTTACCATCAGCAGA | COMMD3-BMI1 | 100532731 |
| AAAGTAGAAACAAACTCAGT | NO_CURRENT_24 | NO_CURRENT_24 |
| AAAGTGATGTAGATCTCCCA | RNF25 | 64320 |
| AAAGTTCTCTGGAGTCATGG | SERPINB2 | 5055 |
| AAAGTTTGACGAGTGTGTGC | NDUFA8 | 4702 |
| AAATAAAATCACTCGTCACG | EP400 | 57634 |
| AAATAAGAAAACTCTAAGGC | NO_CURRENT_25 | NO_CURRENT_25 |
| AAATAAGTATAGTAAAGACA | INO80 | 54617 |
| AAATAATATGCATCTCTCGA | NO_CURRENT_26 | NO_CURRENT_26 |
| AAATACAAGCTATAGCGATA | NO_CURRENT_27 | NO_CURRENT_27 |
| AAATACCTTTATCTGAACCC | NO_CURRENT_28 | NO_CURRENT_28 |
| AAATAGAATGCATCGAGCTG | SEC62 | 7095 |
| AAATATCATGCGTGTTATAG | TBK1 | 29110 |
| AAATATCTCATAGCTGGACA | LTA4H | 4048 |
| AAATCATCCGTAGCAAGACG | RPS6KA4 | 8986 |
| AAATCCTTACAAAACCCTAG | TLR2 | 7097 |
| AAATCTAGAACTTACCCCAA | MEMO1 | 51072 |
| AAATGAACACGTGATCCTGG | LGMN | 5641 |
| AAATGAGCATAATTTAACTG | NO_CURRENT_29 | NO_CURRENT_29 |
| AAATGATTAGCACAGTCATG | NO_CURRENT_30 | NO_CURRENT_30 |

| | | |
|---|---|---|
| AAATGCAAGATAGTTGTGGT | RND3 | 390 |
| AAATGCACAGATCGCTGATC | NO_CURRENT_31 | NO_CURRENT_31 |
| AAATGCTACAGAGCTTTGCC | C4orf17 | 84103 |
| AAATGCTTACCTTGACACCA | ABI1 | 10006 |
| AAATGGTATTACAATGCTGC | UQCRB | 7381 |
| AAATGTATGGAATTATGTAG | NO_CURRENT_32 | NO_CURRENT_32 |
| AAATGTATGGCAGAAATGTA | TMEM70 | 54968 |
| AAATGTTGTGAAAGTAAACA | TLR2 | 7097 |
| AAATTCCGGGTAGATATGCC | FOSL2 | 2355 |
| AAATTTACTAAATCTGGGCA | NO_CURRENT_33 | NO_CURRENT_33 |
| AAATTTCGCTTTGATGAACA | CYP2R1 | 120227 |
| AACAAACTAGCCCTCAAGAC | TRIR | 79002 |
| AACAAATCCTCGCTTCTGAA | INO80C | 125476 |
| AACAAATTACCTTCATCACC | CBLL1 | 79872 |
| AACACAATGATAATATCGCC | DNAJA1 | 3301 |
| AACACAGGAAGCTTGCTGTG | PDHA1 | 5160 |
| AACACATCTGACCTACAGGT | MMP1 | 4312 |
| AACACGATTGATGACAGCAC | EDA2R | 60401 |
| AACACGCTCATCACAATGAG | ATP6V0E1 | 8992 |
| AACACTATTCAACCAAGTGA | BCL2A1 | 597 |
| AACACTCACCTCCAGCACAA | TFPT | 29844 |
| AACAGACCTCCCTATTAAGG | SATB2 | 23314 |
| AACAGCACGTTACTTAACAG | NO_CURRENT_34 | NO_CURRENT_34 |
| AACAGCCACCCTTACCACGC | EDA2R | 60401 |
| AACAGCCTGCACCCACACTG | SEC62 | 7095 |
| AACAGGAAACGTGACTAAAG | NO_CURRENT_35 | NO_CURRENT_35 |
| AACAGGAGCGATGAGTCTGT | IFNAR1 | 3454 |
| AACATACAAGCATATTAATG | DYRK1A | 1859 |
| AACATATCCAGCACTGTTTG | ACOD1 | 730249 |
| AACATCTAATGGACTTCCAG | RB1 | 5925 |
| AACATGAACAATCAGCAGGC | NCOA6 | 23054 |
| AACATTACGTTATTTAGGCC | NO_CURRENT_36 | NO_CURRENT_36 |
| AACCAATACGACTCGATACG | NO_CURRENT_37 | NO_CURRENT_37 |
| AACCAGCTAAGTCTAGCTCG | NFRKB | 4798 |
| AACCATAACAAAACCATCCA | HIF1A | 3091 |
| AACCCCACCTTGGCAAGTCT | UBE2L6 | 9246 |
| AACCTCCTGCCTCAGAATGG | ABCB10 | 23456 |
| AACCTCTGTAATGACGACGG | ABL2 | 27 |
| AACCTGATAGTAGTAGAAAG | NOS2 | 4843 |
| AACCTTGAAAAAATTCCTGG | UCHL3 | 7347 |
| AACGAGTTACCTCGTCCGAG | LCN2 | 3934 |
| AACGCAAGCGGCTCTACCAG | RNF25 | 64320 |
| AACGGACACTGATCTCCCAG | CCAR2 | 57805 |
| AACGGCCAAACCCAGCACCT | SHOC2 | 8036 |
| AACGTGTATTGTTCGTTACC | BMI1 | 648 |
| AACGTGTATTGTTCGTTACC | COMMD3-BMI1 | 100532731 |
| AACGTTCCTGACACAAGTGA | NLRP1 | 22861 |
| AACTAGCCCGAGCAGCTTCG | NO_CURRENT_38 | NO_CURRENT_38 |
| AACTAGTAATCATGACAACA | NO_CURRENT_39 | NO_CURRENT_39 |
| AACTCACATGTCCTCGTCGG | ARID3A | 1820 |

| | | |
|---|---|---|
| AACTCCAAATAAGCTGGCTG | NO_CURRENT_40 | NO_CURRENT_40 |
| AACTCCATGCTTATGACACA | UBA7 | 7318 |
| AACTCGATGTAGTCCCCGTG | ALOX5 | 240 |
| AACTGAACAGGTCACCAACA | ATP6V0E1 | 8992 |
| AACTGAGGCCCCCTTTCAAG | IRF9 | 10379 |
| AACTGCCATTCACACAAATG | NO_CURRENT_41 | NO_CURRENT_41 |
| AACTGCGTGTAGTGGCGCGG | TNFRSF6B | 8771 |
| AACTGGACATATCCCAACAG | CS | 1431 |
| AACTTCTGTATACACAGACG | GBP5 | 115362 |
| AACTTCTTATGAATCAAAAA | FAM107B | 83641 |
| AACTTGTCCTGCATAGCTGA | CYP8B1 | 1582 |
| AAGAAAAAAGTCTCGTTGTG | RNF111 | 54778 |
| AAGAAATTTATGATTCAGGG | PPID | 5481 |
| AAGAACAAGACTCTCGGCTA | COX16 | 51241 |
| AAGAACAGCGCCTCGCGCCG | PLD6 | 201164 |
| AAGAACATAAATCTCTGACA | KMT2C | 58508 |
| AAGAAGAATTGGGGATGATG | NO_CURRENT_42 | NO_CURRENT_42 |
| AAGAAGACAGTTACCTGGCA | CASP1 | 834 |
| AAGAATTCAAAATTAAGGTG | PLD1 | 5337 |
| AAGACAAAACGTATCTTGGT | NO_CURRENT_43 | NO_CURRENT_43 |
| AAGACAAGACCATCAAATCC | TRAF6 | 7189 |
| AAGACAAGTTCTCGAACCTG | ANP32B | 10541 |
| AAGACAGATGACGACCGCTG | BRD1 | 23774 |
| AAGACCCAATAACAATGAGG | SIRT1 | 23411 |
| AAGACCCCGGCGCTGGTGAA | ATP5L | 10632 |
| AAGACCCTTACCAGAAGCAG | BAK1 | 578 |
| AAGACTGGGATGCAGTCTTG | SATB1 | 6304 |
| AAGACTTGCCTCCCGAACAA | SATB1 | 6304 |
| AAGAGAGCTACAGTCGTTAG | PDE5A | 8654 |
| AAGAGAGCTCTTTACAATCG | NO_CURRENT_44 | NO_CURRENT_44 |
| AAGAGATCAACATAGTCAAG | ZNF616 | 90317 |
| AAGAGATCACATCTAGGCCA | NO_CURRENT_45 | NO_CURRENT_45 |
| AAGAGATGACATTTGAGCTA | NO_CURRENT_46 | NO_CURRENT_46 |
| AAGAGCATTACCACCCTTCA | SIAH2 | 6478 |
| AAGAGCGGACCTCAGAACCA | FBXW7 | 55294 |
| AAGAGGCACAAAGAATGTAT | NO_CURRENT_47 | NO_CURRENT_47 |
| AAGAGGTAAGCTGTCTGTCG | CFLAR | 8837 |
| AAGAGTAGTAGACGCCCGGG | NO_CURRENT_48 | NO_CURRENT_48 |
| AAGATAGAGAAATCCATCAA | NO_CURRENT_49 | NO_CURRENT_49 |
| AAGATCTTGATGGTTCCGGG | FAM107B | 83641 |
| AAGATGACTAATGCTAAGAG | NO_CURRENT_50 | NO_CURRENT_50 |
| AAGATGCCGATCCAGAATGG | HCAR2 | 338442 |
| AAGATGGACACGCTGACTAA | ZNF616 | 90317 |
| AAGATGTCAGTGATAGTGGC | CYP1B1 | 1545 |
| AAGCAAATCGACCACAGCTA | CTNNBL1 | 56259 |
| AAGCACACTAAAGCCTCCAT | NO_CURRENT_51 | NO_CURRENT_51 |
| AAGCAGAGCGACTCGGTTAA | SERF2 | 10169 |
| AAGCAGCACTACTTACGTCA | IFNAR1 | 3454 |
| AAGCCAAACAGAAACTCCTG | NOD1 | 10392 |
| AAGCCATTGTATAACTCCAG | NO_CURRENT_52 | NO_CURRENT_52 |

| | | |
|---|---|---|
| AAGCGACTGTGGCCCGCACG | ALOX15 | 246 |
| AAGCGCGCCCACCTCGGGAT | UCHL3 | 7347 |
| AAGCGCTTCCCATAGAGTGA | HTR1D | 3352 |
| AAGCGGAGGTCGGAGCGCCG | BNIP3 | 664 |
| AAGCGGGAGCACATTAACCT | VDAC1 | 7416 |
| AAGCTCAAGAAGCGGATGCA | MAP2K7 | 5609 |
| AAGCTGACGGCTGCATCTGT | SOD2 | 6648 |
| AAGCTGTCTGTCATTCACAG | MAP2K6 | 5608 |
| AAGCTTATACAACTTATGGG | NO_CURRENT_53 | NO_CURRENT_53 |
| AAGGAAGGACGAACGTTCCA | NLRP3 | 114548 |
| AAGGAAGTCAACTTCTACAG | CXCR1 | 3577 |
| AAGGAATAAAAATTTGTAGG | NO_CURRENT_54 | NO_CURRENT_54 |
| AAGGAGCCCTTCCCCGCCGG | IRF8 | 3394 |
| AAGGAGCGTCTATGCAACCA | SPEN | 23013 |
| AAGGAGGACTGCATTGTGTG | TMEM70 | 54968 |
| AAGGATAACATCGTTACCAC | AHCTF1 | 25909 |
| AAGGATATGGAAACAAAAGT | CASP1 | 834 |
| AAGGATGTAAATGAAGACCC | BABAM2 | 9577 |
| AAGGATGTTTGCAAGAACAG | TBK1 | 29110 |
| AAGGCACCTTTATATGAAGT | FBXO38 | 81545 |
| AAGGCACTTTCATTGATGTG | DET1 | 55070 |
| AAGGCAGTCACCTTGACGAG | NO_CURRENT_55 | NO_CURRENT_55 |
| AAGGCATTGGACACACGACT | PPID | 5481 |
| AAGGCCACCAGACATGTAGT | PDHB | 5162 |
| AAGGCGATAACATCTATGAA | UBE2E1 | 7324 |
| AAGGCGCGCGAATGTGGCAG | NO_CURRENT_56 | NO_CURRENT_56 |
| AAGGCGTCCATCATGTCGCG | CYP1B1 | 1545 |
| AAGGCTCTTCATAAATTCGT | NO_CURRENT_57 | NO_CURRENT_57 |
| AAGGCTGAGACCAAGAACTC | NELFE | 7936 |
| AAGGCTGCCCTGATGAACCA | BAHD1 | 22893 |
| AAGGCTTTGGACTCTCGGCA | BCORL1 | 63035 |
| AAGGGAAAGCGCCGAGATGA | SERF2 | 10169 |
| AAGGGATGAATGATCCACGA | ZNF699 | 374879 |
| AAGGGCGTGCTCCAGAACAC | SLC7A11 | 23657 |
| AAGGGGTTCAAAATGGAAGT | CYBB | 1536 |
| AAGGTACAGAAATGGCAGTA | NO_CURRENT_58 | NO_CURRENT_58 |
| AAGGTACGTTGATGCCACTT | NO_CURRENT_59 | NO_CURRENT_59 |
| AAGGTCTACCCCATTGACCA | BAP1 | 8314 |
| AAGGTGCAAAAATGCCCCAA | NO_CURRENT_60 | NO_CURRENT_60 |
| AAGGTTATGCAAGGTCCCAG | CTNNB1 | 1499 |
| AAGGTTGGGGAATCACTTGA | NO_CURRENT_61 | NO_CURRENT_61 |
| AAGTACACAACAGTTTCGCC | UCHL5 | 51377 |
| AAGTAGGAAATCCATAGTGT | NFKB1 | 4790 |
| AAGTATAAGGAGGGGGACAC | IRF9 | 10379 |
| AAGTATGTAGAGAGTTCCAG | MBNL1 | 4154 |
| AAGTCAGGTGCTATTCACAA | NO_CURRENT_62 | NO_CURRENT_62 |
| AAGTCCCAAAAAAGGTGTCC | CBLL1 | 79872 |
| AAGTGAACGACATCTCATCT | RB1 | 5925 |
| AAGTGAATTAGCACAGAAGT | NO_CURRENT_63 | NO_CURRENT_63 |
| AAGTGACAGATGGGCAGGCG | NO_CURRENT_64 | NO_CURRENT_64 |

| | | |
|---|---|---|
| AAGTGAGGACTACCTACACA | NIPBL | 25836 |
| AAGTGATAAACAATTGGGAT | NO_CURRENT_65 | NO_CURRENT_65 |
| AAGTGGGCCATACCAGCCGA | SLC25A19 | 60386 |
| AAGTGTACCCTAACTAGCCG | HIF1A | 3091 |
| AAGTGTCAGTCGCTTCTCCG | FOXO4 | 4303 |
| AAGTGTGTGCATAGCAGGGT | NO_CURRENT_66 | NO_CURRENT_66 |
| AAGTGTTGTTTGATCAGTCA | RAD21 | 5885 |
| AAGTTAACATCCCCAGATGT | VIPR2 | 7434 |
| AAGTTACAAGGGATCGCCAA | LTA4H | 4048 |
| AAGTTAGCTTAATCCTAGCA | NO_CURRENT_67 | NO_CURRENT_67 |
| AAGTTCACATTAGAAGGGCA | NO_CURRENT_68 | NO_CURRENT_68 |
| AAGTTCATACCTGACCCCCA | PTGS1 | 5742 |
| AAGTTGAAGTAGACGAACTG | ARMCX2 | 9823 |
| AAGTTTCCGAGCAAAGCCAA | MOSPD1 | 56180 |
| AAGTTTGATGATCGGCCCAA | NUFIP2 | 57532 |
| AAGTTTGCCTACAATGACCA | SUV39H1 | 6839 |
| AATAAACGTAGCCATCACTT | LRP2 | 4036 |
| AATAAAGACAAGATGCACTT | UCHL3 | 7347 |
| AATAATCAGGCATACCATCT | EZH2 | 2146 |
| AATAATCTTACCTAGACTGG | PIAS1 | 8554 |
| AATAATTACACATAGTGGGC | NO_CURRENT_69 | NO_CURRENT_69 |
| AATACAGAGCTTGTAGGGAA | CHEK2 | 11200 |
| AATACCAAAATGCTCTTGGT | RND3 | 390 |
| AATACCTAGCCACACTCGAG | TYK2 | 7297 |
| AATACCTGACTTCAGGTCAA | SIRT1 | 23411 |
| AATACCTGGAGAATTCCCTA | GBP5 | 115362 |
| AATACTAACCTTGAACCGAC | STAG2 | 10735 |
| AATACTCATTAATATATCAC | PPARGC1A | 10891 |
| AATAGCTACAACCAGCCTAA | PIAS1 | 8554 |
| AATATATTACAACCGGCTGC | NO_CURRENT_70 | NO_CURRENT_70 |
| AATATCAAACGGGTGCACGT | NO_CURRENT_71 | NO_CURRENT_71 |
| AATATCACACAGAATAGTGA | STAG2 | 10735 |
| AATATCTGAATCTACCCCCG | LARP4B | 23185 |
| AATATCTGATATAATAAATG | NO_CURRENT_72 | NO_CURRENT_72 |
| AATATGATTTGAATAGGCAG | TMEM70 | 54968 |
| AATATGGCCCGAGCAGTGTT | TMEM70 | 54968 |
| AATATGGGCTGGAATCTCGT | GLMN | 11146 |
| AATATGTACCTTTGAGACAG | PHIP | 55023 |
| AATATTAGGATGGTTCACAC | MAP3K7 | 6885 |
| AATCAAGAACTTATCTACGA | TBK1 | 29110 |
| AATCAAGCATCCTTAAAATG | DYRK1A | 1859 |
| AATCAAGTAAAAGCCAACGC | NO_CURRENT_73 | NO_CURRENT_73 |
| AATCAATCCCTATATAACAG | NO_CURRENT_74 | NO_CURRENT_74 |
| AATCACATCTTACATCCGAT | NO_CURRENT_75 | NO_CURRENT_75 |
| AATCAGCTACCTTCACCAAG | ABL1 | 25 |
| AATCAGCTATTGCTATGCTA | NO_CURRENT_76 | NO_CURRENT_76 |
| AATCAGTCCAATTGTTCCAC | UCHL3 | 7347 |
| AATCATAACCTGGCAAGATG | NO_CURRENT_77 | NO_CURRENT_77 |
| AATCATATTCTATGGCATCA | TMEM70 | 54968 |
| AATCCAGATAGTCACCGCAG | NR2C2 | 7182 |

| | | |
|---|---|---|
| AATCTACGAGAAAACCGTGG | CYFIP1 | 23191 |
| AATCTCTACAGAGAAGGCAG | CS | 1431 |
| AATCTGCTGTAACTTATATC | NO_CURRENT_78 | NO_CURRENT_78 |
| AATCTGGATGAAACCTCTTG | NAIP | 4671 |
| AATCTTACCTCTCATCGTAG | SNRNP70 | 6625 |
| AATCTTACTCGTCCTCCTTG | NO_CURRENT_79 | NO_CURRENT_79 |
| AATCTTATGATAAATATGGG | NO_CURRENT_80 | NO_CURRENT_80 |
| AATCTTGGTGAAGTACTCAT | BAK1 | 578 |
| AATGAACCTTATGAGAAGGA | PITPNB | 23760 |
| AATGACATTTAGTGCTTGGG | NO_CURRENT_81 | NO_CURRENT_81 |
| AATGAGTACCGTGTTGGTGA | SUV39H1 | 6839 |
| AATGAGTACGGATCTCGGAT | KHSRP | 8570 |
| AATGCAAAGAACACAGTCCA | HSPA4 | 3308 |
| AATGCATGCGGTCTATCCGT | ACACA | 31 |
| AATGCATTTGGAACATTGTG | C4orf17 | 84103 |
| AATGCTCAAGAAAGAACACG | SCN5A | 6331 |
| AATGGAACACAAGATGTATG | PCGF6 | 84108 |
| AATGGACCAATTGACTGGAG | TBK1 | 29110 |
| AATGGAGATCAACCATTCTA | UBE2E1 | 7324 |
| AATGGCAAACACATAGCCGT | CACNA2D2 | 9254 |
| AATGGCTCTAATGAAGATAG | BMI1 | 648 |
| AATGGCTCTAATGAAGATAG | COMMD3-BMI1 | 100532731 |
| AATGGGTTGCCGGGAGTAAA | NO_CURRENT_82 | NO_CURRENT_82 |
| AATGGTCAAGGTCTCAAGCA | OR7C2 | 26658 |
| AATGGTCATCACCAACAGAA | STAG2 | 10735 |
| AATGTACTCATCAATGCTCT | MAP2K6 | 5608 |
| AATGTGAAGCCAAATTCAAA | DNAJB6 | 10049 |
| AATGTTACCACCTTGCAAGA | NO_CURRENT_83 | NO_CURRENT_83 |
| AATTAAAAGAGGCACGAAGC | PHF6 | 84295 |
| AATTAAAGTGATAAGCATAG | NO_CURRENT_84 | NO_CURRENT_84 |
| AATTAGCCTGTCTCATAACT | NO_CURRENT_85 | NO_CURRENT_85 |
| AATTCATATTGAGAGAGATG | SMPD1 | 6609 |
| AATTCTGCCCCGTGACCCGG | ZRSR2 | 8233 |
| AATTCTTGCAGCAAGCGTAG | UTY | 7404 |
| AATTGATGATGCCCTGCACT | SOD1 | 6647 |
| AATTGGGACCCTCAGCAGAG | CHUK | 1147 |
| AATTGTAAATCCCAGTGTTG | NO_CURRENT_86 | NO_CURRENT_86 |
| AATTGTGTTAAAGAAAACCA | RNF111 | 54778 |
| AATTTAAGGAGGTTCCTGAG | IRF9 | 10379 |
| AATTTACGCAAGAAACAAGG | DET1 | 55070 |
| AATTTATTGTGGGCTTCACA | CCL20 | 6364 |
| AATTTCACTGCAAACCCACG | CAT | 847 |
| AATTTCTGCAGACATAGGTG | NO_CURRENT_87 | NO_CURRENT_87 |
| ACAAAAATGCCAATGCCGAT | TMEM30A | 55754 |
| ACAAAAGATGATAGTGTGGG | TRAF6 | 7189 |
| ACAAACCTCAGCCCTAACGG | SOD2 | 6648 |
| ACAAACGACCTTGAGCAGGG | NO_CURRENT_88 | NO_CURRENT_88 |
| ACAAACTAGTAAGTTGCTCT | CDKN2C | 1031 |
| ACAAAGCAAGTTCTCAAAAG | NO_CURRENT_89 | NO_CURRENT_89 |
| ACAAAGCCTAGGTTCTCACA | PDE3A | 5139 |

| | | |
|---|---|---|
| ACAAATCAACTTCACCCACG | CUBN | 8029 |
| ACAACACCACATTTGAAAAG | IRGM | 345611 |
| ACAACAGTCCCACATGAAGG | NO_CURRENT_90 | NO_CURRENT_90 |
| ACAACCAGACATCTTGACAA | PRR14L | 253143 |
| ACAACGGATTCAGAAATGGG | CYP4A11 | 1579 |
| ACAAGAAATCAGGAAACGGT | PDE5A | 8654 |
| ACAATGACTGTCCAGGCCCG | TNFRSF1A | 7132 |
| ACAATGGGTCCCTGACCCCC | ZNF641 | 121274 |
| ACAATTCCACAGGCCAGCCA | IRF9 | 10379 |
| ACACAAAGCTGTCGTAGTCT | CFLAR | 8837 |
| ACACAACAGAATCCAAGCTC | SNRNP70 | 6625 |
| ACACAAGCGAGGTTCCACGC | NUP153 | 9972 |
| ACACACCATCTACCAGTCTG | ACADS | 35 |
| ACACACGGCACTCATGACGT | CYP1B1 | 1545 |
| ACACACTTGAGATTCTGCCG | SUV39H1 | 6839 |
| ACACATCAAGAGATGATAAA | APIP | 51074 |
| ACACATCGCAAAGCTCGCTG | SPEN | 23013 |
| ACACCACTTACCCTTTCCGG | NENF | 29937 |
| ACACCAGATGTGTTCGCAGA | ALOX5 | 240 |
| ACACCATATTAGTCTTGTGA | NO_CURRENT_91 | NO_CURRENT_91 |
| ACACCCAGACCAGTGGACAT | CASP9 | 842 |
| ACACCCATTCTCATAACGGA | NO_CURRENT_92 | NO_CURRENT_92 |
| ACACCCCCCAACGTGTCCCT | JUNB | 3726 |
| ACACCGAAGCACCTGTACGT | NO_CURRENT_93 | NO_CURRENT_93 |
| ACACGAAGTTCTGCTCGTAG | SPINT1 | 6692 |
| ACACGGCAGCTGGCCAATGA | CCL20 | 6364 |
| ACACGGTCATCATGAACCCC | OR7C2 | 26658 |
| ACACTATGTCTTGTAACTGT | PCGF6 | 84108 |
| ACACTATTGAATTAACACAT | PLD1 | 5337 |
| ACACTGCAAGTGGACATCAA | TGFB1 | 7040 |
| ACACTTGAGTCAGACGTCAC | PAXIP1 | 22976 |
| ACAGAAATGCTTGACTTCTG | RHOA | 387 |
| ACAGAAGAAGAAATCTGACT | FAM107B | 83641 |
| ACAGAAGGAGTGTCCTGACA | LTA4H | 4048 |
| ACAGAAGTCAGGATGTGCAC | EZH2 | 2146 |
| ACAGAATTGATACTAACTGG | FBXW7 | 55294 |
| ACAGACAAGGGTGCTGAACA | CASP1 | 834 |
| ACAGACCAAGGAAGCAAACT | LAMTOR3 | 8649 |
| ACAGACCAATATATCAACTG | NO_CURRENT_94 | NO_CURRENT_94 |
| ACAGATAACAACATCCCTCA | RNF25 | 64320 |
| ACAGATGTGATCTTAACTGT | PPARG | 5468 |
| ACAGATTCTGTCTCAGTCCG | RNF111 | 54778 |
| ACAGATTGGCGCCTACTGCG | SPINT1 | 6692 |
| ACAGCAAACAGAATTGGACA | ARNT | 405 |
| ACAGCAAGCCAGTGAACCTG | RAC2 | 5880 |
| ACAGCAATCAGTTCTCAACT | KHSRP | 8570 |
| ACAGCAATGATTGACACTGG | ABCB10 | 23456 |
| ACAGCCACAACATAGCGGGC | SLC7A3 | 84889 |
| ACAGCCCTCACGAGCCCGAA | NO_CURRENT_95 | NO_CURRENT_95 |
| ACAGCGCTCTCGTGTACTAT | NO_CURRENT_96 | NO_CURRENT_96 |

| | | |
|---|---|---|
| ACAGCTATAGACATGCTCGG | PLD1 | 5337 |
| ACAGCTATGCAGTTATCACA | RIPK2 | 8767 |
| ACAGCTCCATCAGAGTACTC | RND1 | 27289 |
| ACAGCTCTACGAACTCGACG | ARID3A | 1820 |
| ACAGCTCTAGGGTCACAGAA | VDR | 7421 |
| ACAGGAATTCCCTACCTACG | GC | 2638 |
| ACAGGACAGAAAGCACGATC | ABCB10 | 23456 |
| ACAGGATGTTTCGCATTAAG | UQCRB | 7381 |
| ACAGGATTGATCAGTTTCTG | FAM107B | 83641 |
| ACAGGCAAAGATCGCAGGCG | GSDMD | 79792 |
| ACAGGCACGGATAAAAGCAA | LARP4B | 23185 |
| ACAGGCCAGCTTCAGGACTT | OR7C2 | 26658 |
| ACAGGCCATTGTAGTCGGCG | BAHD1 | 22893 |
| ACAGGGAAACAGGACTGCCG | SIAH2 | 6478 |
| ACAGGGACCATATGATGTGG | PARK7 | 11315 |
| ACAGGGACTACTTACGATGA | VIPR2 | 7434 |
| ACAGGGCGCTCCATATTCGC | ABI1 | 10006 |
| ACAGGGTGGTGAAGTAGGCC | SLC7A5 | 8140 |
| ACAGGTGATGTCCATTGAAG | CHMP6 | 79643 |
| ACAGGTTCTTATTCATTGAC | NO_CURRENT_97 | NO_CURRENT_97 |
| ACAGTACCTCTTGGCGCTGG | UBE2E1 | 7324 |
| ACAGTAGGGATATATTCTCC | RAC1 | 5879 |
| ACAGTATATTTATGGGAATG | NO_CURRENT_98 | NO_CURRENT_98 |
| ACAGTCGCAGGTCTAGTCCT | NCOA6 | 23054 |
| ACAGTGTATAAACTCCCACA | TBK1 | 29110 |
| ACAGTTACATGCTACAGGTG | SUPT7L | 9913 |
| ACAGTTCCTAGACTTCTATG | ZRSR2 | 8233 |
| ACAGTTGAATCGAAGATACT | SEC62 | 7095 |
| ACATAACAGGGATTTGAACA | PIP4K2A | 5305 |
| ACATACAGTATAGGAAACCT | NO_CURRENT_99 | NO_CURRENT_99 |
| ACATACGGCTCATATGCTGA | MLF2 | 8079 |
| ACATACTCTAAGTCAGGCAG | RAD21 | 5885 |
| ACATCAAAAGATAACCACTC | TGFB1 | 7040 |
| ACATCAACGAATTGCCAGTT | UCHL3 | 7347 |
| ACATCAAGAACCCAGATTGG | KPNB1 | 3837 |
| ACATCCAAAGTGTGAATGTA | CXCL3 | 2921 |
| ACATCCCTGAAGAGTAACTG | MAP2K5 | 5607 |
| ACATCCTCCAGATCAGCAGT | GC | 2638 |
| ACATCCTGAAGCCCTATGTG | CYP4A11 | 1579 |
| ACATCTGAAGGATCCGGACA | CAT | 847 |
| ACATCTTCTTTGGATCGGGC | NO_CURRENT_100 | NO_CURRENT_100 |
| ACATGAACCGTATTTCTACT | PDK4 | 5166 |
| ACATGATCGAAGAGTATCGC | KIF2B | 84643 |
| ACATGCTATATACTTTAGTG | NO_CURRENT_101 | NO_CURRENT_101 |
| ACATGCTGGTGAACAAGATG | FBXW7 | 55294 |
| ACATGGATTACAGTCAATCA | UTY | 7404 |
| ACATGGCACAAACCAGACTT | PITPNB | 23760 |
| ACATGTTTCCCATCCAAATG | PPID | 5481 |
| ACATTACCCTCTTCTCCCAG | CD84 | 8832 |
| ACATTAGTGGGACATACAGG | FBXW7 | 55294 |

| | | |
|---|---|---|
| ACATTAGTTAAACACCTATG | NO_CURRENT_102 | NO_CURRENT_102 |
| ACATTGAGCTCCATAGTGAT | NDUFA8 | 4702 |
| ACATTTATCTACCCACTGGG | ACOD1 | 730249 |
| ACCAAATGAAAACCGATTGG | NIPBL | 25836 |
| ACCAAGCTGTATGGGCCCGA | IRF8 | 3394 |
| ACCACACTTGATGACCCCCT | MYD88 | 4615 |
| ACCACCGCTCTCGGTTCCGG | IKBKB | 3551 |
| ACCAGAGATTCGCAAACCAG | CASP9 | 842 |
| ACCAGATCCCCATAGCCGGG | NR1H3 | 10062 |
| ACCAGCCAGTCCAAAACTCC | LTB4R | 1241 |
| ACCAGCCTTTCAGAATGGCA | NQO1 | 1728 |
| ACCAGGCGTCGTCCTTAAGG | ALOX15 | 246 |
| ACCAGGCTGCTGAATAACGA | LAMTOR2 | 28956 |
| ACCAGTTAATAGCACCAGGT | PHC3 | 80012 |
| ACCATAACCACATTGTCCGA | KIF2B | 84643 |
| ACCATACCCAAATTATTCCG | USP7 | 7874 |
| ACCATCACCCGTATTGGAGT | MLF2 | 8079 |
| ACCATGCAAATCACAATCAC | PPARGC1A | 10891 |
| ACCATGTAATGCAGTTAGCA | ABCB10 | 23456 |
| ACCATTTAGGCTGGGTCACT | APIP | 51074 |
| ACCCAAAGCGCTAATTCACA | MAP3K7 | 6885 |
| ACCCAATGTGGCGGAGCCGA | NO_CURRENT_103 | NO_CURRENT_103 |
| ACCCACCCTGAGTCGCATGA | BAP1 | 8314 |
| ACCCAGACATATTACCCTGT | SCTR | 6344 |
| ACCCAGGTGGGCAAACGCAG | TRIR | 79002 |
| ACCCATGAGTTAAGTTTTCT | NO_CURRENT_104 | NO_CURRENT_104 |
| ACCCATGATGTAACCATAGC | NO_CURRENT_105 | NO_CURRENT_105 |
| ACCCATTGAGAGTCGCCTGA | NO_CURRENT_106 | NO_CURRENT_106 |
| ACCCCCGGTTAGGACCCTTA | ATP6V0E1 | 8992 |
| ACCCGCCAGGCTCAACTCCA | INO80B | 83444 |
| ACCCGGGGGTATCATAACAG | NCOA6 | 23054 |
| ACCCTCGGAGTCCTAAAACC | NNT | 23530 |
| ACCCTGAAGATCCTTGACTT | MAPK9 | 5601 |
| ACCCTGTGGATGAGAGACCA | C6orf15 | 29113 |
| ACCGAATCCTTTAATGAGCT | UCHL5 | 51377 |
| ACCGGCACGTTGCGGTCGAA | ACTR5 | 79913 |
| ACCGGCCAGCCATTGTCAGG | NOD2 | 64127 |
| ACCGGCGCCTCTGTCTGAGA | PCGF6 | 84108 |
| ACCGGGCCGCTCCTCGCGAG | KIAA1211L | 343990 |
| ACCGTCTGTGGATAGGAGAG | RAC2 | 5880 |
| ACCTACCCTTGTAATTGAGG | STAT2 | 6773 |
| ACCTATTGTCCCTTCAAGCT | NO_CURRENT_107 | NO_CURRENT_107 |
| ACCTCAATCGGATTCAGCAA | NO_CURRENT_108 | NO_CURRENT_108 |
| ACCTCCTATAGAAGTGCCAG | RNF111 | 54778 |
| ACCTGATCGCAACAAATAGT | PRKN | 5071 |
| ACCTGCGGAAGACGTTCCGA | TYK2 | 7297 |
| ACCTGCTGCGCGCCGGCTGG | CXCL2 | 2920 |
| ACCTGCTGCGCGCCGGCTGG | CXCL3 | 2921 |
| ACCTGGACAAGTAGTTCCAA | MMP13 | 4322 |
| ACCTGGATTGTTATCCCGAA | PDHB | 5162 |

| | | |
|---|---|---|
| ACCTGTTAGGGGAACACTGT | NO_CURRENT_109 | NO_CURRENT_109 |
| ACCTGTTCAAACGTGCCACT | GC | 2638 |
| ACCTTACAATAAGTTATATT | NO_CURRENT_110 | NO_CURRENT_110 |
| ACCTTCAGAGCAAATGTCAG | BAP1 | 8314 |
| ACCTTGCAGAACGTCGAGCT | ARRB1 | 408 |
| ACGAACCATTCGAACCCTTG | TRAF6 | 7189 |
| ACGACAAGCCCCATCCACTG | FAM214B | 80256 |
| ACGACAAGTTCACAGTGGCG | HSH2D | 84941 |
| ACGACCACTCAGCGTACAAG | CYFIP1 | 23191 |
| ACGAGTGTAATAGCTCACGT | PTGS1 | 5742 |
| ACGATCGGTAATGGTCTGTT | NO_CURRENT_111 | NO_CURRENT_111 |
| ACGATGAAGCCAAAGGTGTG | CXCR1 | 3577 |
| ACGATGTGAACATCGAATCA | RB1 | 5925 |
| ACGCACAGCATGAGTCTGGA | BNIP3 | 664 |
| ACGCCCATGACGGAGAGTCC | NDUFA1 | 4694 |
| ACGCGCACCCACAAGCGGGA | GSDMD | 79792 |
| ACGCGCTGAACGTGTTTGTG | IZUMO2 | 126123 |
| ACGCTCAGCACCCGCTATGC | NO_CURRENT_112 | NO_CURRENT_112 |
| ACGCTGACGAGTAAAAGCGG | NO_CURRENT_113 | NO_CURRENT_113 |
| ACGCTGGATAACATTCTGGG | ZNF641 | 121274 |
| ACGGAAAGACCTCGCTATTC | NO_CURRENT_114 | NO_CURRENT_114 |
| ACGGAAGATTCGTTCCCGGC | MKL1 | 57591 |
| ACGGACTCTCCTATAACCAG | KMT2D | 8085 |
| ACGGAGGCTAAGCGTCGCAA | NO_CURRENT_115 | NO_CURRENT_115 |
| ACGGCAGCTCGCCATCATCG | BAK1 | 578 |
| ACGGCCGCTCGGGAGCCTGG | NDUFC1 | 4717 |
| ACGGCCTCATTATGATACCC | NO_CURRENT_116 | NO_CURRENT_116 |
| ACGGCGCCTTCCTCGACCCG | CYP1B1 | 1545 |
| ACGGGCCAAAGGAGCTTTCG | PDAP1 | 11333 |
| ACGGGCTACCTGACCATCGG | MAP2K7 | 5609 |
| ACGGTACATGCGCATGAGTC | NO_CURRENT_117 | NO_CURRENT_117 |
| ACGGTGCAGTCTCACACCAA | GBP5 | 115362 |
| ACGGTGCCAAAGTGCCTCCA | C4orf17 | 84103 |
| ACGGTGGGGATGGACCTACT | NO_CURRENT_118 | NO_CURRENT_118 |
| ACGGTTATGGTCTCATGGGG | NO_CURRENT_119 | NO_CURRENT_119 |
| ACGTAAAAACAGAACGCTGT | TBXAS1 | 6916 |
| ACGTACACCAGCGTCACGAT | SLC7A5 | 8140 |
| ACGTAGTCTGCGACCTTGTG | CYP27B1 | 1594 |
| ACGTCAACTGCTGGAGTGGG | NO_CURRENT_120 | NO_CURRENT_120 |
| ACGTCCATACTGTCGGCTAC | NO_CURRENT_121 | NO_CURRENT_121 |
| ACGTCGTCCTTATGCAAGGG | MMP9 | 4318 |
| ACGTCGTTTAGCACCCGGCT | NO_CURRENT_122 | NO_CURRENT_122 |
| ACGTGCATGCAGTTTGTGAG | GBP7 | 388646 |
| ACGTGCCTGAAGGCAAACGG | ITGB2 | 3689 |
| ACGTGGCAATGAAGGACACG | EBI3 | 10148 |
| ACGTGGGGACATATACGTGT | NO_CURRENT_123 | NO_CURRENT_123 |
| ACGTGTGTCCTTTAAACACG | NO_CURRENT_124 | NO_CURRENT_124 |
| ACGTTCGAGTACGACCAGCT | NO_CURRENT_125 | NO_CURRENT_125 |
| ACGTTGAGGTCCTAACTCTG | NO_CURRENT_126 | NO_CURRENT_126 |
| ACTAAAGCAATCAGGTAGCA | LTA4H | 4048 |

| | | |
|---|---|---|
| ACTAAAGGACAAGTCACCAC | HIF1A | 3091 |
| ACTAAGCTTATCTACAACCG | NO_CURRENT_127 | NO_CURRENT_127 |
| ACTACATAAGCACCTCACCT | PPARGC1B | 133522 |
| ACTACTCGGAAGACTTGCCG | MMP9 | 4318 |
| ACTAGAGTCATGATCAGCGA | NO_CURRENT_128 | NO_CURRENT_128 |
| ACTAGTGATAAGTTCACCAT | NO_CURRENT_129 | NO_CURRENT_129 |
| ACTATTTAATATTGGTAAGT | NO_CURRENT_130 | NO_CURRENT_130 |
| ACTCACCCGGGGGTACCCAG | NUDT17 | 200035 |
| ACTCACGGTTTCCAGTGACA | GBP7 | 388646 |
| ACTCAGCACACGTTAGAGAG | NO_CURRENT_131 | NO_CURRENT_131 |
| ACTCCAAGAAACGCAAACAG | CCAR2 | 57805 |
| ACTCCAGATATTGCACCAGA | TBK1 | 29110 |
| ACTCCAGGCCAACGCCACAG | ALOX5 | 240 |
| ACTCCATTCGCTGCGACACC | ITGB2 | 3689 |
| ACTCCCAAAAGTTTGACCAG | NOS2 | 4843 |
| ACTCCCATTTGGATGCACAA | HNRNPF | 3185 |
| ACTCCCGGTTAGCCCAGTCC | BRK1 | 55845 |
| ACTCCTACAGGGGAAATTGA | TGFB1 | 7040 |
| ACTCCTCCCATCAAGAACTG | NO_CURRENT_132 | NO_CURRENT_132 |
| ACTCCTTAGATATTGATCAG | MAPK9 | 5601 |
| ACTCGACTACGGCGTCACCG | NFKB2 | 4791 |
| ACTCGTACAAAAGCTGGTAA | SLC7A11 | 23657 |
| ACTCGTTTCACAGGCGCCAC | YPEL5 | 51646 |
| ACTCTCTTGAGATTGCACCA | HSH2D | 84941 |
| ACTCTTCAGTAAATATCTAG | MEMO1 | 51072 |
| ACTCTTCATTACAGAGTGCA | STAG2 | 10735 |
| ACTGAAGGCCAAATCAACCA | STRAP | 11171 |
| ACTGACAATGATGTATATGT | HSPA13 | 6782 |
| ACTGACCCAGAAGCGTCACT | CXCR1 | 3577 |
| ACTGACCTGTCAGGAAGTTG | GSDMD | 79792 |
| ACTGAGAACCGTTTGTGTTG | PPID | 5481 |
| ACTGAGCAGTAGTCACTGCC | TIRAP | 114609 |
| ACTGAGTGGCAGGCTTATCG | IKBKB | 3551 |
| ACTGAGTGGGTAACACGCAT | NO_CURRENT_133 | NO_CURRENT_133 |
| ACTGAGTGTGCAGTTCTGCA | IRF9 | 10379 |
| ACTGCACAATATCATATGCA | HSPA4 | 3308 |
| ACTGCAGAAGTCGGTCTAGG | INO80E | 283899 |
| ACTGCATGACCAGATTCATG | INO80 | 54617 |
| ACTGCATGGTGTCATATGCA | NO_CURRENT_134 | NO_CURRENT_134 |
| ACTGCCAATAGCAATGCTGA | MSL2 | 55167 |
| ACTGCCTACCTGTCGAGCAG | RIPK2 | 8767 |
| ACTGCGGAGCGCCCAATATC | NO_CURRENT_135 | NO_CURRENT_135 |
| ACTGCTACCACATCACAGTT | PDK4 | 5166 |
| ACTGCTCCCGGTCGCCCCTC | NO_CURRENT_136 | NO_CURRENT_136 |
| ACTGCTGCTGACATCTCTTA | NO_CURRENT_137 | NO_CURRENT_137 |
| ACTGCTGCTGTCTTCTAAAT | NO_CURRENT_138 | NO_CURRENT_138 |
| ACTGCTTCGTACACGAGTGA | RIPK3 | 11035 |
| ACTGGAAGCACGAATGACAG | NFKB1 | 4790 |
| ACTGGAATCAACAACAATCG | MGA | 23269 |
| ACTGGAGACAGGCTGGACTC | ATP5E | 514 |

| | | |
|---|---|---|
| ACTGGAGAGGCAATAATTTG | NO_CURRENT_139 | NO_CURRENT_139 |
| ACTGGAGCGGTACCCAAAGG | ARMCX2 | 9823 |
| ACTGGATCTGGACGGGCCAA | PDAP1 | 11333 |
| ACTGGCAACACATCCACTGA | ARNT | 405 |
| ACTGGGACGAGGTGCGTACG | MAP2K6 | 5608 |
| ACTGGGTACTGGCACGCACT | MEX3B | 84206 |
| ACTGGTGAGCAGCCTGCTAG | EDA2R | 60401 |
| ACTGGTGCTGAAGTGTAAGG | AGER | 177 |
| ACTGGTTTAAGCCAGTTCAC | TMEM30A | 55754 |
| ACTGTAGGCTTATTACACTT | ABL2 | 27 |
| ACTGTGAGGACTGTTAAGGC | UTY | 7404 |
| ACTGTGCCCATGGTCATGGT | DLAT | 1737 |
| ACTGTGGTCCATATTTCTTG | NO_CURRENT_140 | NO_CURRENT_140 |
| ACTGTTCTACTCAGCCTCTG | PIGY | 84992 |
| ACTTACAGCAGACAACCTCG | HMGN2 | 3151 |
| ACTTACCCCATCCGATCAGG | VIPR2 | 7434 |
| ACTTACGCTTCATCAATGTT | UBE2H | 7328 |
| ACTTACGGCACTCGCATGCC | NO_CURRENT_141 | NO_CURRENT_141 |
| ACTTCAAAGTCAATCCCACA | ARR3 | 407 |
| ACTTCACCAAGATTGCCACC | BAK1 | 578 |
| ACTTCAGATCGCTCTCAGAA | SLC5A8 | 160728 |
| ACTTCAGTTCGGCGTAGTCA | NO_CURRENT_142 | NO_CURRENT_142 |
| ACTTCGGTTTGGGCGCAGTG | CXCL1 | 2919 |
| ACTTCGGTTTGGGCGCAGTG | CXCL2 | 2920 |
| ACTTCGGTTTGGGCGCAGTG | CXCL3 | 2921 |
| ACTTCTCAGATAATGAGCAC | ALOX15 | 246 |
| ACTTCTGCAGTACGAGAACG | INO80E | 283899 |
| ACTTGCAGAGCTGCAACCCC | HAMP | 57817 |
| ACTTGGGAGTGATGATTCCA | STAT2 | 6773 |
| ACTTGTGATGCATGTAGCCT | PLD6 | 201164 |
| ACTTTAAGATAACACTTAGA | NO_CURRENT_143 | NO_CURRENT_143 |
| ACTTTCTCTCGGTACTTGCG | UBA7 | 7318 |
| ACTTTGCGGAGGAATTCACT | RPS6KA4 | 8986 |
| ACTTTGTCTAGTGCTTCCAT | HIF1A | 3091 |
| AGAAAAACCTGATCATCATG | SEC62 | 7095 |
| AGAAAAACGACCCCGTCCCT | DNAJB6 | 10049 |
| AGAAAAATTAACCCATGAAC | UCHL5 | 51377 |
| AGAAAACTCCGAAACAACGC | BCORL1 | 63035 |
| AGAAAAGTCATCAAAGCAGA | DPY30 | 84661 |
| AGAAACACTCCTGGTACCAT | ABL1 | 25 |
| AGAAACTTCAGCCAACTGTG | MAPK9 | 5601 |
| AGAAATATATCTCTGCACCT | SLC5A8 | 160728 |
| AGAAATCATGTTCACGTGAT | IKZF5 | 64376 |
| AGAAATCCTACTCCACCCAT | EIF4G3 | 8672 |
| AGAACAACAAAAGGAAGAAG | MOSPD1 | 56180 |
| AGAACACCAACTCTGCCCTT | BAP1 | 8314 |
| AGAACACGGAGAGTGAGGGT | RTP1 | 132112 |
| AGAACATGGAAATTTATGCT | BABAM2 | 9577 |
| AGAACCACGAGTTCTCACCC | SLC3A2 | 6520 |
| AGAACCCAGACGCCAGCGGT | NO_CURRENT_144 | NO_CURRENT_144 |

| | | |
|---|---|---|
| AGAACGGAGGAGCTTTGTAG | NO_CURRENT_145 | NO_CURRENT_145 |
| AGAACTCACGGCAAAGTTTG | IZUMO2 | 126123 |
| AGAACTTTCATATAATGCAG | SPEN | 23013 |
| AGAAGAAAAGTAATAAAACC | FBXW7 | 55294 |
| AGAAGAACCAGCAGGCTCTG | UCHL5 | 51377 |
| AGAAGACCGAGGCGCTTCAA | NO_CURRENT_146 | NO_CURRENT_146 |
| AGAAGCCTCCCCCATACCTG | UBE2L6 | 9246 |
| AGAAGGAGATGATCCTTGGA | NAIP | 4671 |
| AGAAGGATTTAAATATTGAG | PRKAA1 | 5562 |
| AGAAGGCCAGGTAATTGTCA | TLR9 | 54106 |
| AGAAGGCCATGCAAAGACGG | NO_CURRENT_147 | NO_CURRENT_147 |
| AGAAGGGACACGTCTTGTAC | PTGES2 | 80142 |
| AGAAGTAAGACTGCACAGAC | UCHL3 | 7347 |
| AGAAGTGAATGGCATCAATC | PIGL | 9487 |
| AGAAGTGGCCACTCTCATGA | NIPBL | 25836 |
| AGAAGTGTGACTACTGGATC | LCN2 | 3934 |
| AGAAGTTGCGCATCATGCTG | CYP1B1 | 1545 |
| AGAATCAACCCACCAAGCAG | POU2F1 | 5451 |
| AGAATCAGTTTAATTAGAAG | NO_CURRENT_148 | NO_CURRENT_148 |
| AGAATCTTCAATAGACACAT | SOD1 | 6647 |
| AGAATTCCCTCGAAGCCGAA | SLC7A3 | 84889 |
| AGACAAAGAGAAATCTAAGG | BMI1 | 648 |
| AGACAAAGAGAAATCTAAGG | COMMD3-BMI1 | 100532731 |
| AGACAAGCTTGCTATTCCAA | PDE3A | 5139 |
| AGACAATATCTTCCGTAACG | RRAGA | 10670 |
| AGACACCAAAGGAGTCACAC | ZNF616 | 90317 |
| AGACACTGCGGCCAGTACGG | PAGR1 | 79447 |
| AGACAGCTCTTCCTTCGATG | MKL1 | 57591 |
| AGACATCAGCTGAGTGGACG | PTGIS | 5740 |
| AGACATGAGTGGTTTCGTAG | KAT2A | 2648 |
| AGACCAAAGAAAATGACCAG | TNFRSF1A | 7132 |
| AGACCAAAGCTGCTTAATCG | NO_CURRENT_149 | NO_CURRENT_149 |
| AGACCAAGTAATAAAGCAGA | FAM107B | 83641 |
| AGACCACGTGGAGCAGATGT | HSP90B1 | 7184 |
| AGACCATCTAGGCAAGAGTT | NO_CURRENT_150 | NO_CURRENT_150 |
| AGACCCAAAATGACAAAGCT | NO_CURRENT_151 | NO_CURRENT_151 |
| AGACCCCGTAGGCAGGACGT | NO_CURRENT_152 | NO_CURRENT_152 |
| AGACCGCAATGCGCGCCTAG | NLRP12 | 91662 |
| AGACGCAGACACACATTGCA | NO_CURRENT_153 | NO_CURRENT_153 |
| AGACGCTTCACAGAAAACTG | NO_CURRENT_154 | NO_CURRENT_154 |
| AGACTCATCTTGTTGCATCA | AHR | 196 |
| AGAGAAAAAAGAGTGCCGGC | SQSTM1 | 8878 |
| AGAGAAAAGGAGAATCCGAA | FOS | 2353 |
| AGAGAATGAATTGATGTATG | PDHB | 5162 |
| AGAGAATTAGGGCTTTGCTA | NO_CURRENT_155 | NO_CURRENT_155 |
| AGAGACATCAGAGATGCGCT | MLF2 | 8079 |
| AGAGACCTGGGAAAATGACT | ASH2L | 9070 |
| AGAGACTAACGAACTGGCAA | METTL3 | 56339 |
| AGAGATGGCTGGAATTGTCC | SIRT1 | 23411 |
| AGAGATGTGTTACTGCACAC | PAXIP1 | 22976 |

| | | |
|---|---|---|
| AGAGATTATTGCCTTCCACG | TP73 | 7161 |
| AGAGATTGATCTCAATCTTG | NLRP3 | 114548 |
| AGAGCCAACCAACTCTTCCG | SATB2 | 23314 |
| AGAGCCTCTGAGGAAAGCAT | PDE3A | 5139 |
| AGAGCTCCATAATTTGCCTA | NO_CURRENT_156 | NO_CURRENT_156 |
| AGAGCTGTGAGAATACCCCA | SATB2 | 23314 |
| AGAGCTGTTTGACAAAGTGG | CHEK2 | 11200 |
| AGAGGACAAGAGCCCATCAG | RAD23A | 5886 |
| AGAGGACTACGAATTGGCCA | SLC7A3 | 84889 |
| AGAGGGACGAATCAGATGAG | POU2F1 | 5451 |
| AGAGGGATTGGGAGCTTGAC | NO_CURRENT_157 | NO_CURRENT_157 |
| AGAGGGCTTAACCTTGCACA | NO_CURRENT_158 | NO_CURRENT_158 |
| AGAGGTGATCTCTAACATCA | IRGM | 345611 |
| AGAGGTGCACGGTCCCATTG | TNFRSF1A | 7132 |
| AGAGTAACAATGGATGTGTA | NO_CURRENT_159 | NO_CURRENT_159 |
| AGAGTAACCAGAAGTATGGG | NO_CURRENT_160 | NO_CURRENT_160 |
| AGAGTCAAAGTCAACCCAAA | NO_CURRENT_161 | NO_CURRENT_161 |
| AGAGTGAGACAGTCTAATAT | NO_CURRENT_162 | NO_CURRENT_162 |
| AGAGTGGCTCACGACCCGGG | NFRKB | 4798 |
| AGATACCCTCAATGTTGTGT | ARNT | 405 |
| AGATATCGTACCCAAAACTG | PPID | 5481 |
| AGATATGCCGAACCAGAACG | SLC7A3 | 84889 |
| AGATATTGTGAATCTGTGTG | CYFIP1 | 23191 |
| AGATCATTCCTCAGTGCCGG | MMP9 | 4318 |
| AGATCCTGTCTAGGCCAGAG | NLRP1 | 22861 |
| AGATCTGATGCTTCAAGTTC | UQCRB | 7381 |
| AGATGAAGTAGAGTTTACTG | CSDE1 | 7812 |
| AGATGATGAGCACGTTTGGG | CYP24A1 | 1591 |
| AGATGATTGAGATGTTCGTG | BABAM1 | 29086 |
| AGATGCCCTGGTGACCAGCG | SLC6A4 | 6532 |
| AGATGCGAACTCACATTATG | HIF1A | 3091 |
| AGATGGAAGTGAACATGAGG | ZNF641 | 121274 |
| AGATGGCTATGGATCAAGCA | RNF111 | 54778 |
| AGATGGTAGAATCACCAAGA | DLAT | 1737 |
| AGATGTAAGCTCGTTCTGGT | RND1 | 27289 |
| AGATGTATGATCCCAAAACG | USP7 | 7874 |
| AGATGTCTAACGATAAGGAG | ARNT | 405 |
| AGATTAAAGACGCAACCGGC | NO_CURRENT_163 | NO_CURRENT_163 |
| AGATTACTGAAGTGGACCAT | NO_CURRENT_164 | NO_CURRENT_164 |
| AGATTCATTCACGAGTTGGG | NO_CURRENT_165 | NO_CURRENT_165 |
| AGATTTATTTAGTCAACGAG | EP400 | 57634 |
| AGCAACATGAGATTGAATCC | STAT2 | 6773 |
| AGCAACTTTGACTGCTGTCT | CCL20 | 6364 |
| AGCAATGGGAAGACCCCAGT | MPC1 | 51660 |
| AGCACACCCCACCTCTGACA | PRKN | 5071 |
| AGCACACGGAGATCACCACC | HTR7 | 3363 |
| AGCACCAAGCAGGTCATAGG | HIF1A | 3091 |
| AGCACCGAATAGGGACAAAG | KDM4A | 9682 |
| AGCACCGGCGTCATTCCCTG | SPHK1 | 8877 |
| AGCACCTGACTCAGATGCCG | ANP32B | 10541 |

| | | |
|---|---|---|
| AGCACTAGAGATATATAGAT | NO_CURRENT_166 | NO_CURRENT_166 |
| AGCACTGATTACTACAAGAG | PDHA1 | 5160 |
| AGCACTGCGAACAGGCAAGG | KMT2D | 8085 |
| AGCACTGTTCCTGTCGATAG | NO_CURRENT_167 | NO_CURRENT_167 |
| AGCAGATGGGAACTTGCCAG | IRGM | 345611 |
| AGCAGATTCAAGCATGCAGT | NO_CURRENT_168 | NO_CURRENT_168 |
| AGCAGCACAACCAGCCAAGG | NELFE | 7936 |
| AGCAGCCAGGAGCAAACTCT | CCL20 | 6364 |
| AGCAGGATGGAGGGACCACA | ASH2L | 9070 |
| AGCAGTGAGATGGCGCTGGA | NFKB1 | 4790 |
| AGCAGTGCTGTGTCTACTAG | ASH2L | 9070 |
| AGCATACCAAGAATAGCTTA | NO_CURRENT_169 | NO_CURRENT_169 |
| AGCATCAAGAAATGGATGAG | AHCTF1 | 25909 |
| AGCATCCAGTACCTTCCAGT | PITPNB | 23760 |
| AGCATCCCGACATGTATGAG | IKBKE | 9641 |
| AGCATCCCGGAGCTCGTCCT | ARF1 | 375 |
| AGCATCGACAAGGCTCCCTG | EP400 | 57634 |
| AGCATTAAAGGACTGACTGA | SOD1 | 6647 |
| AGCATTCTCACCAAGACCGA | NO_CURRENT_170 | NO_CURRENT_170 |
| AGCATTGTAGAATGATACGT | APAF1 | 317 |
| AGCCAAATGACTTTGCATTG | NO_CURRENT_171 | NO_CURRENT_171 |
| AGCCAAGAAGCAGAAGACGG | TRIR | 79002 |
| AGCCAAGAGAAATGTCATCA | NOD2 | 64127 |
| AGCCACAACTTAAAGCCCTG | CYP4F11 | 57834 |
| AGCCACTGGTCAGGCTGGCG | HAMP | 57817 |
| AGCCAGCCATTACCTGTAAG | WASF2 | 10163 |
| AGCCATCGCAGATCACATTG | SQSTM1 | 8878 |
| AGCCATGACATCCACAGCAT | ACO2 | 50 |
| AGCCATGAGAAACTAGACAG | PQBP1 | 10084 |
| AGCCCATGTCCCAAATCACA | PDE4A | 5141 |
| AGCCCCACGGACGAGGACAG | PPARGC1B | 133522 |
| AGCCGTCGATGATGAAAAGG | NLRP12 | 91662 |
| AGCCTACATTTATTAAACTG | ANP32B | 10541 |
| AGCCTATTCAAAAATAAAGG | PIP4K2A | 5305 |
| AGCCTCGATTGGCCACATTT | ATP5L | 10632 |
| AGCCTCTGAGAGCACACAAG | NAIP | 4671 |
| AGCCTGACTACTCGTACCTG | CSNK1D | 1453 |
| AGCCTGTCCGCCATTAGCAG | ARMCX2 | 9823 |
| AGCGAAGGATTGCTATCCAG | RND1 | 27289 |
| AGCGAGGCAAGTCGTACCTG | ACO2 | 50 |
| AGCGATCTGGACACTCTCCA | NO_CURRENT_172 | NO_CURRENT_172 |
| AGCGATTCACGTATTAGATG | NO_CURRENT_173 | NO_CURRENT_173 |
| AGCGCGCAGAGCGCAGTTGA | RHOG | 391 |
| AGCGCGTAGTCCCGGAAGAA | IZUMO2 | 126123 |
| AGCGCTCTGGTTGCATCCCT | NO_CURRENT_174 | NO_CURRENT_174 |
| AGCGGGAGATTCACCAGGAC | BRK1 | 55845 |
| AGCGTGCCACACCAGGACAT | UBE2L6 | 9246 |
| AGCGTGTTCTGGGGCTTCGT | ATP6V0E1 | 8992 |
| AGCGTTCTCGAACCCTTGAG | NELFE | 7936 |
| AGCTAGCGATGGCTCTAAGT | NO_CURRENT_175 | NO_CURRENT_175 |

| | | |
|---|---|---|
| AGCTAGGCGCAATAGTGCCA | NUFIP2 | 57532 |
| AGCTAGTGCTAACCTACCTA | NO_CURRENT_176 | NO_CURRENT_176 |
| AGCTCACATCTCGAACCATG | YPEL5 | 51646 |
| AGCTCCTAGAGACTACAGAC | KPNB1 | 3837 |
| AGCTCGCCATGTCGGTTCTC | NO_CURRENT_177 | NO_CURRENT_177 |
| AGCTCGGATTCCATGAACCT | CTNNBL1 | 56259 |
| AGCTCTAAAGGATCGCCGCA | CREBBP | 1387 |
| AGCTGAAAGTAGATGAGGTG | NDUFA8 | 4702 |
| AGCTGAGGCAGTACCAGAAG | CHMP6 | 79643 |
| AGCTGCGCGCTACTGGATCA | NO_CURRENT_178 | NO_CURRENT_178 |
| AGCTGCGCTCCCGGACTGAG | BNIP3 | 664 |
| AGCTGCTATTGGTAGTGGAG | MKL1 | 57591 |
| AGCTGGACTCTGTAGAAATC | NO_CURRENT_179 | NO_CURRENT_179 |
| AGCTGGAGGACAGATAACTG | POU2F1 | 5451 |
| AGCTTAATGTGCAGGTCAGA | NO_CURRENT_180 | NO_CURRENT_180 |
| AGCTTCGCACGGAGTGTGTG | NUDT17 | 200035 |
| AGCTTCTACCTGGAGACCTA | PYCARD | 29108 |
| AGCTTTGTCTACAACAGCAT | GBP7 | 388646 |
| AGGAAAAACACCCAGGACAA | OR7C2 | 26658 |
| AGGAAAATCGCCAACTCCTG | NO_CURRENT_181 | NO_CURRENT_181 |
| AGGAAATTCCAACTGGCCCA | BAHD1 | 22893 |
| AGGAACTACTACTAAGCCCC | TNFRSF1A | 7132 |
| AGGAACTGAAACATCAGACG | TNFRSF1B | 7133 |
| AGGAAGAGCCTGGCACATTG | TNFRSF6B | 8771 |
| AGGAAGATTATGATCGCCTG | RHOA | 387 |
| AGGAAGCCATGTTTCCTGAG | RXRA | 6256 |
| AGGACAAATGATGCGAAGGT | NDUFA12 | 55967 |
| AGGACACAAAGCCAATGGGC | MCL1 | 4170 |
| AGGACACACTGCGCTCGGGT | KHSRP | 8570 |
| AGGACCCATCCCAAGTCCCT | RIPK1 | 8737 |
| AGGACTAGCCTCATTGTCAG | PPARGC1A | 10891 |
| AGGACTGCCTGATTGACAAG | RXRA | 6256 |
| AGGACTTTAATTGTGAAGGG | NO_CURRENT_182 | NO_CURRENT_182 |
| AGGAGAAACGAAAGAAACGT | ZRSR2 | 8233 |
| AGGAGAAAGGTTCCAAACAG | ZNF616 | 90317 |
| AGGAGAAGATGCCCGGTGCG | BCL2 | 596 |
| AGGAGAAGCGTCGCATCCGG | FOSL2 | 2355 |
| AGGAGATCGACCTCGTTCGG | NO_CURRENT_183 | NO_CURRENT_183 |
| AGGAGATGAATTGATGACCC | STRAP | 11171 |
| AGGAGCTGAGGAAAGCGCAA | INO80E | 283899 |
| AGGAGCTTGTCCGTAATTAG | GC | 2638 |
| AGGAGTACATCAAGACGGGA | IRAK1 | 3654 |
| AGGAGTCCAGCACTCTCATG | TFPT | 29844 |
| AGGAGTCGCTCCACTGTTGA | GBP7 | 388646 |
| AGGAGTTTATCAATAAGACC | NO_CURRENT_184 | NO_CURRENT_184 |
| AGGATAATAACCCAATGGCA | NO_CURRENT_185 | NO_CURRENT_185 |
| AGGATATCATCATGGTAGCG | NO_CURRENT_186 | NO_CURRENT_186 |
| AGGATCAGAATTATTGACTC | CYBB | 1536 |
| AGGATCTATGGGGTTAGGAT | UBE2H | 7328 |
| AGGATGATGAGCAACACCGA | HTR1D | 3352 |

| | | |
|---|---|---|
| AGGATGATTCTAGACACTCT | ACOD1 | 730249 |
| AGGATGCAGAGAGAAAGGAG | DNAJA1 | 3301 |
| AGGATGCTGAACAAGTACGT | NO_CURRENT_187 | NO_CURRENT_187 |
| AGGATGGATTGAGCAGCGGT | NO_CURRENT_188 | NO_CURRENT_188 |
| AGGATGGGCTCAAATACTAC | PDHA1 | 5160 |
| AGGATTCATAGAGAGTACCT | UTY | 7404 |
| AGGATTTGCCAGTGCGATAG | GBP5 | 115362 |
| AGGCAACTAATCACTGCACA | NO_CURRENT_189 | NO_CURRENT_189 |
| AGGCACGAGTTGGCAAAGAC | DPY30 | 84661 |
| AGGCAGCGTGTATGACGCCC | SLC25A19 | 60386 |
| AGGCATCGGGCACCTATGCA | AHDC1 | 27245 |
| AGGCCACATGCAAAACACTT | NO_CURRENT_190 | NO_CURRENT_190 |
| AGGCCCAGCAAGTGTATGGT | PTGES2 | 80142 |
| AGGCCGTAGGAGGAAGATGG | SAP18 | 10284 |
| AGGCCTTGAAGGACAGTCTG | VDR | 7421 |
| AGGCCTTGCAACACTGACAA | GLMN | 11146 |
| AGGCCTTTACAACATTGGCA | CHUK | 1147 |
| AGGCGACGACCCCAGAAACG | PRKN | 5071 |
| AGGCGCATGTGAACTCCCTG | IL10 | 3586 |
| AGGCGCGGCAGTCCTCTCGG | HELZ2 | 85441 |
| AGGCGCTGGAGACCTTACGA | MCL1 | 4170 |
| AGGCGTCAAGACCTCTGGCA | SLC6A4 | 6532 |
| AGGCTACATGCCACGCCAGG | SETD1B | 23067 |
| AGGCTATGCAGCTTGCAAAG | NFKB1 | 4790 |
| AGGCTCTACTTAGAAGACAC | VIPR2 | 7434 |
| AGGCTGAACAGACAGTGCCA | BCORL1 | 63035 |
| AGGCTGGTGACATTGCCACG | TLR9 | 54106 |
| AGGCTGTGAAGCATCGCTGG | TIRAP | 114609 |
| AGGCTTCAACGTGGAAACCG | ARF1 | 375 |
| AGGCTTGTCGTCAAAACGCA | CSNK1D | 1453 |
| AGGGAAACCTCTATGGGTAA | NO_CURRENT_191 | NO_CURRENT_191 |
| AGGGAAAGCGCCGAGATGAC | SERF2 | 10169 |
| AGGGAACCTGCCCTTGCACT | CDKN2C | 1031 |
| AGGGAAGCGTGATGACAAAG | CXCR4 | 7852 |
| AGGGAGCCAGGGTCCTACAC | HTR2A | 3356 |
| AGGGATAGTTAGGAAACTGA | NO_CURRENT_192 | NO_CURRENT_192 |
| AGGGATCGTTAGGAAGGGAA | NO_CURRENT_193 | NO_CURRENT_193 |
| AGGGATTTCCCAGATGACCC | PIGL | 9487 |
| AGGGCATCGACATCTCAATG | CYFIP1 | 23191 |
| AGGGCCGTGCAGTTGCGGTG | TNFRSF6B | 8771 |
| AGGGCTTCGAGAACTTTGGG | NUDT17 | 200035 |
| AGGGGACTATGACTCCATGA | CXCR4 | 7852 |
| AGGGGAGTCTCCATTATCAT | SLC7A11 | 23657 |
| AGGGGCAGCTCTGAGAAGTG | RXRA | 6256 |
| AGGGTCAGGTGGACCACAGG | BCL2 | 596 |
| AGGGTCAGTCTGTCCTTTTT | NO_CURRENT_194 | NO_CURRENT_194 |
| AGGGTGAGAAAACCACGTTG | SERPINE1 | 5054 |
| AGGGTGAGCACGAATATAGA | NO_CURRENT_195 | NO_CURRENT_195 |
| AGGGTGCGCTATGACCAGCA | PIGL | 9487 |
| AGGGTGGAAGGCTGGATCGA | NO_CURRENT_196 | NO_CURRENT_196 |

| | | |
|---|---|---|
| AGGGTTATCGTCCCAGATGG | SBNO2 | 22904 |
| AGGTAAAGCTGGCTTTCGAG | CHEK2 | 11200 |
| AGGTAAGCCCCTTAGAACTG | NO_CURRENT_197 | NO_CURRENT_197 |
| AGGTACAACTCCCATCAGAT | FBXO38 | 81545 |
| AGGTACCGAAGTGGCATCCG | ACACA | 31 |
| AGGTAGACAATTGCAGCCTG | BCL2L11 | 10018 |
| AGGTAGTTGGAAGGTCCAGT | NO_CURRENT_198 | NO_CURRENT_198 |
| AGGTCGAGTACTCCTTACCC | MMP9 | 4318 |
| AGGTCGGTGACGCTGACGAA | HTR7 | 3363 |
| AGGTGAGAAGGATGAATCGA | VDAC1 | 7416 |
| AGGTGATATCTCTAGTGACA | NO_CURRENT_199 | NO_CURRENT_199 |
| AGGTGATCAGCATGACCGTG | GPR119 | 139760 |
| AGGTGCAGTACGTGATTCAA | BABAM2 | 9577 |
| AGGTGGACCAACAATTTCAG | MMP1 | 4312 |
| AGGTGGGCAAACGCAGAGGC | TRIR | 79002 |
| AGGTGGTGACTTTCGATGCA | PIGL | 9487 |
| AGGTGGTTGCTAAGCGACAG | PRKN | 5071 |
| AGGTGTCTTGAGCCCTATGG | UBA7 | 7318 |
| AGGTGTGTGAAGCATGACAC | CACNA2D2 | 9254 |
| AGGTGTTGCGGCTTTATAAG | NDUFB9 | 4715 |
| AGGTTAACAGGACACGGTCA | NOD2 | 64127 |
| AGGTTCAGTATGCCCCAGAG | ARRB1 | 408 |
| AGGTTCGAAGACACCAGACT | PHIP | 55023 |
| AGGTTGACACACTTATAACG | GSDMD | 79792 |
| AGTAAAGCCTCAGTGTTGTG | MOSPD1 | 56180 |
| AGTAATTAAAGGAATAGGAG | PPID | 5481 |
| AGTACAAGCAGATCTTCCTG | SLC25A6 | 293 |
| AGTACAGTGAAGTTCAAGAT | YPEL5 | 51646 |
| AGTACAGTGTAGCCGTGATG | PARK7 | 11315 |
| AGTACCGCACGGTACAGACA | UBA7 | 7318 |
| AGTACCGTGTCACCACGTAA | MAPK9 | 5601 |
| AGTAGAAAAGGGCGACAACC | BAX | 581 |
| AGTAGAAGTAGACCAAACCA | UTY | 7404 |
| AGTAGACGGACGGTGAGCTG | NO_CURRENT_200 | NO_CURRENT_200 |
| AGTAGCAAATCTGCACCAGA | RIPK2 | 8767 |
| AGTAGCCAAGAAGCAGAAGA | TRIR | 79002 |
| AGTAGCGAATGACGTTGGCA | SLC25A6 | 293 |
| AGTATTAGGTACCTGCCCTA | NO_CURRENT_201 | NO_CURRENT_201 |
| AGTATTGCACAGATTACTCA | NO_CURRENT_202 | NO_CURRENT_202 |
| AGTATTGTACTCACACCCGA | NFRKB | 4798 |
| AGTATTGTGACCACATAATG | NO_CURRENT_203 | NO_CURRENT_203 |
| AGTATTGTGGTGTCGTCAAC | NO_CURRENT_204 | NO_CURRENT_204 |
| AGTCACGCTGTTGCGACGAG | DLAT | 1737 |
| AGTCAGAATTGTTGCTGACA | CD84 | 8832 |
| AGTCAGACATATAACCAACG | NO_CURRENT_205 | NO_CURRENT_205 |
| AGTCAGCCTTCCGTATCCAA | NO_CURRENT_206 | NO_CURRENT_206 |
| AGTCATAACTGAGTGAATCG | NO_CURRENT_207 | NO_CURRENT_207 |
| AGTCCACAAACAAGGCAAGC | LARP4B | 23185 |
| AGTCCATGCACCGATGCAGA | VIPR2 | 7434 |
| AGTCCCATGAGCAGAATCCT | RRAGC | 64121 |

| | | |
|---|---|---|
| AGTCCCCGTCATGGAACACC | APEH | 327 |
| AGTCCGGACAGGACGCTACT | KIAA1211L | 343990 |
| AGTCCTCACTGGTGGACACG | PPARGC1A | 10891 |
| AGTCCTGAATTTATAAACAT | PPARGC1A | 10891 |
| AGTCCTGCTAAAGAAACCAG | PIGY | 84992 |
| AGTCCTGTGTCACCAATCAA | HSH2D | 84941 |
| AGTCGCTGGAGATTATCTCT | MCL1 | 4170 |
| AGTCGTCTGTGATGAATACA | HSP90B1 | 7184 |
| AGTCTAAGCAAATCACGTGG | PRKN | 5071 |
| AGTCTCCAGAGATTATCAGT | MPC1 | 51660 |
| AGTCTGCTATATGTAACGTA | NO_CURRENT_208 | NO_CURRENT_208 |
| AGTCTGTTGATAGTAGACAA | JAM2 | 58494 |
| AGTCTTCTAGAGGGGACCGT | DET1 | 55070 |
| AGTCTTGGCCAATGTCACGG | NO_CURRENT_209 | NO_CURRENT_209 |
| AGTGAAGGGCTTGATATCAA | PPARG | 5468 |
| AGTGAGCAATACGGCCAGGT | LTB4R | 1241 |
| AGTGAGTGACAACCAGATCG | NO_CURRENT_210 | NO_CURRENT_210 |
| AGTGAGTGGCTACTCAGACC | INO80 | 54617 |
| AGTGATCGACTTCGAGAACT | NELFE | 7936 |
| AGTGCAAATATCAATCACCT | NO_CURRENT_211 | NO_CURRENT_211 |
| AGTGCACACAAAAACGTCCA | GC | 2638 |
| AGTGCCGACAGCGCCAACGA | HNRNPF | 3185 |
| AGTGCTACTGAAACTTGCCT | NO_CURRENT_212 | NO_CURRENT_212 |
| AGTGCTGGCCATATTGCTCC | CTNNBL1 | 56259 |
| AGTGCTGTTATTCACTGTCG | KHSRP | 8570 |
| AGTGGACAGCGCACTCAAGA | RND1 | 27289 |
| AGTGGCATTCAACCGCACAC | SATB2 | 23314 |
| AGTGGCGCCTGTGAAACGAG | YPEL5 | 51646 |
| AGTGGGGCGCTAAGTGGGGG | NO_CURRENT_213 | NO_CURRENT_213 |
| AGTGGTCAAGACCTAAGGAG | MMP13 | 4322 |
| AGTGGTCATAAGGTTACACA | EP400 | 57634 |
| AGTGGTTTACCCCCCACATG | PIP4K2A | 5305 |
| AGTGTATGCTACGAACGTAA | CSDE1 | 7812 |
| AGTGTCATATCATCCAACTC | SIRT1 | 23411 |
| AGTGTCCAGGAAACCTGCTC | CDKN2C | 1031 |
| AGTGTCCCAACCTTAAAACA | NUFIP2 | 57532 |
| AGTGTCCCCGTAACCAGAGT | LAMTOR2 | 28956 |
| AGTGTGAATGTAAGGTCCCC | CXCL3 | 2921 |
| AGTGTGCAAGATTAGCCCCG | FOSL2 | 2355 |
| AGTGTGTATCACGTGACCCT | NO_CURRENT_214 | NO_CURRENT_214 |
| AGTGTTTGAAAAAAGGGCGG | NO_CURRENT_215 | NO_CURRENT_215 |
| AGTTACTTCGATCACTGCAG | IKZF5 | 64376 |
| AGTTACTTTGAATCCCGCCA | SLC11A1 | 6556 |
| AGTTAGAATGAAAGTACCAA | SPEN | 23013 |
| AGTTATCCAATCCACTCTGA | CASP5 | 838 |
| AGTTCCCAGGGAATCCGCAT | NO_CURRENT_216 | NO_CURRENT_216 |
| AGTTCCTCAGCAACCACATG | SLC25A1 | 6576 |
| AGTTCTGAGTGTGACCGAGA | BCL2L11 | 10018 |
| AGTTCTGTTCGATAGATGCC | NO_CURRENT_217 | NO_CURRENT_217 |
| AGTTGAATGGACCTCGACTA | NO_CURRENT_218 | NO_CURRENT_218 |

| | | |
|---|---|---|
| AGTTGATAAGCAATACAAAT | AIM2 | 9447 |
| AGTTGATCTTTGGAGCATTG | TBK1 | 29110 |
| AGTTGGAAAATGGTACCACA | ZRSR2 | 8233 |
| AGTTTACCGAATTGTTCCTG | IRF8 | 3394 |
| ATAAAACCCTTGGAGTAACT | NO_CURRENT_219 | NO_CURRENT_219 |
| ATAAAAGTCTCATTAACCCA | NO_CURRENT_220 | NO_CURRENT_220 |
| ATAAACCTTCTCGTGCCCAG | HTR3E | 285242 |
| ATAAAGAAATGTACTTCCGG | APIP | 51074 |
| ATAAGCCACACTACCCGCCT | NO_CURRENT_221 | NO_CURRENT_221 |
| ATAAGGGGAGCACAGTTAGG | NO_CURRENT_222 | NO_CURRENT_222 |
| ATAATGGGTAGTTAAAACCG | NO_CURRENT_223 | NO_CURRENT_223 |
| ATAATGTATAGTGTGTCACA | RIPK2 | 8767 |
| ATAATGTTTGTATAACCCGT | NO_CURRENT_224 | NO_CURRENT_224 |
| ATAATTGGATAAAATTGTCT | IFNAR1 | 3454 |
| ATACAATACTTTGGCGCATA | NO_CURRENT_225 | NO_CURRENT_225 |
| ATACACCGTGCCGAACGCAC | EGFR | 1956 |
| ATACAGATGATCGCTCCGAT | DLAT | 1737 |
| ATACAGCTAAAGCCATGGAA | MEMO1 | 51072 |
| ATACAGCTAAGACCATGTTC | IRF9 | 10379 |
| ATACAGGACCTGATTGTGAG | NO_CURRENT_226 | NO_CURRENT_226 |
| ATACAGTTTGTTCCTCGTGG | MMP9 | 4318 |
| ATACCAATCACTTTAGGAGT | HSPA13 | 6782 |
| ATACCAGATGCGTCCGCTTG | NO_CURRENT_227 | NO_CURRENT_227 |
| ATACCATGTAACGTTTAAGG | NO_CURRENT_228 | NO_CURRENT_228 |
| ATACCATTCTCTGTCTCATG | NO_CURRENT_229 | NO_CURRENT_229 |
| ATACCGTTTGATCACATGGG | ARHGAP33 | 115703 |
| ATACCTTGTGGATAGCATGT | STAG2 | 10735 |
| ATACGAGAGAAACCTTCACA | PHF6 | 84295 |
| ATACGAGGCGCTTTTCTTTG | NO_CURRENT_230 | NO_CURRENT_230 |
| ATACGCATGATTGCAAGAGG | NO_CURRENT_231 | NO_CURRENT_231 |
| ATACTATCACATAATCTGAG | NO_CURRENT_232 | NO_CURRENT_232 |
| ATACTCAGAGCCCGTCACTG | RAD23A | 5886 |
| ATACTCTCACAGGTACATAA | NO_CURRENT_233 | NO_CURRENT_233 |
| ATACTTACGAAAAACTACTG | CTNNB1 | 1499 |
| ATAGAGATAAGACTCATGGC | NO_CURRENT_234 | NO_CURRENT_234 |
| ATAGCAGGACGAGGTTCCTT | NO_CURRENT_235 | NO_CURRENT_235 |
| ATAGCCATAGTCGTAACCTG | RBM6 | 10180 |
| ATAGCCGCCGCTCATTACTT | NO_CURRENT_236 | NO_CURRENT_236 |
| ATAGCCTTGTCAGATAAGGA | SIRT1 | 23411 |
| ATAGCGAGGCTTGTTCAAAG | NO_CURRENT_237 | NO_CURRENT_237 |
| ATAGCGGATGTCCTTGGAAA | NO_CURRENT_238 | NO_CURRENT_238 |
| ATAGCTAATGCATGCATGCA | NO_CURRENT_239 | NO_CURRENT_239 |
| ATAGGCACCTTAAGGGTCTC | NO_CURRENT_240 | NO_CURRENT_240 |
| ATAGGGAAGAACATTGTAGT | NO_CURRENT_241 | NO_CURRENT_241 |
| ATAGGTCATCCACTGGGCGG | NO_CURRENT_242 | NO_CURRENT_242 |
| ATAGTAACGTCAGGGAGTAA | NO_CURRENT_243 | NO_CURRENT_243 |
| ATAGTATGTTAGGCAAAGCG | NO_CURRENT_244 | NO_CURRENT_244 |
| ATAGTCAATAGTGCTCAGAC | ATP5L | 10632 |
| ATAGTGAGGACAACTAGTGA | NO_CURRENT_245 | NO_CURRENT_245 |
| ATAGTGTATTTGACTTAGTA | NO_CURRENT_246 | NO_CURRENT_246 |

| | | |
|---|---|---|
| ATAGTTAGATAAGACTGCTA | EGFR | 1956 |
| ATATAAAAAAGAGTTCTGAT | NO_CURRENT_247 | NO_CURRENT_247 |
| ATATAAAAACTAGAAAAGTC | NDUFS4 | 4724 |
| ATATAAACTGTCGCGGTAAA | NO_CURRENT_248 | NO_CURRENT_248 |
| ATATAAAGTTCTTTGAGTTG | TLR2 | 7097 |
| ATATACTCCTTAGGCATGCG | KAT2A | 2648 |
| ATATAGAGGAGAGGATCGTA | NO_CURRENT_249 | NO_CURRENT_249 |
| ATATAGGGTGCTGGCCTGAG | NNT | 23530 |
| ATATATAGGTAAACAGGACA | NAIP | 4671 |
| ATATCATTGATGGCAGCAAT | MPC1 | 51660 |
| ATATCCGTGGTTGAAAGTGT | NFRKB | 4798 |
| ATATCTGTGAAGACACCTGT | DLAT | 1737 |
| ATATCTTTGACTGATGACGA | EP400 | 57634 |
| ATATGATGCGCTAAGCTCCA | NNT | 23530 |
| ATATGGATACTTAAGGAGGA | ZNF699 | 374879 |
| ATATGGCTACTCGTCCAGCA | PHIP | 55023 |
| ATATTCACACTCATTTAGAC | NO_CURRENT_250 | NO_CURRENT_250 |
| ATATTTAGTGACTCACAGAC | PDK4 | 5166 |
| ATATTTCTTCAGCATTAACG | KDM4A | 9682 |
| ATCAACGACCTGGAGAACTT | MAP2K7 | 5609 |
| ATCAACTACCCGTTCGAGAA | NDUFS8 | 4728 |
| ATCAACTCCACGGATCCCGC | SMPD1 | 6609 |
| ATCACCATGAAAATATCAGA | PRKAA1 | 5562 |
| ATCACGTGATCGGATGGTTC | NO_CURRENT_251 | NO_CURRENT_251 |
| ATCACTCCTACATTCCTGGT | NO_CURRENT_252 | NO_CURRENT_252 |
| ATCAGAGACCAAGGACACAC | ABI1 | 10006 |
| ATCAGCTCTCGACTCTAGCA | IKZF5 | 64376 |
| ATCAGTGGGCGGATGACATT | MPC1 | 51660 |
| ATCAGTTCAAAGCTGCCGAG | EP400 | 57634 |
| ATCAGTTGTCAATATAAGGG | NO_CURRENT_253 | NO_CURRENT_253 |
| ATCATAATTCCTCTGCACAT | EGFR | 1956 |
| ATCATACAGACAGCGCAGTG | KDM4A | 9682 |
| ATCATCAATTTCTTGGTGGG | RAD21 | 5885 |
| ATCATCAGATAACAGCGAGA | INO80E | 283899 |
| ATCATCATTATATTGACCAA | EZH2 | 2146 |
| ATCATCTCACAGACGCCCCA | POU2F1 | 5451 |
| ATCATCTCAGCCAACTGGGT | STAT2 | 6773 |
| ATCATCTTCATCATCTACCG | NUFIP2 | 57532 |
| ATCATGACTCGAGTAAATGC | NFRKB | 4798 |
| ATCATGGACTAGACTAGCCT | NO_CURRENT_254 | NO_CURRENT_254 |
| ATCATTTGGCTCATAACTGG | SEC62 | 7095 |
| ATCCACAGGTTCACTAACGG | NDUFA1 | 4694 |
| ATCCAGCAATCGAACCCCAG | JAM3 | 83700 |
| ATCCAGCTCCTGAATACACA | JAM2 | 58494 |
| ATCCAGTGCGGGCTGCGTCA | CDKN2D | 1032 |
| ATCCAGTTCTCTCCGAGACA | PRR14L | 253143 |
| ATCCCAGATACATACTCGGG | NO_CURRENT_255 | NO_CURRENT_255 |
| ATCCCCAGGAACTCACACGG | RAC2 | 5880 |
| ATCCCTAAGGGTCCTAACCG | ATP6V0E1 | 8992 |
| ATCCGCTTAGAATGGAACAA | PDE5A | 8654 |

| | | |
|---|---|---|
| ATCCGTGATTATACAGGCTG | SLC5A8 | 160728 |
| ATCCTAAACCCTTATATGGT | NO_CURRENT_256 | NO_CURRENT_256 |
| ATCCTAATTCTGGCCAATCG | UTY | 7404 |
| ATCCTCTACAAGCTTAAGCT | ARF1 | 375 |
| ATCCTGAGTATCTCAAAGAG | ARRB1 | 408 |
| ATCCTGTCAAAATAGTCCGA | MCTS1 | 28985 |
| ATCGATATACCGCCATAAAA | NO_CURRENT_257 | NO_CURRENT_257 |
| ATCGCAAACAAAGCCAACTG | AHR | 196 |
| ATCGGTACCTCTTCACATAT | NO_CURRENT_258 | NO_CURRENT_258 |
| ATCGTATCATCAGCTAGCGC | NO_CURRENT_259 | NO_CURRENT_259 |
| ATCGTGGACAGCAGAGAGCT | TIFA | 92610 |
| ATCGTTGCTGACAGGATCTA | NO_CURRENT_260 | NO_CURRENT_260 |
| ATCTAAAAAGCTCCTAGACC | NO_CURRENT_261 | NO_CURRENT_261 |
| ATCTACAACGGGCGCGTGGT | IRF9 | 10379 |
| ATCTCCTGCTTAGAGGACAC | CASP9 | 842 |
| ATCTCGGGTCGACTGCGGAT | NO_CURRENT_262 | NO_CURRENT_262 |
| ATCTCTATAAGACATACTCA | NO_CURRENT_263 | NO_CURRENT_263 |
| ATCTCTATCCGAAATCCCCG | RNF25 | 64320 |
| ATCTCTATCTGACACAACAA | LAMTOR3 | 8649 |
| ATCTCTGGGCGCATATCTGC | BCL2L11 | 10018 |
| ATCTCTTCTCATACAACACG | CYP24A1 | 1591 |
| ATCTGAACAGCTGACCCCTG | KDM4A | 9682 |
| ATCTGAGTCATATCAAGTTG | NO_CURRENT_264 | NO_CURRENT_264 |
| ATCTGGTCATTAGCGATCTG | CD84 | 8832 |
| ATCTGGTGACCCATCTCATG | MAPK14 | 1432 |
| ATCTGTGTGACTGCGGTCGG | NO_CURRENT_265 | NO_CURRENT_265 |
| ATCTTATCAACCAGTCACGG | NO_CURRENT_266 | NO_CURRENT_266 |
| ATCTTCCCTGACTCTCCTGG | NDUFB9 | 4715 |
| ATCTTCTCCGAGAGTGTCAG | ACO2 | 50 |
| ATCTTCTCGACGAAAATGCG | NO_CURRENT_267 | NO_CURRENT_267 |
| ATCTTGGCAAATATAACCGA | KPNB1 | 3837 |
| ATCTTTAGAGTCGGAATATG | COX16 | 51241 |
| ATCTTTAGAGTCGGAATATG | SYNJ2BP-COX16 | 100529257 |
| ATGAACCTGGACAAAAATCA | RIPK2 | 8767 |
| ATGAACTTGTAGTACACAGG | BRD1 | 23774 |
| ATGAAGAGTCGGGAGCAGGG | PIGL | 9487 |
| ATGAAGGCATCGAAACGCTC | MYD88 | 4615 |
| ATGAAGGCTGTGAACGAGCA | PTGES2 | 80142 |
| ATGAATGCTTACACAATGAG | APIP | 51074 |
| ATGACAATCCTGAACTCATT | FAM107B | 83641 |
| ATGACATTGCGCGTCTACGG | NO_CURRENT_268 | NO_CURRENT_268 |
| ATGACCAAGACTCTTCCAGG | GBP5 | 115362 |
| ATGAGAAAGGGCATGACTCA | NO_CURRENT_269 | NO_CURRENT_269 |
| ATGAGATAAACCGACTGCTG | KAT2A | 2648 |
| ATGAGCCGTATGTTGTCAGG | MLF2 | 8079 |
| ATGAGCTGCACATCACAATG | NO_CURRENT_270 | NO_CURRENT_270 |
| ATGAGGTCCCAGCTGTCAGC | RTP1 | 132112 |
| ATGAGTATATCAAATTTCTA | HDAC2 | 3066 |
| ATGAGTTAACAGGTTTATCT | C4orf17 | 84103 |
| ATGATAATATTAACCTCATA | NO_CURRENT_271 | NO_CURRENT_271 |

| | | |
|---|---|---|
| ATGATATACAATCTCACTAA | NO_CURRENT_272 | NO_CURRENT_272 |
| ATGATCTCCTTGTTCCGCCG | IKBKE | 9641 |
| ATGATCTGCGCGTTGATGTG | SOD2 | 6648 |
| ATGATGAAAACAGCACAGCG | SCN5A | 6331 |
| ATGATGAACCAACTCGGCCA | ABL1 | 25 |
| ATGATGACCTTGCAGATGCT | RRAGC | 64121 |
| ATGATGGTCTGCCAAGTGGG | CTNNB1 | 1499 |
| ATGATGGTTCTGATCAGTTC | BAX | 581 |
| ATGATTAATTATCTGCACGG | NO_CURRENT_273 | NO_CURRENT_273 |
| ATGATTGAGTCAATGAAGGA | NNT | 23530 |
| ATGATTGGATCAGTGCAGTA | UCHL5 | 51377 |
| ATGCAAGACAGCCTCCCAGC | NO_CURRENT_274 | NO_CURRENT_274 |
| ATGCAATGACTCGAGCTCAG | CTNNB1 | 1499 |
| ATGCACAAAATGGATTTGGA | CDKN2C | 1031 |
| ATGCAGGATATACGTACTGT | MEMO1 | 51072 |
| ATGCAGTCAGTCAGGCTGGG | CHMP5 | 51510 |
| ATGCATCAAACATACCGGAA | NO_CURRENT_275 | NO_CURRENT_275 |
| ATGCATCCAATAATGACCTG | MGA | 23269 |
| ATGCCACAATCACTTAAGGT | MGA | 23269 |
| ATGCCTTAGACTTAACCTCG | NO_CURRENT_276 | NO_CURRENT_276 |
| ATGCGCAGCTCCAGAATTTT | NO_CURRENT_277 | NO_CURRENT_277 |
| ATGCGGGACTTCAGTCCTAG | ATP5J2 | 9551 |
| ATGCGGGACTTCAGTCCTAG | ATP5J2-PTCD1 | 100526740 |
| ATGCTAAGAAACGTGCCCGG | GPT2 | 84706 |
| ATGCTAAGTACCTGTGAAAG | SATB1 | 6304 |
| ATGCTAATGATTGTAGCTCA | NO_CURRENT_278 | NO_CURRENT_278 |
| ATGCTACTAATGATAGATCC | NO_CURRENT_279 | NO_CURRENT_279 |
| ATGCTAGACTTACCCACACA | PRKN | 5071 |
| ATGCTGCAGCTTTACGATCA | NO_CURRENT_280 | NO_CURRENT_280 |
| ATGCTGTTCAAAGAAGAGCC | OXSM | 54995 |
| ATGCTTTCTCAACTAGAAGG | ZNF641 | 121274 |
| ATGGAAAAGGAGTTTGAAGA | BCL2A1 | 597 |
| ATGGAAGAGCGTCATGACTT | NO_CURRENT_281 | NO_CURRENT_281 |
| ATGGACAGCAGCGCGCCCCG | PYCARD | 29108 |
| ATGGACTCTGGCAACTTATG | NO_CURRENT_282 | NO_CURRENT_282 |
| ATGGACTGGACTGGTACCGG | CDKN2D | 1032 |
| ATGGAGCATCTCGTCCACCA | SERPINB2 | 5055 |
| ATGGAGTGTACAGCTGCGAG | RXRA | 6256 |
| ATGGATACCTCGAGGCAGGG | TRERF1 | 55809 |
| ATGGATATACATATTGCAAG | SLC7A11 | 23657 |
| ATGGCCTCTGGACATCCGAG | ACO2 | 50 |
| ATGGCGCCAGAGGGCAACTG | BNIP3 | 664 |
| ATGGCTCGGAAATGTCCAGT | IRF8 | 3394 |
| ATGGGACATCACAGCTTCCA | SIAH2 | 6478 |
| ATGGGATTGTGGAGGATCGG | CCAR2 | 57805 |
| ATGGGTCACCATACATGGAA | APAF1 | 317 |
| ATGGGTTGCAGTCAAACCAC | NO_CURRENT_283 | NO_CURRENT_283 |
| ATGGTAAACAGCATCTGTGT | APAF1 | 317 |
| ATGGTAGTGCAATGGCCAAG | DET1 | 55070 |
| ATGGTATAGGCGATGCTGAT | HTR1D | 3352 |

| | | |
|---|---|---|
| ATGGTCTCAGAAATAATCAA | EIF2AK2 | 5610 |
| ATGGTGAAATGTCCAAATGA | TRAF6 | 7189 |
| ATGGTGCTTACCCACTGACA | PTGIS | 5740 |
| ATGGTTTAGATCCAAACACA | PITPNB | 23760 |
| ATGTAACGAGTTGTAAGTCA | NO_CURRENT_284 | NO_CURRENT_284 |
| ATGTAAGCTGCTTTCTAGTC | C4orf17 | 84103 |
| ATGTAATAGGGCCCCAACAG | NO_CURRENT_285 | NO_CURRENT_285 |
| ATGTACAAGCTGTTACATAA | NO_CURRENT_286 | NO_CURRENT_286 |
| ATGTACCTTAATGAGTGTGG | MKL1 | 57591 |
| ATGTACGCAGTAGCCAGTCC | NDUFA1 | 4694 |
| ATGTACTATATCCTGGACGA | NLRP12 | 91662 |
| ATGTAGAAACTGAATTGCCC | IRAK1 | 3654 |
| ATGTCATCATAGGATGCTCG | KDM4A | 9682 |
| ATGTCTCTGCTGCACGACGT | KIAA1211L | 343990 |
| ATGTCTTTCCCGGAAGCTAA | FBXO38 | 81545 |
| ATGTGACAAGAAGTGATGCG | JAM2 | 58494 |
| ATGTGCCCCACAGATACACA | SNRNP70 | 6625 |
| ATGTGGAAAGCATGTTATCG | MGA | 23269 |
| ATGTGGCCACCAAATAGGGG | TFPT | 29844 |
| ATGTGGTTCGAGATTCTCCC | NDUFA1 | 4694 |
| ATGTGTCTAGTAAGTGACAA | NO_CURRENT_287 | NO_CURRENT_287 |
| ATGTTAAGGCCAGATCAGAG | METTL3 | 56339 |
| ATGTTAGCACTTACCTTTGC | HMGN2 | 3151 |
| ATGTTCCGGATAGTTCCATT | IDH2 | 3418 |
| ATGTTGCTGAAGAGTCTCGC | HSP90B1 | 7184 |
| ATGTTGGCTATGAACCGCCA | HCAR2 | 338442 |
| ATGTTGGGGGTACATTCAGG | EZH2 | 2146 |
| ATGTTGGTGACCGGACGGAA | HTR3E | 285242 |
| ATTAAACTTTAACTAGACAT | NO_CURRENT_288 | NO_CURRENT_288 |
| ATTAAATACTACAACAGCCA | METTL3 | 56339 |
| ATTAAATGAACTCCTACATA | RIPK2 | 8767 |
| ATTACATTGCTCGCGACACC | RRAGA | 10670 |
| ATTACCTATAGTCCGAAACA | CFLAR | 8837 |
| ATTACTGCATTTCGCCAAGG | ACOD1 | 730249 |
| ATTAGCACGGCGACCTTCTA | NO_CURRENT_289 | NO_CURRENT_289 |
| ATTAGCCGTTGCCATATCAA | NO_CURRENT_290 | NO_CURRENT_290 |
| ATTAGGCCTTTTTCTTAACT | NO_CURRENT_291 | NO_CURRENT_291 |
| ATTATAGTAGGAGACTGACT | PHC3 | 80012 |
| ATTATGAACAGTGTGCATCG | EIF2AK2 | 5610 |
| ATTATGTAGACCATGGAATG | NO_CURRENT_292 | NO_CURRENT_292 |
| ATTATTCTCAAACGTAAGCG | AHDC1 | 27245 |
| ATTCAAACTTAGTCTTGAGT | NO_CURRENT_293 | NO_CURRENT_293 |
| ATTCACCTGCCCCTTCAACG | VDR | 7421 |
| ATTCACTGCAGCTTGAAGCG | NUDT17 | 200035 |
| ATTCAGACATTAGGCCATCG | KMT2C | 58508 |
| ATTCAGGGATACTAAAGCCT | CS | 1431 |
| ATTCATGAGATTTGCCTACA | MPC1 | 51660 |
| ATTCATGCGCCGCCTCCTCT | NO_CURRENT_294 | NO_CURRENT_294 |
| ATTCATTGTGTATGAGCTGT | JAM2 | 58494 |
| ATTCCACTGGTTTCTTTAGC | PIGY | 84992 |

| | | |
|---|---|---|
| ATTCCCATGATAAGATGGAC | IFNAR2 | 3455 |
| ATTCCGTTATGGTCAAAGGT | ANKRD50 | 57182 |
| ATTCGAAAGATGGAACATGC | GPR119 | 139760 |
| ATTCGATGTCCACTACAGGA | HTR3E | 285242 |
| ATTCGCAGACAAGTGACCCC | JAM3 | 83700 |
| ATTCGGAGCCTGGACCACTG | NLRC4 | 58484 |
| ATTCGTTGATGTATATGGAA | UCHL3 | 7347 |
| ATTCTATCTGTTAGAACTAA | CCNC | 892 |
| ATTCTGACTGCCATAATCCA | KPNB1 | 3837 |
| ATTCTGTGACTATGGAACCA | METTL3 | 56339 |
| ATTCTTATGTTCTTCAAACC | NDUFB9 | 4715 |
| ATTGAAAGCCCATTATAAGT | MSL2 | 55167 |
| ATTGACTCATTAATGAAGAC | NO_CURRENT_295 | NO_CURRENT_295 |
| ATTGAGAATTCGTTTCAAGG | NO_CURRENT_296 | NO_CURRENT_296 |
| ATTGATATAGATATTCATCA | HDAC2 | 3066 |
| ATTGCCCCCCTCCAAACACG | CREBBP | 1387 |
| ATTGCGGCAAGAAGGAAGCG | JAK1 | 3716 |
| ATTGCTCTGTCGCATCAATC | NO_CURRENT_297 | NO_CURRENT_297 |
| ATTGGAACTAAGCATTAGAC | CTNNBL1 | 56259 |
| ATTGGCTATGCTGTGGACCT | SLC6A4 | 6532 |
| ATTGGGCTAAATTGTTCTGG | NO_CURRENT_298 | NO_CURRENT_298 |
| ATTGGGTTACTTACTGTCCA | PDE5A | 8654 |
| ATTGTAGGATCGGATGCCAA | LARP4B | 23185 |
| ATTGTATCTTCATACTTGTG | NO_CURRENT_299 | NO_CURRENT_299 |
| ATTGTCTCCTCACCTCACTG | ACO2 | 50 |
| ATTGTGATCAACAGGCCCAA | LGMN | 5641 |
| ATTGTGATGAGCGTGTTCTG | ATP6V0E1 | 8992 |
| ATTGTGGAATTGATGCGTGA | CASP3 | 836 |
| ATTGTGTGCTGGACTAAGGT | NO_CURRENT_300 | NO_CURRENT_300 |
| ATTGTTGTGTCAGATAGAGA | LAMTOR3 | 8649 |
| ATTTAACATCATATTACACC | NO_CURRENT_301 | NO_CURRENT_301 |
| ATTTAACTTACTGCTATATG | RB1 | 5925 |
| ATTTAAGATACTTACACAGT | RAC1 | 5879 |
| ATTTACCTGAATTCTGCTGG | WASF2 | 10163 |
| ATTTAGGCCTGGTGTCACTA | PHF6 | 84295 |
| ATTTAGTAATGCACACCCAG | NO_CURRENT_302 | NO_CURRENT_302 |
| ATTTAGTAGACATTGGGTCT | NO_CURRENT_303 | NO_CURRENT_303 |
| ATTTATAGCAACCTGCTCAG | APAF1 | 317 |
| ATTTATCCCACAGAAAGACC | IRAK1 | 3654 |
| ATTTCAAACGTGAAGCTACA | TLR1 | 7096 |
| ATTTCAGCCTCAGCCAAAAG | SPHK1 | 8877 |
| ATTTCAGCGTCAGCTACACT | AHR | 196 |
| ATTTCCAATACACCCTACAT | NO_CURRENT_304 | NO_CURRENT_304 |
| ATTTCCAGAAAGGACATCAC | NQO1 | 1728 |
| ATTTCCCATTCTGCCTCACA | HTR7 | 3363 |
| ATTTCCGGGACTACCTCATG | MPC1 | 51660 |
| ATTTCCTAAACATCCACCAC | NAIP | 4671 |
| ATTTCCTAACCTGCCACCGT | IFNAR2 | 3455 |
| ATTTCGACATACAAGCACCC | STAG2 | 10735 |
| ATTTCTACAGCATGATCGGG | PPID | 5481 |

| | | |
|---|---|---|
| ATTTGAAAATGTCCGTGCAA | RAC1 | 5879 |
| ATTTGAGCGGCTGTACCAAG | CCAR2 | 57805 |
| ATTTGGATATATTTACAGGC | BCL2A1 | 597 |
| ATTTGTCAGCACATAGCCAA | SLC7A11 | 23657 |
| CAAAAAATATCGTGTCAAGT | UCHL5 | 51377 |
| CAAAACTCACTGTACATGGG | ZNF699 | 374879 |
| CAAAAGAAACTAATGAAACC | CYP24A1 | 1591 |
| CAAAAGCCACGAGTTTGCAC | BABAM1 | 29086 |
| CAAAAGTAGAAGGTAACATT | NO_CURRENT_305 | NO_CURRENT_305 |
| CAAAAGTCTCGCTTGGTCCT | NO_CURRENT_306 | NO_CURRENT_306 |
| CAAAATCACTCACTGCGGAA | NELFE | 7936 |
| CAAAATCCTGTCCAGCCCAT | MMP1 | 4312 |
| CAAAATCTAGCAATTTTTGG | NO_CURRENT_307 | NO_CURRENT_307 |
| CAAAATTGTTAGTAAACTGT | NO_CURRENT_308 | NO_CURRENT_308 |
| CAAACAATCGTCCGCAAATG | LTA4H | 4048 |
| CAAACACAAAATGTACCGAA | LGMN | 5641 |
| CAAACACCGAGCCCACAGCG | CYP27B1 | 1594 |
| CAAACTCCTGAAACCGAGCC | JUNB | 3726 |
| CAAACTGGTACTCGAGAACG | CTNNBL1 | 56259 |
| CAAACTTACTGGATTCAAGC | MAP3K7 | 6885 |
| CAAACTTGGACAGCTCTCTA | UBE2H | 7328 |
| CAAAGAGTCTACCAAGATCA | CASP5 | 838 |
| CAAAGCAAAGCTCAGAGCGA | NLRP12 | 91662 |
| CAAAGCAGACATAAATACAC | CD84 | 8832 |
| CAAAGCCAAGTATTGAAGCA | NO_CURRENT_309 | NO_CURRENT_309 |
| CAAAGGAACCCAACTTCATG | CYP24A1 | 1591 |
| CAAAGGACAAGGCCTCCTCG | JAK1 | 3716 |
| CAAAGTGAGCACAAATCAGG | NCKAP1L | 3071 |
| CAAAGTTGCTCTGGGTCACT | VDAC1 | 7416 |
| CAAATGCAATCAGCGTGCGG | CUBN | 8029 |
| CAAATGCCATTTAGGTTATC | NO_CURRENT_310 | NO_CURRENT_310 |
| CAAATGCCGCAACCGGAGGA | FOS | 2353 |
| CAAATGTGACTTCACAGTAC | PIAS1 | 8554 |
| CAAATTAGTGATCAATCTAG | PCGF6 | 84108 |
| CAAATTCCGAAATCATGTCA | WDR26 | 80232 |
| CAACAAGCACACGCCCATGA | NDUFA1 | 4694 |
| CAACAATGGCAACATCAGGT | NOS2 | 4843 |
| CAACACCCAACACTTATAAG | EZH2 | 2146 |
| CAACACCCCGCGTTATGCTA | NO_CURRENT_311 | NO_CURRENT_311 |
| CAACACTCCCATCATCCTAG | RAC1 | 5879 |
| CAACCACCCACAAGTCATTG | CXCR4 | 7852 |
| CAACCGGTATGACTTCATCA | IRGM | 345611 |
| CAACGACCTCAAGTACGATG | SBNO2 | 22904 |
| CAACGACGGGCCTAGTCTCA | NO_CURRENT_312 | NO_CURRENT_312 |
| CAACTGGTAGTCCATAGTGA | CTNNB1 | 1499 |
| CAACTTCTCACAGTCGATGT | CASP9 | 842 |
| CAAGAAGACAGTGGTGCCCA | NDUFC1 | 4717 |
| CAAGACCTGCTACCACATCG | SCN5A | 6331 |
| CAAGACCTTATCGTGCAGCG | NO_CURRENT_313 | NO_CURRENT_313 |
| CAAGAGATTCTATGCCAGGT | CCNC | 892 |

| | | |
|---|---|---|
| CAAGATAATACTGCACCGAT | AHR | 196 |
| CAAGATCCATGTCCAATGTG | IKBKB | 3551 |
| CAAGATCGGGTCATGCTCAC | YPEL5 | 51646 |
| CAAGATCTACCACCCCAACG | UBE2L6 | 9246 |
| CAAGATGCACCTGTATCTCG | ACACA | 31 |
| CAAGATGGAAACCAAGAGGT | HTR1D | 3352 |
| CAAGCCGTCGGAGCGACCGT | MMP9 | 4318 |
| CAAGCTAACCATCTTACGCA | ARNT | 405 |
| CAAGCTGCCTAAATTGAAAA | ANP32B | 10541 |
| CAAGCTGGTTAATGCAAATG | CAT | 847 |
| CAAGGAATGACATCTCGGTC | CASP3 | 836 |
| CAAGGACCGGAACGAGAATG | RXRA | 6256 |
| CAAGGAGACCTCGCTCAACA | NFRKB | 4798 |
| CAAGGAGATGGACCTCAGCG | NFKB1 | 4790 |
| CAAGGATGGAAATTAAAACG | NO_CURRENT_314 | NO_CURRENT_314 |
| CAAGGCGAGTAATACCTGTC | MAPK14 | 1432 |
| CAAGGTCAACTATTCATACG | CACNA2D2 | 9254 |
| CAAGGTCAATTACACCAGGT | KAT2A | 2648 |
| CAAGTACAGTTGATACCGTG | CUBN | 8029 |
| CAAGTACCGTGACTCGAAGC | NR1H3 | 10062 |
| CAAGTGAGCATAAGCGATGT | NO_CURRENT_315 | NO_CURRENT_315 |
| CAATAATACAGTTTATTCAG | NO_CURRENT_316 | NO_CURRENT_316 |
| CAATAGTGATGCCCCACAGA | HCAR2 | 338442 |
| CAATATCGGGTGCTACAGGA | NO_CURRENT_317 | NO_CURRENT_317 |
| CAATCCATTCGCTTTCACCG | NOD2 | 64127 |
| CAATCGGCGACGTTTTAAAT | NO_CURRENT_318 | NO_CURRENT_318 |
| CAATCTCAATATAACTGTGT | NO_CURRENT_319 | NO_CURRENT_319 |
| CAATCTTCTCGACCGACACA | CASP9 | 842 |
| CAATGAGCTCAACTGGACAG | CACNA2D2 | 9254 |
| CAATGATGTACCACTGCCCA | JAM3 | 83700 |
| CAATGCTGCTACAAAATCTG | CASP1 | 834 |
| CAATGGAATTAACACAATAG | TLR2 | 7097 |
| CAATGGCAAGAACTATCCCA | BAHD1 | 22893 |
| CAATGTGGGAGAGACCGTCA | VIPR2 | 7434 |
| CAATTATCAGTTTCAGCATC | UQCRB | 7381 |
| CAATTGATAGAGCAATTTCC | MCTS1 | 28985 |
| CAATTGTTAAGTGTAGACGC | INO80 | 54617 |
| CAATTTCAGCATGACTAGAG | ALOX15 | 246 |
| CAATTTGCTCCTGACCATGA | OR7C2 | 26658 |
| CACAAAAACGGGGTTACGTG | MAPK14 | 1432 |
| CACAAACTCAGATCCCAAGT | OXSM | 54995 |
| CACAACATCTACCACCACTG | APEH | 327 |
| CACAACTACAGGTTGCGTAA | GBP5 | 115362 |
| CACAACTGTCTGATCCAGGT | DPY30 | 84661 |
| CACAAGCAGATTTATCACAA | NO_CURRENT_320 | NO_CURRENT_320 |
| CACAAGCGTTCCACGAGCGA | GSDMD | 79792 |
| CACAAGTAACAGCAATTCCG | AHCTF1 | 25909 |
| CACACAGTGTCCAGATACGA | KPNB1 | 3837 |
| CACACATGACCAATAACAAG | MAP3K7 | 6885 |
| CACACCACAAAGCTGTTGCC | LTB4R | 1241 |

| | | |
|---|---|---|
| CACACCATAGAGGTGTGACA | NO_CURRENT_321 | NO_CURRENT_321 |
| CACACCCCGGCTGTTCAGTG | HELZ2 | 85441 |
| CACACCTCTGACATTCACCA | MMP1 | 4312 |
| CACACTTACTCTCCACGTCG | JAK1 | 3716 |
| CACAGACCACCCAATCAGAA | NO_CURRENT_322 | NO_CURRENT_322 |
| CACAGCAAAACGCAGCAACT | SUPT7L | 9913 |
| CACAGCCCGCACAGGCATCG | NOD1 | 10392 |
| CACAGGTTTAGCGTCCTCTG | SUPT7L | 9913 |
| CACAGTCATTCTGACAGATG | VDR | 7421 |
| CACAGTCTTGCACAAATAGG | FAM214B | 80256 |
| CACAGTGACTGTGTTGTCAT | MBNL1 | 4154 |
| CACAGTGCACCATGTTCCGG | PDE4A | 5141 |
| CACAGTGTCTGGTACACGCA | TLR1 | 7096 |
| CACATACATCAACCAACCCT | NO_CURRENT_323 | NO_CURRENT_323 |
| CACATAGTTCTCAAACACTG | RHOA | 387 |
| CACATCGCAGAAGATGTCAA | CYP4B1 | 1580 |
| CACATGGGGTACAGCACACC | NO_CURRENT_324 | NO_CURRENT_324 |
| CACATGTGTACATCAAATCA | NO_CURRENT_325 | NO_CURRENT_325 |
| CACATTCACACCCACCTGTG | PTGIS | 5740 |
| CACCAATGTTCCTCCGCAAA | ATP5J | 522 |
| CACCACCGGAGGAATCACAA | EP400 | 57634 |
| CACCACGGTCTGTGCGCACG | CEBPD | 1052 |
| CACCAGAAGTAATTTCAGGA | PRKAA1 | 5562 |
| CACCAGTGAGCAGCAACACG | SLC7A3 | 84889 |
| CACCATCACAGACAAACGAT | LRP2 | 4036 |
| CACCATGGGAGGAACTACTG | RBM42 | 79171 |
| CACCATTGAGAGGATCTATG | UBE2H | 7328 |
| CACCATTGCAATCAACACCA | ALOX5 | 240 |
| CACCCAGACGAATTGTACGT | CYBB | 1536 |
| CACCCCAACAGAACAATAGG | HSPA13 | 6782 |
| CACCCCAACGTGGACGAGAA | UBE2L6 | 9246 |
| CACCCGGAACCACTACTGAG | NO_CURRENT_326 | NO_CURRENT_326 |
| CACCCGGAGTATGGCCCCAG | TNFRSF1B | 7133 |
| CACCCTTATATTCAGTAACT | NO_CURRENT_327 | NO_CURRENT_327 |
| CACCGACAAGTACTCCAACA | CYFIP1 | 23191 |
| CACCGCAAACACAGCACGAA | MSL2 | 55167 |
| CACCGGACTCAAATAATGAC | NO_CURRENT_328 | NO_CURRENT_328 |
| CACCGTGAGCAGATACTCCA | RAD23A | 5886 |
| CACCGTGTGTGACTCCTGTG | TNFRSF1B | 7133 |
| CACCTCTGCGGCGAATCAGA | CCL20 | 6364 |
| CACCTGGCAGTCGCCCTACG | RTP1 | 132112 |
| CACCTTAAAGTCCTCCAGCA | IL10 | 3586 |
| CACCTTCGACACCATGTCGC | TP73 | 7161 |
| CACGAACTCACACCGCGCGA | NO_CURRENT_329 | NO_CURRENT_329 |
| CACGACATTCAATTGCCATG | PPARG | 5468 |
| CACGCACAATCCTTCACGCA | NO_CURRENT_330 | NO_CURRENT_330 |
| CACGCACAGCACGTCCACTG | ADRB1 | 153 |
| CACGCCAACTAAAACTGCAG | NO_CURRENT_331 | NO_CURRENT_331 |
| CACGGACATGAACTTCAGAG | HNRNPF | 3185 |
| CACGGACGTATCATCCACCA | BAP1 | 8314 |

| | | |
|---|---|---|
| CACGGAGCTACCGCGACTGG | TLR9 | 54106 |
| CACGTCACCTGCACAGATGG | PARK7 | 11315 |
| CACGTCACCTTGGAGCCGAA | TRMT61A | 115708 |
| CACTAAATGAGGCTATCAGG | ABL2 | 27 |
| CACTACATCTGCCACAGTGG | NR1H3 | 10062 |
| CACTACTTCCAGAACACACA | ARF1 | 375 |
| CACTCAGAACTTACATCAGA | BCL2L11 | 10018 |
| CACTCGATGAGACCACGCTC | TLR9 | 54106 |
| CACTCGTACTCACGCTTCTG | CCAR2 | 57805 |
| CACTCTCCGTGTTCTCACTA | RTP1 | 132112 |
| CACTCTGTGCAATACGCTCC | GNAI2 | 2771 |
| CACTGAGGGGAACCCTCGCT | MMP1 | 4312 |
| CACTGCAGTATTCGTGGCCT | NO_CURRENT_332 | NO_CURRENT_332 |
| CACTGGGCTCTCACCTGTTG | HAMP | 57817 |
| CACTTTCAATGAATCAAAGC | MBNL1 | 4154 |
| CAGAAAACATCCCAGAAAAG | RHOA | 387 |
| CAGAAACACCCACCTACCCC | TNFRSF6B | 8771 |
| CAGAACAGCATGACAAACAG | CXCR1 | 3577 |
| CAGAACCCATGAGTCGCCCT | ACADS | 35 |
| CAGAACCGTGTTCCGGCTCG | RAD23A | 5886 |
| CAGAAGCTATTCGGGCTGTG | CYP8B1 | 1582 |
| CAGAATTCAGAAGGAGCTGG | UBE2E1 | 7324 |
| CAGACAAGGCTTGGCAACCC | IL10 | 3586 |
| CAGACACAGACCTTGTATCC | PQBP1 | 10084 |
| CAGACCAAGTGATCCAACTG | CYP4A11 | 1579 |
| CAGACCAAGTGATCCAACTG | CYP4A22 | 284541 |
| CAGACCCAGTAAAACCACCA | NO_CURRENT_333 | NO_CURRENT_333 |
| CAGACGCGATCTTCGTCCAG | SERPINE1 | 5054 |
| CAGACGGTTGGTAAGGACGC | NO_CURRENT_334 | NO_CURRENT_334 |
| CAGAGAAAACTATTGGGTCA | KIF2B | 84643 |
| CAGAGACAAATCGATATAGG | INO80 | 54617 |
| CAGAGATCCCAGTAAACCAG | KPNB1 | 3837 |
| CAGAGATCCGTCCACAAAAG | NR1H3 | 10062 |
| CAGAGCCTTGCGCAATTTTG | NO_CURRENT_335 | NO_CURRENT_335 |
| CAGAGCGGCGGAACCACGCG | SPHK1 | 8877 |
| CAGAGGGAAGCTACCCTGCA | AGER | 177 |
| CAGAGGTAAAAAAACGACGG | AHDC1 | 27245 |
| CAGAGGTGGCCTCCATCGTG | C6orf15 | 29113 |
| CAGAGTTGGAGGGTTGAAAG | NO_CURRENT_336 | NO_CURRENT_336 |
| CAGATCGGACACTCAAAGTG | FBXW7 | 55294 |
| CAGATGACGGTTTACCATCC | TIFA | 92610 |
| CAGATGCGATGTTCTTGCGG | KIF2B | 84643 |
| CAGATGCTTGTACGCGGAGA | IFNAR1 | 3454 |
| CAGATGGATTGGAAGTGCCC | CDKN2D | 1032 |
| CAGATTCCACTCCTTCAAGA | GPT2 | 84706 |
| CAGCAAAGAACTTTCACATG | PRR14L | 253143 |
| CAGCAAGCTCCATGACTGAG | MMP13 | 4322 |
| CAGCACTTAGATATAAACTC | NO_CURRENT_337 | NO_CURRENT_337 |
| CAGCAGAAAGCCCGTCATCT | SERF2 | 10169 |
| CAGCAGATCACCGGCCACGG | NDUFA12 | 55967 |

| | | |
|---|---|---|
| CAGCAGATTCAAGCAGCTAT | SERPINE1 | 5054 |
| CAGCAGCCATTAAATTACCC | PAXIP1 | 22976 |
| CAGCAGGAATCTATCAGAGA | AIM2 | 9447 |
| CAGCATCATCTTCCTCACGG | HCAR2 | 338442 |
| CAGCATTCGATATAAACTCA | OXSM | 54995 |
| CAGCCAGAGCCACCCAATCG | RPS6KA4 | 8986 |
| CAGCCATAGACAGCAACCGA | ARID3A | 1820 |
| CAGCCATTGGGCCCATACGT | IKBKB | 3551 |
| CAGCCCAGTAGGCAATAGCA | NO_CURRENT_338 | NO_CURRENT_338 |
| CAGCCCAGTTTACTGTCGTG | BTN2A2 | 10385 |
| CAGCCCGGCGAGTTCACGCT | LCN2 | 3934 |
| CAGCCGAACCAAGTACTGGG | CS | 1431 |
| CAGCCGGAGGCATTGTCCGG | HTR7 | 3363 |
| CAGCCTATTTTGCTACCTAC | NO_CURRENT_339 | NO_CURRENT_339 |
| CAGCCTCTGAGAGCACACAA | NAIP | 4671 |
| CAGCCTGAATGTCAATAACG | WDR26 | 80232 |
| CAGCGAGTACACATTCATTG | GSDMD | 79792 |
| CAGCGCCATAGAAGCGGGCC | AKT1 | 207 |
| CAGCGCTGACATAGGGACCT | NUFIP2 | 57532 |
| CAGCGGTCAGTGAATATCCG | DET1 | 55070 |
| CAGCTCGTTAAGGATCAACA | MTOR | 2475 |
| CAGCTTCGGGATCCTAATGT | RIPK3 | 11035 |
| CAGCTTGAATCAGAGTAGAG | RAD21 | 5885 |
| CAGGAAGAGCTCGGGATGAG | BTN2A2 | 10385 |
| CAGGAAGCATGGGTTAGATG | NLRP1 | 22861 |
| CAGGAAGCGGCGGCCATGTG | CYP4B1 | 1580 |
| CAGGACAGGATGACAATACC | CXCR4 | 7852 |
| CAGGACGGTACTTACGTCTG | CREBBP | 1387 |
| CAGGAGAAAATATCCACCCA | NO_CURRENT_340 | NO_CURRENT_340 |
| CAGGAGAGTCAACACCAGAG | NO_CURRENT_341 | NO_CURRENT_341 |
| CAGGATACTGAAAGTTCGCA | NQO1 | 1728 |
| CAGGATGACCCTCTGACCAA | STAT2 | 6773 |
| CAGGATGCCAATCCATGGAA | CYP2R1 | 120227 |
| CAGGATGGACGTGAACGCGT | NO_CURRENT_342 | NO_CURRENT_342 |
| CAGGATGGTAAACCGTCATC | TIFA | 92610 |
| CAGGATGTTTCGCATTAAGA | UQCRB | 7381 |
| CAGGCACGGTACTGCTCAAG | INO80B | 83444 |
| CAGGCCCAATAGGAAACGTG | NLRP1 | 22861 |
| CAGGCCCGTGAACTTAGCCG | SLC3A2 | 6520 |
| CAGGCGGCCCTCATACACGT | TYK2 | 7297 |
| CAGGCGTCAACATCCGACGG | CYP4F11 | 57834 |
| CAGGCGTTCCAGCTCCGAAG | JUNB | 3726 |
| CAGGCTCAGCAGGAACACTA | CXCR1 | 3577 |
| CAGGCTCTCAAATCCAACAG | DLAT | 1737 |
| CAGGCTCTCCGACCTGTACG | EBI3 | 10148 |
| CAGGCTGTAAAAGCAAAGGC | PIGY | 84992 |
| CAGGCTTATTCGATTAAGTG | RBM6 | 10180 |
| CAGGCTTGATACATACTCCA | RBM6 | 10180 |
| CAGGGAGAGGAGGTGACCCA | SUPT7L | 9913 |
| CAGGGCATGGAAGTCGAGGA | PTGES2 | 80142 |

| | | |
|---|---|---|
| CAGGGCTTTGACAAAGCCGT | JUNB | 3726 |
| CAGGGTCAGTCACAGCCATG | HSH2D | 84941 |
| CAGGGTGACGATGTCAACTG | IL2RB | 3560 |
| CAGGGTTCATGCTACTGAGT | MTOR | 2475 |
| CAGGTATAAGAAACCCATGA | HTR3E | 285242 |
| CAGGTCACTCCCACGGAAGA | CYP4F11 | 57834 |
| CAGGTCGCCCGCCGGACAGG | NMU | 10874 |
| CAGGTGGCACAAACCGTGCA | NOD2 | 64127 |
| CAGGTGGGCAAACGCAGAGG | TRIR | 79002 |
| CAGGTTCACTAACGGGGGCA | NDUFA1 | 4694 |
| CAGGTTTGCACGCATAGCTA | NO_CURRENT_343 | NO_CURRENT_343 |
| CAGTAAAGTCAGAAATGTGA | STAT1 | 6772 |
| CAGTACATTGTAGAAGATGG | ACACA | 31 |
| CAGTAGCACGATCACGGTCA | BAHD1 | 22893 |
| CAGTATGGAGAGGTTTACGT | ABL2 | 27 |
| CAGTCACCCAGCGGCCCACA | TRMT61A | 115708 |
| CAGTCAGGTTTGGCATTCGG | NDUFC1 | 4717 |
| CAGTCATCCAGATTTCCCTG | PPID | 5481 |
| CAGTGCAATTGGTTAAAAGC | PDK4 | 5166 |
| CAGTGCATCCTCATACACGT | CACNA2D2 | 9254 |
| CAGTGCTACGGCGAGCGTGG | RTP1 | 132112 |
| CAGTGCTTGCAGACACTGCA | CXCL3 | 2921 |
| CAGTGGTGTTGATGATGACA | CASP3 | 836 |
| CAGTGGTTGAGGGAAATCGT | PLD1 | 5337 |
| CAGTGTCCGCATCGTCATTG | NOD2 | 64127 |
| CAGTGTTTGTGTATAAGGGT | BTN2A2 | 10385 |
| CAGTTAACATCACTATGGCA | IRGM | 345611 |
| CAGTTGCAGGATTCCAGCTA | NO_CURRENT_344 | NO_CURRENT_344 |
| CAGTTTCAGCATCAGGCAAG | UQCRB | 7381 |
| CATAAAAGTCTGTCTGAGAG | ZNF641 | 121274 |
| CATAAGGTTAACTGCGTGGA | NO_CURRENT_345 | NO_CURRENT_345 |
| CATAATGATAAAGCACCGGC | DNTTIP1 | 116092 |
| CATACACGATGTGAATTGGT | PDK4 | 5166 |
| CATACCAAACTATCTGGATC | C4orf17 | 84103 |
| CATACCCACATCAAACACTC | NO_CURRENT_346 | NO_CURRENT_346 |
| CATACCGAGTTTACACGGAA | SERPINB2 | 5055 |
| CATACTACTAGATATTACAC | MAP2K5 | 5607 |
| CATACTCACCATATATGGCT | MMP1 | 4312 |
| CATAGAGACCAACATCCGTG | POU2F1 | 5451 |
| CATAGATGTCCGAAACAGCA | NCKAP1L | 3071 |
| CATAGCCAGCCAATATAGCA | ACOD1 | 730249 |
| CATATAAATTCAGTAGTCCA | NO_CURRENT_347 | NO_CURRENT_347 |
| CATATAAGTGCTTACCAATG | NO_CURRENT_348 | NO_CURRENT_348 |
| CATATCAAAGATGTCCATGG | DNAJA1 | 3301 |
| CATATCATTGACCTTCACAC | DET1 | 55070 |
| CATATCCTTCCCAATAAACA | TMEM70 | 54968 |
| CATATTACATCGGATATCTG | TMEM173 | 340061 |
| CATCAAAAATCCTCTCCATG | PTGIS | 5740 |
| CATCAAATTAAGGTAATATG | NO_CURRENT_349 | NO_CURRENT_349 |
| CATCACAGCAGCTTTAGAGT | APIP | 51074 |

| | | |
|---|---|---|
| CATCACATCAGCATCACCAT | OXSM | 54995 |
| CATCACGGCTACACTGTACT | PARK7 | 11315 |
| CATCACTTGTGCAATCCAGA | NLRC4 | 58484 |
| CATCAGGTGTATTCTAAGTG | NO_CURRENT_350 | NO_CURRENT_350 |
| CATCAGTAGACCAGAGACCC | RND3 | 390 |
| CATCATAGGAGTATCTGTGA | AGER | 177 |
| CATCATATATTGACTAAGGT | NO_CURRENT_351 | NO_CURRENT_351 |
| CATCATCAATTTCGAGCAGA | SOD1 | 6647 |
| CATCATTCAGGAGATGCACG | CTNNBL1 | 56259 |
| CATCCCCAGTCGCTTCCAGT | IZUMO2 | 126123 |
| CATCCCTAGAGAGTACTGCA | CACNA2D2 | 9254 |
| CATCCCTATACAGGCTATAG | NO_CURRENT_352 | NO_CURRENT_352 |
| CATCCCTGGGAATGCAAGCA | TNFRSF1B | 7133 |
| CATCCGAGAAGTGCTGATCA | NLRC4 | 58484 |
| CATCCTCTTTAGATGTATGG | CS | 1431 |
| CATCCTGGCCAGTTGCAGTG | TIFA | 92610 |
| CATCGAGAACCCCAACCGGG | PDAP1 | 11333 |
| CATCGCAACCCATTTCGTGG | KIF2B | 84643 |
| CATCGCCTCTCAACACAGCA | LGMN | 5641 |
| CATCGTTGGACATACCCTGT | CUBN | 8029 |
| CATCTCATGTGCCAAGAAAG | IRAK4 | 51135 |
| CATCTCGCGGCACTTCTACG | HELZ2 | 85441 |
| CATCTGCCAAGGATCCTCAG | C6orf15 | 29113 |
| CATCTGTAGGGTTGCAAGCC | NO_CURRENT_353 | NO_CURRENT_353 |
| CATCTTCCGGAAGACAATCG | UBA7 | 7318 |
| CATGAAAAAGCCAAGATGCG | MMP13 | 4322 |
| CATGAACATGGAATCCATGC | SOD1 | 6647 |
| CATGAACCGAATCCTAGATG | EIF4G3 | 8672 |
| CATGAAGTCAGTGACTGACC | NDUFS8 | 4728 |
| CATGAATCGTGCAGTCAGAG | HNRNPF | 3185 |
| CATGACCCGGAATGCAGTAT | KPNA1 | 3836 |
| CATGAGCACAAAATTTAGGG | NO_CURRENT_354 | NO_CURRENT_354 |
| CATGATCGCCTTGACTAGCG | PDE3A | 5139 |
| CATGCAAGACTTCACAGTTG | PHF8 | 23133 |
| CATGCCTGTTTACAAGAAAG | TP73 | 7161 |
| CATGGCATAAGTATAAGACA | NO_CURRENT_355 | NO_CURRENT_355 |
| CATGGCCTACGGTGTCTTTG | NO_CURRENT_356 | NO_CURRENT_356 |
| CATGGCGGGACAGGAGGATC | BRK1 | 55845 |
| CATGGGACACCGGTACATTG | HNRNPF | 3185 |
| CATGGGAGATAATAGACAAG | NO_CURRENT_357 | NO_CURRENT_357 |
| CATGGGATTAGTGGCTAACG | NO_CURRENT_358 | NO_CURRENT_358 |
| CATGGGCCCCCACTCCATGT | RXRA | 6256 |
| CATGTCGCGCAGCACGTTAG | PYCARD | 29108 |
| CATGTCTCATGGCATCCTAG | CASP5 | 838 |
| CATGTGAGGATCAAAAACTG | NOS2 | 4843 |
| CATGTGCAAGAAGTATGGGG | SETD1B | 23067 |
| CATGTTGGATAAGACCTCCA | FBXO38 | 81545 |
| CATGTTGTACCAAAGGATGG | STAT1 | 6772 |
| CATGTTTACGACGTTTGGAA | ZRSR2 | 8233 |
| CATTACAACAACCTGCTACG | TMEM173 | 340061 |

| | | |
|---|---|---|
| CATTACTTACATATGCTGTG | HTR3E | 285242 |
| CATTCCACAACTGCCGACCT | RRAGC | 64121 |
| CATTCCCAGCATCCCAAAGG | MLF2 | 8079 |
| CATTCGTTGAAAAAGGACTG | NO_CURRENT_359 | NO_CURRENT_359 |
| CATTCTCACCAAGTAGCTCA | JAK1 | 3716 |
| CATTCTCTGGCTTTAAGTCA | CHEK2 | 11200 |
| CATTCTGGGCAAGATGACAG | MAP2K7 | 5609 |
| CATTCTGGGTGACACGCTGG | PRKAA1 | 5562 |
| CATTGAATATGATTGCAAGG | SATB1 | 6304 |
| CATTGACAATTACTTAACCT | NO_CURRENT_360 | NO_CURRENT_360 |
| CATTGAGGTTCGGTTCTATG | NFKB2 | 4791 |
| CATTGCACGCCACAGCATTG | NO_CURRENT_361 | NO_CURRENT_361 |
| CATTGCTACTAACACACTAG | GPR119 | 139760 |
| CATTGCTGTATACAATCATG | IFNAR2 | 3455 |
| CATTGGAAATTAACTATGGG | NO_CURRENT_362 | NO_CURRENT_362 |
| CATTGGCATCATCGCTGGAC | UBE2L6 | 9246 |
| CATTGTAATATTAACAGTCA | UBE2E1 | 7324 |
| CATTGTATGAACGCAATAGC | NO_CURRENT_363 | NO_CURRENT_363 |
| CATTGTCAAGGTGTTGCCGC | HCAR2 | 338442 |
| CATTGTTGAGCGGGCGCGCT | NO_CURRENT_364 | NO_CURRENT_364 |
| CATTTATTACAGGGACTGGA | TMEM30A | 55754 |
| CCAAAAAGATGAATATCTCG | NO_CURRENT_365 | NO_CURRENT_365 |
| CCAAAAGTCGCCCTCCCGGG | MCL1 | 4170 |
| CCAAAATTCTAACAAGCCCA | GBP7 | 388646 |
| CCAAACATAGGTCAACAAGA | AHCTF1 | 25909 |
| CCAAACCAGGTGGAACTGTG | SPINT1 | 6692 |
| CCAAAGACATCTCGGAGCCG | SMPD1 | 6609 |
| CCAAAGGCATGCTCTACTCC | C4orf17 | 84103 |
| CCAAAGGTAACAACTATGCT | NO_CURRENT_366 | NO_CURRENT_366 |
| CCAACAATCTCAAACTCTGG | IFNAR2 | 3455 |
| CCAACCCAAAGGAGCGTTTG | TRERF1 | 55809 |
| CCAACGTTCCTACCACGGAA | ACOD1 | 730249 |
| CCAACTACGACACCGACGAC | MMP9 | 4318 |
| CCAACTCGAGCACAACCCTA | FBXO38 | 81545 |
| CCAAGACCAAATAGCAGTTG | NMRK2 | 27231 |
| CCAAGACCATCTGACCCAGA | NCKAP1L | 3071 |
| CCAAGATCCCCGTATCGCTG | AHDC1 | 27245 |
| CCAAGCGGAAGTCAACCTCG | ACTR5 | 79913 |
| CCAAGCTCAGTGCTCGACGG | KMT2D | 8085 |
| CCAAGGGGTGCTCACAACTG | KAT2A | 2648 |
| CCAAGTACCCTTATTCCAGA | NOD2 | 64127 |
| CCAAGTATTAAAGTAAACAG | NO_CURRENT_367 | NO_CURRENT_367 |
| CCAAGTCTATCACCAAGACG | HSPA13 | 6782 |
| CCAAGTCTTCAATTACGTGA | NLRP12 | 91662 |
| CCAATGATAAGCCCGAACGG | NO_CURRENT_368 | NO_CURRENT_368 |
| CCAATGGGCGCTTACGAAGT | BAHD1 | 22893 |
| CCAATGTCGTATTTCACACA | NFRKB | 4798 |
| CCAATTAAAGATACTACAAG | APAF1 | 317 |
| CCAATTTCTACCATTATCCG | MSL2 | 55167 |
| CCACACACCCCACAGATCCG | VDR | 7421 |

| | | |
|---|---|---|
| CCACACAGCAAGATGTGCTA | GNAI2 | 2771 |
| CCACACCAGCAGGTTGATAA | MCTS1 | 28985 |
| CCACACCTGTCTAGCATGAC | NO_CURRENT_369 | NO_CURRENT_369 |
| CCACAGATGCAACATAAGCT | IL2RB | 3560 |
| CCACAGATTGAGACAAGCTG | HSPA13 | 6782 |
| CCACATCCTCAGGCTCAGAA | CASP1 | 834 |
| CCACATGCTAAAGAACAACG | CASP5 | 838 |
| CCACCAGACGGAACTCCCGT | SUPT7L | 9913 |
| CCACCATTGAACTTCAGTGC | SOD2 | 6648 |
| CCACCCAAAATGGATTCCTG | AHCTF1 | 25909 |
| CCACCGTCACAGCACTTGTG | SMPD1 | 6609 |
| CCACCTCACTCCTTACACTG | CCAR2 | 57805 |
| CCACGATGCCACCTCATCCC | NO_CURRENT_370 | NO_CURRENT_370 |
| CCACGCAGCTCAGAGGCACG | CYP27B1 | 1594 |
| CCACGGAAGCGAGGGCTCAG | NDUFS8 | 4728 |
| CCACGTACACGTTGTCCCCG | GSDMD | 79792 |
| CCACTACCACTGAGTGCCTG | NLRC4 | 58484 |
| CCACTCAAGATGGACTCGAT | FBXO38 | 81545 |
| CCACTCCATAGAATCTCCGG | NFKB2 | 4791 |
| CCAGAAAGCTGGGCGTAGAG | CYP27B1 | 1594 |
| CCAGAAATCGCTCATCCACA | ALOX15 | 246 |
| CCAGAATTGAGTGCAAAAAG | NO_CURRENT_371 | NO_CURRENT_371 |
| CCAGAGAATACATAACACCT | APAF1 | 317 |
| CCAGAGTCCCACATTCCCTG | ACACA | 31 |
| CCAGATCGAAATGAAAGTGA | CHMP6 | 79643 |
| CCAGCAATCCCCGGCTCCTG | CXCL2 | 2920 |
| CCAGCAATCCCCGGCTCCTG | CXCL3 | 2921 |
| CCAGCAGATTCTCTTGCACA | CYP4F11 | 57834 |
| CCAGCAGGATGGGCACATCA | RHOG | 391 |
| CCAGCAGGCACGAGGCGATG | SETD1B | 23067 |
| CCAGCCTTATTAATCATCAG | ZNF616 | 90317 |
| CCAGCGCCAGTACTTCAAGG | RPS6KA4 | 8986 |
| CCAGCGGGTACACGAAGCAG | SLC25A6 | 293 |
| CCAGCGTGGGTAACAGACAG | BAHD1 | 22893 |
| CCAGGAAGCACCTTCTAGGG | VIPR2 | 7434 |
| CCAGGAATTTGCTACCAACA | NO_CURRENT_372 | NO_CURRENT_372 |
| CCAGGACAGAGCTGGAGCCA | HAMP | 57817 |
| CCAGGACTACGAACAGATGG | HELZ2 | 85441 |
| CCAGGAGACACCCTGAGACA | AGER | 177 |
| CCAGGAGGATGTTCCCCGAG | NOD1 | 10392 |
| CCAGGATCCGTGTATGAGGG | UBE2E1 | 7324 |
| CCAGGCTGAAGTTCGTACCT | NO_CURRENT_373 | NO_CURRENT_373 |
| CCAGGGGCTCACCAATTCCA | CSNK1D | 1453 |
| CCAGGTACTGCTTCCAACCA | RTP1 | 132112 |
| CCAGGTCGCAAAGCATGCTC | LAMTOR3 | 8649 |
| CCAGGTGACTCCGATCACCA | RBM42 | 79171 |
| CCAGTACTTATTGCCGAACT | PTGES2 | 80142 |
| CCAGTATGTGCCAATCACAA | LARP4B | 23185 |
| CCAGTCACACATGAGAATGT | TRAF6 | 7189 |
| CCAGTCCAATAACCCCTGAG | TNFRSF1A | 7132 |

| | | |
|---|---|---|
| CCAGTCGACACCCATCATTG | NR2C2 | 7182 |
| CCAGTGCCCTTTTGTCGCAA | NO_CURRENT_374 | NO_CURRENT_374 |
| CCAGTGCTTCCTCCACGGTA | CS | 1431 |
| CCAGTGGACAGACCGCATCA | PDE4A | 5141 |
| CCAGTTATAATTAGGGGTTT | NO_CURRENT_375 | NO_CURRENT_375 |
| CCAGTTCGGTCTTTCAAATC | CDKN2C | 1031 |
| CCAGTTGCTCTGGGGGAACA | NO_CURRENT_376 | NO_CURRENT_376 |
| CCATAAAGATCATCAGCAAA | CHEK2 | 11200 |
| CCATACTGAAAGCCTACGAT | IDH2 | 3418 |
| CCATATCGGGGCGAGACATG | NO_CURRENT_377 | NO_CURRENT_377 |
| CCATATTGATCAGCTTCCAA | GLMN | 11146 |
| CCATCACCGATCGTGAGCCT | NO_CURRENT_378 | NO_CURRENT_378 |
| CCATCATTCACACGAACTGG | VDR | 7421 |
| CCATCGCCGCGGAAGGCCTG | ARID3A | 1820 |
| CCATCTCATCACAGCCTACC | PDHA1 | 5160 |
| CCATCTCGCTGAGATACGAA | ABL1 | 25 |
| CCATGACAAAACCATAGCAA | NO_CURRENT_379 | NO_CURRENT_379 |
| CCATGATCAAGGGGACTGGG | NO_CURRENT_380 | NO_CURRENT_380 |
| CCATGGACTTTGCCACAATG | BRD1 | 23774 |
| CCATGGGCTATGAGCGAGAG | RAD23A | 5886 |
| CCATTCCGTAAGGGCTTGGA | NO_CURRENT_381 | NO_CURRENT_381 |
| CCATTGCAGATAAACTCACG | LRP2 | 4036 |
| CCCAAGGCCGCCACCATTGG | ACO2 | 50 |
| CCCAATGGCTTCTGCGTGAC | NO_CURRENT_382 | NO_CURRENT_382 |
| CCCACACAGTAACACATCAG | SCN5A | 6331 |
| CCCACACATGCGAGTGACAG | TIRAP | 114609 |
| CCCACAGGCCCTGTGCACCA | BAX | 581 |
| CCCACATGACCCCAACTCCG | PQBP1 | 10084 |
| CCCACATTTAAAAACGCAGT | MBNL1 | 4154 |
| CCCACCTCCCGCCCTCACAG | INO80B | 83444 |
| CCCAGCTGGGACTCTAAGAG | NO_CURRENT_383 | NO_CURRENT_383 |
| CCCAGCTTTCCGAATCATGT | MAP3K7 | 6885 |
| CCCAGGATGCTTGACTCACG | IL2RB | 3560 |
| CCCAGTGAAGTAAAACCCAG | NAIP | 4671 |
| CCCATCAGCAAAAGGTGTTG | NDUFB9 | 4715 |
| CCCATCGGGATGGAACGCTG | SLC7A3 | 84889 |
| CCCATCTACTTGGCAGTGAA | NENF | 29937 |
| CCCATGCGACGACCTCCCCA | KMT2C | 58508 |
| CCCATTCCCGACATCCACTG | APEH | 327 |
| CCCCAACTTTCGCGACTCCG | NO_CURRENT_384 | NO_CURRENT_384 |
| CCCCACACGCCAAGTCGGAG | AHDC1 | 27245 |
| CCCCACCATCAGATGGTGCG | CSNK1D | 1453 |
| CCCCAGTAAACGCTCCACAG | NR2C2 | 7182 |
| CCCCAGTCAAAGCTTCGAGG | NELFE | 7936 |
| CCCCATGAACAACGCCACCG | SLC11A1 | 6556 |
| CCCCATGGATACTCACGGGT | UBE2L6 | 9246 |
| CCCCCACTAGTATCCCCATG | RNF25 | 64320 |
| CCCCCTACAATGCCTTGACG | NENF | 29937 |
| CCCCCTGTGAGCTCGCCAAG | SETD1B | 23067 |
| CCCCGCACGTGGATAGACCG | UBA7 | 7318 |

| | | |
|---|---|---|
| CCCCGCCACGTCCCGAACCA | CYP27B1 | 1594 |
| CCCCGGGAAATCTCACAGAG | IRAK1 | 3654 |
| CCCCGTAGCTCATTAGTCTG | NO_CURRENT_385 | NO_CURRENT_385 |
| CCCCGTCAAGGCATTGTAGG | NENF | 29937 |
| CCCCTAAAAGCCCTAACACA | MTOR | 2475 |
| CCCCTATCCTAGTCCTACGG | KMT2D | 8085 |
| CCCCTATGCAGACTACAATT | NO_CURRENT_386 | NO_CURRENT_386 |
| CCCCTCCATGCCCATCAGCA | IZUMO2 | 126123 |
| CCCCTGAACTGTCTACACCC | NO_CURRENT_387 | NO_CURRENT_387 |
| CCCCTGCACCCATACCCGTA | NFKB2 | 4791 |
| CCCCTGGCATTCTCACTCTG | UBA7 | 7318 |
| CCCGACTTGAGCCCCGACGG | HTR7 | 3363 |
| CCCGATGGACTATACCGAAC | NO_CURRENT_388 | NO_CURRENT_388 |
| CCCGCCGAAGACCCTGCTTG | NO_CURRENT_389 | NO_CURRENT_389 |
| CCCGCTTCTCACACTCGTAG | ARID3A | 1820 |
| CCCGGAGCATGATGAGAGCT | ZNF641 | 121274 |
| CCCGGCCGGAGGAACTCACA | BNIP3 | 664 |
| CCCGTGGCGTGCGCACCTGT | NO_CURRENT_390 | NO_CURRENT_390 |
| CCCTCACCTAAACAGGACGG | LCN2 | 3934 |
| CCCTCACGGTCCTTAAGTCT | SPCS1 | 28972 |
| CCCTCATACACGGATCCTGG | UBE2E1 | 7324 |
| CCCTCCATCGCACCCCCTCA | IL2RB | 3560 |
| CCCTCGAGTCTTCTTTGACG | PPID | 5481 |
| CCCTCTACTTCAACGGCCAG | SERPINE1 | 5054 |
| CCCTGACTGCTATGACACCA | GC | 2638 |
| CCCTGCTGGTAGAAAAACTA | NO_CURRENT_391 | NO_CURRENT_391 |
| CCCTGGAATACAGTAAACGC | HTR1D | 3352 |
| CCCTGGGATCACCCCGACCG | NO_CURRENT_392 | NO_CURRENT_392 |
| CCCTTCACTGCCAAGTAGAT | NENF | 29937 |
| CCCTTGTCAGCGTGACCCCG | NO_CURRENT_393 | NO_CURRENT_393 |
| CCGACCGAAGTCCGGGACGG | SLC25A1 | 6576 |
| CCGACTCTTCATTCGTTCTG | PIGL | 9487 |
| CCGAGCACTGGTTCTCCAAG | C16orf72 | 29035 |
| CCGAGGCGGTGTTTAGAGGA | NDUFA8 | 4702 |
| CCGATAAATGGCAGCCCCGG | CYP2R1 | 120227 |
| CCGCAAATGACTGGTCACGC | CREBBP | 1387 |
| CCGCGCATTTCAGAGCACAA | NO_CURRENT_394 | NO_CURRENT_394 |
| CCGCTATGGTTACACTCGGG | MMP9 | 4318 |
| CCGCTATTGAAACCGCCCAC | NO_CURRENT_395 | NO_CURRENT_395 |
| CCGCTCCTGAGACTCCCGCC | SLC25A1 | 6576 |
| CCGCTTCGAGGACGAGGACG | PCGF6 | 84108 |
| CCGGAACAGCAAGAAGAGGC | INO80C | 125476 |
| CCGGAAGCACGGCCTAGACG | NLRP12 | 91662 |
| CCGGAGCACGGTCAAGCAAG | SERPINE1 | 5054 |
| CCGGAGCATCGGTGTTGTGG | C6orf15 | 29113 |
| CCGGATCCCATGGTCAACCA | ACACA | 31 |
| CCGGCCAGCCCTTCCCACAA | INO80C | 125476 |
| CCGGGCTGCGCTTATCGCGA | PYCARD | 29108 |
| CCGGTAAGGCCGAGGACGAG | PAGR1 | 79447 |
| CCGGTTCATGCCATGAATGG | TGFB1 | 7040 |

| | | |
|---|---|---|
| CCGTAGGACCCAACAGACTG | FAM214B | 80256 |
| CCGTCAGGCTGGCATGCGGG | NR1H3 | 10062 |
| CCGTCCTAGAAGGATTCAGC | IFNAR2 | 3455 |
| CCGTCCTGGAAAGAGAAGTG | APIP | 51074 |
| CCGTCCTGGAAAGAGAAGTG | SMG8 | 55181 |
| CCGTCTCCGCATCGTCTTTT | NO_CURRENT_396 | NO_CURRENT_396 |
| CCGTGCCTGACCCCTCAATG | NO_CURRENT_397 | NO_CURRENT_397 |
| CCGTGTTCCACAACTACCCT | NO_CURRENT_398 | NO_CURRENT_398 |
| CCGTTGGACTATGGCGGGTC | NO_CURRENT_399 | NO_CURRENT_399 |
| CCGTTGTACAACCATCCAGC | MPC2 | 25874 |
| CCTAAACTCAGACGCACTAC | NO_CURRENT_400 | NO_CURRENT_400 |
| CCTAAGGGGTACCACCATGG | NO_CURRENT_401 | NO_CURRENT_401 |
| CCTACATGCAGACTAGCAGG | AHCTF1 | 25909 |
| CCTACCATAGCATCACGATA | NO_CURRENT_402 | NO_CURRENT_402 |
| CCTACGCGGTAGGGAACTTT | NO_CURRENT_403 | NO_CURRENT_403 |
| CCTACGTGGAGGCAAACATG | RXRA | 6256 |
| CCTACTCCCGTGTGTTATCC | NO_CURRENT_404 | NO_CURRENT_404 |
| CCTAGAAGGGGTTATCGAGA | RIPK3 | 11035 |
| CCTAGGCTTGCTAAAAGAAG | IZUMO2 | 126123 |
| CCTCAAGAAGGAAGTCATCG | AKT1 | 207 |
| CCTCAAGTGGAACTGCCCGC | TLR9 | 54106 |
| CCTCACCCCTGCACCCGCAG | PCGF6 | 84108 |
| CCTCACCTGATTCTGCCGTG | SQSTM1 | 8878 |
| CCTCACGCCAGACTCCCGGA | MCL1 | 4170 |
| CCTCACTCCACTCACCAACA | COX16 | 51241 |
| CCTCCAAATAGCATCACATG | NO_CURRENT_405 | NO_CURRENT_405 |
| CCTCCACCAAAACCATCCGG | CYBB | 1536 |
| CCTCCGCAAATGGACTGAGA | ATP5J | 522 |
| CCTCCGTGCTAACGCGGACG | NO_CURRENT_406 | NO_CURRENT_406 |
| CCTCCTCCAAGCATCAGTAA | HDAC2 | 3066 |
| CCTCCTGATAGGTCCCGAAG | INO80B | 83444 |
| CCTCGATGGTCACCTGTAGC | NO_CURRENT_407 | NO_CURRENT_407 |
| CCTCGCCCCAGATACCCAAG | SBNO2 | 22904 |
| CCTCGGGCGTAAATACTCAT | NO_CURRENT_408 | NO_CURRENT_408 |
| CCTCGTAAGTCCAGTCGCCG | ARID3A | 1820 |
| CCTCTATCCAGAAAACACGG | SOD1 | 6647 |
| CCTCTCCAGCTCATTTATGG | TRAF6 | 7189 |
| CCTCTGCGTCGAGGGAACCG | KIAA1211L | 343990 |
| CCTCTGGAAGGACACTTCTG | NO_CURRENT_409 | NO_CURRENT_409 |
| CCTCTTACCACAATAGCATG | WDR26 | 80232 |
| CCTGAACCTGTGGGACACTG | RHOG | 391 |
| CCTGAGGAACGCCGACACAG | NOD2 | 64127 |
| CCTGAGGATACCAAAACGTG | NO_CURRENT_410 | NO_CURRENT_410 |
| CCTGATAAATGTGACACCAT | NO_CURRENT_411 | NO_CURRENT_411 |
| CCTGATAAGAACCCAAATGA | DNAJA1 | 3301 |
| CCTGATTAATGATGAACTAG | STAT1 | 6772 |
| CCTGCAAACTCATACGCCTG | NLRP1 | 22861 |
| CCTGCACTCGGAGAAGAACG | AKT1 | 207 |
| CCTGCAGCCCAGCATCAACG | DNTTIP1 | 116092 |
| CCTGCAGTGTCTGCAAGCAC | CXCL3 | 2921 |

| | | |
|---|---|---|
| CCTGCATCGCTTCTACTGTG | RBM42 | 79171 |
| CCTGCGGTGCACGGCTAGCC | NO_CURRENT_412 | NO_CURRENT_412 |
| CCTGCTACCCCCCATATGTG | NO_CURRENT_413 | NO_CURRENT_413 |
| CCTGCTCCTACAGCACCATG | BAK1 | 578 |
| CCTGCTGGATGGTTGTACAA | MPC2 | 25874 |
| CCTGCTGGGAGGCTACACAG | SPHK1 | 8877 |
| CCTGGATAACATCACTGATG | AIM2 | 9447 |
| CCTGGATCCCGACAGCCGGA | AGER | 177 |
| CCTGGATGAGTCTGTGAAAG | CTNNBL1 | 56259 |
| CCTGGCGATACCTCAGCAAC | TGFB1 | 7040 |
| CCTGGCGGGGGCCAGCAGCC | NDUFC1 | 4717 |
| CCTGGCGTGATCAAGACCAT | FOSL2 | 2355 |
| CCTGGCTATTTACTGTGGCT | MAP2K6 | 5608 |
| CCTGGCTCCGGAATTCACTG | PTGS1 | 5742 |
| CCTGGGAGCAAAAGACTCGC | IRAK4 | 51135 |
| CCTGGGGCTACCTCTGCTTG | SERF2 | 10169 |
| CCTGGTCATCAGTCTGGTCA | IDH2 | 3418 |
| CCTGGTGGGAAAACACTGTG | NFKB1 | 4790 |
| CCTGTACCGAGGCCCTGCTG | PLD6 | 201164 |
| CCTGTCCGGCGGGCGACCTG | NMU | 10874 |
| CCTGTGCCTATAGTCTTCAG | PDHB | 5162 |
| CCTGTTTGGACTCCTGAACG | NLRP12 | 91662 |
| CCTTACGAGCTGCAGCCACG | PTGES2 | 80142 |
| CCTTAGCCTCAGAAAGCACT | NO_CURRENT_414 | NO_CURRENT_414 |
| CCTTAGCCTCTAGAGTACCC | NO_CURRENT_415 | NO_CURRENT_415 |
| CCTTATAGACAGGAGAACTA | WDR26 | 80232 |
| CCTTATCCTACCTTAACATG | AIM2 | 9447 |
| CCTTATCTCACGGAATATAG | NO_CURRENT_416 | NO_CURRENT_416 |
| CCTTCACAGAACCTGTCCTG | NLRC4 | 58484 |
| CCTTCATCTGAATACTGACA | MAP3K7 | 6885 |
| CCTTCCGGAGGCTTCGAAGG | FAM214B | 80256 |
| CCTTCCTCTAAACACCGCCT | NDUFA8 | 4702 |
| CCTTCGAAGCGAAAGTACTG | KMT2D | 8085 |
| CCTTCTTGTACCAATAAATG | NO_CURRENT_417 | NO_CURRENT_417 |
| CCTTGAAATCAAATCAAACC | NO_CURRENT_418 | NO_CURRENT_418 |
| CCTTGAAGTAGGGTACAGTA | FOXO4 | 4303 |
| CCTTGAAGTGGGTCACCAAG | EP400 | 57634 |
| CCTTGACAATGAAGTCATCT | HCAR2 | 338442 |
| CCTTGACTTTATAACTATCG | NO_CURRENT_419 | NO_CURRENT_419 |
| CCTTGATCACATTCGCCCTG | IZUMO2 | 126123 |
| CCTTGCAAATGCAATGAATG | ZNF616 | 90317 |
| CCTTGGCTCGACAAAAGCTG | UTY | 7404 |
| CCTTGTCCTGCTCCGTGACG | CHMP6 | 79643 |
| CGAAAACAAGCACACTCCAG | RRAGA | 10670 |
| CGAAAAGAATCTCACAGCCA | CASP5 | 838 |
| CGAAACCCTCTTAAGTTAAC | NO_CURRENT_420 | NO_CURRENT_420 |
| CGAACTTAATCCCGTGGCAA | NO_CURRENT_421 | NO_CURRENT_421 |
| CGAAGACTGAGACTGCTCCA | DYRK1A | 1859 |
| CGAAGTCCGTGAGGACAATG | RPS6KA4 | 8986 |
| CGAAGTTGAGGGCTTGAGTG | SLC25A6 | 293 |

| | | |
|---|---|---|
| CGAATATTATTTCTATCGGG | NO_CURRENT_422 | NO_CURRENT_422 |
| CGAATCGGAACTTTGTACCG | NO_CURRENT_423 | NO_CURRENT_423 |
| CGAATGCCACAAATCCCTGG | SCTR | 6344 |
| CGAATTGAGCAGTGTACACG | CBLL1 | 79872 |
| CGACAACGTGCAGGTGTATC | NO_CURRENT_424 | NO_CURRENT_424 |
| CGACACAAATTACTAACAAC | NO_CURRENT_425 | NO_CURRENT_425 |
| CGACCCGGAGGATGAGATGT | NO_CURRENT_426 | NO_CURRENT_426 |
| CGACCTGACGTGCACCCCAG | KIAA1211L | 343990 |
| CGACGACCAAGATAGTACCT | TBXAS1 | 6916 |
| CGACGGTAATGCACCTACTA | NO_CURRENT_427 | NO_CURRENT_427 |
| CGACTAACCGGAAACTTTTT | NO_CURRENT_428 | NO_CURRENT_428 |
| CGACTCTTCCAACGAGAAGC | SCTR | 6344 |
| CGAGACTTAAGGACCGTGAG | SPCS1 | 28972 |
| CGAGATTACTGTGGAAGATC | VDAC1 | 7416 |
| CGAGCAAAGATTGTTGGATA | NO_CURRENT_429 | NO_CURRENT_429 |
| CGAGCAGATGCTGTCGTCCG | SLC7A5 | 8140 |
| CGAGCTCAGATGCTCCAGCA | SPCS1 | 28972 |
| CGAGGGACCAGCCAAGATCG | NFKB2 | 4791 |
| CGAGGTAACTCGTTAATCCA | LCN2 | 3934 |
| CGAGTGCCAGGACTGTCTGT | SUV39H1 | 6839 |
| CGAGTGGGAAACGGGAATCA | NO_CURRENT_430 | NO_CURRENT_430 |
| CGAGTGTCTCAAGCGCATCG | BAX | 581 |
| CGAGTGTTATACGCACCGTT | NO_CURRENT_431 | NO_CURRENT_431 |
| CGATCGGGTACCGAGAGGCT | EBI3 | 10148 |
| CGATGATCCTGTGGACTACG | PQBP1 | 10084 |
| CGATGCCCGTCTATGGCCCG | NO_CURRENT_432 | NO_CURRENT_432 |
| CGATGCGATAGTCCCTGATA | RAD23A | 5886 |
| CGCAAGGTGTCGGTAACCCT | NO_CURRENT_433 | NO_CURRENT_433 |
| CGCAAGTACATCATCTGTAC | SERPINB2 | 5055 |
| CGCACATCTAAAGTTACTAC | NO_CURRENT_434 | NO_CURRENT_434 |
| CGCACGACCATTGCTGCTGC | NO_CURRENT_435 | NO_CURRENT_435 |
| CGCACGCCCGTTGGCCAGCG | ADRB1 | 153 |
| CGCACGTTCTCATAGGACGG | RHOG | 391 |
| CGCAGACTGGCGCGTCCAGG | RIPK2 | 8767 |
| CGCAGCCATTCCAATACACA | SLC5A8 | 160728 |
| CGCAGGCTAGATGACACCAG | NO_CURRENT_436 | NO_CURRENT_436 |
| CGCAGTACGTATAGACTTAA | NO_CURRENT_437 | NO_CURRENT_437 |
| CGCCACCAGTGCATGAGGAT | ADRB1 | 153 |
| CGCCACGGGCCCGATCCGAG | RPS6KA4 | 8986 |
| CGCCCAACAGCACTGCATGG | PTGIS | 5740 |
| CGCCCGTAGCCCTCGCGCAG | PYCARD | 29108 |
| CGCCGAATGCCAAACCTGAC | NDUFC1 | 4717 |
| CGCCGGCACCTCACGCTCTG | SIRT1 | 23411 |
| CGCCGGCGACCGGCTGAGTG | CDKN2D | 1032 |
| CGCCGGGACCGTTAGGGAAT | NO_CURRENT_438 | NO_CURRENT_438 |
| CGCCGTCCTATGAGAACGTG | RHOG | 391 |
| CGCCTCTCACGTGTAGGCTT | NO_CURRENT_439 | NO_CURRENT_439 |
| CGCCTGGCGAGTAGGCAAAG | ARID3A | 1820 |
| CGCCTGTCCACCCATCAGCA | BAHD1 | 22893 |
| CGCGCACCACGGGCGCGCAC | NO_CURRENT_440 | NO_CURRENT_440 |

| | | |
|---|---|---|
| CGCGGACGACCTCCAACGGG | RBM42 | 79171 |
| CGCGTGTAGCTGGAGACAAG | NO_CURRENT_441 | NO_CURRENT_441 |
| CGCTAAATTGTCACACGTTT | NO_CURRENT_442 | NO_CURRENT_442 |
| CGCTAACACTTGCAACATCG | RND1 | 27289 |
| CGCTAGGTCCGGTAAGTGCG | NO_CURRENT_443 | NO_CURRENT_443 |
| CGCTAGGTTATTTCGTGGCC | NO_CURRENT_444 | NO_CURRENT_444 |
| CGCTAGTACGCTCCTCTATA | NO_CURRENT_445 | NO_CURRENT_445 |
| CGCTCGCACGATTATGACCA | SLC3A2 | 6520 |
| CGCTGTACAAGTCTTCAACG | CYP24A1 | 1591 |
| CGCTTCATAGAGCCAGCGGG | SMPD1 | 6609 |
| CGCTTCCGCGGCCCGTTCAA | NO_CURRENT_446 | NO_CURRENT_446 |
| CGGAACATCCTTCAGACCCA | HTR3E | 285242 |
| CGGAATCCTATGGTTCAACA | NCOA6 | 23054 |
| CGGACACTTTAACCCCGAGG | NFRKB | 4798 |
| CGGACCCAGAAGCTTACCAG | TRERF1 | 55809 |
| CGGACGCCGGAGAAGGCGCG | TMEM70 | 54968 |
| CGGACTCGGTGATCTCGTTG | ITGB2 | 3689 |
| CGGAGACATCTATCTCGGTG | CSNK1D | 1453 |
| CGGAGATCTCGAAGCATGTT | IL10 | 3586 |
| CGGAGCTCTCTGACGAACGT | PHF8 | 23133 |
| CGGAGGTGTCCTGGACATTG | CDKN2D | 1032 |
| CGGATCTCAGAATACGAATG | KHSRP | 8570 |
| CGGATCTTGATGACGTCGAA | SLC25A19 | 60386 |
| CGGATGATGTCCAGCACGCG | MTOR | 2475 |
| CGGCACACCAATGCGTTCGT | NO_CURRENT_447 | NO_CURRENT_447 |
| CGGCAGCACTCACGTAGATG | INO80E | 283899 |
| CGGCCATGGAGAACAGTACC | ZNF641 | 121274 |
| CGGCCGTGGCTCCGATCGAG | SLC25A6 | 293 |
| CGGCCTGAGATGTGTACAGA | CTNNBL1 | 56259 |
| CGGCTATGGAGTCCCCATGT | COX16 | 51241 |
| CGGCTCGCTGTCGTACAGAG | MEX3B | 84206 |
| CGGGAACCTCTGAATACCAT | TBK1 | 29110 |
| CGGGACGTCGCGAAAATGTA | NO_CURRENT_448 | NO_CURRENT_448 |
| CGGGATGCAGCTGGAGAGGA | NO_CURRENT_449 | NO_CURRENT_449 |
| CGGGATGGTCCCTGCCGAGA | NO_CURRENT_450 | NO_CURRENT_450 |
| CGGGCACAACCTCACCGTAG | BTN2A2 | 10385 |
| CGGGCATGACCATAGGCGTG | TRERF1 | 55809 |
| CGGGCCAAGCCTAGCACTGT | PIGL | 9487 |
| CGGGCGCAACATAGGAAGTG | RIPK3 | 11035 |
| CGGGGAATTGCACGGCGGAA | NO_CURRENT_451 | NO_CURRENT_451 |
| CGGGGTGGCATCGCCTACCG | ARMCX2 | 9823 |
| CGGGTAAGAACGGAGCTCGG | BCORL1 | 63035 |
| CGGGTATCCCTTCGACGGGA | MMP9 | 4318 |
| CGGGTCTCCAAAGATCGCTT | NO_CURRENT_452 | NO_CURRENT_452 |
| CGGGTGAGTGGTAGTAAGAG | FOS | 2353 |
| CGGGTGCAGTCCATTTGATG | ABL1 | 25 |
| CGGGTTCAGGTACCGCTTCT | TGFB1 | 7040 |
| CGGTACACAGCCGGTTCCCG | TYK2 | 7297 |
| CGGTACATGCCCAAGAACTG | PIP4K2A | 5305 |
| CGGTAGTATTAATCGCTGAC | NO_CURRENT_453 | NO_CURRENT_453 |

| | | |
|---|---|---|
| CGGTCTCACAGACAACGTTG | DPY30 | 84661 |
| CGGTGAACAATGCTGTGACT | PHC3 | 80012 |
| CGGTGAGCCACACGAAGGAA | NO_CURRENT_454 | NO_CURRENT_454 |
| CGGTGCTGTGAAAGCCGAGC | NO_CURRENT_455 | NO_CURRENT_455 |
| CGGTGGAGTTCCACCAGCAG | TRMT61A | 115708 |
| CGGTGGGAACGACTGTACGT | ARRB1 | 408 |
| CGGTGTCCTCGTGCCGAGCG | KIAA1211L | 343990 |
| CGGTTTACATCTGCCCATCG | NO_CURRENT_456 | NO_CURRENT_456 |
| CGTAATTTTGTAATCGCTTC | NO_CURRENT_457 | NO_CURRENT_457 |
| CGTACCGCGCCGTCCCCGTG | CSNK1D | 1453 |
| CGTAGTAAATATCTAGCTAA | NO_CURRENT_458 | NO_CURRENT_458 |
| CGTATCGTACACGCCGAAGT | SLC25A6 | 293 |
| CGTCAAGTATTAAGCTGCTT | NO_CURRENT_459 | NO_CURRENT_459 |
| CGTCAATGTAGTGGGCACAC | EGFR | 1956 |
| CGTCATATACACAAACGCCC | NO_CURRENT_460 | NO_CURRENT_460 |
| CGTCATTGAAGAACAGTCAG | IFNAR2 | 3455 |
| CGTCCAGAAGAACGGCCCCT | NO_CURRENT_461 | NO_CURRENT_461 |
| CGTCCCTTCGTCTCTGCTTA | NO_CURRENT_462 | NO_CURRENT_462 |
| CGTCGCCATATGCCGGTGGC | NO_CURRENT_463 | NO_CURRENT_463 |
| CGTCGGCGAAAGCGTCCTTG | KHSRP | 8570 |
| CGTCGGGTAGCTATTTCTTT | NO_CURRENT_464 | NO_CURRENT_464 |
| CGTGAACTGCTACAGCGTGA | SLC7A5 | 8140 |
| CGTGCAAAAGCAGAATGGGA | WDR26 | 80232 |
| CGTGCACAAGCAGACCAGTG | IKBKE | 9641 |
| CGTGCCCAGCGGGCAGATCA | MAP2K6 | 5608 |
| CGTGCCTTTACATTCACTTT | NO_CURRENT_465 | NO_CURRENT_465 |
| CGTGCGGTAAATACGAAATA | NO_CURRENT_466 | NO_CURRENT_466 |
| CGTGTAAAAATACCTTTCTA | NO_CURRENT_467 | NO_CURRENT_467 |
| CGTGTGTGGGTAAACGGAAA | NO_CURRENT_468 | NO_CURRENT_468 |
| CGTGTTTGGAATTTGCCGCG | NO_CURRENT_469 | NO_CURRENT_469 |
| CGTTGAAGTAGCTCGGTTAG | C16orf72 | 29035 |
| CGTTGCTCGAAATGACCGCA | JAM3 | 83700 |
| CGTTGGGCATAGCGAACACT | NO_CURRENT_470 | NO_CURRENT_470 |
| CTAAACAGAAAGATCATACA | CASP3 | 836 |
| CTAAATCTGTATCTAACCGG | NO_CURRENT_471 | NO_CURRENT_471 |
| CTAAATGTTATGCAATGCAC | NO_CURRENT_472 | NO_CURRENT_472 |
| CTAAGAAGCGATTGGGCGCG | RPS6KA4 | 8986 |
| CTAAGCCAAGCCAACACTGG | LAMTOR2 | 28956 |
| CTAAGTTTGTTAATGGGCCA | NO_CURRENT_473 | NO_CURRENT_473 |
| CTAATCACGACCTCACCCTA | NO_CURRENT_474 | NO_CURRENT_474 |
| CTAATGCTATCAATCATGAG | NO_CURRENT_475 | NO_CURRENT_475 |
| CTACAATGTTCTGAGCTGCG | NR1H3 | 10062 |
| CTACAGGGGAAACCCGAATG | SETD1B | 23067 |
| CTACATCACCCTTATCCGTG | NCKAP1L | 3071 |
| CTACCATCCTACAAATCTCG | MMP13 | 4322 |
| CTACGTAAATAACACAACTG | IRAK4 | 51135 |
| CTACTGCAACGTCCTCTGCG | TNFRSF6B | 8771 |
| CTAGAAGTCTAGAGTCACGG | NO_CURRENT_476 | NO_CURRENT_476 |
| CTAGCCGCCCAGATCGAGCC | NO_CURRENT_477 | NO_CURRENT_477 |
| CTAGCTCAATATACAATGTG | NO_CURRENT_478 | NO_CURRENT_478 |

| | | |
|---|---|---|
| CTAGCTGAACAGATGTTTCA | SPCS1 | 28972 |
| CTAGCTTAGGAGTTATACCG | NO_CURRENT_479 | NO_CURRENT_479 |
| CTAGTGACCACTCCACACAA | KMT2C | 58508 |
| CTATAGCCATCAGGTTTGGG | PPARG | 5468 |
| CTATATTGTCGCGCAGTGGA | NO_CURRENT_480 | NO_CURRENT_480 |
| CTATCAAACTGGATATCACG | ALOX5 | 240 |
| CTATCCTGACTGATGCCCAA | NO_CURRENT_481 | NO_CURRENT_481 |
| CTATGACTGCCCAAATCGTG | SUV39H1 | 6839 |
| CTATGCTCACCCCAGACACG | AGER | 177 |
| CTATGTGGCAGATATCGAGG | RHOA | 387 |
| CTATTACCTGGTCGGAACCC | NDUFS4 | 4724 |
| CTCAAAATGCTCCCCGAGAG | HSH2D | 84941 |
| CTCAAACATGCTCTGCGACA | BRD1 | 23774 |
| CTCAACAGCTAATTTGGCTG | EIF2AK2 | 5610 |
| CTCAAGCGCGAGCCCGACTG | CEBPD | 1052 |
| CTCAATTACTCACCCAGCCA | CYP4F11 | 57834 |
| CTCACACAGGTCATGATGTT | CDKN2D | 1032 |
| CTCACACATCTAAACTTAAG | ANP32B | 10541 |
| CTCACCCAGTGACAACTCAG | AKT1 | 207 |
| CTCACCGTTCACCAGCGCCG | ATP5L | 10632 |
| CTCACCTGCAATTAGGTGGA | UBE2E1 | 7324 |
| CTCACGGGGACATACAGGGC | NO_CURRENT_482 | NO_CURRENT_482 |
| CTCACTCCACTCACCAACAT | COX16 | 51241 |
| CTCAGCCCTTTATCGCTGTG | JAM3 | 83700 |
| CTCAGCTAGCACCAAGTCAT | GNAI2 | 2771 |
| CTCAGGCAGCAGAACAAGGA | KPNB1 | 3837 |
| CTCAGTAACAGAGAGTCTCC | MSL2 | 55167 |
| CTCAGTGCCGTGTTTCTGTG | ABCB10 | 23456 |
| CTCATCACGAATGAAGGACA | NCKAP1L | 3071 |
| CTCATCATACTGGCTAGTGG | CTNNB1 | 1499 |
| CTCATCATAGTAACAGGCGT | LGMN | 5641 |
| CTCATCCCCTTCTCCCATCG | NOS2 | 4843 |
| CTCATGAGTCGTTTCTTTCA | NO_CURRENT_483 | NO_CURRENT_483 |
| CTCATGGGACTCCGGCGCAG | RRAGC | 64121 |
| CTCATGTCCTGTGTTTCGAA | IRGM | 345611 |
| CTCATTCTCCGCATTGCCTG | TFPT | 29844 |
| CTCATTTCTGCTGGCCAAAG | SEC62 | 7095 |
| CTCCAACCAGTTTCAGACCG | ITGB2 | 3689 |
| CTCCACAAAGCACACACATC | BMI1 | 648 |
| CTCCACAAAGCACACACATC | COMMD3-BMI1 | 100532731 |
| CTCCACCAGGCCCTAAACCC | WASF2 | 10163 |
| CTCCACCATGAAATGCAGCG | CASP9 | 842 |
| CTCCACTAAGCTCAACTGGA | EDA2R | 60401 |
| CTCCACTCCACATTCTCAGG | EZH2 | 2146 |
| CTCCACTTTCAGTGGGAAAG | SUPT7L | 9913 |
| CTCCAGGACTACCCATCGGT | ITGB2 | 3689 |
| CTCCAGTATGTGCAGGACAT | CCNC | 892 |
| CTCCAGTGCTGCCCAACCAG | SHOC2 | 8036 |
| CTCCATGCCACCATAAGGGA | LTA4H | 4048 |
| CTCCCAGTACCAGTCAGTTC | NO_CURRENT_484 | NO_CURRENT_484 |

| | | |
|---|---|---|
| CTCCCATTCTACTGATATCG | MMP1 | 4312 |
| CTCCCATTGATCTACGATGG | NO_CURRENT_485 | NO_CURRENT_485 |
| CTCCCCAGCATGACCAACGG | NMRK2 | 27231 |
| CTCCCGACAAGGCATCGCCG | CYP27B1 | 1594 |
| CTCCCTGCCGGCCGGGTTAG | NO_CURRENT_486 | NO_CURRENT_486 |
| CTCCGCAGACATAGTGACAC | LTB4R | 1241 |
| CTCCGCCAGCGCGGTCCAGT | MYD88 | 4615 |
| CTCCTCTTCTGGTACCACCG | SNRNP70 | 6625 |
| CTCCTTACGTCGGGCATTAA | NO_CURRENT_487 | NO_CURRENT_487 |
| CTCGAACGGGTAGTTGATGG | NDUFS8 | 4728 |
| CTCGACAGTTCGTCCCGAGC | NO_CURRENT_488 | NO_CURRENT_488 |
| CTCGAGCAGTCGGCCTACAG | MYD88 | 4615 |
| CTCGCACCATTGAGGGTAGT | NO_CURRENT_489 | NO_CURRENT_489 |
| CTCGGCCCACTTGCGTTATA | NO_CURRENT_490 | NO_CURRENT_490 |
| CTCGGGAGGGACTTCAACGA | TP73 | 7161 |
| CTCGGGCCCAGAGTCACATG | NO_CURRENT_491 | NO_CURRENT_491 |
| CTCGGGGATTACCATGGTGC | ATP5J2 | 9551 |
| CTCGGTGAAAAGCCGCACTG | NENF | 29937 |
| CTCTAAACTGTAAAGTCCAA | NO_CURRENT_492 | NO_CURRENT_492 |
| CTCTCCGTATCAGACCCCAG | SETD1B | 23067 |
| CTCTCCTATGCAGTACATAG | PIP4K2A | 5305 |
| CTCTCGCTACTGCTCTAGCG | BAHD1 | 22893 |
| CTCTCGGTACCCGATCGCCG | EBI3 | 10148 |
| CTCTCGGTAGTCATTCAGGT | ANP32B | 10541 |
| CTCTCTCCTACCTGTCCACC | NO_CURRENT_493 | NO_CURRENT_493 |
| CTCTGAGCCATACCTATCCG | SIRT1 | 23411 |
| CTCTGCGAGAATTTCTCGGA | PDHA1 | 5160 |
| CTCTGCTCAGAGACTATGTG | ARR3 | 407 |
| CTCTGGTCTGAGCACCACTG | CASP9 | 842 |
| CTCTTATTGGGTGTATATCC | NO_CURRENT_494 | NO_CURRENT_494 |
| CTCTTTGCGAAAGAATCGCG | CASP5 | 838 |
| CTCTTTTGAGATTGACAAGT | NO_CURRENT_495 | NO_CURRENT_495 |
| CTGAACAAACTCAGCAAGCG | USP7 | 7874 |
| CTGAACAACAGCTACAGTTG | NMU | 10874 |
| CTGAACAGATTTCCTGCCCG | CCAR2 | 57805 |
| CTGAACATCAACAAATCTTG | AHCTF1 | 25909 |
| CTGAACTGCTCTACAGAGTG | CFLAR | 8837 |
| CTGAAGAACCATCTACTGTG | FAM214B | 80256 |
| CTGAAGAGCAGGCTGAGAGG | NAIP | 4671 |
| CTGAAGTGCAGCTTGCGACA | METTL3 | 56339 |
| CTGAATCCAGCAGCCCTCAG | ARR3 | 407 |
| CTGACAAATCAGAAGATGGA | TMEM70 | 54968 |
| CTGACACACCGGCCCCTAAG | INO80E | 283899 |
| CTGACATGAAGCTAGCGGTG | SLC5A8 | 160728 |
| CTGACCACGGGAGCTAGTGA | GPT2 | 84706 |
| CTGACGCCCTTCACCGCGCG | BCL2 | 596 |
| CTGACGGAGCTGGTCCACAG | SLC25A19 | 60386 |
| CTGACTGAGCACTGTCATAG | NO_CURRENT_496 | NO_CURRENT_496 |
| CTGAGATCACGTCACTACAC | NO_CURRENT_497 | NO_CURRENT_497 |
| CTGAGATGTCAGATGAACCG | NO_CURRENT_498 | NO_CURRENT_498 |

| | | |
|---|---|---|
| CTGAGGTTGGTGTAAACGGG | JUNB | 3726 |
| CTGAGTAAGTGATCACGAAG | NO_CURRENT_499 | NO_CURRENT_499 |
| CTGAGTGAAAAATAAAAGTT | NO_CURRENT_500 | NO_CURRENT_500 |
| CTGATGAGAAAGGATCCCAG | HSH2D | 84941 |
| CTGATGTCAACTAAACACCA | NO_CURRENT_501 | NO_CURRENT_501 |
| CTGATTAAGAAATACCCATA | STRAP | 11171 |
| CTGATTGTTGGAGGTTCTTT | COX16 | 51241 |
| CTGATTGTTGGAGGTTCTTT | SYNJ2BP-COX16 | 100529257 |
| CTGCACACATTTGAGTCCCA | MTOR | 2475 |
| CTGCAGAGAAAACTGAACTC | TIFA | 92610 |
| CTGCCAGTTTCGCTCCGTGG | PPARG | 5468 |
| CTGCCCCAGGCGTAATCCTC | NO_CURRENT_502 | NO_CURRENT_502 |
| CTGCCCTGAATGGCCACATG | TRERF1 | 55809 |
| CTGCCGGCTCAAACGCTGTG | VDR | 7421 |
| CTGCCTCGCATCCAACAAGG | SPINT1 | 6692 |
| CTGCGTGTCTTGCTCGCATG | NO_CURRENT_503 | NO_CURRENT_503 |
| CTGCGTGTGTTGGGATCGTG | TNFRSF1B | 7133 |
| CTGCTACATCCGGGACAGTG | TYK2 | 7297 |
| CTGCTCATTGAGGATGCCGG | NOD1 | 10392 |
| CTGCTCTCAACATGCGAGTG | MYD88 | 4615 |
| CTGCTCTCTGTAGCACGTGG | RBM42 | 79171 |
| CTGCTGCCGCCCGCAAGCAG | SERF2 | 10169 |
| CTGCTTGTGCTCGGGCTGCG | C16orf72 | 29035 |
| CTGGAACAAGCCCTGAATGG | BCORL1 | 63035 |
| CTGGAATATCACAAGGTCTG | NQO1 | 1728 |
| CTGGACCTGGTACAGAGCAA | NOD1 | 10392 |
| CTGGAGAACCTGAACAAGTG | PDE4A | 5141 |
| CTGGAGCAGATCCCTGACCT | SCN5A | 6331 |
| CTGGAGCGGAAGCCTAGCGG | NUDT17 | 200035 |
| CTGGAGGTTATCGATCCCCA | GPR119 | 139760 |
| CTGGATCGCCCGCAGAAATA | NO_CURRENT_504 | NO_CURRENT_504 |
| CTGGCACTTGCGACGCATGT | NR1H3 | 10062 |
| CTGGCCGAATCTCACTATGT | NO_CURRENT_505 | NO_CURRENT_505 |
| CTGGCCGTGGAGGATACCGG | PAGR1 | 79447 |
| CTGGCCTGCCACAATTGAAG | NOD2 | 64127 |
| CTGGCGACATTCCTCTTACA | C4orf17 | 84103 |
| CTGGCGCCTATGGCTCTGTG | MAPK14 | 1432 |
| CTGGCGGTCAGACTGGACCA | HELZ2 | 85441 |
| CTGGCTCATGCTCAACGAGA | PTGES2 | 80142 |
| CTGGGAGATGAAGACAGACC | C6orf15 | 29113 |
| CTGGGAGTATCGGATGTAGC | ATP5E | 514 |
| CTGGGATGTGGATGGAAGCA | NDUFA12 | 55967 |
| CTGGGATTCAGGGTACCCCA | C6orf15 | 29113 |
| CTGGGCCATATGTGGGCCGA | TLR9 | 54106 |
| CTGGGCTCCAATTATGAAAT | MPC2 | 25874 |
| CTGGGCTTATGGGATACAGC | RAC1 | 5879 |
| CTGGGGAAGAACCCGACCTG | ARHGAP33 | 115703 |
| CTGGGTGACTGTGCGCACCC | TRMT61A | 115708 |
| CTGGTCGCGCACGATCAGGG | IRAK1 | 3654 |
| CTGGTCGGGGACGTGCAGTG | RND1 | 27289 |

| | | |
|---|---|---|
| CTGGTCTTGGTACCATGCTA | UCHL3 | 7347 |
| CTGGTGACCGACAATTACAC | NO_CURRENT_506 | NO_CURRENT_506 |
| CTGGTGCCAGACATCATGTG | SMPD1 | 6609 |
| CTGGTGCGGGAGATGACACA | SLC6A4 | 6532 |
| CTGGTGCTGCTGAACCCGCG | SPHK1 | 8877 |
| CTGGTTCAAAACAGGAGTCA | ARNT | 405 |
| CTGGTTGGAAGACGAAGATG | UBE2E1 | 7324 |
| CTGTAAACTACGGATGGACC | PLD1 | 5337 |
| CTGTAAATATCCAAGAACCA | CD84 | 8832 |
| CTGTACCAAGAGTTTGCTCC | CCL20 | 6364 |
| CTGTAGACAGCAGTGTCCCA | MYD88 | 4615 |
| CTGTATAATAATCATAAATC | NO_CURRENT_507 | NO_CURRENT_507 |
| CTGTCACCCCCAATAGCGGT | SETD1B | 23067 |
| CTGTCACTCGCATGTGTGGG | TIRAP | 114609 |
| CTGTCCAAGGATCACAGTCA | SLC5A8 | 160728 |
| CTGTCCACCTACAGCGATGT | NO_CURRENT_508 | NO_CURRENT_508 |
| CTGTCCAGTCCCCAACGATG | CYBB | 1536 |
| CTGTCCATCAAGACTAGCAG | BAP1 | 8314 |
| CTGTCGCTGAGTAACGTCAA | TNRC18 | 84629 |
| CTGTCGGACAACCGCATCAG | TLR9 | 54106 |
| CTGTCGGATGACATTTGCCA | WASF2 | 10163 |
| CTGTCTAGCAATGAACCATT | IRAK4 | 51135 |
| CTGTCTGCAAACATATCACA | PPARG | 5468 |
| CTGTGAAAACTGATGCGGAT | PAXIP1 | 22976 |
| CTGTGACTCTGTGACTCGGT | NO_CURRENT_509 | NO_CURRENT_509 |
| CTGTGATGATGTCCCGCGCA | IRAK1 | 3654 |
| CTGTGCCGTAGACAGCCCCT | CASP9 | 842 |
| CTGTGGACTGAGCTCTTCCG | NDUFS8 | 4728 |
| CTGTGGCGTGAGATCACCAA | ARID3A | 1820 |
| CTGTGGGAGAGCCATGCACG | KDM4A | 9682 |
| CTGTGTTTCCACAGAACCCA | RIPK1 | 8737 |
| CTGTTACTATAACACCTACC | LAMTOR3 | 8649 |
| CTGTTACTGAAATGTGCGTG | NAIP | 4671 |
| CTGTTATGTCCATCTAACAC | NO_CURRENT_510 | NO_CURRENT_510 |
| CTGTTCTAATTACCATGCAA | CCNC | 892 |
| CTGTTCTAGCAAAAGCGCAA | PDAP1 | 11333 |
| CTGTTGATCCACAAGAATGA | GPR119 | 139760 |
| CTGTTGCCAATACACAATCA | KMT2C | 58508 |
| CTGTTGGAAGACGCTAGGCA | NO_CURRENT_511 | NO_CURRENT_511 |
| CTGTTTGCGGATAGGATAGG | RAC1 | 5879 |
| CTTAACGGCAATGACGTGGC | MAP2K7 | 5609 |
| CTTAAGCTTGTAGAGGATCG | ARF1 | 375 |
| CTTAAGGCGAGAAAAATTAG | NO_CURRENT_512 | NO_CURRENT_512 |
| CTTAATATGCAAGACTCTCA | CASP1 | 834 |
| CTTACACATTAGTGTGAAGC | VDAC1 | 7416 |
| CTTACATGATTCATCCTATA | CCNC | 892 |
| CTTACTTCGGATATGGGACT | AHR | 196 |
| CTTAGAGACAGTGTCCACTC | CHEK2 | 11200 |
| CTTAGCCCACTGATGAACGA | CREBBP | 1387 |
| CTTAGCGCGCTCTCGCCGTG | PLD6 | 201164 |

| | | |
|---|---|---|
| CTTAGCTGACCGACAAGGTG | NO_CURRENT_513 | NO_CURRENT_513 |
| CTTAGGATTCCGAGGTATCT | NO_CURRENT_514 | NO_CURRENT_514 |
| CTTAGGCTATAATCACAATG | ABL1 | 25 |
| CTTAGTCACCCCAACAAGCG | NO_CURRENT_515 | NO_CURRENT_515 |
| CTTATAGGTAGTATCGCAGA | NO_CURRENT_516 | NO_CURRENT_516 |
| CTTATAGGTATCCACATCCG | BCL2A1 | 597 |
| CTTATCTACCAAATTCTCCG | MAPK14 | 1432 |
| CTTATCTGCTACAAGGAAGG | HSP90B1 | 7184 |
| CTTATGTCAGGTTATTCACA | MAPK9 | 5601 |
| CTTCAAAAATTAAACTAGCA | PDE5A | 8654 |
| CTTCAACAGCAATCACAAGG | CEBPD | 1052 |
| CTTCAACTATGCCATGAAGG | NQO1 | 1728 |
| CTTCACGGTTTCGAGAGCAA | FOXO4 | 4303 |
| CTTCAGTCGTAGAGATCCAG | STAG2 | 10735 |
| CTTCATCAACTACCTCACGA | SPINT1 | 6692 |
| CTTCATCCTAAATTTATTGT | CCL20 | 6364 |
| CTTCCCTATGGGACTCAACG | KMT2D | 8085 |
| CTTCCGGGACATCACCCACG | TYK2 | 7297 |
| CTTCCGTTATTCGGAAGTGA | NO_CURRENT_517 | NO_CURRENT_517 |
| CTTCCTTCTGCTTTATTACT | FAM107B | 83641 |
| CTTCCTTGGAGCCAAAATTG | SLC3A2 | 6520 |
| CTTCGACGCCATCGTGCTCA | NO_CURRENT_518 | NO_CURRENT_518 |
| CTTCGGGGCCAATCATGGCA | CTNNBL1 | 56259 |
| CTTCTACGAACCGGGCCGGG | CEBPD | 1052 |
| CTTCTCTCAGTGTCACTGCA | WASF2 | 10163 |
| CTTCTGCATACGTGATGAAG | SLC6A4 | 6532 |
| CTTCTGTGGACGATTACATG | IRF8 | 3394 |
| CTTCTTTCAGCAGGCGGCAA | RIPK3 | 11035 |
| CTTGAAATATACCGTAGCAG | CCNC | 892 |
| CTTGACCGAAGCCTGCTGTG | GC | 2638 |
| CTTGACTTACGAGCCAGTGT | NO_CURRENT_519 | NO_CURRENT_519 |
| CTTGAGTTTATGTTTCACGG | NO_CURRENT_520 | NO_CURRENT_520 |
| CTTGCACTCTTCATACGGGA | NMRK2 | 27231 |
| CTTGCCAAGTCATTAGACCT | C6orf15 | 29113 |
| CTTGCTCTTACACAGAGTGT | METTL3 | 56339 |
| CTTGCTGTCCACCATCACAT | RAC2 | 5880 |
| CTTGCTTTAGACGTGCAGCG | GSDMD | 79792 |
| CTTGGGCGATCCATATCTCT | BCL2L11 | 10018 |
| CTTGGTGCAACTGAAAACAT | MBNL1 | 4154 |
| CTTGGTGTTCTCGTCATACA | JAK1 | 3716 |
| CTTGTGTACAGTTAGTAGTG | ZNF616 | 90317 |
| CTTGTTCCGAAACTGCACAC | SCTR | 6344 |
| CTTGTTGCGTATACGAGACT | NO_CURRENT_521 | NO_CURRENT_521 |
| CTTGTTTCTTATAGGCTGAA | HMGN2 | 3151 |
| CTTTAAACTCCATTCCCTCA | TLR2 | 7097 |
| CTTTAGAAGGACATGTGGAC | NO_CURRENT_522 | NO_CURRENT_522 |
| CTTTAGCTGGATGTCCACGT | IDH2 | 3418 |
| CTTTAGCTTGATATGCAACG | NIPBL | 25836 |
| CTTTAGTTTACGGAGAACTG | MEMO1 | 51072 |
| CTTTATCATGCAATGCACAG | PHF6 | 84295 |

| | | |
|---|---|---|
| CTTTCACACAAAGATTGGGT | PIAS1 | 8554 |
| CTTTCCAAGATATCGGAAAC | DNAJB6 | 10049 |
| CTTTCCAGGTAAATATGCAG | WASF2 | 10163 |
| CTTTCCCACATACCAACAGA | PHF8 | 23133 |
| CTTTCCGTGGCAGGAGACAG | FOSL2 | 2355 |
| CTTTCTTGTTCAGTGATCGG | CAT | 847 |
| CTTTGAGCAGACCTTGACAG | PPARGC1B | 133522 |
| CTTTGAGCGGATAAACAGGA | NLRP12 | 91662 |
| CTTTGCTCAACAATGCAAGG | NNT | 23530 |
| CTTTGGGTGCGTGTACCGGG | IRAK1 | 3654 |
| CTTTTTTTATTTATCGATCG | NO_CURRENT_523 | NO_CURRENT_523 |
| GAAAAAAGGGTTGCTCATTT | NDUFA1 | 4694 |
| GAAAACGTTGGTCTGAGGGT | RHOG | 391 |
| GAAAACTTAGTAGAACTACT | IRAK4 | 51135 |
| GAAAATCCTCACTCTGAGTA | DPY30 | 84661 |
| GAAACAAATAGTGATTACTC | NO_CURRENT_524 | NO_CURRENT_524 |
| GAAACAAGCAAGAAAATGGA | ZNF699 | 374879 |
| GAAACAGATGACGAAAACGT | RHOG | 391 |
| GAAACATAATGTCTTCAGTC | NO_CURRENT_525 | NO_CURRENT_525 |
| GAAACCGATCGACCGCGAGA | SAP18 | 10284 |
| GAAACGAGAAGTTTGTACTA | NO_CURRENT_526 | NO_CURRENT_526 |
| GAAACTCATACTGGGTGTCT | IL2RB | 3560 |
| GAAACTCCCTTCCAGCGAAG | TIFA | 92610 |
| GAAACTGACCAAAGTCACTC | ABCB10 | 23456 |
| GAAACTTACTGCAAAAGAGG | KPNA1 | 3836 |
| GAAAGAAACAAAGTATACTG | EIF2AK2 | 5610 |
| GAAAGACCTTGTAACCATGA | NO_CURRENT_527 | NO_CURRENT_527 |
| GAAAGCAGGTTAAGACATGA | NO_CURRENT_528 | NO_CURRENT_528 |
| GAAAGTACAATTTGCACCAC | ARR3 | 407 |
| GAAAGTACCAGGCACTAGGT | TFPT | 29844 |
| GAAAGTGCACATCGAGAGAG | MEMO1 | 51072 |
| GAAATCCCCGTTTACTACGA | NO_CURRENT_529 | NO_CURRENT_529 |
| GAAATCGACACCTGAAAAAG | TNFRSF1A | 7132 |
| GAAATGACACAGCAAAATGG | CXCR1 | 3577 |
| GAAATGACAGTTATCCGAAG | LRP2 | 4036 |
| GAAATGCGCATCCTCATGGT | ARF1 | 375 |
| GAACAAAAAATCTCAGAATG | CYBB | 1536 |
| GAACAAAAGATAGTGTTATG | IFNAR1 | 3454 |
| GAACAAACCAGAATTGCAGA | FAM107B | 83641 |
| GAACAAACTAGCCCTCAAGA | TRIR | 79002 |
| GAACAACCTGAACGTCACCG | SOD2 | 6648 |
| GAACAAGATTCTCAGAATGA | PAXIP1 | 22976 |
| GAACACCCGCAGGTACACGA | ADRB1 | 153 |
| GAACAGCTGGACATCCGTGA | EBI3 | 10148 |
| GAACATCAGCTTCACTGTGT | ARF1 | 375 |
| GAACATGTAAGGGCCCGACA | PIAS1 | 8554 |
| GAACATTGGTGTTACAGCAG | ATP5J | 522 |
| GAACCAAGACCCAGACATCA | IL10 | 3586 |
| GAACCAGATCAGCAGCCTGG | CHMP6 | 79643 |
| GAACCCAACCTTTTACCGCA | NO_CURRENT_530 | NO_CURRENT_530 |

| | | |
|---|---|---|
| GAACCCAATGTGGCTCGCGT | SMPD1 | 6609 |
| GAACCGTCAGACAGTACTAG | BAP1 | 8314 |
| GAACCTCCCCGAATATCTGG | NO_CURRENT_531 | NO_CURRENT_531 |
| GAACGGCTTATTAACTGTGC | PHC3 | 80012 |
| GAACGGTATGACTGTAAAGG | CBLL1 | 79872 |
| GAACGGTGAGAGTCTTTGGT | NO_CURRENT_532 | NO_CURRENT_532 |
| GAACGTAGAAATTCCCATTT | NO_CURRENT_533 | NO_CURRENT_533 |
| GAACGTCACCTTAGAATCCG | NR2C2 | 7182 |
| GAACGTCCAAGCAAGGGAGC | NO_CURRENT_534 | NO_CURRENT_534 |
| GAACGTGCGAGACAATGTTG | DNTTIP1 | 116092 |
| GAACGTGGATCGCGCTCGAG | MEX3B | 84206 |
| GAACGTTCCCTCAAAATGCA | NO_CURRENT_535 | NO_CURRENT_535 |
| GAACTACATCAGGGGATGCA | IKZF5 | 64376 |
| GAACTCGAACATTCCAAACC | SLC25A1 | 6576 |
| GAACTCTTACTCATTGCCGG | NO_CURRENT_536 | NO_CURRENT_536 |
| GAACTTTAAGGCAAACTCAT | MOSPD1 | 56180 |
| GAAGAAAAGCAGCCTCGCGG | MCL1 | 4170 |
| GAAGAAAATGAAATTAGGCA | NIPBL | 25836 |
| GAAGAAATACAGCCTGACGG | ABL1 | 25 |
| GAAGAACTTTGTGGTCGTCA | RAD23A | 5886 |
| GAAGAAGAATGTGCCCATCG | DNTTIP1 | 116092 |
| GAAGAAGCAGAGTCTGACTG | WASF2 | 10163 |
| GAAGAAGGGGAAGTGACCTC | PAGR1 | 79447 |
| GAAGAAGTCATACACATCAG | CYP4B1 | 1580 |
| GAAGACAGGAATGCCCGTCT | NLRP3 | 114548 |
| GAAGACCCCAGTTGGCTACT | MPC1 | 51660 |
| GAAGACGGCATCATTAACTG | BCL2A1 | 597 |
| GAAGACTCAACAGATATGTT | NO_CURRENT_537 | NO_CURRENT_537 |
| GAAGAGATAGCTGATTCCAA | GLMN | 11146 |
| GAAGAGATTCAAGTATTACG | SLC7A11 | 23657 |
| GAAGAGCTCAGTCCACAGCA | NDUFS8 | 4728 |
| GAAGAGTCTGTTATCTCCAA | GLMN | 11146 |
| GAAGATAGTATGTTCAAACA | NCKAP1L | 3071 |
| GAAGATTCGGAGCCTTGATG | PRKAA1 | 5562 |
| GAAGCAGAAGACGGAGGATG | TRIR | 79002 |
| GAAGCAGAGGCTCTTCAGGG | IKZF5 | 64376 |
| GAAGCAGTGCAAACGCCATG | TRAF6 | 7189 |
| GAAGCATCTATTATCGTACC | NO_CURRENT_538 | NO_CURRENT_538 |
| GAAGCATGGTGCCTCCTGAA | NDUFA12 | 55967 |
| GAAGCCATCAAACGTGACTT | SOD2 | 6648 |
| GAAGCCCGTGATGCACTCAA | IKBKB | 3551 |
| GAAGCGATCCGAGAGTGTAT | NO_CURRENT_539 | NO_CURRENT_539 |
| GAAGCGCTGTTTGCCAGTTA | AIM2 | 9447 |
| GAAGCGGGACCGTGTCTCAC | NO_CURRENT_540 | NO_CURRENT_540 |
| GAAGCTGAGCCAGTACGTAA | EIF4G3 | 8672 |
| GAAGCTTGTTCGTAAAACGC | HSP90B1 | 7184 |
| GAAGGAAGTATAGGTAGCCG | ABL2 | 27 |
| GAAGGACTTTCTAATAACCC | LARP4B | 23185 |
| GAAGGAGCAGGAAAACCGGA | PARK7 | 11315 |
| GAAGGATCACAAAAAAAGTG | OR7C2 | 26658 |

| | | |
|---|---|---|
| GAAGGCCTGAACTTACAAGA | AHR | 196 |
| GAAGGCTCTAGGGAAACACA | HNRNPF | 3185 |
| GAAGGGTGTAACATTACTGA | ANKRD50 | 57182 |
| GAAGGTAATCAAGTTACATG | NO_CURRENT_541 | NO_CURRENT_541 |
| GAAGGTGCGTTCGATGACAG | AKT1 | 207 |
| GAAGGTTGGTACATTAGTGG | NDUFA12 | 55967 |
| GAAGGTTGTTCCCACCCCAG | SLC6A4 | 6532 |
| GAAGTAAAACCAGCCAACGA | NO_CURRENT_542 | NO_CURRENT_542 |
| GAAGTACCTAGAATACCTGG | WDR26 | 80232 |
| GAAGTCAGCATAGTCCAGGC | ABI1 | 10006 |
| GAAGTCCTCTTGCAAGACTG | PPARGC1A | 10891 |
| GAAGTCCTTTGGAAACGAAT | RRAGC | 64121 |
| GAAGTGACGGGATTCACTCT | PDE3A | 5139 |
| GAAGTGGAGTTTGTGTCAGG | TFPT | 29844 |
| GAAGTGGTATGTGGTAGGCC | LCN2 | 3934 |
| GAAGTGGTGTGTATGTGACT | NO_CURRENT_543 | NO_CURRENT_543 |
| GAAGTTACAGAGATGGAGTG | IRF8 | 3394 |
| GAAGTTGCTCACAATTGACC | APEH | 327 |
| GAAGTTTCGCCAGAATACTT | TLR1 | 7096 |
| GAAGTTTGCCCGTGCCCACG | NMRK2 | 27231 |
| GAATAAAATATCTTTAGAGT | COX16 | 51241 |
| GAATAAAATATCTTTAGAGT | SYNJ2BP-COX16 | 100529257 |
| GAATAAATCTGAGTTATACG | NO_CURRENT_544 | NO_CURRENT_544 |
| GAATAACGAGGGATCACTGC | LAMTOR2 | 28956 |
| GAATACAAGCAGAAAATTAA | UBE2H | 7328 |
| GAATACTAATAATCATGTAA | NO_CURRENT_545 | NO_CURRENT_545 |
| GAATAGATTTGTCAGTTAGG | NO_CURRENT_546 | NO_CURRENT_546 |
| GAATCGACCGACACTAATGT | NO_CURRENT_547 | NO_CURRENT_547 |
| GAATCTCCTTGTTTAACACG | NIPBL | 25836 |
| GAATCTTAGGGACAAAAAAT | OXSM | 54995 |
| GAATGAAAAGGATATGGCGA | NDUFB9 | 4715 |
| GAATGAATACCGGTCCCGTG | NOS2 | 4843 |
| GAATGACAGCGAAACCAGTT | CDKN2C | 1031 |
| GAATGACATTCATGTCCCCG | NQO1 | 1728 |
| GAATGGACGTTACAAAGACA | NO_CURRENT_548 | NO_CURRENT_548 |
| GAATGGCAAAAACACATTCT | NDUFA12 | 55967 |
| GAATGTATGCATATGCTGGC | SLC7A11 | 23657 |
| GAATTAGATGGATTAAGAGA | UCHL5 | 51377 |
| GAATTCATGATTCTGAACGC | AHCTF1 | 25909 |
| GAATTCTAGTAAATTCAGGG | NO_CURRENT_549 | NO_CURRENT_549 |
| GAATTCTCAGTCCTACATAC | CYP4A11 | 1579 |
| GAATTTCCTGAAGAACGTTG | SQSTM1 | 8878 |
| GACAAACCAATTGCTCCTTG | TMEM30A | 55754 |
| GACAAAGGACCCTTCCGGGT | NQO1 | 1728 |
| GACAAAGTGGAACGCCTCCA | BAP1 | 8314 |
| GACAACCGCCATCCAGACTG | AKT1 | 207 |
| GACAACTATCTTGTTGCTTG | WDR26 | 80232 |
| GACAACTTGGTGTATAGACA | TLR1 | 7096 |
| GACAATAATCCACACCAGAA | PRKAA1 | 5562 |
| GACAATCATGGTGAAAGCGG | NO_CURRENT_550 | NO_CURRENT_550 |

| | | |
|---|---|---|
| GACACCCAGTGAGCCGGAGT | AHDC1 | 27245 |
| GACACGGGCCACACGTCGTG | TNFRSF6B | 8771 |
| GACAGATCACGTATCAGGGC | NO_CURRENT_551 | NO_CURRENT_551 |
| GACAGATTATTTACCAGTGA | NO_CURRENT_552 | NO_CURRENT_552 |
| GACAGCCTGTGAACAAGACG | NO_CURRENT_553 | NO_CURRENT_553 |
| GACAGGAGGATCCGGTGCAG | BRK1 | 55845 |
| GACAGTGAAATTAGCTCCCA | NO_CURRENT_554 | NO_CURRENT_554 |
| GACAGTGACGAAATAACTGC | NDUFA8 | 4702 |
| GACAGTGCGCACCGTGTACG | CYP27B1 | 1594 |
| GACAGTGGTGCCTACCGAAG | ARMCX2 | 9823 |
| GACAGTTGAGCCCAGTTCCG | ASH2L | 9070 |
| GACATACATGCTGACGCTGG | PDE4A | 5141 |
| GACATATCTGAAATTAGTCG | NO_CURRENT_555 | NO_CURRENT_555 |
| GACATCCCTGAAGATGACGT | PPARGC1B | 133522 |
| GACATCCTGACTCTTGCATG | WDR26 | 80232 |
| GACATCTTCCAACAGAACGT | PHF8 | 23133 |
| GACATGGAGGTGATCACTGT | FOSL2 | 2355 |
| GACATTACTCCAGAGTTGGA | MMP1 | 4312 |
| GACATTGGTCAGATAGCCAG | NO_CURRENT_556 | NO_CURRENT_556 |
| GACCACTCTAGCACCAGTCA | NO_CURRENT_557 | NO_CURRENT_557 |
| GACCAGCCCACACTTGTGAG | KHSRP | 8570 |
| GACCCCAATTCAGAACACCA | INO80 | 54617 |
| GACCCCACACACCGTCCAGA | SLC7A11 | 23657 |
| GACCCCCGATAACTTTTGAC | NO_CURRENT_558 | NO_CURRENT_558 |
| GACCCGTGGGTGCAATGGTG | TRMT61A | 115708 |
| GACCGGACACTCGAAGAGCG | SIAH2 | 6478 |
| GACCGTTTCATAGTAGATGT | NO_CURRENT_559 | NO_CURRENT_559 |
| GACCTACTGCAGGGATACGA | DET1 | 55070 |
| GACCTCACCATTCCCAACGG | NFKB1 | 4790 |
| GACCTGAAACACCAAAACAC | NIPBL | 25836 |
| GACCTTATAAATGAGTCACA | NO_CURRENT_560 | NO_CURRENT_560 |
| GACCTTCATTGAAGAAAAGC | NO_CURRENT_561 | NO_CURRENT_561 |
| GACGAGGGCGGCAGAGCAGT | NO_CURRENT_562 | NO_CURRENT_562 |
| GACGCCAGTGTCCTATGACC | RND3 | 390 |
| GACGCCCTAATGCCCATCGT | NO_CURRENT_563 | NO_CURRENT_563 |
| GACGCCTTGCCCGGCTCACA | NO_CURRENT_564 | NO_CURRENT_564 |
| GACGGCTGACCGAATGACAG | ATP5J | 522 |
| GACGGCTGGACAGCACGCGT | SLC25A1 | 6576 |
| GACGTAGCCTTCCGAAATAT | NO_CURRENT_565 | NO_CURRENT_565 |
| GACGTTCAGGTTGTTCACGT | SOD2 | 6648 |
| GACTAAAAGAAGACTATAGC | CHUK | 1147 |
| GACTACCCTCTCATCCCAGT | CAT | 847 |
| GACTAGCTCAGACCACCCAG | SPINT1 | 6692 |
| GACTCACATCCATTGGACAC | NO_CURRENT_566 | NO_CURRENT_566 |
| GACTCCAAAATGGCGTCAGT | ATP5J2 | 9551 |
| GACTCCAAAATGGCGTCAGT | ATP5J2-PTCD1 | 100526740 |
| GACTCGGAGACCGGATAACA | TMEM30A | 55754 |
| GACTGACTGCATTGGCACGG | CHMP5 | 51510 |
| GACTGCAGATACGGACTCAG | TNRC18 | 84629 |
| GACTGCATGTTATTGCGAGC | NDUFS4 | 4724 |

| | | |
|---|---|---|
| GACTGTCTTGCTAACCCGAA | NUDT17 | 200035 |
| GACTTACTCGAAGCCTCGAT | ATP5L | 10632 |
| GACTTCACTGTGCAGCCCAG | GBP7 | 388646 |
| GACTTCGGGAACTATCACCT | SLC7A5 | 8140 |
| GACTTCTTGAATGATGCACA | NO_CURRENT_567 | NO_CURRENT_567 |
| GACTTGAACAAACCTGAAGC | STRAP | 11171 |
| GACTTGCTCACGGCCAGACA | MBNL1 | 4154 |
| GACTTTGCTAAACAGCTACC | NR1H3 | 10062 |
| GACTTTGGTTGAGCTTCAAT | NO_CURRENT_568 | NO_CURRENT_568 |
| GAGAAAATTCAAGCAAGTAG | DNAJB6 | 10049 |
| GAGAAAGCGGAGCTACGCTA | NO_CURRENT_569 | NO_CURRENT_569 |
| GAGAAAGTGAGAGTAGCCCT | ABL2 | 27 |
| GAGAAATGCTCCACCTCTCG | PDK4 | 5166 |
| GAGAACCTAGAAATCATACG | EGFR | 1956 |
| GAGAAGATGATGACTACTGT | APAF1 | 317 |
| GAGAAGGATGGAAATTAGAA | NO_CURRENT_570 | NO_CURRENT_570 |
| GAGAAGGCAGGTCTTGCCCA | RAC2 | 5880 |
| GAGAAGGGTCTGTACCTCAG | CREBBP | 1387 |
| GAGAAGTGGGGAGCCATTGG | NO_CURRENT_571 | NO_CURRENT_571 |
| GAGACACAAGCGGACCCCAC | MYD88 | 4615 |
| GAGACAGGCTGGACTCAGGT | ATP5E | 514 |
| GAGACCACTCCGAGATAGCA | FOXO4 | 4303 |
| GAGACCACTTTCGTGCAAGC | NO_CURRENT_572 | NO_CURRENT_572 |
| GAGACCTTACCTGGCGTACC | PDE3A | 5139 |
| GAGACTGATTAAATCAACCA | NO_CURRENT_573 | NO_CURRENT_573 |
| GAGAGAGTCGTAATAATGCA | HSPA4 | 3308 |
| GAGAGCCACTGATTCAGGCC | NO_CURRENT_574 | NO_CURRENT_574 |
| GAGAGGGTCAGCACGAAGTG | GPR119 | 139760 |
| GAGAGTACCACTCAGCACGC | CYP27B1 | 1594 |
| GAGAGTCTGTAAACGCCGTG | WDR26 | 80232 |
| GAGAGTGCGCCTTGATAGTA | NO_CURRENT_575 | NO_CURRENT_575 |
| GAGAGTGCTGGACTCCTACG | TFPT | 29844 |
| GAGATAGCAGCAAACTGAAC | ATP6V0E1 | 8992 |
| GAGATCATACGTGCTCGGTG | RIPK2 | 8767 |
| GAGATCCGATGAGGAGCCGG | PAGR1 | 79447 |
| GAGATGAGACCACATGTTAG | NO_CURRENT_576 | NO_CURRENT_576 |
| GAGATGTGCAGCAAGCTCCG | ARHGAP33 | 115703 |
| GAGATTCCAAAACAATGGCA | NO_CURRENT_577 | NO_CURRENT_577 |
| GAGATTTCACGTAACGACGA | TMEM30A | 55754 |
| GAGCACGCCTTGCTCCTCGG | GNAI2 | 2771 |
| GAGCAGCAGTGAGTAGTCTG | IRF9 | 10379 |
| GAGCAGCTCATTCCAGCCTG | RXRA | 6256 |
| GAGCAGCTGTCAGGTCTTGT | NO_CURRENT_578 | NO_CURRENT_578 |
| GAGCAGTGAGCGCTCCGAGG | MKL1 | 57591 |
| GAGCATATCATGTCCCGAGT | MAP2K5 | 5607 |
| GAGCATCAGAGGGGACGACT | PRR14L | 253143 |
| GAGCATCTCGGGGATTACCA | ATP5J2 | 9551 |
| GAGCATTGGGATCATTGTGA | SNRNP70 | 6625 |
| GAGCCACCCCCACTGCGCAT | FOSL2 | 2355 |
| GAGCCAGAGCAGATGCTGGA | DPY30 | 84661 |

| | | |
|---|---|---|
| GAGCCATGACAAGTCGGACA | PQBP1 | 10084 |
| GAGCCCCTTCTCTCGCAAAA | NO_CURRENT_579 | NO_CURRENT_579 |
| GAGCCCTAGGTTCATATCGG | SPEN | 23013 |
| GAGCGCCAAAACCTCTTCAC | AHDC1 | 27245 |
| GAGCTACATCCAGCGTAATG | SHOC2 | 8036 |
| GAGCTCGTAGCAGTTTGCTA | NO_CURRENT_580 | NO_CURRENT_580 |
| GAGCTGATCCCAAAGTTGGT | PPARG | 5468 |
| GAGCTGATGACCGAGAAAGG | RXRA | 6256 |
| GAGCTGTTGCACATTGTGGG | NO_CURRENT_581 | NO_CURRENT_581 |
| GAGCTTGCAGATTTGGAGAT | ZRSR2 | 8233 |
| GAGCTTGTTCGGCTACACGA | CYP8B1 | 1582 |
| GAGCTTTGCTTTATGATTAT | NMU | 10874 |
| GAGGAAGACATACTCTGGCG | SBNO2 | 22904 |
| GAGGACCCTGGAAATAGGGT | NUFIP2 | 57532 |
| GAGGACCTTAAGGTGACATG | NO_CURRENT_582 | NO_CURRENT_582 |
| GAGGAGGAGCAGGAGGCAAG | HAMP | 57817 |
| GAGGATGTCCTAATGCATGG | RNF111 | 54778 |
| GAGGCAAGATCATACACGAG | RNF25 | 64320 |
| GAGGCAAGCGGCCCAGATCT | HAMP | 57817 |
| GAGGCAGTATACAAGCCCTG | PDAP1 | 11333 |
| GAGGCCGGCGCGGTGATTGG | MCL1 | 4170 |
| GAGGCCTCGGCCAACATAGG | STAT2 | 6773 |
| GAGGCCTCTGAATATTTACG | BRD1 | 23774 |
| GAGGCTAAAGAAACAAGCAT | NO_CURRENT_583 | NO_CURRENT_583 |
| GAGGCTCGGCATTGCACCCG | EBI3 | 10148 |
| GAGGGAAAGGGCTTGCACCG | SQSTM1 | 8878 |
| GAGGGAGGCATAAACAGGCG | PPARGC1B | 133522 |
| GAGGGCCATGTAGTCGCAGT | PLD6 | 201164 |
| GAGGGGGACAAAATCAATGA | DLAT | 1737 |
| GAGGGTAGGAGAGTGTACGG | RHOG | 391 |
| GAGGTACTGTACAAGCCCTA | NO_CURRENT_584 | NO_CURRENT_584 |
| GAGGTAGAAATAGACCTGCT | KAT2A | 2648 |
| GAGGTATTAAAACCCAAGCA | PRR14L | 253143 |
| GAGGTCACAAAATGATCTTG | IFNAR2 | 3455 |
| GAGGTCATGAAAACGGATGG | STAT1 | 6772 |
| GAGGTCGGTAATGAGGCCGT | PDE3A | 5139 |
| GAGGTCTCTCCCGAACAGGT | HNRNPF | 3185 |
| GAGGTGACAAGTACAGCTTG | BABAM2 | 9577 |
| GAGGTGGGAGTCTGAACTGA | OR7C2 | 26658 |
| GAGTAATTTCGAACGTATTG | NO_CURRENT_585 | NO_CURRENT_585 |
| GAGTAGTTCGAATCTATCAC | RAD21 | 5885 |
| GAGTCCAGCCTGTCTCCAGT | ATP5E | 514 |
| GAGTCGAAGATGGTCTAGGA | NO_CURRENT_586 | NO_CURRENT_586 |
| GAGTCGCGCAAGTCCCTGTG | SBNO2 | 22904 |
| GAGTCTAGACCGCTCACTGG | LTB4R | 1241 |
| GAGTGATGCTTAGACTCCGT | NO_CURRENT_587 | NO_CURRENT_587 |
| GAGTGCAGGTAGATGTTGTG | SLC11A1 | 6556 |
| GAGTGCATTCTAGCCTGTCG | SPINT1 | 6692 |
| GAGTGCTATAAATTAGTCTG | NO_CURRENT_588 | NO_CURRENT_588 |
| GAGTGCTGTCCCAATTGCCG | DNAJA1 | 3301 |

| | | |
|---|---|---|
| GAGTGGAGATGAGTTCTGAG | YPEL5 | 51646 |
| GAGTTACCAGAACCTGACTC | ABI1 | 10006 |
| GAGTTATTTATTCTCTCGAG | NO_CURRENT_589 | NO_CURRENT_589 |
| GAGTTCACCCACGGTCATGT | ABCB10 | 23456 |
| GAGTTCAGGAATGCGCAGCG | RBM42 | 79171 |
| GAGTTCTTGTAACCATACAG | NO_CURRENT_590 | NO_CURRENT_590 |
| GAGTTGACATCTGTTATTGA | NO_CURRENT_591 | NO_CURRENT_591 |
| GAGTTGATTGAGGTAAAGCG | METTL3 | 56339 |
| GAGTTTGCCTTAAAGTTCAA | MOSPD1 | 56180 |
| GATAAATTTGTACGAACTCA | NO_CURRENT_592 | NO_CURRENT_592 |
| GATAACATCTCAACTACCAG | NUP153 | 9972 |
| GATACAGTTGAATTACACTC | NO_CURRENT_593 | NO_CURRENT_593 |
| GATAGGGCACAGGTACATTG | HNRNPF | 3185 |
| GATATACGTCATTGACGCAC | RRAGC | 64121 |
| GATATCCCGCGAAAAAATCT | NO_CURRENT_594 | NO_CURRENT_594 |
| GATATCGATGTGGAGCACGG | SQSTM1 | 8878 |
| GATCAACCGCAACGCCCGTG | IRAK1 | 3654 |
| GATCACCAGTGTAAAACGTA | CASP5 | 838 |
| GATCACGCCAGGTCTTGGGA | FOSL2 | 2355 |
| GATCACGGGGCAGACGAGTG | NFKB2 | 4791 |
| GATCAGGACAAAGTTATCAT | HTR2A | 3356 |
| GATCAGTAGTACCTGAACCT | RBM6 | 10180 |
| GATCCAACCAAACCCAAATG | NUDT17 | 200035 |
| GATCCACTTTAACTGAATAG | NO_CURRENT_595 | NO_CURRENT_595 |
| GATCCAGCAATATTTCTTAA | NO_CURRENT_596 | NO_CURRENT_596 |
| GATCCAGGAGTGATCGAGTA | NO_CURRENT_597 | NO_CURRENT_597 |
| GATCCAGGCCCCTCCTCAGT | PYCARD | 29108 |
| GATCCCAAATAGTGGCGCGG | MAP2K5 | 5607 |
| GATCCCATCACCCACGATGA | PDE4A | 5141 |
| GATCCTTCTTATTCCCAACC | RHOA | 387 |
| GATCGTGAGATAATGAGAGA | RAD21 | 5885 |
| GATCGTTGTAGAAGTCTAAC | CASP3 | 836 |
| GATCTAATGTTAAAGACTGG | IFNAR1 | 3454 |
| GATCTAGTCCTCTAATCGAT | NO_CURRENT_598 | NO_CURRENT_598 |
| GATCTCGGGAAAGATGACAA | BCORL1 | 63035 |
| GATCTCTGGCTTCGTTAAGG | SLC7A3 | 84889 |
| GATCTGGTTATATAACGATG | NO_CURRENT_599 | NO_CURRENT_599 |
| GATCTGTATGGGCTGAGACA | POU2F1 | 5451 |
| GATGAACAGAATAACCACCA | HSH2D | 84941 |
| GATGAACGACCCATTTCTGT | IRAK4 | 51135 |
| GATGAACTACCACCGACCGT | LRP2 | 4036 |
| GATGAACTGCCAGATAATAC | IDH2 | 3418 |
| GATGAAGTTATCAGCACACC | KPNA1 | 3836 |
| GATGACGAGTTGTGGTCCCT | MMP9 | 4318 |
| GATGAGAGAGGGTCCTGCAA | CHMP5 | 51510 |
| GATGATGGCTGCTGTCCGAT | HCAR2 | 338442 |
| GATGATGGTGATTGTTGTCG | FBXO38 | 81545 |
| GATGATTCAAAATGCTGGGG | AIM2 | 9447 |
| GATGCACGTGCTGAAAGCCG | RIPK1 | 8737 |
| GATGCAGACTGTACGCCGAA | PDHA1 | 5160 |

| | | |
|---|---|---|
| GATGCCAAAAACACACATGT | CCNC | 892 |
| GATGCCATTACCAGTCTCCG | MMP13 | 4322 |
| GATGCTATAACTACGATTCG | MMP1 | 4312 |
| GATGCTATTGAAACATACAA | CYP2R1 | 120227 |
| GATGCTGGAGGGACAAACGC | DPY30 | 84661 |
| GATGCTTAGGAGTGGCCAAG | MTOR | 2475 |
| GATGGAAAACCGGTGAATCT | RAC1 | 5879 |
| GATGGAGTACCTGTTATTAA | LAMTOR3 | 8649 |
| GATGGATACCTGGTAGACCC | FAM214B | 80256 |
| GATGGCGCGCAGTTGAGTCA | NO_CURRENT_600 | NO_CURRENT_600 |
| GATGGTAGTGGATCTGAGAG | NUFIP2 | 57532 |
| GATGGTTTCGAAACATAAAG | DYRK1A | 1859 |
| GATGTAAAAGCTGCAACGGA | DNAJA1 | 3301 |
| GATGTATCAACCTAGTGCTG | HDAC2 | 3066 |
| GATGTCAGACTATTGCCTCA | CSDE1 | 7812 |
| GATGTCCAGGAAGACGGCGT | TRMT61A | 115708 |
| GATGTCGTCACGTTAACATG | NO_CURRENT_601 | NO_CURRENT_601 |
| GATGTCTGCCATCCTAGATG | HTR3E | 285242 |
| GATGTGATCTATGGTTGCGA | NO_CURRENT_602 | NO_CURRENT_602 |
| GATGTGGATTCAAAACTGAA | PLD1 | 5337 |
| GATGTGGTGGTTCTACCAGG | PARK7 | 11315 |
| GATGTTACTGGCTATGGCAG | LAMTOR2 | 28956 |
| GATGTTAGATACTGCTCTGA | NO_CURRENT_603 | NO_CURRENT_603 |
| GATGTTCGATCCTGTCACTA | MOSPD1 | 56180 |
| GATGTTGCAAGTGTTAGCGA | RND1 | 27289 |
| GATTATGTGGAGGAGAATCG | NLRP1 | 22861 |
| GATTCATACTAAACACTCTA | NO_CURRENT_604 | NO_CURRENT_604 |
| GATTCTGGATGGCGACGTAG | HTR2A | 3356 |
| GATTGAAGTTCATGGCATAG | NQO1 | 1728 |
| GATTGATGTCATGTATGAGG | BMI1 | 648 |
| GATTGATGTCATGTATGAGG | COMMD3-BMI1 | 100532731 |
| GATTGTCTTCATTAAACGCT | LAMTOR2 | 28956 |
| GATTTCACAACAATGTTGCT | MAPK9 | 5601 |
| GATTTCTGGATCAGTTAGTG | IKZF5 | 64376 |
| GATTTGATCAGCTTCCTGTG | C16orf72 | 29035 |
| GCAAAAACCACACCAACTGG | IKZF5 | 64376 |
| GCAAAACTACCAGACTGGAG | MGA | 23269 |
| GCAAAAGTGGATAAGAAACC | LAMTOR3 | 8649 |
| GCAAAAGTGGCATAAAACCG | NO_CURRENT_605 | NO_CURRENT_605 |
| GCAAACCCGAGTGACACGTC | NO_CURRENT_606 | NO_CURRENT_606 |
| GCAAACTTGGTCGATGTAAG | ASH2L | 9070 |
| GCAAAGTCTCAGAATATACA | FBXW7 | 55294 |
| GCAAAGTCTGTAAAGTGTCC | CDKN2C | 1031 |
| GCAAAGTGCCTGCATTCGTG | PITPNB | 23760 |
| GCAAATCGGTCTGCTGCGCA | PLD6 | 201164 |
| GCAAATGCTGAGAAGACTTC | ATP5E | 514 |
| GCAACACTCACCATAGCCCT | VDAC1 | 7416 |
| GCAACCTACCTCCTATCATG | EIF2AK2 | 5610 |
| GCAACGTCTGCCATCACAGT | ASH2L | 9070 |
| GCAACGTGACGGTCACCCCA | TRERF1 | 55809 |

| | | |
|---|---|---|
| GCAACTCAGACAACAAGAGG | FBXW7 | 55294 |
| GCAACTCGCAGATGCCTACG | NOD1 | 10392 |
| GCAAGATATTATCGGCACAG | AIM2 | 9447 |
| GCAAGCATCCAAGTGAAGGG | TMEM173 | 340061 |
| GCAAGGGCTTGTCCGACATG | ARRB1 | 408 |
| GCAAGTCCCTGTAATAAGAT | DPY30 | 84661 |
| GCAAGTTCACCGTCCGAACA | STAT2 | 6773 |
| GCAATAAGGATGTGTTGGCT | C16orf72 | 29035 |
| GCAATCACGCCGGCACCAGG | TNFRSF6B | 8771 |
| GCAATCGTATCTCTCATAGG | NDUFB9 | 4715 |
| GCAATGCAATCGCAGGAGCA | NO_CURRENT_607 | NO_CURRENT_607 |
| GCAATGGAATACTGTTCTGG | CHUK | 1147 |
| GCAATTTGGAGCTAACTCCA | NO_CURRENT_608 | NO_CURRENT_608 |
| GCACAAACTGGATGTCGCTG | MYD88 | 4615 |
| GCACAACGCTTGCCATGTAG | NO_CURRENT_609 | NO_CURRENT_609 |
| GCACAATCTCCCTAACCCCG | SLC25A1 | 6576 |
| GCACACAAGTTAATCAACCC | PAXIP1 | 22976 |
| GCACACGGGTGACATCACTG | UBA7 | 7318 |
| GCACACGGTCTAGTTCTCAG | KMT2C | 58508 |
| GCACATATGAAGATTCACAT | ZNF699 | 374879 |
| GCACATGATGAGAAGGCCCA | MEMO1 | 51072 |
| GCACCAGGCGTATTGATGCA | NO_CURRENT_610 | NO_CURRENT_610 |
| GCACCATCCCCCATCCTACG | SERPINE1 | 5054 |
| GCACCGCCACCATTGCACCA | TRMT61A | 115708 |
| GCACGAGGTGAACAGCCGCT | NO_CURRENT_611 | NO_CURRENT_611 |
| GCACGATGTAGGGGCAGTCG | MAP2K7 | 5609 |
| GCACGCTGTACAGACGACAA | NO_CURRENT_612 | NO_CURRENT_612 |
| GCACGGACAGAAAATCTGAG | CCAR2 | 57805 |
| GCACTACTGTGCGGTACGCG | LTB4R | 1241 |
| GCACTGAAGTTCAATGGTGG | SOD2 | 6648 |
| GCACTGACATCAAAGCAGCC | CCL20 | 6364 |
| GCACTGAGCTCGGAGCTAAG | BAHD1 | 22893 |
| GCACTGCCCAGGCTTAAAGG | PDHB | 5162 |
| GCACTGCTAGTCACAAGGCT | POU2F1 | 5451 |
| GCACTTTATAGACCAGCACC | PYCARD | 29108 |
| GCACTTTGTTTGGCCTACTG | NO_CURRENT_613 | NO_CURRENT_613 |
| GCAGAAAAATCACTACAAAG | DNAJB6 | 10049 |
| GCAGAACCGAGGGTACATGC | NOS2 | 4843 |
| GCAGAATCTGCCAAAATAGG | IFNAR2 | 3455 |
| GCAGACTCTATTTGACAGAG | GC | 2638 |
| GCAGACTTTCTCAACTCGTT | BRK1 | 55845 |
| GCAGAGCAAGCCCAACACGG | SLC11A1 | 6556 |
| GCAGAGGCAATGTCATGCCA | PLD1 | 5337 |
| GCAGAGTGTGTCCTCTTCAG | ATP5J2 | 9551 |
| GCAGATGATCCAGATCCGCG | CYFIP1 | 23191 |
| GCAGATGGGATAGACACCAG | NR2C2 | 7182 |
| GCAGATTATTGACAGAGATG | CTNNBL1 | 56259 |
| GCAGATTGTCGCCATGCCTG | ACO2 | 50 |
| GCAGATTTATGGAGCTCACA | SPEN | 23013 |
| GCAGCAAATGAAGGATCAGC | IL10 | 3586 |

| | | |
|---|---|---|
| GCAGCAACATGGCAAGACGG | CS | 1431 |
| GCAGCAGTCTCTGATCGACG | BRD1 | 23774 |
| GCAGCCAATGAGCAAAACGC | RND1 | 27289 |
| GCAGCCTCCTTCTTATCACA | TRERF1 | 55809 |
| GCAGCGCAGTTCAGTGACCA | CXCL3 | 2921 |
| GCAGCGCCGTCTTGCCGAAG | CDKN2D | 1032 |
| GCAGCGCTTCAGACAGAACG | NLRP12 | 91662 |
| GCAGCGTCGACTTTACTCGC | LTA4H | 4048 |
| GCAGCTAGCGACCAGAAGCA | EIF4G3 | 8672 |
| GCAGGCACTAGAGCCGCTCG | AHDC1 | 27245 |
| GCAGGCACTCAGCAGAACCA | TMEM173 | 340061 |
| GCAGGCAGCACAGATCAACA | NLRP1 | 22861 |
| GCAGGCGCAGCAAAGTCGAT | NNT | 23530 |
| GCAGGTAGACAATGCGACAT | APEH | 327 |
| GCAGGTGCCACCGCGCCGTG | PTGES2 | 80142 |
| GCAGGTTGAAGGCTTTCAGG | UBE2L6 | 9246 |
| GCAGTGAAGGGCATAGGATC | LGMN | 5641 |
| GCAGTGCTTTCCATCACCAC | NQO1 | 1728 |
| GCAGTTCCTTTAAGGCATTG | SATB2 | 23314 |
| GCAGTTGCTGCACCAGCGTG | PLD6 | 201164 |
| GCATAAAATATGAGCTCCGT | MOSPD1 | 56180 |
| GCATAAATACCCCTTCCGAG | NO_CURRENT_614 | NO_CURRENT_614 |
| GCATAACCCGACGAGTAGCA | LRP2 | 4036 |
| GCATACAACTCGGATTGCTC | PPARGC1A | 10891 |
| GCATACAGGCTCACGTCCAG | LARP4B | 23185 |
| GCATACATAGAAGATCACAG | CHEK2 | 11200 |
| GCATATATTTAATGCCAAGA | PHF6 | 84295 |
| GCATATCTAGGTCCTAAAGG | KMT2C | 58508 |
| GCATATGAAGTGCTGTCGGA | DNAJB6 | 10049 |
| GCATATGTCTCACCTCAATC | CHMP5 | 51510 |
| GCATCACAAGTTTGCCATCG | PLD6 | 201164 |
| GCATCACAGTCATTGCTGCT | BMI1 | 648 |
| GCATCACAGTCATTGCTGCT | COMMD3-BMI1 | 100532731 |
| GCATCAGCTCCACTTTATCG | MPC2 | 25874 |
| GCATCCTAGAGGAATTGCCG | LRP2 | 4036 |
| GCATCTGCCATACCTTCCAG | HCAR2 | 338442 |
| GCATCTGTCACGTAAGACAG | SATB1 | 6304 |
| GCATGATCTCCGAGTCCCTG | SERF2 | 10169 |
| GCATGGTCACCGGTCTGCTG | TMEM173 | 340061 |
| GCATGTTAGGCAGGTTGCCT | IL10 | 3586 |
| GCATTTATAAGCTTTCTCGC | ZNF616 | 90317 |
| GCATTTGACACGACATTGGG | HSPA4 | 3308 |
| GCCAAGCTGGATCTTGATGC | ATP5J2 | 9551 |
| GCCAAGCTGGATCTTGATGC | ATP5J2-PTCD1 | 100526740 |
| GCCAATCCGAACTTTATGCA | NCOA6 | 23054 |
| GCCACAATCCTGGCCCACGC | SUPT7L | 9913 |
| GCCACACGAATCATAAAGAG | NO_CURRENT_615 | NO_CURRENT_615 |
| GCCACACGTGGCCATCGAAG | NMRK2 | 27231 |
| GCCACCGCAAAGTTGTCCAG | HTR3E | 285242 |
| GCCACGACACCACAGCCAGT | CYP4A11 | 1579 |

| | | |
|---|---|---|
| GCCACGACACCACAGCCAGT | CYP4A22 | 284541 |
| GCCACTACCACCAAATGACG | DNAJB6 | 10049 |
| GCCACTGATCGGAAACGCGG | CYP1B1 | 1545 |
| GCCAGAACGCAGTGTCCCGA | KIAA1211L | 343990 |
| GCCAGCACCAAGCATCTGAG | CYP8B1 | 1582 |
| GCCAGGGTATGGGCATCTCG | NO_CURRENT_616 | NO_CURRENT_616 |
| GCCAGGGTTCTTGGTCCCGA | NO_CURRENT_617 | NO_CURRENT_617 |
| GCCAGTTTCATTGTCAACGG | NCKAP1L | 3071 |
| GCCATAATATATGAGTAACT | NO_CURRENT_618 | NO_CURRENT_618 |
| GCCATCAAATTTGTACTCAG | MCTS1 | 28985 |
| GCCATCGAATCCGGGCCAGT | KIAA1211L | 343990 |
| GCCATGGACAATTCTGCTGG | CBLL1 | 79872 |
| GCCATGGAGTACTGCCAAGG | IKBKB | 3551 |
| GCCATTCATGATCACCACAT | ACACA | 31 |
| GCCATTCTAGTCCCGGCATA | NO_CURRENT_619 | NO_CURRENT_619 |
| GCCATTCTGGTACACCAAAG | WASF2 | 10163 |
| GCCCAAGAGTTGCGGCGTAT | BCL2L11 | 10018 |
| GCCCAAGTGTCCAACAACAA | NO_CURRENT_620 | NO_CURRENT_620 |
| GCCCACAGCCTGTTTGAGAG | BAK1 | 578 |
| GCCCACATCGAGCACACGCA | TLR9 | 54106 |
| GCCCAGAAGAAAACTGTTCT | MPC2 | 25874 |
| GCCCAGACGCCCTAGAATAG | NO_CURRENT_621 | NO_CURRENT_621 |
| GCCCATGGAAGTTTAAAATG | NO_CURRENT_622 | NO_CURRENT_622 |
| GCCCCCGGACCCCTCACGCT | PAGR1 | 79447 |
| GCCCCGTAAATCTCATTACA | NO_CURRENT_623 | NO_CURRENT_623 |
| GCCGAGGATGAACTAAGTAG | KMT2C | 58508 |
| GCCGCAAAAGGGCAGAAAGG | NUDT17 | 200035 |
| GCCGCCGACAAGCAGTACAA | SLC25A6 | 293 |
| GCCGCCGATTTCATAAGTAA | NO_CURRENT_624 | NO_CURRENT_624 |
| GCCGCCGGGAAGAAGCTCCA | CEBPD | 1052 |
| GCCGCTATTACTGCAATCAC | SEC62 | 7095 |
| GCCGCTTACAAACACGCAGA | SNRNP70 | 6625 |
| GCCGGCTAACGGCAATCTGT | SATB2 | 23314 |
| GCCGGGGACGGAGTGTGCTG | SIAH2 | 6478 |
| GCCGGGGGGTCTTTGACCGG | SUV39H1 | 6839 |
| GCCGGTGGTAGGTGGCCCGC | MPC2 | 25874 |
| GCCGGTGTGTCAGACAACTT | INO80E | 283899 |
| GCCGTGGCCCAGCTCTACAA | C16orf72 | 29035 |
| GCCGTGGTATCAAGTCGGTA | NO_CURRENT_625 | NO_CURRENT_625 |
| GCCTAAAAAGGCCCCTGCAA | HMGN2 | 3151 |
| GCCTAGATGTTCGTTCTGAG | NO_CURRENT_626 | NO_CURRENT_626 |
| GCCTATCGGCATTCCCACTG | NO_CURRENT_627 | NO_CURRENT_627 |
| GCCTCACCGTTAGACCAGAA | MGA | 23269 |
| GCCTCCAGGATCCTCTCCAC | RXRA | 6256 |
| GCCTCGGGTGATCTACGCGA | SLC7A3 | 84889 |
| GCCTGAACAGAAACAGATGG | AIM2 | 9447 |
| GCCTGCGGGCCACCTACCAC | MPC2 | 25874 |
| GCCTGGCACTGCAGAATAGG | SIAH2 | 6478 |
| GCCTGGGTTTTGGTGCATAC | NO_CURRENT_628 | NO_CURRENT_628 |
| GCGACTACTGCACAAGCTAG | ARR3 | 407 |

| | | |
|---|---|---|
| GCGAGATGATACAATATACG | UQCRB | 7381 |
| GCGAGCCCTTACATTGACAG | NUP153 | 9972 |
| GCGAGCTCCATGTAAAAGAG | SLC6A4 | 6532 |
| GCGAGGTCAAGGTGCGCAAG | TNRC18 | 84629 |
| GCGAGTTGCAGATCTACACT | SAP18 | 10284 |
| GCGATCGGAGTGCCACGATA | NO_CURRENT_629 | NO_CURRENT_629 |
| GCGCACAAATCCCTTCTACC | MMP1 | 4312 |
| GCGCAGAAGTGGTGGCACAC | SLC3A2 | 6520 |
| GCGCCAGCTACAGGTGCAGT | EBI3 | 10148 |
| GCGCCCCAAGGCTTTGACCC | LAMTOR2 | 28956 |
| GCGCCGGGGTCTTCTCCACA | ATP5L | 10632 |
| GCGCCGGTTACACTGTTGCT | NR1H3 | 10062 |
| GCGCTCACGGAGTCGCGACA | SNRNP70 | 6625 |
| GCGCTGCCGGGAGATCGAGC | TFPT | 29844 |
| GCGCTTTCACTGGACCGCGC | PDHB | 5162 |
| GCGCTTTGAGACTCCGGTAG | JUNB | 3726 |
| GCGGAACACACGCAGCCCAG | NOD1 | 10392 |
| GCGGAATGAACACGAGGTAG | NO_CURRENT_630 | NO_CURRENT_630 |
| GCGGCACAAGTGGCATCCAG | RHOG | 391 |
| GCGGCCCCACACCACGATGG | ADRB1 | 153 |
| GCGGCCGGGGACTCGCACGG | CYP1B1 | 1545 |
| GCGGCGAGGTCCTGGCGACC | BCL2 | 596 |
| GCGGCGGCGATTGGGTACCG | SIRT1 | 23411 |
| GCGGCGGGAGAAGTCGTCGC | BCL2 | 596 |
| GCGGCGTCTGGGAATCGTTC | NO_CURRENT_631 | NO_CURRENT_631 |
| GCGGCTGCTTACTGCTCGAC | INO80B | 83444 |
| GCGGCTGTACAAGTCCACCA | NMRK2 | 27231 |
| GCGGGAGATTCACCAGGACT | BRK1 | 55845 |
| GCGGGCGAGACAGACGCCTC | RRAGA | 10670 |
| GCGGGGAATCCTGCGACAGA | ARHGAP33 | 115703 |
| GCGGGGACGCTTTGCCACGG | BCL2 | 596 |
| GCGGGTACGCGCTCTCGGTG | CYP24A1 | 1591 |
| GCGGTCCCCACAGCAAACCC | ACADS | 35 |
| GCGGTGACATTGAGCACGTA | EBI3 | 10148 |
| GCGGTGGAGTACGCCGTGCG | GPT2 | 84706 |
| GCGTAACCTCGTCTTCCAAG | IRF8 | 3394 |
| GCGTAGCTGATGGAGCCTGT | INO80 | 54617 |
| GCGTCTTCACGTACTCGGTG | SLC25A1 | 6576 |
| GCGTGCGTCCCGGGTTACCC | NO_CURRENT_632 | NO_CURRENT_632 |
| GCGTTCAAATAGCCTCACTT | NO_CURRENT_633 | NO_CURRENT_633 |
| GCGTTCCCCCACTGACGGGG | NO_CURRENT_634 | NO_CURRENT_634 |
| GCTAACGTGCTGCGCGACAT | PYCARD | 29108 |
| GCTACAGCATGGTGGCCTAC | ATP5E | 514 |
| GCTACGTATTTCAAAAAACG | NO_CURRENT_635 | NO_CURRENT_635 |
| GCTACTCGGACATAGGGCGT | LTB4R | 1241 |
| GCTACTGTGTCACAAACGGG | MSL2 | 55167 |
| GCTACTTGAAGATCATGAGT | GPR119 | 139760 |
| GCTAGTAAGAATGAGACTGG | PITPNB | 23760 |
| GCTATTTGAGTAGTTCACTG | NO_CURRENT_636 | NO_CURRENT_636 |
| GCTCAAGAAGAAGCGATACC | CHMP6 | 79643 |

| | | |
|---|---|---|
| GCTCACCCTGAGCTGCAAAG | SLC25A19 | 60386 |
| GCTCACCTGCTAGGTTGCAG | BAK1 | 578 |
| GCTCATCACCACGAGACCTG | NLRP3 | 114548 |
| GCTCATGTACAACGATGAGT | PDE4A | 5141 |
| GCTCCAGAGGAATGCTCCTG | ZNF641 | 121274 |
| GCTCCATGAAAGAATAATTG | NO_CURRENT_637 | NO_CURRENT_637 |
| GCTCCCAGTGAATTAGAACG | C6orf15 | 29113 |
| GCTCCGGGAGCGAGATGAAG | TFPT | 29844 |
| GCTCCTTGTACAGATGCCGG | SERPINE1 | 5054 |
| GCTCGCAAGTATTTAAGGAC | NO_CURRENT_638 | NO_CURRENT_638 |
| GCTCGCAATAACATGCAGTC | NDUFS4 | 4724 |
| GCTCGCTCGCACCTATTGCT | MAP2K5 | 5607 |
| GCTCGGGCAACGCATCCATG | TNRC18 | 84629 |
| GCTCTACCTCACCCAACTGT | ASH2L | 9070 |
| GCTCTAGAGTTTCGTGGTGG | C16orf72 | 29035 |
| GCTCTCAAGAGAATCATCAC | EIF2AK2 | 5610 |
| GCTCTCGTCGTCGTACATGG | CEBPD | 1052 |
| GCTCTGGTCATCCTGGCTAA | PARK7 | 11315 |
| GCTCTGGTGCTAGGTTACGA | VDAC1 | 7416 |
| GCTCTGTGGTGGGATCCAGT | UBA7 | 7318 |
| GCTGAAAGTCATGTACACCG | SCTR | 6344 |
| GCTGAAAGTTGGGTTCACCT | NDUFC1 | 4717 |
| GCTGAACAGGACGTCCACAT | ARHGAP33 | 115703 |
| GCTGAACGTTGGGAATCAGC | RRAGA | 10670 |
| GCTGAAGGCACACTTCATGA | CYP4A11 | 1579 |
| GCTGAAGTCAAGTGCCATTG | VDR | 7421 |
| GCTGAAGTGCAGTTCTACCC | BNIP3 | 664 |
| GCTGAATCAACTGAATCGTA | SUPT7L | 9913 |
| GCTGAATGTCATCCTCCCGT | STAG2 | 10735 |
| GCTGACTGATACACTCCAAG | FOS | 2353 |
| GCTGAGCCCTCGCTTCCGTG | NDUFS8 | 4728 |
| GCTGAGTTCACCACGCCCGA | SERPINE1 | 5054 |
| GCTGATATATACGACAAGCC | NO_CURRENT_639 | NO_CURRENT_639 |
| GCTGATGAAGCTGTCACTCG | HSPA4 | 3308 |
| GCTGATGGCAGAGATTGGTG | CFLAR | 8837 |
| GCTGATGTCAACGTGCCTGA | CDKN2D | 1032 |
| GCTGATTGAATCACTACTGG | SPEN | 23013 |
| GCTGCAAATTCTATTTGTGT | NO_CURRENT_640 | NO_CURRENT_640 |
| GCTGCAATCCTTACTGTTTG | RAC1 | 5879 |
| GCTGCACAAAGGTGCCTGGG | TNFRSF6B | 8771 |
| GCTGCACTACGCTTTCCAGA | RPS6KA4 | 8986 |
| GCTGCAGCGTCACCATTGAG | UBE2H | 7328 |
| GCTGCAGGATTCAATAAACT | UQCRB | 7381 |
| GCTGCAGGCTGAGAAGCAGA | PDAP1 | 11333 |
| GCTGCCCACAGATCGAGGTG | NUDT17 | 200035 |
| GCTGCGCAAATATCCTGGCG | BRD1 | 23774 |
| GCTGCGCCGGTACCCATCCG | NDUFS8 | 4728 |
| GCTGCTCCATGTCTTCGCCA | RND3 | 390 |
| GCTGCTGCGGGACGGCAGGA | CHMP6 | 79643 |
| GCTGCTGCTCCTGCTCCTGG | CXCL2 | 2920 |

| | | |
|---|---|---|
| GCTGCTGCTCCTGCTCCTGG | CXCL3 | 2921 |
| GCTGCTGGAAAAGTTCCGAG | KAT2A | 2648 |
| GCTGCTGGTAGTCATAATAC | TRERF1 | 55809 |
| GCTGGAAAGTGACCTCAAAG | TP73 | 7161 |
| GCTGGACAACAAAATCAATG | CYP24A1 | 1591 |
| GCTGGACATTGGACTTCCTC | BAX | 581 |
| GCTGGACGGTACATTTCCCC | SAP18 | 10284 |
| GCTGGAGAACCTGACCGCCG | PYCARD | 29108 |
| GCTGGAGAGACAATTCTACT | NO_CURRENT_641 | NO_CURRENT_641 |
| GCTGGCCCGCTATGGCGGGG | NENF | 29937 |
| GCTGGGAAAAGAACTTCGAG | IL2RB | 3560 |
| GCTGGGACTGCTGTTAAACG | TMEM173 | 340061 |
| GCTGGGCAGGGAATTTACAG | NCOA6 | 23054 |
| GCTGGGCTACCTAAATAAAA | TIFA | 92610 |
| GCTGGGCTTAGGGCCACCAG | METTL3 | 56339 |
| GCTGGTACACAGCCTCAGGT | MEMO1 | 51072 |
| GCTGGTGGGTGCGTGCATCC | CSNK1D | 1453 |
| GCTGGTTCATATCTTGAACA | IKBKB | 3551 |
| GCTGTAATTGTCGAACACGG | RHOG | 391 |
| GCTGTATATTCTCCTCCCAT | TMEM173 | 340061 |
| GCTGTCCACTGCTGAGACAG | ADRB1 | 153 |
| GCTGTCCCCAGGCCACACCT | IL2RB | 3560 |
| GCTGTGAGGAGACTGTTCGT | SATB2 | 23314 |
| GCTGTGCTAGTGCAGCCCTG | PIGY | 84992 |
| GCTGTTATCTGATGATGCAG | INO80E | 283899 |
| GCTGTTTCCAGTCCTAACCA | UTY | 7404 |
| GCTGTTTGGACCATATCAGT | HTR2A | 3356 |
| GCTTAAGAACTTTCGTCATG | FBXO38 | 81545 |
| GCTTAAGTCACGGCTTTCCA | NO_CURRENT_642 | NO_CURRENT_642 |
| GCTTACACGATTGTGAAGTC | NO_CURRENT_643 | NO_CURRENT_643 |
| GCTTACTTGTTCTACAGTAG | PITPNB | 23760 |
| GCTTCACCCCAGGATTACAA | SPCS1 | 28972 |
| GCTTCATGCTCTCATGCATG | GNAI2 | 2771 |
| GCTTCCCTTGATCTGACTGG | FOS | 2353 |
| GCTTCCTACCTCATAGCCAA | SLC11A1 | 6556 |
| GCTTCCTTGGTCTGTTGCAA | LAMTOR3 | 8649 |
| GCTTCTTTCGAGGGACACTG | ARHGAP33 | 115703 |
| GCTTGAGGTTCACAGTAGGG | TLR1 | 7096 |
| GCTTGCATAATTCTGGCCAG | NO_CURRENT_644 | NO_CURRENT_644 |
| GCTTGCTATATGGGTGCGAG | NO_CURRENT_645 | NO_CURRENT_645 |
| GCTTGGTCAAGAATATGAGC | DYRK1A | 1859 |
| GCTTGTAGACAAAACAACGT | NO_CURRENT_646 | NO_CURRENT_646 |
| GCTTGTATTCTTCTGGTCGG | UBE2H | 7328 |
| GCTTTGGCCCCGCGTCCCCG | SBNO2 | 22904 |
| GCTTTGGTTACGTTCTATGG | PAXIP1 | 22976 |
| GCTTTTTCCAGCGAGAGCAA | NO_CURRENT_647 | NO_CURRENT_647 |
| GGAAAAACTAGAGTTCATCC | FOS | 2353 |
| GGAAAAGTTCAAGAAACCCG | NUP153 | 9972 |
| GGAAACAGATCAGAATGCAA | HSPA4 | 3308 |
| GGAAACTATTCACCAGTTAG | TRAF6 | 7189 |

| | | |
|---|---|---|
| GGAAACTCAAGCCTGCACTC | TNFRSF1B | 7133 |
| GGAAACTTGAGTTTACCGTG | NO_CURRENT_648 | NO_CURRENT_648 |
| GGAAAGGGGACGCAGCAAGG | NDUFC1 | 4717 |
| GGAAAGTAATGGACCAGTGA | SOD1 | 6647 |
| GGAAATGACAGTATGAGTCG | NO_CURRENT_649 | NO_CURRENT_649 |
| GGAACAATGACTGGAGTGTG | EIF4G3 | 8672 |
| GGAACAGAAACGGCTACTGA | MAP2K6 | 5608 |
| GGAACCACTGAAGCGGACAG | UBA7 | 7318 |
| GGAACGAGGCAGTGACAGGG | NO_CURRENT_650 | NO_CURRENT_650 |
| GGAACGCTTGAAGGAAATGG | RBM42 | 79171 |
| GGAACTCTCACTGGACGACG | SETD1B | 23067 |
| GGAACTCTGAGTCATAGCGT | BAK1 | 578 |
| GGAACTTGCCTGCTTCGAGG | LARP4B | 23185 |
| GGAAGAAATATGGAGACAAG | PDHB | 5162 |
| GGAAGAATCCTCTGTACTTG | SLC25A1 | 6576 |
| GGAAGACGGCTTCAAACAGT | NMRK2 | 27231 |
| GGAAGAGCTCGGAGTCATCG | PPARGC1B | 133522 |
| GGAAGAGGATACAGTCACTC | JAM2 | 58494 |
| GGAAGATGCAGAGGTCATAT | SUV39H1 | 6839 |
| GGAAGCATCATAGGACTATG | NO_CURRENT_651 | NO_CURRENT_651 |
| GGAAGCCCTTCGATGTGCAA | NR2C2 | 7182 |
| GGAAGCGAATCAATGGACTC | CASP3 | 836 |
| GGAAGCTATGGGCCTTCACG | SLC11A1 | 6556 |
| GGAAGCTTTATTTACAACCA | SEC62 | 7095 |
| GGAAGGCAAACACCTCCTCG | NOD1 | 10392 |
| GGAAGGGTTCTTACTAGAGA | PPARGC1A | 10891 |
| GGAATAAGAACAGTCAACGT | PIGY | 84992 |
| GGAATGTCCTAGGTTACTGA | NO_CURRENT_652 | NO_CURRENT_652 |
| GGAATTACGACTAACCGATT | NO_CURRENT_653 | NO_CURRENT_653 |
| GGAATTTCAACGTGGCCCAT | TMEM173 | 340061 |
| GGACAACCGGAACCTGTTCG | CACNA2D2 | 9254 |
| GGACACCATCCAGGGAGATG | BTN2A2 | 10385 |
| GGACACCCTGAAGGTCCTAG | CDKN2D | 1032 |
| GGACACTACAGTCAGTCTTG | C16orf72 | 29035 |
| GGACACTATAGCGCCGCCTG | TIRAP | 114609 |
| GGACAGGTAAGGGTCGGAAA | CS | 1431 |
| GGACATGACTACAGAGCTGG | CASP1 | 834 |
| GGACATGGAGAAAGAAGTGT | HTR3E | 285242 |
| GGACCAACTCCAAACATGCA | NCOA6 | 23054 |
| GGACCCACACTCCTACGACG | PTGIS | 5740 |
| GGACCCGCGTAAAGACCACG | GNAI2 | 2771 |
| GGACCCTTAGGGATGAACCA | ATP6V0E1 | 8992 |
| GGACCGCATGGTGACTATCG | RBM6 | 10180 |
| GGACCTGCCATACAGGTGCA | TIFA | 92610 |
| GGACGACCACCTAAGAACAC | MGA | 23269 |
| GGACGCACCATTCCGGGTGA | NO_CURRENT_654 | NO_CURRENT_654 |
| GGACGGCTATTTCCGCCTGA | TYK2 | 7297 |
| GGACGGTCAGGAAGTACCGG | NMRK2 | 27231 |
| GGACGTTCTCATAAGAGGCT | RAC2 | 5880 |
| GGACGTTGGCAGTAGTAAGG | RBM6 | 10180 |

| | | |
|---|---|---|
| GGACTACAACGTCATGGTGA | CSNK1D | 1453 |
| GGACTATCATGTGTAGAACT | PRR14L | 253143 |
| GGACTATCCACCGTTTACTC | NO_CURRENT_655 | NO_CURRENT_655 |
| GGACTCCAAACCACTGCAGG | NQO1 | 1728 |
| GGACTTCTTTGGGAATCGAA | DNAJB6 | 10049 |
| GGAGAAACAGGTCCCAGATA | MBNL1 | 4154 |
| GGAGAAAGACAAGCAGATCG | MKL1 | 57591 |
| GGAGAACAAGCGCATCCTCA | MAP2K7 | 5609 |
| GGAGAACAGGCAATTAAAGA | DNAJA1 | 3301 |
| GGAGAAGCTGGAGTTCATGT | FOSL2 | 2355 |
| GGAGAAGTTTGCCCTAGTGG | CYFIP1 | 23191 |
| GGAGACTTAATAGAACCGTA | CUBN | 8029 |
| GGAGAGGAAAATCGGCACAG | NO_CURRENT_656 | NO_CURRENT_656 |
| GGAGAGTCAGGGAAGATGTA | NDUFB9 | 4715 |
| GGAGATCGACGCGCAGCTGC | PDAP1 | 11333 |
| GGAGATGCGGCCTTCTCAAA | NO_CURRENT_657 | NO_CURRENT_657 |
| GGAGCAAATATCATGTGTCC | MCTS1 | 28985 |
| GGAGCAGTCCATTCAGACAT | IRF9 | 10379 |
| GGAGCATGTTAACACCTGCG | NO_CURRENT_658 | NO_CURRENT_658 |
| GGAGCCCCAGGGGTGGACGG | EBI3 | 10148 |
| GGAGCGAGATCGCATCTACC | SATB2 | 23314 |
| GGAGCGTGAAGTAAAAGCAT | EIF2AK2 | 5610 |
| GGAGCTGAGGTTGACATCAC | CASP1 | 834 |
| GGAGGAATTAGCTTGAAGCA | APIP | 51074 |
| GGAGGACACACAACAAAACC | NO_CURRENT_659 | NO_CURRENT_659 |
| GGAGGAGATGTCGACATCTG | PHC3 | 80012 |
| GGAGGATACTCACGGCCGCT | NDUFC1 | 4717 |
| GGAGGCGGGATATATACCAG | BCORL1 | 63035 |
| GGAGGGTGATCATTAAGTAG | NO_CURRENT_660 | NO_CURRENT_660 |
| GGAGGTGAACCCTGTGCGCA | PTGES2 | 80142 |
| GGAGGTGTCTTAAACATGGT | NO_CURRENT_661 | NO_CURRENT_661 |
| GGAGGTTCACACACCTCTGT | TLR2 | 7097 |
| GGAGTAATGGCGATGATGCA | PIAS1 | 8554 |
| GGAGTCCCGAAGGACTACAC | LGMN | 5641 |
| GGAGTCGCGCGTTACCCAGG | SAP18 | 10284 |
| GGAGTGAAGCCTTGGGTGAG | C6orf15 | 29113 |
| GGAGTGCCTTTAAAACACTC | SATB1 | 6304 |
| GGAGTGTTCCACGTCAATGG | RRAGA | 10670 |
| GGATAAGAAAGAGACACAAG | TNRC18 | 84629 |
| GGATACTCTCGAGCTCTCCG | PTGIS | 5740 |
| GGATATTGAGTAAACCCGAT | NO_CURRENT_662 | NO_CURRENT_662 |
| GGATATTTCCATGAGGACGG | MAP2K6 | 5608 |
| GGATCCACCTCAACAACATG | TRERF1 | 55809 |
| GGATCCATAACCTCGACCCG | SPHK1 | 8877 |
| GGATCTGCGCCGTTTCTGTG | NFKB2 | 4791 |
| GGATCTGTCACGGAACAACC | TLR9 | 54106 |
| GGATCTTCTTGTAATCGCAC | SUV39H1 | 6839 |
| GGATGAAGAAGATGACTACC | PDAP1 | 11333 |
| GGATGCCAGCGACTTTGACT | PPARGC1B | 133522 |
| GGATGTGGGGAACATTGTGC | SLC7A5 | 8140 |

| | | |
|---|---|---|
| GGATGTGTCCAGCGTGAATG | NLRC4 | 58484 |
| GGATGTTGCTGAGCCCTACA | IRF9 | 10379 |
| GGATTACCTGATGTACAGTG | ZNF616 | 90317 |
| GGATTCATACGACGTGACTG | ALOX5 | 240 |
| GGATTCATGGCGAGTCCGCA | NO_CURRENT_663 | NO_CURRENT_663 |
| GGATTCCTAAGAACGGTACT | FBXO38 | 81545 |
| GGATTGAATGGCTAACGCGG | NO_CURRENT_664 | NO_CURRENT_664 |
| GGATTTACAAGACCAGCTAG | CHMP5 | 51510 |
| GGATTTGTCGCTTGCCACAC | NO_CURRENT_665 | NO_CURRENT_665 |
| GGCAACCACAGCGCAGACGG | HELZ2 | 85441 |
| GGCAACCTTTCAGTTCACTG | USP7 | 7874 |
| GGCAACGCTCACCTTGACCC | PRKN | 5071 |
| GGCAAGTTAAAATCATTGCT | CD84 | 8832 |
| GGCACAAGAGATGTCCCTCT | DLAT | 1737 |
| GGCACAGAGCGCGAGTTCCG | SLC25A6 | 293 |
| GGCACCACAAATTGCACAGT | IRAK4 | 51135 |
| GGCACCAGGTAGTCCACCAT | NFKB1 | 4790 |
| GGCACCGCTAAGAAGAATGG | RIPK1 | 8737 |
| GGCAGAAGCAGTAACAAGGG | BCORL1 | 63035 |
| GGCAGACCACCATCCAAGTG | MEX3B | 84206 |
| GGCAGAGCCTGTGATGAGGT | HTR1D | 3352 |
| GGCAGCAACTGAAGCAGCAG | CHMP6 | 79643 |
| GGCAGCAATGAACATGCTGA | SERPINE1 | 5054 |
| GGCAGCAGCTCAACTTCTGG | SIAH2 | 6478 |
| GGCAGCGCACCATAAGTGCA | PHF6 | 84295 |
| GGCAGCGGCAGCATAGGGGA | INO80B | 83444 |
| GGCAGCGTTACCCAGTCCCA | CHEK2 | 11200 |
| GGCAGCTACAGAGCTCGCGG | BABAM1 | 29086 |
| GGCAGGAGCTAATGTTTCGG | IKBKE | 9641 |
| GGCAGGCAGTCAGGCCCAAG | ARMCX2 | 9823 |
| GGCAGGCCTGATCTTCGTGG | ARF1 | 375 |
| GGCAGGTACCTGGTTCGCTG | ACADS | 35 |
| GGCAGGTAGGGTATACATGG | NO_CURRENT_666 | NO_CURRENT_666 |
| GGCAGTAGAACAGCTCGATG | USP7 | 7874 |
| GGCAGTGCTGCCTTCCAAGT | TIRAP | 114609 |
| GGCATCCTCACGGTGAAAGG | ACO2 | 50 |
| GGCATCGGACACACTAATAG | NO_CURRENT_667 | NO_CURRENT_667 |
| GGCATGATTTGATTAAGCCA | MCTS1 | 28985 |
| GGCATGTCCAAAAAGCCAGT | CYP4B1 | 1580 |
| GGCCAAGCCCAAGCACCGCG | AKT1 | 207 |
| GGCCAATAACTGAGGCAGGA | UBE2H | 7328 |
| GGCCACCCAGAAGTACAGTG | IDH2 | 3418 |
| GGCCACGACAAGTCAGACCG | PQBP1 | 10084 |
| GGCCAGCAGACCTTTACTAA | NO_CURRENT_668 | NO_CURRENT_668 |
| GGCCATGACACCACCACCAG | CYP4B1 | 1580 |
| GGCCCATGTAATCCTTCATG | CFLAR | 8837 |
| GGCCCCAAAACGGTAGACCT | SLC5A8 | 160728 |
| GGCCCTCTAGAAAAGTCTCG | NO_CURRENT_669 | NO_CURRENT_669 |
| GGCCGGCTCCTGAAGTGTTG | BABAM1 | 29086 |
| GGCCGTCGTATTCCCCCAAG | NO_CURRENT_670 | NO_CURRENT_670 |

| | | |
|---|---|---|
| GGCCTGTCCACATTTCAGGG | RIPK3 | 11035 |
| GGCCTTAAACAGTGTACGCA | CXCR1 | 3577 |
| GGCCTTCTTTGAGTTCGGTG | BCL2 | 596 |
| GGCGAAGAGGACGCCCTCTG | SBNO2 | 22904 |
| GGCGCATTAAAGTCGAGAGC | NO_CURRENT_671 | NO_CURRENT_671 |
| GGCGCCTGTGCCATGCCGAG | CYFIP1 | 23191 |
| GGCGGACTATGTCCGAAGCA | MPC1 | 51660 |
| GGCGGAGCGCAGGTGACCCA | IZUMO2 | 126123 |
| GGCGGATTACAACACTGCCA | IKBKE | 9641 |
| GGCGGCCATGGAGTCGATGT | CEBPD | 1052 |
| GGCGTCAAAATTAGAAGCCG | NO_CURRENT_672 | NO_CURRENT_672 |
| GGCGTGTGCTTGTTGATTCC | NDUFA1 | 4694 |
| GGCGTTAATTAAACTGTTTT | NO_CURRENT_673 | NO_CURRENT_673 |
| GGCGTTGTGAAGACCATGAC | FOS | 2353 |
| GGCTACTATTCCCGTCTTGA | TRIR | 79002 |
| GGCTATGGAGCTGTCCAAGT | SLC25A19 | 60386 |
| GGCTCACAGTGTCATGGGTG | NENF | 29937 |
| GGCTCACGCCAAAGCCACTG | TRMT61A | 115708 |
| GGCTCAGGGTCATTGAGGAG | FOS | 2353 |
| GGCTCGCGCTTGAGCAGGCG | CEBPD | 1052 |
| GGCTCTGGGGCTCACGGACG | NO_CURRENT_674 | NO_CURRENT_674 |
| GGCTGAGAAGCCTTTACAGG | MYD88 | 4615 |
| GGCTGATTATAGTGCTCATG | CBLL1 | 79872 |
| GGCTGGCATGAACGCCCTGG | TNRC18 | 84629 |
| GGCTGGGAAGAAGCCAACGC | GPR119 | 139760 |
| GGCTGGGATCAGGTCTGAGG | LCN2 | 3934 |
| GGCTGGTTGACGACTCCTGA | NO_CURRENT_675 | NO_CURRENT_675 |
| GGCTTACGTGGGGGGCAAAA | NO_CURRENT_676 | NO_CURRENT_676 |
| GGCTTCAGTGTCTCCGTGCA | SLC25A6 | 293 |
| GGCTTCCAGGCGCACTACCG | SQSTM1 | 8878 |
| GGCTTCGTCCAGATCGGGAA | SLC7A5 | 8140 |
| GGCTTGAACGGATACTCCGG | UBE2L6 | 9246 |
| GGCTTGTATACTGCCTCGCC | PDAP1 | 11333 |
| GGGAAACTAGACATATGTAC | NO_CURRENT_677 | NO_CURRENT_677 |
| GGGAAGCGAATCCGGAAGCT | RIPK1 | 8737 |
| GGGAATATAAGGAGCGCACA | BRD1 | 23774 |
| GGGAATGCTACCAAAGCCTC | NMU | 10874 |
| GGGAATTACCAGCACCTACG | SLC5A8 | 160728 |
| GGGACATCCTTGCCGTCTCA | NO_CURRENT_678 | NO_CURRENT_678 |
| GGGACCATCTACCCTCAATG | NO_CURRENT_679 | NO_CURRENT_679 |
| GGGACCATGTATAGGAGCGA | SLC25A19 | 60386 |
| GGGACCTTACATTCACACTT | CXCL3 | 2921 |
| GGGACGCATTTGACATCCAG | TBXAS1 | 6916 |
| GGGACGCGAAAGAAACCAGT | NO_CURRENT_680 | NO_CURRENT_680 |
| GGGACGCTAACGATATCCTG | NO_CURRENT_681 | NO_CURRENT_681 |
| GGGACTGATATATGGCGAAC | NO_CURRENT_682 | NO_CURRENT_682 |
| GGGAGAACTCGTCCATTCGG | SAP18 | 10284 |
| GGGAGATAAAGCAAAGGTGA | HMGN2 | 3151 |
| GGGAGATCTTGAGCGCCTGG | HTR1D | 3352 |
| GGGAGCCCCAGCATGTGGCG | MAP2K7 | 5609 |

| | | |
|---|---|---|
| GGGAGCTTGAATTGAAACGT | EP400 | 57634 |
| GGGAGGCCGGCAGGACTCGG | PCGF6 | 84108 |
| GGGAGGTCCCAAATCCCATG | MTOR | 2475 |
| GGGAGGTGGCTTTAGGTTTT | NO_CURRENT_683 | NO_CURRENT_683 |
| GGGAGTAAAGGAAATGAAGA | CHMP5 | 51510 |
| GGGAGTAGTTTGTGCCATGA | CSDE1 | 7812 |
| GGGAGTATGTAGGAGACTTG | KPNA1 | 3836 |
| GGGAGTTGATTGTTTCGAGA | NO_CURRENT_684 | NO_CURRENT_684 |
| GGGATCAATTGTTCATGAGT | NO_CURRENT_685 | NO_CURRENT_685 |
| GGGATGCGTCTTGCTAAACC | NO_CURRENT_686 | NO_CURRENT_686 |
| GGGATTCCCAAGGTTGCCTA | ACTR5 | 79913 |
| GGGATTGGTTGAGCAAAGCG | NO_CURRENT_687 | NO_CURRENT_687 |
| GGGCAAGGTAATGTAGACAG | NLRP1 | 22861 |
| GGGCAATAGAAGCAGAGCTT | NO_CURRENT_688 | NO_CURRENT_688 |
| GGGCACATACCGCTCTGGAG | GNAI2 | 2771 |
| GGGCAGAAGTTGCTGTCCTG | NO_CURRENT_689 | NO_CURRENT_689 |
| GGGCAGCACCGAAGACCAGA | SERPINB2 | 5055 |
| GGGCATAAGTTAGCCACTCA | NO_CURRENT_690 | NO_CURRENT_690 |
| GGGCATGACTACATCTGTCA | TP73 | 7161 |
| GGGCATGTACAACACCGACG | IDH2 | 3418 |
| GGGCCACGACAAGTCTGACA | PQBP1 | 10084 |
| GGGCCAGGTTTCGCGTGCTG | KIAA1211L | 343990 |
| GGGCCCATAATCGAGCCCAG | CHEK2 | 11200 |
| GGGCCCGCATAGGATATCGC | NO_CURRENT_691 | NO_CURRENT_691 |
| GGGCCGAGGCAAGATAAGCA | CFLAR | 8837 |
| GGGCCTGGAGGAACAAGTCG | NUFIP2 | 57532 |
| GGGCGACATGATGGCGTCAG | MCL1 | 4170 |
| GGGCGCACAGAATTTATCTG | PARK7 | 11315 |
| GGGCGGAACGGATTCCAGCA | TP73 | 7161 |
| GGGCGTGTATGTTCGTATTG | NO_CURRENT_692 | NO_CURRENT_692 |
| GGGCTACCTCTGCTTGCGGG | SERF2 | 10169 |
| GGGCTATTAAGCGGCGACCA | ARNT | 405 |
| GGGCTCCCCAGTGACGACCG | SBNO2 | 22904 |
| GGGCTCGCCTGTCAACGCGC | FOS | 2353 |
| GGGCTGCATACATTTGCTGA | PHC3 | 80012 |
| GGGCTGCCAGCGTATCGTAG | MTOR | 2475 |
| GGGCTTGACATTAAGATACC | NO_CURRENT_693 | NO_CURRENT_693 |
| GGGCTTTCTCATCGGCGATG | SBNO2 | 22904 |
| GGGGAAACAAGTAGGCTTTG | NO_CURRENT_694 | NO_CURRENT_694 |
| GGGGAAGACTGTAATAGCGG | DLAT | 1737 |
| GGGGAAGGACTCCACTAGAG | NENF | 29937 |
| GGGGACTGAATCCTGACCCA | ARMCX2 | 9823 |
| GGGGATATAAAATGCCACTG | HTR7 | 3363 |
| GGGGATGGAACCGTAGAGCA | SLC25A1 | 6576 |
| GGGGATGTACTCTCCGGGAA | RAC2 | 5880 |
| GGGGCACTGTTATCTGCACG | PDE5A | 8654 |
| GGGGCCACTAGGTGTGACAG | STAT2 | 6773 |
| GGGGCCATAGATGAGTAGCG | HELZ2 | 85441 |
| GGGGCCGAGCCTATGCCAGG | KAT2A | 2648 |
| GGGGCTCACTATGACATCGT | CASP5 | 838 |

| | | |
|---|---|---|
| GGGGCTGCAGCAAACTGGAA | NLRP3 | 114548 |
| GGGGCTTACGTGAAGGGCGG | NO_CURRENT_695 | NO_CURRENT_695 |
| GGGGGACACACCATAGACAG | ABL1 | 25 |
| GGGGGAGTCTGTGTCCACGG | BAX | 581 |
| GGGGGCCAGCAGCCGGGAAA | NDUFC1 | 4717 |
| GGGGGCTACGAGGAGTACAG | HNRNPF | 3185 |
| GGGGTCGTAGGTCTTATGGT | VDR | 7421 |
| GGGGTTGAGGAAAACACTGG | CYP4F11 | 57834 |
| GGGGTTGGTGTGTGCTGGAT | MPC2 | 25874 |
| GGGTAAAAGTACTGTCCCGG | JUNB | 3726 |
| GGGTAACTGCAATTGGCAAA | C4orf17 | 84103 |
| GGGTAATGGTGACTCAGTGT | NO_CURRENT_696 | NO_CURRENT_696 |
| GGGTACAATGTAGACTACGT | NO_CURRENT_697 | NO_CURRENT_697 |
| GGGTACACCTACACAGCGGG | MEX3B | 84206 |
| GGGTACCACCGATATGATCA | YPEL5 | 51646 |
| GGGTACCTCAGCACCCCCAG | RAD23A | 5886 |
| GGGTAGATCCATGAAACAGC | NO_CURRENT_698 | NO_CURRENT_698 |
| GGGTAGGATTCAGCCAAAAG | NO_CURRENT_699 | NO_CURRENT_699 |
| GGGTAGTGGGCTGTCCTGTG | TIRAP | 114609 |
| GGGTATAGACGCGATCCTCA | NO_CURRENT_700 | NO_CURRENT_700 |
| GGGTCAGCACTACTTCGAAG | HIF1A | 3091 |
| GGGTCAGGAGCATTGACTGG | SPEN | 23013 |
| GGGTCCTACAGTGCTAGCCA | GPT2 | 84706 |
| GGGTCGTTCCAAGTCCGCGC | ACTR5 | 79913 |
| GGGTCTACTAATAATAGGAG | TNFRSF1B | 7133 |
| GGGTCTTGGTTCTCAGCTTG | IL10 | 3586 |
| GGGTGCCCACTAATAGCCGC | NO_CURRENT_701 | NO_CURRENT_701 |
| GGGTGCGCAGAACATCCACG | SLC25A19 | 60386 |
| GGGTGGTCATTCTCTACTTG | NO_CURRENT_702 | NO_CURRENT_702 |
| GGGTGTCAGAGCTTGATGTT | PPARGC1B | 133522 |
| GGGTGTCCGCGATCTCAGGA | ARMCX2 | 9823 |
| GGGTTGCATCATGGGAGAGC | MAPK9 | 5601 |
| GGGTTTCAGCGAGGCTTGTG | ITGB2 | 3689 |
| GGTAAACCTATGCTACGCCA | STRAP | 11171 |
| GGTAATAACACCAGTACGGA | MCL1 | 4170 |
| GGTAATAACCACATGGAAGG | INO80 | 54617 |
| GGTACACCCAGTGCCTTAGG | PIGL | 9487 |
| GGTACATTAGGACGGACCCC | CUBN | 8029 |
| GGTACCGGGGCAGCTACTGG | TMEM173 | 340061 |
| GGTAGACGGGGCATCTCAGC | NO_CURRENT_703 | NO_CURRENT_703 |
| GGTAGACGTGTAGGGCCAGA | BAK1 | 578 |
| GGTAGATATATGGAGCAGTG | PRKAA1 | 5562 |
| GGTAGTCAGAAGGGAATGGG | INO80E | 283899 |
| GGTATGAAGACATTCTGTGA | PCGF6 | 84108 |
| GGTATGAGCCTAATTTCCAC | NO_CURRENT_704 | NO_CURRENT_704 |
| GGTATTCGTATGACCCACCA | VIPR2 | 7434 |
| GGTCAACTGTCCAGAGAAAG | BABAM1 | 29086 |
| GGTCACCGATCGAGAGCTAG | NO_CURRENT_705 | NO_CURRENT_705 |
| GGTCACGGGACTCCGTGGCG | SBNO2 | 22904 |
| GGTCAGCAGATATCAGTCCA | PDE5A | 8654 |

| | | |
|---|---|---|
| GGTCAGGGAACAGCCCGTGG | SIAH2 | 6478 |
| GGTCATCCGAGCCAACATCG | GPT2 | 84706 |
| GGTCCAAATTCGGACACCAA | BABAM1 | 29086 |
| GGTCCCTACTCACACCATGC | DNTTIP1 | 116092 |
| GGTCCCTCTGGCTGGGTTAA | NO_CURRENT_706 | NO_CURRENT_706 |
| GGTCCCTGCCAGGCACGAGG | TYK2 | 7297 |
| GGTCCGAGCCGTGATCAGGA | NLRP1 | 22861 |
| GGTCCGCGCACAAGAGCAGG | NO_CURRENT_707 | NO_CURRENT_707 |
| GGTCGCCTGTGCGACATGCT | NO_CURRENT_708 | NO_CURRENT_708 |
| GGTCGCTGACATGCACTCTG | TNRC18 | 84629 |
| GGTCGGCGAAGAGCTCGTCG | CEBPD | 1052 |
| GGTCTAGAAGTACCTAAACC | PCGF6 | 84108 |
| GGTCTCATGAGTAACTCGGA | UCHL3 | 7347 |
| GGTCTGCTCCAATGGGAACC | NO_CURRENT_709 | NO_CURRENT_709 |
| GGTCTGTCTGGATAAAAGCG | USP7 | 7874 |
| GGTGAAATGGATGATAGTCG | KMT2C | 58508 |
| GGTGACACGGAACTACACCT | CACNA2D2 | 9254 |
| GGTGACTATAGCGACGAGAG | LRP2 | 4036 |
| GGTGACTGGGTCAACTGTAC | PDK4 | 5166 |
| GGTGCACAGGGTGTCATGGG | TNFRSF6B | 8771 |
| GGTGCAGCGGGAGATTCACC | BRK1 | 55845 |
| GGTGCATAGTGCCATCACCT | GPT2 | 84706 |
| GGTGCCATTTGTCATACTGA | NO_CURRENT_710 | NO_CURRENT_710 |
| GGTGCCGGCTGTACGCGGAG | TNFRSF6B | 8771 |
| GGTGCCTTTGACGAGCACAT | NLRP3 | 114548 |
| GGTGCTCCAATATTACCTCA | NMU | 10874 |
| GGTGCTCTTGAGAGTTTGAG | CASP9 | 842 |
| GGTGCTGGTCTATAAAGTGC | PYCARD | 29108 |
| GGTGCTTCAGGAGGTACAAG | CYP8B1 | 1582 |
| GGTGGAAATTCCCGCCAACG | KMT2D | 8085 |
| GGTGGAACTCCACCGTGTGC | TRMT61A | 115708 |
| GGTGGATAGTTCTGCTGCAA | RNF25 | 64320 |
| GGTGGATGCTACACGTCCTG | SPEN | 23013 |
| GGTGGTGGCAGAACCGACGG | KMT2D | 8085 |
| GGTGGTGGTGAACGATGACA | BABAM1 | 29086 |
| GGTGTAAAGTCTGGTCCTCG | EIF4G3 | 8672 |
| GGTGTAGCTGATGAGAAGGC | RAC2 | 5880 |
| GGTGTCACCACCGCTTACCA | NO_CURRENT_711 | NO_CURRENT_711 |
| GGTGTCCAGAGAGTGGACGA | EIF4G3 | 8672 |
| GGTGTCCAGGAATCCAGTGC | CDKN2D | 1032 |
| GGTGTCCTTCAAAGACTGGA | CHMP5 | 51510 |
| GGTGTCTTACGTAAAGAGGT | NO_CURRENT_712 | NO_CURRENT_712 |
| GGTTACATGGTTAGATGGAG | NO_CURRENT_713 | NO_CURRENT_713 |
| GGTTCTTTGAGCAACATGGG | RB1 | 5925 |
| GGTTGGGGTGAACCCCATTG | BAP1 | 8314 |
| GGTTGTAAATCCTTCTAACG | NO_CURRENT_714 | NO_CURRENT_714 |
| GGTTTCAGAATCCCACTCTG | SLC11A1 | 6556 |
| GGTTTCTGGTATTGACATCG | KIF2B | 84643 |
| GGTTTGAGTGAGCAATTACG | NO_CURRENT_715 | NO_CURRENT_715 |
| GGTTTGGGAGTATATGACCG | MGA | 23269 |

| | | |
|---|---|---|
| GTAAAACCAGGATAGGTCTG | TBXAS1 | 6916 |
| GTAAAGAAATGACGTACCAA | LARP4B | 23185 |
| GTAAATTCTATATAGCCCAG | NIPBL | 25836 |
| GTAACAACAGGGATTCACTG | PTGS1 | 5742 |
| GTAACACCCAAAGCTAACTC | NO_CURRENT_716 | NO_CURRENT_716 |
| GTAACAGGTCCTCGGAAGGG | FOXO4 | 4303 |
| GTAACAGTCTGATTACGGAC | NO_CURRENT_717 | NO_CURRENT_717 |
| GTAACCAACATTGGTGCAGT | MBNL1 | 4154 |
| GTAACCATATTTGCATTTGA | BCL2A1 | 597 |
| GTAACTTGTCCCATCATCGA | IKZF5 | 64376 |
| GTAAGACACTAACCACAGCC | GPR119 | 139760 |
| GTAAGGAATGACTTGACTGG | NO_CURRENT_718 | NO_CURRENT_718 |
| GTAATAGAGTGGACTGTACG | CUBN | 8029 |
| GTAATTCTCCATTTACTGTA | MCTS1 | 28985 |
| GTAATTTTATGAGTTAAGTG | NO_CURRENT_719 | NO_CURRENT_719 |
| GTACAAATCAAAGGCGAGGT | NO_CURRENT_720 | NO_CURRENT_720 |
| GTACAATTGGCCAAACCGGA | CACNA2D2 | 9254 |
| GTACACACATTGAACCTCAG | EDA2R | 60401 |
| GTACACACTTATGCCATCAC | NO_CURRENT_721 | NO_CURRENT_721 |
| GTACACCATATCAAAAGGGC | SMPD1 | 6609 |
| GTACAGCTCATGGATACCAC | NAIP | 4671 |
| GTACATTGTATAAAGACCAC | IFNAR1 | 3454 |
| GTACCACTCCCTCTAGGTGA | RAD23A | 5886 |
| GTACCATTGCCGGCTCCCTA | NO_CURRENT_722 | NO_CURRENT_722 |
| GTACCCCTATGGCCGTTCTA | NO_CURRENT_723 | NO_CURRENT_723 |
| GTACGAGCTCCCGGTCCCGA | JUNB | 3726 |
| GTACGGCGGCCAATACCGGA | APEH | 327 |
| GTACGGCTGACAGCAGGGGT | FAM214B | 80256 |
| GTACGTACTCTACTGGATAG | ANKRD50 | 57182 |
| GTACTACGCCAAGGTTGAGC | ATP5L | 10632 |
| GTACTTGCTCGGATCCACAC | SLC7A3 | 84889 |
| GTAGAAGGAGACTACGGACG | ADRB1 | 153 |
| GTAGACGTCGTGAGCTTCAC | NO_CURRENT_724 | NO_CURRENT_724 |
| GTAGATACAGAACAATGTGA | NUDT17 | 200035 |
| GTAGATGAAATGGACACCTG | HNRNPF | 3185 |
| GTAGCACCGTGCCAGCCGTA | TMEM30A | 55754 |
| GTAGCCCAGATCTGCCGCAG | ANKRD50 | 57182 |
| GTAGGCGCGCCGCTCTCTAC | NO_CURRENT_725 | NO_CURRENT_725 |
| GTAGGGTACAGCGTCAGCTT | NO_CURRENT_726 | NO_CURRENT_726 |
| GTAGTACATGAAAAGCATGA | HTR7 | 3363 |
| GTAGTCCCTACAATGGCACA | PDK4 | 5166 |
| GTAGTCTCGAGAGATAGACC | SLC11A1 | 6556 |
| GTAGTGCAGGGGGTAACAGA | OR7C2 | 26658 |
| GTATAAACATTACCAAGTGG | NO_CURRENT_727 | NO_CURRENT_727 |
| GTATAATTGTCAATGACAAT | NO_CURRENT_728 | NO_CURRENT_728 |
| GTATACAGAGTTAGACCCAC | NO_CURRENT_729 | NO_CURRENT_729 |
| GTATAGCCTGCTGTACAGTT | NO_CURRENT_730 | NO_CURRENT_730 |
| GTATATAACACAGACTACAC | MSL2 | 55167 |
| GTATATTATCAGCTAAATGT | NO_CURRENT_731 | NO_CURRENT_731 |
| GTATCCCATATCGGCACAGG | NO_CURRENT_732 | NO_CURRENT_732 |

| | | |
|---|---|---|
| GTATCGGCCTCGCCACTGCA | BAP1 | 8314 |
| GTATCTGCTGCGTATAACAT | WDR26 | 80232 |
| GTATGAGAAAGAACTATCAA | NO_CURRENT_733 | NO_CURRENT_733 |
| GTATGGTAATAGGCAAGAGC | PIGY | 84992 |
| GTATGTCGAGAGTACCAACG | MBNL1 | 4154 |
| GTATTAAGATGCGTCTTAGA | NO_CURRENT_734 | NO_CURRENT_734 |
| GTATTCACACACTCAACGGG | CUBN | 8029 |
| GTCAAGAGATTATGAGATTC | NO_CURRENT_735 | NO_CURRENT_735 |
| GTCAAGCCCGACCCTCCAGA | EBI3 | 10148 |
| GTCAATACTAGGCAGATCAA | NO_CURRENT_736 | NO_CURRENT_736 |
| GTCAATGCTGATCACGCACA | HTR7 | 3363 |
| GTCACAGCAGTAGAGTACCA | JAM2 | 58494 |
| GTCACAGGATTTGATCTGTG | SLC5A8 | 160728 |
| GTCACCCCAAGTACGCCTTG | AHDC1 | 27245 |
| GTCACCTCATGAGGTAGTCC | MPC1 | 51660 |
| GTCACTGACAAACAGCAACC | KPNA1 | 3836 |
| GTCAGACTCCCAGATGTGCT | HSP90B1 | 7184 |
| GTCAGAGTCAATAACCAGCT | GSDMD | 79792 |
| GTCAGGGTTACGGTTCCACC | AGER | 177 |
| GTCAGGTAATAGTCGGACTC | NO_CURRENT_737 | NO_CURRENT_737 |
| GTCATCACGGCCATAGCGAA | ARR3 | 407 |
| GTCATCAGCGATTTGACGAG | NO_CURRENT_738 | NO_CURRENT_738 |
| GTCATCCTTGTAATCCATCA | JAK1 | 3716 |
| GTCATCTGCCTCACTGACGT | CXCR4 | 7852 |
| GTCATTGTCATAAAACACGA | EP400 | 57634 |
| GTCCAATAAAAAGTGCCACT | NO_CURRENT_739 | NO_CURRENT_739 |
| GTCCAATGTCCAGCCCATGA | BAX | 581 |
| GTCCAGTAGCAAATATGACA | PHIP | 55023 |
| GTCCAGTTCTGTACCACGGC | CASP3 | 836 |
| GTCCAGTTTAATGAAGTCCA | NLRC4 | 58484 |
| GTCCATCTGATAGATATGAG | KPNA1 | 3836 |
| GTCCATTTGCGGAGGAACAT | ATP5J | 522 |
| GTCCCAGAAGCTACGCACCA | LTB4R | 1241 |
| GTCCCGGATGTGATGCCCAA | MLF2 | 8079 |
| GTCCCGTGATTTTAGCCAGG | NO_CURRENT_740 | NO_CURRENT_740 |
| GTCCGCTTCCAGCTTTGCGC | INO80C | 125476 |
| GTCCGTCGACCCTTATTGGG | NO_CURRENT_741 | NO_CURRENT_741 |
| GTCCGTGCTGGCAAAGACCA | ACADS | 35 |
| GTCCTACAGATACCACAACC | BCL2A1 | 597 |
| GTCCTCATCCGGTCAGGCTG | NO_CURRENT_742 | NO_CURRENT_742 |
| GTCCTCGGCCTTACCGGCCG | PAGR1 | 79447 |
| GTCCTGGGCTGACTTCACTA | STAT2 | 6773 |
| GTCCTGTATGAGTGGGAACA | CTNNB1 | 1499 |
| GTCCTGTCTTTGTCACAGAA | IRAK4 | 51135 |
| GTCCTGTGCCAATCTCCATG | NDUFS4 | 4724 |
| GTCCTTGGTTACAACCAGAA | ABL2 | 27 |
| GTCGAGATGGCAGTGACCGT | FOS | 2353 |
| GTCGGAGACTTCCGCTTCCG | SPCS1 | 28972 |
| GTCGTAATTCTAACTCCAAG | CSDE1 | 7812 |
| GTCGTGAACTCGCCTTGCAG | IL2RB | 3560 |

| | | |
|---|---|---|
| GTCGTTCACATTAACGCCAC | SCTR | 6344 |
| GTCTCAATAGGACATCAACC | ACO2 | 50 |
| GTCTCCAGCAGCTATAACAG | PHC3 | 80012 |
| GTCTCCGAGAACGCCGCACG | PPARGC1B | 133522 |
| GTCTCGTCAGTAGTGAGGGA | RTP1 | 132112 |
| GTCTGATTCTGGTTCTACTG | SLC3A2 | 6520 |
| GTCTGCACGTTCTCTGTGAC | BABAM1 | 29086 |
| GTCTGCGTACTTCCAGACCA | EGFR | 1956 |
| GTCTGGTTTCTGAACCACCA | IKZF5 | 64376 |
| GTCTGTGATCAACCCATCCT | CYP2R1 | 120227 |
| GTCTGTTGCAAGGGCAAAAG | LAMTOR3 | 8649 |
| GTGAAACCCACCCTGAGTCA | NO_CURRENT_743 | NO_CURRENT_743 |
| GTGAAAGGCATTCCTCACAC | CYP4A11 | 1579 |
| GTGAAATTGAACGGCGGCGA | ARNT | 405 |
| GTGAACTGCAATCTTATTAT | NO_CURRENT_744 | NO_CURRENT_744 |
| GTGAAGAGAGAACTTATCCA | ZNF699 | 374879 |
| GTGAAGATCACCAAGTCCGA | SLC25A6 | 293 |
| GTGAAGGCTGCATACAACCC | CCAR2 | 57805 |
| GTGAATAATCAAATCATCCC | NO_CURRENT_745 | NO_CURRENT_745 |
| GTGAATCCTGCGTCTGTAGG | KDM4A | 9682 |
| GTGACAGAGTTGCTATTGGC | CYP27B1 | 1594 |
| GTGACCCTAAGAGTGACTCG | GBP7 | 388646 |
| GTGACCCTGTTGGCACAAGA | NO_CURRENT_746 | NO_CURRENT_746 |
| GTGACGAAGTCGCAGCACTT | ADRB1 | 153 |
| GTGACGCTTTACCTGCGACC | ITGB2 | 3689 |
| GTGACTGAGCTGAGGCTTGG | TIRAP | 114609 |
| GTGAGAAGTATCGGCGTCTG | SPHK1 | 8877 |
| GTGAGACCGTACTCAGAGTG | DPY30 | 84661 |
| GTGAGCAGGTGGCATACTTG | BABAM2 | 9577 |
| GTGAGCCCACAAATAGACGG | PLD1 | 5337 |
| GTGAGGATGATTGCTCCCGA | PITPNB | 23760 |
| GTGAGGTGGTTGTCGCACTG | NO_CURRENT_747 | NO_CURRENT_747 |
| GTGAGTTTGAAGAGCTCGAG | KDM4A | 9682 |
| GTGATAATGATGTATTCTCG | NO_CURRENT_748 | NO_CURRENT_748 |
| GTGATCAGCTTCTACTATGG | IRF8 | 3394 |
| GTGATCGAGTTAACTGTGCA | JAM3 | 83700 |
| GTGATGGTTCAGCCAAACGC | CTNNB1 | 1499 |
| GTGATGTGAAAACATGTCAC | PHF6 | 84295 |
| GTGATGTTGGGCTCGCTATG | POU2F1 | 5451 |
| GTGATTCTGTCCGTTCAGTG | PAXIP1 | 22976 |
| GTGCAGTGTCTGATGAGAGG | ABCB10 | 23456 |
| GTGCATCTCAGGCCGACTGG | SCN5A | 6331 |
| GTGCATCTGTCACATAACTG | SATB2 | 23314 |
| GTGCCAGTCACACCCCAACG | RPS6KA4 | 8986 |
| GTGCCAGTGTACCCTGCTGT | SMPD1 | 6609 |
| GTGCCATGTTTAAGTCAATG | NO_CURRENT_749 | NO_CURRENT_749 |
| GTGCCCATGACCTTTCAAAG | STAG2 | 10735 |
| GTGCCCTCTGAGTTTAGCAC | INO80C | 125476 |
| GTGCCGAAGGCTGTACCAGT | JAM3 | 83700 |
| GTGCCTATGATAAACCACCT | NO_CURRENT_750 | NO_CURRENT_750 |

| | | |
|---|---|---|
| GTGCCTTCAAATGAGAAGTT | SLC7A5 | 8140 |
| GTGCGACGAATTGTCCTGAG | NO_CURRENT_751 | NO_CURRENT_751 |
| GTGCGAGCGAGGGTATGGGT | FOSL2 | 2355 |
| GTGCGCATGGGCTGATGTTA | NO_CURRENT_752 | NO_CURRENT_752 |
| GTGCGGACCACATATGGCGG | BABAM1 | 29086 |
| GTGCGGATACAGTTCAAGCA | KHSRP | 8570 |
| GTGCGTCATGTGAGCATGTG | NO_CURRENT_753 | NO_CURRENT_753 |
| GTGCGTGCATCGACTCGTGA | PAXIP1 | 22976 |
| GTGCGTGCCCTGCAGCGACG | PAGR1 | 79447 |
| GTGCTAATGTACTTCAACCT | CSNK1D | 1453 |
| GTGCTCAAAACATCACAGCC | AGER | 177 |
| GTGCTCACGTAGCCAGAGCG | PTGS1 | 5742 |
| GTGCTCGGGACTGTGCCCAT | FAM214B | 80256 |
| GTGCTGGTGACGGAGAAGGG | ARID3A | 1820 |
| GTGGAACACCTACTGGACCA | ANKRD50 | 57182 |
| GTGGAAGAAAACTTCCCACA | PIGY | 84992 |
| GTGGAATAACAGACTACTGA | RAD21 | 5885 |
| GTGGACAATGGATTGGCACA | SMPD1 | 6609 |
| GTGGACAGTAACCTCAACTG | ARHGAP33 | 115703 |
| GTGGACATGAGCCACGTGCT | SETD1B | 23067 |
| GTGGACCGGGACACCGTGTG | TNFRSF1A | 7132 |
| GTGGAGAACCGAACTGCGAT | CAT | 847 |
| GTGGAGCCTCTTACACCCAG | EGFR | 1956 |
| GTGGAGTCCATCTTCCGCAA | KIF2B | 84643 |
| GTGGATTTGGGTTCACGAAG | NO_CURRENT_754 | NO_CURRENT_754 |
| GTGGCACCTCTTCCAGAACT | C16orf72 | 29035 |
| GTGGCAGACCATATTCCCAA | PDE3A | 5139 |
| GTGGCATACTGGAGTCACTG | NFRKB | 4798 |
| GTGGCATTATAAAACGCTCA | CBLL1 | 79872 |
| GTGGCCCATGACCATACAGG | CYP4B1 | 1580 |
| GTGGCCTACTGGAGACAGGC | ATP5E | 514 |
| GTGGCTGGAACCCGGATCTG | NFRKB | 4798 |
| GTGGGAAGGCGTCTGTACCG | NCOA6 | 23054 |
| GTGGGCATACGGGAACTGGT | CYP4B1 | 1580 |
| GTGGGCCAAAGGATGAAGAG | SOD1 | 6647 |
| GTGGGGCCATTATTGATACT | NO_CURRENT_755 | NO_CURRENT_755 |
| GTGGGGCCTGCAAATATCCG | TNFRSF1B | 7133 |
| GTGGGGTATCATACCGTCTG | NNT | 23530 |
| GTGGGGTCATAGTACCAGCG | SPINT1 | 6692 |
| GTGGTCCGGAATGGAAAAGT | BABAM2 | 9577 |
| GTGGTCTGGTCTTGTGAACT | BMI1 | 648 |
| GTGGTCTGGTCTTGTGAACT | COMMD3-BMI1 | 100532731 |
| GTGGTCTTGTCACACAACCA | DET1 | 55070 |
| GTGGTGGAGAACCCAAAGGT | SOD2 | 6648 |
| GTGGTGGCGGCCGTTATTGG | SAP18 | 10284 |
| GTGGTGGGAGACAGTCAGTG | RND3 | 390 |
| GTGGTGTGGGAAATTTAACA | NO_CURRENT_756 | NO_CURRENT_756 |
| GTGTAAAAAAAATCATGATG | CASP5 | 838 |
| GTGTAAGAAGTTCCCTCCAA | PHF8 | 23133 |
| GTGTAAGTGATCTTGGCACA | HTR2A | 3356 |

| | | |
|---|---|---|
| GTGTAATTCTCAAACACTGT | RND3 | 390 |
| GTGTAATTTAGAGAGCAGCG | RAD21 | 5885 |
| GTGTACATGATGCCAACCGA | USP7 | 7874 |
| GTGTACATTATACAGGTGCT | NO_CURRENT_757 | NO_CURRENT_757 |
| GTGTAGGAAGGCGATATCCA | RIPK3 | 11035 |
| GTGTATGAATGTTAATTCCG | NO_CURRENT_758 | NO_CURRENT_758 |
| GTGTATGATGCTTCGACTTA | NO_CURRENT_759 | NO_CURRENT_759 |
| GTGTGACAAGACTCAATGAT | PRKN | 5071 |
| GTGTGGCAGTACCTTCAAGC | RIPK1 | 8737 |
| GTGTGTCTCCGAGTGCGACC | APEH | 327 |
| GTGTTCAAACATGTCCAGTC | CYP4F11 | 57834 |
| GTGTTCACAGTGCAGCTAGG | CYP8B1 | 1582 |
| GTGTTGATGCACTACCCCCG | PTGS1 | 5742 |
| GTGTTTACTAGCATATTGGA | MAP2K5 | 5607 |
| GTGTTTCCAGTTCTTTACGT | ALOX5 | 240 |
| GTTAAAAGTGGAGCAGCACG | TGFB1 | 7040 |
| GTTAACAAACAGGCTACTGC | NO_CURRENT_760 | NO_CURRENT_760 |
| GTTAACTAATATCCCAAAGG | NO_CURRENT_761 | NO_CURRENT_761 |
| GTTAATCCTTCGTCCAACAC | AHCTF1 | 25909 |
| GTTACCTGCTACGAAAACGA | NO_CURRENT_762 | NO_CURRENT_762 |
| GTTACTGAACTGTACCCACC | UCHL5 | 51377 |
| GTTACTTACCAAAGGTTCCA | ANP32B | 10541 |
| GTTAGAATGTCTGAAAAGCC | ANP32B | 10541 |
| GTTAGCACTTACCTTTGCAG | HMGN2 | 3151 |
| GTTAGCTGCAGAAATAGCAT | PDE3A | 5139 |
| GTTAGTGCCCCATCTGAGCA | JAM2 | 58494 |
| GTTATCATCAATTAATCCTG | NO_CURRENT_763 | NO_CURRENT_763 |
| GTTCAAAGAGCATATGATGG | PDK4 | 5166 |
| GTTCACAGTATTGTACACAA | GBP5 | 115362 |
| GTTCACCTATTACCCTCTGG | NFKB2 | 4791 |
| GTTCACGCCTGTGAACACTT | SLC7A3 | 84889 |
| GTTCAGTAACACACTCACTA | UCHL5 | 51377 |
| GTTCATCAAGGTTCACAACG | FOXO4 | 4303 |
| GTTCATCGAGCCCATCGATG | BRD1 | 23774 |
| GTTCATTCACCACCAAACGT | PHIP | 55023 |
| GTTCATTTCTGTGACAAGTG | CBLL1 | 79872 |
| GTTCCCCGGGAAGTCTATGC | NO_CURRENT_764 | NO_CURRENT_764 |
| GTTCCCGAGTGCTTGAGCTG | TNFRSF1B | 7133 |
| GTTCCCTAGGAATCGGACGT | RRAGA | 10670 |
| GTTCCGATGAGGTTAGATGA | NO_CURRENT_765 | NO_CURRENT_765 |
| GTTCCGGACTATTCCGGGAG | INO80C | 125476 |
| GTTCCGTGAGGGTTACTTCA | NO_CURRENT_766 | NO_CURRENT_766 |
| GTTCCTCTTAGAGATACCGA | SUV39H1 | 6839 |
| GTTCCTGTACGACGAAATTG | CYFIP1 | 23191 |
| GTTCGCTTCGTAACGAGGAA | NO_CURRENT_767 | NO_CURRENT_767 |
| GTTCGTCAGGAATCCCATTG | LGMN | 5641 |
| GTTCTATATCCACTGTCGAG | NLRP3 | 114548 |
| GTTCTCACCGGAAAGCACAG | SCTR | 6344 |
| GTTCTCAGGAAGTCTTCTTA | UQCRB | 7381 |
| GTTCTCAGTCCAAAAAGAAG | BCL2A1 | 597 |

| | | |
|---|---|---|
| GTTCTCCTGAATCAGACTGA | NLRP3 | 114548 |
| GTTCTCTGTGAAGATCTTGA | FAM107B | 83641 |
| GTTCTGATAATCAATACCAC | ZNF699 | 374879 |
| GTTCTGATGCAGCTTCCATG | BCL2L11 | 10018 |
| GTTCTTAGAACTATAGATTA | NO_CURRENT_768 | NO_CURRENT_768 |
| GTTGAATTCTGAAGTGTGAT | PHC3 | 80012 |
| GTTGACACTCATGATGACTC | ACADS | 35 |
| GTTGACTGTTCTTATTCCAC | PIGY | 84992 |
| GTTGATGCCCAACCCATCAA | NDUFS4 | 4724 |
| GTTGCTGCTTATGTGCCAAG | OXSM | 54995 |
| GTTGGAGGCCCCCCTAAACG | KMT2D | 8085 |
| GTTGGAGTAGAACCTAGACC | FBXW7 | 55294 |
| GTTGGCATATTGGCCCAGAC | NO_CURRENT_769 | NO_CURRENT_769 |
| GTTGGCCCGTTCTTATGTCC | NO_CURRENT_770 | NO_CURRENT_770 |
| GTTGTAAAATCAAAGCGCTG | MEX3B | 84206 |
| GTTGTAGAAGCATGTGAACT | IL2RB | 3560 |
| GTTGTCAAAAAACACATATG | JAM3 | 83700 |
| GTTGTCAATAATCATTACAC | PIGL | 9487 |
| GTTGTCCTGAGGGATGGACG | ALOX5 | 240 |
| GTTGTGAATTTGATCATGGG | UBE2L6 | 9246 |
| GTTGTGAGAGGGAATATCGA | NO_CURRENT_771 | NO_CURRENT_771 |
| GTTGTGCGCAAAGCTCAGGG | MEX3B | 84206 |
| GTTGTGGACGATCAACGGGG | JAK1 | 3716 |
| GTTGTGGTCAAAGAGAATGG | RPS6KA4 | 8986 |
| GTTGTGTCGAGATATCTGCC | ABI1 | 10006 |
| GTTGTTATTTCCAAGGAATG | NO_CURRENT_772 | NO_CURRENT_772 |
| GTTTAAAGAAAGGGGCTAAG | NO_CURRENT_773 | NO_CURRENT_773 |
| GTTTAAGGAATACTGCCCGA | PIP4K2A | 5305 |
| GTTTAATACCACAAAGCATC | MAPK9 | 5601 |
| GTTTAATGTATCGCTACCAA | TNFRSF1A | 7132 |
| GTTTACTCATATCCAGTCAC | NO_CURRENT_774 | NO_CURRENT_774 |
| GTTTATACAGATTCACTCCA | ACOD1 | 730249 |
| GTTTCATCCAGGATCGAGCA | BAX | 581 |
| GTTTCCATACATGTCCTACT | NO_CURRENT_775 | NO_CURRENT_775 |
| GTTTCCTGACTGACCGATCG | CBLL1 | 79872 |
| GTTTCGAAACTTGAAGTAAG | NO_CURRENT_776 | NO_CURRENT_776 |
| GTTTCGAAGGGCACTGATGA | IRGM | 345611 |
| GTTTCGCATTAAGAGGGCAC | UQCRB | 7381 |
| GTTTCTCACCAATAGCCGAA | INO80 | 54617 |
| GTTTCTCCAAACACAAATAC | NO_CURRENT_777 | NO_CURRENT_777 |
| GTTTCTCTTGAACCCCGCGA | NO_CURRENT_778 | NO_CURRENT_778 |
| GTTTGATACCAGAGAGCGAT | NDUFS4 | 4724 |
| GTTTGCACAAGAAAACAGAC | YPEL5 | 51646 |
| GTTTGCCTACCTCGTGAACG | SPEN | 23013 |
| GTTTGGGATCGTCTTATCGG | OXSM | 54995 |
| GTTTGTAGTCGTGTAGAGAG | JUNB | 3726 |
| GTTTGTTAACGTAAACCGGA | RIPK3 | 11035 |
| GTTTTCAGTTGCCCAACAGC | NO_CURRENT_779 | NO_CURRENT_779 |
| GTTTTTGGTTAATTGCCTAC | NO_CURRENT_780 | NO_CURRENT_780 |
| TAAAAAATAATTCCTCACAG | RNF111 | 54778 |

| | | |
|---|---|---|
| TAAAAACCAGAATCCCAGTA | TMEM30A | 55754 |
| TAAAAAGACCAATCTGATCG | TIFA | 92610 |
| TAAAAAGGTATGAGCAGCAG | CHMP5 | 51510 |
| TAAAATGAGGACATTAGTGT | NO_CURRENT_781 | NO_CURRENT_781 |
| TAAACACATTGTCAAGACTG | STRAP | 11171 |
| TAAACACCAACTCATTGCGT | MAP3K7 | 6885 |
| TAAACCCTTGGCGCCCTCAC | NO_CURRENT_782 | NO_CURRENT_782 |
| TAAACTATCAGGGCTGTCGA | RHOA | 387 |
| TAAAGCAGAAGAATATACAG | NO_CURRENT_783 | NO_CURRENT_783 |
| TAAAGGCCAGACAATTCCAG | MAP2K6 | 5608 |
| TAAAGTGAGTTACACATCAC | PRR14L | 253143 |
| TAAATAGGCTGTACCAGTGC | SOD1 | 6647 |
| TAAATCTAATGAAGAATCGG | POU2F1 | 5451 |
| TAAATTCATACATATCGATG | NO_CURRENT_784 | NO_CURRENT_784 |
| TAAATTTAGGATGAAGAATA | CCL20 | 6364 |
| TAACAAGATAGCTCAAAACA | NO_CURRENT_785 | NO_CURRENT_785 |
| TAACAGTAGAAATCCTACTG | RNF111 | 54778 |
| TAACATTTCTACTTTACCAG | SHOC2 | 8036 |
| TAACCCAGAAGCCCATTCAG | NO_CURRENT_786 | NO_CURRENT_786 |
| TAACCCATGAAGTGCAGGAC | MSL2 | 55167 |
| TAACCGATACTCCCCACATT | NO_CURRENT_787 | NO_CURRENT_787 |
| TAAGATACATGTCACTAGCA | LARP4B | 23185 |
| TAAGATGACCAGCATCACCA | CXCR1 | 3577 |
| TAAGATGTTCACCTAATGCT | CCNC | 892 |
| TAAGCATCAACAAAATGCTG | CYP2R1 | 120227 |
| TAAGCATGGATTCCATTACT | RB1 | 5925 |
| TAAGCCCGGGAGAGCGATGG | SPCS1 | 28972 |
| TAAGGAACATCTCTTCCCAG | ACADS | 35 |
| TAAGGATTTCTATATGTTCA | MCTS1 | 28985 |
| TAAGGCATAATAATAGACTC | NO_CURRENT_788 | NO_CURRENT_788 |
| TAAGGCCGAATCAGTAGTTG | ARMCX2 | 9823 |
| TAAGGCGACCTGCGCTTGTG | NO_CURRENT_789 | NO_CURRENT_789 |
| TAAGGTGCCAGAGTGTCAAG | NO_CURRENT_790 | NO_CURRENT_790 |
| TAAGTATAATGGAAAAGATG | PDHA1 | 5160 |
| TAAGTCTAATATATCCAAAG | NO_CURRENT_791 | NO_CURRENT_791 |
| TAATAAAGTGAAGCAGCACT | TMEM70 | 54968 |
| TAATAAATCCAGTTGATCTG | CFLAR | 8837 |
| TAATAAGCCAAAGCATGCGT | NO_CURRENT_792 | NO_CURRENT_792 |
| TAATAATAACTACCCAAACC | NO_CURRENT_793 | NO_CURRENT_793 |
| TAATAATAGTTGCCTTAATG | NO_CURRENT_794 | NO_CURRENT_794 |
| TAATACGCTGCGCCACGCCG | SLC25A19 | 60386 |
| TAATATGATTTGAGCACCAG | NO_CURRENT_795 | NO_CURRENT_795 |
| TAATATTGATTGTTAAAATG | NO_CURRENT_796 | NO_CURRENT_796 |
| TAATCATGCACATTCGGGAC | NO_CURRENT_797 | NO_CURRENT_797 |
| TAATCCATTCGATCTCACCA | CAT | 847 |
| TAATCCATTTAATCCCAATG | NO_CURRENT_798 | NO_CURRENT_798 |
| TAATGAGAGCAAGACGTGTG | CASP1 | 834 |
| TAATGCGAAAGGGGATACGA | TBK1 | 29110 |
| TAATGCTGCACACGCCGAAT | NO_CURRENT_799 | NO_CURRENT_799 |
| TAATGTGTGATTTGGCCAGT | GBP5 | 115362 |

| | | |
|---|---|---|
| TAATGTTGGATTAACCATAG | NNT | 23530 |
| TAATTAATCAGGCTTCACAA | CREBBP | 1387 |
| TAATTAGCTGCATAACTCCA | SLC7A11 | 23657 |
| TACAACAGATCGTGTAATGA | HDAC2 | 3066 |
| TACAACCAATGTGCTACGGT | CYP2R1 | 120227 |
| TACAAGTGTTCCGTGAAGCA | PHF8 | 23133 |
| TACAATCGTCTTGGCTATAC | NO_CURRENT_800 | NO_CURRENT_800 |
| TACACAGCCTGTAGTAGTGG | GBP7 | 388646 |
| TACACCCTGGACCTGACAGG | AHCTF1 | 25909 |
| TACAGTCACCAAAACAGTTT | NO_CURRENT_801 | NO_CURRENT_801 |
| TACAGTTGTGGAATGAGGTA | NMU | 10874 |
| TACATACCAGTGTCTCAATG | GLMN | 11146 |
| TACATAGTTTCTACAAACCA | NO_CURRENT_802 | NO_CURRENT_802 |
| TACATATGTTACCAGAATGG | SLC6A4 | 6532 |
| TACATCAAGACCCCAGTTCG | MEX3B | 84206 |
| TACATCACCAGGAGGGATGG | GPT2 | 84706 |
| TACATTACTTACGTAATAGA | MEMO1 | 51072 |
| TACATTATAAAAGACACTGT | NO_CURRENT_803 | NO_CURRENT_803 |
| TACCAAAATGTGGCCAATCG | ATP5L | 10632 |
| TACCAATAAAATAATCAACC | SEC62 | 7095 |
| TACCAGAACCTGTCTCCAGT | MAPK14 | 1432 |
| TACCAGCTCAGTAGGAGGTG | IRGM | 345611 |
| TACCATCATACTGATGCCTA | ABCB10 | 23456 |
| TACCATGAGACATGAACACC | CASP1 | 834 |
| TACCCCATTAGGTATTAAGC | NO_CURRENT_804 | NO_CURRENT_804 |
| TACCCTCCGGATACGGACTG | NO_CURRENT_805 | NO_CURRENT_805 |
| TACCCTGGATTGTCCTTGCG | NO_CURRENT_806 | NO_CURRENT_806 |
| TACCTGAGTGAATTCCACGT | NOS2 | 4843 |
| TACCTTCACTGTGATCCCAG | INO80B | 83444 |
| TACCTTCTGATTCGCCGCAG | CCL20 | 6364 |
| TACGCTTGCGTTTAGCGTCC | NO_CURRENT_807 | NO_CURRENT_807 |
| TACGTCATTAAGAGTTCAAC | NO_CURRENT_808 | NO_CURRENT_808 |
| TACGTCGGCTTAAAGAATCA | USP7 | 7874 |
| TACTACCTGAAGGAGAAGCA | MAP2K7 | 5609 |
| TACTAGTATTGAAAATCAGT | TLR2 | 7097 |
| TACTCAGACAGCTTAATGCT | NO_CURRENT_809 | NO_CURRENT_809 |
| TACTCATACACAAAAATCAG | NO_CURRENT_810 | NO_CURRENT_810 |
| TACTCCAGAAGATAAACCAT | NUP153 | 9972 |
| TACTCTGAACTAGCATCAAC | ATP5J | 522 |
| TACTCTTCACAGCATCATAT | COX16 | 51241 |
| TACTCTTCACAGCATCATAT | SYNJ2BP-COX16 | 100529257 |
| TACTGCAACTATGCCAGCAA | IKZF5 | 64376 |
| TACTGGAGTTTGCGACTCGG | NO_CURRENT_811 | NO_CURRENT_811 |
| TACTTACCAAACAATCCCTT | MAPK9 | 5601 |
| TACTTACTGCACTAGTCCCA | PRKN | 5071 |
| TACTTGTAATCTGCAACCAC | PPARG | 5468 |
| TAGAAGAAAACTGACCTGGG | POU2F1 | 5451 |
| TAGAATTTGACCAAAGGCAC | NO_CURRENT_812 | NO_CURRENT_812 |
| TAGACAACCGCGGAGAATGC | NO_CURRENT_813 | NO_CURRENT_813 |
| TAGACAGGGACATACCAAGG | ANKRD50 | 57182 |

| | | |
|---|---|---|
| TAGACTATACAGAATACATG | NO_CURRENT_814 | NO_CURRENT_814 |
| TAGAGATATCCGATCGTGGT | NO_CURRENT_815 | NO_CURRENT_815 |
| TAGAGCCGCTGGATTTCGGA | PAGR1 | 79447 |
| TAGAGCGGGTCATGAATCAC | ALOX5 | 240 |
| TAGAGTGCATAAGAGAACCA | NO_CURRENT_816 | NO_CURRENT_816 |
| TAGAGTGGAAGAAACTGGGT | JAM2 | 58494 |
| TAGATCGAGTTTATTTTCCT | NO_CURRENT_817 | NO_CURRENT_817 |
| TAGATCTGCAACTCGCTGGA | SAP18 | 10284 |
| TAGATCTTTGCTGCAACACA | CS | 1431 |
| TAGATGTTAACAGAGTGATC | NO_CURRENT_818 | NO_CURRENT_818 |
| TAGATTCATTCAGGCGAGCT | DNTTIP1 | 116092 |
| TAGCACCAGAGCACCCCGGA | VDAC1 | 7416 |
| TAGCCGGGCCTTCTAAGCCC | SPCS1 | 28972 |
| TAGCCTGAAGCAGACTGTAA | NUFIP2 | 57532 |
| TAGCTATACATATTACAACG | NO_CURRENT_819 | NO_CURRENT_819 |
| TAGGACTGATCCGCTCCCCA | DNAJA1 | 3301 |
| TAGGACTTGAATGTACGTTG | PIAS1 | 8554 |
| TAGGAGCTGTATCTAGTGGC | NO_CURRENT_820 | NO_CURRENT_820 |
| TAGGAGGAATGGTATACTAG | NO_CURRENT_821 | NO_CURRENT_821 |
| TAGGAGGCAAAACGTCAGAG | PLD1 | 5337 |
| TAGGCAAAGATCCGTTCTGT | CCNC | 892 |
| TAGGCCGGATAATGTCAGGG | CYP4F11 | 57834 |
| TAGGCTGACCCACCCCAATG | IKBKB | 3551 |
| TAGGCTGCAAGCATGTTGTG | PHIP | 55023 |
| TAGGGGATTAGCTGACAGTC | NO_CURRENT_822 | NO_CURRENT_822 |
| TAGGGGCCAAGAGTTTGTGT | NO_CURRENT_823 | NO_CURRENT_823 |
| TAGGTGTTGATACGAGCCCA | SATB1 | 6304 |
| TAGTAATTATTTGTTAACCG | STRAP | 11171 |
| TAGTACAAAATACTGATGCA | NO_CURRENT_824 | NO_CURRENT_824 |
| TAGTACATGTGTGGTATTTA | NO_CURRENT_825 | NO_CURRENT_825 |
| TAGTAGAGCTCTCCAATCAC | RND3 | 390 |
| TAGTAGGTACGCTAAAGGCA | NO_CURRENT_826 | NO_CURRENT_826 |
| TAGTATGGAATATATTCCTG | STRAP | 11171 |
| TAGTATTTATTAGTGACCAC | NO_CURRENT_827 | NO_CURRENT_827 |
| TAGTCAACATTCGCAAGAGG | NO_CURRENT_828 | NO_CURRENT_828 |
| TAGTCATTAGGAACTGTTGG | ABI1 | 10006 |
| TAGTCCTTAGGGTGGGCTGA | NO_CURRENT_829 | NO_CURRENT_829 |
| TAGTCGCGATAGGCCTTCCA | CYP24A1 | 1591 |
| TAGTCTCAGTAGATACCATG | NO_CURRENT_830 | NO_CURRENT_830 |
| TAGTGGCTGAGCTTCATCAA | SLC6A4 | 6532 |
| TAGTGGGAATGGTCGCGTAG | NO_CURRENT_831 | NO_CURRENT_831 |
| TAGTGTACTTGATAAACAGG | DET1 | 55070 |
| TAGTGTTCTGGCAGTGTCTA | BCL2A1 | 597 |
| TAGTTATACGATTAAAGCGA | SHOC2 | 8036 |
| TAGTTCTAATCGTTCCTTGA | NO_CURRENT_832 | NO_CURRENT_832 |
| TAGTTTAGTAGTAGAACCCA | CHUK | 1147 |
| TATAAACAGAAAATTATTCG | PDHA1 | 5160 |
| TATAACATCATATACCACAT | NO_CURRENT_833 | NO_CURRENT_833 |
| TATAACATCGGTATCTGGGT | RHOA | 387 |
| TATACAGCTGCGTAAAGTGT | CHUK | 1147 |

| | | |
|---|---|---|
| TATACCAGACCACAGCGCCG | NO_CURRENT_834 | NO_CURRENT_834 |
| TATACGAACTAACCTCCATG | NIPBL | 25836 |
| TATACTGCGGATCAATCTGA | NO_CURRENT_835 | NO_CURRENT_835 |
| TATAGCTGTTTCGAAGGCGC | NO_CURRENT_836 | NO_CURRENT_836 |
| TATATACAACCCATCGGTGA | NDUFA12 | 55967 |
| TATCAATCGTCCGGGTCACT | NO_CURRENT_837 | NO_CURRENT_837 |
| TATCAGTAGAGGTAAACTCT | NO_CURRENT_838 | NO_CURRENT_838 |
| TATCATTAGACTCCCACACT | NO_CURRENT_839 | NO_CURRENT_839 |
| TATCCAGGAGTTGATCACAG | NLRC4 | 58484 |
| TATCGAATGGAACAGCCTGA | ABL2 | 27 |
| TATCGCCGTTCACTGGCCTG | ABCB10 | 23456 |
| TATCGCTTCCGATTAGTCCG | NO_CURRENT_840 | NO_CURRENT_840 |
| TATCGGGACACTCTTGGTCA | MAP2K5 | 5607 |
| TATCTCCAAACTCATGAACA | SOD1 | 6647 |
| TATCTCCAGATCAACATCTG | METTL3 | 56339 |
| TATCTCGAACAACTACACTG | KPNB1 | 3837 |
| TATGAAAAAGCAGAGCGACT | SERF2 | 10169 |
| TATGAACGGATACATGCCTT | CD84 | 8832 |
| TATGACAAATATGGCAAAGA | DNAJB6 | 10049 |
| TATGACCCTGTTACATTGCC | NO_CURRENT_841 | NO_CURRENT_841 |
| TATGACTGCTATTTCTGACG | MGA | 23269 |
| TATGAGACATCCCCACTGCA | PPARG | 5468 |
| TATGCCAAAGATGTTGATCA | CHUK | 1147 |
| TATGCCAAGAACTTCTACGG | PDHA1 | 5160 |
| TATGCTCTTACCGTATTATG | SLC7A11 | 23657 |
| TATGGATCGCCCAAGAGTTG | BCL2L11 | 10018 |
| TATGGCTCCTAATATTCGGG | NO_CURRENT_842 | NO_CURRENT_842 |
| TATGGGCAACAGATGATGGG | PHIP | 55023 |
| TATGGGGGTACAATTAATGT | NO_CURRENT_843 | NO_CURRENT_843 |
| TATGGTCAATGAATTCAAGT | HIF1A | 3091 |
| TATGTAACAATTAGTAAACC | NO_CURRENT_844 | NO_CURRENT_844 |
| TATGTGAAACACTTCCGTAG | ACOD1 | 730249 |
| TATGTTGTCGTTGATGCTGC | MOSPD1 | 56180 |
| TATTAGACGTGTCTTACCAG | EZH2 | 2146 |
| TATTATGTGGACCTCATCGG | APEH | 327 |
| TATTCATAGATCTACTGACA | SATB1 | 6304 |
| TATTGAACGCACCTTAAGTG | PPARGC1A | 10891 |
| TATTGACACTCCCATATCAG | PDHB | 5162 |
| TATTGACCTAGGATATGCCA | IKBKB | 3551 |
| TATTGATGACAACAACAAGA | PIP4K2A | 5305 |
| TATTTACTAATTGGGTCGGG | PDK4 | 5166 |
| TATTTATGTATGGAGCAGCA | HSP90B1 | 7184 |
| TATTTGCAGCTCGTTTGTGG | STAT1 | 6772 |
| TATTTTGACTTGACGCAGGC | NO_CURRENT_845 | NO_CURRENT_845 |
| TCAAAAAAGTCCATTTGCCC | RRAGC | 64121 |
| TCAAACAGTAATTCAGAAGT | IRAK4 | 51135 |
| TCAAACGCATACATATAATG | NO_CURRENT_846 | NO_CURRENT_846 |
| TCAAAGAGTCCATCAAACAC | HDAC2 | 3066 |
| TCAAAGTCTTGCACGCCCAT | TBXAS1 | 6916 |
| TCAAATAGAGACAAGGAATG | PHF6 | 84295 |

| | | |
|---|---|---|
| TCAAATATGGGTAACACTTG | NO_CURRENT_847 | NO_CURRENT_847 |
| TCAAATCAGAATGGGACAGG | PPID | 5481 |
| TCAAATCTTCCAGACTCCTG | CD84 | 8832 |
| TCAACACTACCAGCTACCTG | IKBKE | 9641 |
| TCAACTTCTGCATTGAACCG | PDE5A | 8654 |
| TCAACTTGAACACCTTCCAA | SHOC2 | 8036 |
| TCAAGCTGATGTTAACATCG | CDKN2C | 1031 |
| TCAAGGCACAATATCAGCAG | KPNB1 | 3837 |
| TCAATCACCCAGAAACAATG | NO_CURRENT_848 | NO_CURRENT_848 |
| TCAATCAGACACTGTAGCAG | KPNB1 | 3837 |
| TCAATGGGAGAATCACCCCA | ACACA | 31 |
| TCAATGTGATAAGCCCAACA | NDUFA8 | 4702 |
| TCAATTAAATCTGACGTCTG | ABL2 | 27 |
| TCAATTCTCACTCACGACCA | NO_CURRENT_849 | NO_CURRENT_849 |
| TCACAAAGGTGATTCTTCCC | BTN2A2 | 10385 |
| TCACAACCAATTGTTGCTCC | PHF6 | 84295 |
| TCACAATGAGAGGCACAGTG | ATP6V0E1 | 8992 |
| TCACAGAGATACAGAAACAT | CYBB | 1536 |
| TCACAGATGGCAACATGCCA | MLF2 | 8079 |
| TCACATCTAGTGGTATCCTG | RAC1 | 5879 |
| TCACCACCATCTTACTCACC | HTR1D | 3352 |
| TCACCAGCCATCTTGCCTAG | TIRAP | 114609 |
| TCACCAGGACTGGGCTAACC | BRK1 | 55845 |
| TCACCTCCGAACGAACACCT | NO_CURRENT_850 | NO_CURRENT_850 |
| TCACGTTTCCAATTAGGCAC | NO_CURRENT_851 | NO_CURRENT_851 |
| TCACTCCACTCACCAACATG | COX16 | 51241 |
| TCACTGCCAATTTAATCACA | NO_CURRENT_852 | NO_CURRENT_852 |
| TCACTTGTCTATACGGGGTA | NO_CURRENT_853 | NO_CURRENT_853 |
| TCACTTTCTATCCTATACTG | NO_CURRENT_854 | NO_CURRENT_854 |
| TCAGAACTTCTGTTGAACTG | UTY | 7404 |
| TCAGAATGGGACCACAAACG | PDE3A | 5139 |
| TCAGAGCAAGGCTTCGCACG | PHF8 | 23133 |
| TCAGAGCAGCCAATGGCCAA | FBXW7 | 55294 |
| TCAGAGGTAGACACCGATCT | RBM6 | 10180 |
| TCAGAGTCCACACTAAGTCT | GBP5 | 115362 |
| TCAGATAGTACAGGTCGATG | ITGB2 | 3689 |
| TCAGATTCCGCAAGGGTCCA | NO_CURRENT_855 | NO_CURRENT_855 |
| TCAGCAAAGGACGAAACAAA | NO_CURRENT_856 | NO_CURRENT_856 |
| TCAGCAACCTCTAACTAACC | DYRK1A | 1859 |
| TCAGCACAGCATCCGAATCA | RND3 | 390 |
| TCAGCATGGTCAATGAACAC | NO_CURRENT_857 | NO_CURRENT_857 |
| TCAGCCAAAGATTCTCTCGA | NO_CURRENT_858 | NO_CURRENT_858 |
| TCAGCCGGCAAAACTGACAG | NR2C2 | 7182 |
| TCAGCCTCTTTATTACAGAG | NO_CURRENT_859 | NO_CURRENT_859 |
| TCAGCTGTAGATGTCTCCAA | RNF25 | 64320 |
| TCAGGAAATGATCCGCACAG | MTOR | 2475 |
| TCAGGAAATTACTACAATCA | PIGL | 9487 |
| TCAGGATTGGAGCGAGTGCG | BABAM1 | 29086 |
| TCAGGCTCTGAATAGCTCTA | ATP5L | 10632 |
| TCAGGCTCTGAATAGCTCTA | ATP5L2 | 267020 |

| | | |
|---|---|---|
| TCAGGGTGCGTGTTCACGAA | ITGB2 | 3689 |
| TCAGGTTGAAAAAACTGTCT | HSP90B1 | 7184 |
| TCAGGTTGGCCAAAGCACTA | ASH2L | 9070 |
| TCAGTAAAAGACACCATGCT | HSPA13 | 6782 |
| TCAGTCAGAAGCTGATCAAG | OXSM | 54995 |
| TCAGTCAGCAGTTATGCAGG | FOXO4 | 4303 |
| TCAGTCCCTTGCAGTTCCAG | LCN2 | 3934 |
| TCAGTGATAATGAAAACCCT | USP7 | 7874 |
| TCAGTGATGATATAGAACGG | ABL1 | 25 |
| TCAGTTATGTTCCTGTTAAT | UCHL5 | 51377 |
| TCATACACAGCTACGGGATA | JUNB | 3726 |
| TCATACAGAGATTTCCTCTG | NELFE | 7936 |
| TCATACCTAACTGGAGTGTG | RB1 | 5925 |
| TCATACCTATAGTATCTGGG | SHOC2 | 8036 |
| TCATACTGTCCCCATACATA | LTA4H | 4048 |
| TCATATACATACCTCCTCAG | HSH2D | 84941 |
| TCATATGAACTACAGAGACA | RBM6 | 10180 |
| TCATCAGACAAGGTTTGCTG | KIF2B | 84643 |
| TCATCAGAGACAAGTATGTG | BTN2A2 | 10385 |
| TCATCCAAAAACATGAATGG | NO_CURRENT_860 | NO_CURRENT_860 |
| TCATCCATATCGTCATGTGC | RAD21 | 5885 |
| TCATCCATGGAGACATCAAG | IRAK1 | 3654 |
| TCATCCCTGTAGATGTCATG | PARK7 | 11315 |
| TCATCTGTCAATCAACTACT | NO_CURRENT_861 | NO_CURRENT_861 |
| TCATCTTACATCTGGGAGAC | NO_CURRENT_862 | NO_CURRENT_862 |
| TCATGAAAAGAGGAGTACAC | APIP | 51074 |
| TCATGAGCCAACAGAGCAGT | PARK7 | 11315 |
| TCATGCTTGCTTGGGCAAAA | NO_CURRENT_863 | NO_CURRENT_863 |
| TCATGGTCTCAAGCTACATG | NO_CURRENT_864 | NO_CURRENT_864 |
| TCATGTGTAGCATTGTTCAA | CBLL1 | 79872 |
| TCATTAGGTGCTACACTACA | EIF4G3 | 8672 |
| TCATTATCTGGTGATAGACT | HDAC2 | 3066 |
| TCATTTACACCATTTCGCAA | IFNAR1 | 3454 |
| TCATTTGAACTCCAGGCCGA | PHIP | 55023 |
| TCATTTGCCGTGAATACAGT | ACTR5 | 79913 |
| TCCAACACTCACCTTCACCT | SLC3A2 | 6520 |
| TCCAAGCTGTCACACTCCAA | CXCR4 | 7852 |
| TCCAAGGCATACCCCAACCA | PTGIS | 5740 |
| TCCAATCGTAAAATCTACTG | NUP153 | 9972 |
| TCCAATGGGTAGCCATGAGG | ASH2L | 9070 |
| TCCAATTCACAAGAACCAGA | SLC3A2 | 6520 |
| TCCACAAAGAGTTTCTGTAT | ATP5J | 522 |
| TCCACAACCATAAGGGGAGA | INO80B | 83444 |
| TCCACCTCCAGAATTTGCAA | KPNA1 | 3836 |
| TCCACGCCACCAACAGTCAG | CXCR4 | 7852 |
| TCCACGTTATGATTTAGACG | TBK1 | 29110 |
| TCCACTCTGCTACAGCGACT | RND1 | 27289 |
| TCCAGAACTGTGTAATCGAT | ABI1 | 10006 |
| TCCAGAGAGTTCCAGCACAG | SQSTM1 | 8878 |
| TCCAGCACACGACACCACCT | CREBBP | 1387 |

| | | |
|---|---|---|
| TCCAGCCAGGGACTAGGCGT | IDH2 | 3418 |
| TCCAGCCTATACACCACTGA | SHOC2 | 8036 |
| TCCAGGGCGCCGTAGTCGTA | SOD2 | 6648 |
| TCCAGTGATGCCTGCCAACA | CYBB | 1536 |
| TCCATACCATGCAGTCCATG | LAMTOR2 | 28956 |
| TCCATCTAGAACACATTGAG | CHUK | 1147 |
| TCCATTATAAATAGAAACCG | BNIP3 | 664 |
| TCCCAAGGGTTTAAGTCGGG | NO_CURRENT_865 | NO_CURRENT_865 |
| TCCCAATTTAGGTCAGTTCA | NO_CURRENT_866 | NO_CURRENT_866 |
| TCCCACAGCCCATGTTCCAG | HAMP | 57817 |
| TCCCACTGGGAGATCAGATG | ZNF699 | 374879 |
| TCCCATTACAGGCTCAGTCG | STAT1 | 6772 |
| TCCCCCTGTGTCATACCTTG | TMEM173 | 340061 |
| TCCCCGAGACCATCTTAGGG | NO_CURRENT_867 | NO_CURRENT_867 |
| TCCCCTCTTTAAGGCATCGT | NDUFA12 | 55967 |
| TCCCGAAAACATTAAAACCC | EIF4G3 | 8672 |
| TCCCGCAAACGTGGATTACG | HSH2D | 84941 |
| TCCCGCATCAAGATCCAGCT | ATP5J2 | 9551 |
| TCCCGCATCAAGATCCAGCT | ATP5J2-PTCD1 | 100526740 |
| TCCCGCATCTGACAGCGCCG | BCORL1 | 63035 |
| TCCCGCGGTACCCCAGCGAA | LTB4R | 1241 |
| TCCCGGCTTAGAAGGACTGA | RIPK3 | 11035 |
| TCCCGGCTTGCCAGTCAGTG | MKL1 | 57591 |
| TCCCGGTTGGTGAACGATAC | NO_CURRENT_868 | NO_CURRENT_868 |
| TCCCGTATGAAGAGTGCAAG | NMRK2 | 27231 |
| TCCCTACAGACAGAGCCACA | BCL2L11 | 10018 |
| TCCCTGCATTCATGGTTTTA | NO_CURRENT_869 | NO_CURRENT_869 |
| TCCGGTCAAATATGCACCGG | ARHGAP33 | 115703 |
| TCCGTAAAGAGTCAGGTAGC | PTGIS | 5740 |
| TCCGTAATGTTGCTCGATGT | HDAC2 | 3066 |
| TCCGTGGGGAACACGAAAAC | MOSPD1 | 56180 |
| TCCGTGTACATCACACGGAG | HELZ2 | 85441 |
| TCCTACCTCCTTATCCAGAG | ARRB1 | 408 |
| TCCTAGAACTCAGTCAATCG | LRP2 | 4036 |
| TCCTATACACAGACAAACGG | PIP4K2A | 5305 |
| TCCTATACAGAAACTCTTTG | ATP5J | 522 |
| TCCTCACCTAAAGTGCAATA | NO_CURRENT_870 | NO_CURRENT_870 |
| TCCTCCAGCTGAAGTACCCT | ACTR5 | 79913 |
| TCCTCGAAGCCATCTACACA | SCN5A | 6331 |
| TCCTGAAAAGTCTCCACCCA | KIF2B | 84643 |
| TCCTGAATGTCTCTTACTTG | BABAM2 | 9577 |
| TCCTGACTTCAGAGCACGAG | ATP5J2 | 9551 |
| TCCTGATGGACAACTATGTG | HCAR2 | 338442 |
| TCCTGCCCACAGGCACAACG | RIPK1 | 8737 |
| TCCTGGACAGGAGCTATCCA | EDA2R | 60401 |
| TCCTGTATGTCTGAGAAACC | NUP153 | 9972 |
| TCCTTACCCCGACACCAGGT | PTGES2 | 80142 |
| TCCTTCCGTGTGGTGTCCGG | CYP1B1 | 1545 |
| TCGAACACCTTGTACCAGCT | PQBP1 | 10084 |
| TCGAACTGCCTCTCTGCGGA | TMEM70 | 54968 |

| | | |
|---|---|---|
| TCGACATACTTGGTCCAGGG | SCN5A | 6331 |
| TCGACCTGTACGATGTCCGA | CACNA2D2 | 9254 |
| TCGAGACCGGGAACGGGACA | NELFE | 7936 |
| TCGAGAGCCAACATATTCTG | HSPA13 | 6782 |
| TCGAGAGCGTGGCTATGACA | PQBP1 | 10084 |
| TCGAGAGGAAAAACACACTG | NO_CURRENT_871 | NO_CURRENT_871 |
| TCGAGATCCCAACACAAAGG | DNTTIP1 | 116092 |
| TCGAGTCGTGGTGCGTCCAG | NDUFB9 | 4715 |
| TCGATGAAGGCATAGCCACG | SNRNP70 | 6625 |
| TCGATGTAGCCCCGCCCAAG | NO_CURRENT_872 | NO_CURRENT_872 |
| TCGATGTGCACTTTCCAGTG | MEMO1 | 51072 |
| TCGCACCCGCCGCGGTACCG | SLC25A1 | 6576 |
| TCGCCACCCACTATTGGACC | ABI1 | 10006 |
| TCGCGGAAAACGGTGCGCAC | CYP1B1 | 1545 |
| TCGCGGTCGATCGGTTTCTC | SAP18 | 10284 |
| TCGCTTGACAGGATGTGTCC | DNTTIP1 | 116092 |
| TCGGAAAAAGACCTCTCGGG | BAX | 581 |
| TCGGAACCCTGGAAACGGAA | NDUFS4 | 4724 |
| TCGGACTTTCCATCGCCAAG | NLRC4 | 58484 |
| TCGGAGGAATCGTTTGACCA | TNRC18 | 84629 |
| TCGGATCTTTCTCTTCCCAG | NDUFA8 | 4702 |
| TCGGCAAATACCAATAATGA | NO_CURRENT_873 | NO_CURRENT_873 |
| TCGGCAGGCGAAGCATGAAG | VDR | 7421 |
| TCGGCATACGGGACACACGC | NO_CURRENT_874 | NO_CURRENT_874 |
| TCGGCCCGGCGAGAGATAGG | MCL1 | 4170 |
| TCGGCTACGGCGTGGAGAAG | NO_CURRENT_875 | NO_CURRENT_875 |
| TCGGCTGGCGCAACGTCACG | ITGB2 | 3689 |
| TCGGGACCGCGGGTATGACA | PQBP1 | 10084 |
| TCGGGCAGTGAGTACAATAC | NO_CURRENT_876 | NO_CURRENT_876 |
| TCGGGGACCACCCACGATCC | NO_CURRENT_877 | NO_CURRENT_877 |
| TCGGTACAGGTCACCTGAGA | PLD6 | 201164 |
| TCGGTACTGACCATGTGCAA | TNRC18 | 84629 |
| TCGGTCATAGCCATCCAAGT | ANP32B | 10541 |
| TCGGTCGACCCTCTCAGCAA | WASF2 | 10163 |
| TCGGTCTTCACCCCAAATCG | PDE4A | 5141 |
| TCGGTTACATGGCTGAGATG | SLC6A4 | 6532 |
| TCGTAAACACACGACCAAGT | NO_CURRENT_878 | NO_CURRENT_878 |
| TCGTACAAGTTGTCGGCCAG | SIRT1 | 23411 |
| TCGTCAAAAACAGCTTGACT | PPARGC1A | 10891 |
| TCGTCACTGTCGCAAACAGC | NDUFA8 | 4702 |
| TCGTCGGCTTCTTGGTGCCT | ATP6V0E1 | 8992 |
| TCTAACGAGACTGTTACACT | NO_CURRENT_879 | NO_CURRENT_879 |
| TCTAACGTAAAATCCTCCAG | NO_CURRENT_880 | NO_CURRENT_880 |
| TCTACGTGTAGTTGTACATA | NO_CURRENT_881 | NO_CURRENT_881 |
| TCTAGACTCAGATGAGAGTG | PDAP1 | 11333 |
| TCTAGGCTAGTAATGTCAGT | HSP90B1 | 7184 |
| TCTATCCCACTCCTATCAAG | NUP153 | 9972 |
| TCTATTACACAAATTGACCT | NO_CURRENT_882 | NO_CURRENT_882 |
| TCTATTTTGTCTGCGCAGAA | NO_CURRENT_883 | NO_CURRENT_883 |
| TCTGAAAAATAGGCCCAACC | NO_CURRENT_884 | NO_CURRENT_884 |

| | | |
|---|---|---|
| TCTGACCACACCAGAAAATC | ALOX15 | 246 |
| TCTGACCATTGGGTGCGACA | NO_CURRENT_885 | NO_CURRENT_885 |
| TCTGAGCAGACAGGTACCTG | METTL3 | 56339 |
| TCTGATCGTTGATCTTTCTG | NO_CURRENT_886 | NO_CURRENT_886 |
| TCTGATGGAACACTCGGACT | DET1 | 55070 |
| TCTGCATCAGTATAGAAACA | HSPA4 | 3308 |
| TCTGCCTCATAGGGTACCGG | HTR2A | 3356 |
| TCTGCGTACAGTGCTCCCGG | HELZ2 | 85441 |
| TCTGCTCTACCCTCTAATGG | PRR14L | 253143 |
| TCTGCTTCTAGAACACAGGC | AGER | 177 |
| TCTGGAAATTACTATCCAGT | GBP7 | 388646 |
| TCTGGAACTACAGATAAGGC | NO_CURRENT_887 | NO_CURRENT_887 |
| TCTGGACGCACCACGACTCG | NDUFB9 | 4715 |
| TCTGGCGCCATGTCGCAGAA | BNIP3 | 664 |
| TCTGGCGGGCGAGCTCACGC | SERF2 | 10169 |
| TCTGGCTTGACACGACCGTT | NO_CURRENT_888 | NO_CURRENT_888 |
| TCTGGGACAACATCACGTGC | VIPR2 | 7434 |
| TCTGGGGGAAAAGGTACACA | FOXO4 | 4303 |
| TCTGGTCAATCTCAATGCTG | MAP2K7 | 5609 |
| TCTGGTGATAAATGTCACAA | CYP24A1 | 1591 |
| TCTGGTTCTCCGTCCTGTCC | CHMP6 | 79643 |
| TCTGGTTTCATGATAGCAAG | SIRT1 | 23411 |
| TCTGTCAACAACACCCTGCT | BTN2A2 | 10385 |
| TCTGTGCAATCTTCTCATTG | MKL1 | 57591 |
| TCTGTGGGTCACAGTAACTA | CD84 | 8832 |
| TCTGTTCAGACTCTAATAGG | RAD21 | 5885 |
| TCTGTTCAGTGGTGCCAATG | DYRK1A | 1859 |
| TCTGTTGAAGGAGCGCCAAA | CCNC | 892 |
| TCTGTTGCTTATGTAACACC | CD84 | 8832 |
| TCTTAATTGTAGGTTAGTCG | PDHB | 5162 |
| TCTTACAATCTAAAGGCAGG | SUPT7L | 9913 |
| TCTTACCCGAAGTGTCCCAC | RND3 | 390 |
| TCTTACCTTATGTTATAAAG | PCGF6 | 84108 |
| TCTTACGGTAGACACGGACT | AGER | 177 |
| TCTTATCACTGTCAAAACTG | TLR1 | 7096 |
| TCTTATTAATTGAGGGACCT | NO_CURRENT_889 | NO_CURRENT_889 |
| TCTTCAATGGACATCACCTG | CHMP6 | 79643 |
| TCTTCAATTGGAATTACCTG | NMU | 10874 |
| TCTTCATAAATAATCAGACG | PAXIP1 | 22976 |
| TCTTCATTAAAATCGAAACA | PITPNB | 23760 |
| TCTTCATTGGCAGTAACCTG | RNF111 | 54778 |
| TCTTCTAGCTCATCAACTGA | RNF111 | 54778 |
| TCTTCTGGTAACCCATGACC | CXCR4 | 7852 |
| TCTTCTTTCTAGAACAACCG | NO_CURRENT_890 | NO_CURRENT_890 |
| TCTTCTTTGCGTGTTAGGTG | JAM3 | 83700 |
| TCTTGACAAAGGACGCCACA | IZUMO2 | 126123 |
| TCTTGATCATCTCTCCACTC | KHSRP | 8570 |
| TCTTGCCGGAATGTCAGCCG | EGFR | 1956 |
| TCTTGGAGGCATATCAACTG | CYP2R1 | 120227 |
| TCTTGTATAAAAAGTTACCA | LAMTOR3 | 8649 |

| | | |
|---|---|---|
| TCTTGTGGTGTCTGCGAGCG | BNIP3 | 664 |
| TCTTGTTTCTTATAGGCTGA | HMGN2 | 3151 |
| TCTTTACAACAGAAATCACC | PRKAA1 | 5562 |
| TCTTTACTGAATAAACAAGG | HSPA13 | 6782 |
| TCTTTCAAAAAGCATACAGT | GC | 2638 |
| TCTTTGTCTGCCACAAACGT | NR2C2 | 7182 |
| TGAAAACCCCAATAGACAGG | NO_CURRENT_891 | NO_CURRENT_891 |
| TGAAAAGCAGCAACTAATTG | NNT | 23530 |
| TGAAAATCCTATTGCTCAGT | NO_CURRENT_892 | NO_CURRENT_892 |
| TGAAACACGGACACTTCGGG | SQSTM1 | 8878 |
| TGAAACCCGAGAGGAGCGCA | SNRNP70 | 6625 |
| TGAAACTCTTACCCAGTGTT | VDAC1 | 7416 |
| TGAAAGAGACCGAATATACC | SATB1 | 6304 |
| TGAAAGTGATAGCGATACTG | IFNAR2 | 3455 |
| TGAAATATATGCAAAAATTG | NO_CURRENT_893 | NO_CURRENT_893 |
| TGAAATGAAACCGTGCCCTT | CSDE1 | 7812 |
| TGAACATGCGGCAGCGAAAG | NOD1 | 10392 |
| TGAACCCGTGTTGCTCTCCC | TGFB1 | 7040 |
| TGAACGGTGAAGAGATAGGG | NO_CURRENT_894 | NO_CURRENT_894 |
| TGAACGTGTCCACCTCAGCA | CYP4A11 | 1579 |
| TGAACGTGTCCACCTCAGCA | CYP4A22 | 284541 |
| TGAACTACTTACGAACTGCT | RB1 | 5925 |
| TGAACTGCTTCGTGGACGGA | FOXO4 | 4303 |
| TGAAGAAATACCTTAACACA | PPARGC1B | 133522 |
| TGAAGAACGAGCCAGATACC | UCHL3 | 7347 |
| TGAAGACACGCAGCATGTGT | MTOR | 2475 |
| TGAAGACCCGTAGCAACAGT | SAP18 | 10284 |
| TGAAGAGCATCAGCCGGCAA | HCAR2 | 338442 |
| TGAAGAGCTGATGTAAGATG | PIAS1 | 8554 |
| TGAAGCACAATACCAATACT | NO_CURRENT_895 | NO_CURRENT_895 |
| TGAAGCGAGACCCATCGTCC | NO_CURRENT_896 | NO_CURRENT_896 |
| TGAAGGTGTCTATACACTGT | NO_CURRENT_897 | NO_CURRENT_897 |
| TGAAGTAAGGAGAGACACGA | HTR2A | 3356 |
| TGAAGTCAAAGCCTTCTGCG | ARRB1 | 408 |
| TGAAGTCGGGAATAGTCAGC | CXCR4 | 7852 |
| TGAAGTTGGACAGGGCACGA | SCTR | 6344 |
| TGAATAAGTCCATCAGACAG | JAK1 | 3716 |
| TGAATACCAGACATTACTGG | BABAM2 | 9577 |
| TGAATCAGAGCGACATCGAG | NIPBL | 25836 |
| TGAATCGAATACAAACGATG | NO_CURRENT_898 | NO_CURRENT_898 |
| TGAATCGTAACCTCGCCATT | NO_CURRENT_899 | NO_CURRENT_899 |
| TGAATCTGGACGTTTAAGCT | MAP3K7 | 6885 |
| TGAATGACGTGGCATCACTG | TYK2 | 7297 |
| TGAATGCCACTGGCAATCGC | SLC3A2 | 6520 |
| TGAATGGTGCGCGTCGTAGG | IRF8 | 3394 |
| TGAATGTGAATGAGCTCCGG | GPT2 | 84706 |
| TGAATTTGTATGACCAAACA | APAF1 | 317 |
| TGACAAAGCCTCCAACGAGT | KIF2B | 84643 |
| TGACAACTACTGGTTTGGGA | CHEK2 | 11200 |
| TGACACATTGGCTGGGTGTT | NO_CURRENT_900 | NO_CURRENT_900 |

| | | |
|---|---|---|
| TGACAGAAGTAAAATGACTG | NCKAP1L | 3071 |
| TGACAGAGAGCGTGTGAACG | ARF1 | 375 |
| TGACAGATCGTGTTATTCAG | ACOD1 | 730249 |
| TGACAGGTTCAACTTACGTG | RNF25 | 64320 |
| TGACATCATTACGAGAGTGC | RIPK3 | 11035 |
| TGACATGGACTGCCTTGCAT | CXCR4 | 7852 |
| TGACATTACCAAATACTCCA | CAT | 847 |
| TGACCACCAGAGCGTCCCAG | SPHK1 | 8877 |
| TGACCTCTGAGGAATTCACA | NO_CURRENT_901 | NO_CURRENT_901 |
| TGACGATATATGCCTTCCGA | ANKRD50 | 57182 |
| TGACGCCCAAGGCCAACCGG | KAT2A | 2648 |
| TGACGCCGGACAGGTCATCA | GNAI2 | 2771 |
| TGACGCGATAGAGTTGGCTT | NO_CURRENT_902 | NO_CURRENT_902 |
| TGACGTATTATTCACAGTGC | UBE2E1 | 7324 |
| TGACTCGGGCAATATCGGTT | NO_CURRENT_903 | NO_CURRENT_903 |
| TGACTGACTGGAATAATGCT | PITPNB | 23760 |
| TGACTGTCAGCATCCCCCTG | UTY | 7404 |
| TGAGAAACACCAATTTCCGA | DYRK1A | 1859 |
| TGAGAACCCTGACTTAGCCA | NCKAP1L | 3071 |
| TGAGAATGTTAAGGGCACTG | TNFRSF1A | 7132 |
| TGAGACTACATGGCTCTGAG | SUPT7L | 9913 |
| TGAGAGAATGCATCACCATG | NO_CURRENT_904 | NO_CURRENT_904 |
| TGAGAGATACGATTGCTACA | NDUFB9 | 4715 |
| TGAGAGCCCCCCATCTTGTG | NO_CURRENT_905 | NO_CURRENT_905 |
| TGAGCACAGACTCACAGACG | PLD1 | 5337 |
| TGAGCACAGCTACATCGCTG | ACTR5 | 79913 |
| TGAGCACTGTTGTGACAGGA | PDHA1 | 5160 |
| TGAGCATGTCGGGAGTAACT | NO_CURRENT_906 | NO_CURRENT_906 |
| TGAGCATTCGTAGCCCAGCA | NO_CURRENT_907 | NO_CURRENT_907 |
| TGAGCCAGACATGATCAAGC | TBXAS1 | 6916 |
| TGAGCCAGCTGAGTTTCGAT | JAK1 | 3716 |
| TGAGCCCAGGCGTTGGCATG | ACADS | 35 |
| TGAGCCCATCCAAAGAGTGG | HTR7 | 3363 |
| TGAGCGGCCTCTAATTAATC | NO_CURRENT_908 | NO_CURRENT_908 |
| TGAGCGTTATACCCGACTCT | TRAF6 | 7189 |
| TGAGCTAACAGAGCTAATCA | GBP7 | 388646 |
| TGAGCTCATCTCCCGTCAGT | NOS2 | 4843 |
| TGAGCTGGCATCAAGGAGAG | JAK1 | 3716 |
| TGAGGACTGTCGCATCCCCA | ACADS | 35 |
| TGAGGATTGAGGTGTATGAA | NO_CURRENT_909 | NO_CURRENT_909 |
| TGAGGCCATCCTACGCCTCG | BTN2A2 | 10385 |
| TGAGGCTCAGATTAGTAACA | KPNA1 | 3836 |
| TGAGGTCTATTGATCTGCCG | CYFIP1 | 23191 |
| TGAGGTTGGGGTCTATGATG | RTP1 | 132112 |
| TGAGTACAAAGGCATTGGAG | HTR1D | 3352 |
| TGAGTCTACACGTATCATGC | CYP4B1 | 1580 |
| TGAGTCTTACTAGGTCCTGT | NO_CURRENT_910 | NO_CURRENT_910 |
| TGAGTGGAGAAGCACACACG | IFNAR2 | 3455 |
| TGAGTGGCAAAATATCACCA | SLC3A2 | 6520 |
| TGAGTGTGGACCCTAACACC | GSDMD | 79792 |

| | | |
|---|---|---|
| TGAGTTATTTAGTGAAAATG | CYP2R1 | 120227 |
| TGAGTTCTGAGCGGTTGGTC | YPEL5 | 51646 |
| TGAGTTGGGGCAGTAAACTG | NCOA6 | 23054 |
| TGAGTTTCTGTCCAGCCACG | RAD23A | 5886 |
| TGATACCTTCCTTGCACACA | PDE5A | 8654 |
| TGATAGGCAAAGATTACATG | ZRSR2 | 8233 |
| TGATCCAGGTAGGCACGAGT | DPY30 | 84661 |
| TGATGAAATACAATTTAAGG | PIAS1 | 8554 |
| TGATGAAATGACAGGCTACG | MAPK14 | 1432 |
| TGATGAGAGCCAGGCGTCTG | CASP5 | 838 |
| TGATGATGAGAAGTTCGTCT | IKBKE | 9641 |
| TGATGATTTCCTCAAGAACA | CYP4F11 | 57834 |
| TGATGCAACTCTGACATCTG | EDA2R | 60401 |
| TGATGCTGCCCGAGAAATTG | MPC2 | 25874 |
| TGATGGAAACGTGATAGTGT | PHIP | 55023 |
| TGATGGAAGAAATGCACTGC | ANKRD50 | 57182 |
| TGATGGAGTACTGCTCCAGT | IKBKE | 9641 |
| TGATGGATCACGCAGCAGTT | NMRK2 | 27231 |
| TGATGGATCTGAGTATAGTG | NFKB2 | 4791 |
| TGATGGCCTAGTGATGTCAT | NO_CURRENT_911 | NO_CURRENT_911 |
| TGATGGGTCCATGATAACCG | ACACA | 31 |
| TGATGTGCTGAAGCCCTATG | CYP4B1 | 1580 |
| TGATGTGGAATTCACAATCA | CSDE1 | 7812 |
| TGATTATCTCAATGTACTCC | BRK1 | 55845 |
| TGATTCCAGCACATTAATGG | EZH2 | 2146 |
| TGATTTCATAGCCAGCGATG | TBXAS1 | 6916 |
| TGATTTGGGCAAAATTCCTG | RRAGA | 10670 |
| TGCAAAACCTTGCAGAACAG | STAT1 | 6772 |
| TGCAACAGGTCATAAATACA | NO_CURRENT_912 | NO_CURRENT_912 |
| TGCAAGAGTTTCGTTTATGG | SPINT1 | 6692 |
| TGCAAGCGCATCAATCTTGG | ACTR5 | 79913 |
| TGCAATTAAATGATGACACC | HTR2A | 3356 |
| TGCAATTCTGGTTTGTTCTG | FAM107B | 83641 |
| TGCACAGTGGCAGTAACACT | NO_CURRENT_913 | NO_CURRENT_913 |
| TGCACCCACCGTAGTTATCG | SLC11A1 | 6556 |
| TGCAGAAAGTTCAAACTGAA | ZNF616 | 90317 |
| TGCAGAGGGACCTTGCTCAG | LCN2 | 3934 |
| TGCAGCAGTAAGCGGTTCCG | DLAT | 1737 |
| TGCAGCCCCAATGGCCATGA | OXSM | 54995 |
| TGCAGCGCCGGATCAACACA | ALOX5 | 240 |
| TGCAGGAAGCACCAGGACAT | SIAH2 | 6478 |
| TGCAGGAAGCATTTGATGGG | FAM214B | 80256 |
| TGCAGGATCCAAGGACATCT | NENF | 29937 |
| TGCATCGCGACATCAAGCCG | IKBKE | 9641 |
| TGCATGCCGAGCATTTTCAA | NO_CURRENT_914 | NO_CURRENT_914 |
| TGCCAATATCCAGATTGCAT | OXSM | 54995 |
| TGCCACAATAAGGTTGATTG | SNRNP70 | 6625 |
| TGCCAGCTACTGCGCCACCG | INO80C | 125476 |
| TGCCATAACTTAGAAACCGG | NO_CURRENT_915 | NO_CURRENT_915 |
| TGCCATAGCCAGTAACATCT | LAMTOR2 | 28956 |

| | | |
|---|---|---|
| TGCCATGGTACTTTGCGCTG | SLC25A19 | 60386 |
| TGCCATTATACTGTGAACAG | CSDE1 | 7812 |
| TGCCCAACAGTCATTCAACA | CHMP5 | 51510 |
| TGCCCCCGTTAGTGAACCTG | NDUFA1 | 4694 |
| TGCCGACAAAAGGATCAAGG | IDH2 | 3418 |
| TGCCGCGCCTTGCATGACCA | NLRP12 | 91662 |
| TGCCGGATAACATCAATGGT | LGMN | 5641 |
| TGCCGTACTCAATCCACCGA | KDM4A | 9682 |
| TGCCGTGAAAAGACGCTGCG | NO_CURRENT_916 | NO_CURRENT_916 |
| TGCCGTTAGCATGCGATCCC | NO_CURRENT_917 | NO_CURRENT_917 |
| TGCCTCTCCCTTACCCGGAC | NO_CURRENT_918 | NO_CURRENT_918 |
| TGCCTGAACAGCTTTACCGC | SPINT1 | 6692 |
| TGCCTTCAATTCAAGCACTC | INO80C | 125476 |
| TGCCTTTGTCACGTCCCCAG | ACO2 | 50 |
| TGCGACTACATGGCCCTCAA | PLD6 | 201164 |
| TGCGCCCGAACTCTTCGTTG | CYP1B1 | 1545 |
| TGCGCCCTCACCATCTGCGT | SPCS1 | 28972 |
| TGCGCGCTGAGCAAGCAGGA | TNFRSF1B | 7133 |
| TGCGGCCCTCAAAGGACCGG | TP73 | 7161 |
| TGCGTGCTGGATGTCGAGAG | APEH | 327 |
| TGCTAAGGCATTAATACTGG | SLC5A8 | 160728 |
| TGCTAAGGGAGATAAAGCAA | HMGN2 | 3151 |
| TGCTACCTTCGGGACCACCA | NO_CURRENT_919 | NO_CURRENT_919 |
| TGCTATTAGTGTGCACCTAG | NO_CURRENT_920 | NO_CURRENT_920 |
| TGCTCACTCCACTCCTCAAC | NO_CURRENT_921 | NO_CURRENT_921 |
| TGCTCATTTGAATATCAGCA | TLR1 | 7096 |
| TGCTCGCTCACTTGAAGTGT | GNAI2 | 2771 |
| TGCTCTGCATAATCAACCCT | GPT2 | 84706 |
| TGCTGAAGAAGCCCAAGAAG | TIRAP | 114609 |
| TGCTGAGTGCTGCACCAAAG | GC | 2638 |
| TGCTGCGTAGGCACTATGAT | EIF4G3 | 8672 |
| TGCTGGATAGGTTGCAGGCA | NO_CURRENT_922 | NO_CURRENT_922 |
| TGCTGGCACAGCACTATGTG | NO_CURRENT_923 | NO_CURRENT_923 |
| TGCTGGCACCAGAACGAATG | STAT1 | 6772 |
| TGCTGGGTCTAGCTTACCTG | CS | 1431 |
| TGCTGTACCAAGTGCCACAA | TNFRSF1A | 7132 |
| TGCTTCACCAGGGATAGGAA | ABL2 | 27 |
| TGCTTCCAGTTGTATCCAGT | SHOC2 | 8036 |
| TGCTTCCATACAGCTTAGAG | NO_CURRENT_924 | NO_CURRENT_924 |
| TGCTTGATGGACATCCACAG | SCN5A | 6331 |
| TGCTTTCTCATGTAGACATG | CYP2R1 | 120227 |
| TGGAAAAACAACATACGGTG | NUP153 | 9972 |
| TGGAAAAGGCGTGATACACA | RIPK1 | 8737 |
| TGGAAACATACTCGTCATCA | HTR2A | 3356 |
| TGGAAACGTTAACAATCCGG | TLR2 | 7097 |
| TGGAAAGAGTGGTTTCACAT | NO_CURRENT_925 | NO_CURRENT_925 |
| TGGAAAGCGAGCACACCGTC | NO_CURRENT_926 | NO_CURRENT_926 |
| TGGAAATAATCACCACAGGG | STAT2 | 6773 |
| TGGAACATGGGGATTCCGAG | GPR119 | 139760 |
| TGGAACCCACTCTTCCTACA | SPHK1 | 8877 |

| | | |
|---|---|---|
| TGGAAGTACCGGAGTATCTG | ALOX15 | 246 |
| TGGAAGTAGTGGCGCCACAG | ARF1 | 375 |
| TGGAATACCGACAATACACT | VDAC1 | 7416 |
| TGGAATGCCCGAGATTCCTC | SCTR | 6344 |
| TGGAATTGCATGCTTCAGGC | RTP1 | 132112 |
| TGGAATTGGGAAGTCAACAC | NLRP1 | 22861 |
| TGGACATCTCGGCGAAGTCG | BCL2 | 596 |
| TGGACCATTTAAGAGTTCGA | MMP13 | 4322 |
| TGGACCCAAAAGATCCAGTG | WASF2 | 10163 |
| TGGACCCACTCAAGAACCTG | TNRC18 | 84629 |
| TGGAGAAATGTACCCGCCCT | PTGS1 | 5742 |
| TGGAGAAGCTGTGTTGTATG | AHR | 196 |
| TGGAGAGGAACAGCCCATGG | OR7C2 | 26658 |
| TGGAGATAGTACCATGTGCA | KPNA1 | 3836 |
| TGGAGCGTCAGTATCAACTG | PTGS1 | 5742 |
| TGGAGCTTGATCAGAAACCC | NO_CURRENT_927 | NO_CURRENT_927 |
| TGGAGGAGAAGATCCACAAG | ARRB1 | 408 |
| TGGAGGCTCCGGCCACCAGG | NDUFS8 | 4728 |
| TGGAGTAGAGGATCTGCGTG | TRMT61A | 115708 |
| TGGAGTCAAGTGTAATGGCG | NO_CURRENT_928 | NO_CURRENT_928 |
| TGGATAACTCCCTGTCCCGG | NO_CURRENT_929 | NO_CURRENT_929 |
| TGGATACCGTTCAGTGCAAG | CYP8B1 | 1582 |
| TGGATCATCCCCGTCTTCGT | SLC7A5 | 8140 |
| TGGATGATTGCACTTCACTC | CYBB | 1536 |
| TGGATGGGTGAATTAACGGG | NO_CURRENT_930 | NO_CURRENT_930 |
| TGGATGTAAAAAGTCCAGGA | CAT | 847 |
| TGGATGTCCTCGATCATATG | CASP9 | 842 |
| TGGATTGTGTTGGAGAGTGT | ARNT | 405 |
| TGGATTTATCTACGGGTACG | SPCS1 | 28972 |
| TGGCAACTGGACGTTCCCCA | CREBBP | 1387 |
| TGGCAAGTTCCCATCTGCTG | IRGM | 345611 |
| TGGCAATTAGCACACAACAA | NO_CURRENT_931 | NO_CURRENT_931 |
| TGGCACTGACTAAAGAGATA | LTA4H | 4048 |
| TGGCACTGTGTCTCGCCATG | ARHGAP33 | 115703 |
| TGGCATCAATTGGGGCCGTG | BAK1 | 578 |
| TGGCATTACATTTATTGATC | NO_CURRENT_932 | NO_CURRENT_932 |
| TGGCCAAACCACTCTCTGTG | NOD2 | 64127 |
| TGGCCACGAATTCCGCCGCC | NO_CURRENT_933 | NO_CURRENT_933 |
| TGGCCAGCAATTCTAGACCA | NO_CURRENT_934 | NO_CURRENT_934 |
| TGGCCAGTGTGATGACGGAA | HTR1D | 3352 |
| TGGCCATCGCCAAGACGCCG | ADRB1 | 153 |
| TGGCCCACACGGTGCACACG | ADRB1 | 153 |
| TGGCCGCATTCCCCCCAATG | ACACA | 31 |
| TGGCCGTGAAAATTGCAGGG | PHIP | 55023 |
| TGGCGTAGGATGAGTCCATG | MKL1 | 57591 |
| TGGCGTAGGCCAGGTAATCG | ACADS | 35 |
| TGGCGTCAGGAGAGTTCTGG | RRAGA | 10670 |
| TGGCTACAATGCCACGACCC | NCKAP1L | 3071 |
| TGGCTACTATTCCCGTCTTG | TRIR | 79002 |
| TGGCTACTGCGTACATCCAC | NDUFA1 | 4694 |

| | | |
|---|---|---|
| TGGCTATGACCGAGAGGACC | ANP32B | 10541 |
| TGGGAAGGAATTGTCTGATG | CYP4F11 | 57834 |
| TGGGACTTTCATGATCCCCG | HSH2D | 84941 |
| TGGGAGATACGCACAGTCGA | NO_CURRENT_935 | NO_CURRENT_935 |
| TGGGAGCAGCACACCGTCTG | RAC2 | 5880 |
| TGGGAGCCCCCTCTGGACAA | CYP8B1 | 1582 |
| TGGGATTCAGCAATATGTGT | NO_CURRENT_936 | NO_CURRENT_936 |
| TGGGCAAAATCCGGGCCAGT | TBXAS1 | 6916 |
| TGGGCATCACATCCGGGACA | MLF2 | 8079 |
| TGGGCCAATGTGTACAACAG | PHC3 | 80012 |
| TGGGCCCCATCAAGTCACTG | SCN5A | 6331 |
| TGGGCCGCCTACGACCGGAA | LAMTOR2 | 28956 |
| TGGGCTACAGGACGATTCGA | HTR2A | 3356 |
| TGGGCTATAGATTCCATGTG | NO_CURRENT_937 | NO_CURRENT_937 |
| TGGGCTCCATGGCTGCACTG | RBM42 | 79171 |
| TGGGGAACCTCACTTCGTGG | ARNT | 405 |
| TGGGGACGTTTATCAATATA | NO_CURRENT_938 | NO_CURRENT_938 |
| TGGGGACTGGTCCCTTCTCG | NELFE | 7936 |
| TGGGGATGTCACCCATCACA | KAT2A | 2648 |
| TGGGGCCGGAATAGCCCGTG | PTGS1 | 5742 |
| TGGGGCTATGAAGAACGCAG | NELFE | 7936 |
| TGGGGCTGGACCCGGCAAAG | IRAK1 | 3654 |
| TGGGGTCTATTGATGAAGGC | NO_CURRENT_939 | NO_CURRENT_939 |
| TGGGGTCTTTATCCGCTCAG | EZH2 | 2146 |
| TGGGTAAAACATGCGTACAC | ACTR5 | 79913 |
| TGGGTAAATCAAGTTAACAC | NO_CURRENT_940 | NO_CURRENT_940 |
| TGGGTCAAGACTCAAACCAA | SERPINB2 | 5055 |
| TGGGTGAAGACACACACAAA | SERPINE1 | 5054 |
| TGGGTGGTCAGCGGTCTGAG | GNAI2 | 2771 |
| TGGTACAGAAATGACAGCAT | HSPA13 | 6782 |
| TGGTACTGAGGCAGACGTTG | NDUFS4 | 4724 |
| TGGTAGCGGGAAGGGATAGC | TRERF1 | 55809 |
| TGGTATCATATACGTGAATG | HIF1A | 3091 |
| TGGTCAACACTGACCCAGGT | NOD1 | 10392 |
| TGGTCAGGCTGGCGAGGAGG | HAMP | 57817 |
| TGGTGAATCTCCCGCTGCAC | BRK1 | 55845 |
| TGGTGAGCTTGGCGGCCGCA | CEBPD | 1052 |
| TGGTGAGGTTGACAACGCCG | NFKB2 | 4791 |
| TGGTGCTACACGAATCCCTG | HSPA4 | 3308 |
| TGGTGGCGAGACTCAGAACG | CYP24A1 | 1591 |
| TGGTGGTGGGTTTGTTATAT | NO_CURRENT_941 | NO_CURRENT_941 |
| TGGTGGTTGAAATCTAGCCA | CUBN | 8029 |
| TGGTGTATGATAATGCATCA | NO_CURRENT_942 | NO_CURRENT_942 |
| TGGTTAGCGATCTCTGGTCG | FOXO4 | 4303 |
| TGGTTGTGTAACTGATCAGT | RAC1 | 5879 |
| TGGTTTCTCCGTATGGTGCA | SIAH2 | 6478 |
| TGGTTTGGTAATTGACCTCG | STAT1 | 6772 |
| TGTAAATCATGGTCCACAAG | RIPK2 | 8767 |
| TGTAACCAAGATCATAAGAC | NO_CURRENT_943 | NO_CURRENT_943 |
| TGTACACGAAGGCCCGCGGA | SUV39H1 | 6839 |

| | | |
|---|---|---|
| TGTACTTCATCACTATCTCC | BCL2 | 596 |
| TGTACTTGTCTAGTGAATAC | ARR3 | 407 |
| TGTAGAGTTCATCTCTCTGC | LTB4R | 1241 |
| TGTAGAGTTTGACCCACCCC | TRAF6 | 7189 |
| TGTAGATATAGGGTGTCTAC | NO_CURRENT_944 | NO_CURRENT_944 |
| TGTAGATGGGGATACCAGGA | SERPINB2 | 5055 |
| TGTAGATGTTACAAGTTCCG | TLR2 | 7097 |
| TGTAGCACGTCACATAGTCG | SCTR | 6344 |
| TGTAGCTAAGTGAGTATGCC | NO_CURRENT_945 | NO_CURRENT_945 |
| TGTAGTCAATAAAACCCACC | PDE4A | 5141 |
| TGTCAAAGAATAGCAACAGA | MPC1 | 51660 |
| TGTCAATGTCAGTGGTACTG | NDUFS4 | 4724 |
| TGTCACCACATAATTACCTG | EGFR | 1956 |
| TGTCATAGTAAGTGCCAATG | EZH2 | 2146 |
| TGTCATCAATCGTGTTCAGA | EDA2R | 60401 |
| TGTCATGGAGTACGCCAACG | AKT1 | 207 |
| TGTCATGGGTCGAAAACTTG | PHF8 | 23133 |
| TGTCCAGGCTTAACTTCTCC | MCTS1 | 28985 |
| TGTCCATGAAACCACCCGAC | MLF2 | 8079 |
| TGTCCTCAAAACTAACAGAG | BABAM2 | 9577 |
| TGTCCTTCTAGGCCGCACTG | ALOX15 | 246 |
| TGTCGATGACCATCTTGCGT | PDE4A | 5141 |
| TGTCGATGCAGCAAACCTCA | CASP3 | 836 |
| TGTCGCAGAGGGGCTACGAG | BCL2 | 596 |
| TGTCTCTGCACACGGCTTCG | SPHK1 | 8877 |
| TGTCTCTGTTCTTGAACGTG | MYD88 | 4615 |
| TGTCTCTTGGTAGACCTTGG | MLF2 | 8079 |
| TGTCTGCTCTAGTAATAAGC | CTNNB1 | 1499 |
| TGTCTGGTACCATGTACTCA | PHF8 | 23133 |
| TGTCTTCCGAAATCTACCAG | SATB1 | 6304 |
| TGTGAAAACGAGCTCCACTG | PIP4K2A | 5305 |
| TGTGAAACAGATGTCAGCAA | OR7C2 | 26658 |
| TGTGAAAGTTTATGGTGTTA | NO_CURRENT_946 | NO_CURRENT_946 |
| TGTGAACCAATTCGCCGGGA | DNTTIP1 | 116092 |
| TGTGAAGCACTTATTGAACA | ANKRD50 | 57182 |
| TGTGACGTTATGTATGACCA | BTN2A2 | 10385 |
| TGTGAGAAGATTTAGTGACT | NO_CURRENT_947 | NO_CURRENT_947 |
| TGTGAGCTCAATGATGCCGC | MEX3B | 84206 |
| TGTGATAGGGTCATGTTCGT | STAT1 | 6772 |
| TGTGATCCAAGCTGTCCCAA | EGFR | 1956 |
| TGTGCACAAGTCGCAACGAA | NO_CURRENT_948 | NO_CURRENT_948 |
| TGTGCAGGTAATTTACACCA | RIPK2 | 8767 |
| TGTGCTGGATGTACCGAGCA | RNF25 | 64320 |
| TGTGGAAGAAAACTTCCCAC | PIGY | 84992 |
| TGTGGACAGTCGGACAAGTG | PIAS1 | 8554 |
| TGTGGACCACATCGACCTCG | ARRB1 | 408 |
| TGTGGCATTGAGACAGACAG | CASP3 | 836 |
| TGTGGGAAACAGTTTGTATG | ZNF641 | 121274 |
| TGTGGGGAACAGATCAACTA | HTR7 | 3363 |
| TGTGGTTCCACCTAATGTCA | CFLAR | 8837 |

| | | |
|---|---|---|
| TGTGTCACCTGCCAACGGTG | EDA2R | 60401 |
| TGTGTGACCTTCATTATATG | NO_CURRENT_949 | NO_CURRENT_949 |
| TGTGTTAGCCGAGATCTCTG | NO_CURRENT_950 | NO_CURRENT_950 |
| TGTGTTATCCCTGCTGTCAC | TGFB1 | 7040 |
| TGTGTTTCAGGGCTCTACAG | PIP4K2A | 5305 |
| TGTTAAAATCACCCGGTCTG | NO_CURRENT_951 | NO_CURRENT_951 |
| TGTTAATACAGGGTCCACAA | PRR14L | 253143 |
| TGTTACACATGAAGTTGTAC | TP73 | 7161 |
| TGTTAGCACTTACCTTTGCA | HMGN2 | 3151 |
| TGTTCAGGGTTCCGGCGTAG | CYP27B1 | 1594 |
| TGTTCCAAAACACTACGAAG | HSPA4 | 3308 |
| TGTTCTATAGAGCCGCGGTG | BRD1 | 23774 |
| TGTTCTTATTCCGGGAGGCG | MEX3B | 84206 |
| TGTTCTTCCAGTCAACAGCT | MAPK14 | 1432 |
| TGTTGAGGGGAGCCTCACGT | AKT1 | 207 |
| TGTTGCGCTCACCATCCAGC | HAMP | 57817 |
| TGTTGGTCGCGATAATTGGG | RBM42 | 79171 |
| TGTTGTACTTGAGTGACAGG | TLR9 | 54106 |
| TGTTTCAGGCCGTCACTATG | NR2C2 | 7182 |
| TGTTTCCATACTGCATGACG | LGMN | 5641 |
| TGTTTGGATGGTAAGCCTGG | CXCR1 | 3577 |
| TGTTTTGCATGTTGCATAGG | NO_CURRENT_952 | NO_CURRENT_952 |
| TTAAATATAGATTCTGCCAA | NO_CURRENT_953 | NO_CURRENT_953 |
| TTAACAATGTACCGTCTGGT | NO_CURRENT_954 | NO_CURRENT_954 |
| TTAACACAGTGTTAAGAACA | NO_CURRENT_955 | NO_CURRENT_955 |
| TTAACATCGAGGATAATGAA | CDKN2C | 1031 |
| TTAAGAAGCCTTCATAAGCG | CUBN | 8029 |
| TTAAGACATTGCGAATCGAT | NO_CURRENT_956 | NO_CURRENT_956 |
| TTAAGGCAATGAATTTCCAA | AHR | 196 |
| TTAAGTACTCAGGATCAACC | ARR3 | 407 |
| TTAATGAATGAAACCACTGC | HSPA4 | 3308 |
| TTAATGGGTAACTCTCTGAA | IZUMO2 | 126123 |
| TTAATTCCAAACCGGATGAG | PRKN | 5071 |
| TTACACAAGAAAACTTCTTG | CYBB | 1536 |
| TTACACCCATTATCACAGGG | NO_CURRENT_957 | NO_CURRENT_957 |
| TTACAGGAGTACAAACCACC | DYRK1A | 1859 |
| TTACAGTCTTACATGAGAGG | NO_CURRENT_958 | NO_CURRENT_958 |
| TTACCAGGGCCTACAAAAGG | RBM42 | 79171 |
| TTACCTGATATCTTTGATTG | PRKAA1 | 5562 |
| TTACCTGCACATATAGCAGT | ACTR5 | 79913 |
| TTACCTTCGAACTATAAGCG | CCAR2 | 57805 |
| TTACTCGACCGCTGGTTGCG | KIAA1211L | 343990 |
| TTACTGTATTCTACATGCCC | NO_CURRENT_959 | NO_CURRENT_959 |
| TTACTTAACAGGACATCTGG | ATP5J | 522 |
| TTACTTGAATTTCGTCACAG | SLC7A5 | 8140 |
| TTACTTTCTTGTTCGAGCAT | NO_CURRENT_960 | NO_CURRENT_960 |
| TTAGAGAGTCTTTGATAGCG | ALOX15 | 246 |
| TTAGCAGCAGTAGTTAGCAG | NUFIP2 | 57532 |
| TTAGCCCTCGATTGGTTGCG | NO_CURRENT_961 | NO_CURRENT_961 |
| TTAGCGTGGCTAGATCCACA | NLRP3 | 114548 |

| | | |
|---|---|---|
| TTAGGAATGTGATTGCCTTG | TBXAS1 | 6916 |
| TTAGGATAGGCCAATAACTG | UBE2H | 7328 |
| TTAGGATCGTTCAGTTGCCA | INO80C | 125476 |
| TTAGGCATCCAACGCACTGG | NMU | 10874 |
| TTAGGTGTAAGGGCTGCTTA | NO_CURRENT_962 | NO_CURRENT_962 |
| TTAGTATTAGTGACAAACAG | CSDE1 | 7812 |
| TTAGTGCAGGTCCTGAAACG | NDUFA12 | 55967 |
| TTATAACAGACTACTGTCTG | AHR | 196 |
| TTATAGGTGAATACTCGGGC | MAP2K5 | 5607 |
| TTATATAAATAACCAAACCA | NO_CURRENT_963 | NO_CURRENT_963 |
| TTATCCATGGGGAATTACAT | EIF2AK2 | 5610 |
| TTATCGTAGTAGGGAGATCC | RND1 | 27289 |
| TTATGAGAACGTCCGCGCCA | RAC2 | 5880 |
| TTATGAGAGTCAGAAATGAT | NO_CURRENT_964 | NO_CURRENT_964 |
| TTATGGCGCACTGTTTCGGG | MAP2K6 | 5608 |
| TTATGTACACAATTCTCCCG | SETD1B | 23067 |
| TTATTACAGTAGGGCCCCGT | CAT | 847 |
| TTATTCCCCCGGCTTGACTG | TP73 | 7161 |
| TTATTCCTTTGAGCACATTG | NO_CURRENT_965 | NO_CURRENT_965 |
| TTATTCGAAGACACAGAAGT | NMU | 10874 |
| TTATTGTAGAGTGTTGCTCA | NO_CURRENT_966 | NO_CURRENT_966 |
| TTCAACCAAAATACACCCGA | DYRK1A | 1859 |
| TTCAAGATAACGCTCCTGCT | HTR3E | 285242 |
| TTCAATCACCTCACGGTAAG | NO_CURRENT_967 | NO_CURRENT_967 |
| TTCACAACATGAAATCGCAC | NO_CURRENT_968 | NO_CURRENT_968 |
| TTCACACATACAATGCACTG | HIF1A | 3091 |
| TTCACAGATGCATAAGCCAG | NO_CURRENT_969 | NO_CURRENT_969 |
| TTCACCCTCGATACCTTGAG | FBXO38 | 81545 |
| TTCACCGTCCACGTGCGCAT | NO_CURRENT_970 | NO_CURRENT_970 |
| TTCACGTCGTCTCGCGACCA | NO_CURRENT_971 | NO_CURRENT_971 |
| TTCACGTTTGAGACCCAAGA | AIM2 | 9447 |
| TTCACTCTTGTCGACCAAGG | NO_CURRENT_972 | NO_CURRENT_972 |
| TTCACTTATTCTTGACAAAG | NO_CURRENT_973 | NO_CURRENT_973 |
| TTCAGAATGAGTCATATCAG | MMP13 | 4322 |
| TTCAGAGTATCAGCAAGAGC | ATP5J | 522 |
| TTCAGCAATAATGGGAACGG | BNIP3 | 664 |
| TTCAGCTGACATAATTCACA | MAPK14 | 1432 |
| TTCAGCTGGGACCACTTGCA | ARR3 | 407 |
| TTCAGCTTCATGCAGAAGCG | ARID3A | 1820 |
| TTCATGCAGCAGATCCAGAA | SERPINB2 | 5055 |
| TTCATGTCATAGATAACGAA | PPARG | 5468 |
| TTCCAAATTGCAGCTGACCT | ZRSR2 | 8233 |
| TTCCAATAGTACCAAATACT | NO_CURRENT_974 | NO_CURRENT_974 |
| TTCCACGACAACGCCAACGG | KHSRP | 8570 |
| TTCCACGGTAAAATCGGTCA | NO_CURRENT_975 | NO_CURRENT_975 |
| TTCCAGGGTGTATCGCAGGT | ALOX15 | 246 |
| TTCCAGTCCAGTTTGCTATG | CXCR1 | 3577 |
| TTCCATTGGCTGGAATCTGA | NO_CURRENT_976 | NO_CURRENT_976 |
| TTCCCACAGATACTGGTTGG | PTGIS | 5740 |
| TTCCCAGCGCCCTTCAATGG | MAP3K7 | 6885 |

| | | |
|---|---|---|
| TTCCCATAAGTATAATCTTG | NO_CURRENT_977 | NO_CURRENT_977 |
| TTCCCATCAATGCTAAATGG | RBM6 | 10180 |
| TTCCCATTGACTGAACACCG | BCORL1 | 63035 |
| TTCCCCACAGGCTTGCGGTG | ARRB1 | 408 |
| TTCCCGGAACAGGTAGCTCA | NDUFS8 | 4728 |
| TTCCCTAGGACTCTCACAGA | ZRSR2 | 8233 |
| TTCCGCCGCAGCGTCATCAA | NR1H3 | 10062 |
| TTCCGTTTCAGTACCAGTGA | ATP5J2 | 9551 |
| TTCCGTTTCAGTACCAGTGA | ATP5J2-PTCD1 | 100526740 |
| TTCCTAAGGAACTCTCCACA | APAF1 | 317 |
| TTCCTGCAATTCCCCAGGAG | ZNF641 | 121274 |
| TTCCTGCCCGAACTGCAGAA | NO_CURRENT_978 | NO_CURRENT_978 |
| TTCCTGTCTCTGTCGCAAAG | NOS2 | 4843 |
| TTCGAGGTCCGGACAGGTCG | NO_CURRENT_979 | NO_CURRENT_979 |
| TTCGAGTAAGTCACAGCAGC | ATP5L | 10632 |
| TTCGCAATAAATAATATACC | NO_CURRENT_980 | NO_CURRENT_980 |
| TTCGCACGATTGCACCTTGG | NO_CURRENT_981 | NO_CURRENT_981 |
| TTCGGAACTTACTCAGGGTA | NO_CURRENT_982 | NO_CURRENT_982 |
| TTCGGAATGATGAGCACACA | RHOA | 387 |
| TTCGGGGTAGACCTCCATAG | KMT2D | 8085 |
| TTCGGTATGATGTTGATTGT | NO_CURRENT_983 | NO_CURRENT_983 |
| TTCGTAGGAACTAAACTGTA | NO_CURRENT_984 | NO_CURRENT_984 |
| TTCGTGGTAGGTATAACTAT | NO_CURRENT_985 | NO_CURRENT_985 |
| TTCTAAGCGCCCTGGGGACA | NO_CURRENT_986 | NO_CURRENT_986 |
| TTCTACCGAAAGACACGCAT | EDA2R | 60401 |
| TTCTAGGAAGTCTACTGCCT | SERPINB2 | 5055 |
| TTCTCCTAATCCATGAACAG | NO_CURRENT_987 | NO_CURRENT_987 |
| TTCTCTGGTCATTCCTACAC | HSP90B1 | 7184 |
| TTCTCTGTTCCACAACACAG | FAM214B | 80256 |
| TTCTGAACAAGACGTTGACT | CTNNB1 | 1499 |
| TTCTGCGATTTAATTCCCGC | TFPT | 29844 |
| TTCTGCTCTGAGTGGAACTG | JAM2 | 58494 |
| TTCTGCTGTGCTTAAAGCTG | NDUFA8 | 4702 |
| TTCTGGTCGGTGGAGGTACA | UBE2H | 7328 |
| TTCTGGTTCCCCTAGAACAG | ARMCX2 | 9823 |
| TTCTGTAGGCAAACTAGGCG | IRF8 | 3394 |
| TTCTTAATCAGCGCTTGCAA | STRAP | 11171 |
| TTCTTACAAAAAACAACCGA | AHR | 196 |
| TTCTTCATGAACTCTGCTGA | MAPK9 | 5601 |
| TTCTTGAGGAGGAAGTAGCG | AKT1 | 207 |
| TTCTTGCATCTCAGTATGCA | CFLAR | 8837 |
| TTCTTGGTGCTCATGTACAG | IDH2 | 3418 |
| TTCTTGGTGGACGCATCCTG | BAX | 581 |
| TTCTTTAGCTGGTCAAAAGG | NO_CURRENT_988 | NO_CURRENT_988 |
| TTCTTTCAGCTCATAGATGG | LCN2 | 3934 |
| TTGAAACAGTTTCTAAGCTG | MSL2 | 55167 |
| TTGAAGTATCATCGGACCGA | CSDE1 | 7812 |
| TTGAATTTCTGAAGCGCATG | CYP8B1 | 1582 |
| TTGACAACAATGGGTCTAAG | PDE5A | 8654 |
| TTGACCTAAAATATGGGTGT | NO_CURRENT_989 | NO_CURRENT_989 |

| | | |
|---|---|---|
| TTGACTGACATGAATAAATG | NO_CURRENT_990 | NO_CURRENT_990 |
| TTGAGTTGAGTGATGATCAT | NO_CURRENT_991 | NO_CURRENT_991 |
| TTGATCAAAGAGGTGCACGT | NO_CURRENT_992 | NO_CURRENT_992 |
| TTGATGACGACGTGGTGTCA | USP7 | 7874 |
| TTGATGTCACCGGAGTTGTG | TGFB1 | 7040 |
| TTGCAAAGCTGATCGGCTGT | NO_CURRENT_993 | NO_CURRENT_993 |
| TTGCAATGCTGCTATAGAAG | NO_CURRENT_994 | NO_CURRENT_994 |
| TTGCACAGATCTGGGAGTAT | ATP5E | 514 |
| TTGCACCAGACTATATCCAA | INO80 | 54617 |
| TTGCAGAGTGACATATCTGC | NCOA6 | 23054 |
| TTGCAGCTCCAGAGTACCGT | PAGR1 | 79447 |
| TTGCCGCACAGAGCTATGGG | KDM4A | 9682 |
| TTGCCTCCCCATAGCCAATG | CYP4A11 | 1579 |
| TTGCCTGGTCCTCCTGACTG | IL10 | 3586 |
| TTGCGATACCAGCTGTAGTG | JAM3 | 83700 |
| TTGCGGTGCTGCAACCACTT | CYP4B1 | 1580 |
| TTGCGTGAAAAGCTAAACAT | SERPINB2 | 5055 |
| TTGCGTTTGCAGGTTACTAC | ATP5J2 | 9551 |
| TTGCTAGAAAATCCCAGACT | JAM2 | 58494 |
| TTGCTCAAAGGTGCCGAACG | HELZ2 | 85441 |
| TTGCTCCCTCGAAAAGAGCG | ABL1 | 25 |
| TTGCTGTGCAAAAGGGTCAT | RIPK1 | 8737 |
| TTGCTTCAGCCATTACAAAC | GLMN | 11146 |
| TTGGAAAACGAGCCAATGAG | STAG2 | 10735 |
| TTGGAAATCCCGAGATTGCC | CDKN2C | 1031 |
| TTGGAATGTTATTCAAGCCT | C4orf17 | 84103 |
| TTGGACACCAAACATCACTT | NO_CURRENT_995 | NO_CURRENT_995 |
| TTGGAGATGATAGTTTGGGG | NO_CURRENT_996 | NO_CURRENT_996 |
| TTGGAGCACCGATATAGATA | IFNAR1 | 3454 |
| TTGGAGCTGTCAGCGCCCCC | RRAGC | 64121 |
| TTGGAGTCTAAATCTCGAGG | MAP2K6 | 5608 |
| TTGGATTACTAGCTTTAGGT | NO_CURRENT_997 | NO_CURRENT_997 |
| TTGGCAAACATGAATTGACT | PRKAA1 | 5562 |
| TTGGCAATGCTGCCAAGCAT | ACOD1 | 730249 |
| TTGGCACAGATGTTGAACTG | SLC11A1 | 6556 |
| TTGGCCTATTTCAGTTAGCT | GLMN | 11146 |
| TTGGCTTGCAAATGTGCAGG | MBNL1 | 4154 |
| TTGGGAAGGACACCATTCCG | IL2RB | 3560 |
| TTGGGAGAAAGTTTCATCTG | RB1 | 5925 |
| TTGGGATCCTTAAATGGCAA | INO80C | 125476 |
| TTGGGCCTGAGCATCGAAGA | TYK2 | 7297 |
| TTGGGGCACCCACGCAAACG | AHDC1 | 27245 |
| TTGGTAAATGCATTGGACGC | SATB2 | 23314 |
| TTGGTAGTGTGTATAAACAC | NO_CURRENT_998 | NO_CURRENT_998 |
| TTGGTCCGAGTCTGGAGAAA | NO_CURRENT_999 | NO_CURRENT_999 |
| TTGGTTTAAGCTTAAAAGTG | NO_CURRENT_100( | NO_CURRENT_1000 |
| TTGGTTTACACAAAACCTTG | NO_CURRENT_100: | NO_CURRENT_1001 |
| TTGTAATAGAAGAAATTACG | NO_CURRENT_100: | NO_CURRENT_1002 |
| TTGTACCATTAACTTCACAC | TMEM30A | 55754 |
| TTGTAGCGGGTTCCTACCAC | SQSTM1 | 8878 |

| | | |
|---|---|---|
| TTGTATACAAATTAGTTCCA | BMI1 | 648 |
| TTGTATACAAATTAGTTCCA | COMMD3-BMI1 | 100532731 |
| TTGTATCTATCATCACCAGA | IRAK4 | 51135 |
| TTGTATGCCAAACCAGCTGG | TLR1 | 7096 |
| TTGTCAATGGGATATATGAG | NO_CURRENT_1003 | NO_CURRENT_1003 |
| TTGTCCCTGAGAAAACGCGG | NO_CURRENT_1004 | NO_CURRENT_1004 |
| TTGTCCGAGCTGAGGCAAGT | C6orf15 | 29113 |
| TTGTCCGTGACCCTGATTAA | NO_CURRENT_1005 | NO_CURRENT_1005 |
| TTGTCTATGAACATCTGTGG | NFKB1 | 4790 |
| TTGTGAAGTAATCTTAGGGT | MCTS1 | 28985 |
| TTGTGAATTTGGTTACCATG | NO_CURRENT_1006 | NO_CURRENT_1006 |
| TTGTGGCTCTGTCTGTAGGG | BCL2L11 | 10018 |
| TTGTGGGAAAACAGAGCCAC | HAMP | 57817 |
| TTGTGGGCACCTTACTGAGT | TLR1 | 7096 |
| TTGTGTGTAGATTATCTCAA | HDAC2 | 3066 |
| TTGTTAGCCCTACATGATCG | ASH2L | 9070 |
| TTGTTGTCATCTGTGAGCGG | DNTTIP1 | 116092 |
| TTGTTTCAACAGGGTTTATT | MPC2 | 25874 |
| TTGTTTCTTATAGGCTGAAG | HMGN2 | 3151 |
| TTTAAATTTGGAGATCAGGG | NUP153 | 9972 |
| TTTACACAACACTTCGATGG | CSNK1D | 1453 |
| TTTACACTTACCAGTAGAGT | LTA4H | 4048 |
| TTTACCGAGTAAAAACAGCA | DNAJA1 | 3301 |
| TTTACGAAGTATACCAGGTC | NO_CURRENT_1007 | NO_CURRENT_1007 |
| TTTAGACAATAAGTAGTCTC | IFNAR1 | 3454 |
| TTTAGCTTCTTCGATGGCGA | APIP | 51074 |
| TTTAGTAAATCTCATGACAC | SHOC2 | 8036 |
| TTTATAGTTATGCAATCCAA | GLMN | 11146 |
| TTTCAAGTTGGATGCATTGG | NFKB1 | 4790 |
| TTTCACACCCGCTACCACTG | PHC3 | 80012 |
| TTTCATAAGAGTGGGACACG | NO_CURRENT_1008 | NO_CURRENT_1008 |
| TTTCCCATGATCATTTAGTG | NO_CURRENT_1009 | NO_CURRENT_1009 |
| TTTCCTAATATTGTACAACT | APAF1 | 317 |
| TTTCCTATCTACACCTACAC | MMP13 | 4322 |
| TTTCGCAAGAACAAGACTCT | COX16 | 51241 |
| TTTCGCCCAAGAGGCTTGGG | NO_CURRENT_1010 | NO_CURRENT_1010 |
| TTTCGTGCCGATGTAACACA | NO_CURRENT_1011 | NO_CURRENT_1011 |
| TTTCTAGTTACTACTGGACG | NO_CURRENT_1012 | NO_CURRENT_1012 |
| TTTCTGCACCACCTTACTAA | RRAGC | 64121 |
| TTTCTGTGCTGAAAACCCAG | ARR3 | 407 |
| TTTCTTAAAACTTAAACCGG | NO_CURRENT_1013 | NO_CURRENT_1013 |
| TTTGAAGGACACCAAGACCA | CHMP5 | 51510 |
| TTTGAATAATTTCTAATAGC | GLMN | 11146 |
| TTTGAGACTCGAATTCAACG | NO_CURRENT_1014 | NO_CURRENT_1014 |
| TTTGAGCTGAGGCTTGGCAG | INO80B | 83444 |
| TTTGATTTACGTGTTTGACG | RRAGA | 10670 |
| TTTGCAAGTCACTATAATTG | NO_CURRENT_1015 | NO_CURRENT_1015 |
| TTTGCACACATACCAATCCA | CYP4A11 | 1579 |
| TTTGCAGGCATTCAAAAGTG | IKBKB | 3551 |
| TTTGCATGTGAAGAGATGTC | CHUK | 1147 |

| | | |
|---|---|---|
| TTTGCCATACAAGCTAACAT | NO_CURRENT_1016 | NO_CURRENT_1016 |
| TTTGCCTTTCGAGAGACAGG | MGA | 23269 |
| TTTGCTCCACATCTTAAAGG | RIPK1 | 8737 |
| TTTGCTCGGAAACTTGGAGA | MOSPD1 | 56180 |
| TTTGCTGTGGACACCCAAGT | APEH | 327 |
| TTTGGACGTCAGCATTACAA | CYP8B1 | 1582 |
| TTTGGATTCAAAGTGGGCAA | SEC62 | 7095 |
| TTTGGCAAAACGTCTTCAGG | AIM2 | 9447 |
| TTTGGCAAGTTAATCTCTAT | NO_CURRENT_1017 | NO_CURRENT_1017 |
| TTTGGCCTCCAGTACATGAG | NO_CURRENT_1018 | NO_CURRENT_1018 |
| TTTGGGAAGTGATAACGCGT | MAP3K7 | 6885 |
| TTTGGGTCACTAGGTCACGG | DNAJB6 | 10049 |
| TTTGGTCTTGATAGGCTCGA | MKL1 | 57591 |
| TTTGGTTACTCTGACAGTCA | PRR14L | 253143 |
| TTTGTAGAGATTGGTGACGG | C16orf72 | 29035 |
| TTTGTCGTGAAGATGCACAA | CREBBP | 1387 |
| TTTGTGAAATACTATTACAG | PHF8 | 23133 |
| TTTGTGTTCTGTGAGCGATG | ZNF699 | 374879 |
| TTTGTTTATGTTATGACGCA | NO_CURRENT_1019 | NO_CURRENT_1019 |
| TTTTACCTTGTTCACATGGA | NO_CURRENT_1020 | NO_CURRENT_1020 |
