## Supplementary material for "Illuminating host-mycobacterial interactions with functional genomic screening to inhibit mycobacterial pathogenesis": Table S5

**Table S5 Secondary CRISPRi target genes and sgRNA target sequences. Related to Figure 3.**

| Barcode Sequence | Gene Symbol | Gene ID |
|---|---|---|
| AAAAAACCCGTAGATAGCCT | NDUFA12 | 55967 |
| AAAAATCAGAAATAATCCAT | NO_CURRENT_1 | NO_CURRENT_1 |
| AAAACAAAAGGGAAACCCGG | FOXO4 | 4303 |
| AAAACAGCTACTATAATCCC | NO_CURRENT_2 | NO_CURRENT_2 |
| AAAACAGGACGATGTGCGGC | NO_CURRENT_3 | NO_CURRENT_3 |
| AAAACATCGACCGAAAGCGT | NO_CURRENT_4 | NO_CURRENT_4 |
| AAAAGAAGTCAGCTCTGGTT | RIPK2 | 8767 |
| AAAAGACATGTCCCCTCGCT | NO_CURRENT_5 | NO_CURRENT_5 |
| AAAAGGGACTCCCGGAGCTA | CFLAR | 8837 |
| AAAAGGGATAGAACCAGAGG | APAF1 | 317 |
| AAAATAGCAGTAAACTCAAC | NO_CURRENT_6 | NO_CURRENT_6 |
| AAAATATTCAAAATGGCGGA | POU2F1 | 5451 |
| AAAATCACTTAAATCACTCT | NO_CURRENT_7 | NO_CURRENT_7 |
| AAAATCCTTTCGCCAGATTA | NO_CURRENT_8 | NO_CURRENT_8 |
| AAAATGCGGGAATCTTAACG | IKBKB | 3551 |
| AAAATGGCGACGGACCGACC | DYRK1A | 1859 |
| AAAATGGCTGGTAAGCAGGC | UQCRB | 7381 |
| AAAATGGGATGGTGCTAACA | NO_CURRENT_9 | NO_CURRENT_9 |
| AAAATTAAAGCAGAGAGGAG | TNFRSF1A | 7132 |
| AAACAAAAATCCCTAGATTG | NO_CURRENT_10 | NO_CURRENT_10 |
| AAACAAAAGGGAAACCCGGG | FOXO4 | 4303 |
| AAACCAAGGTAAGCGCCGTA | ZRSR2 | 8233 |
| AAACCAGCTAGTTATTTACA | NO_CURRENT_11 | NO_CURRENT_11 |
| AAACCCATACCATTGACTCT | NO_CURRENT_12 | NO_CURRENT_12 |
| AAACCTCTGGGTTTCACAAG | NO_CURRENT_13 | NO_CURRENT_13 |
| AAACCTGGCAAATAGAGCCG | NO_CURRENT_14 | NO_CURRENT_14 |
| AAACGAGATCGAGAAAGGTA | NO_CURRENT_15 | NO_CURRENT_15 |
| AAACTCTATTTCCAAAACCA | NO_CURRENT_16 | NO_CURRENT_16 |
| AAACTGTAGTGCAGGGTCAG | NO_CURRENT_17 | NO_CURRENT_17 |
| AAAGAAAAGAGGTTTACACT | NO_CURRENT_18 | NO_CURRENT_18 |
| AAAGAAAAGGGCAAGGAGAA | SERF2 | 10169 |
| AAAGAAAGAGGAATAGTAGC | NO_CURRENT_19 | NO_CURRENT_19 |
| AAAGAACTGCATGGGAAAGT | HTR2A | 3356 |
| AAAGAGAAAGAAGTCGGTCT | PHF6 | 84295 |
| AAAGCAGTAGGATCGGCCAG | KDM4A | 9682 |
| AAAGCCCCATTGGAGTGAGG | DYRK1A | 1859 |
| AAAGCCGAGCTCTGGGCAGG | SEC62 | 7095 |
| AAAGCGACGTAGGCATACTT | NO_CURRENT_20 | NO_CURRENT_20 |
| AAAGGAGCAAGATCCATAGG | CD84 | 8832 |
| AAAGGAGCGTTCGCTAGCAG | LRP2 | 4036 |
| AAAGGAGGGGCGAAGGAGGA | OXSM | 54995 |
| AAAGGAGTATATGACACTAA | NO_CURRENT_21 | NO_CURRENT_21 |
| AAAGGGACTCCCGGAGCTAG | CFLAR | 8837 |
| AAAGGGATAGAACCAGAGGT | APAF1 | 317 |
| AAAGGTTGGGCAAAGTCAAG | NO_CURRENT_22 | NO_CURRENT_22 |
| AAAGTAGAAACAAACTCAGT | NO_CURRENT_23 | NO_CURRENT_23 |
| AAAGTAGCGCCAGTTACCTG | LAMTOR3 | 8649 |
| AAAGTCCTAGCAAACAGAGG | CXCR1 | 3577 |

| | | |
|---|---|---|
| AAAGTTCCAGGTGTGGGTGA | HTR1D | 3352 |
| AAATAAGAAAACTCTAAGGC | NO_CURRENT_24 | NO_CURRENT_24 |
| AAATAATATGCATCTCTCGA | NO_CURRENT_25 | NO_CURRENT_25 |
| AAATACAAGCTATAGCGATA | NO_CURRENT_26 | NO_CURRENT_26 |
| AAATACCTTTATCTGAACCC | NO_CURRENT_27 | NO_CURRENT_27 |
| AAATCAGCCGCCGACTCACT | BABAM1 | 29086 |
| AAATCCCATGCTCAGTTGGT | CUBN | 8029 |
| AAATCCTCATCGACAAGACC | RTP1 | 132112 |
| AAATGAGCATAATTTAACTG | NO_CURRENT_28 | NO_CURRENT_28 |
| AAATGATTAGCACAGTCATG | NO_CURRENT_29 | NO_CURRENT_29 |
| AAATGCACAGATCGCTGATC | NO_CURRENT_30 | NO_CURRENT_30 |
| AAATGCGGGAATCTTAACGC | IKBKB | 3551 |
| AAATGGCTGCCCGAGGGAGA | TRIR | 79002 |
| AAATGTATGGAATTATGTAG | NO_CURRENT_31 | NO_CURRENT_31 |
| AAATGTTCCTCAGACGCGGG | ZNF699 | 374879 |
| AAATTCCCCACCGGGCACCG | PTGS1 | 5742 |
| AAATTTAAAAGGTGAAGCAG | SUV39H1 | 6839 |
| AAATTTACTAAATCTGGGCA | NO_CURRENT_32 | NO_CURRENT_32 |
| AACACACACTGACAGCTATA | PHIP | 55023 |
| AACAGACAAGCAGGTTGTCT | TLR1 | 7096 |
| AACAGCACGTTACTTAACAG | NO_CURRENT_33 | NO_CURRENT_33 |
| AACAGCACTCCCAAAGAACT | CCL20 | 6364 |
| AACAGCGCGCTCGCAGCGGG | ACADS | 35 |
| AACAGGAAACGTGACTAAAG | NO_CURRENT_34 | NO_CURRENT_34 |
| AACAGGCGGAGGGTCGGCGT | FBXO38 | 81545 |
| AACATTACGTTATTTAGGCC | NO_CURRENT_35 | NO_CURRENT_35 |
| AACCAAGAGGGTGGGAAAGA | MKL1 | 57591 |
| AACCAAGGCCAAGCGAATAA | UBA7 | 7318 |
| AACCAAGGTAAGCGCCGTAC | ZRSR2 | 8233 |
| AACCAATACGACTCGATACG | NO_CURRENT_36 | NO_CURRENT_36 |
| AACCAATTCTCCTGTCGGCG | NIPBL | 25836 |
| AACCAGCGGTTACCATGGAG | CXCR4 | 7852 |
| AACCAGCTCTCCCGAAGCCG | EIF2AK2 | 5610 |
| AACCCTATTTCCCATAGACA | SERPINB2 | 5055 |
| AACCGACGGAGGACCGCGGG | RHOA | 387 |
| AACCGCTGGTTCTCCAGATG | CXCR4 | 7852 |
| AACCGGCCTTGGAACAACTG | CHUK | 1147 |
| AACGAGCAGAGCATCCAACA | EIF4G3 | 8672 |
| AACGCAGGAGACACAAGAAG | SEC62 | 7095 |
| AACGGGACCCACGGAACTAC | FAM214B | 80256 |
| AACTAGCCCGAGCAGCTTCG | NO_CURRENT_37 | NO_CURRENT_37 |
| AACTAGGCTCTCGGTGAGCT | NQO1 | 1728 |
| AACTAGTAATCATGACAACA | NO_CURRENT_38 | NO_CURRENT_38 |
| AACTCAGGTAAGATCAGCCA | IRF9 | 10379 |
| AACTCCAAATAAGCTGGCTG | NO_CURRENT_39 | NO_CURRENT_39 |
| AACTCGAGGCCTCACTGAAA | NLRC4 | 58484 |
| AACTGAAAGGTCTGGGGAGA | PAGR1 | 79447 |
| AACTGCCATTCACACAAATG | NO_CURRENT_40 | NO_CURRENT_40 |
| AACTTGTCAGCTCCTTGTCA | MBNL1 | 4154 |
| AAGAAAAGGGCAAGGAGAAG | SERF2 | 10169 |

| | | |
|---|---|---|
| AAGAACGAACGGCTTGGGCG | DPY30 | 84661 |
| AAGAACGAACGGCTTGGGCG | MEMO1 | 51072 |
| AAGAAGAATTGGGGATGATG | NO_CURRENT_41 | NO_CURRENT_41 |
| AAGACAAAACGTATCTTGGT | NO_CURRENT_42 | NO_CURRENT_42 |
| AAGAGATCACATCTAGGCCA | NO_CURRENT_43 | NO_CURRENT_43 |
| AAGAGATGACATTTGAGCTA | NO_CURRENT_44 | NO_CURRENT_44 |
| AAGAGCCGCGTGAGGAAACG | LTB4R | 1241 |
| AAGAGGCACAAAGAATGTAT | NO_CURRENT_45 | NO_CURRENT_45 |
| AAGAGGGATGCTAAGAGCAA | ARR3 | 407 |
| AAGAGTAGTAGACGCCCGGG | NO_CURRENT_46 | NO_CURRENT_46 |
| AAGATAGAGAAATCCATCAA | NO_CURRENT_47 | NO_CURRENT_47 |
| AAGATCGGCCACTACATTCT | PRKAA1 | 5562 |
| AAGATGACAGACTGTGAATT | BCL2A1 | 597 |
| AAGATGACTAATGCTAAGAG | NO_CURRENT_48 | NO_CURRENT_48 |
| AAGATGCCCGCGACGGAAAA | MBNL1 | 4154 |
| AAGATGGCGGAAAATTTAAA | SUV39H1 | 6839 |
| AAGATGGCGGGGAACGACTG | PPARGC1B | 133522 |
| AAGCACACTAAAGCCTCCAT | NO_CURRENT_49 | NO_CURRENT_49 |
| AAGCAGAGGGCCTTACCCGA | NELFE | 7936 |
| AAGCAGGCCGGTAAGTAACT | UQCRB | 7381 |
| AAGCAGTGGAGGCAGAGGCG | SUV39H1 | 6839 |
| AAGCAGTGGCGTCCGCAGCT | TRAF6 | 7189 |
| AAGCCATTGTATAACTCCAG | NO_CURRENT_50 | NO_CURRENT_50 |
| AAGCCCGGGTTCAGGCTCTC | FLCN | 201163 |
| AAGCCCGGGTTCAGGCTCTC | PLD6 | 201164 |
| AAGCCGGCGCGCCTGCAGCT | KIAA1211L | 343990 |
| AAGCGGAGGTCGGAGCGCCG | BNIP3 | 664 |
| AAGCGGAGTGAGTGACTAGA | PHC3 | 80012 |
| AAGCGGATCGGCGGCTCCTG | CASP9 | 842 |
| AAGCGGCGGCGGCGGCTGTA | RNF111 | 54778 |
| AAGCGGGAGAGCTGGCGGGG | CXCL3 | 2921 |
| AAGCTCAGGGAGAGGAAGCT | CYP4B1 | 1580 |
| AAGCTGGAGAGCGAACGAGC | IDH2 | 3418 |
| AAGCTTATACAACTTATGGG | NO_CURRENT_51 | NO_CURRENT_51 |
| AAGCTTGACTGGCGTCATTC | PPARGC1A | 10891 |
| AAGGAATAAAAATTTGTAGG | NO_CURRENT_52 | NO_CURRENT_52 |
| AAGGAGCCTGTGTGGCTCCC | C4orf17 | 84103 |
| AAGGAGGTGGCATGTCGGTC | TNFRSF6B | 8771 |
| AAGGCAGAAGCTCGCAGTGC | UTY | 7404 |
| AAGGCAGTCACCTTGACGAG | NO_CURRENT_53 | NO_CURRENT_53 |
| AAGGCCGAGGAGCTCAAGCC | MLF2 | 8079 |
| AAGGCGCGCGAATGTGGCAG | NO_CURRENT_54 | NO_CURRENT_54 |
| AAGGCTCTTCATAAATTCGT | NO_CURRENT_55 | NO_CURRENT_55 |
| AAGGGACTCCTGGTTTCCAC | HTR2A | 3356 |
| AAGGGAGGAGAGAAAAGCCA | CARD17 | 440068 |
| AAGGGAGGAGAGAAAAGCCA | CASP1 | 834 |
| AAGGGATAGAACCAGAGGTG | APAF1 | 317 |
| AAGGGATATAGGGCTTAGTA | ARR3 | 407 |
| AAGGGCCAGGAGAAGCTGCG | EBI3 | 10148 |
| AAGGGGACGCAGCGAAACCG | TP73 | 7161 |

| | | |
|---|---|---|
| AAGGGGTGTGGAGGCTGGTG | RAC2 | 5880 |
| AAGGTACAGAAATGGCAGTA | NO_CURRENT_56 | NO_CURRENT_56 |
| AAGGTACGTTGATGCCACTT | NO_CURRENT_57 | NO_CURRENT_57 |
| AAGGTCAGAGTTCGTGACCC | NR2C2 | 7182 |
| AAGGTGCAAAAATGCCCCAA | NO_CURRENT_58 | NO_CURRENT_58 |
| AAGGTGGGAGGGGAAGAGTG | TNFRSF1A | 7132 |
| AAGGTTGGGGAATCACTTGA | NO_CURRENT_59 | NO_CURRENT_59 |
| AAGTAGGGGATGGGGAGAAG | STRAP | 11171 |
| AAGTCAGAGGAAGTGGGTGA | MOSPD1 | 56180 |
| AAGTCAGGTGCTATTCACAA | NO_CURRENT_60 | NO_CURRENT_60 |
| AAGTGAATTAGCACAGAAGT | NO_CURRENT_61 | NO_CURRENT_61 |
| AAGTGACAGATGGGCAGGCG | NO_CURRENT_62 | NO_CURRENT_62 |
| AAGTGACAGCGGAGAGAACC | LAMTOR3 | 8649 |
| AAGTGATAAACAATTGGGAT | NO_CURRENT_63 | NO_CURRENT_63 |
| AAGTGGAGTATTTGCTGGGC | CTNNBL1 | 56259 |
| AAGTGTCCTGAGTCTCGAGG | TBK1 | 29110 |
| AAGTGTGTGCATAGCAGGGT | NO_CURRENT_64 | NO_CURRENT_64 |
| AAGTTAGCTTAATCCTAGCA | NO_CURRENT_65 | NO_CURRENT_65 |
| AAGTTCACATTAGAAGGGCA | NO_CURRENT_66 | NO_CURRENT_66 |
| AAGTTCCTGAGGAGATGTTC | PAXIP1 | 22976 |
| AATAATTACACATAGTGGGC | NO_CURRENT_67 | NO_CURRENT_67 |
| AATATATTACAACCGGCTGC | NO_CURRENT_68 | NO_CURRENT_68 |
| AATATCAAACGGGTGCACGT | NO_CURRENT_69 | NO_CURRENT_69 |
| AATATCTGATATAATAAATG | NO_CURRENT_70 | NO_CURRENT_70 |
| AATCAACAAACCAGCTTTCG | SUPT7L | 9913 |
| AATCAAGTAAAAGCCAACGC | NO_CURRENT_71 | NO_CURRENT_71 |
| AATCAATCCCTATATAACAG | NO_CURRENT_72 | NO_CURRENT_72 |
| AATCACACTCGTCAATGTTA | NLRC4 | 58484 |
| AATCACATCTTACATCCGAT | NO_CURRENT_73 | NO_CURRENT_73 |
| AATCAGCTATTGCTATGCTA | NO_CURRENT_74 | NO_CURRENT_74 |
| AATCAGGTACGGGCCCTGAG | STAT2 | 6773 |
| AATCATAACCTGGCAAGATG | NO_CURRENT_75 | NO_CURRENT_75 |
| AATCATGGATCACACTGGTA | CBLL1 | 79872 |
| AATCATGTGACGGTGGCTTG | LAMTOR3 | 8649 |
| AATCCAATTGCCCTCTTCTT | NLRC4 | 58484 |
| AATCCGCAGGAGCATAGCCA | CYP8B1 | 1582 |
| AATCGCGATGTTTGGAGTTG | BTN2A2 | 10385 |
| AATCTGCTGTAACTTATATC | NO_CURRENT_76 | NO_CURRENT_76 |
| AATCTTACTCGTCCTCCTTG | NO_CURRENT_77 | NO_CURRENT_77 |
| AATCTTATGATAAATATGGG | NO_CURRENT_78 | NO_CURRENT_78 |
| AATGACATTTAGTGCTTGGG | NO_CURRENT_79 | NO_CURRENT_79 |
| AATGAGACCTTCTCTCAACG | PQBP1 | 10084 |
| AATGAGTCCAGCTCAAGAAG | MMP13 | 4322 |
| AATGCAGAAATGCAGGCTTC | PPARG | 5468 |
| AATGGCGCCCTACAGCCTAC | ACO2 | 50 |
| AATGGGTTGCCGGGAGTAAA | NO_CURRENT_80 | NO_CURRENT_80 |
| AATTAAAGTGATAAGCATAG | NO_CURRENT_81 | NO_CURRENT_81 |
| AATTAGCCTGTCTCATAACT | NO_CURRENT_82 | NO_CURRENT_82 |
| AATTGCCTACCACCGCCACT | PHF8 | 23133 |
| AATTGTAAATCCCAGTGTTG | NO_CURRENT_83 | NO_CURRENT_83 |

| | | |
|---|---|---|
| AATTTCTGCAGACATAGGTG | NO_CURRENT_84 | NO_CURRENT_84 |
| ACAAACGACCTTGAGCAGGG | NO_CURRENT_85 | NO_CURRENT_85 |
| ACAAAGCAAGTTCTCAAAAG | NO_CURRENT_86 | NO_CURRENT_86 |
| ACAAAGGTTCCACAGCCTGG | GBP5 | 115362 |
| ACAAAGTTGTAATTGTGTCA | UTY | 7404 |
| ACAACAACTTTATTAGCACC | TFPT | 29844 |
| ACAACAGTCCCACATGAAGG | NO_CURRENT_87 | NO_CURRENT_87 |
| ACAACCCAGGAGGCACGACG | PDHA1 | 5160 |
| ACAAGGGCCAGGAGAAGCTG | EBI3 | 10148 |
| ACACAAGTAGTTTACATTGT | RXRA | 6256 |
| ACACAATTACAACTTTGTGC | UTY | 7404 |
| ACACAGCCACCTAAGAGGCC | GBP7 | 388646 |
| ACACCAAACACCCACGACGT | LGMN | 5641 |
| ACACCACCAAGCCGCAGGGA | NIPBL | 25836 |
| ACACCATATTAGTCTTGTGA | NO_CURRENT_88 | NO_CURRENT_88 |
| ACACCCATTCTCATAACGGA | NO_CURRENT_89 | NO_CURRENT_89 |
| ACACCGAAGCACCTGTACGT | NO_CURRENT_90 | NO_CURRENT_90 |
| ACACCTGAGCAGATGAGAAC | SLC5A8 | 160728 |
| ACACCTGGACCGCCGCGCCG | ALOX5 | 240 |
| ACACGAGGTGTTATTCCCAG | TLR1 | 7096 |
| ACACGGCCCGAAAAGTCGCT | PHF8 | 23133 |
| ACACGTGCCTCCGCGACGCG | TRMT61A | 115708 |
| ACACTCACCGCGTGCGGCGG | PTGIS | 5740 |
| ACACTCTGGAGGCGTACTTG | CYP27B1 | 1594 |
| ACAGAATAACCCGCCCCCAG | MGA | 23269 |
| ACAGACACCCTCCGCCTCAT | SHOC2 | 8036 |
| ACAGACCAATATATCAACTG | NO_CURRENT_91 | NO_CURRENT_91 |
| ACAGAGTCCAGCGGAGTTGT | DNTTIP1 | 116092 |
| ACAGAGTCTTGGACTGGGCT | SLC7A3 | 84889 |
| ACAGCCCTCACGAGCCCGAA | NO_CURRENT_92 | NO_CURRENT_92 |
| ACAGCGATAAGGCCTTTCCC | ZNF616 | 90317 |
| ACAGCGCTCTCGTGTACTAT | NO_CURRENT_93 | NO_CURRENT_93 |
| ACAGCGGAAAGGAAGTTCCA | INO80C | 125476 |
| ACAGCTCCGACTGTCAGTGC | MCTS1 | 28985 |
| ACAGGACCGGGAGAGCAGTG | NLRP3 | 114548 |
| ACAGGAGGATCCGGTGCAGC | BRK1 | 55845 |
| ACAGGCAGATTTGCCTGCTG | CAT | 847 |
| ACAGGTTCTTATTCATTGAC | NO_CURRENT_94 | NO_CURRENT_94 |
| ACAGTATATTTATGGGAATG | NO_CURRENT_95 | NO_CURRENT_95 |
| ACAGTCAGACAGCCAATAAA | HTR3E | 285242 |
| ACAGTCTGTCATCTTCTGCC | BCL2A1 | 597 |
| ACAGTGCGGCCAGGAGGCCG | PTGIS | 5740 |
| ACAGTGCTGCCTCGTCTGAG | HIF1A | 3091 |
| ACATACAGTATAGGAAACCT | NO_CURRENT_96 | NO_CURRENT_96 |
| ACATCACCCAGGAGTCTGCT | CYP4A11 | 1579 |
| ACATCTCTGCCCCGTTACCT | NDUFA1 | 4694 |
| ACATGAGTGAACTCTTCCAT | COX16 | 51241 |
| ACATGCAGCTTCTCCTCCAT | OR7C2 | 26658 |
| ACATGCTATATACTTTAGTG | NO_CURRENT_97 | NO_CURRENT_97 |
| ACATTAGTTAAACACCTATG | NO_CURRENT_98 | NO_CURRENT_98 |

| | | |
|---|---|---|
| ACATTCTGGGTGACACGCTG | PRKAA1 | 5562 |
| ACATTGCCTCAACAGCTTCA | BCL2A1 | 597 |
| ACCAACCAACAAAATGGCGA | DYRK1A | 1859 |
| ACCAAGAGGGTGGGAAAGAT | MKL1 | 57591 |
| ACCAAGGTAAGCGCCGTACG | ZRSR2 | 8233 |
| ACCAATTCTCCTGTCGGCGG | NIPBL | 25836 |
| ACCACAGACACTGCCGCCGC | MKL1 | 57591 |
| ACCAGCAGAGCTCCCAGCAC | CYP8B1 | 1582 |
| ACCAGCCTCCCCGCGCAGCC | SLC6A4 | 6532 |
| ACCAGGAGCACCAGGACCAG | MMP9 | 4318 |
| ACCAGGCTGCATGATGGCAT | CDKN2C | 1031 |
| ACCAGGGCCAGCGCTGCCAG | NENF | 29937 |
| ACCATGGTGCTATCTCCCAG | AGER | 177 |
| ACCATTCAAGATGCATCCAG | MMP13 | 4322 |
| ACCCAATCCGAGGGTCATGG | ERCC6L | 54821 |
| ACCCAATGTGGCGGAGCCGA | NO_CURRENT_99 | NO_CURRENT_99 |
| ACCCACCTGCCAGGCTGCGC | SLC6A4 | 6532 |
| ACCCACCTGCCCGAATCTGG | ARMCX2 | 9823 |
| ACCCACGGAACTACAGGTGT | FAM214B | 80256 |
| ACCCACTGAGGTCACCCTCC | GBP5 | 115362 |
| ACCCAGCTGCTAGTCAGAAA | CASP5 | 838 |
| ACCCATGAGTTAAGTTTTCT | NO_CURRENT_100 | NO_CURRENT_100 |
| ACCCATGATGTAACCATAGC | NO_CURRENT_101 | NO_CURRENT_101 |
| ACCCATTGAGAGTCGCCTGA | NO_CURRENT_102 | NO_CURRENT_102 |
| ACCCCACCTTGAGGAAGACG | HNRNPF | 3185 |
| ACCCCCGCCTCACTCCAATG | DYRK1A | 1859 |
| ACCCGACACCACCAAGCCGC | NIPBL | 25836 |
| ACCCGAGGGGGCGGCAACCG | NELFE | 7936 |
| ACCCGAGTGCTTCCCGCAGA | GNAI2 | 2771 |
| ACCCGCCATGCTCTGCGCAC | SLC7A5 | 8140 |
| ACCCGGCTGCGCGGGTGGTG | PTGES2 | 80142 |
| ACCCGGCTGGGTAAGTGAGA | TBK1 | 29110 |
| ACCCGGGGCTGGAGAGGCCG | RRAGA | 10670 |
| ACCCGTTAGAGTGGCGAGTA | NO_CURRENT_103 | NO_CURRENT_103 |
| ACCGACGAGTCTCAACTAAA | CFLAR | 8837 |
| ACCGAGACTTAAGGACCGTG | SPCS1 | 28972 |
| ACCGAGAGCGCTGCTCAGGC | HSPA4 | 3308 |
| ACCGCATCCGCGTGTCCACT | ALOX15 | 246 |
| ACCGCGCGGCTGGAGGTGTG | HSP90B1 | 7184 |
| ACCGCGGGGACCTAGGACGG | VIPR2 | 7434 |
| ACCGCGGTGGTGGCGCCGGC | LGMN | 5641 |
| ACCGGCCACGGCGGTCTCCG | NDUFA12 | 55967 |
| ACCGGGGGCCCCACGGTCTC | NELFE | 7936 |
| ACCGGGGGGACAAAGACACC | CCAR2 | 57805 |
| ACCGTGAAGGCCTACCTTCT | SQSTM1 | 8878 |
| ACCTATCATGGTCACTGGGT | PRKN | 5071 |
| ACCTATTGTCCCTTCAAGCT | NO_CURRENT_104 | NO_CURRENT_104 |
| ACCTCATGAGTAAGCTGTGG | INO80B | 83444 |
| ACCTCCCTGAACATCTCCTC | PAXIP1 | 22976 |
| ACCTCCTAGCAGAGGGGGAG | CYP4F11 | 57834 |

| | | |
|---|---|---|
| ACCTCGAGAGGCCTGAGGCA | SUPT7L | 9913 |
| ACCTCGGGATTGGCCTCCAG | UCHL3 | 7347 |
| ACCTCTCGAATAAGGCGCGG | METTL3 | 56339 |
| ACCTGCCAGGCTGCGCGGGG | SLC6A4 | 6532 |
| ACCTGCGACGCTCCTGCTGC | TRMT61A | 115708 |
| ACCTGCGTCCGCGGCGGCTT | CYFIP1 | 23191 |
| ACCTGCTTGTCTGTTCCATT | TLR1 | 7096 |
| ACCTGGCCGGTGCTTCAGGT | CXCR1 | 3577 |
| ACCTGGGTTCAGAACAAGTC | HTR3E | 285242 |
| ACCTGGTGCCCTCCCCGAGG | RHOG | 391 |
| ACCTGTTAGGGGAACACTGT | NO_CURRENT_105 | NO_CURRENT_105 |
| ACCTTACAATAAGTTATATT | NO_CURRENT_106 | NO_CURRENT_106 |
| ACCTTGTCAACCCGTCGGCG | CS | 1431 |
| ACGACGACCGTTACCCCAAC | MSL2 | 55167 |
| ACGACGACGACGACGGCGGC | NUP153 | 9972 |
| ACGACGCCCCTCTCCACACC | PAGR1 | 79447 |
| ACGATCGGTAATGGTCTGTT | NO_CURRENT_107 | NO_CURRENT_107 |
| ACGATCTTAGGTGGGTTCCC | HSPA13 | 6782 |
| ACGATTCACAGCGTCCACAC | STRAP | 11171 |
| ACGCAAGATGGCGGACAGTG | PIAS1 | 8554 |
| ACGCAGCGCGGCCCGAAGAC | AKT1 | 207 |
| ACGCCAGAAACAGCATCACG | TMEM70 | 54968 |
| ACGCCGAGGTCCTGGTTCCG | SOD1 | 6647 |
| ACGCGCTGGAGATGATGGGG | STAG2 | 10735 |
| ACGCGGCGCCCCGGCCCGTT | PARK7 | 11315 |
| ACGCGGGTGACGAGCGGCTG | TMEM70 | 54968 |
| ACGCTCAGCACCCGCTATGC | NO_CURRENT_108 | NO_CURRENT_108 |
| ACGCTCCCTCCTGCCTGCTG | UBE2H | 7328 |
| ACGCTGACGAGTAAAAGCGG | NO_CURRENT_109 | NO_CURRENT_109 |
| ACGGAAAGACCTCGCTATTC | NO_CURRENT_110 | NO_CURRENT_110 |
| ACGGAGCCGACAGAGGCCGG | PRR14L | 253143 |
| ACGGAGGCTAAGCGTCGCAA | NO_CURRENT_111 | NO_CURRENT_111 |
| ACGGCCGCTCGGGAGCCTGG | NDUFC1 | 4717 |
| ACGGCCTCATTATGATACCC | NO_CURRENT_112 | NO_CURRENT_112 |
| ACGGCGCCGCTGCTGCCAGG | PIAS1 | 8554 |
| ACGGCGGCAGGTAGGCACAG | IRF8 | 3394 |
| ACGGCGGCGGGGGAGTCTCG | MSL2 | 55167 |
| ACGGCTCGGCGGGTGCGGCC | SLC25A19 | 60386 |
| ACGGCTGGATGCGGAGAGTT | TRERF1 | 55809 |
| ACGGGAGGACAGCCGGAGAA | SLC25A6 | 293 |
| ACGGGCCGCAGCGCCTCTGC | CACNA2D2 | 9254 |
| ACGGGCGGAGCCTCAGGGCC | TRIR | 79002 |
| ACGGGCGGCCGGGCGGTGCG | MAP2K7 | 5609 |
| ACGGGGCGATGGCGGCTGAG | IFNAR1 | 3454 |
| ACGGGTCATGGCGACGGCAG | SERF2 | 10169 |
| ACGGTAACTCGGGGCCGATG | SHOC2 | 8036 |
| ACGGTACATGCGCATGAGTC | NO_CURRENT_113 | NO_CURRENT_113 |
| ACGGTGAAGCGCGACGATGC | UQCRB | 7381 |
| ACGGTGGGGATGGACCTACT | NO_CURRENT_114 | NO_CURRENT_114 |
| ACGGTGGGGGACTCTCACCT | MOSPD1 | 56180 |

| | | |
|---|---|---|
| ACGGTTATGGTCTCATGGGG | NO_CURRENT_115 | NO_CURRENT_115 |
| ACGTCAACTGCTGGAGTGGG | NO_CURRENT_116 | NO_CURRENT_116 |
| ACGTCCATACTGTCGGCTAC | NO_CURRENT_117 | NO_CURRENT_117 |
| ACGTCCTCTCGCGGCCCTCA | NNT | 23530 |
| ACGTCGTTTAGCACCCGGCT | NO_CURRENT_118 | NO_CURRENT_118 |
| ACGTGCCTGTCGCCCCTGGT | GPT2 | 84706 |
| ACGTGGGGACATATACGTGT | NO_CURRENT_119 | NO_CURRENT_119 |
| ACGTGTGGAGCCCCGGCAGC | TRMT61A | 115708 |
| ACGTGTGTCCTTTAAACACG | NO_CURRENT_120 | NO_CURRENT_120 |
| ACGTTCGAGTACGACCAGCT | NO_CURRENT_121 | NO_CURRENT_121 |
| ACGTTGAGGTCCTAACTCTG | NO_CURRENT_122 | NO_CURRENT_122 |
| ACTAACGAGAGAACCGACGG | RHOA | 387 |
| ACTAAGCTTATCTACAACCG | NO_CURRENT_123 | NO_CURRENT_123 |
| ACTAGAGTCATGATCAGCGA | NO_CURRENT_124 | NO_CURRENT_124 |
| ACTAGTGATAAGTTCACCAT | NO_CURRENT_125 | NO_CURRENT_125 |
| ACTATCTCGGGCATGGCTCT | LTA4H | 4048 |
| ACTATTTAATATTGGTAAGT | NO_CURRENT_126 | NO_CURRENT_126 |
| ACTCACAGGTTTGCAGGCAA | CD84 | 8832 |
| ACTCACCGCGTGAGCCCACC | CHEK2 | 11200 |
| ACTCAGCACACGTTAGAGAG | NO_CURRENT_127 | NO_CURRENT_127 |
| ACTCAGCATGGTCTGGCAGC | PIGY | 84992 |
| ACTCAGCATGGTCTGGCAGC | PYURF | 100996939 |
| ACTCATGGTGCAGCACCTGC | CYP4A11 | 1579 |
| ACTCATGGTGCAGCACCTGC | CYP4A22 | 284541 |
| ACTCCAGGGGACCCACAGCT | RIPK1 | 8737 |
| ACTCCATTCCCGTGGAAACC | HTR2A | 3356 |
| ACTCCCGGGCCGGTCATGAA | INO80E | 283899 |
| ACTCCCGGTTAGCCCAGTCC | BRK1 | 55845 |
| ACTCCTCCCATCAAGAACTG | NO_CURRENT_128 | NO_CURRENT_128 |
| ACTCCTGGTTTCCACGGGAA | HTR2A | 3356 |
| ACTCGAGGTCCCTTCCCACA | TLR9 | 54106 |
| ACTCGCGTCCGCCTCTCGCC | MPC2 | 25874 |
| ACTCGGTTAGCTCGCGCGCC | GPT2 | 84706 |
| ACTCTCCCCCTTCGTCCGGG | RBM42 | 79171 |
| ACTCTGGAGTTTGGGCGGCC | PPID | 5481 |
| ACTGACAGCTATAGGGCAGG | PHIP | 55023 |
| ACTGACTGCATTGGCACGGT | CHMP5 | 51510 |
| ACTGAGTGGGTAACACGCAT | NO_CURRENT_129 | NO_CURRENT_129 |
| ACTGATGTCGTAACTAGGGG | PHC3 | 80012 |
| ACTGCAACCCTAATCAGGTA | STAT2 | 6773 |
| ACTGCATGGTGTCATATGCA | NO_CURRENT_130 | NO_CURRENT_130 |
| ACTGCCTTCTGAGTCCCACC | EZH2 | 2146 |
| ACTGCGACGCAGCCCGGAGT | MAPK14 | 1432 |
| ACTGCGCCGCGTCTTCCTCA | HNRNPF | 3185 |
| ACTGCGCTCGGCTCGCGCCC | PIP4K2A | 5305 |
| ACTGCGGAGCGCCCAATATC | NO_CURRENT_131 | NO_CURRENT_131 |
| ACTGCTCCCGGTCGCCCCTC | NO_CURRENT_132 | NO_CURRENT_132 |
| ACTGCTCGTTCGCTGCGCCG | FOS | 2353 |
| ACTGCTCTATCCCGGGCAAA | APAF1 | 317 |
| ACTGCTGCTGACATCTCTTA | NO_CURRENT_133 | NO_CURRENT_133 |

| | | |
|---|---|---|
| ACTGCTGCTGCTGCTGTTCT | MMP1 | 4312 |
| ACTGCTGCTGTCTTCTAAAT | NO_CURRENT_134 | NO_CURRENT_134 |
| ACTGGAGAGGCAATAATTTG | NO_CURRENT_135 | NO_CURRENT_135 |
| ACTGGAGCGCCCCGCTGGAG | SPHK1 | 8877 |
| ACTGGCCGAAGCGACGAAGA | HIF1A | 3091 |
| ACTGGGGCCTCGCTCTATGC | ALOX15 | 246 |
| ACTGGTAACTTTGGGGGCAT | RNF25 | 64320 |
| ACTGTGGTCCATATTTCTTG | NO_CURRENT_136 | NO_CURRENT_136 |
| ACTTACACTGATCCCCTCCA | CXCR4 | 7852 |
| ACTTACGGCACTCGCATGCC | NO_CURRENT_137 | NO_CURRENT_137 |
| ACTTCAGTTCGGCGTAGTCA | NO_CURRENT_138 | NO_CURRENT_138 |
| ACTTCCAAGGCAGCCCGGGT | NMRK2 | 27231 |
| ACTTGAACTTCGATTCAGAA | RND1 | 27289 |
| ACTTGCAGACGTGTTTAGTA | PPID | 5481 |
| ACTTGGTGGGTCCCGGGCCA | CHMP6 | 79643 |
| ACTTGTCAGCTCCTTGTCAT | MBNL1 | 4154 |
| ACTTGTCCCTGTGCAGCCCG | BAK1 | 578 |
| ACTTTAAGATAACACTTAGA | NO_CURRENT_139 | NO_CURRENT_139 |
| AGAAAAGGGCAAGGAGAAGG | SERF2 | 10169 |
| AGAAAATTAAAGCAGAGAGG | TNFRSF1A | 7132 |
| AGAACAAAGGTTCCACAGCC | GBP5 | 115362 |
| AGAACCCAGACGCCAGCGGT | NO_CURRENT_140 | NO_CURRENT_140 |
| AGAACGAACGGCTTGGGCGC | DPY30 | 84661 |
| AGAACGAACGGCTTGGGCGC | MEMO1 | 51072 |
| AGAACGGAGGAGCTTTGTAG | NO_CURRENT_141 | NO_CURRENT_141 |
| AGAAGACCGAGGCGCTTCAA | NO_CURRENT_142 | NO_CURRENT_142 |
| AGAAGCAGAAACACGACGGG | PRKAA1 | 5562 |
| AGAAGCAGGTAGTTCTTAAA | ACOD1 | 730249 |
| AGAAGCCAAGACTGAGCCGG | FOS | 2353 |
| AGAAGCCCACAGGCCCAAGG | CYP4B1 | 1580 |
| AGAAGGCAGTGGAGCCCCGG | EZH2 | 2146 |
| AGAAGGCCATGCAAAGACGG | NO_CURRENT_143 | NO_CURRENT_143 |
| AGAAGGGAGCGGGACCGGAA | MAPK9 | 5601 |
| AGAAGGGGGCTCGGTGCTCA | SERF2 | 10169 |
| AGAAGTGCACCAGCGAGCCG | NFKB1 | 4790 |
| AGAATAACCAGAGAACAACC | SERPINB2 | 5055 |
| AGAATCAGTTTAATTAGAAG | NO_CURRENT_144 | NO_CURRENT_144 |
| AGACAGACAGCCAGAGAGAG | AHDC1 | 27245 |
| AGACAGCCAGAGAGAGCGGG | AHDC1 | 27245 |
| AGACCAAAGCTGCTTAATCG | NO_CURRENT_145 | NO_CURRENT_145 |
| AGACCATCTAGGCAAGAGTT | NO_CURRENT_146 | NO_CURRENT_146 |
| AGACCCAAAATGACAAAGCT | NO_CURRENT_147 | NO_CURRENT_147 |
| AGACCCCGTAGGCAGGACGT | NO_CURRENT_148 | NO_CURRENT_148 |
| AGACCCGGGCTTGGGCCCCG | NNT | 23530 |
| AGACCGCCAACGCCACACAC | PIGL | 9487 |
| AGACGCAGACACACATTGCA | NO_CURRENT_149 | NO_CURRENT_149 |
| AGACGCTTCACAGAAAACTG | NO_CURRENT_150 | NO_CURRENT_150 |
| AGACTATGGAACGTCAGGTG | APEH | 327 |
| AGAGAACCGACGGAGGACCG | RHOA | 387 |
| AGAGAAGGCGCTGGGCTGCG | TNFRSF1B | 7133 |

| | | |
|---|---|---|
| AGAGAATTAGGGCTTTGCTA | NO_CURRENT_151 | NO_CURRENT_151 |
| AGAGACCATCCCTCTCCCCC | SPEN | 23013 |
| AGAGAGAGACACGAGTGGCC | UBE2E1 | 7324 |
| AGAGAGATGACGATCTTAGG | HSPA13 | 6782 |
| AGAGAGCGAAGGAGTGATCG | AHDC1 | 27245 |
| AGAGAGCTACTCCATTCCCG | HTR2A | 3356 |
| AGAGCATGCCCAAATAACCT | AIM2 | 9447 |
| AGAGCCGCTGGTGGGAGGCG | PRKN | 5071 |
| AGAGCTCCATAATTTGCCTA | NO_CURRENT_152 | NO_CURRENT_152 |
| AGAGCTGGCCCGCGCCTCGG | KPNA1 | 3836 |
| AGAGGAAACCAAACAGAAGT | OR7C2 | 26658 |
| AGAGGAGACTATGGAACGTC | APEH | 327 |
| AGAGGATGTGGCGGCGGGCT | VDAC1 | 7416 |
| AGAGGCAGCAAGGTCCGCCG | ABI1 | 10006 |
| AGAGGCGACCGCGGCTGCCT | SATB2 | 23314 |
| AGAGGCTGCTGCTCCGAGCC | ARF1 | 375 |
| AGAGGGATTGGGAGCTTGAC | NO_CURRENT_153 | NO_CURRENT_153 |
| AGAGGGCTTAACCTTGCACA | NO_CURRENT_154 | NO_CURRENT_154 |
| AGAGGGGGCGCTGAGCTGTT | CSDE1 | 7812 |
| AGAGGGTCTGGCGGGCAGAC | TLR9 | 54106 |
| AGAGTAACAATGGATGTGTA | NO_CURRENT_155 | NO_CURRENT_155 |
| AGAGTAACCAGAAGTATGGG | NO_CURRENT_156 | NO_CURRENT_156 |
| AGAGTACCTAGTGGCCGCCC | PPID | 5481 |
| AGAGTAGACAGCAGAACCAG | RBM42 | 79171 |
| AGAGTCAAAGTCAACCCAAA | NO_CURRENT_157 | NO_CURRENT_157 |
| AGAGTCCAGCGGAGTTGTGG | DNTTIP1 | 116092 |
| AGAGTCCGCCGCCAACGCGC | PLD1 | 5337 |
| AGAGTGAGACAGTCTAATAT | NO_CURRENT_158 | NO_CURRENT_158 |
| AGAGTGAGCCAACTCACCTA | PPARG | 5468 |
| AGATCTGGGTGCAAAAGCCC | LAMTOR2 | 28956 |
| AGATGAGAGTTCAGCCGCGG | POU2F1 | 5451 |
| AGATGGCGGCGGCGCTGAGG | INO80 | 54617 |
| AGATGGCTCAGCACCACCTA | CD84 | 8832 |
| AGATGGCTCTTGCTCCCAAA | EDA2R | 60401 |
| AGATTAAAGACGCAACCGGC | NO_CURRENT_159 | NO_CURRENT_159 |
| AGATTACTGAAGTGGACCAT | NO_CURRENT_160 | NO_CURRENT_160 |
| AGATTCATTCACGAGTTGGG | NO_CURRENT_161 | NO_CURRENT_161 |
| AGATTTGGGATTCTTTAAAC | TNRC18 | 84629 |
| AGCAAAACAAAAGGGAAACC | FOXO4 | 4303 |
| AGCAAAGCCGAGCTCTGGGC | SEC62 | 7095 |
| AGCAAGCGTGGGAACGCGGG | IRF8 | 3394 |
| AGCAATGTGGCTCCTGTGTG | PIGL | 9487 |
| AGCACAGGCAGTGGCGGCAG | PDHA1 | 5160 |
| AGCACAGGCCTGCCAGCAGG | SERPINA2 | 390502 |
| AGCACTAGAGATATATAGAT | NO_CURRENT_162 | NO_CURRENT_162 |
| AGCACTGTTCCTGTCGATAG | NO_CURRENT_163 | NO_CURRENT_163 |
| AGCAGAGCGGAGCGGATCCG | WDR26 | 80232 |
| AGCAGAGCTGGGTAGGAGCA | FOS | 2353 |
| AGCAGAGGGCCTTACCCGAG | NELFE | 7936 |
| AGCAGATGTCAAAGACATGA | NCKAP1L | 3071 |

| | | |
|---|---|---|
| AGCAGATTCAAGCATGCAGT | NO_CURRENT_164 | NO_CURRENT_164 |
| AGCAGCAGCAGGAAGAAGTC | JAM3 | 83700 |
| AGCAGCAGCCTCTGAGGTGA | ARF1 | 375 |
| AGCAGCCAGGAGCAAACTCT | CCL20 | 6364 |
| AGCAGCCCGCAGCCTCAGCC | NUFIP2 | 57532 |
| AGCAGCCTCCGCCCCCCGCA | EGFR | 1956 |
| AGCAGCGGCAGTGGCGGCGA | RNF111 | 54778 |
| AGCAGCGTCCCTTATTCGCT | UBA7 | 7318 |
| AGCAGGACACTCACCGCGTG | PTGIS | 5740 |
| AGCAGGACCTGGCCAGGAAG | ASH2L | 9070 |
| AGCAGGCCGGTAAGTAACTG | UQCRB | 7381 |
| AGCAGGGAGCTGGGAGCTGG | JUNB | 3726 |
| AGCAGTGGAGGCAGAGGCGC | SUV39H1 | 6839 |
| AGCAGTGGCGTCCGCAGCTG | TRAF6 | 7189 |
| AGCATACCAAGAATAGCTTA | NO_CURRENT_165 | NO_CURRENT_165 |
| AGCATAGCCATGGTTCTCTG | CYP8B1 | 1582 |
| AGCATCACGCGGGTGACGAG | TMEM70 | 54968 |
| AGCATTCTCACCAAGACCGA | NO_CURRENT_166 | NO_CURRENT_166 |
| AGCCAAATGACTTTGCATTG | NO_CURRENT_167 | NO_CURRENT_167 |
| AGCCAACTCACCTAAGGAAA | PPARG | 5468 |
| AGCCAAGTTACTCACGGGAG | ANP32B | 10541 |
| AGCCAATAAAAGGCAGAAGC | HTR3E | 285242 |
| AGCCAGAGGCAGGCGCACCC | PDE3A | 5139 |
| AGCCAGCGATACCTGTAACG | OXSM | 54995 |
| AGCCAGGCGGCGGCGGCGAC | C14orf28 | 122525 |
| AGCCAGGCGGCGGCGGCGAC | MCL1 | 4170 |
| AGCCCAGCACCAGGAGCACC | MMP9 | 4318 |
| AGCCCAGGGTTAGGAACCGT | LAMTOR2 | 28956 |
| AGCCCCCTTGCTGGGAGAAG | NR1H3 | 10062 |
| AGCCCCGGGGGCCGCTCCAT | CHUK | 1147 |
| AGCCCCGTTACAGGTATCGC | OXSM | 54995 |
| AGCCCGGAGTCGGCCTTGTA | MAPK14 | 1432 |
| AGCCCGGGTTCAGGCTCTCA | FLCN | 201163 |
| AGCCCGGGTTCAGGCTCTCA | PLD6 | 201164 |
| AGCCCTCTGCGGGAAGCACT | GNAI2 | 2771 |
| AGCCGAATCATGGATCACAC | CBLL1 | 79872 |
| AGCCGACATTGCCGGCGTCT | ATP5E | 514 |
| AGCCGCAGCAGCAGCCGCCG | UBE2E1 | 7324 |
| AGCCGCAGCGGCCAGGCAGG | ABCB10 | 23456 |
| AGCCGCCCTCGCGCCGCCTC | FAM107B | 83641 |
| AGCCGCCGCCGCTGCACCAG | CS | 1431 |
| AGCCGCCTGTCCGGGTGTGG | NFKB2 | 4791 |
| AGCCGCGCAGTCCTACTACC | MAP3K7 | 6885 |
| AGCCGCGGCTGGAAGATGGC | PPARGC1B | 133522 |
| AGCCGCGGGAAGGGTACTCC | MPC2 | 25874 |
| AGCCGCGGGTCTCCGGCCGG | EIF2AK2 | 5610 |
| AGCCGCGGGTGCAGGAGTCC | DET1 | 55070 |
| AGCCGGAGAGGCCAACGAAC | LTA4H | 4048 |
| AGCCGGTCAGAGGGTGAGTG | UBE2H | 7328 |
| AGCCTCTGAGGTGAGGGCGA | ARF1 | 375 |

| | | |
|---|---|---|
| AGCCTGGAGGGTGACCTCAG | GBP5 | 115362 |
| AGCCTTGGGCTGTGCCCTGA | NNT | 23530 |
| AGCCTTTGTAGATCTCAGAA | RAD21 | 5885 |
| AGCGAACGAGCGGCGCTCGG | FOSL2 | 2355 |
| AGCGAGAAAGCGCAGCCAGG | TLR2 | 7097 |
| AGCGAGGGGTCGAGCGCGGC | ARHGAP33 | 115703 |
| AGCGATCTGGACACTCTCCA | NO_CURRENT_168 | NO_CURRENT_168 |
| AGCGATTCACGTATTAGATG | NO_CURRENT_169 | NO_CURRENT_169 |
| AGCGCAGCCAGGCGGCTGCT | TLR2 | 7097 |
| AGCGCAGGCCGCGGCGGCCG | JAK1 | 3716 |
| AGCGCCGAGCTGTGCGCAGC | DNAJA1 | 3301 |
| AGCGCCTAAGTCCTTCCAGT | HSPA4 | 3308 |
| AGCGCGCCGGGCGCAGGGCC | KIAA1211L | 343990 |
| AGCGCGGTTACCGGACGGGC | CCNC | 892 |
| AGCGCGTCGCCGGGCCGCTC | NFKB1 | 4790 |
| AGCGCTCTGGTTGCATCCCT | NO_CURRENT_170 | NO_CURRENT_170 |
| AGCGCTGACGGCCGCCGGCA | BCL2 | 596 |
| AGCGCTGCTCAGGCCGGCTC | HSPA4 | 3308 |
| AGCGGAGTTGTGGGGGCCGG | DNTTIP1 | 116092 |
| AGCGGATCCGAGGACAGAGG | WDR26 | 80232 |
| AGCGGATGGGTGCTATTGTG | CASP3 | 836 |
| AGCGGCCGCAGCCGCCGCCG | NENF | 29937 |
| AGCGGCGGCGGCGGCTGTAG | RNF111 | 54778 |
| AGCGGCGGTACCGGTGCTGG | MTOR | 2475 |
| AGCGGCTCCAGCAGCAGCGG | ADRB1 | 153 |
| AGCGGGAGCAGAGGAGGCGA | SIRT1 | 23411 |
| AGCGGGAGGTCGCGAAGCCT | ACADS | 35 |
| AGCGGGATGTGAAGGACTCC | MGA | 23269 |
| AGCGGGGTGAGCGGCGGCAG | PYCARD | 29108 |
| AGCGTGACTACAGGGTATGG | DLAT | 1737 |
| AGCGTTGAGATTGAGACTGG | CYP1B1 | 1545 |
| AGCTAGCGATGGCTCTAAGT | NO_CURRENT_171 | NO_CURRENT_171 |
| AGCTAGTGCTAACCTACCTA | NO_CURRENT_172 | NO_CURRENT_172 |
| AGCTCACCGACTCCGGACGC | NUDT17 | 200035 |
| AGCTCCAGGAGGGTGAGCGC | HELZ2 | 85441 |
| AGCTCCCAGCTCCCAGCTGC | KIAA1211L | 343990 |
| AGCTCGCCATGTCGGTTCTC | NO_CURRENT_173 | NO_CURRENT_173 |
| AGCTCTTCAGGCGGCGAGTC | IRGM | 345611 |
| AGCTGCAGGAATTCAGCTGC | SERPINE1 | 5054 |
| AGCTGCATTTGCCTTTACTG | FBXW7 | 55294 |
| AGCTGCGCGCTACTGGATCA | NO_CURRENT_174 | NO_CURRENT_174 |
| AGCTGCGGGCGGTGGGAAAG | BNIP3 | 664 |
| AGCTGCTGGAGGGGGGCGTG | SERPINE1 | 5054 |
| AGCTGCTTCTGATAGGAGCC | CYP4F11 | 57834 |
| AGCTGGACTCTGTAGAAATC | NO_CURRENT_175 | NO_CURRENT_175 |
| AGCTGGAGAGCGAACGAGCA | IDH2 | 3418 |
| AGCTGGGGACCAGGCAGCAC | SERPINA2 | 390502 |
| AGCTGTAACTCTCACTTCGA | IRGM | 345611 |
| AGCTGTCCCTGAAGTGACAG | LAMTOR3 | 8649 |
| AGCTGTGGGAGGAGGCGGCG | MLF2 | 8079 |

| | | |
|---|---|---|
| AGCTTAATGTGCAGGTCAGA | NO_CURRENT_176 | NO_CURRENT_176 |
| AGCTTCATGCCGGTGCGGAG | NMRK2 | 27231 |
| AGCTTCGCACGGAGTGTGTG | NUDT17 | 200035 |
| AGGAAAATCGCCAACTCCTG | NO_CURRENT_177 | NO_CURRENT_177 |
| AGGAAACGGGGACCTGCCCG | LTB4R | 1241 |
| AGGAAGAGCTGTCTGCACCA | HTR2A | 3356 |
| AGGAAGCGTTGGCTCTTCTC | NO_CURRENT_178 | NO_CURRENT_178 |
| AGGAAGGCAGCCAGGACCCC | MMP13 | 4322 |
| AGGAATTCAGCTGCTGGAGG | SERPINE1 | 5054 |
| AGGACACTCACCGCGTGCGG | PTGIS | 5740 |
| AGGACAGCGGGGAAGGCGGG | MTOR | 2475 |
| AGGACCCAGAAGTAGGGTTT | NDUFA1 | 4694 |
| AGGACCTCATGAGTAAGCTG | INO80B | 83444 |
| AGGACTTTAATTGTGAAGGG | NO_CURRENT_179 | NO_CURRENT_179 |
| AGGAGAGAAAGGAAAGCGCG | DNAJB6 | 10049 |
| AGGAGATCGACCTCGTTCGG | NO_CURRENT_180 | NO_CURRENT_180 |
| AGGAGATTCTTCAAATGGTG | KMT2D | 8085 |
| AGGAGGACTGGTAACTTTGG | RNF25 | 64320 |
| AGGAGGAGAAGCTCAGGGAG | CYP4B1 | 1580 |
| AGGAGGAGGGAGGGTGAGTT | CDKN2D | 1032 |
| AGGAGGGCCAGAGAGGCAGT | SIRT1 | 23411 |
| AGGAGGTTTCGCCACCGGAG | NFKB1 | 4790 |
| AGGAGTCCTGGGGCATGGCG | DET1 | 55070 |
| AGGAGTTTATCAATAAGACC | NO_CURRENT_181 | NO_CURRENT_181 |
| AGGATAATAACCCAATGGCA | NO_CURRENT_182 | NO_CURRENT_182 |
| AGGATATCATCATGGTAGCG | NO_CURRENT_183 | NO_CURRENT_183 |
| AGGATGCAGGGCCGCGTGGC | C6orf15 | 29113 |
| AGGATGCTGAACAAGTACGT | NO_CURRENT_184 | NO_CURRENT_184 |
| AGGATGGATTGAGCAGCGGT | NO_CURRENT_185 | NO_CURRENT_185 |
| AGGATTATTATAATTAACGG | POU2F1 | 5451 |
| AGGCAACTAATCACTGCACA | NO_CURRENT_186 | NO_CURRENT_186 |
| AGGCAGAGGCGCGGGCCCGC | SUV39H1 | 6839 |
| AGGCAGGGCGGGGCGGGGCA | SRI | 6717 |
| AGGCAGGGCGGGGCGGGGCA | ZNF699 | 374879 |
| AGGCAGTGGAGCCCCGGCGG | EZH2 | 2146 |
| AGGCCAAGAAAATTCCCCAC | PTGS1 | 5742 |
| AGGCCACATGCAAAACACTT | NO_CURRENT_187 | NO_CURRENT_187 |
| AGGCCAGCCTCGGAGCCAGC | JUNB | 3726 |
| AGGCCAGGCCAGGCCGAGCC | RRAGC | 64121 |
| AGGCCCGAGAGTCAGAACCT | SPEN | 23013 |
| AGGCCGCGCGCGCGGGAGCG | TIFA | 92610 |
| AGGCCGTAGGAGGAAGATGG | SAP18 | 10284 |
| AGGCCTGGTGAGCACCGCCG | KPNA1 | 3836 |
| AGGCGCCGTGTGGCACTCGG | NDUFA8 | 4702 |
| AGGCGCGGGCCAGCTCTTCG | KPNA1 | 3836 |
| AGGCGGACCGGGGGCAAAGG | TNFRSF6B | 8771 |
| AGGCGGAGTTAAGCCGAGTT | SLC7A3 | 84889 |
| AGGCGGCCACAGCGCCATGT | EIF4G3 | 8672 |
| AGGCGGCGGAGAGCGAGGCC | KPNA1 | 3836 |
| AGGCGGCTCCTGCGATCGAA | HSP90B1 | 7184 |

| | | |
|---|---|---|
| AGGCGGTTGTAGAAGGTATG | CASP3 | 836 |
| AGGCTACGGTGAGCCGAAGG | BABAM1 | 29086 |
| AGGCTCGCGCGCCCGCGAAG | TP73 | 7161 |
| AGGCTCTCAAGCCCGGGTTC | FLCN | 201163 |
| AGGCTCTCAAGCCCGGGTTC | PLD6 | 201164 |
| AGGCTCTTGGGTCACATACC | HSH2D | 84941 |
| AGGCTGAGGACTGAAAAGAG | GPR119 | 139760 |
| AGGCTGGGCGGCGGAGCCTT | CHMP5 | 51510 |
| AGGCTGTTTGGACTCCGTGG | ATP5J2 | 9551 |
| AGGCTGTTTGGACTCCGTGG | ATP5J2-PTCD1 | 100526740 |
| AGGCTGTTTGGACTCCGTGG | PTCD1 | 26024 |
| AGGCTTAAAACTTGTTGGAG | RND3 | 390 |
| AGGCTTCGCCGCCTAGGTAA | RBM6 | 10180 |
| AGGCTTCTGGTCACCAGCAG | PPARG | 5468 |
| AGGGAAACCTCTATGGGTAA | NO_CURRENT_188 | NO_CURRENT_188 |
| AGGGAAGTGGAGTATTTGCT | CTNNBL1 | 56259 |
| AGGGACCTCTGGGTTCACGG | ACO2 | 50 |
| AGGGACGACGACGACGACGG | NUP153 | 9972 |
| AGGGAGCCACACAGGCTCCT | C4orf17 | 84103 |
| AGGGAGGAAAGTAGGGGATG | STRAP | 11171 |
| AGGGATAGTTAGGAAACTGA | NO_CURRENT_189 | NO_CURRENT_189 |
| AGGGATATAGGGCTTAGTAT | ARR3 | 407 |
| AGGGATCGTTAGGAAGGGAA | NO_CURRENT_190 | NO_CURRENT_190 |
| AGGGCAAGGAGAAGGGGGCT | SERF2 | 10169 |
| AGGGCAGGACAGGAAGCGGG | ZNF699 | 374879 |
| AGGGCCAGGCGCGTTTCTGC | RAC2 | 5880 |
| AGGGCCGGGCACCGCGGCCA | UCHL3 | 7347 |
| AGGGCGAGCGAGGAGGATGG | SHOC2 | 8036 |
| AGGGCGGGCAGCTGGCCCAG | EBI3 | 10148 |
| AGGGCTTTGGAATATTACCT | ARR3 | 407 |
| AGGGGAAGGACCCAGAAGCG | AHCTF1 | 25909 |
| AGGGGACGCAGCAAGGCGGA | NDUFC1 | 4717 |
| AGGGGCCTCCATAGCCCCAG | INO80B | 83444 |
| AGGGGCCTCCGGGGCTGCAC | CACNA2D2 | 9254 |
| AGGGGCGATGCTTCAGGTGG | KMT2D | 8085 |
| AGGGGCTCGGCTGCACCGGG | STAT1 | 6772 |
| AGGGGGAAAGAGGCTCGGAG | POU2F1 | 5451 |
| AGGGGGAGAGGAGGGTATGT | CYP4F11 | 57834 |
| AGGGGTGGCCGTGATGGCGG | ASH2L | 9070 |
| AGGGTACTCCAGGCGAGAGG | MPC2 | 25874 |
| AGGGTCAGTCTGTCCTTTTT | NO_CURRENT_191 | NO_CURRENT_191 |
| AGGGTCGCGGCCCGGAACGT | MLF2 | 8079 |
| AGGGTCTGGCTGGAGCCACG | CYP24A1 | 1591 |
| AGGGTGAGCACGAATATAGA | NO_CURRENT_192 | NO_CURRENT_192 |
| AGGGTGGAAGGCTGGATCGA | NO_CURRENT_193 | NO_CURRENT_193 |
| AGGTAAGATCAGCCAAGGAT | IRF9 | 10379 |
| AGGTAAGCCCCTTAGAACTG | NO_CURRENT_194 | NO_CURRENT_194 |
| AGGTACTCTGGGAGTGTGAG | SLC11A1 | 6556 |
| AGGTAGACCTGGCGACGACG | PRR14L | 253143 |
| AGGTAGCAAAGCCGAGCTCT | SEC62 | 7095 |

| | | |
|---|---|---|
| AGGTAGCAAAGTGACGCCGA | CXCR4 | 7852 |
| AGGTAGTTCTTAAAAGGAGA | ACOD1 | 730249 |
| AGGTAGTTGGAAGGTCCAGT | NO_CURRENT_195 | NO_CURRENT_195 |
| AGGTCCCGGCGCGGGGTCTG | MYD88 | 4615 |
| AGGTCCTCAGGAAGAAGCCG | TYK2 | 7297 |
| AGGTCGAGGGTCCCGGCGGC | JAM2 | 58494 |
| AGGTGAAGCAGTGGAGGCAG | SUV39H1 | 6839 |
| AGGTGAAGGTCTACACATCA | IL10 | 3586 |
| AGGTGAGGCGGCGGCTGAGT | FBXO38 | 81545 |
| AGGTGATATCTCTAGTGACA | NO_CURRENT_196 | NO_CURRENT_196 |
| AGGTGCCGAAAGCTCGGAAT | HDAC2 | 3066 |
| AGGTGGATCGGGGCGGTGTG | UCHL5 | 51377 |
| AGGTGTGAGGATCCGAACCC | HSP90B1 | 7184 |
| AGGTTACTCACACGGCTCGG | SLC25A19 | 60386 |
| AGGTTAGACCCTGAAAGAGA | CD84 | 8832 |
| AGGTTCTCTATCGACGAGTC | LTA4H | 4048 |
| AGGTTGTTTCCTTTCATTCG | PHF6 | 84295 |
| AGGTTTCTTCAGACCTCTCC | NLRP1 | 22861 |
| AGTAAAAGACTCACCGGCCA | CARD16 | 114769 |
| AGTAAAAGACTCACCGGCCA | CARD17 | 440068 |
| AGTAAAAGACTCACCGGCCA | CASP1 | 834 |
| AGTAAGAGCCAGCCCGTCCG | RRAGA | 10670 |
| AGTAATTCTTCATGACCTGT | GPR119 | 139760 |
| AGTAGACGGACGGTGAGCTG | NO_CURRENT_197 | NO_CURRENT_197 |
| AGTATTAGGTACCTGCCCTA | NO_CURRENT_198 | NO_CURRENT_198 |
| AGTATTGCACAGATTACTCA | NO_CURRENT_199 | NO_CURRENT_199 |
| AGTATTGTGACCACATAATG | NO_CURRENT_200 | NO_CURRENT_200 |
| AGTATTGTGGTGTCGTCAAC | NO_CURRENT_201 | NO_CURRENT_201 |
| AGTCACGCTGTTGCGACGAG | DLAT | 1737 |
| AGTCAGACATATAACCAACG | NO_CURRENT_202 | NO_CURRENT_202 |
| AGTCAGAGGAAGTGGGTGAC | MOSPD1 | 56180 |
| AGTCAGAGTCAGACATTTGG | BCL2L11 | 10018 |
| AGTCAGCCTTCCGTATCCAA | NO_CURRENT_203 | NO_CURRENT_203 |
| AGTCATAACTGAGTGAATCG | NO_CURRENT_204 | NO_CURRENT_204 |
| AGTCCAAACAGCCTTACCTG | ATP5J2 | 9551 |
| AGTCCAAACAGCCTTACCTG | ATP5J2-PTCD1 | 100526740 |
| AGTCCAAACAGCCTTACCTG | PTCD1 | 26024 |
| AGTCCAGAGGATCTCTACTC | GBP7 | 388646 |
| AGTCCAGCTCAAGAAGAGGA | MMP13 | 4322 |
| AGTCCAGTTACTTTCAGGCT | PQBP1 | 10084 |
| AGTCCGTCGCCCCGAACCGC | ATP5L | 10632 |
| AGTCCTCGGAACCAGGACCT | SOD1 | 6647 |
| AGTCCTCTGCGAATCCCAGG | OR7C2 | 26658 |
| AGTCCTTCACATCCCGCTGG | MGA | 23269 |
| AGTCGCACAAAAGAACTGCA | HTR2A | 3356 |
| AGTCGCCGCGCTCCGGCAGC | RIPK1 | 8737 |
| AGTCGCTATCGGAGGCCGCG | SNRNP70 | 6625 |
| AGTCGCTGGGAGAAGCCCAG | PHF8 | 23133 |
| AGTCGGCCTTGTAGGGGCGA | MAPK14 | 1432 |
| AGTCGTGGACTCGTGCAGCT | TMEM70 | 54968 |

| | | |
|---|---|---|
| AGTCTGCCCGCCAGACCCTC | TLR9 | 54106 |
| AGTCTGCTATATGTAACGTA | NO_CURRENT_205 | NO_CURRENT_205 |
| AGTCTTGGCCAATGTCACGG | NO_CURRENT_206 | NO_CURRENT_206 |
| AGTGACCTCTCCAAGAGCAG | DLAT | 1737 |
| AGTGAGAACACGGAGAGTGA | RTP1 | 132112 |
| AGTGAGAGTTACAGCTCTTC | IRGM | 345611 |
| AGTGAGCGGCGGCAGCTGCG | SBNO2 | 22904 |
| AGTGAGGCCGTGATACGCCA | ATP6V0E1 | 8992 |
| AGTGAGGGTAGGTGCAGGGC | RTP1 | 132112 |
| AGTGAGTGACAACCAGATCG | NO_CURRENT_207 | NO_CURRENT_207 |
| AGTGATCATAGAGGTGCCAG | BAK1 | 578 |
| AGTGCAAATATCAATCACCT | NO_CURRENT_208 | NO_CURRENT_208 |
| AGTGCAGACGCGGCTCCTAG | CASP3 | 836 |
| AGTGCCCAGAGAGGGGGTGC | SLC11A1 | 6556 |
| AGTGCGACCCCGGTGCCCGG | PTGS1 | 5742 |
| AGTGCGGCGAGGGCCTACCA | GPT2 | 84706 |
| AGTGCGGCTAGTGAGTCCGA | HCAR2 | 338442 |
| AGTGCTACTGAAACTTGCCT | NO_CURRENT_209 | NO_CURRENT_209 |
| AGTGCTCATGCTCGCTGCAG | SAP18 | 10284 |
| AGTGCTGGGAGCTCTGCTGG | CYP8B1 | 1582 |
| AGTGCTGTACAAAGAGACAG | CASP5 | 838 |
| AGTGCTGTGCCCAGCGCCTG | NOD2 | 64127 |
| AGTGGAGTATTTGCTGGGCC | CTNNBL1 | 56259 |
| AGTGGCGAGTAAGGAAGCGT | NO_CURRENT_210 | NO_CURRENT_210 |
| AGTGGCTCCCATGGCGCCCC | DNTTIP1 | 116092 |
| AGTGGGGCGCTAAGTGGGGG | NO_CURRENT_211 | NO_CURRENT_211 |
| AGTGGGTGACGGGACGGTGG | MOSPD1 | 56180 |
| AGTGGGTGTGAGGGGCGCCG | MAPK9 | 5601 |
| AGTGTCTGTGGTGGCCGCGG | MKL1 | 57591 |
| AGTGTGTATCACGTGACCCT | NO_CURRENT_212 | NO_CURRENT_212 |
| AGTGTTTGAAAAAAGGGCGG | NO_CURRENT_213 | NO_CURRENT_213 |
| AGTTAAGCTCTCTGAAAAGA | AIM2 | 9447 |
| AGTTAAGGACGTACTCGTCT | COX16 | 51241 |
| AGTTACTGTGTAGTTTGTAC | TLR1 | 7096 |
| AGTTCCCAGGGAATCCGCAT | NO_CURRENT_214 | NO_CURRENT_214 |
| AGTTCCGGCCAGGGTCGCCC | NQO1 | 1728 |
| AGTTCCTGAGGAGATGTTCA | PAXIP1 | 22976 |
| AGTTCTGTTCGATAGATGCC | NO_CURRENT_215 | NO_CURRENT_215 |
| AGTTGAATGGACCTCGACTA | NO_CURRENT_216 | NO_CURRENT_216 |
| AGTTGCTATAGCGACCGGGT | DYRK1A | 1859 |
| AGTTGCTGCCTTTAGAGAGC | GPR119 | 139760 |
| AGTTGCTTGCGGCGGAGGGC | TRERF1 | 55809 |
| AGTTGGAAACTTTCTCCTCC | OR7C2 | 26658 |
| AGTTGTTCGTTTCTCCCCGG | IRGM | 345611 |
| AGTTTCCTGTCCAGAAGTCT | NR1H3 | 10062 |
| ATAAAACCCTTGGAGTAACT | NO_CURRENT_217 | NO_CURRENT_217 |
| ATAAAAGTCTCATTAACCCA | NO_CURRENT_218 | NO_CURRENT_218 |
| ATAAGAGAAGTAGCCAGCAG | GBP5 | 115362 |
| ATAAGCCACACTACCCGCCT | NO_CURRENT_219 | NO_CURRENT_219 |
| ATAAGGGGAGCACAGTTAGG | NO_CURRENT_220 | NO_CURRENT_220 |

| | | |
|---|---|---|
| ATAATAATAAAAAGCCCCAT | DYRK1A | 1859 |
| ATAATGGGTAGTTAAAACCG | NO_CURRENT_221 | NO_CURRENT_221 |
| ATAATGTTTGTATAACCCGT | NO_CURRENT_222 | NO_CURRENT_222 |
| ATACAATACTTTGGCGCATA | NO_CURRENT_223 | NO_CURRENT_223 |
| ATACAGGACCTGATTGTGAG | NO_CURRENT_224 | NO_CURRENT_224 |
| ATACCAGATGCGTCCGCTTG | NO_CURRENT_225 | NO_CURRENT_225 |
| ATACCATGTAACGTTTAAGG | NO_CURRENT_226 | NO_CURRENT_226 |
| ATACCTGGGAGACCCTTGGA | HSH2D | 84941 |
| ATACCTTTCCTTCCACTGCA | EBI3 | 10148 |
| ATACGAGGCGCTTTTCTTTG | NO_CURRENT_227 | NO_CURRENT_227 |
| ATACGCATGATTGCAAGAGG | NO_CURRENT_228 | NO_CURRENT_228 |
| ATACTATCACATAATCTGAG | NO_CURRENT_229 | NO_CURRENT_229 |
| ATACTCTCACAGGTACATAA | NO_CURRENT_230 | NO_CURRENT_230 |
| ATAGAGATAAGACTCATGGC | NO_CURRENT_231 | NO_CURRENT_231 |
| ATAGCAGGACGAGGTTCCTT | NO_CURRENT_232 | NO_CURRENT_232 |
| ATAGCCGCCGCTCATTACTT | NO_CURRENT_233 | NO_CURRENT_233 |
| ATAGCGAGGCTTGTTCAAAG | NO_CURRENT_234 | NO_CURRENT_234 |
| ATAGCGGATGTCCTTGGAAA | NO_CURRENT_235 | NO_CURRENT_235 |
| ATAGCTAATGCATGCATGCA | NO_CURRENT_236 | NO_CURRENT_236 |
| ATAGGATGACCCCTTTCTCT | UBE2L6 | 9246 |
| ATAGGCACCTTAAGGGTCTC | NO_CURRENT_237 | NO_CURRENT_237 |
| ATAGGGAAGAACATTGTAGT | NO_CURRENT_238 | NO_CURRENT_238 |
| ATAGGTCATCCACTGGGCGG | NO_CURRENT_239 | NO_CURRENT_239 |
| ATAGTAACGTCAGGGAGTAA | NO_CURRENT_240 | NO_CURRENT_240 |
| ATAGTATGTTAGGCAAAGCG | NO_CURRENT_241 | NO_CURRENT_241 |
| ATAGTGAGGACAACTAGTGA | NO_CURRENT_242 | NO_CURRENT_242 |
| ATAGTGTATTTGACTTAGTA | NO_CURRENT_243 | NO_CURRENT_243 |
| ATATAAAAAAGAGTTCTGAT | NO_CURRENT_244 | NO_CURRENT_244 |
| ATATAAACTGTCGCGGTAAA | NO_CURRENT_245 | NO_CURRENT_245 |
| ATATAGAGGAGAGGATCGTA | NO_CURRENT_246 | NO_CURRENT_246 |
| ATATATTTACAGGCTGGCTC | BCL2A1 | 597 |
| ATATTCAAAATGGCGGACGG | POU2F1 | 5451 |
| ATATTCACACTCATTTAGAC | NO_CURRENT_247 | NO_CURRENT_247 |
| ATATTGGAGCAGCAAGAGGC | MMP1 | 4312 |
| ATCACCCACACCTGGAACTT | HTR1D | 3352 |
| ATCACGTGATCGGATGGTTC | NO_CURRENT_248 | NO_CURRENT_248 |
| ATCACTCCTACATTCCTGGT | NO_CURRENT_249 | NO_CURRENT_249 |
| ATCAGGTACGGGCCCTGAGA | STAT2 | 6773 |
| ATCAGTTGTCAATATAAGGG | NO_CURRENT_250 | NO_CURRENT_250 |
| ATCATCTCCAGCGCGTGGTG | STAG2 | 10735 |
| ATCATGCCTTAGTCTTTCAG | NAIP | 4671 |
| ATCATGGACTAGACTAGCCT | NO_CURRENT_251 | NO_CURRENT_251 |
| ATCCCAGATACATACTCGGG | NO_CURRENT_252 | NO_CURRENT_252 |
| ATCCGCTCCGCTCTGCTCCC | WDR26 | 80232 |
| ATCCGGTGGAGAGCGAGATC | NFKB2 | 4791 |
| ATCCTAAACCCTTATATGGT | NO_CURRENT_253 | NO_CURRENT_253 |
| ATCCTCCCGCCCTCCTGACG | EP400 | 57634 |
| ATCCTCTGGACTTGGTAAGA | GBP7 | 388646 |
| ATCCTGGAGCGAGTGCTGGG | METTL3 | 56339 |
| ATCGATATACCGCCATAAAA | NO_CURRENT_254 | NO_CURRENT_254 |

| | | |
|---|---|---|
| ATCGCGCTCGAAGCCCCGGT | CREBBP | 1387 |
| ATCGGCCCCGAGTTACCGTC | SHOC2 | 8036 |
| ATCGGCTCAGAACTCCAAGC | COX16 | 51241 |
| ATCGGCTTCCAGTCCGCGGA | RAC1 | 5879 |
| ATCGGGGCGGTGTGTGGCCA | UCHL5 | 51377 |
| ATCGGTAACCGGCGACTACG | BABAM2 | 9577 |
| ATCGGTACCTCTTCACATAT | NO_CURRENT_255 | NO_CURRENT_255 |
| ATCGTATCATCAGCTAGCGC | NO_CURRENT_256 | NO_CURRENT_256 |
| ATCGTTGCTGACAGGATCTA | NO_CURRENT_257 | NO_CURRENT_257 |
| ATCTAAAAAGCTCCTAGACC | NO_CURRENT_258 | NO_CURRENT_258 |
| ATCTCGGGTCGACTGCGGAT | NO_CURRENT_259 | NO_CURRENT_259 |
| ATCTCTATAAGACATACTCA | NO_CURRENT_260 | NO_CURRENT_260 |
| ATCTGAGTCATATCAAGTTG | NO_CURRENT_261 | NO_CURRENT_261 |
| ATCTGGTCTACAAGAACTCG | NLRC4 | 58484 |
| ATCTGTGTGACTGCGGTCGG | NO_CURRENT_262 | NO_CURRENT_262 |
| ATCTTATCAACCAGTCACGG | NO_CURRENT_263 | NO_CURRENT_263 |
| ATCTTCTCGACGAAAATGCG | NO_CURRENT_264 | NO_CURRENT_264 |
| ATCTTTGTCAGTGCACAAAA | ACO2 | 50 |
| ATGACAGTGGCTTTGCCCGT | MSL2 | 55167 |
| ATGACATTGCGCGTCTACGG | NO_CURRENT_265 | NO_CURRENT_265 |
| ATGAGAAAGGGCATGACTCA | NO_CURRENT_266 | NO_CURRENT_266 |
| ATGAGCCGTGAGTGCGACCC | PTGS1 | 5742 |
| ATGAGCTGCACATCACAATG | NO_CURRENT_267 | NO_CURRENT_267 |
| ATGAGTAAGCTGTGGCGGCG | INO80B | 83444 |
| ATGATAATATTAACCTCATA | NO_CURRENT_268 | NO_CURRENT_268 |
| ATGATATACAATCTCACTAA | NO_CURRENT_269 | NO_CURRENT_269 |
| ATGATTAATTATCTGCACGG | NO_CURRENT_270 | NO_CURRENT_270 |
| ATGCAAGACAGCCTCCCAGC | NO_CURRENT_271 | NO_CURRENT_271 |
| ATGCAATGGAAGGTGTGCTG | NLRP12 | 91662 |
| ATGCACCGCTGAAGGCAGAG | BMI1 | 648 |
| ATGCAGGGATCGCTCCCCCG | SCN5A | 6331 |
| ATGCCTACGGTTCCTAACCC | LAMTOR2 | 28956 |
| ATGCCTTACCTGCCCGGGCG | IRAK4 | 51135 |
| ATGCCTTAGACTTAACCTCG | NO_CURRENT_272 | NO_CURRENT_272 |
| ATGCGAACGGGAAGGAGCGT | LARP4B | 23185 |
| ATGCGCAGCTCCAGAATTTT | NO_CURRENT_273 | NO_CURRENT_273 |
| ATGCTAATGATTGTAGCTCA | NO_CURRENT_274 | NO_CURRENT_274 |
| ATGCTACTAATGATAGATCC | NO_CURRENT_275 | NO_CURRENT_275 |
| ATGCTCGCTGCAGATGCGGT | FOS | 2353 |
| ATGCTCGCTGCAGGGGTCGG | SAP18 | 10284 |
| ATGCTGCAGCTTTACGATCA | NO_CURRENT_276 | NO_CURRENT_276 |
| ATGGAAATCATTGAAATACC | SERPINB2 | 5055 |
| ATGGAAGAGCGTCATGACTT | NO_CURRENT_277 | NO_CURRENT_277 |
| ATGGACTCTGGCAACTTATG | NO_CURRENT_278 | NO_CURRENT_278 |
| ATGGCCACGGTCCCAGCACC | IL2RB | 3560 |
| ATGGCGGCTTCCTAGTGAGT | BABAM1 | 29086 |
| ATGGCGGGGCCCGAGCGACG | RAD23A | 5886 |
| ATGGCTGCCGGAACAGCAGT | AGER | 177 |
| ATGGCTTGGGCCGCGCTCCT | PTGIS | 5740 |
| ATGGGGAGAAGCGGAGAACC | STRAP | 11171 |

| | | |
|---|---|---|
| ATGGGGCTCAGCAGACGCGT | CXCL3 | 2921 |
| ATGGGTTGCAGTCAAACCAC | NO_CURRENT_279 | NO_CURRENT_279 |
| ATGGTATAGGCGATGCTGAT | HTR1D | 3352 |
| ATGGTGCCGATTCATGAGTG | HCAR2 | 338442 |
| ATGGTGGCTCAACCCCTACC | SLC3A2 | 6520 |
| ATGTAACGAGTTGTAAGTCA | NO_CURRENT_280 | NO_CURRENT_280 |
| ATGTAATAGGGCCCCAACAG | NO_CURRENT_281 | NO_CURRENT_281 |
| ATGTACAAGCTGTTACATAA | NO_CURRENT_282 | NO_CURRENT_282 |
| ATGTGTCTAGTAAGTGACAA | NO_CURRENT_283 | NO_CURRENT_283 |
| ATGTTCATCATCAACCTCCC | CUBN | 8029 |
| ATTAAACTTTAACTAGACAT | NO_CURRENT_284 | NO_CURRENT_284 |
| ATTACGGCGCTGACCTTCCC | NDUFB9 | 4715 |
| ATTACTTCGCTCCGTCTGCC | ZNF616 | 90317 |
| ATTAGCACGGCGACCTTCTA | NO_CURRENT_285 | NO_CURRENT_285 |
| ATTAGCCGTTGCCATATCAA | NO_CURRENT_286 | NO_CURRENT_286 |
| ATTAGGCCTTTTTCTTAACT | NO_CURRENT_287 | NO_CURRENT_287 |
| ATTATGTAGACCATGGAATG | NO_CURRENT_288 | NO_CURRENT_288 |
| ATTCAAACTTAGTCTTGAGT | NO_CURRENT_289 | NO_CURRENT_289 |
| ATTCAACCTCTGCCACCATG | CYBB | 1536 |
| ATTCACAGCCCAGTTCCCCA | CYBB | 1536 |
| ATTCAGGAGCTGGATGGCGT | PPARGC1A | 10891 |
| ATTCATGCGCCGCCTCCTCT | NO_CURRENT_290 | NO_CURRENT_290 |
| ATTCCAACAATAATAGGGCA | NCKAP1L | 3071 |
| ATTCCGGAGCCCCTGCGTGG | ACACA | 31 |
| ATTCCTGCTTTAAAATCTCT | NOS2 | 4843 |
| ATTGACTCATTAATGAAGAC | NO_CURRENT_291 | NO_CURRENT_291 |
| ATTGAGAATTCGTTTCAAGG | NO_CURRENT_292 | NO_CURRENT_292 |
| ATTGCATCATTCACAGGCAG | NLRP12 | 91662 |
| ATTGCTCTGTCGCATCAATC | NO_CURRENT_293 | NO_CURRENT_293 |
| ATTGGGCCATGGTTCTGGAA | ATP5L | 10632 |
| ATTGGGCTAAATTGTTCTGG | NO_CURRENT_294 | NO_CURRENT_294 |
| ATTGTATCTTCATACTTGTG | NO_CURRENT_295 | NO_CURRENT_295 |
| ATTGTGGGATGTGCGGCTAC | SETD1B | 23067 |
| ATTGTGTGCTGGACTAAGGT | NO_CURRENT_296 | NO_CURRENT_296 |
| ATTTAACATCATATTACACC | NO_CURRENT_297 | NO_CURRENT_297 |
| ATTTAGTAATGCACACCCAG | NO_CURRENT_298 | NO_CURRENT_298 |
| ATTTAGTAGACATTGGGTCT | NO_CURRENT_299 | NO_CURRENT_299 |
| ATTTCCAATACACCCTACAT | NO_CURRENT_300 | NO_CURRENT_300 |
| ATTTCTTACAGACTGCCAAA | TLR1 | 7096 |
| ATTTGCTGGGCCGGGTACCA | CTNNBL1 | 56259 |
| ATTTGCTTTCCGTTTCAGTG | NLRC4 | 58484 |
| ATTTGGATATATTTACAGGC | BCL2A1 | 597 |
| ATTTGTACCGGAGTCCCATT | EDA2R | 60401 |
| CAAAAATGTTCCTCAGACGC | ZNF699 | 374879 |
| CAAAACAAAAGGGAAACCCG | FOXO4 | 4303 |
| CAAAAGTAGAAGGTAACATT | NO_CURRENT_301 | NO_CURRENT_301 |
| CAAAAGTCTCGCTTGGTCCT | NO_CURRENT_302 | NO_CURRENT_302 |
| CAAAATCTAGCAATTTTTGG | NO_CURRENT_303 | NO_CURRENT_303 |
| CAAAATTGTTAGTAAACTGT | NO_CURRENT_304 | NO_CURRENT_304 |
| CAAACAGCCTTACCTGGGGC | ATP5J2 | 9551 |

| | | |
|---|---|---|
| CAAACAGCCTTACCTGGGGC | ATP5J2-PTCD1 | 100526740 |
| CAAACAGCCTTACCTGGGGC | PTCD1 | 26024 |
| CAAACATCGCGATTAATAGG | BTN2A2 | 10385 |
| CAAACTCTTGGTACAGCACA | CCL20 | 6364 |
| CAAAGCAGCAGCCCAGCACC | MMP9 | 4318 |
| CAAAGCCAAGTATTGAAGCA | NO_CURRENT_305 | NO_CURRENT_305 |
| CAAAGCCGAGCTCTGGGCAG | SEC62 | 7095 |
| CAAAGCCGTGCGGAGATTGG | APIP | 51074 |
| CAAAGGTTCCACAGCCTGGA | GBP5 | 115362 |
| CAAAGTTGTAATTGTGTCAC | UTY | 7404 |
| CAAATCTCCATATACAGCCC | TNRC18 | 84629 |
| CAAATGCCATTTAGGTTATC | NO_CURRENT_306 | NO_CURRENT_306 |
| CAAATGGGCTGCTGGCGGCG | KPNB1 | 3837 |
| CAACAAGCACACGCCCATGA | NDUFA1 | 4694 |
| CAACACACCAGGGAGCAGAG | WDR26 | 80232 |
| CAACACCCCGCGTTATGCTA | NO_CURRENT_307 | NO_CURRENT_307 |
| CAACCACAATAACAGGCGGA | FBXO38 | 81545 |
| CAACCCAGGAGGCACGACGC | PDHA1 | 5160 |
| CAACCCGTCGGCGCGGCCTC | CS | 1431 |
| CAACGACGGGCCTAGTCTCA | NO_CURRENT_308 | NO_CURRENT_308 |
| CAACGCCGCTGCTCTTGGAG | DLAT | 1737 |
| CAAGACAGACTTGCAAAAGA | IL10 | 3586 |
| CAAGACCTTATCGTGCAGCG | NO_CURRENT_309 | NO_CURRENT_309 |
| CAAGACTGAGCCGGCGGCCG | FOS | 2353 |
| CAAGAGCAACAAGGACCCCC | HSPA13 | 6782 |
| CAAGATGCATCCAGGGGTCC | MMP13 | 4322 |
| CAAGATTTGGCCAAAGTTCC | HTR1D | 3352 |
| CAAGCTCAAGACCCAGCAGT | HAMP | 57817 |
| CAAGCTCAGCCCACGTCTGA | SPHK1 | 8877 |
| CAAGCTCCAAACTCCCGCCG | ERCC6L | 54821 |
| CAAGGAGGTGTGTGTCTGTG | HCAR2 | 338442 |
| CAAGGATGGAAATTAAAACG | NO_CURRENT_310 | NO_CURRENT_310 |
| CAAGGTGGGTGTTCACGTTT | NDUFS8 | 4728 |
| CAAGTAGCGGCGGCGCTTCA | NDUFS8 | 4728 |
| CAAGTGAGCATAAGCGATGT | NO_CURRENT_311 | NO_CURRENT_311 |
| CAATAATACAGTTTATTCAG | NO_CURRENT_312 | NO_CURRENT_312 |
| CAATATCGGGTGCTACAGGA | NO_CURRENT_313 | NO_CURRENT_313 |
| CAATCGGCGACGTTTTAAAT | NO_CURRENT_314 | NO_CURRENT_314 |
| CAATCTCAATATAACTGTGT | NO_CURRENT_315 | NO_CURRENT_315 |
| CAATCTGCTTCCGGGAGTGA | ZNF616 | 90317 |
| CAATGCCTAAAGGAGGTGAG | PDAP1 | 11333 |
| CACAAGCAGATTTATCACAA | NO_CURRENT_316 | NO_CURRENT_316 |
| CACAAGGTTACGGACAAATT | ATP5L | 10632 |
| CACAAGTAGTTTACATTGTT | RXRA | 6256 |
| CACAATAACAGGCGGAGGGT | FBXO38 | 81545 |
| CACAATAGCACCCATCCGCT | CASP3 | 836 |
| CACACCATAGAGGTGTGACA | NO_CURRENT_317 | NO_CURRENT_317 |
| CACACGGCCCGAAAAGTCGC | PHF8 | 23133 |
| CACACTCACCCTCGGTGTGC | ITGB2 | 3689 |
| CACACTCGGTCCCGACATGA | UBE2L6 | 9246 |

| | | |
|---|---|---|
| CACACTGACAGCTATAGGGC | PHIP | 55023 |
| CACAGACCACCCAATCAGAA | NO_CURRENT_318 | NO_CURRENT_318 |
| CACAGAGCACACACTCATGC | PPARGC1A | 10891 |
| CACAGCAGTGCCGACGTCGT | LGMN | 5641 |
| CACAGCCATGGCGGGCGCGT | MPC1 | 51660 |
| CACAGCCCAGTTCCCCATGG | CYBB | 1536 |
| CACATACATCAACCAACCCT | NO_CURRENT_319 | NO_CURRENT_319 |
| CACATGGGGTACAGCACACC | NO_CURRENT_320 | NO_CURRENT_320 |
| CACCACAGCGCCTGCTTCCT | LCN2 | 3934 |
| CACCACCAGCGCAGCAGTCC | DNAJB6 | 10049 |
| CACCACGCGCTGGAGATGAT | STAG2 | 10735 |
| CACCAGGAGCACCAGGACCA | MMP9 | 4318 |
| CACCAGGGAGCAGAGCGGAG | WDR26 | 80232 |
| CACCAGGTGAGGCCCGGCCG | RHOG | 391 |
| CACCATTCAAGATGCATCCA | MMP13 | 4322 |
| CACCCCTGAAAAGCAGCAGC | JAM3 | 83700 |
| CACCCGGAACCACTACTGAG | NO_CURRENT_321 | NO_CURRENT_321 |
| CACCCTTATATTCAGTAACT | NO_CURRENT_322 | NO_CURRENT_322 |
| CACCGCCGCCGCGCACCGCC | MAP2K7 | 5609 |
| CACCGGACTCAAATAATGAC | NO_CURRENT_323 | NO_CURRENT_323 |
| CACCGTGAAGGCCTACCTTC | SQSTM1 | 8878 |
| CACCTCCAGCTCCTGCTCGC | GSDMD | 79792 |
| CACCTCCCAGCACTCGCTCC | METTL3 | 56339 |
| CACCTCCGATTCCGAGCTTT | HDAC2 | 3066 |
| CACCTGCGTCCGCGGCGGCT | CYFIP1 | 23191 |
| CACCTTCGCTGGCCGCCCGC | TRAF6 | 7189 |
| CACGAACTCACACCGCGCGA | NO_CURRENT_324 | NO_CURRENT_324 |
| CACGACGGGCGGGTGAAGAT | PRKAA1 | 5562 |
| CACGACGGGCGGGTGAAGAT | PRKAA2 | 5563 |
| CACGCACAATCCTTCACGCA | NO_CURRENT_325 | NO_CURRENT_325 |
| CACGCAGCGCGGCCCGAAGA | AKT1 | 207 |
| CACGCCAACTAAAACTGCAG | NO_CURRENT_326 | NO_CURRENT_326 |
| CACGGGCCGCAGCGCCTCTG | CACNA2D2 | 9254 |
| CACGGTGTGAGCGCCCGACG | EGFR | 1956 |
| CACTAGCAGCATGTTGAGCC | SOD2 | 6648 |
| CACTAGCCGCACTCATGAAT | HCAR2 | 338442 |
| CACTAGGCGAGGCGCTCCAT | HCAR2 | 338442 |
| CACTAGGCGAGGCGCTCCAT | HCAR3 | 8843 |
| CACTATCTCGGGCATGGCTC | LTA4H | 4048 |
| CACTCTCCGCGCCCGTTCTC | SLC25A6 | 293 |
| CACTCTGGAGGCGTACTTGA | CYP27B1 | 1594 |
| CACTGCAGCTGAATGAGTTG | MBNL1 | 4154 |
| CACTGCAGTATTCGTGGCCT | NO_CURRENT_327 | NO_CURRENT_327 |
| CACTGCTGCTGCTGCTGTTC | MMP1 | 4312 |
| CACTGGACCCCAGAGAACCA | CYP8B1 | 1582 |
| CACTGGCCGAAGCGACGAAG | HIF1A | 3091 |
| CACTGGCTACAGCAGGGCAC | UBA7 | 7318 |
| CACTGGGTCCGGTGCGGCAC | ACTR5 | 79913 |
| CAGAAAGTCCTAGCAAACAG | CXCR1 | 3577 |
| CAGAACACTGCAGGAGGCCT | CAT | 847 |

| | | |
|---|---|---|
| CAGAAGTTCGCCCACGTCCA | CTNNBL1 | 56259 |
| CAGAATAACCCGCCCCCAGC | MGA | 23269 |
| CAGACCCAGTAAAACCACCA | NO_CURRENT_328 | NO_CURRENT_328 |
| CAGACGCGGCTCCTAGCGGA | CASP3 | 836 |
| CAGACGGTTGGTAAGGACGC | NO_CURRENT_329 | NO_CURRENT_329 |
| CAGACTCTGGGCGCCACTCC | INO80E | 283899 |
| CAGACTGGACAGCAGCTACA | TLR9 | 54106 |
| CAGAGAGGGGGTGCAGGCTG | SLC11A1 | 6556 |
| CAGAGCAGCCAGTGTCCGGG | TMEM173 | 340061 |
| CAGAGCCATTGGAGGGCGCG | STAT2 | 6773 |
| CAGAGCCTTGCGCAATTTTG | NO_CURRENT_330 | NO_CURRENT_330 |
| CAGAGCTCGGCTTTGCTACC | SEC62 | 7095 |
| CAGAGGCTGTTAGCTATGGC | CASP5 | 838 |
| CAGAGTACCTAGTGGCCGCC | PPID | 5481 |
| CAGAGTCCAGCGGAGTTGTG | DNTTIP1 | 116092 |
| CAGAGTCCTCTGCGAATCCC | OR7C2 | 26658 |
| CAGAGTTGGAGGGTTGAAAG | NO_CURRENT_331 | NO_CURRENT_331 |
| CAGATCCCCTCACACGAGGA | MCTS1 | 28985 |
| CAGATGATGGTCGTCCTCCT | IFNAR1 | 3454 |
| CAGATGGCTGCCCCGGCGAG | FOXO4 | 4303 |
| CAGCAAGGGTCAGAACTCAC | CD84 | 8832 |
| CAGCACCTCCTCTCTCCTGT | GC | 2638 |
| CAGCACTTAGATATAAACTC | NO_CURRENT_332 | NO_CURRENT_332 |
| CAGCAGATCACCGGCCACGG | NDUFA12 | 55967 |
| CAGCAGATCGGCGGCATCAG | SOD2 | 6648 |
| CAGCAGCCGCCGCGGCGGCA | UBE2E1 | 7324 |
| CAGCAGCGTTGGCCCGGCCC | CTNNB1 | 1499 |
| CAGCAGGGAGCTGGGAGCTG | JUNB | 3726 |
| CAGCCAGACAGACGGCACGA | HAMP | 57817 |
| CAGCCAGACAGCGAGGGCCC | TGFB1 | 7040 |
| CAGCCAGCCGGGCACTCGGG | MEX3B | 84206 |
| CAGCCAGCGTTGACCCTCCA | UCHL3 | 7347 |
| CAGCCCAGCGCCTTCTCTCC | TNFRSF1B | 7133 |
| CAGCCCAGTAGGCAATAGCA | NO_CURRENT_333 | NO_CURRENT_333 |
| CAGCCCGCTCCGAGCGCTGA | BCL2 | 596 |
| CAGCCCGGAGTCGGCCTTGT | MAPK14 | 1432 |
| CAGCCCTGCCACTGCCAGCC | CYP4F11 | 57834 |
| CAGCCGCAGCGGCCAGGCAG | ABCB10 | 23456 |
| CAGCCGCCGCGGCGGCACGG | UBE2E1 | 7324 |
| CAGCCGCGGCTGGAAGATGG | PPARGC1B | 133522 |
| CAGCCGGCGGTTCGGGGCGA | ATP5L | 10632 |
| CAGCCTATTTTGCTACCTAC | NO_CURRENT_334 | NO_CURRENT_334 |
| CAGCCTCTGAGGTGAGGGCG | ARF1 | 375 |
| CAGCGAGCCGGGGCAGGAAG | NFKB1 | 4790 |
| CAGCGCCGTAATGGCGTTCT | NDUFB9 | 4715 |
| CAGCGGAACGGGAGGACAGC | SLC25A6 | 293 |
| CAGCGGAGTTGTGGGGGCCG | DNTTIP1 | 116092 |
| CAGCGGCTTCAGCAGATCGG | SOD2 | 6648 |
| CAGCGGGAGGTCGCGAAGCC | ACADS | 35 |
| CAGCGGGATGTGAAGGACTC | MGA | 23269 |

| | | |
|---|---|---|
| CAGCGGGGAAGGCGGGCGGT | MTOR | 2475 |
| CAGCGTTGAGATTGAGACTG | CYP1B1 | 1545 |
| CAGCTCCAGGACTGCTGCGC | DNAJB6 | 10049 |
| CAGCTCCAGGAGGGTGAGCG | HELZ2 | 85441 |
| CAGCTCCCCCGCCTCGGGGA | RHOG | 391 |
| CAGCTCGCTGGCTCCCGCCG | RB1 | 5925 |
| CAGCTGCAGGGGAGGAGGAC | RNF25 | 64320 |
| CAGCTGCTTTCCGAGGAAGC | MOSPD1 | 56180 |
| CAGCTTGTCCACCCGCCGGC | VDR | 7421 |
| CAGGAAGAAGCCGCGGGGAC | TYK2 | 7297 |
| CAGGAATTCAGCTGCTGGAG | SERPINE1 | 5054 |
| CAGGACCGGGAGAGCAGTGG | NLRP3 | 114548 |
| CAGGACTGCTGCGCTGGTGG | DNAJB6 | 10049 |
| CAGGAGAAAATATCCACCCA | NO_CURRENT_335 | NO_CURRENT_335 |
| CAGGAGAGTCAACACCAGAG | NO_CURRENT_336 | NO_CURRENT_336 |
| CAGGAGCTGAGCCTAAGCCC | KDM4A | 9682 |
| CAGGAGTCCTGGGGCATGGC | DET1 | 55070 |
| CAGGATGGACGTGAACGCGT | NO_CURRENT_337 | NO_CURRENT_337 |
| CAGGCAGATTTGCCTGCTGA | CAT | 847 |
| CAGGCAGCCCCAGCCTCCGG | IZUMO2 | 126123 |
| CAGGCAGCCGAGCGCAGAGC | JAM3 | 83700 |
| CAGGCAGCCGGACGGGCGGG | BAHD1 | 22893 |
| CAGGCAGCGTTGCAAGGGGA | AHCTF1 | 25909 |
| CAGGCCAAGGCGCACCCACC | PTGES2 | 80142 |
| CAGGCCCGGCACCCGCTTCC | ASH2L | 9070 |
| CAGGCGCTGTGGTGGCTGCT | LCN2 | 3934 |
| CAGGCTGGGCGGCGGAGCCT | CHMP5 | 51510 |
| CAGGCTTAAAACTTGTTGGA | RND3 | 390 |
| CAGGGAAGTGGAGTATTTGC | CTNNBL1 | 56259 |
| CAGGGACCTCTGGGTTCACG | ACO2 | 50 |
| CAGGGAGGTCAAGTATTACG | PAXIP1 | 22976 |
| CAGGGCAGGCAGCTCCAGGA | HELZ2 | 85441 |
| CAGGGCCAGGCGCGTTTCTG | RAC2 | 5880 |
| CAGGGCCCAGCCCGACACAC | HIF1A | 3091 |
| CAGGGCCGGGGCCTGAACCG | NO_CURRENT_338 | NO_CURRENT_338 |
| CAGGGCTCTGCGCACCGCTG | SIAH2 | 6478 |
| CAGGGGGGTGTCATAGCCCC | IKBKB | 3551 |
| CAGGGTGGACGCATGCCCTC | CYP2R1 | 120227 |
| CAGGTAAGATCAGCCAAGGA | IRF9 | 10379 |
| CAGGTAGACCTGGCGACGAC | PRR14L | 253143 |
| CAGGTAGCAAAGCCGAGCTC | SEC62 | 7095 |
| CAGGTAGCAAAGTGACGCCG | CXCR4 | 7852 |
| CAGGTAGTTCTTAAAAGGAG | ACOD1 | 730249 |
| CAGGTCGGTGCGTCTGTCGG | BABAM2 | 9577 |
| CAGGTTGGCAGGTAAGAGTG | CUBN | 8029 |
| CAGGTTTGCACGCATAGCTA | NO_CURRENT_339 | NO_CURRENT_339 |
| CAGTCAGTCAGGCTGGGCGG | CHMP5 | 51510 |
| CAGTCGTGGACTCGTGCAGC | TMEM70 | 54968 |
| CAGTGGCGGCGACGGCGAGG | RNF111 | 54778 |
| CAGTGGGACAGCCAGACAGA | HAMP | 57817 |

| | | |
|---|---|---|
| CAGTGGGTGTGAGGGGCGCC | MAPK9 | 5601 |
| CAGTGTCTCTACACAGTCTC | CASP5 | 838 |
| CAGTTGCGATGAGGGCTGCG | TIRAP | 114609 |
| CAGTTGGAAGATGGCGGACG | SIRT1 | 23411 |
| CAGTTTCCCGGCTCTCCGCG | NDUFB9 | 4715 |
| CATAAGGTTAACTGCGTGGA | NO_CURRENT_340 | NO_CURRENT_340 |
| CATACCCACATCAAACACTC | NO_CURRENT_341 | NO_CURRENT_341 |
| CATACCTTTCCTTCCACTGC | EBI3 | 10148 |
| CATATAAATTCAGTAGTCCA | NO_CURRENT_342 | NO_CURRENT_342 |
| CATATAAGTGCTTACCAATG | NO_CURRENT_343 | NO_CURRENT_343 |
| CATCAAATTAAGGTAATATG | NO_CURRENT_344 | NO_CURRENT_344 |
| CATCAACCTCCCAGGTTGGC | CUBN | 8029 |
| CATCACCCAGGAGTCTGCTG | CYP4A11 | 1579 |
| CATCAGGTGTATTCTAAGTG | NO_CURRENT_345 | NO_CURRENT_345 |
| CATCATATATTGACTAAGGT | NO_CURRENT_346 | NO_CURRENT_346 |
| CATCATCTCCAGCGCGTGGT | STAG2 | 10735 |
| CATCATTCACAGGCAGCGGG | NLRP12 | 91662 |
| CATCGCCTTCCTCCCCAGGG | SPINT1 | 6692 |
| CATCGGCTCAGAACTCCAAG | COX16 | 51241 |
| CATCGGCTTCCAGTCCGCGG | RAC1 | 5879 |
| CATCTGTAGGGTTGCAAGCC | NO_CURRENT_347 | NO_CURRENT_347 |
| CATGAGCACAAAATTTAGGG | NO_CURRENT_348 | NO_CURRENT_348 |
| CATGAGCCTCTGGCAGCCCC | MMP9 | 4318 |
| CATGATGGCGAGCATGCGAG | UBE2L6 | 9246 |
| CATGCCACCTCCTTTGCCCC | TNFRSF6B | 8771 |
| CATGGATCACACTGGTAAGG | CBLL1 | 79872 |
| CATGGCATAAGTATAAGACA | NO_CURRENT_349 | NO_CURRENT_349 |
| CATGGCCTACGGTGTCTTTG | NO_CURRENT_350 | NO_CURRENT_350 |
| CATGGCGGGACAGGAGGATC | BRK1 | 55845 |
| CATGGGAGATAATAGACAAG | NO_CURRENT_351 | NO_CURRENT_351 |
| CATGGGATTAGTGGCTAACG | NO_CURRENT_352 | NO_CURRENT_352 |
| CATGGTCACCTACAGGAGAG | GC | 2638 |
| CATGGTCCGCGACGGTCGCA | ARRB1 | 408 |
| CATGTAACTGGTGGGATCTG | IFNAR1 | 3454 |
| CATGTCGGTCAGGCACAGCA | TNFRSF6B | 8771 |
| CATGTGACGGTGGCTTGAGG | LAMTOR3 | 8649 |
| CATGTTGTGTGAGGATCCCG | RAD23A | 5886 |
| CATTCAACCTCTGCCACCAT | CYBB | 1536 |
| CATTCCCCGCCTTAATGCCT | IRAK4 | 51135 |
| CATTCGTTGAAAAAGGACTG | NO_CURRENT_353 | NO_CURRENT_353 |
| CATTCTGGGTGACACGCTGG | PRKAA1 | 5562 |
| CATTGACAATTACTTAACCT | NO_CURRENT_354 | NO_CURRENT_354 |
| CATTGCACGCCACAGCATTG | NO_CURRENT_355 | NO_CURRENT_355 |
| CATTGGAAATTAACTATGGG | NO_CURRENT_356 | NO_CURRENT_356 |
| CATTGTATGAACGCAATAGC | NO_CURRENT_357 | NO_CURRENT_357 |
| CATTGTTGAGCGGGCGCGCT | NO_CURRENT_358 | NO_CURRENT_358 |
| CCAAAAAGATGAATATCTCG | NO_CURRENT_359 | NO_CURRENT_359 |
| CCAAAGCCCCGCCAGGGCTT | KDM4A | 9682 |
| CCAAAGGTAACAACTATGCT | NO_CURRENT_360 | NO_CURRENT_360 |
| CCAAATCTCCATATACAGCC | TNRC18 | 84629 |

| | | |
|---|---|---|
| CCAACAAACTGCAGACCAGG | BCL2L11 | 10018 |
| CCAAGATGGCGGCGGCGCTG | INO80 | 54617 |
| CCAAGCAGCCGGACACACGG | FBXW7 | 55294 |
| CCAAGCCAGCCTGAAGCGCC | KIF2B | 84643 |
| CCAAGCTCCAAACTCCCGCC | ERCC6L | 54821 |
| CCAAGGATGGGAGTCAGCCT | IRF9 | 10379 |
| CCAAGTATTAAAGTAAACAG | NO_CURRENT_361 | NO_CURRENT_361 |
| CCAATGATAAGCCCGAACGG | NO_CURRENT_362 | NO_CURRENT_362 |
| CCACACCTGTCTAGCATGAC | NO_CURRENT_363 | NO_CURRENT_363 |
| CCACAGCAAGCAGTAGTACC | GC | 2638 |
| CCACCATGCTGTAGCGAAAG | ATP5E | 514 |
| CCACCCCCCACCCCTGGGTT | HSP90B1 | 7184 |
| CCACCCGAGGCCGCGCGGAC | MAP2K5 | 5607 |
| CCACCGCCATCTTCCTCCTA | SAP18 | 10284 |
| CCACCTTTGATGAGGGGACT | NOS2 | 4843 |
| CCACGATGCCACCTCATCCC | NO_CURRENT_364 | NO_CURRENT_364 |
| CCACGCCGCCTCGCCCTCCG | RAC1 | 5879 |
| CCACGGGGAGGTGTCAAGGA | CYP24A1 | 1591 |
| CCACGGTCCCAGCACCGGGG | IL2RB | 3560 |
| CCACTCACCCCCGGTAGGCT | ZNF641 | 121274 |
| CCACTGCTTGGGGTAGCGGG | INO80E | 283899 |
| CCAGAAAGTGATCCTGCAGA | HCAR2 | 338442 |
| CCAGAAAGTGATCCTGCAGA | HCAR3 | 8843 |
| CCAGAACGCTCGGTGAGAGG | DNAJA1 | 3301 |
| CCAGAATTGAGTGCAAAAAG | NO_CURRENT_365 | NO_CURRENT_365 |
| CCAGCACCCTCGGAACCCAG | SPINT1 | 6692 |
| CCAGCAGTGGCAACCCACTG | GBP5 | 115362 |
| CCAGCAGTTCCTCCACGCAG | ACACA | 31 |
| CCAGCCAGCCGGGCACTCGG | MEX3B | 84206 |
| CCAGCCCACCCCCAGGCGCT | NOD2 | 64127 |
| CCAGCCCGCCTGCCCAGCCC | SLC6A4 | 6532 |
| CCAGCCCTCAGGGCCCCGGG | SMPD1 | 6609 |
| CCAGCGAGCGAGCGAACGAG | FOSL2 | 2355 |
| CCAGCTCCCCCGCCTCGGGG | RHOG | 391 |
| CCAGCTCCCTGCTGGCTCCG | JUNB | 3726 |
| CCAGCTCCGCGGGGCAGTGT | NLRP3 | 114548 |
| CCAGCTTGGAGAGCAGAGAA | AHDC1 | 27245 |
| CCAGGAATTTGCTACCAACA | NO_CURRENT_366 | NO_CURRENT_366 |
| CCAGGACCAGGGGCTGCCAG | MMP9 | 4318 |
| CCAGGAGAGCCGCTGGTGGG | PRKN | 5071 |
| CCAGGCAGGGGGGCCTCGCA | ABCB10 | 23456 |
| CCAGGCGCCCAAAGTTCCTG | PAXIP1 | 22976 |
| CCAGGCTGAAGTTCGTACCT | NO_CURRENT_367 | NO_CURRENT_367 |
| CCAGGGCTCTGCGCACCGCT | SIAH2 | 6478 |
| CCAGGGGCGGGCGCCGCCAT | CHMP6 | 79643 |
| CCAGGTGTGGGTGATGGTAT | HTR1D | 3352 |
| CCAGTCCGCGCGGCCTCGGG | MAP2K5 | 5607 |
| CCAGTCCGCGGAGGGCGAGG | RAC1 | 5879 |
| CCAGTGATCTTGAACCCCAA | TNFRSF1A | 7132 |
| CCAGTGCCCTTTTGTCGCAA | NO_CURRENT_368 | NO_CURRENT_368 |

| | | |
|---|---|---|
| CCAGTTATAATTAGGGGTTT | NO_CURRENT_369 | NO_CURRENT_369 |
| CCAGTTGCTCTGGGGGAACA | NO_CURRENT_370 | NO_CURRENT_370 |
| CCATATCGGGGCGAGACATG | NO_CURRENT_371 | NO_CURRENT_371 |
| CCATCACCGATCGTGAGCCT | NO_CURRENT_372 | NO_CURRENT_372 |
| CCATCATCTCCAGCGCGTGG | STAG2 | 10735 |
| CCATCCAGAAGAAGGCACAC | PDE3A | 5139 |
| CCATCGCCTTCCTCCCCAGG | SPINT1 | 6692 |
| CCATCTACTCGTCCGTTACC | EDA2R | 60401 |
| CCATCTTCCAACTGCCTCTC | SIRT1 | 23411 |
| CCATGACAAAACCATAGCAA | NO_CURRENT_373 | NO_CURRENT_373 |
| CCATGCGAGGCCCCCCTGCC | ABCB10 | 23456 |
| CCATGCTCAGTTGGTTGGAG | CUBN | 8029 |
| CCATGCTCTGCGCACCGGCC | SLC7A5 | 8140 |
| CCATGGACGAAGCGGATCGG | CASP9 | 842 |
| CCATGGCGGGCGGGTCGCTG | RPS6KA4 | 8986 |
| CCATGGGCGACAAAGGGACC | ARRB1 | 408 |
| CCATGGGGCGGGAGGGCAAG | CHUK | 1147 |
| CCATGGTCGGGAGAGACACC | PCGF6 | 84108 |
| CCATGGTGGGCCCCGCGCCG | NENF | 29937 |
| CCATGTCGAGGGTTGCTGAG | JAM3 | 83700 |
| CCATTCCGTAAGGGCTTGGA | NO_CURRENT_374 | NO_CURRENT_374 |
| CCATTGTAGTGACTCATCTC | FOSL2 | 2355 |
| CCCAAGTGCAACGGGACCCA | FAM214B | 80256 |
| CCCAATGGCTTCTGCGTGAC | NO_CURRENT_375 | NO_CURRENT_375 |
| CCCACAATGCTCCCATGACA | MBNL1 | 4154 |
| CCCACCGCCACAAGGAGGCA | IKBKE | 9641 |
| CCCACCTGCCAGGCTGCGCG | SLC6A4 | 6532 |
| CCCACGGAACTACAGGTGTG | FAM214B | 80256 |
| CCCAGAACTGGAGTCCCAAA | ATP5J | 522 |
| CCCAGAGGCCCGCGCTCTGA | SPINT1 | 6692 |
| CCCAGCAGTTCCTCCACGCA | ACACA | 31 |
| CCCAGCCAGGAGCACCGCCG | ITGB2 | 3689 |
| CCCAGCCCACCCCCAGGCGC | NOD2 | 64127 |
| CCCAGCTGCTAGTCAGAAAA | CASP5 | 838 |
| CCCAGCTGGGACTCTAAGAG | NO_CURRENT_376 | NO_CURRENT_376 |
| CCCAGGCTGACTCCCATCCT | IRF9 | 10379 |
| CCCAGTCCCCTCATCAAAGG | NOS2 | 4843 |
| CCCAGTTGCAACGTTAGGGA | MAP2K6 | 5608 |
| CCCATCCCCCCTGGCAGCAG | PIAS1 | 8554 |
| CCCATGGCGGCGCCCGCCCC | CHMP6 | 79643 |
| CCCATGGCGGGCGGGTCGCT | RPS6KA4 | 8986 |
| CCCCAACTTTCGCGACTCCG | NO_CURRENT_377 | NO_CURRENT_377 |
| CCCCACACCTGTAGTTCCGT | FAM214B | 80256 |
| CCCCAGGAGCTGGGACTAGT | IKBKE | 9641 |
| CCCCAGTGGACACGCGGATG | ALOX15 | 246 |
| CCCCATGGCTCCAGGATCCC | PYCARD | 29108 |
| CCCCCAGCTCCCCCGCCTCG | RHOG | 391 |
| CCCCCGGTAGGCTTGGCCCG | ZNF641 | 121274 |
| CCCCCTGACCGCGATCATTC | NCKAP1L | 3071 |
| CCCCGACGACGACGCCGAGG | USP7 | 7874 |

| | | |
|---|---|---|
| CCCCGCCCCCGCCATGCCCC | DET1 | 55070 |
| CCCCGCCCCCGTGAACCCAG | ACO2 | 50 |
| CCCCGCTGGAGCGGACTCCC | SPHK1 | 8877 |
| CCCCGGACCATGGCGCTCTC | NOD1 | 10392 |
| CCCCGGAGCCAAGTTACTCA | ANP32B | 10541 |
| CCCCGGCCGCGCGGAGAGCC | NDUFB9 | 4715 |
| CCCCGGGAGCGGAGAGCGAG | CTNNB1 | 1499 |
| CCCCGGGCTCACCTGCGCGC | TIRAP | 114609 |
| CCCCGTAGCTCATTAGTCTG | NO_CURRENT_378 | NO_CURRENT_378 |
| CCCCTATGCAGACTACAATT | NO_CURRENT_379 | NO_CURRENT_379 |
| CCCCTCCGCAGGCGGACCGG | TNFRSF6B | 8771 |
| CCCCTCGCTCTCCGCTCCCG | CTNNB1 | 1499 |
| CCCCTCTCCACACCCGGACA | PAGR1 | 79447 |
| CCCCTGAAAACATTAAAATG | IKBKB | 3551 |
| CCCCTGAACTGTCTACACCC | NO_CURRENT_380 | NO_CURRENT_380 |
| CCCCTGGAGTCGCCGCGCTC | RIPK1 | 8737 |
| CCCGAAAATGTCTCAACCCC | LARP4B | 23185 |
| CCCGAATGTCGTTAGCCGTG | SUV39H1 | 6839 |
| CCCGACACCACCAAGCCGCA | NIPBL | 25836 |
| CCCGACCATGGAGGGGGTCG | PCGF6 | 84108 |
| CCCGACCTGCGGCGGGCCCC | SCTR | 6344 |
| CCCGACTCTCAGCTCCATGG | CSNK1D | 1453 |
| CCCGACTGTGGAGAAGTGTC | OXSM | 54995 |
| CCCGAGGCCGCGCGGACTGG | MAP2K5 | 5607 |
| CCCGAGTCCAGCCCCTCCCG | YPEL5 | 51646 |
| CCCGATGGACTATACCGAAC | NO_CURRENT_381 | NO_CURRENT_381 |
| CCCGCACTGTCTGCAGCTCC | JAK1 | 3716 |
| CCCGCCCTCACCTGAACCGC | NFRKB | 4798 |
| CCCGCCGAAGACCCTGCTTG | NO_CURRENT_382 | NO_CURRENT_382 |
| CCCGCCGCCCGGGCCCGGGC | RXRA | 6256 |
| CCCGCCTCACCCCCGCCTCT | BAX | 581 |
| CCCGCCTCGGGGAGGGCACC | RHOG | 391 |
| CCCGCCTCTGGGCTGCTCCC | BAX | 581 |
| CCCGCGCGCAGGTGAGCCCG | TIRAP | 114609 |
| CCCGCGCGCTGGTAGGAGGA | NR2C2 | 7182 |
| CCCGCGGGACTCACCCTGAG | YPEL5 | 51646 |
| CCCGCTCGCACGCACACACA | BMI1 | 648 |
| CCCGGAAGCAGATTGCGCCA | ZNF616 | 90317 |
| CCCGGACCATGGCGCTCTCC | NOD1 | 10392 |
| CCCGGACGCGGCGGATGAGC | SATB1 | 6304 |
| CCCGGAGAGCGCCATGGTCC | NOD1 | 10392 |
| CCCGGAGCCAAGTTACTCAC | ANP32B | 10541 |
| CCCGGAGCCTCCCGGCCCTG | TRIR | 79002 |
| CCCGGAGTGGGCCTTGGAGT | NOD2 | 64127 |
| CCCGGCAGCAGGAGCGTCGC | TRMT61A | 115708 |
| CCCGGCAGGTCCCGGCCCGG | IRAK1 | 3654 |
| CCCGGCCCCCACAACTCCGC | DNTTIP1 | 116092 |
| CCCGGCGCTACGAGCAGCCA | TMEM173 | 340061 |
| CCCGGCGCTCGCTGCTCCCG | ALOX5 | 240 |
| CCCGGCGGCAGGAGCGCAGG | JAM2 | 58494 |

| | | |
|---|---|---|
| CCCGGCGGCTCTTCCAGCAG | ADRB1 | 153 |
| CCCGGCTCTCCGCGCGGCCG | NDUFB9 | 4715 |
| CCCGGCTGAGCCGACATTGC | ATP5E | 514 |
| CCCGGGCTGCAGGGACCACG | APEH | 327 |
| CCCGGGGCCGCCGCGTCGCT | RAD23A | 5886 |
| CCCGGTAACGGACGAGTAGA | EDA2R | 60401 |
| CCCGTATCCTGCGCCCGCCG | PIP4K2A | 5305 |
| CCCGTCGTGTTTCTGCTTCT | PRKAA1 | 5562 |
| CCCGTGAGTAACTTGGCTCC | ANP32B | 10541 |
| CCCGTGGCGTGCGCACCTGT | NO_CURRENT_383 | NO_CURRENT_383 |
| CCCTCACGGTCCTTAAGTCT | SPCS1 | 28972 |
| CCCTCCAAATGGGCTGCTGG | KPNB1 | 3837 |
| CCCTCCTCCTACCAGCGCGC | NR2C2 | 7182 |
| CCCTCCTTGACACCTCCCCG | CYP24A1 | 1591 |
| CCCTCGACATGGCGCTGAGG | JAM3 | 83700 |
| CCCTCGACTCCGCGCGTCCC | HTR7 | 3363 |
| CCCTGAAAACATTAAAATGC | IKBKB | 3551 |
| CCCTGCGTGGAGGAACTGCT | ACACA | 31 |
| CCCTGCTGGTAGAAAAACTA | NO_CURRENT_384 | NO_CURRENT_384 |
| CCCTGGGATCACCCCGACCG | NO_CURRENT_385 | NO_CURRENT_385 |
| CCCTTGCTGGGAGAAGAGGA | NR1H3 | 10062 |
| CCCTTGGCTGCTCGTAGCGC | TMEM173 | 340061 |
| CCCTTGTCAGCGTGACCCCG | NO_CURRENT_386 | NO_CURRENT_386 |
| CCGAACAATGAAGCGGCGCG | MBNL1 | 4154 |
| CCGAATCTCCTCACAGCTCC | DNAJB6 | 10049 |
| CCGACGAGTCTCAACTAAAA | CFLAR | 8837 |
| CCGACTCCGGACGCCGGGAC | NUDT17 | 200035 |
| CCGACTGTGGAGAAGTGTCC | OXSM | 54995 |
| CCGAGAAGCAGAAACACGAC | PRKAA1 | 5562 |
| CCGAGACTTAAGGACCGTGA | SPCS1 | 28972 |
| CCGAGAGCCTAGTTCCGGCC | NQO1 | 1728 |
| CCGAGCGACGCGGCGGCCCC | RAD23A | 5886 |
| CCGAGGACTCCAGCACACCG | ITGB2 | 3689 |
| CCGAGTCACCAGTAGGCTGT | ACO2 | 50 |
| CCGAGTGAGGCGACGGGGTA | ATP6V0E1 | 8992 |
| CCGAGTTCGCCAAGGCGCCG | SLC5A8 | 160728 |
| CCGCACCAACGCGCCCGCCA | MPC1 | 51660 |
| CCGCACGCTGCAGCCGCGGC | PPARGC1B | 133522 |
| CCGCAGCCGCCGCCGCGGCG | NENF | 29937 |
| CCGCAGCTTTGAAGCCTGAG | SLC7A3 | 84889 |
| CCGCATCCGCGTGTCCACTG | ALOX15 | 246 |
| CCGCATTGTTTGCTTCGCTG | BRD1 | 23774 |
| CCGCCATGGAGCTGAGAGTC | CSNK1D | 1453 |
| CCGCCCCCAGCGGGATGTGA | MGA | 23269 |
| CCGCCCGCCATGGCTGCAGG | MYD88 | 4615 |
| CCGCCCGCGCACCCGCCCGG | TP73 | 7161 |
| CCGCCCGGCGAGGAGGAGGG | CDKN2D | 1032 |
| CCGCCGCCGCCGCCGCCGGC | EIF2AK2 | 5610 |
| CCGCCGCCGCCGCCGCCGGC | HCN2 | 610 |
| CCGCCTCGCCCTCCGCGGAC | RAC1 | 5879 |

| | | |
|---|---|---|
| CCGCCTCTTCCGCTCTACAG | TMEM30A | 55754 |
| CCGCCTGCGCGCCGACTCCA | NOD2 | 64127 |
| CCGCCTGGCTGAGAAAACTG | MCL1 | 4170 |
| CCGCCTGGTCTGCAGTTTGT | BCL2L11 | 10018 |
| CCGCGACCCCCTCCATGGTC | PCGF6 | 84108 |
| CCGCGACGCCCGTGCCGCAC | ACTR5 | 79913 |
| CCGCGCATTTCAGAGCACAA | NO_CURRENT_387 | NO_CURRENT_387 |
| CCGCGCCATGCCCTCCTACA | ALOX5 | 240 |
| CCGCGCGGGTGGCTGAGCAG | SNRNP70 | 6625 |
| CCGCGCTGGGAAAAAGGTGG | INO80C | 125476 |
| CCGCGGAAGCAGTGCAGACG | CASP3 | 836 |
| CCGCGGAGGGCGAGGCGGCG | RAC1 | 5879 |
| CCGCGGCCATGGGCGGGCGC | ABL1 | 25 |
| CCGCGGCGCGGGGCCCACCA | NENF | 29937 |
| CCGCGGCGGGCGCAGGATAC | PIP4K2A | 5305 |
| CCGCGGCTGCTGCCAGTGTG | BCORL1 | 63035 |
| CCGCGGGAGCCCGCCGGCGG | TBK1 | 29110 |
| CCGCGGGCCAAGCCTACCGG | ZNF641 | 121274 |
| CCGCGGGGCCCGGAGCCTCC | TRIR | 79002 |
| CCGCGGGTGCAGGAGTCCTG | DET1 | 55070 |
| CCGCGTCCTCCTTGCCCAGA | SQSTM1 | 8878 |
| CCGCTATTGAAACCGCCCAC | NO_CURRENT_388 | NO_CURRENT_388 |
| CCGCTCAGGGTGAGTCCCGC | YPEL5 | 51646 |
| CCGCTGCCAGCGAAGGTGCC | BCL2 | 596 |
| CCGCTGCTGGAAGAGCCGCC | ADRB1 | 153 |
| CCGCTGGAGATCTGCTGCCC | LTB4R | 1241 |
| CCGCTGGGGAAGGCTCCGGG | SATB1 | 6304 |
| CCGCTGGTGGGAGGCGCGGC | PRKN | 5071 |
| CCGCTTGCCCTCCCGCCCCA | CHUK | 1147 |
| CCGGAACAGCGCGCTCGCAG | ACADS | 35 |
| CCGGAACGTGGGGATTGGTC | MLF2 | 8079 |
| CCGGAAGGGTCGCAGCGCCC | MAPK9 | 5601 |
| CCGGACCAATCCCCACGTTC | MLF2 | 8079 |
| CCGGACCGCCAGCTCAGAAC | AHR | 196 |
| CCGGAGAGCGCCATGGTCCG | NOD1 | 10392 |
| CCGGAGCGCGGCGACTCCAG | RIPK1 | 8737 |
| CCGGAGTCCGAACAATGAAG | MBNL1 | 4154 |
| CCGGCAATGTCGGCTCAGCC | ATP5E | 514 |
| CCGGCAGATCCCCTCACACG | MCTS1 | 28985 |
| CCGGCCCATGCTGGAGGCGG | VIPR2 | 7434 |
| CCGGCCGGGCGCGGAGCCCG | PXMP2 | 5827 |
| CCGGCCGGGCGCGGAGCCCG | SBNO2 | 22904 |
| CCGGCCTCCAGCACTGGGTC | ACTR5 | 79913 |
| CCGGCCTCTGTCGGCTCCGT | PRR14L | 253143 |
| CCGGCCTGGGCTGGGGTTTG | PDK4 | 5166 |
| CCGGCGACTACGTGGGGTAC | BABAM2 | 9577 |
| CCGGCGCTACGAGCAGCCAA | TMEM173 | 340061 |
| CCGGCGGCAGCTGGTGCGGG | UCHL5 | 51377 |
| CCGGCGGGATTCCTACCCGC | IFNAR2 | 3455 |
| CCGGCGGGTAGGAATCCCGC | IFNAR2 | 3455 |

| | | |
|---|---|---|
| CCGGCGTGACAGCGATTCGG | HNRNPF | 3185 |
| CCGGCTCATCCGCCGCGTCC | SATB1 | 6304 |
| CCGGCTCCGTGAGGCCCTGC | NCOA6 | 23054 |
| CCGGGAAAGGGGACGCAGCA | NDUFC1 | 4717 |
| CCGGGAACCAATTCTCCTGT | NIPBL | 25836 |
| CCGGGACGCCGCCGGGAGGA | SIAH2 | 6478 |
| CCGGGAGGGCAGGAGCGAAT | CSNK1D | 1453 |
| CCGGGCACCGCGGCCATGGA | UCHL3 | 7347 |
| CCGGGCCCCGAGGCACAGCC | NMU | 10874 |
| CCGGGCGCTTCAGGCTGGCT | KIF2B | 84643 |
| CCGGGGAGTCCGCTCCAGCG | SPHK1 | 8877 |
| CCGGGGATCCTGGAGCCATG | PYCARD | 29108 |
| CCGGGGCAGCAGATCTCCAG | LTB4R | 1241 |
| CCGGGGCCGCCGCGTCGCTC | RAD23A | 5886 |
| CCGGGGCGCTGCGACCCTTC | MAPK9 | 5601 |
| CCGGGTCCCTTTGTCGCCCA | ARRB1 | 408 |
| CCGGGTGCCTCGAGGGCGCG | SLC6A4 | 6532 |
| CCGGTGCGGCACGGGCGTCG | ACTR5 | 79913 |
| CCGTAGGAGGAAGATGGCGG | SAP18 | 10284 |
| CCGTCCTGCCGGACTGCTGG | EIF4G3 | 8672 |
| CCGTCTCCGCATCGTCTTTT | NO_CURRENT_389 | NO_CURRENT_389 |
| CCGTGAGGCCCTGCCGGGTC | NCOA6 | 23054 |
| CCGTGAGTAACTTGGCTCCG | ANP32B | 10541 |
| CCGTGCCTGACCCCTCAATG | NO_CURRENT_390 | NO_CURRENT_390 |
| CCGTGCGTTTAACCCAGGCG | ARMCX2 | 9823 |
| CCGTGCTCTGCAGGTAGAAA | TBXAS1 | 6916 |
| CCGTGGAGGCTTCGCCGCCT | RBM6 | 10180 |
| CCGTGTAGGAGGGCATGGCG | ALOX5 | 240 |
| CCGTGTGTGTGCGTGCGAGC | BMI1 | 648 |
| CCGTGTTCCACAACTACCCT | NO_CURRENT_391 | NO_CURRENT_391 |
| CCGTTGCAGTCCTCGGAACC | SOD1 | 6647 |
| CCGTTGCCACTGACAGCCGC | RRAGA | 10670 |
| CCGTTGGACTATGGCGGGTC | NO_CURRENT_392 | NO_CURRENT_392 |
| CCTAAACTCAGACGCACTAC | NO_CURRENT_393 | NO_CURRENT_393 |
| CCTAAGGGGTACCACCATGG | NO_CURRENT_394 | NO_CURRENT_394 |
| CCTACAGCCTACTGGTGACT | ACO2 | 50 |
| CCTACCATAGCATCACGATA | NO_CURRENT_395 | NO_CURRENT_395 |
| CCTACGCGGTAGGGAACTTT | NO_CURRENT_396 | NO_CURRENT_396 |
| CCTACTCCCGTGTGTTATCC | NO_CURRENT_397 | NO_CURRENT_397 |
| CCTACTGGTGACTCGGCTGC | ACO2 | 50 |
| CCTAGAATAAGTGAGAAAGT | DPY30 | 84661 |
| CCTAGAATAAGTGAGAAAGT | MEMO1 | 51072 |
| CCTAGCAGAGGGGGAGAGGA | CYP4F11 | 57834 |
| CCTAGGCGGCGAAGCCTCCA | RBM6 | 10180 |
| CCTATACCATCACCCACACC | HTR1D | 3352 |
| CCTCACTCTTTCGCTCGCCG | BCORL1 | 63035 |
| CCTCAGGAACTTTGGGCGCC | PAXIP1 | 22976 |
| CCTCAGGGCCGGGAGGCTCC | TRIR | 79002 |
| CCTCCAAATAGCATCACATG | NO_CURRENT_398 | NO_CURRENT_398 |
| CCTCCCCGGTGCTGGGACCG | IL2RB | 3560 |

| | | |
|---|---|---|
| CCTCCCTCCAAATGGGCTGC | KPNB1 | 3837 |
| CCTCCGCCCTCACTTACCCT | EP400 | 57634 |
| CCTCCGCGTCGGACCCCGCG | SIAH2 | 6478 |
| CCTCCGTGCTAACGCGGACG | NO_CURRENT_399 | NO_CURRENT_399 |
| CCTCCTCCTCAGTTGGAGGG | RND1 | 27289 |
| CCTCCTCGGCGTCGTCGTCG | USP7 | 7874 |
| CCTCCTCTTCCACCCCTGCC | LCN2 | 3934 |
| CCTCCTGCAGCCATGGCGGG | MYD88 | 4615 |
| CCTCCTGCGCTCCTGCCGCC | JAM2 | 58494 |
| CCTCGAGGGCTGAGGCTCTT | HSH2D | 84941 |
| CCTCGATGGTCACCTGTAGC | NO_CURRENT_400 | NO_CURRENT_400 |
| CCTCGCCTGGGTTAAACGCA | ARMCX2 | 9823 |
| CCTCGGCCATAACGCGACTC | NUDT17 | 200035 |
| CCTCGGCGGTGCTCCTGGCT | ITGB2 | 3689 |
| CCTCGGGCGTAAATACTCAT | NO_CURRENT_401 | NO_CURRENT_401 |
| CCTCGGTGTGCTGGAGTCCT | ITGB2 | 3689 |
| CCTCGTGTGAGGGGATCTGC | MCTS1 | 28985 |
| CCTCTCTGGAGGTCGAGGCG | NAIP | 4671 |
| CCTCTGCCACCATGGGGAAC | CYBB | 1536 |
| CCTCTGGAAGGACACTTCTG | NO_CURRENT_402 | NO_CURRENT_402 |
| CCTCTGGCAGCCCCTGGTCC | MMP9 | 4318 |
| CCTGACTGACTGCATTGGCA | CHMP5 | 51510 |
| CCTGAGCCCCGCTACCGAGA | EIF4G3 | 8672 |
| CCTGAGGATACCAAAACGTG | NO_CURRENT_403 | NO_CURRENT_403 |
| CCTGATAAATGTGACACCAT | NO_CURRENT_404 | NO_CURRENT_404 |
| CCTGCACCTTCGCCCCTACA | MAPK14 | 1432 |
| CCTGCAGCCGAGTCACCAGT | ACO2 | 50 |
| CCTGCGACGCTCCTGCTGCC | TRMT61A | 115708 |
| CCTGCGCCGGACTGGCTTGC | FAM107B | 83641 |
| CCTGCGGTGCACGGCTAGCC | NO_CURRENT_405 | NO_CURRENT_405 |
| CCTGCGTCCGCGGCGGCTTG | CYFIP1 | 23191 |
| CCTGCTACCCCCCATATGTG | NO_CURRENT_406 | NO_CURRENT_406 |
| CCTGCTCCAGTCGCTATCGG | SNRNP70 | 6625 |
| CCTGGAGCTGCAGACAGTGC | JAK1 | 3716 |
| CCTGGCCGGAACTAGGCTCT | NQO1 | 1728 |
| CCTGGGACACCCGGGAGCCG | SLC7A5 | 8140 |
| CCTGGGACGCGCGGAGTCGA | HTR7 | 3363 |
| CCTGGGCGCTAAGATGGCGG | RAD23A | 5886 |
| CCTGGTACTACTGCTTGCTG | GC | 2638 |
| CCTGGTGTCTCTCCCGACCA | PCGF6 | 84108 |
| CCTGGTTAGGAGCAAAGGAA | CDKN2C | 1031 |
| CCTGGTTCCGAGGACTGCAA | SOD1 | 6647 |
| CCTGTACCCCACGTAGTCGC | BABAM2 | 9577 |
| CCTGTCCCGGCGTCCGGAGT | NUDT17 | 200035 |
| CCTGTGAATGATGCAATGGA | NLRP12 | 91662 |
| CCTGTGCTCCCTATATGAGA | HTR3E | 285242 |
| CCTGTTCTGAGCTGGCGGTC | AHR | 196 |
| CCTTAGCCTCAGAAAGCACT | NO_CURRENT_407 | NO_CURRENT_407 |
| CCTTAGCCTCTAGAGTACCC | NO_CURRENT_408 | NO_CURRENT_408 |
| CCTTATCTCACGGAATATAG | NO_CURRENT_409 | NO_CURRENT_409 |

| | | |
|---|---|---|
| CCTTCACATCCCGCTGGGGG | MGA | 23269 |
| CCTTCAGTTCACTCTCAGTA | PPARGC1A | 10891 |
| CCTTCCATTGCATCATTCAC | NLRP12 | 91662 |
| CCTTCCCTAACGTTGCAACT | MAP2K6 | 5608 |
| CCTTCTCGGTAGCGGGGCTC | EIF4G3 | 8672 |
| CCTTCTTGTACCAATAAATG | NO_CURRENT_410 | NO_CURRENT_410 |
| CCTTGAAATCAAATCAAACC | NO_CURRENT_411 | NO_CURRENT_411 |
| CCTTGACTTTATAACTATCG | NO_CURRENT_412 | NO_CURRENT_412 |
| CCTTGCTGCGTCCCCTTTCC | NDUFC1 | 4717 |
| CCTTGGAGTCGGCGCGCAGG | NOD2 | 64127 |
| CCTTGGCAGGATCAAGACCT | COX16 | 51241 |
| CCTTGGCTGCTCGTAGCGCC | TMEM173 | 340061 |
| CCTTGTCATGGGAGCATTGT | MBNL1 | 4154 |
| CCTTTAGACAACTTCAGGAT | NAIP | 4671 |
| CCTTTCAAGGACCCAGAAGT | NDUFA1 | 4694 |
| CCTTTCCTTCCACTGCAGGG | EBI3 | 10148 |
| CCTTTCCTTTGCTCCTAACC | CDKN2C | 1031 |
| CCTTTCGCAGAGAGGGGAAG | RIPK3 | 11035 |
| CCTTTCTCTGCTCTCCAAGC | AHDC1 | 27245 |
| CCTTTGGGGTTCAAGATCAC | TNFRSF1A | 7132 |
| CCTTTGTGTCGCCATGGCGG | ARHGAP33 | 115703 |
| CGAAACCCGGAAGTGAGCGG | SBNO2 | 22904 |
| CGAAACCCTCTTAAGTTAAC | NO_CURRENT_413 | NO_CURRENT_413 |
| CGAACGAGCGGCGCTCGGCG | FOSL2 | 2355 |
| CGAACTTAATCCCGTGGCAA | NO_CURRENT_414 | NO_CURRENT_414 |
| CGAAGAGCCGCGTGAGGAAA | LTB4R | 1241 |
| CGAAGGGGACGCAGCGAAAC | TP73 | 7161 |
| CGAATATTATTTCTATCGGG | NO_CURRENT_415 | NO_CURRENT_415 |
| CGAATCGGAACTTTGTACCG | NO_CURRENT_416 | NO_CURRENT_416 |
| CGACAACGTGCAGGTGTATC | NO_CURRENT_417 | NO_CURRENT_417 |
| CGACACAAATTACTAACAAC | NO_CURRENT_418 | NO_CURRENT_418 |
| CGACCCGGAGGATGAGATGT | NO_CURRENT_419 | NO_CURRENT_419 |
| CGACCCTCCGCCTGTTATTG | FBXO38 | 81545 |
| CGACGACGACGACGGCGGCG | NUP153 | 9972 |
| CGACGACGACGCCGAGGAGG | USP7 | 7874 |
| CGACGACGCCGAGGAGGCGG | USP7 | 7874 |
| CGACGAGAACGGCGAGCGAG | ARHGAP33 | 115703 |
| CGACGCCAAGAACGCCATTA | NDUFB9 | 4715 |
| CGACGCGCAAGAGGCGACCG | SATB2 | 23314 |
| CGACGGACGCCTGGGACGCG | HTR7 | 3363 |
| CGACGGCGGCGGGGGAGTCT | MSL2 | 55167 |
| CGACGGTAATGCACCTACTA | NO_CURRENT_420 | NO_CURRENT_420 |
| CGACTAACCGGAAACTTTTT | NO_CURRENT_421 | NO_CURRENT_421 |
| CGACTACAGAGAAGGGTAAT | SMPD1 | 6609 |
| CGACTGTGGAGAAGTGTCCG | OXSM | 54995 |
| CGAGAAGCGGGAGAGCTGGC | CXCL3 | 2921 |
| CGAGACTCCAGTGATCATAG | BAK1 | 578 |
| CGAGACTTAAGGACCGTGAG | SPCS1 | 28972 |
| CGAGAGAGGAAGCTCTTTCG | RIPK2 | 8767 |
| CGAGAGCCTAGTTCCGGCCA | NQO1 | 1728 |

| | | |
|---|---|---|
| CGAGAGGGCGGCGAGAAGGA | SLC25A19 | 60386 |
| CGAGCAAAGATTGTTGGATA | NO_CURRENT_422 | NO_CURRENT_422 |
| CGAGCACGACCGATTCTGCT | ANKRD50 | 57182 |
| CGAGCAGGCAGCGTTGCAAG | AHCTF1 | 25909 |
| CGAGCCGCCGCGAGTTCTCA | LGMN | 5641 |
| CGAGCCGGGGCAGGAAGAGG | NFKB1 | 4790 |
| CGAGCGCAGAGCCGGAGTCG | JAM3 | 83700 |
| CGAGCTAACCGAGTGCGGCG | GPT2 | 84706 |
| CGAGCTTTCGGCACCTCTGC | HDAC2 | 3066 |
| CGAGGAAGCAGGCGCTGTGG | LCN2 | 3934 |
| CGAGGAAGCCGGTGTTGCCG | MOSPD1 | 56180 |
| CGAGGACTCCAGCACACCGA | ITGB2 | 3689 |
| CGAGGAGCGCGGTTACCGGA | CCNC | 892 |
| CGAGGAGGATGGCGGAGTCG | SHOC2 | 8036 |
| CGAGGCACAGCCAGGGCACC | NMU | 10874 |
| CGAGGCCCGGGCGAGCAGCG | ACADS | 35 |
| CGAGGGCCTCGGCCATCCAA | HSH2D | 84941 |
| CGAGGGGAGCCAAGAGAAAG | UBE2L6 | 9246 |
| CGAGGGTTCTGGCCGGTAAG | GLMN | 11146 |
| CGAGGTCATGAATCATGTGA | LAMTOR3 | 8649 |
| CGAGTCACCAGTAGGCTGTA | ACO2 | 50 |
| CGAGTCCCCACCCCTAGCTC | CFLAR | 8837 |
| CGAGTCGAAAACTGAGGCTC | NOS2 | 4843 |
| CGAGTCTGGTAGCTGAGCGT | LTA4H | 4048 |
| CGAGTGAGGCGACGGGGTAG | ATP6V0E1 | 8992 |
| CGAGTGAGTGTGGTCGCTCC | FLCN | 201163 |
| CGAGTGAGTGTGGTCGCTCC | PLD6 | 201164 |
| CGAGTGGGAAACGGGAATCA | NO_CURRENT_423 | NO_CURRENT_423 |
| CGAGTGTTATACGCACCGTT | NO_CURRENT_424 | NO_CURRENT_424 |
| CGATCTTAGGTGGGTTCCCG | HSPA13 | 6782 |
| CGATGCCCGTCTATGGCCCG | NO_CURRENT_425 | NO_CURRENT_425 |
| CGCAAGACGCCCGCACCTGA | CYP2R1 | 120227 |
| CGCAAGGTGTCGGTAACCCT | NO_CURRENT_426 | NO_CURRENT_426 |
| CGCAATGCCTAAAGGAGGTG | PDAP1 | 11333 |
| CGCACATCTAAAGTTACTAC | NO_CURRENT_427 | NO_CURRENT_427 |
| CGCACCGGACCCAGTGCTGG | ACTR5 | 79913 |
| CGCACGACCATTGCTGCTGC | NO_CURRENT_428 | NO_CURRENT_428 |
| CGCACTCACGGCTCATGGCG | PTGS1 | 5742 |
| CGCAGCAGCAGCCGCCGCGG | UBE2E1 | 7324 |
| CGCAGCCCGCAGAGGCGCTG | CACNA2D2 | 9254 |
| CGCAGCGGAGCCTGGAGAGA | TNFRSF1B | 7133 |
| CGCAGGCCAAGCCCCAGCTG | TRAF6 | 7189 |
| CGCAGGCCGTAGGAGGAAGA | SAP18 | 10284 |
| CGCAGGCGGACCGGGGGCAA | TNFRSF6B | 8771 |
| CGCAGGCTAGATGACACCAG | NO_CURRENT_429 | NO_CURRENT_429 |
| CGCAGGCTCAGCGGCCCCGG | SCN5A | 6331 |
| CGCAGGTCGGTGCGTCTGTC | BABAM2 | 9577 |
| CGCAGTACGTATAGACTTAA | NO_CURRENT_430 | NO_CURRENT_430 |
| CGCATGCTCGCCATCATGTC | UBE2L6 | 9246 |
| CGCCAACGCGCAGGTGCTAG | PLD1 | 5337 |

| | | |
|---|---|---|
| CGCCAAGACGCCGGCAATGT | ATP5E | 514 |
| CGCCAGAAACAGCATCACGC | TMEM70 | 54968 |
| CGCCAGCTCGCCGCTCGCTA | SQSTM1 | 8878 |
| CGCCATCTTTGGGATCCTAC | GPR119 | 139760 |
| CGCCATGGCCTCCCCCGCAG | KAT2A | 2648 |
| CGCCCATGACGGAGAGTCCG | NDUFA1 | 4694 |
| CGCCCCAGCTGTGGGTCCCC | RIPK1 | 8737 |
| CGCCCCGAGGGTGGCGCGTC | GSDMD | 79792 |
| CGCCCCTACAAGGCCGACTC | MAPK14 | 1432 |
| CGCCCGCAGCCCGACCCGGC | NCOA6 | 23054 |
| CGCCCGCGCCCTGACAGGCC | RRAGC | 64121 |
| CGCCCGGAGCCCGAGCCGCG | RPS6KA4 | 8986 |
| CGCCCGGCGAGGAGGAGGGA | CDKN2D | 1032 |
| CGCCGAGCTGTGCGCAGCCG | DNAJA1 | 3301 |
| CGCCGCCCGCGGTCCTCCGT | RHOA | 387 |
| CGCCGCCGCCGCCCGCGCGC | NR2C2 | 7182 |
| CGCCGCCGTTGCTTGAATGG | PHIP | 55023 |
| CGCCGCCTGGCTGAGAAAAC | MCL1 | 4170 |
| CGCCGCGGCGGCTGCTGCTG | BBC3 | 27113 |
| CGCCGCGGCGGCTGCTGCTG | UBE2E1 | 7324 |
| CGCCGCGTCTTCCTCAAGGT | HNRNPF | 3185 |
| CGCCGGACTGGCTTGCGGGC | FAM107B | 83641 |
| CGCCGGGACCGTTAGGGAAT | NO_CURRENT_431 | NO_CURRENT_431 |
| CGCCGGGCCGGGACCTGCCG | IRAK1 | 3654 |
| CGCCGGGGTGCGCCTGCCTC | PDE3A | 5139 |
| CGCCTCCTCGGCGTCGTCGT | USP7 | 7874 |
| CGCCTCTCACGTGTAGGCTT | NO_CURRENT_432 | NO_CURRENT_432 |
| CGCCTCTTCTTCACACTGCT | HMGN2 | 3151 |
| CGCCTGCTCATCTCCCCGTA | ZRSR2 | 8233 |
| CGCCTGGAGTACCCTTCCCG | MPC2 | 25874 |
| CGCCTTGGCCCCTCAGCCCG | TBK1 | 29110 |
| CGCGACTCCGGCTCTGCGCT | JAM3 | 83700 |
| CGCGAGAGGAAGCGATGCAG | ABI1 | 10006 |
| CGCGATCTACTCGGCCCCGC | CREBBP | 1387 |
| CGCGCACAACCTCGAAGAGC | KPNA1 | 3836 |
| CGCGCACCACGGGCGCGCAC | NO_CURRENT_433 | NO_CURRENT_433 |
| CGCGCAGCCCTCATCGCAAC | TIRAP | 114609 |
| CGCGCCTGCAGCTGGGAGCT | KIAA1211L | 343990 |
| CGCGCCTGGCCGGCGGGCTG | ABCB10 | 23456 |
| CGCGCGCGCCCCATGGCTCC | PYCARD | 29108 |
| CGCGCGGAGTCGAGGGAGCT | HTR7 | 3363 |
| CGCGCGGCCTGCCTTCCCAC | TIFA | 92610 |
| CGCGCGGCTCGGGCTCCGGG | KLHL29 | 114818 |
| CGCGCGGCTCGGGCTCCGGG | RPS6KA4 | 8986 |
| CGCGCGGGCTTTGGTCGGTC | HMGN2 | 3151 |
| CGCGCGTCGAGCGGGAGCAG | SIRT1 | 23411 |
| CGCGCTCTGAGCTGCCCCCA | KPNB1 | 3837 |
| CGCGCTGGGAAAAAGGTGGG | INO80C | 125476 |
| CGCGGAAGGAGCGCGGCCGG | TYK2 | 7297 |
| CGCGGACCATGGGCGACAAA | ARRB1 | 408 |

| | | |
|---|---|---|
| CGCGGAGCAGCCAGACAGCG | TGFB1 | 7040 |
| CGCGGAGGCTGAAGCTGAGG | KHSRP | 8570 |
| CGCGGATCCGCGGCGGGCGC | PIP4K2A | 5305 |
| CGCGGATCTCACCGCCGCTC | YPEL5 | 51646 |
| CGCGGCACAGCCCCGTGGGT | ARID3A | 1820 |
| CGCGGCCACCTGGTCCGAGG | SCTR | 6344 |
| CGCGGCCCGAAGACGGGAGC | AKT1 | 207 |
| CGCGGCCGCCGGCTCAGTCT | FOS | 2353 |
| CGCGGCCGGAGAAGGGCTGC | NFRKB | 4798 |
| CGCGGCCTCCAATCTCCGCA | APIP | 51074 |
| CGCGGCCTCTGGTGCAGCGG | CS | 1431 |
| CGCGGCGCCGCTCTCCGCTG | KAT2A | 2648 |
| CGCGGCGCTCATGGCGGCGT | CEBPD | 1052 |
| CGCGGCGGCTTGGGGGTCCT | CYFIP1 | 23191 |
| CGCGGCTGAGGGCTTCTCGT | NDUFA8 | 4702 |
| CGCGGGACGCGGGGCTGGCT | BRD1 | 23774 |
| CGCGGGCCCAGTTGCGATGA | TIRAP | 114609 |
| CGCGGGCCTCTGGGTTCCGA | SPINT1 | 6692 |
| CGCGGGCTGCAGCAGATCAC | NDUFA12 | 55967 |
| CGCGGGCTTTGGTCGGTCCG | HMGN2 | 3151 |
| CGCGGGTCTCCGGCCGGCGG | EIF2AK2 | 5610 |
| CGCGGGTGGTGCGGGCGCTG | PTGES2 | 80142 |
| CGCGGTCGCCTCTTGCGCGT | SATB2 | 23314 |
| CGCGGTGCCAACCATGGAGC | PDE5A | 8654 |
| CGCGTAGCTGGTGCTCCACC | NMU | 10874 |
| CGCGTCCGCAGGAAGAGGCG | PARK7 | 11315 |
| CGCGTGCGCGAACATGTAAC | IFNAR1 | 3454 |
| CGCGTGTAGCTGGAGACAAG | NO_CURRENT_434 | NO_CURRENT_434 |
| CGCGTTTCTGCGGGCGCAAG | RAC2 | 5880 |
| CGCTAAATTGTCACACGTTT | NO_CURRENT_435 | NO_CURRENT_435 |
| CGCTAGCACCTGCGCGTTGG | PLD1 | 5337 |
| CGCTAGGTCCGGTAAGTGCG | NO_CURRENT_436 | NO_CURRENT_436 |
| CGCTAGGTTATTTCGTGGCC | NO_CURRENT_437 | NO_CURRENT_437 |
| CGCTAGTACGCTCCTCTATA | NO_CURRENT_438 | NO_CURRENT_438 |
| CGCTCACACCGTGCGGGGGG | EGFR | 1956 |
| CGCTCATGGCGGCGTCGGGC | CEBPD | 1052 |
| CGCTCCAGGGCCCTTGGGGT | KIF2B | 84643 |
| CGCTCCCTGTGTAGATTCGC | SLC5A8 | 160728 |
| CGCTCGAGCGGTTCCTGTCA | TMEM30A | 55754 |
| CGCTCGCTCGCTGGTTCGCT | FOSL2 | 2355 |
| CGCTCTGAAGGTGACCCCCC | SPINT1 | 6692 |
| CGCTGACAGACGCAAGATGG | PIAS1 | 8554 |
| CGCTGCAGGGGTCGGAGGTC | SAP18 | 10284 |
| CGCTGCCAGCGAAGGTGCCG | BCL2 | 596 |
| CGCTGCCGTGCGTTTAACCC | ARMCX2 | 9823 |
| CGCTGCCTCTACCCCCGCCA | MAP3K7 | 6885 |
| CGCTGCGCGCTCACCATGGT | NENF | 29937 |
| CGCTGCGGCCCGTGCAGCCC | CACNA2D2 | 9254 |
| CGCTGCGGGGGAGGCCATGG | KAT2A | 2648 |
| CGCTGCTCGGTGCGAGGCGG | KPNA1 | 3836 |

| | | |
|---|---|---|
| CGCTGCTGCCTACCTCCCGA | PDHB | 5162 |
| CGCTGCTGCTGCTGCTGCCC | SETD1B | 23067 |
| CGCTGCTGCTGCTGCTGCCC | TM9SF3 | 56889 |
| CGCTGGAGATCTGCTGCCCC | LTB4R | 1241 |
| CGCTGGAGATGATGGGGCGG | STAG2 | 10735 |
| CGCTGGCTCCCGCCGCGGAA | RB1 | 5925 |
| CGCTGGGAGAAGCCCAGTGG | PHF8 | 23133 |
| CGCTGGGCTGCGAGGGCGCG | TNFRSF1B | 7133 |
| CGCTGGGGCCCGCGCGGCTC | RPS6KA4 | 8986 |
| CGCTGGTTCTCCAGATGCGG | CXCR4 | 7852 |
| CGCTTATAGTGGAGGCTGCT | UBE2H | 7328 |
| CGCTTCCGCGGCCCGTTCAA | NO_CURRENT_439 | NO_CURRENT_439 |
| CGCTTGGAGTTCTGAGCCGA | COX16 | 51241 |
| CGGAACGTGGGGATTGGTCC | MLF2 | 8079 |
| CGGAACTACAGGTGTGGGGC | FAM214B | 80256 |
| CGGAATATCAGAGTACCTAG | PPID | 5481 |
| CGGACCCAGAACGCTCTCCC | SLC3A2 | 6520 |
| CGGACCGCCAGCTCAGAACA | AHR | 196 |
| CGGACGCGAGTCGTCGTGGC | MPC2 | 25874 |
| CGGACTGACGGGCGGCCGGG | MAP2K7 | 5609 |
| CGGACTGGCGGCGGCTGCGG | MAP2K5 | 5607 |
| CGGAGAAGCAGCGGCTGGCG | RIPK2 | 8767 |
| CGGAGACACAACAAAGATGG | IKZF5 | 64376 |
| CGGAGACCCCTTCGGGAGGT | PDHB | 5162 |
| CGGAGAGTGAGGGTAGGTGC | RTP1 | 132112 |
| CGGAGCGGATCCGAGGACAG | WDR26 | 80232 |
| CGGAGGAGCGGTAACTACCC | DNAJA1 | 3301 |
| CGGAGGAGGTGGCGGCGGCT | GLMN | 11146 |
| CGGAGGCACGTGTGGAGCCC | TRMT61A | 115708 |
| CGGAGGGGCTCGGCTGCACC | STAT1 | 6772 |
| CGGAGTCGGTGAGCTTCGCA | NUDT17 | 200035 |
| CGGATCCTCCTGTCCCGCCA | BRK1 | 55845 |
| CGGATCGGCGGCTCCTGCGG | CASP9 | 842 |
| CGGATCTCACCCGCCACACC | NFKB2 | 4791 |
| CGGATGAGCCGGCCCCGCTG | SATB1 | 6304 |
| CGGCACACCAATGCGTTCGT | NO_CURRENT_440 | NO_CURRENT_440 |
| CGGCACAGCCCCGTGGGTCG | ARID3A | 1820 |
| CGGCACGGGCGTCGCGGAAC | ACTR5 | 79913 |
| CGGCAGAGGTGCCGAAAGCT | HDAC2 | 3066 |
| CGGCAGCAAGCGTGGGAACG | IRF8 | 3394 |
| CGGCAGCGGCTTCAGCAGAT | SOD2 | 6648 |
| CGGCAGCTGTGAGGGGGTTC | PITPNB | 23760 |
| CGGCAGGTAGGCACAGTGGG | IRF8 | 3394 |
| CGGCAGTGGCGGCGACGGCG | RNF111 | 54778 |
| CGGCATTGCGGCCAGCCAGC | MEX3B | 84206 |
| CGGCATTTAACAAGGCGGTG | IKZF5 | 64376 |
| CGGCCAAGGGGGCTTGGAAC | CBLL1 | 79872 |
| CGGCCATCCAAGGGTCTCCC | HSH2D | 84941 |
| CGGCCATGGAGGGTCAACGC | UCHL3 | 7347 |
| CGGCCCCAGGTAAGGCTGTT | ATP5J2 | 9551 |

| | | |
|---|---|---|
| CGGCCCCAGGTAAGGCTGTT | ATP5J2-PTCD1 | 100526740 |
| CGGCCCCAGGTAAGGCTGTT | PTCD1 | 26024 |
| CGGCCCCTCCGCAGGCGGAC | TNFRSF6B | 8771 |
| CGGCCCGGACCCATGCAGCT | RAD21 | 5885 |
| CGGCCCGTTCATGACCGGCC | INO80E | 283899 |
| CGGCCCTGAGAGGCCCCGGC | IRAK1 | 3654 |
| CGGCCGCCTGCTCCCGTCTT | AKT1 | 207 |
| CGGCCGGAGCTGTTTGTGCT | DPY30 | 84661 |
| CGGCCGGAGCTGTTTGTGCT | MEMO1 | 51072 |
| CGGCCGGGGCCCTCGCTGTC | TGFB1 | 7040 |
| CGGCCGTCAGCGCTCGGAGC | BCL2 | 596 |
| CGGCCTCCTCCTCAGTTGGA | RND1 | 27289 |
| CGGCCTGCCTTCCCACTGGC | TIFA | 92610 |
| CGGCCTGGCCTGGCCTGTCA | RRAGC | 64121 |
| CGGCCTGGGCTGGGGTTTGA | PDK4 | 5166 |
| CGGCGAGCGAAAGAGTGAGG | BCORL1 | 63035 |
| CGGCGAGGACAGCCGGGACT | JAK1 | 3716 |
| CGGCGCAGGAGCGGCACTCG | SOD2 | 6648 |
| CGGCGCAGGTGAGCAGGGCA | EIF2AK2 | 5610 |
| CGGCGCCCGCGGTTCAGGTG | NFRKB | 4798 |
| CGGCGCCCTTCTCGGTAGCG | EIF4G3 | 8672 |
| CGGCGCCGCCCGGGGCCCTG | SMPD1 | 6609 |
| CGGCGCCGCTCTCCGCTGCG | KAT2A | 2648 |
| CGGCGCCGCTGCTGCCAGGG | PIAS1 | 8554 |
| CGGCGCGCACACTGCTCGCT | SLC7A5 | 8140 |
| CGGCGCGCGGCACAGCCCCG | ARID3A | 1820 |
| CGGCGCGGCCTCTGGTGCAG | CS | 1431 |
| CGGCGCGGCTCCTACCTGCA | PLD1 | 5337 |
| CGGCGCTGGGTCGGTGGCGG | RBM6 | 10180 |
| CGGCGCTGTCCGAGGCGAGC | AHCTF1 | 25909 |
| CGGCGCTTACCTTGGTTTCT | ZRSR2 | 8233 |
| CGGCGGAACCGCGGGGACCT | VIPR2 | 7434 |
| CGGCGGAGGCTTTGGCAGCT | ATP5J | 522 |
| CGGCGGCAGAGAGGAGACTA | APEH | 327 |
| CGGCGGCAGCGGCGCTCGAG | TMEM30A | 55754 |
| CGGCGGCAGGTAGGCACAGT | IRF8 | 3394 |
| CGGCGGCCCGAGGCGGCGCG | FAM107B | 83641 |
| CGGCGGCGAAGCGGCGGCGG | RNF111 | 54778 |
| CGGCGGCGCCCGTAGGATGC | SCN5A | 6331 |
| CGGCGGCGCTGAGGCGGCTG | INO80 | 54617 |
| CGGCGGCGGCGAAGCGGCGG | RNF111 | 54778 |
| CGGCGGCGGCGCAGGTGAGC | EIF2AK2 | 5610 |
| CGGCGGCGGCGGCGAAGCGG | RNF111 | 54778 |
| CGGCGGCGGCGGCGAAGCGG | TRAF5 | 7188 |
| CGGCGGCGGCGGCGGCACCG | NR2C2 | 7182 |
| CGGCGGCGGCGGCGGTTGGC | ANKRD50 | 57182 |
| CGGCGGCGGCGGCTCAACGC | KHSRP | 8570 |
| CGGCGGCGGCGGTTGGCCGG | ANKRD50 | 57182 |
| CGGCGGCGGTTGGCCGGTGG | ANKRD50 | 57182 |
| CGGCGGCTGTAGGGGAGCAG | RNF111 | 54778 |

| | | |
|---|---|---|
| CGGCGGGAGCCAGCGAGCTG | RB1 | 5925 |
| CGGCGGGATTCCTACCCGCC | IFNAR2 | 3455 |
| CGGCGGGGGAGGGACCCTGG | KPNB1 | 3837 |
| CGGCGGGGGCTGCGACGCGG | VDAC1 | 7416 |
| CGGCGGTGATGGACGGGTCC | BAX | 581 |
| CGGCGGTGTCTGGCTTGGTG | PDHB | 5162 |
| CGGCGTCGTCGTCGGGGCTC | USP7 | 7874 |
| CGGCTCATCCGCCGCGTCCG | SATB1 | 6304 |
| CGGCTCCACCATTCAAGCAA | PHIP | 55023 |
| CGGCTCCACTCACCCCCGGT | ZNF641 | 121274 |
| CGGCTCCAGGTGGGCTCACG | CHEK2 | 11200 |
| CGGCTCCGTGAGGCCCTGCC | NCOA6 | 23054 |
| CGGCTCGGTGAGTCGGCGCG | ARID3A | 1820 |
| CGGCTGAGGGCTTCTCGTCG | NDUFA8 | 4702 |
| CGGCTGCCCCTGTTCTGAGC | AHR | 196 |
| CGGCTGCTCGGCGTTCTCTC | TLR2 | 7097 |
| CGGCTGGCGGCTCGGTGAGT | ARID3A | 1820 |
| CGGCTGGCGTGGGCCATCCG | RIPK2 | 8767 |
| CGGCTGGGACTCCCTGGCTG | NUFIP2 | 57532 |
| CGGCTGTCAGCCATAGCGTG | CAT | 847 |
| CGGCTTCAGCAGCCCGCGCC | IRAK4 | 51135 |
| CGGGAAGATGGTGCTGATCA | PITPNB | 23760 |
| CGGGAAGCTGTGCGAGAAGC | CXCL3 | 2921 |
| CGGGAAGGCTCGGTACCACC | HDAC2 | 3066 |
| CGGGACGTCGCGAAAATGTA | NO_CURRENT_441 | NO_CURRENT_441 |
| CGGGAGCAGCAGCAGGTACC | KMT2C | 58508 |
| CGGGAGCCAGCGAGCTGTGG | RB1 | 5925 |
| CGGGAGCCCACCCGGACGAA | RBM42 | 79171 |
| CGGGAGCGGAGAGCGAGGGG | CTNNB1 | 1499 |
| CGGGAGCGGCGGTGATGGAC | BAX | 581 |
| CGGGAGGACAGCCGGAGAAC | SLC25A6 | 293 |
| CGGGAGGAGCGGCGCGCGCT | IDH2 | 3418 |
| CGGGAGGCACCTCGGAGATC | LAMTOR2 | 28956 |
| CGGGAGGGCAGGAGCGAATC | CSNK1D | 1453 |
| CGGGATGCAGCTGGAGAGGA | NO_CURRENT_442 | NO_CURRENT_442 |
| CGGGATGCCTCCATGACCCT | ERCC6L | 54821 |
| CGGGATGGTCCCTGCCGAGA | NO_CURRENT_443 | NO_CURRENT_443 |
| CGGGCCAGGGCAGCGACACC | CCAR2 | 57805 |
| CGGGCCCCGAGGCACAGCCA | NMU | 10874 |
| CGGGCCGCGCTGCGTGCGCT | AKT1 | 207 |
| CGGGCCGGTCATGAACGGGC | INO80E | 283899 |
| CGGGCCGTGTGCCTCGGACG | PHF8 | 23133 |
| CGGGCGCAGGATACGGGCCG | PIP4K2A | 5305 |
| CGGGCGCGCGAGCCTCCGGG | TP73 | 7161 |
| CGGGCGCGGAGAGTGAATGG | SLC25A6 | 293 |
| CGGGCGCGGCCGGAGACCGT | NELFE | 7936 |
| CGGGCGCGGGCTGCTGAAGC | IRAK4 | 51135 |
| CGGGCGCTCACACCGTGCGG | EGFR | 1956 |
| CGGGCGCTTCAGGCTGGCTT | KIF2B | 84643 |
| CGGGCTGGCTCTTACTCAGA | RRAGA | 10670 |

| | | |
|---|---|---|
| CGGGGAACTGTTCGGCTCGC | IFNAR2 | 3455 |
| CGGGGAATTGCACGGCGGAA | NO_CURRENT_444 | NO_CURRENT_444 |
| CGGGGACTGCAACCCTAATC | STAT2 | 6773 |
| CGGGGAGCAGCCCAGAGGCG | BAX | 581 |
| CGGGGAGCCGGCGGCTGCGG | RXRA | 6256 |
| CGGGGCAGGCAGGGCGGGGC | ZNF699 | 374879 |
| CGGGGCTCAGGCGATGCCGG | EIF4G3 | 8672 |
| CGGGGCTCCGGCAGCGGACG | USP7 | 7874 |
| CGGGGCTCCTGACGGTAACT | SHOC2 | 8036 |
| CGGGGCTGGCTCGGACTCCA | BRD1 | 23774 |
| CGGGGGAAAAATAAAACAGA | SLC7A11 | 23657 |
| CGGGGGCACAGAATCACTGA | FBXW7 | 55294 |
| CGGGGGCCAGCAGCCGGGAA | NDUFC1 | 4717 |
| CGGGGGCCCTTGAACTTCTG | JAM2 | 58494 |
| CGGGGGCCGCTCCATGGGGC | CHUK | 1147 |
| CGGGGTAGGGGTTGGCGCTC | ATP6V0E1 | 8992 |
| CGGGGTGCGCGGGCGGTTGT | CREBBP | 1387 |
| CGGGGTGTCCGCGTGAGAAT | METTL3 | 56339 |
| CGGGTCCCTCGGGCTGCACA | BAK1 | 578 |
| CGGGTCTCCAAAGATCGCTT | NO_CURRENT_445 | NO_CURRENT_445 |
| CGGGTGGCGAGGCTGCGGTG | PIGY | 84992 |
| CGGGTGGCGAGGCTGCGGTG | PYURF | 100996939 |
| CGGGTGGCTGAGCAGCGGCC | SNRNP70 | 6625 |
| CGGGTGGTACCGAGCCTTCC | HDAC2 | 3066 |
| CGGTAACCGGCGACTACGTG | BABAM2 | 9577 |
| CGGTAGTATTAATCGCTGAC | NO_CURRENT_446 | NO_CURRENT_446 |
| CGGTCCGCGAGTCGCAAACT | RHOA | 387 |
| CGGTCGCGTCCGACACCCGG | EZH2 | 2146 |
| CGGTCTCCGAGGCTATCTAC | NDUFA12 | 55967 |
| CGGTGAGCCACACGAAGGAA | NO_CURRENT_447 | NO_CURRENT_447 |
| CGGTGCCGGGGGTTCCGCGG | RB1 | 5925 |
| CGGTGCTCCTGGCTGGGCGT | ITGB2 | 3689 |
| CGGTGCTGTGAAAGCCGAGC | NO_CURRENT_448 | NO_CURRENT_448 |
| CGGTGGCTACTGGAGCACTC | CXCR4 | 7852 |
| CGGTGGGACTCAGAAGGCAG | EZH2 | 2146 |
| CGGTGTGCTGGAGTCCTCGG | ITGB2 | 3689 |
| CGGTTGCAGTACCCACTGGA | HSPA4 | 3308 |
| CGGTTGGCCGGTGGGGGCTG | ANKRD50 | 57182 |
| CGGTTGTAGAAGGTATGAGG | CASP3 | 836 |
| CGGTTTACATCTGCCCATCG | NO_CURRENT_449 | NO_CURRENT_449 |
| CGGTTTGTATCCGGGCTGTG | ABI1 | 10006 |
| CGTAACTAGGGGGGGCGAGT | PHC3 | 80012 |
| CGTAATGGCGGACACAGGCA | WASF2 | 10163 |
| CGTAATTTTGTAATCGCTTC | NO_CURRENT_450 | NO_CURRENT_450 |
| CGTAGCGCGCTGAGGGGGTC | WASF2 | 10163 |
| CGTAGGCATGCTGCGCCCCA | LAMTOR2 | 28956 |
| CGTAGTAAATATCTAGCTAA | NO_CURRENT_451 | NO_CURRENT_451 |
| CGTCAAGTATTAAGCTGCTT | NO_CURRENT_452 | NO_CURRENT_452 |
| CGTCAATTCCCATGCCCTTG | CYP1B1 | 1545 |
| CGTCATATACACAAACGCCC | NO_CURRENT_453 | NO_CURRENT_453 |

| | | |
|---|---|---|
| CGTCCAGAAGAACGGCCCCT | NO_CURRENT_454 | NO_CURRENT_454 |
| CGTCCCTGGCTGACAACGAA | MPC2 | 25874 |
| CGTCCCTTCGTCTCTGCTTA | NO_CURRENT_455 | NO_CURRENT_455 |
| CGTCCGCAGGAAGAGGCGCG | PARK7 | 11315 |
| CGTCGCCATATGCCGGTGGC | NO_CURRENT_456 | NO_CURRENT_456 |
| CGTCGCCGCTGCCGCCGCCA | ARHGAP33 | 115703 |
| CGTCGCCGCTGCCGCCGCCA | KIF5B | 3799 |
| CGTCGCTTCGGCCAGTGTGT | HIF1A | 3091 |
| CGTCGGGCGCTCACACCGTG | EGFR | 1956 |
| CGTCGGGTAGCTATTTCTTT | NO_CURRENT_457 | NO_CURRENT_457 |
| CGTCGTCGGGGCTCCGGCAG | USP7 | 7874 |
| CGTCGTGGGTGTTTGGTGTG | LGMN | 5641 |
| CGTCTGCGTCTGCAGCTGCA | RNF25 | 64320 |
| CGTCTGTCGGGGGCGCGCTC | BABAM2 | 9577 |
| CGTGAGAATTGGCTATATCC | METTL3 | 56339 |
| CGTGAGTAACTTGGCTCCGG | ANP32B | 10541 |
| CGTGAGTTGCATGTTGTGTG | RAD23A | 5886 |
| CGTGCCTTTACATTCACTTT | NO_CURRENT_458 | NO_CURRENT_458 |
| CGTGCGAAGCTCACCGACTC | NUDT17 | 200035 |
| CGTGCGAGCGGGGGGAGGGG | BMI1 | 648 |
| CGTGCGGTAAATACGAAATA | NO_CURRENT_459 | NO_CURRENT_459 |
| CGTGCTCTGCAGGTAGAAAA | TBXAS1 | 6916 |
| CGTGCTGCTTATGGCGGCGC | CSDE1 | 7812 |
| CGTGGAAACCAGGAGTCCCT | HTR2A | 3356 |
| CGTGTAAAAATACCTTTCTA | NO_CURRENT_460 | NO_CURRENT_460 |
| CGTGTAGGAGGGCATGGCGC | ALOX5 | 240 |
| CGTGTCGCCGCTACGATACC | PQBP1 | 10084 |
| CGTGTGCGGTCGGGCGATTC | ACACA | 31 |
| CGTGTGTGGGTAAACGGAAA | NO_CURRENT_461 | NO_CURRENT_461 |
| CGTGTGTGTGCGTGCGAGCG | BMI1 | 648 |
| CGTGTTTGGAATTTGCCGCG | NO_CURRENT_462 | NO_CURRENT_462 |
| CGTTCTCTCAGGTGACTGCT | TLR2 | 7097 |
| CGTTCTGGGTCCGAGGGTCC | SLC3A2 | 6520 |
| CGTTGAGAGAAGGTCTCATT | PQBP1 | 10084 |
| CGTTGGCCCGGCCCCGGGAG | CTNNB1 | 1499 |
| CGTTGGGCATAGCGAACACT | NO_CURRENT_463 | NO_CURRENT_463 |
| CGTTGGGGATCTGGAGTCCC | LARP4B | 23185 |
| CTAAAGACGCTTCTTCCCGG | IFNAR2 | 3455 |
| CTAAATCTGTATCTAACCGG | NO_CURRENT_464 | NO_CURRENT_464 |
| CTAAATGTTATGCAATGCAC | NO_CURRENT_465 | NO_CURRENT_465 |
| CTAAGTTTGTTAATGGGCCA | NO_CURRENT_466 | NO_CURRENT_466 |
| CTAATATTCGGCCGCGGAGA | PIAS1 | 8554 |
| CTAATCACGACCTCACCCTA | NO_CURRENT_467 | NO_CURRENT_467 |
| CTAATGCTATCAATCATGAG | NO_CURRENT_468 | NO_CURRENT_468 |
| CTACCCCCAGTTGGACCCTG | CSDE1 | 7812 |
| CTACCCCCGCCACGGATCGC | MAP3K7 | 6885 |
| CTACCCGGCGATCCGTGGCG | MAP3K7 | 6885 |
| CTACCCTCCCGCGGCCGGCC | KIAA1211L | 343990 |
| CTACGCCCGCAAGAGCAACA | HSPA13 | 6782 |
| CTACGCGCGCCCACCTCCCG | LRP2 | 4036 |

| | | |
|---|---|---|
| CTACGGCGGGGGCTGCGACG | VDAC1 | 7416 |
| CTACTGCTTGCTGTGGCATT | GC | 2638 |
| CTAGAAGTCTAGAGTCACGG | NO_CURRENT_469 | NO_CURRENT_469 |
| CTAGAATAAGTGAGAAAGTG | DPY30 | 84661 |
| CTAGAATAAGTGAGAAAGTG | MEMO1 | 51072 |
| CTAGCAGTTTCCGTCACTCC | ZNF616 | 90317 |
| CTAGCCGCCCAGATCGAGCC | NO_CURRENT_470 | NO_CURRENT_470 |
| CTAGCGAGCGCAGCGGAGCC | TNFRSF1B | 7133 |
| CTAGCTCAATATACAATGTG | NO_CURRENT_471 | NO_CURRENT_471 |
| CTAGCTTAGGAGTTATACCG | NO_CURRENT_472 | NO_CURRENT_472 |
| CTAGGAACTGAGAGGAAGAA | BCORL1 | 63035 |
| CTAGGCCAAAACCCTACTTC | NDUFA1 | 4694 |
| CTAGGGTCTGGCTGGAGCCA | CYP24A1 | 1591 |
| CTAGTGTTTGGGTTTCTTCG | CHMP5 | 51510 |
| CTATATCCTGGAGCGAGTGC | METTL3 | 56339 |
| CTATATTGTCGCGCAGTGGA | NO_CURRENT_473 | NO_CURRENT_473 |
| CTATCCTGACTGATGCCCAA | NO_CURRENT_474 | NO_CURRENT_474 |
| CTATCTCGGGCATGGCTCTG | LTA4H | 4048 |
| CTATGATCACTGGAGTCTCG | BAK1 | 578 |
| CTATGCCGGTTCCAACAACC | ALOX15 | 246 |
| CTATTGTACAAAGTTAGGAA | ACOD1 | 730249 |
| CTCAAGATGAACCGACTCTT | CHMP5 | 51510 |
| CTCAAGTCCCCTTCGATCGC | HSP90B1 | 7184 |
| CTCACACGGCTCGGCGGGTG | SLC25A19 | 60386 |
| CTCACCCGCCACACCCGGAC | NFKB2 | 4791 |
| CTCACCTCCTTTAGGCATTG | PDAP1 | 11333 |
| CTCACCTGAACCGCGGGCGC | NFRKB | 4798 |
| CTCACCTGGTGCCCTCCCCG | RHOG | 391 |
| CTCACGGCCGCTCGGGAGCC | NDUFC1 | 4717 |
| CTCACGGGGACATACAGGGC | NO_CURRENT_475 | NO_CURRENT_475 |
| CTCACTATCCTGAATGATCG | NCKAP1L | 3071 |
| CTCACTCTTACCTGCCAACC | CUBN | 8029 |
| CTCACTTACCCTGGGCCTCA | EP400 | 57634 |
| CTCAGAAGGCAGTGGAGCCC | EZH2 | 2146 |
| CTCAGCACTGCTCTGTTGCC | IL10 | 3586 |
| CTCAGCCTGCACCCCCTCTC | SLC11A1 | 6556 |
| CTCAGCTCCAGCTCACTGGC | NR1H3 | 10062 |
| CTCAGGGCCCGTACCTGATT | STAT2 | 6773 |
| CTCAGGGTGAGTCCCGCGGG | YPEL5 | 51646 |
| CTCAGTATCAAACCGCTTCT | FBXW7 | 55294 |
| CTCATCTCGGGCAGAGCGCT | FOSL2 | 2355 |
| CTCATCTGCTCAGGTGTCCC | SLC5A8 | 160728 |
| CTCATGAGTCGTTTCTTTCA | NO_CURRENT_476 | NO_CURRENT_476 |
| CTCATGGGGCGTCCTCCTGC | SERPINA2 | 390502 |
| CTCCAAACATCGCGATTAAT | BTN2A2 | 10385 |
| CTCCACCATTCAAGCAACGG | PHIP | 55023 |
| CTCCAGCACTGGGTCCGGTG | ACTR5 | 79913 |
| CTCCAGCAGTGCCTGAGAGC | PHF6 | 84295 |
| CTCCAGGACTGCTGCGCTGG | DNAJB6 | 10049 |
| CTCCAGGAGGGTGAGCGCGG | HELZ2 | 85441 |

| | | |
|---|---|---|
| CTCCAGGGGACCCACAGCTG | RIPK1 | 8737 |
| CTCCAGTCCTCCCCGGTGCT | IL2RB | 3560 |
| CTCCAGTTGTGAGAGCCGCA | CYP1B1 | 1545 |
| CTCCATATACAGCCCGGGAT | TNRC18 | 84629 |
| CTCCATGTTGCCGCCTCTCC | KMT2C | 58508 |
| CTCCCACCACCAGCCCTCTG | GNAI2 | 2771 |
| CTCCCAGTACCAGTCAGTTC | NO_CURRENT_477 | NO_CURRENT_477 |
| CTCCCATTGATCTACGATGG | NO_CURRENT_478 | NO_CURRENT_478 |
| CTCCCCACTCACCCTCTGAC | UBE2H | 7328 |
| CTCCCCAGACCTTTCAGTTG | PAGR1 | 79447 |
| CTCCCCGGACTCTCCGTCAT | NDUFA1 | 4694 |
| CTCCCCGGCCTCGCTCCCTC | SPHK1 | 8877 |
| CTCCCCTCGCTCTCCGCTCC | CTNNB1 | 1499 |
| CTCCCCTGAGAGCCTGAACC | FLCN | 201163 |
| CTCCCCTGAGAGCCTGAACC | PLD6 | 201164 |
| CTCCCGCGCTCCCGCGCGCG | TIFA | 92610 |
| CTCCCGGCCCCCGACCCACG | ARID3A | 1820 |
| CTCCCGGGCCGGTCATGAAC | INO80E | 283899 |
| CTCCCGTTCCCCGGACGCGG | SATB1 | 6304 |
| CTCCCTCCCACCTCCCTCCG | TGFB1 | 7040 |
| CTCCCTGAGCTTCTCCTCCT | CYP4B1 | 1580 |
| CTCCCTGCCGGCCGGGTTAG | NO_CURRENT_479 | NO_CURRENT_479 |
| CTCCCTGCTGGCTCCGAGGC | JUNB | 3726 |
| CTCCGCCGCCAGCCGCCTCC | SETD1B | 23067 |
| CTCCGCCGCCGTTGCTTGAA | PHIP | 55023 |
| CTCCGCTCCCGTGAGTAACT | ANP32B | 10541 |
| CTCCGCTGCGGGGGAGGCCA | KAT2A | 2648 |
| CTCCGGATCTCGCTCTCCAC | NFKB2 | 4791 |
| CTCCGGCCGCGTCAGGAGGG | EP400 | 57634 |
| CTCCGGGCCCTCCCTGCCGG | BCL2 | 596 |
| CTCCGTCCTTTCGGTCCAGG | SLC25A6 | 293 |
| CTCCTAACCAGGCTGCATGA | CDKN2C | 1031 |
| CTCCTACCTGCAGGGCGAAG | PLD1 | 5337 |
| CTCCTACTGTTGGACACACC | CXCR1 | 3577 |
| CTCCTATTAATCGCGATGTT | BTN2A2 | 10385 |
| CTCCTCCATTGAAAGCGGCT | NUFIP2 | 57532 |
| CTCCTCCCTCCCGCTCTCTC | AHDC1 | 27245 |
| CTCCTCTCAGTGAGTGAAAG | INO80 | 54617 |
| CTCCTCTGCGGCCACTGAGC | HSPA4 | 3308 |
| CTCCTTACGTCGGGCATTAA | NO_CURRENT_480 | NO_CURRENT_480 |
| CTCCTTCCACCTAATAGTTA | ABL2 | 27 |
| CTCGACAGTTCGTCCCGAGC | NO_CURRENT_481 | NO_CURRENT_481 |
| CTCGAGGTCCCTTCCCACAG | TLR9 | 54106 |
| CTCGCACCATTGAGGGTAGT | NO_CURRENT_482 | NO_CURRENT_482 |
| CTCGCATTTCCCGCAGCCCC | KMT2C | 58508 |
| CTCGCCGTTCCTTCCCCGGG | LRP2 | 4036 |
| CTCGCGGGCTGCCAAACGGC | WASF2 | 10163 |
| CTCGCTGGGCCGCGGCTCCC | SLC7A5 | 8140 |
| CTCGCTGTCTGGCTGCTCCG | TGFB1 | 7040 |
| CTCGGACCCGAAGCCGCCAC | SLC25A1 | 6576 |

| | | |
|---|---|---|
| CTCGGAGAAACAGGCGCCGC | SBNO2 | 22904 |
| CTCGGAGCCAGCAGGGAGCT | JUNB | 3726 |
| CTCGGAGGCGGCTACGGCGG | VDAC1 | 7416 |
| CTCGGCAACACCGGCTTCCT | MOSPD1 | 56180 |
| CTCGGCCCACTTGCGTTATA | NO_CURRENT_483 | NO_CURRENT_483 |
| CTCGGGAAACGTCATCTTCT | ZRSR2 | 8233 |
| CTCGGGCCCAGAGTCACATG | NO_CURRENT_484 | NO_CURRENT_484 |
| CTCGGTGAGAGGCGGAGGAG | DNAJA1 | 3301 |
| CTCGTGAAAAAACCCAGAAC | ATP5J | 522 |
| CTCTAAACTGTAAAGTCCAA | NO_CURRENT_485 | NO_CURRENT_485 |
| CTCTAAAGGGGAGGGGCCGA | ARNT | 405 |
| CTCTCAACCACAATAACAGG | FBXO38 | 81545 |
| CTCTCCACAACTCCACTTAC | GLMN | 11146 |
| CTCTCCCCCTTCGTCCGGGT | RBM42 | 79171 |
| CTCTCGCGTTAAAGAGACAG | ABI1 | 10006 |
| CTCTCGGCCACCTTTGATGA | NOS2 | 4843 |
| CTCTCGGGCTTGGGCGCCCC | IZUMO2 | 126123 |
| CTCTCGGTTGCAGTACCCAC | HSPA4 | 3308 |
| CTCTCTCCTACCTGTCCACC | NO_CURRENT_486 | NO_CURRENT_486 |
| CTCTGCCACCATGGGGAACT | CYBB | 1536 |
| CTCTGCGCACCGCTGGGGTT | SIAH2 | 6478 |
| CTCTGCTCTCCAAGCTGGGT | AHDC1 | 27245 |
| CTCTGGAGTTTGGGCGGCCC | PPID | 5481 |
| CTCTGGCGCGATAGCTTTCC | JUNB | 3726 |
| CTCTGGTAGAAAAATGAAGA | GC | 2638 |
| CTCTGTCTCTTTAACGCGAG | ABI1 | 10006 |
| CTCTTAAGAACGAACGGCTT | DPY30 | 84661 |
| CTCTTAAGAACGAACGGCTT | MEMO1 | 51072 |
| CTCTTACCTGCCAACCTGGG | CUBN | 8029 |
| CTCTTATTGGGTGTATATCC | NO_CURRENT_487 | NO_CURRENT_487 |
| CTCTTTTGAGATTGACAAGT | NO_CURRENT_488 | NO_CURRENT_488 |
| CTGAAGCACATCCCGCAGCC | BNIP3 | 664 |
| CTGACGGCCGCCGGCAGGGA | BCL2 | 596 |
| CTGACTGAGCACTGTCATAG | NO_CURRENT_489 | NO_CURRENT_489 |
| CTGACTGCAGCTCCTACTGT | CXCR1 | 3577 |
| CTGAGAAAATTAAAGCAGAG | TNFRSF1A | 7132 |
| CTGAGAAGCCAAGACTGAGC | FOS | 2353 |
| CTGAGAGCGAGAGGTGGATC | UCHL5 | 51377 |
| CTGAGAGCTAGAGAAGAGCC | NOD1 | 10392 |
| CTGAGATCACGTCACTACAC | NO_CURRENT_490 | NO_CURRENT_490 |
| CTGAGATGTCAGATGAACCG | NO_CURRENT_491 | NO_CURRENT_491 |
| CTGAGCCCCAGCAGACTCCT | CYP4A11 | 1579 |
| CTGAGCGGCGGTGAGATCCG | YPEL5 | 51646 |
| CTGAGCTGAGTCTCGCTGCT | KPNA1 | 3836 |
| CTGAGCTGGTTTGTAATGAT | SLC7A11 | 23657 |
| CTGAGGCGGCTGCGGGCGCT | INO80 | 54617 |
| CTGAGGCTGCGGAGAGTGTG | CHEK2 | 11200 |
| CTGAGGGCATGCGTCCACCC | CYP2R1 | 120227 |
| CTGAGTAAGTGATCACGAAG | NO_CURRENT_492 | NO_CURRENT_492 |
| CTGAGTGAAAAATAAAAGTT | NO_CURRENT_493 | NO_CURRENT_493 |

| | | |
|---|---|---|
| CTGAGTGCAAGGTGACTGGG | ATP5J | 522 |
| CTGATGTCAACTAAACACCA | NO_CURRENT_494 | NO_CURRENT_494 |
| CTGCAACCCTAATCAGGTAC | STAT2 | 6773 |
| CTGCAGACCAGGCGGCTGCG | BCL2L11 | 10018 |
| CTGCAGCAGATCACCGGCCA | NDUFA12 | 55967 |
| CTGCAGCCGCGGCTGGAAGA | PPARGC1B | 133522 |
| CTGCAGGAGGCCTCGGCTCG | CAT | 847 |
| CTGCAGGAGGTCCCGGCGCG | MYD88 | 4615 |
| CTGCAGGCAGCCCCAGCCTC | IZUMO2 | 126123 |
| CTGCAGGTAGAAAAGGGAAC | TBXAS1 | 6916 |
| CTGCCAGCCAGTGGGAAGGC | TIFA | 92610 |
| CTGCCCAAGCCCGACTTTCA | NDUFS8 | 4728 |
| CTGCCCAGACTCCCCACCTC | APAF1 | 317 |
| CTGCCCAGTCCCCTCATCAA | NOS2 | 4843 |
| CTGCCCCAGGCGTAATCCTC | NO_CURRENT_495 | NO_CURRENT_495 |
| CTGCCCCCCTCCCATGGGCA | PDE4A | 5141 |
| CTGCCCCTGTTCTGAGCTGG | AHR | 196 |
| CTGCCCTCACCATGAGCCTC | MMP9 | 4318 |
| CTGCCGGACTGACGGGCGGC | MAP2K7 | 5609 |
| CTGCCGGGTCGGGCTGCGGG | NCOA6 | 23054 |
| CTGCCTTCCTCTTCTTGAGC | MMP13 | 4322 |
| CTGCCTTTAGAGAGCTGGAG | GPR119 | 139760 |
| CTGCGACCGTCGCGGACCAT | ARRB1 | 408 |
| CTGCGACGCTCCTGCTGCCG | TRMT61A | 115708 |
| CTGCGCACCGCTGGGGTTCG | SIAH2 | 6478 |
| CTGCGGGAGGCTCGGTGCTA | NMU | 10874 |
| CTGCGGGCGGCCAGCGAAGG | TRAF6 | 7189 |
| CTGCGGTGAGGCCTGGTCTC | PIGY | 84992 |
| CTGCGGTGAGGCCTGGTCTC | PYURF | 100996939 |
| CTGCGTCCGCGGCGGCTTGG | CYFIP1 | 23191 |
| CTGCGTGGTCGCACCCTACC | NMRK2 | 27231 |
| CTGCGTGTCTTGCTCGCATG | NO_CURRENT_496 | NO_CURRENT_496 |
| CTGCTAGCGAACGCTCCTTT | LRP2 | 4036 |
| CTGCTCCTCCTCGGACCAGG | SCTR | 6344 |
| CTGCTCGCCGGACGGCTCCC | GSDMD | 79792 |
| CTGCTCTATCCCGGGCAAAA | APAF1 | 317 |
| CTGCTGCCCGAAGCTGGGGC | PDE5A | 8654 |
| CTGCTGCCTACCTCCCGAAG | PDHB | 5162 |
| CTGCTGCTGCTGCTGTTCTG | MMP1 | 4312 |
| CTGCTGTAGCCAGTGCAAAC | UBA7 | 7318 |
| CTGCTTAATTCCCTTTCCTT | PPARG | 5468 |
| CTGCTTGGAGCAAAACAAAA | FOXO4 | 4303 |
| CTGCTTGGCAAGGCAAGTAG | NDUFS8 | 4728 |
| CTGGACAGCAGCTACAGGGA | TLR9 | 54106 |
| CTGGACCCTAGCCAGCCCTC | SMPD1 | 6609 |
| CTGGACTGAGGCTCCAGTTC | TNFRSF1A | 7132 |
| CTGGAGGCCAATCCCGAGGT | UCHL3 | 7347 |
| CTGGAGGCGGCGGAACCGCG | VIPR2 | 7434 |
| CTGGATCGCCCGCAGAAATA | NO_CURRENT_497 | NO_CURRENT_497 |
| CTGGCCAGGAAGCGGGTGCC | ASH2L | 9070 |

| | | |
|---|---|---|
| CTGGCCCAGCGCACGCAGCG | AKT1 | 207 |
| CTGGCCGAATCTCACTATGT | NO_CURRENT_498 | NO_CURRENT_498 |
| CTGGCCGGTAAGTGGAGTTG | GLMN | 11146 |
| CTGGCCTGTCAGGGCGCGGG | RRAGC | 64121 |
| CTGGCGCGTCTGCTCTCCCT | GSDMD | 79792 |
| CTGGCTCCTCCGTGCCGCCG | UBE2E1 | 7324 |
| CTGGCTCGGACTCCAGGGCC | BRD1 | 23774 |
| CTGGCTGAGACATGGGGCGG | IL2RB | 3560 |
| CTGGCTGGTGCCCCAGGGCC | NLRP1 | 22861 |
| CTGGGAGATGAAGACAGACC | C6orf15 | 29113 |
| CTGGGAGCCGTCCGGCGAGC | GSDMD | 79792 |
| CTGGGAGCGCGCCGGGCGCA | KIAA1211L | 343990 |
| CTGGGAGCTGGGAGCGCGCC | KIAA1211L | 343990 |
| CTGGGCAGAGGTATGGTCCT | ABL2 | 27 |
| CTGGGCATAGGAACATGGAG | CYP24A1 | 1591 |
| CTGGGCATGCGCCACTTGTG | TMEM70 | 54968 |
| CTGGGGGCGTAGCGCGCTGA | WASF2 | 10163 |
| CTGGGTAAGTCCAGCTCCGC | NLRP3 | 114548 |
| CTGGGTCATGGTCTGGTTCA | CYP27B1 | 1594 |
| CTGGGTCCGAGGGTCCAGGT | SLC3A2 | 6520 |
| CTGGGTGCCGCGTCTGCCGG | SATB2 | 23314 |
| CTGGGTGGCGGGCGGCGTGC | ANP32B | 10541 |
| CTGGTCACCAGCAGAGGTTA | PPARG | 5468 |
| CTGGTGACCGACAATTACAC | NO_CURRENT_499 | NO_CURRENT_499 |
| CTGGTGCCCTCCCCGAGGCG | RHOG | 391 |
| CTGGTGGGTGCGCCTTGGCC | PTGES2 | 80142 |
| CTGGTTAGGAGCAAAGGAAA | CDKN2C | 1031 |
| CTGTACCAAGAGTTTGCTCC | CCL20 | 6364 |
| CTGTATAATAATCATAAATC | NO_CURRENT_500 | NO_CURRENT_500 |
| CTGTCCACCTACAGCGATGT | NO_CURRENT_501 | NO_CURRENT_501 |
| CTGTCGCCCACACTGCCCCG | NLRP3 | 114548 |
| CTGTCTGGCTGCTCCGCGGA | TGFB1 | 7040 |
| CTGTCTGGCTGTCCCACTGC | HAMP | 57817 |
| CTGTGAAACAACGAACCCCC | IRGM | 345611 |
| CTGTGACTCTGTGACTCGGT | NO_CURRENT_502 | NO_CURRENT_502 |
| CTGTGAGGTGCTCGGAGCCT | ABI1 | 10006 |
| CTGTGCTCCCTATATGAGAG | HTR3E | 285242 |
| CTGTGGCAGCAGCGTTGGCC | CTNNB1 | 1499 |
| CTGTTATGTCCATCTAACAC | NO_CURRENT_503 | NO_CURRENT_503 |
| CTGTTCCAAGAACAGGTCCC | CARD16 | 114769 |
| CTGTTCCAAGAACAGGTCCC | CASP1 | 834 |
| CTGTTCTGAGCTGGCGGTCC | AHR | 196 |
| CTGTTGGAAGACGCTAGGCA | NO_CURRENT_504 | NO_CURRENT_504 |
| CTGTTTGCACTGGCTACAGC | UBA7 | 7318 |
| CTTAAGGACCGTGAGGGGTA | SPCS1 | 28972 |
| CTTAAGGCGAGAAAAATTAG | NO_CURRENT_505 | NO_CURRENT_505 |
| CTTACTCCAAGGAGCCTGTG | C4orf17 | 84103 |
| CTTAGCTGACCGACAAGGTG | NO_CURRENT_506 | NO_CURRENT_506 |
| CTTAGGATTCCGAGGTATCT | NO_CURRENT_507 | NO_CURRENT_507 |
| CTTATAGGTAGTATCGCAGA | NO_CURRENT_508 | NO_CURRENT_508 |

| | | |
|---|---|---|
| CTTATGGCGGCGCTGGAGAG | CSDE1 | 7812 |
| CTTCACACTGCTCGGGCTCC | HMGN2 | 3151 |
| CTTCAGTTCACTCTCAGTAA | PPARGC1A | 10891 |
| CTTCATGGTTGCAGTGTCCG | RND1 | 27289 |
| CTTCCAAGGCAGCCCGGGTA | NMRK2 | 27231 |
| CTTCCAGTCCGCGGAGGGCG | RAC1 | 5879 |
| CTTCCCTAACGTTGCAACTG | MAP2K6 | 5608 |
| CTTCCGTTATTCGGAAGTGA | NO_CURRENT_509 | NO_CURRENT_509 |
| CTTCCTAGTCCCGGGCAGGC | TLR2 | 7097 |
| CTTCCTGGGGATCCTCAACC | SERPINA2 | 390502 |
| CTTCCTTCCAGTAAGGAGTC | MCL1 | 4170 |
| CTTCGACGCCATCGTGCTCA | NO_CURRENT_510 | NO_CURRENT_510 |
| CTTCGCTCCGCGCAGCCGCC | BCL2L11 | 10018 |
| CTTCGGCCAGTGTGTCGGGC | HIF1A | 3091 |
| CTTCGGGAAAGCGAAACCCA | CHMP5 | 51510 |
| CTTCGTCGCCATAACTCGCT | SOD1 | 6647 |
| CTTCTCACCCGCACACTGCC | SOD2 | 6648 |
| CTTGACTGAGAAGAACTCGA | FBXW7 | 55294 |
| CTTGACTTACGAGCCAGTGT | NO_CURRENT_511 | NO_CURRENT_511 |
| CTTGAGCTGGACTCATTGTC | MMP13 | 4322 |
| CTTGAGTTTATGTTTCACGG | NO_CURRENT_512 | NO_CURRENT_512 |
| CTTGATCAGCACCATCTTCC | PITPNB | 23760 |
| CTTGCACCAGTGCCCAGAGA | SLC11A1 | 6556 |
| CTTGCGGCGGAGGGCAGGAA | TRERF1 | 55809 |
| CTTGCTCCCAAATGGGACTC | EDA2R | 60401 |
| CTTGGCAAGGCAAGTAGCGG | NDUFS8 | 4728 |
| CTTGGCATGGGGCTTCCTCT | PIGL | 9487 |
| CTTGGGCGCCCCCGGAGGCT | IZUMO2 | 126123 |
| CTTGGGGCGCAGCATGCCTA | LAMTOR2 | 28956 |
| CTTGGTCCTTTCGCAGAGAG | RIPK3 | 11035 |
| CTTGGTGCGGAGACCCCTTC | PDHB | 5162 |
| CTTGGTGGGTCCCGGGCCAG | CHMP6 | 79643 |
| CTTGGTGTCCCACCGCCACA | IKBKE | 9641 |
| CTTGTACCTCCTGCGCGCCG | PDE3A | 5139 |
| CTTGTCCCTGTGCAGCCCGA | BAK1 | 578 |
| CTTGTTGCGTATACGAGACT | NO_CURRENT_513 | NO_CURRENT_513 |
| CTTTAAGCAGGGATTCGGGG | TFPT | 29844 |
| CTTTAGAAGGACATGTGGAC | NO_CURRENT_514 | NO_CURRENT_514 |
| CTTTCAAGGACCCAGAAGTA | NDUFA1 | 4694 |
| CTTTCAATGGAGGAGAAGCC | NUFIP2 | 57532 |
| CTTTCGGCACCTCTGCCGGG | HDAC2 | 3066 |
| CTTTCTCTGCTCTCCAAGCT | AHDC1 | 27245 |
| CTTTGGGATGCTTTGTTGTC | BTN2A2 | 10385 |
| CTTTGGGGTTCAAGATCACT | TNFRSF1A | 7132 |
| CTTTTTTTATTTATCGATCG | NO_CURRENT_515 | NO_CURRENT_515 |
| GAAAAAGGTGGGGGACCAG | INO80C | 125476 |
| GAAAACAAAACTTTCAACTT | SLC7A11 | 23657 |
| GAAAAGTTCTAGGACGACTG | AGER | 177 |
| GAAAATTAAAGCAGAGAGGA | TNFRSF1A | 7132 |
| GAAAATTCCCCACCGGGCAC | PTGS1 | 5742 |

| | | |
|---|---|---|
| GAAACAAATAGTGATTACTC | NO_CURRENT_516 | NO_CURRENT_516 |
| GAAACACACACACAGCGATA | ZNF616 | 90317 |
| GAAACATAATGTCTTCAGTC | NO_CURRENT_517 | NO_CURRENT_517 |
| GAAACATCTTTGAGCAAGAT | ALOX15 | 246 |
| GAAACATGCTGAAGTCCCGG | ADRB1 | 153 |
| GAAACGAGAAGTTTGTACTA | NO_CURRENT_518 | NO_CURRENT_518 |
| GAAACTCCCAGGGCCCGCCC | CYFIP1 | 23191 |
| GAAACTGCGCGGAGGCACAG | SLC3A2 | 6520 |
| GAAAGAAATTGAATTTGCAG | MCTS1 | 28985 |
| GAAAGACCTTGTAACCATGA | NO_CURRENT_519 | NO_CURRENT_519 |
| GAAAGACGTTGGCAACTCAG | TFPT | 29844 |
| GAAAGCAGGTTAAGACATGA | NO_CURRENT_520 | NO_CURRENT_520 |
| GAAAGGGGAGTTCAAGGAGA | NDUFA8 | 4702 |
| GAAAGGGGTCATCCTATACG | UBE2L6 | 9246 |
| GAAAGTCACCTAGGTTATTT | AIM2 | 9447 |
| GAAAGTCCCCACACTCCTCT | TLR1 | 7096 |
| GAAAGTTTCCAACTTCTGTT | OR7C2 | 26658 |
| GAAATCCCCGTTTACTACGA | NO_CURRENT_521 | NO_CURRENT_521 |
| GAACAAGGCCTCGCTCCACA | KHSRP | 8570 |
| GAACACACACTGACAGCTAT | PHIP | 55023 |
| GAACAGGCCCATGCGCGCAG | PRKN | 5071 |
| GAACATCTCCTCAGGAACTT | PAXIP1 | 22976 |
| GAACCAAGGCCAAGCGAATA | UBA7 | 7318 |
| GAACCAATTCTCCTGTCGGC | NIPBL | 25836 |
| GAACCAGCGGTTACCATGGA | CXCR4 | 7852 |
| GAACCATGGCTATGCTCCTG | CYP8B1 | 1582 |
| GAACCCAACCTTTTACCGCA | NO_CURRENT_522 | NO_CURRENT_522 |
| GAACCTCCCCGAATATCTGG | NO_CURRENT_523 | NO_CURRENT_523 |
| GAACGACTGCGGCGCGCTGC | PPARGC1B | 133522 |
| GAACGCTCGGTGAGAGGCGG | DNAJA1 | 3301 |
| GAACGGGCGCGGAGAGTGAA | SLC25A6 | 293 |
| GAACGGTGAGAGTCTTTGGT | NO_CURRENT_524 | NO_CURRENT_524 |
| GAACGTAGAAATTCCCATTT | NO_CURRENT_525 | NO_CURRENT_525 |
| GAACGTCCAAGCAAGGGAGC | NO_CURRENT_526 | NO_CURRENT_526 |
| GAACGTTCCCTCAAAATGCA | NO_CURRENT_527 | NO_CURRENT_527 |
| GAACTAGGCTCTCGGTGAGC | NQO1 | 1728 |
| GAACTCTTACTCATTGCCGG | NO_CURRENT_528 | NO_CURRENT_528 |
| GAACTGAGAGGAAGAAGGGA | BCORL1 | 63035 |
| GAACTTCGATTCAGAAGGGA | RND1 | 27289 |
| GAAGACCCCCAGTTACTTAC | UQCRB | 7381 |
| GAAGACGGCGGCGGCGGCGA | IKZF5 | 64376 |
| GAAGACTCAACAGATATGTT | NO_CURRENT_529 | NO_CURRENT_529 |
| GAAGAGCCCGGAGAGCGCCA | NOD1 | 10392 |
| GAAGAGCCGCGTGAGGAAAC | LTB4R | 1241 |
| GAAGAGCGCGGCGCTCATGG | CEBPD | 1052 |
| GAAGATCGGCCACTACATTC | PRKAA1 | 5562 |
| GAAGCAGAAACACGACGGGC | PRKAA1 | 5562 |
| GAAGCAGAGGGCCTTACCCG | NELFE | 7936 |
| GAAGCATCTATTATCGTACC | NO_CURRENT_530 | NO_CURRENT_530 |
| GAAGCCCCCTGCCCAGAGCT | SEC62 | 7095 |

| | | |
|---|---|---|
| GAAGCCGCCATCTTTACAGC | BABAM1 | 29086 |
| GAAGCCGGCGCGCCTGCAGC | KIAA1211L | 343990 |
| GAAGCCGGGAGCTCGGCCAC | PIGY | 84992 |
| GAAGCCGGGAGCTCGGCCAC | PYURF | 100996939 |
| GAAGCCGTGTCTCGCAGTCG | TMEM70 | 54968 |
| GAAGCCTGAGCGGCCGAACT | SLC7A3 | 84889 |
| GAAGCGATCCGAGAGTGTAT | NO_CURRENT_531 | NO_CURRENT_531 |
| GAAGCGGGACCGTGTCTCAC | NO_CURRENT_532 | NO_CURRENT_532 |
| GAAGCTCAGGGAGAGGAAGC | CYP4B1 | 1580 |
| GAAGCTCTTTCGCGGCGCTA | RIPK2 | 8767 |
| GAAGCTGTCAGCGCGTCGCC | NFKB1 | 4790 |
| GAAGGAAGCAGGATGGCTGC | AGER | 177 |
| GAAGGACCCAGAAGCGCGGC | AHCTF1 | 25909 |
| GAAGGAGGAGCGGGCCGTGG | RBM6 | 10180 |
| GAAGGCCTACCTTCTGGGCA | SQSTM1 | 8878 |
| GAAGGGAGGGCTCAGCAAAG | BCORL1 | 63035 |
| GAAGGGGACGCAGCGAAACC | TP73 | 7161 |
| GAAGGGGCCTTAACCTCTGC | PPARG | 5468 |
| GAAGGGGGCTCGGTGCTCAC | SERF2 | 10169 |
| GAAGGGGTCTGTCTGTGCAG | RIPK3 | 11035 |
| GAAGGGTAATCGGGTGTCCC | SMPD1 | 6609 |
| GAAGGTAATCAAGTTACATG | NO_CURRENT_533 | NO_CURRENT_533 |
| GAAGGTATGAGGAGGCTGTG | CASP3 | 836 |
| GAAGGTCTCATTCGGTGTTT | PQBP1 | 10084 |
| GAAGTAAAACCAGCCAACGA | NO_CURRENT_534 | NO_CURRENT_534 |
| GAAGTGGGTGACGGGACGGT | MOSPD1 | 56180 |
| GAAGTGGTGTGTATGTGACT | NO_CURRENT_535 | NO_CURRENT_535 |
| GAAGTTCCAAGGCCCGCGCT | INO80C | 125476 |
| GAATAAATCTGAGTTATACG | NO_CURRENT_536 | NO_CURRENT_536 |
| GAATACTAATAATCATGTAA | NO_CURRENT_537 | NO_CURRENT_537 |
| GAATAGATTTGTCAGTTAGG | NO_CURRENT_538 | NO_CURRENT_538 |
| GAATCGAAGTTCAAGTCCCG | RND1 | 27289 |
| GAATCGACCGACACTAATGT | NO_CURRENT_539 | NO_CURRENT_539 |
| GAATCGGGCCGCCGCCGCCA | CSNK1D | 1453 |
| GAATCTTAACGCGGGCAGCG | IKBKB | 3551 |
| GAATGAAAGGAAACAACCTC | PHF6 | 84295 |
| GAATGAGGCTGAGTTCTCTG | NOS2 | 4843 |
| GAATGGACGTTACAAAGACA | NO_CURRENT_540 | NO_CURRENT_540 |
| GAATTCTAGTAAATTCAGGG | NO_CURRENT_541 | NO_CURRENT_541 |
| GAATTGCCTACCACCGCCAC | PHF8 | 23133 |
| GACAAATTGGGCCATGGTTC | ATP5L | 10632 |
| GACAAGCTGTTCCGCGCTCC | VDR | 7421 |
| GACAATCATGGTGAAAGCGG | NO_CURRENT_542 | NO_CURRENT_542 |
| GACACCACCAAGCCGCAGGG | NIPBL | 25836 |
| GACACCCGGTGGGACTCAGA | EZH2 | 2146 |
| GACACCTCTCGAATAAGGCG | METTL3 | 56339 |
| GACAGACAGCCAGAGAGAGC | AHDC1 | 27245 |
| GACAGAGTCCAGCGGAGTTG | DNTTIP1 | 116092 |
| GACAGATCACGTATCAGGGC | NO_CURRENT_543 | NO_CURRENT_543 |
| GACAGATTATTTACCAGTGA | NO_CURRENT_544 | NO_CURRENT_544 |

| | | |
|---|---|---|
| GACAGCCAGAGAGAGCGGGA | AHDC1 | 27245 |
| GACAGCCTGGACAGCAACTC | IRF9 | 10379 |
| GACAGCCTGTGAACAAGACG | NO_CURRENT_545 | NO_CURRENT_545 |
| GACAGGAGGATCCGGTGCAG | BRK1 | 55845 |
| GACAGGGGCGATGCTTCAGG | KMT2D | 8085 |
| GACAGTGAAATTAGCTCCCA | NO_CURRENT_546 | NO_CURRENT_546 |
| GACAGTGGCTTTGCCCGTTG | MSL2 | 55167 |
| GACATATCTGAAATTAGTCG | NO_CURRENT_547 | NO_CURRENT_547 |
| GACATCACCCAGGAGTCTGC | CYP4A11 | 1579 |
| GACATGATGGCCACTGAGAG | NCKAP1L | 3071 |
| GACATTCAGCCGGCGGTTCG | ATP5L | 10632 |
| GACATTGGTCAGATAGCCAG | NO_CURRENT_548 | NO_CURRENT_548 |
| GACCACTCGGGGTCTGGTGT | MPC1 | 51660 |
| GACCACTCTAGCACCAGTCA | NO_CURRENT_549 | NO_CURRENT_549 |
| GACCATAGACCCAGCCCGTC | CCNC | 892 |
| GACCCACCTGCCAGGCTGCG | SLC6A4 | 6532 |
| GACCCACGGAACTACAGGTG | FAM214B | 80256 |
| GACCCCCGATAACTTTTGAC | NO_CURRENT_550 | NO_CURRENT_550 |
| GACCCGAGTGCTTCCCGCAG | GNAI2 | 2771 |
| GACCGAGCGCGGCAGGCGGC | AKT1 | 207 |
| GACCGCGGTGGTGGCGCCGG | LGMN | 5641 |
| GACCGCTCCTGGGCCAGGCC | ASH2L | 9070 |
| GACCGTCGCTCTCGCCGCCG | MKL1 | 57591 |
| GACCGTTTCATAGTAGATGT | NO_CURRENT_551 | NO_CURRENT_551 |
| GACCTAGGGGCATGATTTCA | LCN2 | 3934 |
| GACCTGTAGGATCCCAAAGA | GPR119 | 139760 |
| GACCTTATAAATGAGTCACA | NO_CURRENT_552 | NO_CURRENT_552 |
| GACCTTCATTGAAGAAAAGC | NO_CURRENT_553 | NO_CURRENT_553 |
| GACGACGACCGTTACCCCAA | MSL2 | 55167 |
| GACGACGACGACGACGGCGG | NUP153 | 9972 |
| GACGACTTCCAAGGCAGCCC | NMRK2 | 27231 |
| GACGAGGGCGGCAGAGCAGT | NO_CURRENT_554 | NO_CURRENT_554 |
| GACGATCTTAGGTGGGTTCC | HSPA13 | 6782 |
| GACGCAGACCGACGCGCAAG | SATB2 | 23314 |
| GACGCCCTAATGCCCATCGT | NO_CURRENT_555 | NO_CURRENT_555 |
| GACGCCGGAGGAGGTGGCGG | GLMN | 11146 |
| GACGCCTTGCCCGGCTCACA | NO_CURRENT_556 | NO_CURRENT_556 |
| GACGCGATGGAACACTCTGG | CYP27B1 | 1594 |
| GACGCGTCTGCTGAGCCCCA | CXCL3 | 2921 |
| GACGCTTCTTCCCGGCGGGT | IFNAR2 | 3455 |
| GACGGACTCTGGAGTTTGGG | PPID | 5481 |
| GACGGCAACGCCGCTGCTCT | DLAT | 1737 |
| GACGGCGGCAGGACGGCGGC | IRF8 | 3394 |
| GACGGCGGCGGGGGAGTCTC | MSL2 | 55167 |
| GACGGGCGGAGCCTCAGGGC | TRIR | 79002 |
| GACGGGGGCGACGCGGCTGA | NDUFA8 | 4702 |
| GACGGTCGCAGGGAGGTCCG | ARRB1 | 408 |
| GACGGTGACGAAGACGGCGG | IKZF5 | 64376 |
| GACGTAGCCTTCCGAAATAT | NO_CURRENT_557 | NO_CURRENT_557 |
| GACGTCCTCTCGCGGCCCTC | NNT | 23530 |

| | | |
|---|---|---|
| GACGTCGCGCTGGCAGACCC | NNT | 23530 |
| GACGTGCTCTGCCAGCCAGT | TIFA | 92610 |
| GACTACAGAGAAGGGTAATC | SMPD1 | 6609 |
| GACTACTGCCAACCCCGGTG | ARNT | 405 |
| GACTATGGAACGTCAGGTGA | APEH | 327 |
| GACTCACATCCATTGGACAC | NO_CURRENT_558 | NO_CURRENT_558 |
| GACTCCAAAATGGCGTCAGT | ATP5J2 | 9551 |
| GACTCCAAAATGGCGTCAGT | ATP5J2-PTCD1 | 100526740 |
| GACTCCAAAATGGCGTCAGT | PTCD1 | 26024 |
| GACTCCAGGGGACCCACAGC | RIPK1 | 8737 |
| GACTCCATTAGGCAGCAGCT | WDR26 | 80232 |
| GACTCCCGGAGCTAGGGGTG | CFLAR | 8837 |
| GACTCCCTGGCTGAGGCTGC | NUFIP2 | 57532 |
| GACTCTCACCTCGGCAACAC | MOSPD1 | 56180 |
| GACTCTCAGCTCCATGGCGG | CSNK1D | 1453 |
| GACTCTCCATTCCAGAACCA | ATP5L | 10632 |
| GACTCTTCCGGTGGAGACAA | KMT2D | 8085 |
| GACTGATGTCGTAACTAGGG | PHC3 | 80012 |
| GACTGCTCGGCGGCGGCCCG | FAM107B | 83641 |
| GACTGCTGCGCTGGTGGTGG | DNAJB6 | 10049 |
| GACTGGTAACTTTGGGGGCA | RNF25 | 64320 |
| GACTGTGACGGTGACGAAGA | IKZF5 | 64376 |
| GACTTAGGACGGGGCGATGG | IFNAR1 | 3454 |
| GACTTCTTGAATGATGCACA | NO_CURRENT_559 | NO_CURRENT_559 |
| GACTTGAACTTCGATTCAGA | RND1 | 27289 |
| GACTTTGGTTGAGCTTCAAT | NO_CURRENT_560 | NO_CURRENT_560 |
| GAGAAAGCGGAGCTACGCTA | NO_CURRENT_561 | NO_CURRENT_561 |
| GAGAACCGACGGAGGACCGC | RHOA | 387 |
| GAGAACGCCGAGCAGCCGCC | TLR2 | 7097 |
| GAGAAGCGGGAGAGCTGGCG | CXCL3 | 2921 |
| GAGAAGGATGGAAATTAGAA | NO_CURRENT_562 | NO_CURRENT_562 |
| GAGAAGGCGCTGGGCTGCGA | TNFRSF1B | 7133 |
| GAGAAGGGAGCGGGACCGGA | MAPK9 | 5601 |
| GAGAAGGGCTGCGGGTTAGG | NFRKB | 4798 |
| GAGAAGTGGGGAGCCATTGG | NO_CURRENT_563 | NO_CURRENT_563 |
| GAGACACAAGAAGCGGATCC | SEC62 | 7095 |
| GAGACAGAGGCTGTTAGCTA | CASP5 | 838 |
| GAGACCACTTTCGTGCAAGC | NO_CURRENT_564 | NO_CURRENT_564 |
| GAGACCCGGCCCCCAAGGGG | CDKN2D | 1032 |
| GAGACTGATTAAATCAACCA | NO_CURRENT_565 | NO_CURRENT_565 |
| GAGAGACACCAGGCGAGGCG | PCGF6 | 84108 |
| GAGAGACAGTCTGAAAGCAG | SLC7A11 | 23657 |
| GAGAGAGGCGGTAAGCCGCG | TFPT | 29844 |
| GAGAGATGACGATCTTAGGT | HSPA13 | 6782 |
| GAGAGCCACTGATTCAGGCC | NO_CURRENT_566 | NO_CURRENT_566 |
| GAGAGCGGCCAGGCCAGCCT | JUNB | 3726 |
| GAGAGGCAGTTGGAAGATGG | SIRT1 | 23411 |
| GAGAGGGATGGTCTCTGCAC | SPEN | 23013 |
| GAGAGTCAGAGTCAGACATT | BCL2L11 | 10018 |
| GAGAGTGCGCCTTGATAGTA | NO_CURRENT_567 | NO_CURRENT_567 |

| | | |
|---|---|---|
| GAGAGTGTGCGGCTCCAGGT | CHEK2 | 11200 |
| GAGATAGCACCATGGTAGAG | AGER | 177 |
| GAGATGAGACCACATGTTAG | NO_CURRENT_568 | NO_CURRENT_568 |
| GAGATTCCAAAACAATGGCA | NO_CURRENT_569 | NO_CURRENT_569 |
| GAGATTTGGGAGTCTGCGCT | COX16 | 51241 |
| GAGCACCTCTGGGGCTATGG | INO80B | 83444 |
| GAGCACGGTTCCCATAAGGG | TBXAS1 | 6916 |
| GAGCAGCAGCCTCTGAGGTG | ARF1 | 375 |
| GAGCAGCTGTCAGGTCTTGT | NO_CURRENT_570 | NO_CURRENT_570 |
| GAGCAGGATGCAGGGCCGCG | C6orf15 | 29113 |
| GAGCATAGCCATGGTTCTCT | CYP8B1 | 1582 |
| GAGCATCCAACATGGCGCTG | EIF4G3 | 8672 |
| GAGCCAACTCACCTAAGGAA | PPARG | 5468 |
| GAGCCCCGACGACGACGCCG | USP7 | 7874 |
| GAGCCCCTTCTCTCGCAAAA | NO_CURRENT_571 | NO_CURRENT_571 |
| GAGCCGAGCTGATTTGATCG | CCNC | 892 |
| GAGCCGGTCAGAGGGTGAGT | UBE2H | 7328 |
| GAGCCTGGAGAGAAGGCGCT | TNFRSF1B | 7133 |
| GAGCGAGAGGTGGATCGGGG | UCHL5 | 51377 |
| GAGCGATCCCTGCATCCTAC | SCN5A | 6331 |
| GAGCGCAGCCCTGAGGCCCA | EP400 | 57634 |
| GAGCGCAGCTCCCTGCCACG | C6orf15 | 29113 |
| GAGCGCCGCGCTCTTCAGCC | CEBPD | 1052 |
| GAGCGCCGGGAGCCTCGGCG | ALOX5 | 240 |
| GAGCGCCTAAGTCCTTCCAG | HSPA4 | 3308 |
| GAGCGCGGCCGGAGGTCCTC | TYK2 | 7297 |
| GAGCGGGAGCAGAGGAGGCG | SIRT1 | 23411 |
| GAGCGGGCCGGCCCCAGCTT | PDE5A | 8654 |
| GAGCGGGCTGGGCGGCTGTC | SUPT7L | 9913 |
| GAGCGGGGCTCTGTCGCCGG | PHF6 | 84295 |
| GAGCGTGAGCTGCTGAGATT | COX16 | 51241 |
| GAGCTAACCGAGTGCGGCGA | GPT2 | 84706 |
| GAGCTCGGGCTGGGCTCCGC | HTR7 | 3363 |
| GAGCTCGTAGCAGTTTGCTA | NO_CURRENT_572 | NO_CURRENT_572 |
| GAGCTGAGCCTAAGCCCTGG | KDM4A | 9682 |
| GAGCTGGGGGAAACGACGCC | JUNB | 3726 |
| GAGCTGTTGCACATTGTGGG | NO_CURRENT_573 | NO_CURRENT_573 |
| GAGCTTTCGGCACCTCTGCC | HDAC2 | 3066 |
| GAGGAAACGGGGACCTGCCC | LTB4R | 1241 |
| GAGGAAGCGATGCAGAGGGG | ABI1 | 10006 |
| GAGGACCTTAAGGTGACATG | NO_CURRENT_574 | NO_CURRENT_574 |
| GAGGAGAGAAAAGCCATGGC | CARD16 | 114769 |
| GAGGAGAGAAAAGCCATGGC | CARD17 | 440068 |
| GAGGAGAGAAAAGCCATGGC | CASP1 | 834 |
| GAGGAGCGCGGTTACCGGAC | CCNC | 892 |
| GAGGAGGACTGGTAACTTTG | RNF25 | 64320 |
| GAGGAGGTCGAGGGTCCCGG | JAM2 | 58494 |
| GAGGATCCGAACCCAGGGGT | HSP90B1 | 7184 |
| GAGGCAAGCGGCCCAGATCT | HAMP | 57817 |
| GAGGCCCGAGAGTCAGAACC | SPEN | 23013 |

| | | |
|---|---|---|
| GAGGCCGCGCGGACTGGCGG | MAP2K5 | 5607 |
| GAGGCCGGGGAGAGCGTTCT | SLC3A2 | 6520 |
| GAGGCGCGCGGTGCCAACCA | PDE5A | 8654 |
| GAGGCGGACGCGAGTCGTCG | MPC2 | 25874 |
| GAGGCGGACTGGGGAGGCGG | RRAGC | 64121 |
| GAGGCGGAGTTAAGCCGAGT | SLC7A3 | 84889 |
| GAGGCGGCTCCTGCGATCGA | HSP90B1 | 7184 |
| GAGGCTAAAGAAACAAGCAT | NO_CURRENT_575 | NO_CURRENT_575 |
| GAGGCTCGCGCGCCCGCGAA | TP73 | 7161 |
| GAGGCTGAGTTCTCTGCGGC | NOS2 | 4843 |
| GAGGCTTCGCCGCCTAGGTA | RBM6 | 10180 |
| GAGGGACTGGAGCGCCCCGC | SPHK1 | 8877 |
| GAGGGAGGAAAGTAGGGGAT | STRAP | 11171 |
| GAGGGAGGAGGGAATGCGGG | UBE2L6 | 9246 |
| GAGGGATGGTCTCTGCACGG | SPEN | 23013 |
| GAGGGCAACTGAAGAAATTG | ACOD1 | 730249 |
| GAGGGCGAGTAAAATTTAAG | SERPINB2 | 5055 |
| GAGGGCGGCGAGAAGGACGG | SLC25A19 | 60386 |
| GAGGGCTCGGCGATCAGCGG | C16orf72 | 29035 |
| GAGGGCTTTGGAATATTACC | ARR3 | 407 |
| GAGGGGCCTCCGGGGCTGCA | CACNA2D2 | 9254 |
| GAGGGGCTCGGCTGCACCGG | STAT1 | 6772 |
| GAGGGGGAGAGGAGGGTATG | CYP4F11 | 57834 |
| GAGGGGGCGCCGGGAAGGCT | HDAC2 | 3066 |
| GAGGGGGCGCTGAGCTGTTG | CSDE1 | 7812 |
| GAGGGTAAAGAGAAAGAAGT | PHF6 | 84295 |
| GAGGGTAGATGAGATGCTGC | CYP24A1 | 1591 |
| GAGGTACTGTACAAGCCCTA | NO_CURRENT_576 | NO_CURRENT_576 |
| GAGGTCAAGTATTACGCGGT | PAXIP1 | 22976 |
| GAGGTCGCGGCGGCTGCGTG | UTY | 7404 |
| GAGGTGAAGGTCTACACATC | IL10 | 3586 |
| GAGGTGAGGGCGAGGGGCGC | ARF1 | 375 |
| GAGTAATTTCGAACGTATTG | NO_CURRENT_577 | NO_CURRENT_577 |
| GAGTAGAGATCCTCTGGACT | GBP7 | 388646 |
| GAGTCAGAACCTGGGGGAGA | SPEN | 23013 |
| GAGTCAGAGTCAGACATTTG | BCL2L11 | 10018 |
| GAGTCAGGCTTAAAACTTGT | RND3 | 390 |
| GAGTCCAAACAGCCTTACCT | ATP5J2 | 9551 |
| GAGTCCAAACAGCCTTACCT | ATP5J2-PTCD1 | 100526740 |
| GAGTCCAAACAGCCTTACCT | PTCD1 | 26024 |
| GAGTCCACGACTGCGAGACA | TMEM70 | 54968 |
| GAGTCCCCACCCCTAGCTCC | CFLAR | 8837 |
| GAGTCGAAGATGGTCTAGGA | NO_CURRENT_578 | NO_CURRENT_578 |
| GAGTCGAGGGAGCTCGGGCT | HTR7 | 3363 |
| GAGTCGGCCTCGCAGCTCTG | CYP2R1 | 120227 |
| GAGTCGGCGGCTGATTTAGA | BABAM1 | 29086 |
| GAGTCTCGCTGCTCGGTGCG | KPNA1 | 3836 |
| GAGTCTGCGCTAGGCCCGCT | COX16 | 51241 |
| GAGTCTGGTAGCTGAGCGTT | LTA4H | 4048 |
| GAGTGAATGGAGGGCGTCGC | SLC25A6 | 293 |

| | | |
|---|---|---|
| GAGTGATGCTTAGACTCCGT | NO_CURRENT_579 | NO_CURRENT_579 |
| GAGTGCGGCGAGGGCCTACC | GPT2 | 84706 |
| GAGTGCTATAAATTAGTCTG | NO_CURRENT_580 | NO_CURRENT_580 |
| GAGTGCTTCCCGCAGAGGGC | GNAI2 | 2771 |
| GAGTGGTACCCAACGGGCCG | PARK7 | 11315 |
| GAGTTATTTATTCTCTCGAG | NO_CURRENT_581 | NO_CURRENT_581 |
| GAGTTCTTGTAACCATACAG | NO_CURRENT_582 | NO_CURRENT_582 |
| GAGTTGACATCTGTTATTGA | NO_CURRENT_583 | NO_CURRENT_583 |
| GAGTTGCGGACTCGGTATCT | BTN2A2 | 10385 |
| GAGTTGTTCGTTTCTCCCCG | IRGM | 345611 |
| GAGTTTGGAGTTGCTTGCGG | TRERF1 | 55809 |
| GATAAATTTGTACGAACTCA | NO_CURRENT_584 | NO_CURRENT_584 |
| GATACAGTTGAATTACACTC | NO_CURRENT_585 | NO_CURRENT_585 |
| GATAGCCTCGGAGACCGCCG | NDUFA12 | 55967 |
| GATAGGAGCCAGGCTGGCAG | CYP4F11 | 57834 |
| GATATCCCGCGAAAAAATCT | NO_CURRENT_586 | NO_CURRENT_586 |
| GATCACCCTCTTCGTCGCTT | HIF1A | 3091 |
| GATCCACTTTAACTGAATAG | NO_CURRENT_587 | NO_CURRENT_587 |
| GATCCAGCAATATTTCTTAA | NO_CURRENT_588 | NO_CURRENT_588 |
| GATCCAGGAGTGATCGAGTA | NO_CURRENT_589 | NO_CURRENT_589 |
| GATCCGAACCCAGGGGTGGG | HSP90B1 | 7184 |
| GATCCTCAACCAGGGAGCTG | SERPINA2 | 390502 |
| GATCGGCCAGTGGCGACAGC | KDM4A | 9682 |
| GATCTAGTCCTCTAATCGAT | NO_CURRENT_590 | NO_CURRENT_590 |
| GATCTGGGTGCAAAAGCCCA | LAMTOR2 | 28956 |
| GATCTGGTTATATAACGATG | NO_CURRENT_591 | NO_CURRENT_591 |
| GATCTTAGGTGGGTTCCCGG | HSPA13 | 6782 |
| GATGAAGACAGACCAGGAGC | C6orf15 | 29113 |
| GATGACGAGGAGAGAAGGCC | TMEM173 | 340061 |
| GATGCAATGGAAGGTGTGCT | NLRP12 | 91662 |
| GATGCAGGGATCGCTCCCCC | SCN5A | 6331 |
| GATGCGGGCCCAAGCTGCAT | RAD21 | 5885 |
| GATGGCCACGGTCCCAGCAC | IL2RB | 3560 |
| GATGGCGCGCAGTTGAGTCA | NO_CURRENT_592 | NO_CURRENT_592 |
| GATGGCGGCGGTGTCTGGCT | PDHB | 5162 |
| GATGGCTCTTGCTCCCAAAT | EDA2R | 60401 |
| GATGGGGAGAAGCGGAGAAC | STRAP | 11171 |
| GATGTCGTCACGTTAACATG | NO_CURRENT_593 | NO_CURRENT_593 |
| GATGTGAAGGACTCCGGGTG | MGA | 23269 |
| GATGTGATCTATGGTTGCGA | NO_CURRENT_594 | NO_CURRENT_594 |
| GATGTTAGATACTGCTCTGA | NO_CURRENT_595 | NO_CURRENT_595 |
| GATGTTTGGAGTTGCGGACT | BTN2A2 | 10385 |
| GATTATTATAATTAACGGCG | POU2F1 | 5451 |
| GATTCATACTAAACACTCTA | NO_CURRENT_596 | NO_CURRENT_596 |
| GATTCGCTGGAAGCAGCTGG | SLC5A8 | 160728 |
| GATTTCAGGGCCGAGGAAGC | LCN2 | 3934 |
| GATTTCCATGCCATGTCTAT | SERPINB2 | 5055 |
| GCAAAAATGTTCCTCAGACG | ZNF699 | 374879 |
| GCAAAACAAAAGGGAAACCC | FOXO4 | 4303 |
| GCAAAAGTGGCATAAAACCG | NO_CURRENT_597 | NO_CURRENT_597 |

| | | |
|---|---|---|
| GCAAACCCGAGTGACACGTC | NO_CURRENT_598 | NO_CURRENT_598 |
| GCAAAGCCGAGCTCTGGGCA | SEC62 | 7095 |
| GCAAAGTGCACGTTTAATCC | ZNF616 | 90317 |
| GCAACAAAACTCCACAGTAA | FBXW7 | 55294 |
| GCAACGCTGCCTGCTCGCCT | AHCTF1 | 25909 |
| GCAACTGAAGAAATTGAGGA | ACOD1 | 730249 |
| GCAAGAGCAACAAGGACCCC | HSPA13 | 6782 |
| GCAATAGAGTTGGTACCCAG | ARMCX2 | 9823 |
| GCAATGCAATCGCAGGAGCA | NO_CURRENT_599 | NO_CURRENT_599 |
| GCAATTTGGAGCTAACTCCA | NO_CURRENT_600 | NO_CURRENT_600 |
| GCACAACGCTTGCCATGTAG | NO_CURRENT_601 | NO_CURRENT_601 |
| GCACACCTTGCAAAACAGAC | MAP2K6 | 5608 |
| GCACACTGCTCGCTGGGCCG | SLC7A5 | 8140 |
| GCACAGCAAACCGCACGCTA | CAT | 847 |
| GCACCAAGGGGTCGCACTGC | IZUMO2 | 126123 |
| GCACCAGAGGCCGCGCCGAC | CS | 1431 |
| GCACCAGGAGCACCAGGACC | MMP9 | 4318 |
| GCACCAGGCGTATTGATGCA | NO_CURRENT_602 | NO_CURRENT_602 |
| GCACCCCGCGCCTCTTCCTG | PARK7 | 11315 |
| GCACCCGCGTCGTGCCTCCT | PDHA1 | 5160 |
| GCACCCTACCCGGGCTGCCT | NMRK2 | 27231 |
| GCACCTCCGCCAGCTCCTAG | LRP2 | 4036 |
| GCACGAGGTGAACAGCCGCT | NO_CURRENT_603 | NO_CURRENT_603 |
| GCACGCTGTACAGACGACAA | NO_CURRENT_604 | NO_CURRENT_604 |
| GCACGTTTAATCCAGGCAGA | ZNF616 | 90317 |
| GCACTAGCAGCATGTTGAGC | SOD2 | 6648 |
| GCACTCAGAGAAGCTGCCCT | TMEM173 | 340061 |
| GCACTGACATCAAAGCAGCC | CCL20 | 6364 |
| GCACTGGGTCCGGTGCGGCA | ACTR5 | 79913 |
| GCACTTTGTTTGGCCTACTG | NO_CURRENT_605 | NO_CURRENT_605 |
| GCAGACCCCAGACCGAGCAG | STAT1 | 6772 |
| GCAGACGTTGTGGCGGAGAA | NDUFS4 | 4724 |
| GCAGACTGGACAGCAGCTAC | TLR9 | 54106 |
| GCAGAGCACGGTTCCCATAA | TBXAS1 | 6916 |
| GCAGAGGCAGGGACCACTCG | MPC1 | 51660 |
| GCAGAGGGCCTTACCCGAGG | NELFE | 7936 |
| GCAGCAACAGTGCGGCCAGG | PTGIS | 5740 |
| GCAGCAGCAACAGTGCGGCC | PTGIS | 5740 |
| GCAGCAGCAGGTACCGGGAG | KMT2C | 58508 |
| GCAGCAGCAGTGGTGGAACG | SLC7A11 | 23657 |
| GCAGCAGGCGTGCAGTTTCC | NDUFB9 | 4715 |
| GCAGCAGGTACCGGGAGAGG | KMT2C | 58508 |
| GCAGCAGTCCTGGAGCTGTG | DNAJB6 | 10049 |
| GCAGCAGTGGTGGAACGAGG | SLC7A11 | 23657 |
| GCAGCCATGGCGGGCGGTCC | MYD88 | 4615 |
| GCAGCCCATGTTAGTGATGG | PHC3 | 80012 |
| GCAGCCCGCAGCCTCAGCCA | NUFIP2 | 57532 |
| GCAGCCGCAGCGGCCAGGCA | ABCB10 | 23456 |
| GCAGCCGCCGCCAGTCCGCG | MAP2K5 | 5607 |
| GCAGCGAGCGCCGGGAGCCT | ALOX5 | 240 |

| | | |
|---|---|---|
| GCAGCGCGACAGCCAGAGGC | PDE3A | 5139 |
| GCAGCGGCGGAGAAGGGAGC | MAPK9 | 5601 |
| GCAGCGGCTGTGGTGGTTCC | CCAR2 | 57805 |
| GCAGCGTGCTCCACTCAGCA | PIGY | 84992 |
| GCAGCGTGCTCCACTCAGCA | PYURF | 100996939 |
| GCAGCTCGGGACTGAGTGCA | ATP5J | 522 |
| GCAGCTGCCGCCTCTGTCCT | WDR26 | 80232 |
| GCAGCTGGCCCAGAGGACAA | EBI3 | 10148 |
| GCAGCTTCCACAGAGCTGCG | CYP2R1 | 120227 |
| GCAGGAATTAGAGCACACTA | GBP7 | 388646 |
| GCAGGAATTCAGCTGCTGGA | SERPINE1 | 5054 |
| GCAGGACAGGAAGCGGGCGG | ZNF699 | 374879 |
| GCAGGACCTGGCCAGGAAGC | ASH2L | 9070 |
| GCAGGAGTCCTGGGGCATGG | DET1 | 55070 |
| GCAGGCAGCCCCAGCCTCCG | IZUMO2 | 126123 |
| GCAGGCCGGTAAGTAACTGG | UQCRB | 7381 |
| GCAGGCGAGCGAGCCACAGC | UBE2H | 7328 |
| GCAGGCGCACCCCGGCGCGC | PDE3A | 5139 |
| GCAGGGACCACGGGGACCGC | APEH | 327 |
| GCAGGGAGCTGCGCTCCTCT | C6orf15 | 29113 |
| GCAGGGCAGCGCAGTCTCCA | RTP1 | 132112 |
| GCAGGGCAGGCAGCTCCAGG | HELZ2 | 85441 |
| GCAGGGCTCTTAAGAACGAA | DPY30 | 84661 |
| GCAGGGCTCTTAAGAACGAA | MEMO1 | 51072 |
| GCAGGTCGGTGCGTCTGTCG | BABAM2 | 9577 |
| GCAGTCGCAGCTGCTTTCCG | MOSPD1 | 56180 |
| GCAGTGGTGGAACGAGGAGG | SLC7A11 | 23657 |
| GCAGTGTGGGCGACAGGACC | NLRP3 | 114548 |
| GCAGTTCCCCCACCACGCGC | STAG2 | 10735 |
| GCATAAATACCCCTTCCGAG | NO_CURRENT_606 | NO_CURRENT_606 |
| GCATCACGGCCTCCATACCT | C6orf15 | 29113 |
| GCATGACGCCTTTCCGCGGC | RB1 | 5925 |
| GCATGTCGGTCAGGCACAGC | TNFRSF6B | 8771 |
| GCATGTTGTGTGAGGATCCC | RAD23A | 5886 |
| GCCAAATCTTGTGTGACATC | HTR1D | 3352 |
| GCCAAATGGAACAGACAAGC | TLR1 | 7096 |
| GCCAAGCAGGCCAGTCAATC | NDUFS8 | 4728 |
| GCCAAGGATGGGAGTCAGCC | IRF9 | 10379 |
| GCCACACGAATCATAAAGAG | NO_CURRENT_607 | NO_CURRENT_607 |
| GCCACCTTTGATGAGGGGAC | NOS2 | 4843 |
| GCCACGGATCGCCGGGTAGT | MAP3K7 | 6885 |
| GCCACGGGGAGGTGTCAAGG | CYP24A1 | 1591 |
| GCCACGGTGACCGTGTAGGA | ALOX5 | 240 |
| GCCACTGCCTGTGCTTCATG | PDHA1 | 5160 |
| GCCAGACATGGCCCAGACCA | APIP | 51074 |
| GCCAGAGAGATGACGATCTT | HSPA13 | 6782 |
| GCCAGAGCCGGACACGGCTG | MAP3K7 | 6885 |
| GCCAGATGTCACACAAGATT | HTR1D | 3352 |
| GCCAGCAGCGCGACAGCCAG | PDE3A | 5139 |
| GCCAGCCAGCCGGGCACTCG | MEX3B | 84206 |

| | | |
|---|---|---|
| GCCAGCGCGACGTCCTCTCG | NNT | 23530 |
| GCCAGCTGCCCGCCCTGCAG | EBI3 | 10148 |
| GCCAGGGCAGCGACACCGGG | CCAR2 | 57805 |
| GCCAGGGCTCTGCGCACCGC | SIAH2 | 6478 |
| GCCAGGGGCGGGCGCCGCCA | CHMP6 | 79643 |
| GCCAGGGTATGGGCATCTCG | NO_CURRENT_608 | NO_CURRENT_608 |
| GCCAGGGTTCTTGGTCCCGA | NO_CURRENT_609 | NO_CURRENT_609 |
| GCCAGGTCTACCTGAGCCTA | PRR14L | 253143 |
| GCCAGTGCAAACAGGAACCA | UBA7 | 7318 |
| GCCATAATATATGAGTAACT | NO_CURRENT_610 | NO_CURRENT_610 |
| GCCATCGCCTTCCTCCCCAG | SPINT1 | 6692 |
| GCCATGCTCTGCGCACCGGC | SLC7A5 | 8140 |
| GCCATGGTCCGGGGACGCCG | NOD1 | 10392 |
| GCCATGTCGAGGGTTGCTGA | JAM3 | 83700 |
| GCCATTCTAGTCCCGGCATA | NO_CURRENT_611 | NO_CURRENT_611 |
| GCCCAAAACCCCCCGAAAAA | RB1 | 5925 |
| GCCCAAGGAGGAGAAGCTCA | CYP4B1 | 1580 |
| GCCCAAGTGTCCAACAACAA | NO_CURRENT_612 | NO_CURRENT_612 |
| GCCCACAACTGAAAGGTCTG | PAGR1 | 79447 |
| GCCCAGAAGGTAGGCCTTCA | SQSTM1 | 8878 |
| GCCCAGACCAGGGACCCGCG | APIP | 51074 |
| GCCCAGACGCCCTAGAATAG | NO_CURRENT_613 | NO_CURRENT_613 |
| GCCCAGCAGTTCCTCCACGC | ACACA | 31 |
| GCCCAGGCCCGCGCCCGCCG | IRAK1 | 3654 |
| GCCCATGGAAGTTTAAAATG | NO_CURRENT_614 | NO_CURRENT_614 |
| GCCCCACACCTGTAGTTCCG | FAM214B | 80256 |
| GCCCCATCATCTCCAGCGCG | STAG2 | 10735 |
| GCCCCATGGAGCGGCCCCCG | CHUK | 1147 |
| GCCCCCCGAACCCCACCTTG | HNRNPF | 3185 |
| GCCCCCGCCAGGCTCCCGAG | NDUFC1 | 4717 |
| GCCCCCGCCGCCCGCCCGTC | BAHD1 | 22893 |
| GCCCCGGCGTCCCCGGACCA | NOD1 | 10392 |
| GCCCCGGGACTGCTCCTCCT | SCTR | 6344 |
| GCCCCGGGAGCGGAGAGCGA | CTNNB1 | 1499 |
| GCCCCGGGCTCACCTGCGCG | TIRAP | 114609 |
| GCCCCGGGCTGCTGGCTCCC | NLRP1 | 22861 |
| GCCCCGGGGGCCGCTCCATG | CHUK | 1147 |
| GCCCCGTAAATCTCATTACA | NO_CURRENT_615 | NO_CURRENT_615 |
| GCCCCTACAAGGCCGACTCC | MAPK14 | 1432 |
| GCCCCTCCGCAGGCGGACCG | TNFRSF6B | 8771 |
| GCCCCTCGCCCTCACCTCAG | ARF1 | 375 |
| GCCCCTGGTCCTGGTGCTCC | MMP9 | 4318 |
| GCCCGAAAATGTCTCAACCC | LARP4B | 23185 |
| GCCCGAAACCCGGAAGTGAG | SBNO2 | 22904 |
| GCCCGAATGTCGTTAGCCGT | SUV39H1 | 6839 |
| GCCCGAGAGTCAGAACCTGG | SPEN | 23013 |
| GCCCGAGCAGTGTGAAGAAG | HMGN2 | 3151 |
| GCCCGCAGCCCGACCCGGCA | NCOA6 | 23054 |
| GCCCGCCCCTCACCTCCTTT | PDAP1 | 11333 |
| GCCCGCCCTCACCTGAACCG | NFRKB | 4798 |

| | | |
|---|---|---|
| GCCCGCCGGCGGTGGCGCGG | TBK1 | 29110 |
| GCCCGCGCGGCTCGGGCTCC | RPS6KA4 | 8986 |
| GCCCGCGCTGGGAAAAAGGT | INO80C | 125476 |
| GCCCGCGGTTCAGGTGAGGG | NFRKB | 4798 |
| GCCCGCTAAGCGAGCGCACC | SNRNP70 | 6625 |
| GCCCGGAGCCCGAGCCGCGC | RPS6KA4 | 8986 |
| GCCCGGAGCCTTCCCCAGCG | SATB1 | 6304 |
| GCCCGGAGTCGGCCTTGTAG | MAPK14 | 1432 |
| GCCCGGCGCCCGCCCCTTGG | CDKN2D | 1032 |
| GCCCGGGTTCAGGCTCTCAG | FLCN | 201163 |
| GCCCGGGTTCAGGCTCTCAG | PLD6 | 201164 |
| GCCCTCCTCCTACCAGCGCG | NR2C2 | 7182 |
| GCCCTCTGCGGGAAGCACTC | GNAI2 | 2771 |
| GCCCTTGCGGCTCTCACAAC | CYP1B1 | 1545 |
| GCCGAACAGGTTACCCATGG | CHMP6 | 79643 |
| GCCGAGAAGCAGAAACACGA | PRKAA1 | 5562 |
| GCCGAGCCCTCCGCGCAGCT | C16orf72 | 29035 |
| GCCGCAGCGGCCAGGCAGGG | ABCB10 | 23456 |
| GCCGCCACAGCTTACTCATG | INO80B | 83444 |
| GCCGCCACGTCCGAGGCACA | PHF8 | 23133 |
| GCCGCCATGGAGCTGAGAGT | CSNK1D | 1453 |
| GCCGCCATGGGTAACCTGTT | CHMP6 | 79643 |
| GCCGCCCGCGCGCTGGTAGG | NR2C2 | 7182 |
| GCCGCCGATTTCATAAGTAA | NO_CURRENT_616 | NO_CURRENT_616 |
| GCCGCCGCAGTGGGTGTGAG | MAPK9 | 5601 |
| GCCGCCGCCACCGCAGCCGC | RXRA | 6256 |
| GCCGCCGCCCGCGCCCTGAC | RRAGC | 64121 |
| GCCGCCGCCGCCGCTGCTGC | ADRB1 | 153 |
| GCCGCCGCCGCCGCTGCTGC | CDYL2 | 124359 |
| GCCGCCGCCGCCGCTGCTGC | LIN54 | 132660 |
| GCCGCCGCCGCGCGTCGCGG | TRMT61A | 115708 |
| GCCGCCGCCTTCTCCGCGAG | CACNA2D2 | 9254 |
| GCCGCCGGCTCCCCGCCGCC | RXRA | 6256 |
| GCCGCCTGGCTGAGAAAACT | MCL1 | 4170 |
| GCCGCGAGAGGACGTCGCGC | NNT | 23530 |
| GCCGCGCCGACGGGTTGACA | CS | 1431 |
| GCCGCGCGGGTCCCTGGTCT | APIP | 51074 |
| GCCGCGCTCTTCAGCCTGGA | CEBPD | 1052 |
| GCCGCGGCCATGGGCGGGCG | ABL1 | 25 |
| GCCGCGGCTGGAAGATGGCG | PPARGC1B | 133522 |
| GCCGCGGGCCAAGCCTACCG | ZNF641 | 121274 |
| GCCGCGGGTGCAGGAGTCCT | DET1 | 55070 |
| GCCGCGTCTTCCTCAAGGTG | HNRNPF | 3185 |
| GCCGCTCAGGGTGAGTCCCG | YPEL5 | 51646 |
| GCCGCTCTCCGCTGCGGGGG | KAT2A | 2648 |
| GCCGCTGCCAGCGAAGGTGC | BCL2 | 596 |
| GCCGCTGGCAGCGCTGGCCC | NENF | 29937 |
| GCCGGACCAGGTGCGAACCC | VDR | 7421 |
| GCCGGACTGGCTTGCGGGCC | FAM107B | 83641 |
| GCCGGAGCCGCAATGCCTAA | PDAP1 | 11333 |

| | | |
|---|---|---|
| GCCGGAGCGCGGCGACTCCA | RIPK1 | 8737 |
| GCCGGCCTGAGCAGCGCTCT | HSPA4 | 3308 |
| GCCGGCCTTCCTCGTGTGAG | MCTS1 | 28985 |
| GCCGGCGGCGGCAGTGTCTG | MKL1 | 57591 |
| GCCGGCGGGCTGAGGCGTAC | ABCB10 | 23456 |
| GCCGGCTCATCCGCCGCGTC | SATB1 | 6304 |
| GCCGGGACCTCCTGCAGCCA | MYD88 | 4615 |
| GCCGGGAGGAGGGCACCGCG | SIAH2 | 6478 |
| GCCGGGCACCGCGGCCATGG | UCHL3 | 7347 |
| GCCGGGGAGTCCGCTCCAGC | SPHK1 | 8877 |
| GCCGGGGATCCTGGAGCCAT | PYCARD | 29108 |
| GCCGGGGCCGGGAAGCGCCG | NOD1 | 10392 |
| GCCGGGGCCTCTCAGGGCCG | IRAK1 | 3654 |
| GCCGGGGCCTGAACCGGGGA | NO_CURRENT_617 | NO_CURRENT_617 |
| GCCGGTCATGAACGGGCCGG | INO80E | 283899 |
| GCCGTCCAGGCTGAAGAGCG | CEBPD | 1052 |
| GCCGTCCGGCGAGCAGGAGC | GSDMD | 79792 |
| GCCGTGGTATCAAGTCGGTA | NO_CURRENT_618 | NO_CURRENT_618 |
| GCCGTGTGCCTCGGACGTGG | PHF8 | 23133 |
| GCCGTTGCCACTGACAGCCG | RRAGA | 10670 |
| GCCTAAAGGAGGTGAGGGGC | PDAP1 | 11333 |
| GCCTAGATGTTCGTTCTGAG | NO_CURRENT_619 | NO_CURRENT_619 |
| GCCTAGTGAATGCTCCAGCA | HCAR2 | 338442 |
| GCCTATCGGCATTCCCACTG | NO_CURRENT_620 | NO_CURRENT_620 |
| GCCTCAGGGCCGGGAGGCTC | TRIR | 79002 |
| GCCTCCCAGCCTCCAGGCCA | PTGES2 | 80142 |
| GCCTCCCCCGCAGCGGAGAG | KAT2A | 2648 |
| GCCTCCGTCCTAGGTCCCCG | VIPR2 | 7434 |
| GCCTCCTCGGCGTCGTCGTC | USP7 | 7874 |
| GCCTCCTGCGCTCCTGCCGC | JAM2 | 58494 |
| GCCTCGCCCCGGCGGCAGAG | APEH | 327 |
| GCCTCGGAGACCGCCGTGGC | NDUFA12 | 55967 |
| GCCTCTGAGGTGAGGGCGAG | ARF1 | 375 |
| GCCTCTTCTTCACACTGCTC | HMGN2 | 3151 |
| GCCTGGAGGGTGACCTCAGT | GBP5 | 115362 |
| GCCTGGGACGCGCGGAGTCG | HTR7 | 3363 |
| GCCTGGGTTTTGGTGCATAC | NO_CURRENT_621 | NO_CURRENT_621 |
| GCCTGGTGCGCTCGCTTAGC | SNRNP70 | 6625 |
| GCCTTATTCGAGAGGTGTCA | METTL3 | 56339 |
| GCCTTCCCCGGTTCAGGCCC | NO_CURRENT_622 | NO_CURRENT_622 |
| GCCTTCCCTAACGTTGCAAC | MAP2K6 | 5608 |
| GCCTTGGTTCCTGTTTGCAC | UBA7 | 7318 |
| GCGAACGAGCAGGGCGGGAG | IDH2 | 3418 |
| GCGAACGAGCGGCGCTCGGC | FOSL2 | 2355 |
| GCGAACGGGAAGGAGCGTTG | LARP4B | 23185 |
| GCGAACTGTGGAGCTGCCAG | HTR7 | 3363 |
| GCGACAAAATGGCTGCCCGA | TRIR | 79002 |
| GCGACGAGAACGGCGAGCGA | ARHGAP33 | 115703 |
| GCGACGGAATCAGACGGACG | SNRNP70 | 6625 |
| GCGACTCCTCACAACCCAGG | PDHA1 | 5160 |

| | | |
|---|---|---|
| GCGAGAAGCGGGAGAGCTGG | CXCL3 | 2921 |
| GCGAGAGGAAGCGATGCAGA | ABI1 | 10006 |
| GCGAGATCCGGAGTTGGGCT | NFKB2 | 4791 |
| GCGAGCACGACCGATTCTGC | ANKRD50 | 57182 |
| GCGAGCAGCGCGGCGGCCAT | ACADS | 35 |
| GCGAGCAGGCAGCGTTGCAA | AHCTF1 | 25909 |
| GCGAGCCGAGCGCAGTCTGC | PIP4K2A | 5305 |
| GCGAGCGAACGAGCGGCGCT | FOSL2 | 2355 |
| GCGAGCGAGCGACCCGGCCC | BRD1 | 23774 |
| GCGAGCGGCGTAGCGGATCC | BAHD1 | 22893 |
| GCGAGGCCCCAGTGGACACG | ALOX15 | 246 |
| GCGAGGGAGGAGGGCCAGAG | SIRT1 | 23411 |
| GCGAGGGCCTCGGCCATCCA | HSH2D | 84941 |
| GCGAGGGCGACCGAGACTTA | SPCS1 | 28972 |
| GCGAGGGCGCGAGGGCGCGA | LYST | 1130 |
| GCGAGGGCGCGAGGGCGCGA | TNFRSF1B | 7133 |
| GCGAGGGGCGCGGGCCGGTG | ARF1 | 375 |
| GCGAGGGGTCGAGCGCGGCC | ARHGAP33 | 115703 |
| GCGAGTTCTCACGGTCCCGC | LGMN | 5641 |
| GCGATCGGAGTGCCACGATA | NO_CURRENT_623 | NO_CURRENT_623 |
| GCGATTCCGGAGCCCCTGCG | ACACA | 31 |
| GCGCAAAGCCGTGCGGAGAT | APIP | 51074 |
| GCGCAGAGCCGGAGTCGCGG | JAM3 | 83700 |
| GCGCAGCCCTCATCGCAACT | TIRAP | 114609 |
| GCGCAGCTCGGCCTGCACCG | C16orf72 | 29035 |
| GCGCAGGCTCAGCGGCCCCG | SCN5A | 6331 |
| GCGCAGGTCGGTGCGTCTGT | BABAM2 | 9577 |
| GCGCCATGGTCCGGGGACGC | NOD1 | 10392 |
| GCGCCCGCCGCGGCCCTGAG | IRAK1 | 3654 |
| GCGCCCGGGAGGGAGGCACC | ZNF641 | 121274 |
| GCGCCCTGACAGGCCAGGCC | RRAGC | 64121 |
| GCGCCGAGCTGTGCGCAGCC | DNAJA1 | 3301 |
| GCGCCGCCACTTCAGGTCCA | CYP2R1 | 120227 |
| GCGCCGCGCGCAAAGCCGTG | APIP | 51074 |
| GCGCCGCGTCTTCCTCAAGG | HNRNPF | 3185 |
| GCGCCTCGGCGGTGCTCACC | KPNA1 | 3836 |
| GCGCGCAGGTGAGCCCGGGG | TIRAP | 114609 |
| GCGCGCCCACCTCCCGGGGA | LRP2 | 4036 |
| GCGCGCCCCCCCGGCGCCCC | ABL1 | 25 |
| GCGCGCCTGGAGCAGCTACC | GPT2 | 84706 |
| GCGCGCGAGCTAACCGAGTG | GPT2 | 84706 |
| GCGCGCGGCCACCTGGTCCG | SCTR | 6344 |
| GCGCGCGTAGCACACCGCAC | LRP2 | 4036 |
| GCGCGCTCTCCAGTCCTCCC | IL2RB | 3560 |
| GCGCGGAGTCGAGGGAGCTC | HTR7 | 3363 |
| GCGCGGGCATGGCGGGCGGG | SLC25A1 | 6576 |
| GCGCGGGCCCAGTTGCGATG | TIRAP | 114609 |
| GCGCGGGCGGCGGGACAGAC | NCOA6 | 23054 |
| GCGCGGGGCGCGGGCATGGC | SLC25A1 | 6576 |
| GCGCGGGGCGGCGGTGAGGG | ABL1 | 25 |

| | | |
|---|---|---|
| GCGCGGTGCCAACCATGGAG | PDE5A | 8654 |
| GCGCGGTGCCCTCCTCCCGG | SIAH2 | 6478 |
| GCGCGGTTACCGGACGGGCT | CCNC | 892 |
| GCGCGTCGCGGAGGCACGTG | TRMT61A | 115708 |
| GCGCGTTTCTGCGGGCGCAA | RAC2 | 5880 |
| GCGCTCCGGCCGCGTCAGGA | EP400 | 57634 |
| GCGCTCGAGCGGTTCCTGTC | TMEM30A | 55754 |
| GCGCTCTGGCGCCCCAGCTG | RIPK1 | 8737 |
| GCGCTGAGGTGGATTTGTAC | EDA2R | 60401 |
| GCGCTGCGCGCTCACCATGG | NENF | 29937 |
| GCGCTGTGCGCACAGGGGAG | NUP153 | 9972 |
| GCGGAACTTTCCAGAACGCT | DNAJA1 | 3301 |
| GCGGAATGAACACGAGGTAG | NO_CURRENT_624 | NO_CURRENT_624 |
| GCGGACGCGATGGAACACTC | CYP27B1 | 1594 |
| GCGGACTGGAAGCCGATGCC | RAC1 | 5879 |
| GCGGAGATTGGAGGCCGCGC | APIP | 51074 |
| GCGGAGCAGCCAGACAGCGA | TGFB1 | 7040 |
| GCGGAGGGCTCGGCGATCAG | C16orf72 | 29035 |
| GCGGAGGGGCTCGGCTGCAC | STAT1 | 6772 |
| GCGGAGGGTCGGCGTAGGTG | FBXO38 | 81545 |
| GCGGATCTCACCGCCGCTCA | YPEL5 | 51646 |
| GCGGCACAGCCCCGTGGGTC | ARID3A | 1820 |
| GCGGCACGGGCGTCGCGGAA | ACTR5 | 79913 |
| GCGGCAGCGGCCGGGGATCC | PYCARD | 29108 |
| GCGGCAGGACGGCGGCAGGT | IRF8 | 3394 |
| GCGGCCAGCGAAGGTGGCGA | TRAF6 | 7189 |
| GCGGCCAGGAACATCGCGAG | TFPT | 29844 |
| GCGGCCCGGAACGTGGGGAT | MLF2 | 8079 |
| GCGGCCGAACAGGTTACCCA | CHMP6 | 79643 |
| GCGGCCGGAGCGCAGCCCTG | EP400 | 57634 |
| GCGGCCGTCAGCGCTCGGAG | BCL2 | 596 |
| GCGGCGACAAGGCAGTCGCT | SATB2 | 23314 |
| GCGGCGACTACTGCCAACCC | ARNT | 405 |
| GCGGCGACTCCTCACAACCC | PDHA1 | 5160 |
| GCGGCGAGGACAGCCGGGAC | JAK1 | 3716 |
| GCGGCGCAGGTGAGCAGGGC | EIF2AK2 | 5610 |
| GCGGCGCGAGGGCGGCTCCG | FAM107B | 83641 |
| GCGGCGCGCACACTGCTCGC | SLC7A5 | 8140 |
| GCGGCGCTCATGGCGGCGTC | CEBPD | 1052 |
| GCGGCGGCAGCGGCGGAGAA | MAPK9 | 5601 |
| GCGGCGGCAGCTGCGAGGCT | SBNO2 | 22904 |
| GCGGCGGCAGGAGCAGGACC | ASH2L | 9070 |
| GCGGCGGCCGCGGAGTATCC | JAK1 | 3716 |
| GCGGCGGCGCGCGGAGGCCG | PPP2R3A | 5523 |
| GCGGCGGCGCGCGGAGGCCG | RXRA | 6256 |
| GCGGCGGCGGCGGCTCAACG | KHSRP | 8570 |
| GCGGCGGCGGTTGGCCGGTG | ANKRD50 | 57182 |
| GCGGCGGCTGGGAGAGGCGA | GLMN | 11146 |
| GCGGCGGGGGAGGGACCCTG | KPNB1 | 3837 |
| GCGGCGGTGAGGGCGGCTGG | ABL1 | 25 |

| | | |
|---|---|---|
| GCGGCGGTGATGGACGGGTC | BAX | 581 |
| GCGGCGTCTGGGAATCGTTC | NO_CURRENT_625 | NO_CURRENT_625 |
| GCGGCGTGGGAGCACCTCTG | INO80B | 83444 |
| GCGGCTGGTGGAAGAGCTGC | CASP9 | 842 |
| GCGGCTGTCTGGACCTCGAG | SUPT7L | 9913 |
| GCGGCTTCCTAGTGAGTCGG | BABAM1 | 29086 |
| GCGGCTTGGGGGTCCTGGGC | CYFIP1 | 23191 |
| GCGGGACTCACCCTGAGCGG | YPEL5 | 51646 |
| GCGGGAGATTCACCAGGACT | BRK1 | 55845 |
| GCGGGAGCGGCGGTGATGGA | BAX | 581 |
| GCGGGAGGCGGGGCGCCCTG | SLC25A1 | 6576 |
| GCGGGATGTGCTTCAGCTGC | BNIP3 | 664 |
| GCGGGCCAGCTGCCGGAGCG | RIPK1 | 8737 |
| GCGGGCCAGGGCAGCGACAC | CCAR2 | 57805 |
| GCGGGCGCAGCAGCTGGAAC | MAPK14 | 1432 |
| GCGGGCGCAGGATACGGGCC | PIP4K2A | 5305 |
| GCGGGCGCGGGACGCGGGGC | BRD1 | 23774 |
| GCGGGCGCGGGCGCGCGGGG | ABL1 | 25 |
| GCGGGCTTTGGTCGGTCCGG | HMGN2 | 3151 |
| GCGGGGACCTAGGACGGAGG | VIPR2 | 7434 |
| GCGGGGCCCGAGCGACGCGG | RAD23A | 5886 |
| GCGGGGCTGGCTCGGACTCC | BRD1 | 23774 |
| GCGGGTCCCTCGGGCTGCAC | BAK1 | 578 |
| GCGGGTCGCTGGGGCCCGCG | RPS6KA4 | 8986 |
| GCGGTCGAAAGGGGAGTTCA | NDUFA8 | 4702 |
| GCGGTCTGGGCGTGAGTGCA | CTNNBL1 | 56259 |
| GCGGTGCAGGCCGAGCTGCG | C16orf72 | 29035 |
| GCGGTGCCGGCTGCGGCTGG | CASP9 | 842 |
| GCGGTGCTCCTGGCTGGGCG | ITGB2 | 3689 |
| GCGGTGGCGCGGCGGAGACC | TBK1 | 29110 |
| GCGGTTGTGGGGCCCGGGAC | CREBBP | 1387 |
| GCGTAGCGGATCCCGGAGCC | BAHD1 | 22893 |
| GCGTCCGCAGGAAGAGGCGC | PARK7 | 11315 |
| GCGTCGAGCGGGAGCAGAGG | SIRT1 | 23411 |
| GCGTCGCAGAGCGGGCTGGG | SUPT7L | 9913 |
| GCGTCGCAGGTGAGACTCCG | TRMT61A | 115708 |
| GCGTCGCCGGGCCGCTCCGG | NFKB1 | 4790 |
| GCGTGCGTCCCGGGTTACCC | NO_CURRENT_626 | NO_CURRENT_626 |
| GCGTGGAGGAACTGCTGGGC | ACACA | 31 |
| GCGTGGTGGGGGAACTGCTG | STAG2 | 10735 |
| GCGTGTCAGCGAGATAGTTG | PPARGC1B | 133522 |
| GCGTTCAAATAGCCTCACTT | NO_CURRENT_627 | NO_CURRENT_627 |
| GCGTTCCCCCACTGACGGGG | NO_CURRENT_628 | NO_CURRENT_628 |
| GCGTTGACCCTCCATGGCCG | UCHL3 | 7347 |
| GCGTTGGCGGTCTTGGCATG | PIGL | 9487 |
| GCGTTGGGGATCTGGAGTCC | LARP4B | 23185 |
| GCTAACTAAGGTGCAATAAG | HTR3E | 285242 |
| GCTACGTATTTCAAAAAACG | NO_CURRENT_629 | NO_CURRENT_629 |
| GCTAGGCCACGCCGAGGTCC | SOD1 | 6647 |
| GCTATTTGAGTAGTTCACTG | NO_CURRENT_630 | NO_CURRENT_630 |

| | | |
|---|---|---|
| GCTCAACATGCTGCTAGTGC | SOD2 | 6648 |
| GCTCACCGACTCCGGACGCC | NUDT17 | 200035 |
| GCTCACCGAGAGCCTAGTTC | NQO1 | 1728 |
| GCTCACGCGGTGAGTCATAT | CHEK2 | 11200 |
| GCTCAGAGCGCGCTGGTGCT | KPNB1 | 3837 |
| GCTCAGCACAGAGACACTCA | CYP4A11 | 1579 |
| GCTCATGGCGGCGTCGGGCC | CEBPD | 1052 |
| GCTCCAACAAACTGCAGACC | BCL2L11 | 10018 |
| GCTCCAGGAGGGTGAGCGCG | HELZ2 | 85441 |
| GCTCCAGTAGCCACCGCATC | CXCR4 | 7852 |
| GCTCCAGTTCTCATCTGCTC | SLC5A8 | 160728 |
| GCTCCATGAAAGAATAATTG | NO_CURRENT_631 | NO_CURRENT_631 |
| GCTCCCGGCCCCCGACCCAC | ARID3A | 1820 |
| GCTCCCGGGTGTCCCAGGCC | SLC7A5 | 8140 |
| GCTCCTACCTGCAGGGCGAA | PLD1 | 5337 |
| GCTCCTCCGTGCCGCCGCGG | UBE2E1 | 7324 |
| GCTCCTCGATCAAATCAGCT | CCNC | 892 |
| GCTCCTGGGCAGGCTCGGCA | HELZ2 | 85441 |
| GCTCCTGTGTGTGGCGTTGG | PIGL | 9487 |
| GCTCGCAAGTATTTAAGGAC | NO_CURRENT_632 | NO_CURRENT_632 |
| GCTCGGAGAAACAGGCGCCG | SBNO2 | 22904 |
| GCTCGGAGCAGCAGCCTCTG | ARF1 | 375 |
| GCTCGGCGGGGACAGAAAGA | FOSL2 | 2355 |
| GCTCGTTCGCTCTCCAGCTT | IDH2 | 3418 |
| GCTCTCCGCGCGGCCGGGGA | NDUFB9 | 4715 |
| GCTCTGGTTCGGAGAAGCAG | RIPK2 | 8767 |
| GCTCTTAAGAACGAACGGCT | DPY30 | 84661 |
| GCTCTTAAGAACGAACGGCT | MEMO1 | 51072 |
| GCTCTTCAGCCTGGACGGCC | CEBPD | 1052 |
| GCTCTTCCACCAGCCGCAGC | CASP9 | 842 |
| GCTGAAGAGCGCGGCGCTCA | CEBPD | 1052 |
| GCTGACCTTCCCCGGCCGCG | NDUFB9 | 4715 |
| GCTGACGGCCGCCGGCAGGG | BCL2 | 596 |
| GCTGAGAGCGAGAGGTGGAT | UCHL5 | 51377 |
| GCTGAGCCCCAGCAGACTCC | CYP4A11 | 1579 |
| GCTGAGCCTAAGCCCTGGCG | KDM4A | 9682 |
| GCTGAGGGGGTCCGGCCGTT | WASF2 | 10163 |
| GCTGATATATACGACAAGCC | NO_CURRENT_633 | NO_CURRENT_633 |
| GCTGATCTCGCCGCCCTTAT | TBXAS1 | 6916 |
| GCTGATTTAGAAGGAGGTTC | BABAM1 | 29086 |
| GCTGATTTGATCGAGGAGCG | CCNC | 892 |
| GCTGCAAATTCTATTTGTGT | NO_CURRENT_634 | NO_CURRENT_634 |
| GCTGCAGGGGTCGGAGGTCA | SAP18 | 10284 |
| GCTGCCCGCCCTGCAGTGGA | EBI3 | 10148 |
| GCTGCCCGGGAGGCGGCTGG | SETD1B | 23067 |
| GCTGCCTGTGAATGATGCAA | NLRP12 | 91662 |
| GCTGCGAGGCTCGGAGAAAC | SBNO2 | 22904 |
| GCTGCGGCTGCGGCTGGGCC | UBE2E1 | 7324 |
| GCTGCTAGGAGCCGGTCAGA | UBE2H | 7328 |
| GCTGCTCCAATATCCCAGCT | MMP1 | 4312 |

| | | |
|---|---|---|
| GCTGCTCCCGCCGCGCTTCT | AHCTF1 | 25909 |
| GCTGCTCCCGGGTTCGCACC | VDR | 7421 |
| GCTGCTCCGCGGAGGGAGGT | TGFB1 | 7040 |
| GCTGCTCCTGTCCCGGCGTC | NUDT17 | 200035 |
| GCTGCTGCCCGGGAGGCGGC | SETD1B | 23067 |
| GCTGCTGCCTACCTCCCGAA | PDHB | 5162 |
| GCTGCTGCTCCCGGGGCTGC | KMT2C | 58508 |
| GCTGCTGGAGGGGGGCGTGT | SERPINE1 | 5054 |
| GCTGCTTGGAGCAAAACAAA | FOXO4 | 4303 |
| GCTGGAGAGACAATTCTACT | NO_CURRENT_635 | NO_CURRENT_635 |
| GCTGGAGCATTCACTAGGCG | HCAR2 | 338442 |
| GCTGGAGCATTCACTAGGCG | HCAR3 | 8843 |
| GCTGGAGCCGCTGCCGCTGC | ADRB1 | 153 |
| GCTGGAGGCGGCGGAACCGC | VIPR2 | 7434 |
| GCTGGCACCTCTATGATCAC | BAK1 | 578 |
| GCTGGCAGGTCGCCATCTTT | GPR119 | 139760 |
| GCTGGCGCCGCTGCGCGCAT | PRKN | 5071 |
| GCTGGCTCGGACTCCAGGGC | BRD1 | 23774 |
| GCTGGCTGGCTTGTGCGCCC | PDK4 | 5166 |
| GCTGGGAGAGGCGAGGGTTC | GLMN | 11146 |
| GCTGGGAGCCGAGGCAGCCG | SATB2 | 23314 |
| GCTGGGAGCTGGGAGCGCGC | KIAA1211L | 343990 |
| GCTGGGCTGCGAGGGCGCGA | TNFRSF1B | 7133 |
| GCTGGGGCCGGCCCGCTCCA | PDE5A | 8654 |
| GCTGGGGCTTGGCCTGCGGG | TRAF6 | 7189 |
| GCTGGGGGCGTAGCGCGCTG | WASF2 | 10163 |
| GCTGGGGTTTGAGGGTGCCG | PDK4 | 5166 |
| GCTGGTGGAAGAGCTGCAGG | CASP9 | 842 |
| GCTGGTTTGTAATGATAGGG | SLC7A11 | 23657 |
| GCTGTCTGGCTGCTCCGCGG | TGFB1 | 7040 |
| GCTGTGAATCGTGGCTGGCC | STRAP | 11171 |
| GCTGTGCAGAACACTGCAGG | CAT | 847 |
| GCTGTGCTGCAGCTACATGA | MAP2K6 | 5608 |
| GCTGTGGTGGCTGCTGGGCC | LCN2 | 3934 |
| GCTTAACTAGCAGAGGACTC | AIM2 | 9447 |
| GCTTAAGTCACGGCTTTCCA | NO_CURRENT_636 | NO_CURRENT_636 |
| GCTTACACGATTGTGAAGTC | NO_CURRENT_637 | NO_CURRENT_637 |
| GCTTAGCGCGCGCGGCCACC | SCTR | 6344 |
| GCTTATGGCGGCGCTGGAGA | CSDE1 | 7812 |
| GCTTCAGGTGGTGGGGATAG | KMT2D | 8085 |
| GCTTCCAGCGAATCTACACA | SLC5A8 | 160728 |
| GCTTCCTTCCAGTAAGGAGT | MCL1 | 4170 |
| GCTTCTCCAGAGGGTCTGGC | TLR9 | 54106 |
| GCTTCTCTGCTAATTTATGC | NCKAP1L | 3071 |
| GCTTCTGATAGGAGCCAGGC | CYP4F11 | 57834 |
| GCTTGACTGAGAAGAACTCG | FBXW7 | 55294 |
| GCTTGCATAATTCTGGCCAG | NO_CURRENT_638 | NO_CURRENT_638 |
| GCTTGCGGCGGAGGGCAGGA | TRERF1 | 55809 |
| GCTTGCTATATGGGTGCGAG | NO_CURRENT_639 | NO_CURRENT_639 |
| GCTTGGTGCGGAGACCCCTT | PDHB | 5162 |

| | | |
|---|---|---|
| GCTTGTACCTCCTGCGCGCC | PDE3A | 5139 |
| GCTTGTAGACAAAACAACGT | NO_CURRENT_640 | NO_CURRENT_640 |
| GCTTTTTCCAGCGAGAGCAA | NO_CURRENT_641 | NO_CURRENT_641 |
| GGAAAAAGGTGGGGGGACCA | INO80C | 125476 |
| GGAAACTTGAGTTTACCGTG | NO_CURRENT_642 | NO_CURRENT_642 |
| GGAAAGAAATTGAATTTGCA | MCTS1 | 28985 |
| GGAAAGGGGACGCAGCAAGG | NDUFC1 | 4717 |
| GGAAAGTAGACTCACCGTCT | RIPK3 | 11035 |
| GGAAATGACAGTATGAGTCG | NO_CURRENT_643 | NO_CURRENT_643 |
| GGAACAGCTTGTCCACCCGC | VDR | 7421 |
| GGAACCAATTCTCCTGTCGG | NIPBL | 25836 |
| GGAACCCAATCCGAGGGTCA | ERCC6L | 54821 |
| GGAACCGCGGGGACCTAGGA | VIPR2 | 7434 |
| GGAACGAGGCAGTGACAGGG | NO_CURRENT_644 | NO_CURRENT_644 |
| GGAACTACAGGTGTGGGGCA | FAM214B | 80256 |
| GGAACTGAGAGGAAGAAGGG | BCORL1 | 63035 |
| GGAACTGGGCTGTGAATGAG | CYBB | 1536 |
| GGAACTGTTCGGCTCGCCGG | IFNAR2 | 3455 |
| GGAAGAAGCGTCTTTAGTGC | IFNAR2 | 3455 |
| GGAAGACCCCGACTCCTTAC | MCL1 | 4170 |
| GGAAGAGCTGTCTGCACCAA | HTR2A | 3356 |
| GGAAGAGGAGGTTTCGCCAC | NFKB1 | 4790 |
| GGAAGCATCATAGGACTATG | NO_CURRENT_645 | NO_CURRENT_645 |
| GGAAGCCACCTCTGTTTGCT | CXCR1 | 3577 |
| GGAAGCGTTGGCTCTTCTCC | NO_CURRENT_646 | NO_CURRENT_646 |
| GGAAGCTGTCAGCGCGTCGC | NFKB1 | 4790 |
| GGAAGGACCCAGAAGCGCGG | AHCTF1 | 25909 |
| GGAAGGATGCTTCACACTCG | TLR9 | 54106 |
| GGAAGGCAGGCCGCGCGCGC | TIFA | 92610 |
| GGAAGGCCGGCACTGACAGT | MCTS1 | 28985 |
| GGAAGGGGTCTGTCTGTGCA | RIPK3 | 11035 |
| GGAAGGGTACTCCAGGCGAG | MPC2 | 25874 |
| GGAATGTCCTAGGTTACTGA | NO_CURRENT_647 | NO_CURRENT_647 |
| GGAATTACGACTAACCGATT | NO_CURRENT_648 | NO_CURRENT_648 |
| GGAATTCAGCTGCTGGAGGG | SERPINE1 | 5054 |
| GGAATTGGACTTGGGAGGCG | RND3 | 390 |
| GGACACAGGCAGGGCGAGCG | WASF2 | 10163 |
| GGACAGCCGGAGAACGGGCG | SLC25A6 | 293 |
| GGACAGCCGGGACTGGGCGC | JAK1 | 3716 |
| GGACCGCCAGCTCAGAACAG | AHR | 196 |
| GGACCGTGAGGGGTAAGGTC | SPCS1 | 28972 |
| GGACCTAGGACGGAGGCGGC | VIPR2 | 7434 |
| GGACCTCCCTGCGACCGTCG | ARRB1 | 408 |
| GGACCTCCTGCAGCCATGGC | MYD88 | 4615 |
| GGACCTGCCGGGGCCTCTCA | IRAK1 | 3654 |
| GGACGACTTCCAAGGCAGCC | NMRK2 | 27231 |
| GGACGCACCATTCCGGGTGA | NO_CURRENT_649 | NO_CURRENT_649 |
| GGACGCATGCCCTCAGGTGC | CYP2R1 | 120227 |
| GGACGTCGCGCTGGCAGACC | NNT | 23530 |
| GGACGTGACTGCTCTATCCC | APAF1 | 317 |

| | | |
|---|---|---|
| GGACGTGCTCTGCCAGCCAG | TIFA | 92610 |
| GGACTATCCACCGTTTACTC | NO_CURRENT_650 | NO_CURRENT_650 |
| GGACTCCCGGAGCTAGGGGT | CFLAR | 8837 |
| GGACTCCCTGGCTGAGGCTG | NUFIP2 | 57532 |
| GGACTGAGTGCAAGGTGACT | ATP5J | 522 |
| GGACTGCTCCTCCTCGGACC | SCTR | 6344 |
| GGACTGGGGCGCCTCTAAAG | ARNT | 405 |
| GGAGAAGAAGAGGTAGCGAG | APAF1 | 317 |
| GGAGAAGCAGCGGCTGGCGT | RIPK2 | 8767 |
| GGAGAAGGAGGAGCGGGCCG | RBM6 | 10180 |
| GGAGAAGGGCTGCGGGTTAG | NFRKB | 4798 |
| GGAGAGACAGAGCCACACAC | BCORL1 | 63035 |
| GGAGAGCGAACGAGCAGGGC | IDH2 | 3418 |
| GGAGAGCGAGAGGAGCAGGC | RND3 | 390 |
| GGAGAGCGAGATCCGGAGTT | NFKB2 | 4791 |
| GGAGAGCGTTCTGGGTCCGA | SLC3A2 | 6520 |
| GGAGAGGAAAATCGGCACAG | NO_CURRENT_651 | NO_CURRENT_651 |
| GGAGAGGAAGCTGGGCACCA | CYP4B1 | 1580 |
| GGAGAGGCGAGGGTTCTGGC | GLMN | 11146 |
| GGAGAGGGATGGTCTCTGCA | SPEN | 23013 |
| GGAGAGGTCTGAAGAAACCT | NLRP1 | 22861 |
| GGAGAGTGAGGGTAGGTGCA | RTP1 | 132112 |
| GGAGAGTGTGCGGCTCCAGG | CHEK2 | 11200 |
| GGAGAGTTTGGAGTTGCTTG | TRERF1 | 55809 |
| GGAGATCAGACATTGCTGTC | NCKAP1L | 3071 |
| GGAGATGCGGCCTTCTCAAA | NO_CURRENT_652 | NO_CURRENT_652 |
| GGAGCATAGCCATGGTTCTC | CYP8B1 | 1582 |
| GGAGCATGTTAACACCTGCG | NO_CURRENT_653 | NO_CURRENT_653 |
| GGAGCCACGGGGAGGTGTCA | CYP24A1 | 1591 |
| GGAGCCCACCCGGACGAAGG | RBM42 | 79171 |
| GGAGCCCGCCGGCGGTGGCG | TBK1 | 29110 |
| GGAGCCGCAATGCCTAAAGG | PDAP1 | 11333 |
| GGAGCCGGCGGCTGCGGTGG | RXRA | 6256 |
| GGAGCCGGTCAGAGGGTGAG | UBE2H | 7328 |
| GGAGCCTGGAGAGAAGGCGC | TNFRSF1B | 7133 |
| GGAGCCTTCGCCACCTTCGC | TRAF6 | 7189 |
| GGAGCGATCCCTGCATCCTA | SCN5A | 6331 |
| GGAGCGCCCCCGGCCTCTGT | PRR14L | 253143 |
| GGAGCGCCCGCGGCTGTCAG | RRAGA | 10670 |
| GGAGCGCTCCAGGGCCCTTG | KIF2B | 84643 |
| GGAGCTGGTGGCGCGGTGCA | DPY30 | 84661 |
| GGAGCTGGTGGCGCGGTGCA | MEMO1 | 51072 |
| GGAGGACACACAACAAAACC | NO_CURRENT_654 | NO_CURRENT_654 |
| GGAGGAGGACTGGTAACTTT | RNF25 | 64320 |
| GGAGGAGGGAGGGTGAGTTA | CDKN2D | 1032 |
| GGAGGCAAGCGGCCCAGATC | HAMP | 57817 |
| GGAGGCGGCGTGGAAGGCCG | MLF2 | 8079 |
| GGAGGCGGGGCGCCCTGTGG | SLC25A1 | 6576 |
| GGAGGCGTACTTGAGGGTCT | CYP27B1 | 1594 |
| GGAGGCTGAAGCTGAGGAGG | KHSRP | 8570 |

| | | |
|---|---|---|
| GGAGGGAGGAAAGTAGGGGA | STRAP | 11171 |
| GGAGGGCACCAGGTGAGGCC | RHOG | 391 |
| GGAGGGCAGGAAGGGGAGCA | TRERF1 | 55809 |
| GGAGGGGCTCGGCTGCACCG | STAT1 | 6772 |
| GGAGGGTAGCTAGATCAGCG | ATP5J | 522 |
| GGAGGGTGATCATTAAGTAG | NO_CURRENT_655 | NO_CURRENT_655 |
| GGAGGTAGAGGTTACTCACA | SLC25A19 | 60386 |
| GGAGGTCAAGTATTACGCGG | PAXIP1 | 22976 |
| GGAGGTCGCGGCGGCTGCGT | UTY | 7404 |
| GGAGGTGCTGCAAGACTCTC | GC | 2638 |
| GGAGGTGTCTTAAACATGGT | NO_CURRENT_656 | NO_CURRENT_656 |
| GGAGTAAGAGTGTGAGAAAC | C4orf17 | 84103 |
| GGAGTCCAAACAGCCTTACC | ATP5J2 | 9551 |
| GGAGTCCAAACAGCCTTACC | ATP5J2-PTCD1 | 100526740 |
| GGAGTCCAAACAGCCTTACC | PTCD1 | 26024 |
| GGAGTCCCAGCCGCTTTCAA | NUFIP2 | 57532 |
| GGAGTCCTGGGGCATGGCGG | DET1 | 55070 |
| GGAGTCCTTCACATCCCGCT | MGA | 23269 |
| GGAGTCGAGGGAGCTCGGGC | HTR7 | 3363 |
| GGAGTCGCGCGTTACCCAGG | SAP18 | 10284 |
| GGAGTTAAGCCGAGTTGGGT | SLC7A3 | 84889 |
| GGATATAGCCAATTCTCACG | METTL3 | 56339 |
| GGATATTGAGTAAACCCGAT | NO_CURRENT_657 | NO_CURRENT_657 |
| GGATCACACTGGTAAGGAGG | CBLL1 | 79872 |
| GGATCCTCAACCAGGGAGCT | SERPINA2 | 390502 |
| GGATGCAGGGATCGCTCCCC | SCN5A | 6331 |
| GGATGCAGGGCCGCGTGGCA | C6orf15 | 29113 |
| GGATGGGTGCTATTGTGAGG | CASP3 | 836 |
| GGATGTGCTTCAGCTGCGGG | BNIP3 | 664 |
| GGATGTGGCGGCGGGCTCGG | VDAC1 | 7416 |
| GGATTCATGGCGAGTCCGCA | NO_CURRENT_658 | NO_CURRENT_658 |
| GGATTCCAGAGACATCACCC | CYP4A11 | 1579 |
| GGATTGAATGGCTAACGCGG | NO_CURRENT_659 | NO_CURRENT_659 |
| GGATTGGGTTCCAGTTACCC | ERCC6L | 54821 |
| GGATTTGTCGCTTGCCACAC | NO_CURRENT_660 | NO_CURRENT_660 |
| GGCAAAAGGGATAGAACCAG | APAF1 | 317 |
| GGCAAAGGAGCAAGATCCAT | CD84 | 8832 |
| GGCACAGCCAGGGCACCAGG | HPF1 | 54969 |
| GGCACAGCCAGGGCACCAGG | NMU | 10874 |
| GGCACAGCCCCGTGGGTCGG | ARID3A | 1820 |
| GGCACCCGCGTCGTGCCTCC | PDHA1 | 5160 |
| GGCACTGAGCTCCCAGATCT | HAMP | 57817 |
| GGCACTGGGCGCAGGCTCAG | SCN5A | 6331 |
| GGCAGACGTTGTGGCGGAGA | NDUFS4 | 4724 |
| GGCAGCAAGCGTGGGAACGC | IRF8 | 3394 |
| GGCAGCGAGAAAGCGCAGCC | TLR2 | 7097 |
| GGCAGCGGCGGAGAAGGGAG | MAPK9 | 5601 |
| GGCAGCGGCTCCAGCAGCAG | ADRB1 | 153 |
| GGCAGCTGGCCCAGAGGACA | EBI3 | 10148 |
| GGCAGCTGTGAGGGGGTTCC | PITPNB | 23760 |

| | | |
|---|---|---|
| GGCAGGAGCAGGACCTGGCC | ASH2L | 9070 |
| GGCAGGGAATCTGGCTTGAT | RND1 | 27289 |
| GGCAGGGACCACTCGGGGTC | MPC1 | 51660 |
| GGCAGGGAGCTGCGCTCCTC | C6orf15 | 29113 |
| GGCAGGGCAAGGGTCCCAGG | PDE4A | 5141 |
| GGCAGGGCAGGACAGGAAGC | ZNF699 | 374879 |
| GGCAGGGCGAGCGCGGCTGG | WASF2 | 10163 |
| GGCAGGTAGGCACAGTGGGC | IRF8 | 3394 |
| GGCAGGTAGGGTATACATGG | NO_CURRENT_661 | NO_CURRENT_661 |
| GGCAGGTCCCGGCCCGGCGG | IRAK1 | 3654 |
| GGCAGTGTCTGTGGTGGCCG | MKL1 | 57591 |
| GGCAGTGTGGGCGACAGGAC | NLRP3 | 114548 |
| GGCATAGAGCGAGGCCCCAG | ALOX15 | 246 |
| GGCATCGGACACACTAATAG | NO_CURRENT_662 | NO_CURRENT_662 |
| GGCATGACGCCTTTCCGCGG | RB1 | 5925 |
| GGCATGGCGGGCGGGAGGCG | SLC25A1 | 6576 |
| GGCATTGCGGCCAGCCAGCC | MEX3B | 84206 |
| GGCCAAGAAAATTCCCCACC | PTGS1 | 5742 |
| GGCCAAGGGGGCTTGGAACC | CBLL1 | 79872 |
| GGCCACCGAGACTTCTGGAC | NR1H3 | 10062 |
| GGCCACGGGTGGCGAGGCTG | PIGY | 84992 |
| GGCCACGGGTGGCGAGGCTG | PYURF | 100996939 |
| GGCCACGGTGACCGTGTAGG | ALOX5 | 240 |
| GGCCACTTTCACTCACTGAG | INO80 | 54617 |
| GGCCAGCAGACCTTTACTAA | NO_CURRENT_663 | NO_CURRENT_663 |
| GGCCAGCCAGCCGGGCACTC | MEX3B | 84206 |
| GGCCAGCCTCGGAGCCAGCA | JUNB | 3726 |
| GGCCAGGGCAGCGACACCGG | CCAR2 | 57805 |
| GGCCAGGTGTGTCCAACAGT | CXCR1 | 3577 |
| GGCCATGGCGGAACCTTCCC | KAT2A | 2648 |
| GGCCCAAGCTGCATGGGTCC | RAD21 | 5885 |
| GGCCCAAGGAGGAGAAGCTC | CYP4B1 | 1580 |
| GGCCCACAACTGAAAGGTCT | PAGR1 | 79447 |
| GGCCCCGGCAGGTCCCGGCC | IRAK1 | 3654 |
| GGCCCCGGGAGCGGAGAGCG | CTNNB1 | 1499 |
| GGCCCCTTCGCCCTGCAGGT | PLD1 | 5337 |
| GGCCCGAAGACGGGAGCAGG | AKT1 | 207 |
| GGCCCGAGAGTCAGAACCTG | SPEN | 23013 |
| GGCCCGAGGCGGCGCGAGGG | FAM107B | 83641 |
| GGCCCGCATCATCGCTTGTT | RAD21 | 5885 |
| GGCCCGCGCGGCTCGGGCTC | RPS6KA4 | 8986 |
| GGCCCGCGCTGGGAAAAAGG | INO80C | 125476 |
| GGCCCGGACCCATGCAGCTT | RAD21 | 5885 |
| GGCCCGGGCGAGCAGCGCGG | ACADS | 35 |
| GGCCCGGGTCCGGCCGGGCG | SBNO2 | 22904 |
| GGCCCTCTAGAAAAGTCTCG | NO_CURRENT_664 | NO_CURRENT_664 |
| GGCCCTGAGAGGGTGTGCTG | STAT2 | 6773 |
| GGCCCTGCCGGGTCGGGCTG | NCOA6 | 23054 |
| GGCCGAGGAAGCAGGCGCTG | LCN2 | 3934 |
| GGCCGCCTGCTCCCGTCTTC | AKT1 | 207 |

| | | |
|---|---|---|
| GGCCGCGCTCCTCGGCCTCC | PTGIS | 5740 |
| GGCCGCGGCGGCGAGAGCGA | MKL1 | 57591 |
| GGCCGCGGGCCAAGCCTACC | ZNF641 | 121274 |
| GGCCGCGTCAGGAGGGCGGG | EP400 | 57634 |
| GGCCGCTAGCACCTGCGCGT | PLD1 | 5337 |
| GGCCGCTCCATGGGGCGGGA | CHUK | 1147 |
| GGCCGGACCAGGTGCGAACC | VDR | 7421 |
| GGCCGGCCCGGCCCTGCGCC | KIAA1211L | 343990 |
| GGCCGGCGGCAGCTGGTGCG | UCHL5 | 51377 |
| GGCCGGCGGGCTGAGGCGTA | ABCB10 | 23456 |
| GGCCGGGGAGTCCGCTCCAG | SPHK1 | 8877 |
| GGCCGGGGATCCTGGAGCCA | PYCARD | 29108 |
| GGCCGTCGTATTCCCCCAAG | NO_CURRENT_665 | NO_CURRENT_665 |
| GGCCTACCTTCTGGGCAAGG | SQSTM1 | 8878 |
| GGCCTCTGGTGCAGCGGCGG | CS | 1431 |
| GGCCTGCGGGCGGCCAGCGA | TRAF6 | 7189 |
| GGCCTGGCCTGTCAGGGCGC | RRAGC | 64121 |
| GGCCTGGGAAGGTTCCGCCA | KAT2A | 2648 |
| GGCCTGGTGCGCTCGCTTAG | SNRNP70 | 6625 |
| GGCCTTTAAGCAGGGATTCG | TFPT | 29844 |
| GGCGACAAAATGGCTGCCCG | TRIR | 79002 |
| GGCGACAGAGCCCCGCTCTC | PHF6 | 84295 |
| GGCGACGAGAACGGCGAGCG | ARHGAP33 | 115703 |
| GGCGACGGCATTTAACAAGG | IKZF5 | 64376 |
| GGCGAGAAGGACGGAGGTAG | SLC25A19 | 60386 |
| GGCGAGCAGCGCGGCGGCCA | ACADS | 35 |
| GGCGAGCAGGCAGCGTTGCA | AHCTF1 | 25909 |
| GGCGAGCATGCGAGTGGTGA | UBE2L6 | 9246 |
| GGCGAGCGAGGGGTCGAGCG | ARHGAP33 | 115703 |
| GGCGAGGCTGCGGTGAGGCC | PIGY | 84992 |
| GGCGAGGCTGCGGTGAGGCC | PYURF | 100996939 |
| GGCGCACCCCGGCGCGCAGG | PDE3A | 5139 |
| GGCGCAGGTGAGCAGGGCAG | EIF2AK2 | 5610 |
| GGCGCATTAAAGTCGAGAGC | NO_CURRENT_666 | NO_CURRENT_666 |
| GGCGCCCGCGGTTCAGGTGA | NFRKB | 4798 |
| GGCGCCGCCCGGGGCCCTGA | SMPD1 | 6609 |
| GGCGCCGCTCTCCGCTGCGG | KAT2A | 2648 |
| GGCGCGCGCAGGCTGAGCTC | MAP2K5 | 5607 |
| GGCGCGCGGAGGCCGCGGCG | RXRA | 6256 |
| GGCGCGCGGGGCGGCGGTGA | ABL1 | 25 |
| GGCGCGGCGGAGACCCGGCT | TBK1 | 29110 |
| GGCGCGGGGCGCGGGCATGG | SLC25A1 | 6576 |
| GGCGCGTTTCTGCGGGCGCA | RAC2 | 5880 |
| GGCGCTCCTCCACAGCTCGC | RB1 | 5925 |
| GGCGCTGGTGGCTGCGGCGG | TMEM30A | 55754 |
| GGCGCTGTGCGCACAGGGGA | NUP153 | 9972 |
| GGCGCTTACCTTGGTTTCTC | ZRSR2 | 8233 |
| GGCGGACAGTGCGGAACTAA | PIAS1 | 8554 |
| GGCGGAGCCTCAGGGCCGGG | TRIR | 79002 |
| GGCGGAGGCAGTGGGGGTCC | CBLL1 | 79872 |

| | | |
|---|---|---|
| GGCGGAGGCTTTGGCAGCTC | ATP5J | 522 |
| GGCGGAGTCGGGGCTCCTGA | SHOC2 | 8036 |
| GGCGGCAACATGGAGCGAGC | KMT2C | 58508 |
| GGCGGCAACCGGGGGCCCCA | NELFE | 7936 |
| GGCGGCAGCAGCGGCTGTGG | CCAR2 | 57805 |
| GGCGGCAGCAGCGGCTGTGG | L1CAM | 3897 |
| GGCGGCAGCGGCAGCAGCTC | SETD1B | 23067 |
| GGCGGCAGCGGCAGCGGCGC | BCL2L11 | 10018 |
| GGCGGCAGCGGCGACGAGAA | ARHGAP33 | 115703 |
| GGCGGCATGACGCCTTTCCG | RB1 | 5925 |
| GGCGGCCATGGCGGGACAGG | BRK1 | 55845 |
| GGCGGCCCGAGGCGGCGCGA | FAM107B | 83641 |
| GGCGGCCGGGCGGTGCGCGG | MAP2K7 | 5609 |
| GGCGGCGACGGCATTTAACA | IKZF5 | 64376 |
| GGCGGCGCCCGTAGGATGCA | SCN5A | 6331 |
| GGCGGCGCTGAGGCGGCTGC | EGLN1 | 54583 |
| GGCGGCGCTGAGGCGGCTGC | INO80 | 54617 |
| GGCGGCGGCAGTGTCTGTGG | MKL1 | 57591 |
| GGCGGCGGCCATGGCGGGAC | BRK1 | 55845 |
| GGCGGCGGCGCAGGTGAGCA | EIF2AK2 | 5610 |
| GGCGGCGGCGGTGGTATCGG | PITPNB | 23760 |
| GGCGGCGGCGGTTGGCCGGT | ANKRD50 | 57182 |
| GGCGGCGGGCGCGGGACGCG | BRD1 | 23774 |
| GGCGGCGGGGACGACTTCCA | NMRK2 | 27231 |
| GGCGGCGGGGGCCGGGCTCC | BAHD1 | 22893 |
| GGCGGCGGTGTCAATGTCAG | NDUFS4 | 4724 |
| GGCGGCGTGGGAGCACCTCT | INO80B | 83444 |
| GGCGGCTCAACGCGGGAACA | KHSRP | 8570 |
| GGCGGCTCCTGCGGCGGTGC | CASP9 | 842 |
| GGCGGCTGCAGCGGGGTGAG | PYCARD | 29108 |
| GGCGGCTGCGGCGGCGGCGA | GMPR | 2766 |
| GGCGGCTGCGGCGGCGGCGA | MAP2K5 | 5607 |
| GGCGGCTGCGGGCCAGGAAC | PRKN | 5071 |
| GGCGGCTGCGGGCGCTGGGC | DOCK4 | 9732 |
| GGCGGCTGCGGGCGCTGGGC | INO80 | 54617 |
| GGCGGCTGCTGCTGCGGCTG | UBE2E1 | 7324 |
| GGCGGCTTGGGGGTCCTGGG | CYFIP1 | 23191 |
| GGCGGGACCGTGAGAACTCG | LGMN | 5641 |
| GGCGGGCGCAGGATACGGGC | PIP4K2A | 5305 |
| GGCGGGCGCTGTGCGCACAG | NUP153 | 9972 |
| GGCGGGCGGCTTAGCGCGCG | SCTR | 6344 |
| GGCGGGCTCGGAGGCGGCTA | VDAC1 | 7416 |
| GGCGGGGCAGGCAGCCGGAC | BAHD1 | 22893 |
| GGCGGGGGCGTCCCTCCCTG | NIPBL | 25836 |
| GGCGGGGGCTCAATGCTCCC | POU2F1 | 5451 |
| GGCGGTACCGGTGCTGGCGG | MTOR | 2475 |
| GGCGGTCCGGGGCACGCTCT | AHR | 196 |
| GGCGGTGAGGGCGGCTGGCG | ABL1 | 25 |
| GGCGGTGATGGACGGGTCCG | BAX | 581 |
| GGCGGTGGCATCTGCGGCCA | ARNT | 405 |

| | | |
|---|---|---|
| GGCGGTGGCGCTGGTGGCTG | TMEM30A | 55754 |
| GGCGGTGGCGGCGGGGAAGA | MAP2K7 | 5609 |
| GGCGGTGGGAAAGCGGAGGT | BNIP3 | 664 |
| GGCGGTTCGTCCTGCTCTGG | SERPINE1 | 5054 |
| GGCGTCAAAATTAGAAGCCG | NO_CURRENT_667 | NO_CURRENT_667 |
| GGCGTCAGTTGGTGAGTGTC | ATP5J2 | 9551 |
| GGCGTCAGTTGGTGAGTGTC | ATP5J2-PTCD1 | 100526740 |
| GGCGTCAGTTGGTGAGTGTC | PTCD1 | 26024 |
| GGCGTCGCCAGTGGCTCCCA | DNTTIP1 | 116092 |
| GGCGTCGCGCGCGCGCCCCA | PYCARD | 29108 |
| GGCGTTAATTAAACTGTTTT | NO_CURRENT_668 | NO_CURRENT_668 |
| GGCGTTGGCGGTCTTGGCAT | PIGL | 9487 |
| GGCTAAGACGCCGGAGGAGG | GLMN | 11146 |
| GGCTACGGTGAGCCGAAGGT | BABAM1 | 29086 |
| GGCTATCCCCTGAAAGACTA | NAIP | 4671 |
| GGCTCACCTTGAAGCTGTTG | BCL2A1 | 597 |
| GGCTCACGCGGTGAGTCATA | CHEK2 | 11200 |
| GGCTCCAGCAGCAGCGGCGG | ADRB1 | 153 |
| GGCTCCAGGACCGCCCGCCA | MYD88 | 4615 |
| GGCTCCCGGCCCCCGACCCA | ARID3A | 1820 |
| GGCTCCTACCTGCAGGGCGA | PLD1 | 5337 |
| GGCTCGGCCCGCGGTCCCCG | APEH | 327 |
| GGCTCGGCGATCAGCGGCGG | C16orf72 | 29035 |
| GGCTCGGTACCACCCGGCAG | HDAC2 | 3066 |
| GGCTCGGTGCTCACGGGTCA | SERF2 | 10169 |
| GGCTCTGGGGCTCACGGACG | NO_CURRENT_669 | NO_CURRENT_669 |
| GGCTCTTCCAGCAGCGGCAG | ADRB1 | 153 |
| GGCTCTTGGGTCACATACCT | HSH2D | 84941 |
| GGCTGAAGCTGAGGAGGCGG | KHSRP | 8570 |
| GGCTGAGCTCCGGCCACCCG | MAP2K5 | 5607 |
| GGCTGAGGACTGAAAAGAGA | GPR119 | 139760 |
| GGCTGATCTCGCCGCCCTTA | TBXAS1 | 6916 |
| GGCTGCAAGGAAAAAAGCTG | UTY | 7404 |
| GGCTGCAGCGGGGTGAGCGG | PYCARD | 29108 |
| GGCTGCAGGAGGTCCCGGCG | MYD88 | 4615 |
| GGCTGCCAGACCATGCTGAG | PIGY | 84992 |
| GGCTGCCAGACCATGCTGAG | PYURF | 100996939 |
| GGCTGCGGGCGCTGGGCCGG | INO80 | 54617 |
| GGCTGCTAGGAGCCGGTCAG | UBE2H | 7328 |
| GGCTGCTCTGAGGAGGTCGA | JAM2 | 58494 |
| GGCTGCTCTGGATGATGACG | TMEM173 | 340061 |
| GGCTGGCAGGTCGCCATCTT | GPR119 | 139760 |
| GGCTGGCGCCGCTGCGCGCA | PRKN | 5071 |
| GGCTGGGCTCGTGGTTGCCG | NQO1 | 1728 |
| GGCTGGGGCGGCCAAACCCG | IKBKB | 3551 |
| GGCTGGTGAGGCGCCTGCTG | RAC2 | 5880 |
| GGCTGGTTGACGACTCCTGA | NO_CURRENT_670 | NO_CURRENT_670 |
| GGCTGTGGCGGTGGCGCTGG | TMEM30A | 55754 |
| GGCTGTGGTGGCTTCGGCAG | SOD2 | 6648 |
| GGCTTACGTGGGGGGCAAAA | NO_CURRENT_671 | NO_CURRENT_671 |

| | | |
|---|---|---|
| GGCTTCAGCAGCCCGCGCCC | IRAK4 | 51135 |
| GGCTTCGAGCGCGATCTACT | CREBBP | 1387 |
| GGCTTCTCCAGAGGGTCTGG | TLR9 | 54106 |
| GGCTTCTCCTCCATTGAAAG | NUFIP2 | 57532 |
| GGCTTCTTCCTGAGGACCTC | TYK2 | 7297 |
| GGCTTGTACCTCCTGCGCGC | PDE3A | 5139 |
| GGCTTTGCCCGTTGGGGTAA | MSL2 | 55167 |
| GGGAAACTAGACATATGTAC | NO_CURRENT_672 | NO_CURRENT_672 |
| GGGAACCGCGAGCGGCGTAG | BAHD1 | 22893 |
| GGGAACTGGGCTGTGAATGA | CYBB | 1536 |
| GGGAAGAAGCGTCTTTAGTG | IFNAR2 | 3455 |
| GGGAAGCGGCAGCAGAGGCA | MPC1 | 51660 |
| GGGAAGGAGCGTTGGGGATC | LARP4B | 23185 |
| GGGAAGGCAGGCCGCGCGCG | TIFA | 92610 |
| GGGAAGGGGTCTGTCTGTGC | RIPK3 | 11035 |
| GGGACATCCTTGCCGTCTCA | NO_CURRENT_673 | NO_CURRENT_673 |
| GGGACCATCTACCCTCAATG | NO_CURRENT_674 | NO_CURRENT_674 |
| GGGACCGTGAGAACTCGCGG | LGMN | 5641 |
| GGGACCTCCTGCAGCCATGG | MYD88 | 4615 |
| GGGACGCGAAAGAAACCAGT | NO_CURRENT_675 | NO_CURRENT_675 |
| GGGACGCTAACGATATCCTG | NO_CURRENT_676 | NO_CURRENT_676 |
| GGGACGTAACGGAGGCAGGT | SERF2 | 10169 |
| GGGACTCCCGGAGCTAGGGG | CFLAR | 8837 |
| GGGACTGAGTGCAAGGTGAC | ATP5J | 522 |
| GGGACTGATATATGGCGAAC | NO_CURRENT_677 | NO_CURRENT_677 |
| GGGACTGTGTCTGTCGCCCA | ACADS | 35 |
| GGGAGACCCTTGGATGGCCG | HSH2D | 84941 |
| GGGAGAGCGTTCTGGGTCCG | SLC3A2 | 6520 |
| GGGAGCAGAGGAGGCGAGGG | SIRT1 | 23411 |
| GGGAGCCCACCCGGACGAAG | RBM42 | 79171 |
| GGGAGCCCCCACCGCCCCCG | BAP1 | 8314 |
| GGGAGCGCTCCAGGGCCCTT | KIF2B | 84643 |
| GGGAGGAGGACTGGTAACTT | RNF25 | 64320 |
| GGGAGGCACCTCGGAGATCT | LAMTOR2 | 28956 |
| GGGAGGCGCGGTGAGGAGTC | RND3 | 390 |
| GGGAGGGCTCAGCAAAGCGG | BCORL1 | 63035 |
| GGGAGGTAGGCAGCAGCGCG | PDHB | 5162 |
| GGGAGGTCGCGGCGGCTGCG | UTY | 7404 |
| GGGAGGTGGCTTTAGGTTTT | NO_CURRENT_678 | NO_CURRENT_678 |
| GGGAGTACTGCGACGCAGCC | MAPK14 | 1432 |
| GGGAGTCTCGGGGATGACAG | MSL2 | 55167 |
| GGGAGTGTGAGCGGTGCTCT | SLC11A1 | 6556 |
| GGGAGTTGATTGTTTCGAGA | NO_CURRENT_679 | NO_CURRENT_679 |
| GGGATCAATTGTTCATGAGT | NO_CURRENT_680 | NO_CURRENT_680 |
| GGGATCCGAGTGAGGCGACG | ATP6V0E1 | 8992 |
| GGGATCCTCAACCAGGGAGC | SERPINA2 | 390502 |
| GGGATCCTTTAGACAACTTC | NAIP | 4671 |
| GGGATGCGTCTTGCTAAACC | NO_CURRENT_681 | NO_CURRENT_681 |
| GGGATGGCGCGCTTTGACTC | CYP1B1 | 1545 |
| GGGATGGTCTCTGCACGGGG | SPEN | 23013 |

| | | |
|---|---|---|
| GGGATTGGTTGAGCAAAGCG | NO_CURRENT_682 | NO_CURRENT_682 |
| GGGCAAAGGAGGTGGCATGT | TNFRSF6B | 8771 |
| GGGCAATAGAAGCAGAGCTT | NO_CURRENT_683 | NO_CURRENT_683 |
| GGGCACCGGGGTCGCACTCA | PTGS1 | 5742 |
| GGGCACTCGGGGGGCACGCG | MEX3B | 84206 |
| GGGCAGAAGTTGCTGTCCTG | NO_CURRENT_684 | NO_CURRENT_684 |
| GGGCAGAGTCGGAGATGCAG | PDK4 | 5166 |
| GGGCAGCCCCCGGCGCAGCG | EGFR | 1956 |
| GGGCAGCTGAGAGCGAGAGG | UCHL5 | 51377 |
| GGGCAGGCAGCCGGACGGGC | BAHD1 | 22893 |
| GGGCAGGGCAGGACAGGAAG | ZNF699 | 374879 |
| GGGCAGGGGGCCTGAAGCGG | MTOR | 2475 |
| GGGCATAAGTTAGCCACTCA | NO_CURRENT_685 | NO_CURRENT_685 |
| GGGCATCGGCTTCCAGTCCG | RAC1 | 5879 |
| GGGCCAGGGCAGCGACACCG | CCAR2 | 57805 |
| GGGCCCAAGCTGCATGGGTC | RAD21 | 5885 |
| GGGCCCGCATAGGATATCGC | NO_CURRENT_686 | NO_CURRENT_686 |
| GGGCCCTGGAGAGATGGCCA | IL2RB | 3560 |
| GGGCCGAATCGCCAAGACGC | ATP5E | 514 |
| GGGCCGCGGGCCAAGCCTAC | ZNF641 | 121274 |
| GGGCCGGCCCGCTCCATGGT | PDE5A | 8654 |
| GGGCCGGGTACCATGGACGT | CTNNBL1 | 56259 |
| GGGCCTTGGAGTCGGCGCGC | NOD2 | 64127 |
| GGGCGACCCTGGCCGGAACT | NQO1 | 1728 |
| GGGCGATGCTTCAGGTGGTG | KMT2D | 8085 |
| GGGCGCCCTGTGGCGGCTTC | SLC25A1 | 6576 |
| GGGCGCGCGAGCCTCCGGGC | TP73 | 7161 |
| GGGCGCGGAGAGTGAATGGA | SLC25A6 | 293 |
| GGGCGCGGCCGGAGACCGTG | NELFE | 7936 |
| GGGCGCTCACACCGTGCGGG | EGFR | 1956 |
| GGGCGCTCAGTCCTGTGTCC | NMU | 10874 |
| GGGCGCTGTGGAGCAGAGCT | FOS | 2353 |
| GGGCGGCCCGGGCGGCCACT | PPID | 5481 |
| GGGCGGGCGCTGTGCGCACA | NUP153 | 9972 |
| GGGCGGGGCAGGCAGCCGGA | BAHD1 | 22893 |
| GGGCGGTGGGGGCTCCCGGT | BAP1 | 8314 |
| GGGCGGTGTGACTTAGGACG | IFNAR1 | 3454 |
| GGGCGTGTATGTTCGTATTG | NO_CURRENT_687 | NO_CURRENT_687 |
| GGGCTACCTAATTAACTGCA | ACOD1 | 730249 |
| GGGCTAGAAGTAGAGGGCTT | ARR3 | 407 |
| GGGCTCCTGACGGTAACTCG | SHOC2 | 8036 |
| GGGCTCGGAGGCGGCTACGG | VDAC1 | 7416 |
| GGGCTGCTGGCGGCGGGGGA | KPNB1 | 3837 |
| GGGCTGGAGAGGCCGCGGAC | RRAGA | 10670 |
| GGGCTGGCCTCGCGCCCTCG | SLC6A4 | 6532 |
| GGGCTGGCTCTTACTCAGAT | RRAGA | 10670 |
| GGGCTTGACATTAAGATACC | NO_CURRENT_688 | NO_CURRENT_688 |
| GGGCTTTGGAATATTACCTG | ARR3 | 407 |
| GGGGAAACAAGTAGGCTTTG | NO_CURRENT_689 | NO_CURRENT_689 |
| GGGGAACTGGGCTGTGAATG | CYBB | 1536 |

| | | |
|---|---|---|
| GGGGAAGAATCAACACACCA | WDR26 | 80232 |
| GGGGAAGGTCAGCGCCGTAA | NDUFB9 | 4715 |
| GGGGAATGCCTTACCTGCCC | IRAK4 | 51135 |
| GGGGACAGGGGCGATGCTTC | KMT2D | 8085 |
| GGGGACTGGCTGCGCTTGAC | TYK2 | 7297 |
| GGGGAGAAGGCGCCGTGTCC | PAGR1 | 79447 |
| GGGGAGACCCGGCCCCCAAG | CDKN2D | 1032 |
| GGGGAGCAGCCCAGAGGCGG | BAX | 581 |
| GGGGAGCGCACGACGTACCT | TLR2 | 7097 |
| GGGGAGGCGGCGGCCTGGCT | RRAGC | 64121 |
| GGGGAGGCTGGTCCCGGGCT | SLC6A4 | 6532 |
| GGGGCACTCAGAGATCCAGC | CYP4A11 | 1579 |
| GGGGCACTCAGAGATCCAGC | CYP4A22 | 284541 |
| GGGGCAGGCAGCCGGACGGG | BAHD1 | 22893 |
| GGGGCAGGCAGGGCGGGGCG | ZNF699 | 374879 |
| GGGGCAGGCGGCGCCCGGGA | ZNF641 | 121274 |
| GGGGCATGATTTCAGGGCCG | LCN2 | 3934 |
| GGGGCCAGCAGCCGGGAAAG | NDUFC1 | 4717 |
| GGGGCCCAAATATTAAAATC | NLRC4 | 58484 |
| GGGGCCCTGAGGGCTGGCTA | SMPD1 | 6609 |
| GGGGCCTGAAGCGGCGGTAC | MTOR | 2475 |
| GGGGCGATGCTTCAGGTGGT | KMT2D | 8085 |
| GGGGCGCCCTGTGGCGGCTT | SLC25A1 | 6576 |
| GGGGCGCCTCTAAAGGGGAG | ARNT | 405 |
| GGGGCGGCGGTGAGGGCGGC | ABL1 | 25 |
| GGGGCGGGCGCTGTGCGCAC | NUP153 | 9972 |
| GGGGCGGTGTGACTTAGGAC | IFNAR1 | 3454 |
| GGGGCTCCTGACGGTAACTC | SHOC2 | 8036 |
| GGGGCTGCGACGCGGAGGCA | VDAC1 | 7416 |
| GGGGCTGGAGAGGCCGCGGA | RRAGA | 10670 |
| GGGGCTGGGGCGGCCAAACC | IKBKB | 3551 |
| GGGGCTTACGTGAAGGGCGG | NO_CURRENT_690 | NO_CURRENT_690 |
| GGGGGAAGAATCAACACACC | WDR26 | 80232 |
| GGGGGAGACCCGGCCCCCAA | CDKN2D | 1032 |
| GGGGGAGGCGCGCACAGAAC | DET1 | 55070 |
| GGGGGCACAGAATCACTGAA | FBXW7 | 55294 |
| GGGGGCAGCTCAGAGCGCGC | KPNB1 | 3837 |
| GGGGGCCAGCAGCCGGGAAA | NDUFC1 | 4717 |
| GGGGGCGTAGCGCGCTGAGG | WASF2 | 10163 |
| GGGGGCTGCGACGCGGAGGC | VDAC1 | 7416 |
| GGGGGGACCAGGGGAAGACT | INO80C | 125476 |
| GGGGGGCAGCTGAGAGCGAG | UCHL5 | 51377 |
| GGGGGGTCACCTTCAGAGCG | SPINT1 | 6692 |
| GGGGGGTCCGTGCCCATGGG | PDE4A | 5141 |
| GGGGGGTTGGTGGCACTATG | DLAT | 1737 |
| GGGGGTCACCTTCAGAGCGC | SPINT1 | 6692 |
| GGGGGTCCGTGCCCATGGGA | PDE4A | 5141 |
| GGGGTCCGTGCCCATGGGAG | PDE4A | 5141 |
| GGGGTCGCACTCACGGCTCA | PTGS1 | 5742 |
| GGGGTGCGCGGGCGGTTGTG | CREBBP | 1387 |

| | | |
|---|---|---|
| GGGGTGTTCCCACACTCACC | EDA2R | 60401 |
| GGGGTTTCCGTTGCAGTCCT | SOD1 | 6647 |
| GGGTAAGACTATGACTTATA | GBP7 | 388646 |
| GGGTAATGGTGACTCAGTGT | NO_CURRENT_691 | NO_CURRENT_691 |
| GGGTACAATGTAGACTACGT | NO_CURRENT_692 | NO_CURRENT_692 |
| GGGTAGATCCATGAAACAGC | NO_CURRENT_693 | NO_CURRENT_693 |
| GGGTAGGATTCAGCCAAAAG | NO_CURRENT_694 | NO_CURRENT_694 |
| GGGTATAGACGCGATCCTCA | NO_CURRENT_695 | NO_CURRENT_695 |
| GGGTCAACGCTGGCTGCCGC | UCHL3 | 7347 |
| GGGTCCGAGGGTCCAGGTAG | SLC3A2 | 6520 |
| GGGTCCGGGGAGCAGCCCAG | BAX | 581 |
| GGGTCCGTGCCCATGGGAGG | PDE4A | 5141 |
| GGGTCGCGGCCCGGAACGTG | MLF2 | 8079 |
| GGGTCTCCGGCCGGCGGCGG | EIF2AK2 | 5610 |
| GGGTCTGCCTTCCCCGGTTC | NO_CURRENT_696 | NO_CURRENT_696 |
| GGGTGAGCGCGGGGGTCCCA | HELZ2 | 85441 |
| GGGTGAGTTAGGGGGAGACC | CDKN2D | 1032 |
| GGGTGCAAAAGCCCAGGGTT | LAMTOR2 | 28956 |
| GGGTGCAGCTGCTCCTGTCC | NUDT17 | 200035 |
| GGGTGCCCACTAATAGCCGC | NO_CURRENT_697 | NO_CURRENT_697 |
| GGGTGGAGACCCACGAGCCG | CAT | 847 |
| GGGTGGGGGCAGAGCTCTAG | MEX3B | 84206 |
| GGGTGGTCATTCTCTACTTG | NO_CURRENT_698 | NO_CURRENT_698 |
| GGGTGTCATAGCCCCGGGTT | IKBKB | 3551 |
| GGGTTAAAGTGCTTTCTGCA | AGER | 177 |
| GGGTTCCTCACCTGCGTCCG | CYFIP1 | 23191 |
| GGGTTCGCACCTGGTCCGGC | VDR | 7421 |
| GGTAACTGGAACCCAATCCG | ERCC6L | 54821 |
| GGTACCTGCTGCTGCTCCCG | KMT2C | 58508 |
| GGTACTTACAATGACAAAAA | CYBB | 1536 |
| GGTAGACGGGGCATCTCAGC | NO_CURRENT_699 | NO_CURRENT_699 |
| GGTAGCCGGCCATCCCAAGC | IDH2 | 3418 |
| GGTAGCGGGAGGGCAGACTC | INO80E | 283899 |
| GGTAGCTGAAGACCAGACCG | RHOA | 387 |
| GGTAGGCTTGGCCCGCGGCC | ZNF641 | 121274 |
| GGTAGGGGTTGGCGCTCAGG | ATP6V0E1 | 8992 |
| GGTATCCACTATCTCGGGCA | LTA4H | 4048 |
| GGTATCGGCGGCAGCTGTGA | PITPNB | 23760 |
| GGTATCTGGGACCTGTTCTC | BTN2A2 | 10385 |
| GGTATGAGCCTAATTTCCAC | NO_CURRENT_700 | NO_CURRENT_700 |
| GGTCAAAATGGCTGGTAAGC | UQCRB | 7381 |
| GGTCACCGATCGAGAGCTAG | NO_CURRENT_701 | NO_CURRENT_701 |
| GGTCACCTACAGGAGAGAGG | GC | 2638 |
| GGTCAGTCTCTAGACCAAGA | RIPK3 | 11035 |
| GGTCATGAATCATGTGACGG | LAMTOR3 | 8649 |
| GGTCCCCAGCTCCCTGGTTG | SERPINA2 | 390502 |
| GGTCCCGGCGGCAGGAGCGC | JAM2 | 58494 |
| GGTCCCTCTGGCTGGGTTAA | NO_CURRENT_702 | NO_CURRENT_702 |
| GGTCCGAGGAGGAGCAGTCC | SCTR | 6344 |
| GGTCCGCGACGGTCGCAGGG | ARRB1 | 408 |

| | | |
|---|---|---|
| GGTCCGCGCACAAGAGCAGG | NO_CURRENT_703 | NO_CURRENT_703 |
| GGTCCGTGCCCATGGGAGGG | PDE4A | 5141 |
| GGTCCTCAGGAAGAAGCCGC | TYK2 | 7297 |
| GGTCGCCTCTGCTCGGTCTG | STAT1 | 6772 |
| GGTCGCCTGTGCGACATGCT | NO_CURRENT_704 | NO_CURRENT_704 |
| GGTCGCGTCCGACACCCGGT | EZH2 | 2146 |
| GGTCTCCGCCGCGCCACCGC | TBK1 | 29110 |
| GGTCTGCTCCAATGGGAACC | NO_CURRENT_705 | NO_CURRENT_705 |
| GGTCTTCCAGATGAGACAGG | TBXAS1 | 6916 |
| GGTCTTCTTGACAGCGCTCT | SERPINE1 | 5054 |
| GGTGAAGGTCTACACATCAG | IL10 | 3586 |
| GGTGACCGTGTAGGAGGGCA | ALOX5 | 240 |
| GGTGACGAAGACGGCGGCGG | IKZF5 | 64376 |
| GGTGACTGATGTCGTAACTA | PHC3 | 80012 |
| GGTGATCCGTCCTTTCATCC | NDUFS4 | 4724 |
| GGTGCAGCGGGAGATTCACC | BRK1 | 55845 |
| GGTGCAGGAGTCCTGGGGCA | DET1 | 55070 |
| GGTGCCATTTGTCATACTGA | NO_CURRENT_706 | NO_CURRENT_706 |
| GGTGCGGAGACCCCTTCGGG | PDHB | 5162 |
| GGTGCGGGCGTCTTGCGAGT | CYP2R1 | 120227 |
| GGTGCTCCTGGCTGGGCGTG | ITGB2 | 3689 |
| GGTGGAGCACCAGCTACGCG | NMU | 10874 |
| GGTGGCATCTGCGGCCATGG | ARNT | 405 |
| GGTGGCGAGGCTCTCAAGCC | FLCN | 201163 |
| GGTGGCGAGGCTCTCAAGCC | PLD6 | 201164 |
| GGTGGCGCTGGTGGCTGCGG | TMEM30A | 55754 |
| GGTGGCGGAGGCTGAGGAGA | RBM6 | 10180 |
| GGTGGCGGCGGGGAAGATGG | MAP2K7 | 5609 |
| GGTGGCTGCTGGGCCTGGCA | LCN2 | 3934 |
| GGTGGCTGGAACTGCAACCA | CYP4B1 | 1580 |
| GGTGGGGGGTCCGTGCCCAT | PDE4A | 5141 |
| GGTGGTAGTGGCGGCGGCGG | BCL2L11 | 10018 |
| GGTGGTAGTGGCGGCGGCGG | ERC1 | 23085 |
| GGTGTCACCACCGCTTACCA | NO_CURRENT_707 | NO_CURRENT_707 |
| GGTGTCCCACCGCCACAAGG | IKBKE | 9641 |
| GGTGTCCCCGGCGCCGCCCG | SMPD1 | 6609 |
| GGTGTCGGCACAGCCATGGC | MPC1 | 51660 |
| GGTGTCTCTCCCGACCATGG | PCGF6 | 84108 |
| GGTGTCTTACGTAAAGAGGT | NO_CURRENT_708 | NO_CURRENT_708 |
| GGTGTGAGGATCCGAACCCA | HSP90B1 | 7184 |
| GGTGTGGACGCTGTGAATCG | STRAP | 11171 |
| GGTTAAAATATTCAAAATGG | POU2F1 | 5451 |
| GGTTACATGGTTAGATGGAG | NO_CURRENT_709 | NO_CURRENT_709 |
| GGTTACTCACACGGCTCGGC | SLC25A19 | 60386 |
| GGTTAGGGGGCCGGCGCCCG | NFRKB | 4798 |
| GGTTGCCGCCCCCTCGGGTA | NELFE | 7936 |
| GGTTGTAAATCCTTCTAACG | NO_CURRENT_710 | NO_CURRENT_710 |
| GGTTTGAGTGAGCAATTACG | NO_CURRENT_711 | NO_CURRENT_711 |
| GTAACACCCAAAGCTAACTC | NO_CURRENT_712 | NO_CURRENT_712 |
| GTAACAGTCTGATTACGGAC | NO_CURRENT_713 | NO_CURRENT_713 |

| | | |
|---|---|---|
| GTAACATATCTTAACAACCA | C4orf17 | 84103 |
| GTAACTAGGGGGGGCGAGTT | PHC3 | 80012 |
| GTAACTCGGGGCCGATGAGG | SHOC2 | 8036 |
| GTAACTGGAACCCAATCCGA | ERCC6L | 54821 |
| GTAACTGGTGGGATCTGCGG | IFNAR1 | 3454 |
| GTAAGCAAACTCCAAAGAAA | CUBN | 8029 |
| GTAAGGAATGACTTGACTGG | NO_CURRENT_714 | NO_CURRENT_714 |
| GTAAGGAGGCGGAGGCAGTG | CBLL1 | 79872 |
| GTAAGGCTGTTTGGACTCCG | ATP5J2 | 9551 |
| GTAAGGCTGTTTGGACTCCG | ATP5J2-PTCD1 | 100526740 |
| GTAAGGCTGTTTGGACTCCG | PTCD1 | 26024 |
| GTAATTTTATGAGTTAAGTG | NO_CURRENT_715 | NO_CURRENT_715 |
| GTACACACTTATGCCATCAC | NO_CURRENT_716 | NO_CURRENT_716 |
| GTACCATTGCCGGCTCCCTA | NO_CURRENT_717 | NO_CURRENT_717 |
| GTACCCCTATGGCCGTTCTA | NO_CURRENT_718 | NO_CURRENT_718 |
| GTACTTGAGGGTCTGGGTCA | CYP27B1 | 1594 |
| GTAGACAGCAGAACCAGCGG | RBM42 | 79171 |
| GTAGACGTCGTGAGCTTCAC | NO_CURRENT_719 | NO_CURRENT_719 |
| GTAGATCGCGCTCGAAGCCC | CREBBP | 1387 |
| GTAGATTCGCTGGAAGCAGC | SLC5A8 | 160728 |
| GTAGCCAGCGATACCTGTAA | OXSM | 54995 |
| GTAGCGAAAGCGGAGCTCGT | ATP5E | 514 |
| GTAGCGGCGGCGCTTCAAGG | NDUFS8 | 4728 |
| GTAGCTGAGCGTTGGGCTGT | LTA4H | 4048 |
| GTAGGCGCGCCGCTCTCTAC | NO_CURRENT_720 | NO_CURRENT_720 |
| GTAGGGTACAGCGTCAGCTT | NO_CURRENT_721 | NO_CURRENT_721 |
| GTAGGTGGCGGCTGCGGGCC | PRKN | 5071 |
| GTATAAACATTACCAAGTGG | NO_CURRENT_722 | NO_CURRENT_722 |
| GTATAATTGTCAATGACAAT | NO_CURRENT_723 | NO_CURRENT_723 |
| GTATACAGAGTTAGACCCAC | NO_CURRENT_724 | NO_CURRENT_724 |
| GTATAGCCTGCTGTACAGTT | NO_CURRENT_725 | NO_CURRENT_725 |
| GTATATTATCAGCTAAATGT | NO_CURRENT_726 | NO_CURRENT_726 |
| GTATCCCATATCGGCACAGG | NO_CURRENT_727 | NO_CURRENT_727 |
| GTATCGGCGGCAGCTGTGAG | PITPNB | 23760 |
| GTATCTGGGACCTGTTCTCT | BTN2A2 | 10385 |
| GTATGAGAAAGAACTATCAA | NO_CURRENT_728 | NO_CURRENT_728 |
| GTATTAAGATGCGTCTTAGA | NO_CURRENT_729 | NO_CURRENT_729 |
| GTCAAGAGATTATGAGATTC | NO_CURRENT_730 | NO_CURRENT_730 |
| GTCAATACTAGGCAGATCAA | NO_CURRENT_731 | NO_CURRENT_731 |
| GTCAGAACTCACAGGTTTGC | CD84 | 8832 |
| GTCAGAGCAGGAAGTGTTTG | IKBKB | 3551 |
| GTCAGGAGGGGGCTGCTGCT | FOXO4 | 4303 |
| GTCAGGTAATAGTCGGACTC | NO_CURRENT_732 | NO_CURRENT_732 |
| GTCATATGGGGAACTTCTGT | CHEK2 | 11200 |
| GTCATCAGCGATTTGACGAG | NO_CURRENT_733 | NO_CURRENT_733 |
| GTCATCCTCGGGAGCCCACC | RBM42 | 79171 |
| GTCCAATAAAAAGTGCCACT | NO_CURRENT_734 | NO_CURRENT_734 |
| GTCCAGAGGATCTCTACTCT | GBP7 | 388646 |
| GTCCCAGCACCGGGGAGGAC | IL2RB | 3560 |
| GTCCCAGCCGCTTTCAATGG | NUFIP2 | 57532 |

| | | |
|---|---|---|
| GTCCCCGCGGCTTCTTCCTG | TYK2 | 7297 |
| GTCCCGGGCCCTTACCTAGG | RBM6 | 10180 |
| GTCCCGGTTCCAAGCCCCCT | CBLL1 | 79872 |
| GTCCCGTGATTTTAGCCAGG | NO_CURRENT_735 | NO_CURRENT_735 |
| GTCCCTGGCTGACAACGAAG | MPC2 | 25874 |
| GTCCCTTATTCGCTTGGCCT | UBA7 | 7318 |
| GTCCGGGGTAGCCCCGTTAC | OXSM | 54995 |
| GTCCGGTCGCGTCCGACACC | EZH2 | 2146 |
| GTCCGTCGACCCTTATTGGG | NO_CURRENT_736 | NO_CURRENT_736 |
| GTCCTCAGGAAGAAGCCGCG | TYK2 | 7297 |
| GTCCTCATCCGGTCAGGCTG | NO_CURRENT_737 | NO_CURRENT_737 |
| GTCCTCCTTGCCCAGAAGGT | SQSTM1 | 8878 |
| GTCCTGCTGCTCCAACCCCA | KIF2B | 84643 |
| GTCCTTTCGCAGAGAGGGGA | RIPK3 | 11035 |
| GTCGCACAAAAGAACTGCAT | HTR2A | 3356 |
| GTCGCACGCAGCGCCATGCG | ABCB10 | 23456 |
| GTCGCCCTTCAGCACGCACA | SOD1 | 6647 |
| GTCGCTGCTGCCCGAAGCTG | PDE5A | 8654 |
| GTCGCTTCGGCCAGTGTGTC | HIF1A | 3091 |
| GTCGGCGCGCAGGCGGCTCG | NOD2 | 64127 |
| GTCGGCTCGGAATTGGACTT | RND3 | 390 |
| GTCGGGAGAGACACCAGGCG | PCGF6 | 84108 |
| GTCGGGCGCTCACACCGTGC | EGFR | 1956 |
| GTCGTGCCTCCTGGGTTGTG | PDHA1 | 5160 |
| GTCGTGGACTCGTGCAGCTG | TMEM70 | 54968 |
| GTCTCAAGTGCAGGAGATGG | KMT2D | 8085 |
| GTCTCCGAGTTTGCGACTCG | RHOA | 387 |
| GTCTCTCCCGACCATGGAGG | PCGF6 | 84108 |
| GTCTCTGGAGCCCCCTTGCT | NR1H3 | 10062 |
| GTCTCTGTTTACCACTCGCC | FOXO4 | 4303 |
| GTCTGAGCCCTGCGACAGCC | IRF9 | 10379 |
| GTCTGCGTCTGCAGCTGCAG | RNF25 | 64320 |
| GTCTGGCTGGAGCCACGGGG | CYP24A1 | 1591 |
| GTCTGGTGCTCCTCTCTCAG | NCKAP1L | 3071 |
| GTCTGGTGTCGGCACAGCCA | MPC1 | 51660 |
| GTCTGTCATCTTCTGCCTGG | BCL2A1 | 597 |
| GTCTTGGACTGGGCTCGGTG | SLC7A3 | 84889 |
| GTGAAACAAAGCAGTCGGCT | RND3 | 390 |
| GTGAAACAACGAACCCCCCG | IRGM | 345611 |
| GTGAAACCCACCCTGAGTCA | NO_CURRENT_738 | NO_CURRENT_738 |
| GTGAACTGCAATCTTATTAT | NO_CURRENT_739 | NO_CURRENT_739 |
| GTGAAGAGTCTGGGCAGAGT | PDK4 | 5166 |
| GTGAAGGCCTCGTTGAGAGA | PQBP1 | 10084 |
| GTGAATAATCAAATCATCCC | NO_CURRENT_740 | NO_CURRENT_740 |
| GTGAATTTGGATATATTTAC | BCL2A1 | 597 |
| GTGACATTCAGCCGGCGGTT | ATP5L | 10632 |
| GTGACCCTGTTGGCACAAGA | NO_CURRENT_741 | NO_CURRENT_741 |
| GTGACTGATGTCGTAACTAG | PHC3 | 80012 |
| GTGAGATCCGCGGCTGCCAG | YPEL5 | 51646 |
| GTGAGCGACGCCATAGCGAG | SQSTM1 | 8878 |

| | | |
|---|---|---|
| GTGAGGGGGTTCCGGGAAGA | PITPNB | 23760 |
| GTGAGGGTAGGTGCAGGGCA | RTP1 | 132112 |
| GTGAGGTGGTTGTCGCACTG | NO_CURRENT_742 | NO_CURRENT_742 |
| GTGAGTGGTACCCAACGGGC | PARK7 | 11315 |
| GTGATAATGATGTATTCTCG | NO_CURRENT_743 | NO_CURRENT_743 |
| GTGCAGGCCGAGCTGCGCGG | C16orf72 | 29035 |
| GTGCCAACCATGGAGCGGGC | PDE5A | 8654 |
| GTGCCAGACCTTACCCCTCA | SPCS1 | 28972 |
| GTGCCATCGTGCCGTCTGTC | HAMP | 57817 |
| GTGCCATGTTTAAGTCAATG | NO_CURRENT_744 | NO_CURRENT_744 |
| GTGCCCATGGGAGGGGGGCA | PDE4A | 5141 |
| GTGCCCCCCGAGTGCCCGGC | MEX3B | 84206 |
| GTGCCGACGTCGTGGGTGTT | LGMN | 5641 |
| GTGCCGCGGAGTGAAGAGTC | PDK4 | 5166 |
| GTGCCGGCCTTCCTCGTGTG | MCTS1 | 28985 |
| GTGCGACCCCTTGGTGCTGG | IZUMO2 | 126123 |
| GTGCGACGAATTGTCCTGAG | NO_CURRENT_745 | NO_CURRENT_745 |
| GTGCGAGGCGGCGGAGAGCG | KPNA1 | 3836 |
| GTGCGCATGGGCTGATGTTA | NO_CURRENT_746 | NO_CURRENT_746 |
| GTGCGCGAACATGTAACTGG | IFNAR1 | 3454 |
| GTGCGGCGAGGGCCTACCAG | GPT2 | 84706 |
| GTGCGGGCGCTGTGGCCTGG | PTGES2 | 80142 |
| GTGCGTCATGTGAGCATGTG | NO_CURRENT_747 | NO_CURRENT_747 |
| GTGCGTGCGAGCGGGGGGAG | BMI1 | 648 |
| GTGCTCAAAACATCACAGCC | AGER | 177 |
| GTGCTCACGGGTCATGGCGA | SERF2 | 10169 |
| GTGCTCCACTCAGCATGGTC | PIGY | 84992 |
| GTGCTCCACTCAGCATGGTC | PYURF | 100996939 |
| GTGCTCCGGGCGCTTCAGGC | KIF2B | 84643 |
| GTGCTCGCCCCAGCCCCCAC | ANKRD50 | 57182 |
| GTGCTCTACGTCTTCACCGC | LTB4R | 1241 |
| GTGCTGAGCCATCTCTTTCA | CD84 | 8832 |
| GTGCTGTGCCCAGCGCCTGG | NOD2 | 64127 |
| GTGCTTCAGCTGCGGGCGGT | BNIP3 | 664 |
| GTGGAAGGAGAACTGTCCCA | ABL2 | 27 |
| GTGGATTTGGGTTCACGAAG | NO_CURRENT_748 | NO_CURRENT_748 |
| GTGGCACTCGGCGGTCGAAA | NDUFA8 | 4702 |
| GTGGCCGTGATGGCGGCGGC | ASH2L | 9070 |
| GTGGCCTGGTGGGTGCGCCT | PTGES2 | 80142 |
| GTGGCGAGGCTCTCAAGCCC | FLCN | 201163 |
| GTGGCGAGGCTCTCAAGCCC | PLD6 | 201164 |
| GTGGCGCGTCTGTGCGCGAC | DLAT | 1737 |
| GTGGCGGGCGGCGTGCGGGC | ANP32B | 10541 |
| GTGGCTCACCCCACAGACTG | AGER | 177 |
| GTGGCTCCTGTGTGTGGCGT | PIGL | 9487 |
| GTGGGACTGAGGGGCCCCGG | BAP1 | 8314 |
| GTGGGATCCGAGTGAGGCGA | ATP6V0E1 | 8992 |
| GTGGGGCCATTATTGATACT | NO_CURRENT_749 | NO_CURRENT_749 |
| GTGGGGTGAGTGGTACCCAA | PARK7 | 11315 |
| GTGGGTGCGCCTTGGCCTGG | PTGES2 | 80142 |

| | | |
|---|---|---|
| GTGGTGCGGGCGCTGTGGCC | PTGES2 | 80142 |
| GTGGTGGGAGCGGAGTGGGT | GNAI2 | 2771 |
| GTGGTGTGGGAAATTTAACA | NO_CURRENT_750 | NO_CURRENT_750 |
| GTGTACATTATACAGGTGCT | NO_CURRENT_751 | NO_CURRENT_751 |
| GTGTATGAATGTTAATTCCG | NO_CURRENT_752 | NO_CURRENT_752 |
| GTGTATGATGCTTCGACTTA | NO_CURRENT_753 | NO_CURRENT_753 |
| GTGTCGCCGCTACGATACCT | PQBP1 | 10084 |
| GTGTCTCTCCCGACCATGGA | PCGF6 | 84108 |
| GTGTGAGGATCCGAACCCAG | HSP90B1 | 7184 |
| GTGTGCGTGCGAGCGGGGGG | BMI1 | 648 |
| GTGTGGGACTGAGGGGCCCC | BAP1 | 8314 |
| GTGTGTGCTCTGTGTCACTG | PPARGC1A | 10891 |
| GTGTGTGGCGTTGGCGGTCT | PIGL | 9487 |
| GTGTGTGTGCGTGCGAGCGG | BMI1 | 648 |
| GTGTTATTCCCAGAGGAGTG | TLR1 | 7096 |
| GTGTTCTCATGCGAACGGGA | LARP4B | 23185 |
| GTGTTGAGTACCCAGTTCTT | CCL20 | 6364 |
| GTTAACTAATATCCCAAAGG | NO_CURRENT_754 | NO_CURRENT_754 |
| GTTACCTGCTACGAAAACGA | NO_CURRENT_755 | NO_CURRENT_755 |
| GTTACGGACAAATTGGGCCA | ATP5L | 10632 |
| GTTAGCTCGTTTATTACATG | HTR3E | 285242 |
| GTTATCATCAATTAATCCTG | NO_CURRENT_756 | NO_CURRENT_756 |
| GTTATTCCCAGAGGAGTGTG | TLR1 | 7096 |
| GTTCACTCTCAGTAAGGGGC | PPARGC1A | 10891 |
| GTTCCCCGGGAAGTCTATGC | NO_CURRENT_757 | NO_CURRENT_757 |
| GTTCCCGACTCTCAGCTCCA | CSNK1D | 1453 |
| GTTCCGATGAGGTTAGATGA | NO_CURRENT_758 | NO_CURRENT_758 |
| GTTCCGTGAGGGTTACTTCA | NO_CURRENT_759 | NO_CURRENT_759 |
| GTTCGCTCTCCAGCTTGGGA | IDH2 | 3418 |
| GTTCGCTGCGCCGCGGCCGC | FOS | 2353 |
| GTTCGCTTCGTAACGAGGAA | NO_CURRENT_760 | NO_CURRENT_760 |
| GTTCGTGGTTTCACTGACTC | IRGM | 345611 |
| GTTCTCTCCGCTGTCACTTC | LAMTOR3 | 8649 |
| GTTCTCTGGGGTCCAGTGCT | CYP8B1 | 1582 |
| GTTCTTAGAACTATAGATTA | NO_CURRENT_761 | NO_CURRENT_761 |
| GTTGAGTGAAGAAAATCCAC | EIF4G3 | 8672 |
| GTTGCCTGGTCCTCCTGACT | IL10 | 3586 |
| GTTGGAAACTTTCTCCTCCT | OR7C2 | 26658 |
| GTTGGAACCGGCATAGAGCG | ALOX15 | 246 |
| GTTGGAGGCGCAGCGCCGCG | NNT | 23530 |
| GTTGGCAGTAGTCGCCGCCA | ARNT | 405 |
| GTTGGCATATTGGCCCAGAC | NO_CURRENT_762 | NO_CURRENT_762 |
| GTTGGCCCGTTCTTATGTCC | NO_CURRENT_763 | NO_CURRENT_763 |
| GTTGGGGATCTGGAGTCCCG | LARP4B | 23185 |
| GTTGGGGTGAGTACGACCTC | CSDE1 | 7812 |
| GTTGTCTGTGTGTGGGACTG | BAP1 | 8314 |
| GTTGTGAGAGGGAATATCGA | NO_CURRENT_764 | NO_CURRENT_764 |
| GTTGTGGGGCCCGGGACCGG | CREBBP | 1387 |
| GTTGTTATTTCCAAGGAATG | NO_CURRENT_765 | NO_CURRENT_765 |
| GTTGTTCGTTTCTCCCCGGG | IRGM | 345611 |

| | | |
|---|---|---|
| GTTTAAAGAAAGGGGCTAAG | NO_CURRENT_766 | NO_CURRENT_766 |
| GTTTAACCCAGGCGAGGAGG | ARMCX2 | 9823 |
| GTTTACTCATATCCAGTCAC | NO_CURRENT_767 | NO_CURRENT_767 |
| GTTTCCATACATGTCCTACT | NO_CURRENT_768 | NO_CURRENT_768 |
| GTTTCGAAACTTGAAGTAAG | NO_CURRENT_769 | NO_CURRENT_769 |
| GTTTCGCCACCGGAGCGGCC | NFKB1 | 4790 |
| GTTTCGCTGCGTCCCCTTCG | TP73 | 7161 |
| GTTTCGCTTTCCCGAAGAGT | CHMP5 | 51510 |
| GTTTCTCCAAACACAAATAC | NO_CURRENT_770 | NO_CURRENT_770 |
| GTTTCTCTTGAACCCCGCGA | NO_CURRENT_771 | NO_CURRENT_771 |
| GTTTCTTCCCTGCCTCCTTG | IKBKE | 9641 |
| GTTTGACCGAGGCATTAAGG | IRAK4 | 51135 |
| GTTTGAGGCCGGAGGGAGCG | SPHK1 | 8877 |
| GTTTGCCTGCAGCAAGATGG | NDUFS4 | 4724 |
| GTTTGGCCTCAAAAGAAACG | MCL1 | 4170 |
| GTTTTCAGTTGCCCAACAGC | NO_CURRENT_772 | NO_CURRENT_772 |
| GTTTTTGGTTAATTGCCTAC | NO_CURRENT_773 | NO_CURRENT_773 |
| TAAAAACTTTATAAACCCCC | HMGN2 | 3151 |
| TAAAAAGCCCCATTGGAGTG | DYRK1A | 1859 |
| TAAAAAGCGGGGAGCTCGGC | TNRC18 | 84629 |
| TAAAAGGGACTCCCGGAGCT | CFLAR | 8837 |
| TAAAATGAGGACATTAGTGT | NO_CURRENT_774 | NO_CURRENT_774 |
| TAAACCCTTGGCGCCCTCAC | NO_CURRENT_775 | NO_CURRENT_775 |
| TAAAGACGCTTCTTCCCGGC | IFNAR2 | 3455 |
| TAAAGCAGAAGAATATACAG | NO_CURRENT_776 | NO_CURRENT_776 |
| TAAATTCATACATATCGATG | NO_CURRENT_777 | NO_CURRENT_777 |
| TAACAAAATTTGGACAGGGT | AGER | 177 |
| TAACAACCAGGGAGCCACAC | C4orf17 | 84103 |
| TAACAAGATAGCTCAAAACA | NO_CURRENT_778 | NO_CURRENT_778 |
| TAACAGCACTCCCAAAGAAC | CCL20 | 6364 |
| TAACCCAGAAGCCCATTCAG | NO_CURRENT_779 | NO_CURRENT_779 |
| TAACCCAGGCGAGGAGGAGG | ARMCX2 | 9823 |
| TAACCGATACTCCCCACATT | NO_CURRENT_780 | NO_CURRENT_780 |
| TAACCTTCTGTGGAAAAGCA | CD84 | 8832 |
| TAACTCGCTAGGCCACGCCG | SOD1 | 6647 |
| TAACTGGGGGTCTTCTTGGC | UQCRB | 7381 |
| TAAGACGCCGGAGGAGGTGG | GLMN | 11146 |
| TAAGCAGAATGCAGAAATGC | PPARG | 5468 |
| TAAGCAGGCCGGTAAGTAAC | UQCRB | 7381 |
| TAAGCTCTCTGAAAAGAAGG | AIM2 | 9447 |
| TAAGGCATAATAATAGACTC | NO_CURRENT_781 | NO_CURRENT_781 |
| TAAGGCATGATTAGAGGTCT | NAIP | 4671 |
| TAAGGCGACCTGCGCTTGTG | NO_CURRENT_782 | NO_CURRENT_782 |
| TAAGGTGCCAGAGTGTCAAG | NO_CURRENT_783 | NO_CURRENT_783 |
| TAAGTAACTGGGGGTCTTCT | UQCRB | 7381 |
| TAAGTCTAATATATCCAAAG | NO_CURRENT_784 | NO_CURRENT_784 |
| TAATAAAATCCAGGTAGCGC | TNRC18 | 84629 |
| TAATAAGCCAAAGCATGCGT | NO_CURRENT_785 | NO_CURRENT_785 |
| TAATAATAACTACCCAAACC | NO_CURRENT_786 | NO_CURRENT_786 |
| TAATAATAGTTGCCTTAATG | NO_CURRENT_787 | NO_CURRENT_787 |

| | | |
|---|---|---|
| TAATAGGGCAGGGAACCCTG | NCKAP1L | 3071 |
| TAATAGTTATGGCAGAGATC | ABL2 | 27 |
| TAATATGATTTGAGCACCAG | NO_CURRENT_788 | NO_CURRENT_788 |
| TAATATTGATTGTTAAAATG | NO_CURRENT_789 | NO_CURRENT_789 |
| TAATCATGCACATTCGGGAC | NO_CURRENT_790 | NO_CURRENT_790 |
| TAATCCATTTAATCCCAATG | NO_CURRENT_791 | NO_CURRENT_791 |
| TAATGCTGCACACGCCGAAT | NO_CURRENT_792 | NO_CURRENT_792 |
| TACAAAGACTCTGACACTTC | AIM2 | 9447 |
| TACAACTCCCACAAGGCCTA | TFPT | 29844 |
| TACAATCGTCTTGGCTATAC | NO_CURRENT_793 | NO_CURRENT_793 |
| TACAGCCCAAAGCCCCGCCA | KDM4A | 9682 |
| TACAGTCACCAAAACAGTTT | NO_CURRENT_794 | NO_CURRENT_794 |
| TACATAGTTTCTACAAACCA | NO_CURRENT_795 | NO_CURRENT_795 |
| TACATTATAAAAGACACTGT | NO_CURRENT_796 | NO_CURRENT_796 |
| TACCAGCGCGCGGGCGGCGG | NR2C2 | 7182 |
| TACCCAGAGTAGAGATCCTC | GBP7 | 388646 |
| TACCCATCATGGAAGCAATG | PIGL | 9487 |
| TACCCCAGCACACCCTCTCA | STAT2 | 6773 |
| TACCCCATTAGGTATTAAGC | NO_CURRENT_797 | NO_CURRENT_797 |
| TACCCCCGCCACGGATCGCC | MAP3K7 | 6885 |
| TACCCCGGCTGCGCACAGCT | DNAJA1 | 3301 |
| TACCCGGCGATCCGTGGCGG | MAP3K7 | 6885 |
| TACCCTCCGGATACGGACTG | NO_CURRENT_798 | NO_CURRENT_798 |
| TACCCTGGATTGTCCTTGCG | NO_CURRENT_799 | NO_CURRENT_799 |
| TACCGAACAAGCGATGATGC | RAD21 | 5885 |
| TACCGCATCCGCGTGTCCAC | ALOX15 | 246 |
| TACCGGACGGGCTGGGTCTA | CCNC | 892 |
| TACCGGGAGAGGCGGCAACA | KMT2C | 58508 |
| TACCTGTAACGGGGCTACCC | OXSM | 54995 |
| TACGCTTGCGTTTAGCGTCC | NO_CURRENT_800 | NO_CURRENT_800 |
| TACGTCATTAAGAGTTCAAC | NO_CURRENT_801 | NO_CURRENT_801 |
| TACTACCCGGCGATCCGTGG | MAP3K7 | 6885 |
| TACTCAGACAGCTTAATGCT | NO_CURRENT_802 | NO_CURRENT_802 |
| TACTCAGATGGGAGCGCCCG | RRAGA | 10670 |
| TACTCATACACAAAAATCAG | NO_CURRENT_803 | NO_CURRENT_803 |
| TACTCCATGCAGTTGTCAGG | IKZF5 | 64376 |
| TACTCGGCCCCGCCGGTCCC | CREBBP | 1387 |
| TACTGAGCCGCCGCCGCACG | PTGIS | 5740 |
| TACTGAGGCAGACGTTGTGG | NDUFS4 | 4724 |
| TACTGGAGCACTCAGGCCCT | CXCR4 | 7852 |
| TACTGGAGTTTGCGACTCGG | NO_CURRENT_804 | NO_CURRENT_804 |
| TACTGTTGGACACACCTGGC | CXCR1 | 3577 |
| TAGAAAAATGAAGAGGGTCC | GC | 2638 |
| TAGAATAAGTGAGAAAGTGG | DPY30 | 84661 |
| TAGAATAAGTGAGAAAGTGG | MEMO1 | 51072 |
| TAGAATTTGACCAAAGGCAC | NO_CURRENT_805 | NO_CURRENT_805 |
| TAGACAACCGCGGAGAATGC | NO_CURRENT_806 | NO_CURRENT_806 |
| TAGACTATACAGAATACATG | NO_CURRENT_807 | NO_CURRENT_807 |
| TAGAGAGCTGGAGAGGCAGG | GPR119 | 139760 |
| TAGAGATATCCGATCGTGGT | NO_CURRENT_808 | NO_CURRENT_808 |

| | | |
|---|---|---|
| TAGAGGACAGCGGGGAAGGC | MTOR | 2475 |
| TAGAGGTTACTCACACGGCT | SLC25A19 | 60386 |
| TAGAGTGCATAAGAGAACCA | NO_CURRENT_809 | NO_CURRENT_809 |
| TAGATCGAGTTTATTTTCCT | NO_CURRENT_810 | NO_CURRENT_810 |
| TAGATGTTAACAGAGTGATC | NO_CURRENT_811 | NO_CURRENT_811 |
| TAGCACCTGCGCGTTGGCGG | PLD1 | 5337 |
| TAGCAGAGGCGCTGCCGGTG | LRP2 | 4036 |
| TAGCAGCAGCAGCAACAGTG | PTGIS | 5740 |
| TAGCAGCGCGCAGCCCGCAG | CACNA2D2 | 9254 |
| TAGCAGGATCGGTCCACAGC | TFPT | 29844 |
| TAGCCAGCGATACCTGTAAC | OXSM | 54995 |
| TAGCCGGGCCTTCTAAGCCC | SPCS1 | 28972 |
| TAGCGAAAGCGGAGCTCGTC | ATP5E | 514 |
| TAGCGGCGGCGCTTCAAGGT | NDUFS8 | 4728 |
| TAGCGGGCGACGGAATCAGA | SNRNP70 | 6625 |
| TAGCTATACATATTACAACG | NO_CURRENT_812 | NO_CURRENT_812 |
| TAGCTATGGCTGGTGAGTCT | CASP5 | 838 |
| TAGGAGCTGTATCTAGTGGC | NO_CURRENT_813 | NO_CURRENT_813 |
| TAGGAGGAATGGTATACTAG | NO_CURRENT_814 | NO_CURRENT_814 |
| TAGGCCAAAACCCTACTTCT | NDUFA1 | 4694 |
| TAGGCTCTCGGTGAGCTGGG | NQO1 | 1728 |
| TAGGGCAGGGAACCCTGAGG | NCKAP1L | 3071 |
| TAGGGGAGCAGCGGCAGTGG | RNF111 | 54778 |
| TAGGGGATTAGCTGACAGTC | NO_CURRENT_815 | NO_CURRENT_815 |
| TAGGGTCTGGCTGGAGCCAC | CYP24A1 | 1591 |
| TAGGTAACGGGGCAGAGATG | NDUFA1 | 4694 |
| TAGGTGAGGCGGCGGCTGAG | FBXO38 | 81545 |
| TAGTACAAAATACTGATGCA | NO_CURRENT_816 | NO_CURRENT_816 |
| TAGTACATGTGTGGTATTTA | NO_CURRENT_817 | NO_CURRENT_817 |
| TAGTAGGTACGCTAAAGGCA | NO_CURRENT_818 | NO_CURRENT_818 |
| TAGTATGGGCTAGAAGTAGA | ARR3 | 407 |
| TAGTATTTATTAGTGACCAC | NO_CURRENT_819 | NO_CURRENT_819 |
| TAGTCAACATTCGCAAGAGG | NO_CURRENT_820 | NO_CURRENT_820 |
| TAGTCCTTAGGGTGGGCTGA | NO_CURRENT_821 | NO_CURRENT_821 |
| TAGTCTCAGTAGATACCATG | NO_CURRENT_822 | NO_CURRENT_822 |
| TAGTGAATGCTCCAGCAAGG | HCAR2 | 338442 |
| TAGTGAGAACACGGAGAGTG | RTP1 | 132112 |
| TAGTGCAGGTCCTGAAACGC | NDUFA12 | 55967 |
| TAGTGCTCATGCTCGCTGCA | SAP18 | 10284 |
| TAGTGGAGGCTGCTAGGAGC | UBE2H | 7328 |
| TAGTGGGAATGGTCGCGTAG | NO_CURRENT_823 | NO_CURRENT_823 |
| TAGTTCTAATCGTTCCTTGA | NO_CURRENT_824 | NO_CURRENT_824 |
| TATAACATCATATACCACAT | NO_CURRENT_825 | NO_CURRENT_825 |
| TATACCAGACCACAGCGCCG | NO_CURRENT_826 | NO_CURRENT_826 |
| TATACTGCGGATCAATCTGA | NO_CURRENT_827 | NO_CURRENT_827 |
| TATAGCTGTTTCGAAGGCGC | NO_CURRENT_828 | NO_CURRENT_828 |
| TATATCCTGGAGCGAGTGCT | METTL3 | 56339 |
| TATCAATCGTCCGGGTCACT | NO_CURRENT_829 | NO_CURRENT_829 |
| TATCAGTAGAGGTAAACTCT | NO_CURRENT_830 | NO_CURRENT_830 |
| TATCATTAGACTCCCACACT | NO_CURRENT_831 | NO_CURRENT_831 |

| | | |
|---|---|---|
| TATCGCTTCCGATTAGTCCG | NO_CURRENT_832 | NO_CURRENT_832 |
| TATCGGCGGCAGCTGTGAGG | PITPNB | 23760 |
| TATCTCGGGCATGGCTCTGG | LTA4H | 4048 |
| TATGACCCTGTTACATTGCC | NO_CURRENT_833 | NO_CURRENT_833 |
| TATGATCACTGGAGTCTCGC | BAK1 | 578 |
| TATGGCGTCGCTCACCGTGA | SQSTM1 | 8878 |
| TATGGCTCCTAATATTCGGG | NO_CURRENT_834 | NO_CURRENT_834 |
| TATGGGGGTACAATTAATGT | NO_CURRENT_835 | NO_CURRENT_835 |
| TATGTAACAATTAGTAAACC | NO_CURRENT_836 | NO_CURRENT_836 |
| TATTCGGGCGGGAAGCGAAA | DYRK1A | 1859 |
| TATTGGAGCAGCAAGAGGCT | MMP1 | 4312 |
| TATTGTGAGGCGGTTGTAGA | CASP3 | 836 |
| TATTTCCCATAGACATGGCA | SERPINB2 | 5055 |
| TATTTTGACTTGACGCAGGC | NO_CURRENT_837 | NO_CURRENT_837 |
| TCAAACCCCAGCCCAGGCCG | PDK4 | 5166 |
| TCAAACGCATACATATAATG | NO_CURRENT_838 | NO_CURRENT_838 |
| TCAAATATGGGTAACACTTG | NO_CURRENT_839 | NO_CURRENT_839 |
| TCAACCACAATAACAGGCGG | FBXO38 | 81545 |
| TCAACCCCTACCTGGACCCT | SLC3A2 | 6520 |
| TCAACGCTGGCTGCCGCTGG | UCHL3 | 7347 |
| TCAAGACTTTGCTCTCCACC | BCL2A1 | 597 |
| TCAAGATGAACCGACTCTTC | CHMP5 | 51510 |
| TCAAGATGAGAGTTCAGCCG | POU2F1 | 5451 |
| TCAAGCGCAGCCAGTCCCCG | TYK2 | 7297 |
| TCAATCACCCAGAAACAATG | NO_CURRENT_840 | NO_CURRENT_840 |
| TCAATTCTCACTCACGACCA | NO_CURRENT_841 | NO_CURRENT_841 |
| TCACACCGTGCGGGGGGCGG | EGFR | 1956 |
| TCACACTGGTAAGGAGGCGG | CBLL1 | 79872 |
| TCACAGAACAAGAAGGTATC | NLRC4 | 58484 |
| TCACATACCTGGGAGACCCT | HSH2D | 84941 |
| TCACCATTCAAGATGCATCC | MMP13 | 4322 |
| TCACCCCAGTCAGGAGGACC | IL10 | 3586 |
| TCACCTCCGAACGAACACCT | NO_CURRENT_842 | NO_CURRENT_842 |
| TCACGAGGGCGCCCATTCCC | RIPK2 | 8767 |
| TCACGTTTCCAATTAGGCAC | NO_CURRENT_843 | NO_CURRENT_843 |
| TCACTCTTACCTGCCAACCT | CUBN | 8029 |
| TCACTGCCAATTTAATCACA | NO_CURRENT_844 | NO_CURRENT_844 |
| TCACTTCTAATATTCGGCCG | PIAS1 | 8554 |
| TCACTTGTCTATACGGGGTA | NO_CURRENT_845 | NO_CURRENT_845 |
| TCACTTTCTATCCTATACTG | NO_CURRENT_846 | NO_CURRENT_846 |
| TCAGAAACATGCTGAAGTCC | ADRB1 | 153 |
| TCAGAAGGGAGGGTTGCGCC | RND1 | 27289 |
| TCAGACCTCTCCAGGCCCTG | NLRP1 | 22861 |
| TCAGAGAGCTTAACTAGCAG | AIM2 | 9447 |
| TCAGATGCTCGCTGCAGATG | FOS | 2353 |
| TCAGATTCCGCAAGGGTCCA | NO_CURRENT_847 | NO_CURRENT_847 |
| TCAGCAAAGGACGAAACAAA | NO_CURRENT_848 | NO_CURRENT_848 |
| TCAGCATGGTCAATGAACAC | NO_CURRENT_849 | NO_CURRENT_849 |
| TCAGCCAAAGATTCTCTCGA | NO_CURRENT_850 | NO_CURRENT_850 |
| TCAGCCCACGTCTGAGGGAC | SPHK1 | 8877 |

| | | |
|---|---|---|
| TCAGCCTCTTTATTACAGAG | NO_CURRENT_851 | NO_CURRENT_851 |
| TCAGCCTGCACCCCCTCTCT | SLC11A1 | 6556 |
| TCAGCTCCTGCTGTCGCCAC | KDM4A | 9682 |
| TCAGCTGCCGCCGCCAGCAC | MTOR | 2475 |
| TCAGGCCGGCTCCGGCTCAG | HSPA4 | 3308 |
| TCAGGCTTAAAACTTGTTGG | RND3 | 390 |
| TCAGGGACAGCTTTAAAGAC | LAMTOR3 | 8649 |
| TCAGGGCCCGTACCTGATTA | STAT2 | 6773 |
| TCAGGGGAGCTGGCAGAACC | FLCN | 201163 |
| TCAGGGGAGCTGGCAGAACC | PLD6 | 201164 |
| TCAGGTAGACCTGGCGACGA | PRR14L | 253143 |
| TCAGGTGAGGGCTCGGCCCG | APEH | 327 |
| TCAGTATCAAACCGCTTCTC | FBXW7 | 55294 |
| TCATATGGGGAACTTCTGTT | CHEK2 | 11200 |
| TCATCATCAACCTCCCAGGT | CUBN | 8029 |
| TCATCCAAAAACATGAATGG | NO_CURRENT_852 | NO_CURRENT_852 |
| TCATCTCCAGCGCGTGGTGG | STAG2 | 10735 |
| TCATCTCGGGCAGAGCGCTA | FOSL2 | 2355 |
| TCATCTGTCAATCAACTACT | NO_CURRENT_853 | NO_CURRENT_853 |
| TCATCTTACATCTGGGAGAC | NO_CURRENT_854 | NO_CURRENT_854 |
| TCATGAAGCACAGGCAGTGG | PDHA1 | 5160 |
| TCATGCTTGCTTGGGCAAAA | NO_CURRENT_855 | NO_CURRENT_855 |
| TCATGGTCTCAAGCTACATG | NO_CURRENT_856 | NO_CURRENT_856 |
| TCATTCACACTTGATTAATG | ABL2 | 27 |
| TCCAAAACTCTTTCTTCCCA | CASP1 | 834 |
| TCCAAATGGGCTGCTGGCGG | KPNB1 | 3837 |
| TCCAAGCTCCAAACTCCCGC | ERCC6L | 54821 |
| TCCAGCAGTGCCTGAGAGCG | PHF6 | 84295 |
| TCCAGCCGCCTGTCCGGGTG | NFKB2 | 4791 |
| TCCAGCGGAGTTGTGGGGGC | DNTTIP1 | 116092 |
| TCCAGCTCCGCGGGGCAGTG | NLRP3 | 114548 |
| TCCAGCTCCTGCTCGCCGGA | GSDMD | 79792 |
| TCCAGCTTTCCCAGCAGAAT | ANKRD50 | 57182 |
| TCCAGTGCTGGGAGCTCTGC | CYP8B1 | 1582 |
| TCCAGTTACTTTCAGGCTCG | PQBP1 | 10084 |
| TCCAGTTGTGAGAGCCGCAA | CYP1B1 | 1545 |
| TCCCAAGGGTTTAAGTCGGG | NO_CURRENT_857 | NO_CURRENT_857 |
| TCCCAATTTAGGTCAGTTCA | NO_CURRENT_858 | NO_CURRENT_858 |
| TCCCACCACCAGCCCTCTGC | GNAI2 | 2771 |
| TCCCACCGCCACAAGGAGGC | IKBKE | 9641 |
| TCCCAGCTCTGTTAACTGCG | SATB2 | 23314 |
| TCCCCACCGCTGCCCCCACC | BAP1 | 8314 |
| TCCCCAGACCTTTCAGTTGT | PAGR1 | 79447 |
| TCCCCCAGATTCGGGCAGGT | ARMCX2 | 9823 |
| TCCCCCCGGTGTCGCTGCCC | CCAR2 | 57805 |
| TCCCCGAGACCATCTTAGGG | NO_CURRENT_859 | NO_CURRENT_859 |
| TCCCCGAGAGGGCGGCGAGA | SLC25A19 | 60386 |
| TCCCCGAGCCTGAAAGTAAC | PQBP1 | 10084 |
| TCCCCGCCATCTTCCAGCCG | PPARGC1B | 133522 |
| TCCCCGGCCAAGCCCAACTC | NFKB2 | 4791 |

| | | |
|---|---|---|
| TCCCCGTACGGCGCTTACCT | ZRSR2 | 8233 |
| TCCCCTCACACGAGGAAGGC | MCTS1 | 28985 |
| TCCCCTGAGAGCCTGAACCC | FLCN | 201163 |
| TCCCCTGAGAGCCTGAACCC | PLD6 | 201164 |
| TCCCGCAGAGGGCTGGTGGT | GNAI2 | 2771 |
| TCCCGCCCTCCTGACGCGGC | EP400 | 57634 |
| TCCCGGAAGCAGATTGCGCC | ZNF616 | 90317 |
| TCCCGGGCCCCGAACCCCAG | SIAH2 | 6478 |
| TCCCGGGTTCGCACCTGGTC | VDR | 7421 |
| TCCCGGTTGGTGAACGATAC | NO_CURRENT_860 | NO_CURRENT_860 |
| TCCCGTGAGTAACTTGGCTC | ANP32B | 10541 |
| TCCCTCTCAACCACAATAAC | FBXO38 | 81545 |
| TCCCTGAGCTTCTCCTCCTT | CYP4B1 | 1580 |
| TCCCTGCATTCATGGTTTTA | NO_CURRENT_861 | NO_CURRENT_861 |
| TCCCTGCCTCCTTGTGGCGG | IKBKE | 9641 |
| TCCGAGGAGGAGCAGTCCCG | SCTR | 6344 |
| TCCGAGTGAGGCGACGGGGT | ATP6V0E1 | 8992 |
| TCCGATGCCATCATGCAGCC | CDKN2C | 1031 |
| TCCGCCGCACGCTGCAGCCG | PPARGC1B | 133522 |
| TCCGCCGCCAGCCGCCTCCC | BCL7A | 605 |
| TCCGCCGCCAGCCGCCTCCC | SETD1B | 23067 |
| TCCGCCGGCCCGTTCATGAC | INO80E | 283899 |
| TCCGCGCAGTTAACAGAGCT | SATB2 | 23314 |
| TCCGCGGCGGGCGCAGGATA | PIP4K2A | 5305 |
| TCCGGCCACCCGAGGCCGCG | MAP2K5 | 5607 |
| TCCGGGTGTCTTTGTCCCCC | CCAR2 | 57805 |
| TCCGTAGGCTCAGGTAGACC | PRR14L | 253143 |
| TCCGTGAGGCCCTGCCGGGT | NCOA6 | 23054 |
| TCCTACCAGCGCGCGGGCGG | NR2C2 | 7182 |
| TCCTACCTGAAGCACCGGCC | CXCR1 | 3577 |
| TCCTACTACCCGGCGATCCG | MAP3K7 | 6885 |
| TCCTAGCAGAGGGGGAGAGG | CYP4F11 | 57834 |
| TCCTCAAGGTGGGGTTCGGG | HNRNPF | 3185 |
| TCCTCACCTAAAGTGCAATA | NO_CURRENT_862 | NO_CURRENT_862 |
| TCCTCCTACCAGCGCGCGGG | NR2C2 | 7182 |
| TCCTCCTGGTCTTGTCGATG | RTP1 | 132112 |
| TCCTCCTGTCCCCTCAGACG | HIF1A | 3091 |
| TCCTCGAGGGCTGAGGCTCT | HSH2D | 84941 |
| TCCTGAGGAGATGTTCAGGG | PAXIP1 | 22976 |
| TCCTGCGGCGGTGCCGGCTG | CASP9 | 842 |
| TCCTGCTGCTCCAACCCCAA | KIF2B | 84643 |
| TCCTGGAGCTGCAGACAGTG | JAK1 | 3716 |
| TCCTGTTTGTTGTTGGAGAA | DNAJB6 | 10049 |
| TCCTTGCCTCAGGCCTCTCG | SUPT7L | 9913 |
| TCCTTGCTGGAGCATTCACT | HCAR2 | 338442 |
| TCCTTGCTGGAGCATTCACT | HCAR3 | 8843 |
| TCCTTGTCATGGGAGCATTG | MBNL1 | 4154 |
| TCCTTTAGACAACTTCAGGA | NAIP | 4671 |
| TCCTTTAGGCATTGCGGCTC | PDAP1 | 11333 |
| TCCTTTCATTCGAGGAGAGA | PHF6 | 84295 |

| | | |
|---|---|---|
| TCCTTTCGCAGAGAGGGGAA | RIPK3 | 11035 |
| TCGAATTCACTTCTAATATT | PIAS1 | 8554 |
| TCGAGAGGAAAAACACACTG | NO_CURRENT_863 | NO_CURRENT_863 |
| TCGATGTAGCCCCGCCCAAG | NO_CURRENT_864 | NO_CURRENT_864 |
| TCGCAACCAGAGCCTCACCT | SBNO2 | 22904 |
| TCGCAACTGGGCCCGCGCGC | TIRAP | 114609 |
| TCGCAAGACGCCCGCACCTG | CYP2R1 | 120227 |
| TCGCACCTGGTCCGGCCGGC | VDR | 7421 |
| TCGCACTCCGAGTCTTCCCC | INO80C | 125476 |
| TCGCATGCTCGCCATCATGT | UBE2L6 | 9246 |
| TCGCCATGGACGAAGCGGAT | CASP9 | 842 |
| TCGCCCACGTCCATGGTACC | CTNNBL1 | 56259 |
| TCGCCTGCTGGAAAAGCAGT | KDM4A | 9682 |
| TCGCGCTCGAAGCCCCGGTC | CREBBP | 1387 |
| TCGCGGACCATGGGCGACAA | ARRB1 | 408 |
| TCGCGTCAGACCCCGAAAGC | SUPT7L | 9913 |
| TCGCTCCCGTTCCCCGGACG | SATB1 | 6304 |
| TCGCTGGGGCCCGCGCGGCT | RPS6KA4 | 8986 |
| TCGCTGGGTGCCGCGTCTGC | SATB2 | 23314 |
| TCGGAAAAAGGGTAACAACC | RIPK3 | 11035 |
| TCGGAACCAGGACCTCGGCG | SOD1 | 6647 |
| TCGGACCCGAAGCCGCCACA | SLC25A1 | 6576 |
| TCGGAGTCCGCGCCGGGCTG | BNIP3 | 664 |
| TCGGCAAATACCAATAATGA | NO_CURRENT_865 | NO_CURRENT_865 |
| TCGGCAGCTCCAACCCAACT | SLC7A3 | 84889 |
| TCGGCATACGGGACACACGC | NO_CURRENT_866 | NO_CURRENT_866 |
| TCGGCCCGGCAGACTGCGCT | PIP4K2A | 5305 |
| TCGGCGATCAGCGGCGGCGG | C16orf72 | 29035 |
| TCGGCGGCTGATTTAGAAGG | BABAM1 | 29086 |
| TCGGCTACGGCGTGGAGAAG | NO_CURRENT_867 | NO_CURRENT_867 |
| TCGGGAAACGTCATCTTCTC | ZRSR2 | 8233 |
| TCGGGAAGCTGTGCGAGAAG | CXCL3 | 2921 |
| TCGGGAGCCCACCCGGACGA | RBM42 | 79171 |
| TCGGGCAGTGAGTACAATAC | NO_CURRENT_868 | NO_CURRENT_868 |
| TCGGGCCGCGCTGCGTGCGC | AKT1 | 207 |
| TCGGGCGCTCACACCGTGCG | EGFR | 1956 |
| TCGGGCTTGGGCGCCCCCGG | IZUMO2 | 126123 |
| TCGGGGACCACCCACGATCC | NO_CURRENT_869 | NO_CURRENT_869 |
| TCGGTAACCGGCGACTACGT | BABAM2 | 9577 |
| TCGGTCTGTGGCAGCAGCGT | CTNNB1 | 1499 |
| TCGGTGGCCAGCCAGTGAGC | NR1H3 | 10062 |
| TCGGTTCTCTCGTTAGTCCA | RHOA | 387 |
| TCGTAAACACACGACCAAGT | NO_CURRENT_870 | NO_CURRENT_870 |
| TCGTCCCTGGCTGACAACGA | MPC2 | 25874 |
| TCTAACGAGACTGTTACACT | NO_CURRENT_871 | NO_CURRENT_871 |
| TCTAACGTAAAATCCTCCAG | NO_CURRENT_872 | NO_CURRENT_872 |
| TCTACCCGACTTAACAGGTA | ABL2 | 27 |
| TCTACGTGTAGTTGTACATA | NO_CURRENT_873 | NO_CURRENT_873 |
| TCTAGAAAAGAAGTCAGCTC | RIPK2 | 8767 |
| TCTAGACATATTATGAGGCT | C4orf17 | 84103 |

| | | |
|---|---|---|
| TCTAGGAACTGAGAGGAAGA | BCORL1 | 63035 |
| TCTATTACACAAATTGACCT | NO_CURRENT_874 | NO_CURRENT_874 |
| TCTATTGTACAAAGTTAGGA | ACOD1 | 730249 |
| TCTATTTTGTCTGCGCAGAA | NO_CURRENT_875 | NO_CURRENT_875 |
| TCTGAAAAATAGGCCCAACC | NO_CURRENT_876 | NO_CURRENT_876 |
| TCTGACCATTGGGTGCGACA | NO_CURRENT_877 | NO_CURRENT_877 |
| TCTGACGCGATCCTTGCCTC | SUPT7L | 9913 |
| TCTGAGGAGGTCGAGGGTCC | JAM2 | 58494 |
| TCTGATCGTTGATCTTTCTG | NO_CURRENT_878 | NO_CURRENT_878 |
| TCTGCCAGGCGCGATCCCCC | STAT1 | 6772 |
| TCTGCGCACCGGCCGGGCCT | SLC7A5 | 8140 |
| TCTGCTGAAAAAGACAACTG | AIM2 | 9447 |
| TCTGCTTCTTGCGGCCGAAC | CHMP6 | 79643 |
| TCTGGAACTACAGATAAGGC | NO_CURRENT_879 | NO_CURRENT_879 |
| TCTGGACCTCGAGAGGCCTG | SUPT7L | 9913 |
| TCTGGCGCGTCTGCTCTCCC | GSDMD | 79792 |
| TCTGGCTGCTCCGCGGAGGG | TGFB1 | 7040 |
| TCTGGCTTGACACGACCGTT | NO_CURRENT_880 | NO_CURRENT_880 |
| TCTGGGCGCCACTCCCGGGC | INO80E | 283899 |
| TCTGGGTAAGTCCAGCTCCG | NLRP3 | 114548 |
| TCTGGGTCATGGTCTGGTTC | CYP27B1 | 1594 |
| TCTGTCCCGCCGCCCGCGCC | NCOA6 | 23054 |
| TCTGTCGGCTCCGTAGGCTC | PRR14L | 253143 |
| TCTGTCTCTTTGTACAGCAC | CASP5 | 838 |
| TCTGTGCCACCCCAGGAGCT | IKBKE | 9641 |
| TCTGTGGTCTGATCTTCCTG | ATP5E | 514 |
| TCTGTTCCGAGAGCGTGCCC | AHR | 196 |
| TCTTACCTGAGTTGCTGTCC | IRF9 | 10379 |
| TCTTATTAATTGAGGGACCT | NO_CURRENT_881 | NO_CURRENT_881 |
| TCTTCAAATGGTGGGGACAG | KMT2D | 8085 |
| TCTTCACACTGCTCGGGCTC | HMGN2 | 3151 |
| TCTTCATCTCCCAGGTATGG | C6orf15 | 29113 |
| TCTTCCCAGGGACCTGTTCT | CARD16 | 114769 |
| TCTTCCCAGGGACCTGTTCT | CASP1 | 834 |
| TCTTCCCTGCCTCCTTGTGG | IKBKE | 9641 |
| TCTTCCTCAAGGTGGGGTTC | HNRNPF | 3185 |
| TCTTCTTTCTAGAACAACCG | NO_CURRENT_882 | NO_CURRENT_882 |
| TCTTGAGCTGGACTCATTGT | MMP13 | 4322 |
| TCTTGCTGCAGGCAAACGCC | NDUFS4 | 4724 |
| TCTTGCTGGCTGGTGCCCCA | NLRP1 | 22861 |
| TGAAAACCCCAATAGACAGG | NO_CURRENT_883 | NO_CURRENT_883 |
| TGAAAAGAAGGTGGCAGACC | AIM2 | 9447 |
| TGAAAATCCTATTGCTCAGT | NO_CURRENT_884 | NO_CURRENT_884 |
| TGAAATACCTGGTTGTTCTC | SERPINB2 | 5055 |
| TGAAATATATGCAAAAATTG | NO_CURRENT_885 | NO_CURRENT_885 |
| TGAAATCCTGGCCTCTTAGG | GBP7 | 388646 |
| TGAACCCCAAAGGCCAGAAC | TNFRSF1A | 7132 |
| TGAACCGGCCGCCACGTCCG | PHF8 | 23133 |
| TGAACCGGGGAAGGCAGACC | NO_CURRENT_886 | NO_CURRENT_886 |
| TGAACGGTGAAGAGATAGGG | NO_CURRENT_887 | NO_CURRENT_887 |

| | | |
|---|---|---|
| TGAACTGTAACAACTCTCAG | SERPINB2 | 5055 |
| TGAACTTCATCTCTCTCCCC | ARR3 | 407 |
| TGAACTTCGATTCAGAAGGG | RND1 | 27289 |
| TGAAGACAGCTGTATTCTTT | C4orf17 | 84103 |
| TGAAGCACAATACCAATACT | NO_CURRENT_888 | NO_CURRENT_888 |
| TGAAGCGAGACCCATCGTCC | NO_CURRENT_889 | NO_CURRENT_889 |
| TGAAGCGGCGGTACCGGTGC | MTOR | 2475 |
| TGAAGCTGAGGAGGCGGCGG | HTT | 3064 |
| TGAAGCTGAGGAGGCGGCGG | KHSRP | 8570 |
| TGAAGGTGACCCCCCTGGGG | SPINT1 | 6692 |
| TGAAGGTGTCTATACACTGT | NO_CURRENT_890 | NO_CURRENT_890 |
| TGAATCGAATACAAACGATG | NO_CURRENT_891 | NO_CURRENT_891 |
| TGAATCGTAACCTCGCCATT | NO_CURRENT_892 | NO_CURRENT_892 |
| TGAATGGTGGAGCCGAAGCT | PHIP | 55023 |
| TGACACATTGGCTGGGTGTT | NO_CURRENT_893 | NO_CURRENT_893 |
| TGACAGTGGCTTTGCCCGTT | MSL2 | 55167 |
| TGACAGTTGCTATAGCGACC | DYRK1A | 1859 |
| TGACATTCAGCCGGCGGTTC | ATP5L | 10632 |
| TGACCTCTGAGGAATTCACA | NO_CURRENT_894 | NO_CURRENT_894 |
| TGACGCGATAGAGTTGGCTT | NO_CURRENT_895 | NO_CURRENT_895 |
| TGACGCGGCTCTGCTTCTTG | CHMP6 | 79643 |
| TGACGTGGCCCACAACTGAA | PAGR1 | 79447 |
| TGACGTTTCCCGAGAAACCA | ZRSR2 | 8233 |
| TGACTAGCAGCTGGGTACTC | CASP5 | 838 |
| TGACTCGGGCAATATCGGTT | NO_CURRENT_896 | NO_CURRENT_896 |
| TGACTGATGTCGTAACTAGG | PHC3 | 80012 |
| TGACTGCTCGGAGTTCTCCC | TLR2 | 7097 |
| TGAGAGAATGCATCACCATG | NO_CURRENT_897 | NO_CURRENT_897 |
| TGAGAGCACGATGCATACAC | SLC7A11 | 23657 |
| TGAGAGCCCCCCATCTTGTG | NO_CURRENT_898 | NO_CURRENT_898 |
| TGAGAGCGAGAGGTGGATCG | UCHL5 | 51377 |
| TGAGAGCGGGGCTCTGTCGC | PHF6 | 84295 |
| TGAGAGCTCCTGGGCAGGCT | HELZ2 | 85441 |
| TGAGAGGCCCCGGCAGGTCC | IRAK1 | 3654 |
| TGAGAGTCGGGAACAGGTAC | CSNK1D | 1453 |
| TGAGCATGTCGGGAGTAACT | NO_CURRENT_899 | NO_CURRENT_899 |
| TGAGCATTCGTAGCCCAGCA | NO_CURRENT_900 | NO_CURRENT_900 |
| TGAGCCGGCCCCGCTGGGGA | SATB1 | 6304 |
| TGAGCCTACCTTCTGCCTCC | TMEM173 | 340061 |
| TGAGCCTACGGAGCCGACAG | PRR14L | 253143 |
| TGAGCGCGGGGGTCCCAGGG | HELZ2 | 85441 |
| TGAGCGGCCTCTAATTAATC | NO_CURRENT_901 | NO_CURRENT_901 |
| TGAGCGTGGAAGAAGGGACG | FAM214B | 80256 |
| TGAGCTGGTTTGTAATGATA | SLC7A11 | 23657 |
| TGAGGAAACGGGGACCTGCC | LTB4R | 1241 |
| TGAGGATTGAGGTGTATGAA | NO_CURRENT_902 | NO_CURRENT_902 |
| TGAGGCGACGGGGTAGGGGT | ATP6V0E1 | 8992 |
| TGAGGCGGCCACCGCGACTC | JAM3 | 83700 |
| TGAGGGTCTGGGTCATGGTC | CYP27B1 | 1594 |
| TGAGGTCACCCTCCAGGCTG | GBP5 | 115362 |

| | | |
|---|---|---|
| TGAGGTGCTCGGAGCCTCGG | ABI1 | 10006 |
| TGAGTAAGCTGTGGCGGCGT | INO80B | 83444 |
| TGAGTCTTACTAGGTCCTGT | NO_CURRENT_903 | NO_CURRENT_903 |
| TGAGTGCAAGGTGACTGGGA | ATP5J | 522 |
| TGAGTGCTGTGCCCAGCGCC | NOD2 | 64127 |
| TGAGTGGTACCCAACGGGCC | PARK7 | 11315 |
| TGAGTGTCCGGCCCCAGGTA | ATP5J2 | 9551 |
| TGAGTGTCCGGCCCCAGGTA | ATP5J2-PTCD1 | 100526740 |
| TGAGTGTCCGGCCCCAGGTA | PTCD1 | 26024 |
| TGATCGAGGAGCGCGGTTAC | CCNC | 892 |
| TGATCTCTGCCATAACTATT | ABL2 | 27 |
| TGATGCAATGGAAGGTGTGC | NLRP12 | 91662 |
| TGATGCGGGCCCAAGCTGCA | RAD21 | 5885 |
| TGATGGCCTAGTGATGTCAT | NO_CURRENT_904 | NO_CURRENT_904 |
| TGATTCCAAGATGCCCGCGA | MBNL1 | 4154 |
| TGATTCTATTGTACAAAGTT | ACOD1 | 730249 |
| TGATTGGAGCCGGGTGCCGC | BAK1 | 578 |
| TGATTTCCATGCCATGTCTA | SERPINB2 | 5055 |
| TGCAAAGAGTGAAAGAACAA | GBP5 | 115362 |
| TGCAACAACTCGCCGCGCCG | RXRA | 6256 |
| TGCAACAGGTCATAAATACA | NO_CURRENT_905 | NO_CURRENT_905 |
| TGCAACCGAGAGCGCTGCTC | HSPA4 | 3308 |
| TGCAAGTCTGTCTTGTGGTT | IL10 | 3586 |
| TGCAATGTCTGTGCCACCCC | IKBKE | 9641 |
| TGCACAGTGGCAGTAACACT | NO_CURRENT_906 | NO_CURRENT_906 |
| TGCACCAGAGGCCGCGCCGA | CS | 1431 |
| TGCACCAGTGCCCAGAGAGG | SLC11A1 | 6556 |
| TGCACCGGGGGGATCGCGCC | STAT1 | 6772 |
| TGCACTGAGAAAGAAGACAA | MMP1 | 4312 |
| TGCAGAGCACGGTTCCCATA | TBXAS1 | 6916 |
| TGCAGCCTGGTTAGGAGCAA | CDKN2C | 1031 |
| TGCAGCTACATGATGGGCCA | MAP2K6 | 5608 |
| TGCAGCTCCAGGATACTCCG | JAK1 | 3716 |
| TGCAGGAATTCAGCTGCTGG | SERPINE1 | 5054 |
| TGCAGGAGGCCTCGGCTCGT | CAT | 847 |
| TGCAGGCAGCCCCAGCCTCC | IZUMO2 | 126123 |
| TGCAGGCCGAGCTGCGCGGA | C16orf72 | 29035 |
| TGCAGGGACGACGACGACGA | NUP153 | 9972 |
| TGCATGCCGAGCATTTTCAA | NO_CURRENT_907 | NO_CURRENT_907 |
| TGCATGTTGTGTGAGGATCC | RAD23A | 5886 |
| TGCCAACCCCGGTGAGGAGA | ARNT | 405 |
| TGCCAGTCGCCGCCGCCGCC | MCL1 | 4170 |
| TGCCATAACTATTAGGTGGA | ABL2 | 27 |
| TGCCATAACTTAGAAACCGG | NO_CURRENT_908 | NO_CURRENT_908 |
| TGCCATCATGCAGCCTGGTT | CDKN2C | 1031 |
| TGCCCAGACTCTTCACTCCG | PDK4 | 5166 |
| TGCCCGGCGCCCGCCCCTTG | CDKN2D | 1032 |
| TGCCCTCCCGCCCCATGGAG | CHUK | 1147 |
| TGCCGACACCAGACCCCGAG | MPC1 | 51660 |
| TGCCGCACCGGACCCAGTGC | ACTR5 | 79913 |

| | | |
|---|---|---|
| TGCCGCGGAGTGAAGAGTCT | PDK4 | 5166 |
| TGCCGGACTGACGGGCGGCC | MAP2K7 | 5609 |
| TGCCGTGAAAAGACGCTGCG | NO_CURRENT_909 | NO_CURRENT_909 |
| TGCCGTTAGCATGCGATCCC | NO_CURRENT_910 | NO_CURRENT_910 |
| TGCCTAAAGGAGGTGAGGGG | PDAP1 | 11333 |
| TGCCTACGGTTCCTAACCCT | LAMTOR2 | 28956 |
| TGCCTCTCCCTTACCCGGAC | NO_CURRENT_911 | NO_CURRENT_911 |
| TGCCTGCAGCAAGATGGCGG | NDUFS4 | 4724 |
| TGCCTTACCTGCCCGGGCGC | IRAK4 | 51135 |
| TGCCTTCTTACCAAGTCCAG | GBP7 | 388646 |
| TGCGAACGGGAAGGAGCGTT | LARP4B | 23185 |
| TGCGACCCCGGTGCCCGGTG | PTGS1 | 5742 |
| TGCGCCCGCAGAAACGCGCC | RAC2 | 5880 |
| TGCGCGAACATGTAACTGGT | IFNAR1 | 3454 |
| TGCGCGCATGGGCCTGTTCC | PRKN | 5071 |
| TGCGCTCGCTTAGCGGGCGA | SNRNP70 | 6625 |
| TGCGCTGCGCCACGCGTAGC | NMU | 10874 |
| TGCGGAGAGTGTGCGGCTCC | CHEK2 | 11200 |
| TGCGGAGATTGGAGGCCGCG | APIP | 51074 |
| TGCGGCCCGTGCAGCCCCGG | CACNA2D2 | 9254 |
| TGCGGGATGTGCTTCAGCTG | BNIP3 | 664 |
| TGCGGGCCTGCGCCCAGTCC | JAK1 | 3716 |
| TGCGGGCGCAAGGGGTGTGG | RAC2 | 5880 |
| TGCGGGCGCTGGGCCGGCGG | INO80 | 54617 |
| TGCGGGCGCTGTGGCCTGGT | PTGES2 | 80142 |
| TGCGGGCGGTGGGAAAGCGG | BNIP3 | 664 |
| TGCGTCTGCAGCTGCAGGGG | RNF25 | 64320 |
| TGCGTGGAGGAACTGCTGGG | ACACA | 31 |
| TGCGTGGTCGCACCCTACCC | NMRK2 | 27231 |
| TGCGTTTAACCCAGGCGAGG | ARMCX2 | 9823 |
| TGCTAAGCAGAGAATCGCTG | TFPT | 29844 |
| TGCTACCTTCGGGACCACCA | NO_CURRENT_912 | NO_CURRENT_912 |
| TGCTATTAGTGTGCACCTAG | NO_CURRENT_913 | NO_CURRENT_913 |
| TGCTCACTCCACTCCTCAAC | NO_CURRENT_914 | NO_CURRENT_914 |
| TGCTCCAACCCCAAGGGCCC | KIF2B | 84643 |
| TGCTCGCCGGACGGCTCCCA | GSDMD | 79792 |
| TGCTCGGCGGCGGCCCGAGG | FAM107B | 83641 |
| TGCTCGGTCTGGGGTCTGCC | STAT1 | 6772 |
| TGCTCGTTCGCTCTCCAGCT | IDH2 | 3418 |
| TGCTCTCCCTGGGAGCCGTC | GSDMD | 79792 |
| TGCTCTGCCAGCCAGTGGGA | TIFA | 92610 |
| TGCTCTGTGTCACTGTGGAT | PPARGC1A | 10891 |
| TGCTCTTGGAGAGGTCACTC | DLAT | 1737 |
| TGCTGCGCTGGTGGTGGCGG | DNAJB6 | 10049 |
| TGCTGCTGCTCCCGGGGCTG | KMT2C | 58508 |
| TGCTGCTGCTGCCCGGGAGG | SETD1B | 23067 |
| TGCTGCTGCTGCTGCCCGGG | SETD1B | 23067 |
| TGCTGGAAAAGCAGTAGGAT | KDM4A | 9682 |
| TGCTGGAGGCGGCGGAACCG | VIPR2 | 7434 |
| TGCTGGATAGGTTGCAGGCA | NO_CURRENT_915 | NO_CURRENT_915 |

| | | |
|---|---|---|
| TGCTTATGGCGGCGCTGGAG | CSDE1 | 7812 |
| TGCTTCCAGCGAATCTACAC | SLC5A8 | 160728 |
| TGCTTCCATACAGCTTAGAG | NO_CURRENT_916 | NO_CURRENT_916 |
| TGCTTCCCGCAGAGGGCTGG | GNAI2 | 2771 |
| TGGAAAGAGTGGTTTCACAT | NO_CURRENT_917 | NO_CURRENT_917 |
| TGGAAAGCGAGCACACCGTC | NO_CURRENT_918 | NO_CURRENT_918 |
| TGGAACAACTGTGGAACCTG | CHUK | 1147 |
| TGGAACAGTAAAAGACTCAC | CARD16 | 114769 |
| TGGAACAGTAAAAGACTCAC | CASP1 | 834 |
| TGGAACGTCAGGTGAGGGCT | APEH | 327 |
| TGGAAGCCGATGCCCGGCAG | RAC1 | 5879 |
| TGGACCCTAGCCAGCCCTCA | SMPD1 | 6609 |
| TGGACGCATGCCCTCAGGTG | CYP2R1 | 120227 |
| TGGACGCTGTGAATCGTGGC | STRAP | 11171 |
| TGGACGTGACTGCTCTATCC | APAF1 | 317 |
| TGGACTAACGAGAGAACCGA | RHOA | 387 |
| TGGACTTGGGAGGCGCGGTG | RND3 | 390 |
| TGGAGAGCAAAGTCTTGAGC | BCL2A1 | 597 |
| TGGAGAGCGAACGAGCAGGG | IDH2 | 3418 |
| TGGAGAGCGAGATCCGGAGT | NFKB2 | 4791 |
| TGGAGAGGAACAGCCCATGG | OR7C2 | 26658 |
| TGGAGAGGTCACTCCGGAGA | DLAT | 1737 |
| TGGAGAGGTCTGAAGAAACC | NLRP1 | 22861 |
| TGGAGCTGAGAGTCGGGAAC | CSNK1D | 1453 |
| TGGAGCTTGATCAGAAACCC | NO_CURRENT_919 | NO_CURRENT_919 |
| TGGAGGAAAGCTGTGCATAC | MMP1 | 4312 |
| TGGAGGAACTGCTGGGCGGG | ACACA | 31 |
| TGGAGGCGTACTTGAGGGTC | CYP27B1 | 1594 |
| TGGAGTCAAGTGTAATGGCG | NO_CURRENT_920 | NO_CURRENT_920 |
| TGGAGTGGCCGAGTTCGCCA | SLC5A8 | 160728 |
| TGGAGTTGAAAAAGCTTGAC | PPARGC1A | 10891 |
| TGGAGTTTGGGCGGCCCGGG | PPID | 5481 |
| TGGATGATGACGAGGAGAGA | TMEM173 | 340061 |
| TGGATGGGTGAATTAACGGG | NO_CURRENT_921 | NO_CURRENT_921 |
| TGGCAATTAGCACACAACAA | NO_CURRENT_922 | NO_CURRENT_922 |
| TGGCACTCGGCGGTCGAAAG | NDUFA8 | 4702 |
| TGGCACTGAGCTCCCAGATC | HAMP | 57817 |
| TGGCACTGCGCGTCAGTAGC | SPCS1 | 28972 |
| TGGCATCGGAGACTGACAGA | CDKN2C | 1031 |
| TGGCATTACATTTATTGATC | NO_CURRENT_923 | NO_CURRENT_923 |
| TGGCCACGAATTCCGCCGCC | NO_CURRENT_924 | NO_CURRENT_924 |
| TGGCCACGGTCCCAGCACCG | IL2RB | 3560 |
| TGGCCAGCAATTCTAGACCA | NO_CURRENT_925 | NO_CURRENT_925 |
| TGGCCCACAACTGAAAGGTC | PAGR1 | 79447 |
| TGGCCCTCACCCCAGTCAGG | IL10 | 3586 |
| TGGCCGGTCAGCGTCGCTGC | PPID | 5481 |
| TGGCCGTGAAAGTCGGGCTT | NDUFS8 | 4728 |
| TGGCCTGGCCTGTCAGGGCG | RRAGC | 64121 |
| TGGCGCGGAGGCTGAAGCTG | KHSRP | 8570 |
| TGGCGCGGCGGAGACCCGGC | TBK1 | 29110 |

| | | |
|---|---|---|
| TGGCGGAGGTGCAGACCTAA | LRP2 | 4036 |
| TGGCGTCCGCAGCTGGGGCT | TRAF6 | 7189 |
| TGGCGTTGGCGGTCTTGGCA | PIGL | 9487 |
| TGGCTGGCCACCGAGACTTC | NR1H3 | 10062 |
| TGGCTTCCTACCTGAAGCAC | CXCR1 | 3577 |
| TGGGAACGCGGGCGGCGAGA | IRF8 | 3394 |
| TGGGACAAGCTGCTTCTGAT | CYP4F11 | 57834 |
| TGGGAGATACGCACAGTCGA | NO_CURRENT_926 | NO_CURRENT_926 |
| TGGGAGCACCTCTGGGGCTA | INO80B | 83444 |
| TGGGAGCGGGGAGGTCGCGG | UTY | 7404 |
| TGGGAGTGTGAGCGGTGCTC | SLC11A1 | 6556 |
| TGGGATATTGGAGCAGCAAG | MMP1 | 4312 |
| TGGGATCCGAGTGAGGCGAC | ATP6V0E1 | 8992 |
| TGGGATGCTTTGTTGTCTGG | BTN2A2 | 10385 |
| TGGGATTCAGCAATATGTGT | NO_CURRENT_927 | NO_CURRENT_927 |
| TGGGCAGAGGTATGGTCCTT | ABL2 | 27 |
| TGGGCCCCGCGCCGCGGCGG | NENF | 29937 |
| TGGGCCGGGTACCATGGACG | CTNNBL1 | 56259 |
| TGGGCCTCAGGGCTGCGCTC | EP400 | 57634 |
| TGGGCGACAGACACAGTCCC | ACADS | 35 |
| TGGGCGGGCGCGGGCGCGCG | ABL1 | 25 |
| TGGGCGGGCGCGGGCGCGCG | PXN | 5829 |
| TGGGCTATAGATTCCATGTG | NO_CURRENT_928 | NO_CURRENT_928 |
| TGGGGACGTTTATCAATATA | NO_CURRENT_929 | NO_CURRENT_929 |
| TGGGGAGCGAGCGAGCGACC | BRD1 | 23774 |
| TGGGGAGCGCACGACGTACC | TLR2 | 7097 |
| TGGGGCGCCTCTAAAGGGGA | ARNT | 405 |
| TGGGGCGTCCTCCTGCTGGC | SERPINA2 | 390502 |
| TGGGGCTCAGCAGACGCGTC | CXCL3 | 2921 |
| TGGGGGCAGAGCTCTAGCGG | MEX3B | 84206 |
| TGGGGGCGGGTTATTCTGTT | MGA | 23269 |
| TGGGGGCGTAGCGCGCTGAG | WASF2 | 10163 |
| TGGGGTCTATTGATGAAGGC | NO_CURRENT_930 | NO_CURRENT_930 |
| TGGGGTGAGTGGTACCCAAC | PARK7 | 11315 |
| TGGGGTGGGCTCCGCGGTGC | C16orf72 | 29035 |
| TGGGTAAATCAAGTTAACAC | NO_CURRENT_931 | NO_CURRENT_931 |
| TGGGTAAGTCCAGCTCCGCG | NLRP3 | 114548 |
| TGGGTCGGTGGCGGAGGCTG | RBM6 | 10180 |
| TGGGTTCCCAGAGTGCTCCG | KHSRP | 8570 |
| TGGGTTGGAGCTGCCGAGTT | SLC7A3 | 84889 |
| TGGTAACCAGGTACATGGAG | OR7C2 | 26658 |
| TGGTACTGAGGCAGACGTTG | NDUFS4 | 4724 |
| TGGTAGGCCCTCGCCGCACT | GPT2 | 84706 |
| TGGTATCGGCGGCAGCTGTG | PITPNB | 23760 |
| TGGTCCCGGGCTGGGCAGGC | SLC6A4 | 6532 |
| TGGTCCGGGCTTGAGCTCCT | MLF2 | 8079 |
| TGGTCTGGCAGCCGGAGACC | PIGY | 84992 |
| TGGTCTGGCAGCCGGAGACC | PYURF | 100996939 |
| TGGTGAAGGTAACCGCGTAT | UBE2L6 | 9246 |
| TGGTGAATCTCCCGCTGCAC | BRK1 | 55845 |

| | | |
|---|---|---|
| TGGTGACTGATGTCGTAACT | PHC3 | 80012 |
| TGGTGCCCCAGGGCCTGGAG | NLRP1 | 22861 |
| TGGTGCCCTCCCCGAGGCGG | RHOG | 391 |
| TGGTGCTCCACCTGGTGCCC | NMU | 10874 |
| TGGTGGGCCCCGCGCCGCGG | NENF | 29937 |
| TGGTGGTGGGTTTGTTATAT | NO_CURRENT_932 | NO_CURRENT_932 |
| TGGTGTATGATAATGCATCA | NO_CURRENT_933 | NO_CURRENT_933 |
| TGGTGTCGGCACAGCCATGG | MPC1 | 51660 |
| TGGTTAGGAGCAAAGGAAAG | CDKN2C | 1031 |
| TGGTTCGGAGAAGCAGCGGC | RIPK2 | 8767 |
| TGTAACATATCTTAACAACC | C4orf17 | 84103 |
| TGTAACCAAGATCATAAGAC | NO_CURRENT_934 | NO_CURRENT_934 |
| TGTAATAAACGAGCTAACTA | HTR3E | 285242 |
| TGTAGATATAGGGTGTCTAC | NO_CURRENT_935 | NO_CURRENT_935 |
| TGTAGCTAAGTGAGTATGCC | NO_CURRENT_936 | NO_CURRENT_936 |
| TGTAGCTTTGAAAGTCACCT | AIM2 | 9447 |
| TGTAGGTGACCATGTAAAAG | GC | 2638 |
| TGTATGTGTGTCCCGAGAAG | FBXW7 | 55294 |
| TGTCAATGTCAGTGGTACTG | NDUFS4 | 4724 |
| TGTCCTCTCACAGAACAAGA | NLRC4 | 58484 |
| TGTCGCCATGGCGGCGGCAG | ARHGAP33 | 115703 |
| TGTCGCCATGGCGGCGGCAG | LRRC47 | 57470 |
| TGTCGCTGCTGCCCGAAGCT | PDE5A | 8654 |
| TGTCTCTCCCGACCATGGAG | PCGF6 | 84108 |
| TGTCTCTGGAATCCTCCAAG | CYP4A11 | 1579 |
| TGTCTGGCTGTCCCACTGCT | HAMP | 57817 |
| TGTCTGTGTGTGGGACTGAG | BAP1 | 8314 |
| TGTCTTGGTCTCCCAAGAAG | NLRC4 | 58484 |
| TGTGAAACAACGAACCCCCC | IRGM | 345611 |
| TGTGAAAGTTTATGGTGTTA | NO_CURRENT_937 | NO_CURRENT_937 |
| TGTGACGGTGACGAAGACGG | IKZF5 | 64376 |
| TGTGAGAAGATTTAGTGACT | NO_CURRENT_938 | NO_CURRENT_938 |
| TGTGAGAGCCGCAAGGGCAT | CYP1B1 | 1545 |
| TGTGCACAAGTCGCAACGAA | NO_CURRENT_939 | NO_CURRENT_939 |
| TGTGCCCTGAGGGCCGCGAG | NNT | 23530 |
| TGTGCGAGAAGCGGGAGAGC | CXCL3 | 2921 |
| TGTGCGTGCGAGCGGGGGGA | BMI1 | 648 |
| TGTGCTCAGCTCCAGCTCAC | NR1H3 | 10062 |
| TGTGCTTCTGGGCAGAGGTA | ABL2 | 27 |
| TGTGGCACTCGGCGGTCGAA | NDUFA8 | 4702 |
| TGTGGCCAGCGGCAACCAAG | MKL1 | 57591 |
| TGTGGCCTCCGCCCTAGGTG | AGER | 177 |
| TGTGGCGCGTCTGTGCGCGA | DLAT | 1737 |
| TGTGGCGGCGGGCTCGGAGG | VDAC1 | 7416 |
| TGTGGGACTGAGGGGCCCCG | BAP1 | 8314 |
| TGTGGGCCCAGGTATCGTAG | PQBP1 | 10084 |
| TGTGTGACCTTCATTATATG | NO_CURRENT_940 | NO_CURRENT_940 |
| TGTGTGGGACTGAGGGGCCC | BAP1 | 8314 |
| TGTGTGTGCGTGCGAGCGGG | BMI1 | 648 |
| TGTGTGTGTTTCAGTCTACG | ZNF616 | 90317 |

| | | |
|---|---|---|
| TGTGTTAGCCGAGATCTCTG | NO_CURRENT_941 | NO_CURRENT_941 |
| TGTGTTCACGAGCCGAGCTT | PHIP | 55023 |
| TGTTAAAATCACCCGGTCTG | NO_CURRENT_942 | NO_CURRENT_942 |
| TGTTATTCCCAGAGGAGTGT | TLR1 | 7096 |
| TGTTCCAAGAACAGGTCCCT | CARD16 | 114769 |
| TGTTCCAAGAACAGGTCCCT | CASP1 | 834 |
| TGTTCTGAGCTGGCGGTCCG | AHR | 196 |
| TGTTGAGCCGGGCAGTGTGC | SOD2 | 6648 |
| TGTTGAGTACCCAGTTCTTT | CCL20 | 6364 |
| TGTTTGCACTGGCTACAGCA | UBA7 | 7318 |
| TGTTTGTCTGCCGGACTGAC | MAP2K7 | 5609 |
| TGTTTTGCATGTTGCATAGG | NO_CURRENT_943 | NO_CURRENT_943 |
| TTAAAAAGCGGGGAGCTCGG | TNRC18 | 84629 |
| TTAAAGAGACAGAGGCAGCA | ABI1 | 10006 |
| TTAAATATAGATTCTGCCAA | NO_CURRENT_944 | NO_CURRENT_944 |
| TTAACAATGTACCGTCTGGT | NO_CURRENT_945 | NO_CURRENT_945 |
| TTAACACAGTGTTAAGAACA | NO_CURRENT_946 | NO_CURRENT_946 |
| TTACAACGAAATGATGCTCA | ACOD1 | 730249 |
| TTACAACTTTGTGCTGGTGC | KDM6A | 7403 |
| TTACAACTTTGTGCTGGTGC | UTY | 7404 |
| TTACACCCATTATCACAGGG | NO_CURRENT_947 | NO_CURRENT_947 |
| TTACAGTCTTACATGAGAGG | NO_CURRENT_948 | NO_CURRENT_948 |
| TTACCGAACAAGCGATGATG | RAD21 | 5885 |
| TTACTGTATTCTACATGCCC | NO_CURRENT_949 | NO_CURRENT_949 |
| TTACTTTCTTGTTCGAGCAT | NO_CURRENT_950 | NO_CURRENT_950 |
| TTAGAAGGAGGTTCAGGCTA | BABAM1 | 29086 |
| TTAGAGGACAGCGGGGAAGG | MTOR | 2475 |
| TTAGCAGGATCGGTCCACAG | TFPT | 29844 |
| TTAGCCCTCGATTGGTTGCG | NO_CURRENT_951 | NO_CURRENT_951 |
| TTAGCCGTGGGGAAAGATGG | SUV39H1 | 6839 |
| TTAGCCTTGAGCTCAACTGG | CCL20 | 6364 |
| TTAGCTATGGCTGGTGAGTC | CASP5 | 838 |
| TTAGCTCGTTTATTACATGA | HTR3E | 285242 |
| TTAGGTGTAAGGGCTGCTTA | NO_CURRENT_952 | NO_CURRENT_952 |
| TTAGTATGGGCTAGAAGTAG | ARR3 | 407 |
| TTAGTGCAGGTCCTGAAACG | NDUFA12 | 55967 |
| TTATATAAATAACCAAACCA | NO_CURRENT_953 | NO_CURRENT_953 |
| TTATGAGAGTCAGAAATGAT | NO_CURRENT_954 | NO_CURRENT_954 |
| TTATGGCGGCGCTGGAGAGG | CSDE1 | 7812 |
| TTATGGGAACCGTGCTCTGC | TBXAS1 | 6916 |
| TTATTAAAAAGCGGGGAGCT | TNRC18 | 84629 |
| TTATTCCTTTGAGCACATTG | NO_CURRENT_955 | NO_CURRENT_955 |
| TTATTCGGGCGGGAAGCGAA | DYRK1A | 1859 |
| TTATTGTAGAGTGTTGCTCA | NO_CURRENT_956 | NO_CURRENT_956 |
| TTCAATCACCTCACGGTAAG | NO_CURRENT_957 | NO_CURRENT_957 |
| TTCACAACATGAAATCGCAC | NO_CURRENT_958 | NO_CURRENT_958 |
| TTCACACTGCTCGGGCTCCG | HMGN2 | 3151 |
| TTCACAGATGCATAAGCCAG | NO_CURRENT_959 | NO_CURRENT_959 |
| TTCACAGGCAGCGGGCGGAG | NLRP12 | 91662 |
| TTCACCGTCCACGTGCGCAT | NO_CURRENT_960 | NO_CURRENT_960 |

| | | |
|---|---|---|
| TTCACGTCGTCTCGCGACCA | NO_CURRENT_961 | NO_CURRENT_961 |
| TTCACTCATGTTTGCACCCG | COX16 | 51241 |
| TTCACTCTTGTCGACCAAGG | NO_CURRENT_962 | NO_CURRENT_962 |
| TTCACTGCTGCGTCGCAGAG | SUPT7L | 9913 |
| TTCACTTATTCTTGACAAAG | NO_CURRENT_963 | NO_CURRENT_963 |
| TTCAGACCTCTCCAGGCCCT | NLRP1 | 22861 |
| TTCAGTTCACTCTCAGTAAG | PPARGC1A | 10891 |
| TTCATGCCGGTGCGGAGGGG | NMRK2 | 27231 |
| TTCATTTAAAGCTTGTGTCT | NLRC4 | 58484 |
| TTCCAAAACTCTTTCTTCCC | CASP1 | 834 |
| TTCCAATAGTACCAAATACT | NO_CURRENT_964 | NO_CURRENT_964 |
| TTCCAATCCCGGGCTGTATA | TNRC18 | 84629 |
| TTCCACGGTAAAATCGGTCA | NO_CURRENT_965 | NO_CURRENT_965 |
| TTCCATTGGCTGGAATCTGA | NO_CURRENT_966 | NO_CURRENT_966 |
| TTCCCACACTCACCCGGTAA | EDA2R | 60401 |
| TTCCCATAAGTATAATCTTG | NO_CURRENT_967 | NO_CURRENT_967 |
| TTCCCCCAGTTGCAACGTTA | MAP2K6 | 5608 |
| TTCCCGCCCCAGAAGTTCAA | JAM2 | 58494 |
| TTCCCTAACGTTGCAACTGG | MAP2K6 | 5608 |
| TTCCCTTTCCTTAGGTGAGT | PPARG | 5468 |
| TTCCTCAAGGTGGGGTTCGG | HNRNPF | 3185 |
| TTCCTCACCTGCGTCCGCGG | CYFIP1 | 23191 |
| TTCCTGCCCGAACTGCAGAA | NO_CURRENT_968 | NO_CURRENT_968 |
| TTCCTGGGGATCCTCAACCA | SERPINA2 | 390502 |
| TTCCTGTCCAGAAGTCTCGG | NR1H3 | 10062 |
| TTCCTTCCAGTAAGGAGTCG | MCL1 | 4170 |
| TTCGACCGCCGAGTGCCACA | NDUFA8 | 4702 |
| TTCGAGGTCCGGACAGGTCG | NO_CURRENT_969 | NO_CURRENT_969 |
| TTCGCAATAAATAATATACC | NO_CURRENT_970 | NO_CURRENT_970 |
| TTCGCACCTGGTCCGGCCGG | VDR | 7421 |
| TTCGCACGATTGCACCTTGG | NO_CURRENT_971 | NO_CURRENT_971 |
| TTCGCTAGCAGAGGCGCTGC | LRP2 | 4036 |
| TTCGGAACTTACTCAGGGTA | NO_CURRENT_972 | NO_CURRENT_972 |
| TTCGGCCAGTGTGTCGGGCT | HIF1A | 3091 |
| TTCGGTATGATGTTGATTGT | NO_CURRENT_973 | NO_CURRENT_973 |
| TTCGTAGGAACTAAACTGTA | NO_CURRENT_974 | NO_CURRENT_974 |
| TTCGTCCGGGTGGGCTCCCG | RBM42 | 79171 |
| TTCGTGGTAGGTATAACTAT | NO_CURRENT_975 | NO_CURRENT_975 |
| TTCTAAGCGCCCTGGGGACA | NO_CURRENT_976 | NO_CURRENT_976 |
| TTCTCCGCGAGGGGCCTCCG | CACNA2D2 | 9254 |
| TTCTCCGGGTCTGCCTTCCC | NO_CURRENT_977 | NO_CURRENT_977 |
| TTCTCCTAATCCATGAACAG | NO_CURRENT_978 | NO_CURRENT_978 |
| TTCTCCTCCTCCTCCTCGCC | ARMCX2 | 9823 |
| TTCTCTCCGCTGTCACTTCA | LAMTOR3 | 8649 |
| TTCTCTGAGTTGCTGTCTGA | SERPINB2 | 5055 |
| TTCTGAGTCCCACCGGGTGT | EZH2 | 2146 |
| TTCTGCGGGCGCAAGGGGTG | RAC2 | 5880 |
| TTCTTGCTGGCTGGTGCCCC | NLRP1 | 22861 |
| TTCTTTAAACCGGCGCTACC | TNRC18 | 84629 |
| TTCTTTAGCTGGTCAAAAGG | NO_CURRENT_979 | NO_CURRENT_979 |

| | | |
|---|---|---|
| TTGACCGAGGCATTAAGGCG | IRAK4 | 51135 |
| TTGACCTAAAATATGGGTGT | NO_CURRENT_980 | NO_CURRENT_980 |
| TTGACTGACATGAATAAATG | NO_CURRENT_981 | NO_CURRENT_981 |
| TTGAGATTGAGACTGGGGGT | CYP1B1 | 1545 |
| TTGAGTTGAGTGATGATCAT | NO_CURRENT_982 | NO_CURRENT_982 |
| TTGATCAAAGAGGTGCACGT | NO_CURRENT_983 | NO_CURRENT_983 |
| TTGCAAAGCTGATCGGCTGT | NO_CURRENT_984 | NO_CURRENT_984 |
| TTGCAATGCTGCTATAGAAG | NO_CURRENT_985 | NO_CURRENT_985 |
| TTGCACCAGTGCCCAGAGAG | SLC11A1 | 6556 |
| TTGCAGGTGGACCACCCAGG | TMEM30A | 55754 |
| TTGCATCATTCACAGGCAGC | NLRP12 | 91662 |
| TTGCCCTGCCCCCCTCCCAT | PDE4A | 5141 |
| TTGCCGGCGTCTTGGCGATT | ATP5E | 514 |
| TTGCCTGGTCCTCCTGACTG | IL10 | 3586 |
| TTGCGCTGACAGACGCAAGA | PIAS1 | 8554 |
| TTGCGGCGGAGGGCAGGAAG | TRERF1 | 55809 |
| TTGCTGTCCAGGCTGTCGCA | IRF9 | 10379 |
| TTGGACACCAAACATCACTT | NO_CURRENT_986 | NO_CURRENT_986 |
| TTGGACTGGGCTCGGTGAGG | SLC7A3 | 84889 |
| TTGGAGATGATAGTTTGGGG | NO_CURRENT_987 | NO_CURRENT_987 |
| TTGGAGGCCGCGCGGGTCCC | APIP | 51074 |
| TTGGAGTTGCTTGCGGCGGA | TRERF1 | 55809 |
| TTGGATTACTAGCTTTAGGT | NO_CURRENT_988 | NO_CURRENT_988 |
| TTGGCCAAAGTTCCAGGTGT | HTR1D | 3352 |
| TTGGCCGTGAAAGTCGGGCT | NDUFS8 | 4728 |
| TTGGGCGCCCCCGGAGGCTG | IZUMO2 | 126123 |
| TTGGGGTGAGTACGACCTCA | CSDE1 | 7812 |
| TTGGGTTCCAGTTACCCCGG | ERCC6L | 54821 |
| TTGGTAGTGTGTATAAACAC | NO_CURRENT_989 | NO_CURRENT_989 |
| TTGGTCCGAGTCTGGAGAAA | NO_CURRENT_990 | NO_CURRENT_990 |
| TTGGTTTAAGCTTAAAAGTG | NO_CURRENT_991 | NO_CURRENT_991 |
| TTGGTTTACACAAAACCTTG | NO_CURRENT_992 | NO_CURRENT_992 |
| TTGTAATAGAAGAAATTACG | NO_CURRENT_993 | NO_CURRENT_993 |
| TTGTAGCGAGTCGAAAACTG | NOS2 | 4843 |
| TTGTCAATGGGATATATGAG | NO_CURRENT_994 | NO_CURRENT_994 |
| TTGTCCCTGAGAAAACGCGG | NO_CURRENT_995 | NO_CURRENT_995 |
| TTGTCCGTGACCCTGATTAA | NO_CURRENT_996 | NO_CURRENT_996 |
| TTGTCGCCCATGGTCCGCGA | ARRB1 | 408 |
| TTGTCTGCCGGACTGACGGG | MAP2K7 | 5609 |
| TTGTCTGTGTGTGGGACTGA | BAP1 | 8314 |
| TTGTCTTCTTTCTCAGTGCA | MMP1 | 4312 |
| TTGTGAATTTGGTTACCATG | NO_CURRENT_997 | NO_CURRENT_997 |
| TTGTGAGAGCCGCAAGGGCA | CYP1B1 | 1545 |
| TTGTGGGATGTGCGGCTACT | SETD1B | 23067 |
| TTTAAAAGGTGAAGCAGTGG | SUV39H1 | 6839 |
| TTTACCCTCTCTGGAGGTCG | NAIP | 4671 |
| TTTACGAAGTATACCAGGTC | NO_CURRENT_998 | NO_CURRENT_998 |
| TTTAGACAACTTCAGGATGG | NAIP | 4671 |
| TTTAGCCTTGAGCTCAACTG | CCL20 | 6364 |
| TTTAGGCATTGCGGCTCCGG | PDAP1 | 11333 |

| | | |
|---|---|---|
| TTTATTAGAATACTTGGTCT | NAIP | 4671 |
| TTTATTGGCTGTCTGACTGT | HTR3E | 285242 |
| TTTCAACCTTGTCAACCCGT | CS | 1431 |
| TTTCAGGGTCTAACCTTCTG | CD84 | 8832 |
| TTTCAGTCACACAAGAAGGG | CASP1 | 834 |
| TTTCATAAGAGTGGGACACG | NO_CURRENT_999 | NO_CURRENT_999 |
| TTTCCACCTTAGTGAGAACA | RTP1 | 132112 |
| TTTCCAGAACGCTCGGTGAG | DNAJA1 | 3301 |
| TTTCCCATGATCATTTAGTG | NO_CURRENT_1000 | NO_CURRENT_1000 |
| TTTCCCCCAGTTGCAACGTT | MAP2K6 | 5608 |
| TTTCCGAGCAGCCGTCCTGC | EIF4G3 | 8672 |
| TTTCGCCCAAGAGGCTTGGG | NO_CURRENT_1001 | NO_CURRENT_1001 |
| TTTCGCTGCGTCCCCTTCGC | TP73 | 7161 |
| TTTCGTGCCGATGTAACACA | NO_CURRENT_1002 | NO_CURRENT_1002 |
| TTTCTAGTTACTACTGGACG | NO_CURRENT_1003 | NO_CURRENT_1003 |
| TTTCTCACACTCTTACTCCA | C4orf17 | 84103 |
| TTTCTTAAAACTTAAACCGG | NO_CURRENT_1004 | NO_CURRENT_1004 |
| TTTGACCGAGGCATTAAGGC | IRAK4 | 51135 |
| TTTGCAAGTCACTATAATTG | NO_CURRENT_1005 | NO_CURRENT_1005 |
| TTTGCAGGTGGACCACCCAG | TMEM30A | 55754 |
| TTTGCCATACAAGCTAACAT | NO_CURRENT_1006 | NO_CURRENT_1006 |
| TTTGCGACCGCTCCTGGGCC | ASH2L | 9070 |
| TTTGCTGTGCAGAACACTGC | CAT | 847 |
| TTTGGAGTTGCTTGCGGCGG | TRERF1 | 55809 |
| TTTGGCAAGTTAATCTCTAT | NO_CURRENT_1007 | NO_CURRENT_1007 |
| TTTGGCCAAAGTTCCAGGTG | HTR1D | 3352 |
| TTTGGCCTCCAGTACATGAG | NO_CURRENT_1008 | NO_CURRENT_1008 |
| TTTGGTTAAAATATTCAAAA | POU2F1 | 5451 |
| TTTGGTTTCCTCTTTCCATG | OR7C2 | 26658 |
| TTTGTACCGGAGTCCCATTT | EDA2R | 60401 |
| TTTGTTTATGTTATGACGCA | NO_CURRENT_1009 | NO_CURRENT_1009 |
| TTTTACCTTGTTCACATGGA | NO_CURRENT_1010 | NO_CURRENT_1010 |
