## Supplementary material for "Illuminating host-mycobacterial interactions with functional genomic screening to inhibit mycobacterial pathogenesis": Table S6

**Table S6. Positive genetic hits from secondary CRISPR-Cas9 knockout screen in *M. bovis* BCG infection. Related to Figure 3.**

| Gene Symbol | Average LFC | Average -log (p-values) | Number of perturbations | Perturbations | Individual LFCs |
|---|---|---|---|---|---|
| IFNAR2 | 5.070 | 12.332 | 10 | ATTCCCATGATAAGATGGAC;CGTCATTGAAGAACAGTCAG;TGAAAGTGATAGCGATACTG;ATTTCCTAACCTGCCACCGT;GCAGAATCTGCCAAAATAGG;GAGGTCACAAAATGATCTTG;TGAGTGGAGAAGCACACACG;CATTGCTGTATACAATCATG;CCGTCCTAGAAGGATTCAGC;CCAACAATCTCAAACTCTGG | 4.61;4.79;4.95;5.1;5.11;5.16;5.19;5.2;5.27;5.32 |
| IFNAR1 | 4.964 | 11.782 | 10 | AACAGGAGCGATGAGTCTGT;TTTAGACAATAAGTAGTCTC;AAGCAGCACTACTTACGTCA;ATAATTGGATAAAATTGTCT;GAACAAAAGATAGTGTTATG;CAGATGCTTGTACGCGGAGA;TTGGAGCACCGATATAGATA;TCATTTACACCATTTCGCAA;GTACATTGTATAAAGACCAC;GATCTAATGTTAAAGACTGG | 4.17;4.57;4.69;4.78;5.05;5.17;5.21;5.23;5.26;5.52 |
| JAK1 | 4.893 | 11.167 | 10 | TGAGCCAGCTGAGTTTCGAT;CACACTTACTCTCCACGTCG;TGAGCTGGCATCAAGGAGAG;ATTGCGGCAAGAAGGAAGCG;GTTGTGGACGATCAACGGGG;TGAATAAGTCCATCAGACAG;GTCATCCTTGTAATCCATCA;CAAAGGACAAGGCCTCCTCG;CTTGGTGTTCTCGTCATACA;CATTCTCACCAAGTAGCTCA | 4.34;4.36;4.68;4.95;4.96;4.97;4.98;5.09;5.16;5.45 |
| STAT2 | 4.238 | 9.560 | 10 | ACCTACCCTTGTAATTGAGG;CAGGATGACCCTCTGACCAA;AGCAACATGAGATTGAATCC;GGGGCCACTAGGTGTGACAG;ATCATCTCAGCCAACTGGGT;ACTTGGGAGTGATGATTCCA;GCAAGTTCACCGTCCGAACA;GAGGCCTCGGCCAACATAGG;GTCCTGGGCTGACTTCACTA;TGGAAATAATCACCACAGGG | 2.0;3.89;4.06;4.11;4.38;4.57;4.6;4.78;4.89;5.13 |
| TYK2 | 3.979 | 9.234 | 10 | GGACGGCTATTTCCGCCTGA;CGGTACACAGCCGGTTCCCG;GGTCCCTGCCAGGCACGAGG;ACCTGCGGAAGACGTTCCGA;CTGCTACATCCGGGACAGTG;T | 2.0;3.21;3.55;4.07;4.2;4.26;4.42;4.61;4.68;4.8 |

| | | | | | |
|---|---|---|---|---|---|
| | | | | GAATGACGTGGCATCACTG;CTTCCGGGACATCACCCACG;TTGGGCCTGAGCATCGAAGA;CAGGCGGCCCTCATACACGT;AATACCTAGCCACACTCGAG | |
| STAT1 | 3.607 | 8.852 | 10 | TCCCATTACAGGCTCAGTCG;GAGGTCATGAAAACGGATGG;CAGTAAAGTCAGAAATGTGA;TGCAAAACCTTGCAGAACAG;TGTGATAGGGTCATGTTCGT;CCTGATTAATGATGAACTAG;TGGTTTGGTAATTGACCTCG;TGCTGGCACCAGAACGAATG;CATGTTGTACCAAAGGATGG;TATTTGCAGCTCGTTTGTGG | 2.4;2.97;2.99;3.01;3.45;3.96;4.03;4.16;4.52;4.58 |
| IRF9 | 3.098 | 8.497 | 10 | AACTGAGGCCCCCTTTCAAG;ATCTACAACGGGCGCGTGGT;ATACAGCTAAGACCATGTTC;ACAATTCCACAGGCCAGCCA;GGAGCAGTCCATTCAGACAT;AAGTATAAGGAGGGGGACAC;GGATGTTGCTGAGCCCTACA;AATTTAAGGAGGTTCCTGAG;ACTGAGTGTGCAGTTCTGCA;GAGCAGCAGTGAGTAGTCTG | 2.01;2.62;2.64;2.64;3.11;3.29;3.53;3.59;3.76;3.79 |
| TRERF1 | 2.253 | 6.985 | 10 | CGGGCATGACCATAGGCGTG;TGGTAGCGGGAAGGGATAGC;GCTGCTGGTAGTCATAATAC;GCAGCCTCCTTCTTATCACA;ATGGATACCTCGAGGCAGGG;GGATCCACCTCAACAACATG;CGGACCCAGAAGCTTACCAG;CTGCCCTGAATGGCCACATG;CCAACCCAAAGGAGCGTTTG;GCAACGTGACGGTCACCCCA | 0.15;0.45;1.84;2.24;2.79;2.83;2.88;2.97;3.1;3.29 |
| CDKN2C | 1.995 | 7.613 | 10 | TTGGAAATCCCGAGATTGCC;AGGGAACCTGCCCTTGCACT;AGTGTCCAGGAAACCTGCTC;GAATGACAGCGAAACCAGTT;TCAAGCTGATGTTAACATCG;GCAAAGTCTGTAAAGTGTCC;ATGCACAAAATGGATTTGGA;TTAACATCGAGGATAATGAA;ACAAACTAGTAAGTTGCTCT;CCAGTTCGGTCTTTCAAATC | 1.35;1.73;1.79;1.87;1.92;2.19;2.21;2.23;2.27;2.38 |
| DNAJB6 | 1.521 | 4.451 | 10 | AATGTGAAGCCAAATTCAAA;AGAAAAACGACCCCGTCCCT;TATGACAAATATGGCAAAGA;CTTTCCAAGATATCGGAAAC;GGACTTCTTTGGGAATCGAA;GCCACTACCACCAAATGACG;TTTGGGTCACTAGGTCACGG;GCATATGAAGTGCTGTCGGA;GCAGAAAAATCACTACAAAG;GAGAAAATTCAAGCAAGTAG | -1.1;-0.29;-0.07;0.06;0.39;2.38;2.64;3.72;3.72;3.73 |

| | | | | | |
|---|---|---|---|---|---|
| SUV39H1 | 1.249 | 6.336 | 10 | GGAAGATGCAGAGGTCATAT;GCCGGGGGGTCTTTGACCGG;TGTACACGAAGGCCCGCGGA;AAGTTTGCCTACAATGACCA;GGATCTTCTTGTAATCGCAC;CGAGTGCCAGGACTGTCTGT;ACACACTTGAGATTCTGCCG;AATGAGTACCGTGTTGGTGA;GTTCCTCTTAGAGATACCGA;CTATGACTGCCCAAATCGTG | 0.67;0.79;1.1;1.13;1.14;1.27;1.38;1.59;1.68;1.72 |
| TMEM30A | 1.182 | 4.631 | 10 | ACAAAAATGCCAATGCCGAT;ACTGGTTTAAGCCAGTTCAC;GACTCGGAGACCGGATAACA;GTAGCACCGTGCCAGCCGTA;GACAAACCAATTGCTCCTTG;AAAAGAAAGGTATTGCTTGG;GAGATTTCACGTAACGACGA;TAAAAACCAGAATCCCAGTA;TTGTACCATTAACTTCACAC;CATTTATTACAGGGACTGGA | -1.09;-0.74;0.07;1.12;1.62;1.75;1.93;1.95;2.6;2.6 |
| KHSRP | 1.071 | 5.906 | 10 | TTCCACGACAACGCCAACGG;AATGAGTACGGATCTCGGAT;CGTCGGCGAAAGCGTCCTTG;CGGATCTCAGAATACGAATG;AGTGCTGTTATTCACTGTCG;AGGACACACTGCGCTCGGGT;ACAGCAATCAGTTCTCAACT;TCTTGATCATCTCTCCACTC;GTGCGGATACAGTTCAAGCA;GACCAGCCCACACTTGTGAG | 0.23;0.72;0.9;0.94;1.11;1.14;1.25;1.29;1.5;1.63 |
| SLC7A5 | 0.970 | 5.564 | 10 | TTACTTGAATTTCGTCACAG;ACAGGGTGGTGAAGTAGGCC;GGCTTCGTCCAGATCGGGAA;GACTTCGGGAACTATCACCT;TGGATCATCCCCGTCTTCGT;ACGTACACCAGCGTCACGAT;GGATGTGGGGAACATTGTGC;CGAGCAGATGCTGTCGTCCG;CGTGAACTGCTACAGCGTGA;GTGCCTTCAAATGAGAAGTT | 0.1;0.32;0.75;0.93;0.95;1.01;1.1;1.41;1.41;1.72 |
| DNTTIP1 | 0.960 | 4.716 | 10 | TCGAGATCCCAACACAAAGG;TGTGAACCAATTCGCCGGGA;CATAATGATAAAGCACCGGC;GAAGAAGAATGTGCCCATCG;TAGATTCATTCAGGCGAGCT;GGTCCCTACTCACACCATGC;TTGTTGTCATCTGTGAGCGG;TCGCTTGACAGGATGTGTCC;GAACGTGCGAGACAATGTTG;CCTGCAGCCCAGCATCAACG | -0.71;-0.48;0.23;0.45;1.28;1.51;1.52;1.66;1.94;2.21 |
| HSPA13 | 0.945 | 5.133 | 10 | AAACAATAAACTTGGAGGAC;ATACCAATCACTTTAGGAGT;CCACAGATTGAGACAAGCTG;TCTTTACTGAATAAACAAGG;TGGTACAGAAATGACAGCAT;ACTGAC | -1.07;0.38;0.93;0.95;1.18;1.23; |

| | | | | | |
|---|---|---|---|---|---|
| | | | | AATGATGTATATGT;TCAGTAAAAGACACCATGCT;CCAAGTCTATCACCAAGACG;CACCCCAACAGAACAATAGG;TCGAGAGCCAACATATTCTG | 1.42;1.43;1.5;1.51 |
| HDAC2 | 0.918 | 5.628 | 10 | ATGAGTATATCAAATTTCTA;AAACCGACAACAGACTGATA;TTGTGTGTAGATTATCTCAA;ATTGATATAGATATTCATCA;TCCGTAATGTTGCTCGATGT;TCATTATCTGGTGATAGACT;CCTCCTCCAAGCATCAGTAA;GATGTATCAACCTAGTGCTG;TCAAAGAGTCCATCAAACAC;TACAACAGATCGTGTAATGA | -0.08;0.68;0.92;0.92;0.95;1.12;1.14;1.16;1.18;1.18 |
| FOSL2 | 0.878 | 3.998 | 10 | GAGCCACCCCCACTGCGCAT;AGTGTGCAAGATTAGCCCCG;AAATTCCGGGTAGATATGCC;CCTGGCGTGATCAAGACCAT;GACATGGAGGTGATCACTGT;GTGCGAGCGAGGGTATGGGT;GGAGAAGCTGGAGTTCATGT;AGGAGAAGCGTCGCATCCGG;GATCACGCCAGGTCTTGGGA;CTTTCCGTGGCAGGAGACAG | -2.86;-1.78;-0.06;1.55;1.56;1.74;1.82;2.16;2.23;2.42 |
| JUNB | 0.845 | 5.500 | 10 | CAAACTCCTGAAACCGAGCC;CAGGGCTTTGACAAAGCCGT;TCATACACAGCTACGGGATA;GGGTAAAAGTACTGTCCCGG;GCGCTTTGAGACTCCGGTAG;GTTTGTAGTCGTGTAGAGAG;CAGGCGTTCCAGCTCCGAAG;CTGAGGTTGGTGTAAACGGG;GTACGAGCTCCCGGTCCCGA;ACACCCCCCAACGTGTCCCT | 0.13;0.58;0.69;0.73;0.92;0.95;0.96;1.04;1.06;1.37 |
| UCHL5 | 0.786 | 4.511 | 10 | CAAAAAATATCGTGTCAAGT;TCAGTTATGTTCCTGTTAAT;AGAAAAATTAACCCATGAAC;AAGTACACAACAGTTTCGCC;AGAAGAACCAGCAGGCTCTG;ATGATTGGATCAGTGCAGTA;ACCGAATCCTTTAATGAGCT;GTTACTGAACTGTACCCACC;GAATTAGATGGATTAAGAGA;GTTCAGTAACACACTCACTA | -0.86;-0.79;0.77;0.83;1.02;1.16;1.22;1.42;1.47;1.61 |
| SEC62 | 0.763 | 5.139 | 10 | ACAGTTGAATCGAAGATACT;CTCATTTCTGCTGGCCAAAG;TTTGGATTCAAAGTGGGCAA;GCCGCTATTACTGCAATCAC;AGAAAAACCTGATCATCATG;AACAGCCTGCACCCACACTG;ATCATTTGGCTCATAACTGG;TACCAATAAAATAATCAACC;AAATAGAATGCATCGAGCTG;GGAAGCTTTATTTACAACCA | 0.13;0.14;0.21;0.57;0.73;0.83;1.19;1.22;1.24;1.36 |

| | | | | | |
|---|---|---|---|---|---|
| BMI1 | 0.760 | 4.850 | 10 | AACGTGTATTGTTCGTTACC;GTGGTCTGGTCTTGTGAACT;CTCCACAAAGCACACACATC;AAAGGATTATTATACACTAA;AGACAAAGAGAAATCTAAGG;GATTGATGTCATGTATGAGG;AAAGGTTTACCATCAGCAGA;TTGTATACAAATTAGTTCCA;GCATCACAGTCATTGCTGCT;AATGGCTCTAATGAAGATAG | -0.73;0.41;0.57;0.61;0.8;0.82;0.87;1.18;1.29;1.77 |
| COMMD3-BMI1 | 0.760 | 4.850 | 10 | AACGTGTATTGTTCGTTACC;GTGGTCTGGTCTTGTGAACT;CTCCACAAAGCACACACATC;AAAGGATTATTATACACTAA;AGACAAAGAGAAATCTAAGG;GATTGATGTCATGTATGAGG;AAAGGTTTACCATCAGCAGA;TTGTATACAAATTAGTTCCA;GCATCACAGTCATTGCTGCT;AATGGCTCTAATGAAGATAG | -0.73;0.41;0.57;0.61;0.8;0.82;0.87;1.18;1.29;1.77 |
| SLC3A2 | 0.590 | 4.072 | 10 | AGAACCACGAGTTCTCACCC;TGAATGCCACTGGCAATCGC;GTCTGATTCTGGTTCTACTG;CAGGCCCGTGAACTTAGCCG;CTTCCTTGGAGCCAAAATTG;TCCAACACTCACCTTCACCT;TCCAATTCACAAGAACCAGA;TGAGTGGCAAAATATCACCA;CGCTCGCACGATTATGACCA;GCGCAGAAGTGGTGGCACAC | -1.19;-0.21;0.1;0.2;0.6;1.02;1.07;1.14;1.39;1.77 |
| MSL2 | 0.549 | 4.883 | 10 | CTCAGTAACAGAGAGTCTCC;ATTGAAAGCCCATTATAAGT;TTGAAACAGTTTCTAAGCTG;CACCGCAAACACAGCACGAA;GCTACTGTGTCACAAACGGG;CCAATTTCTACCATTATCCG;AAAAGGTCCATTGGATACAG;ACTGCCAATAGCAATGCTGA;GTATATAACACAGACTACAC;TAACCCATGAAGTGCAGGAC | 0.18;0.23;0.28;0.37;0.4;0.49;0.68;0.69;0.89;1.29 |
