## Supplementary material for "Illuminating host-mycobacterial interactions with functional genomic screening to inhibit mycobacterial pathogenesis": Table S7

**Table S7. Positive genetic hits from secondary CRISPRi screen in *M. bovis* BCG infection. Related to Figure 3.**

| Gene Symbol | Average LFC | Average -log (p-values) | Number of perturbations | Perturbations | Individual LFCs |
|---|---|---|---|---|---|
| TYK2 | 2.916131534 | 11.99062628 | 10 | CGCGGAAGGAGCGCGGCCGG;CAGGAAGAAGCCGCGGGGAC;AGGTCCTCAGGAAGAAGCCG;GGTCCTCAGGAAGAAGCCGC;GTCCTCAGGAAGAAGCCGCG;GGGGACTGGCTGCGCTTGAC;TCAAGCGCAGCCAGTCCCCG;GGCTTCTTCCTGAGGACCTC;GTCCCCGCGGCTTCTTCCTG;GAGCGCGGCCGGAGGTCCTC | 1.3;2.56;2.96;3.02;3.04;3.09;3.18;3.19;3.36;3.46 |
| IFNAR1 | 2.150968318 | 9.060590547 | 10 | TGCGCGAACATGTAACTGGT;GTGCGCGAACATGTAACTGG;CGCGTGCGCGAACATGTAAC;GGGGCGGTGTGACTTAGGAC;GGGCGGTGTGACTTAGGACG;ACGGGGCGATGGCGGCTGAG;GACTTAGGACGGGGCGATGG;CATGTAACTGGTGGGATCTG;CAGATGATGGTCGTCCTCCT;GTAACTGGTGGGATCTGCGG | 0.36;0.9;1.06;1.26;1.47;2.5;3.32;3.38;3.43;3.84 |
| IRF9 | 1.941531649 | 6.147083593 | 10 | GTCTGAGCCCTGCGACAGCC;CCAAGGATGGGAGTCAGCCT;GCCAAGGATGGGAGTCAGCC;CAGGTAAGATCAGCCAAGGA;AACTCAGGTAAGATCAGCCA;CCCAGGCTGACTCCCATCCT;AGGTAAGATCAGCCAAGGAT;GACAGCCTGGACAGCAACTC;TCTTACCTGAGTTGCTGTCC;TTGCTGTCCAGGCTGTCGCA | -0.31;-0.21;0.05;1.85;2.09;2.93;3.14;3.16;3.31;3.42 |
| IFNAR2 | 1.937382184 | 8.500670649 | 10 | CGGCGGGATTCCTACCCGCC;CGGGGAACTGTTCGGCTCGC;TAAAGACGCTTCTTCCCGGC;CCGGCGGGATTCCTACCCGC;GGAACTGTTCGGCTCGCCGG;CCGGCGGGTAGGAATCCCGC;GGAAGAAGCGTCTTTAGTGC;GACGCTTCTTCCCGGCGGGT;GGGAAGAAGCGTCTTTAGTG;CTAAAGACGCTTCTTCCCGG | 0.07;1.0;1.23;1.99;2.01;2.19;2.46;2.48;2.92;3.02 |
| STAT2 | 1.404108072 | 6.255647125 | 10 | TCAGGGCCCGTACCTGATTA;CTCAGGGCCCGTACCTGATT;CTGCAACCCTAATCAGGTAC;CAGAGCCATTGGAGGGCGCG;ACTGCAACCCTAATCAGGTA;ATCAGGTACGGGCCCTGAGA;AATCAGGTACGGGCCCTG | -0.23;0.0;0.84;1.14;1.32;1.68;2 |

| | | | | | |
|---|---|---|---|---|---|
| | | | | AG;CGGGGACTGCAACCCTAATC;TACCCCAGCACACCCTCTCA;GGCCCTGAGAGGGTGTGCTG | .01;2.16;2.49;2.63 |
| STAG2 | 1.033339595 | 6.909632078 | 10 | CCATCATCTCCAGCGCGTGG;CATCATCTCCAGCGCGTGGT;ACGCGCTGGAGATGATGGGG;GCAGTTCCCCCACCACGCGC;ATCATCTCCAGCGCGTGGTG;CGCTGGAGATGATGGGGCGG;TCATCTCCAGCGCGTGGTGG;GCCCCATCATCTCCAGCGCG;GCGTGGTGGGGGAACTGCTG;CACCACGCGCTGGAGATGAT | 0.33;0.56;0.97;1.04;1.08;1.11;1.14;1.25;1.36;1.49 |
| NFRKB | 0.866522622 | 5.45936147 | 10 | GGTTAGGGGGCCGGCGCCCG;CGCGGCCGGAGAAGGGCTGC;CTCACCTGAACCGCGGGCGC;GCCCGCCCTCACCTGAACCG;GCCCGCGGTTCAGGTGAGGG;GGAGAAGGGCTGCGGGTTAG;CCCGCCCTCACCTGAACCGC;GAGAAGGGCTGCGGGTTAGG;GGCGCCCGCGGTTCAGGTGA;CGGCGCCCGCGGTTCAGGTG | -0.25;0.52;0.7;0.8;0.82;1.09;1.15;1.21;1.26;1.35 |
| DPY30 | 0.861234604 | 5.911668158 | 10 | CTAGAATAAGTGAGAAAGTG;CCTAGAATAAGTGAGAAAGT;GGAGCTGGTGGCGCGGTGCA;GCAGGGCTCTTAAGAACGAA;CGGCCGGAGCTGTTTGTGCT;AGAACGAACGGCTTGGGCGC;GCTCTTAAGAACGAACGGCT;TAGAATAAGTGAGAAAGTGG;CTCTTAAGAACGAACGGCTT;AAGAACGAACGGCTTGGGCG | -0.01;0.63;0.65;0.83;1.06;1.06;1.07;1.09;1.11;1.12 |
| MEMO1 | 0.861234604 | 5.911668158 | 10 | CTAGAATAAGTGAGAAAGTG;CCTAGAATAAGTGAGAAAGT;GGAGCTGGTGGCGCGGTGCA;GCAGGGCTCTTAAGAACGAA;CGGCCGGAGCTGTTTGTGCT;AGAACGAACGGCTTGGGCGC;GCTCTTAAGAACGAACGGCT;TAGAATAAGTGAGAAAGTGG;CTCTTAAGAACGAACGGCTT;AAGAACGAACGGCTTGGGCG | -0.01;0.63;0.65;0.83;1.06;1.06;1.07;1.09;1.11;1.12 |
| TRERF1 | 0.792164862 | 4.618105349 | 10 | TTGGAGTTGCTTGCGGCGGA;TTTGGAGTTGCTTGCGGCGG;AGTTGCTTGCGGCGGAGGGC;GGAGAGTTTGGAGTTGCTTG;GAGTTTGGAGTTGCTTGCGG;GGAGGGCAGGAAGGGGAGCA;CTTGCGGCGGAGGGCAGGAA;GCTTGCGGCGGAGGGCAGGA;ACGGCTGGATGCGGAGAGTT;TTGCGGCGGAGGGCAGGAAG | -0.24;-0.09;0.48;0.85;0.85;0.9;0.94;1.3;1.3;1.64 |

| | | | | | |
|---|---|---|---|---|---|
| UCHL5 | 0.771126276 | 4.904484138 | 10 | ATCGGGGCGGTGTGTGGCCA;AGGTGGATCGGGGCGGTGTG;GCTGAGAGCGAGAGGTGGAT;GAGCGAGAGGTGGATCGGGG;CTGAGAGCGAGAGGTGGATC;GGGGGGCAGCTGAGAGCGAG;GGCCGGCGGCAGCTGGTGCG;GGGCAGCTGAGAGCGAGAGG;TGAGAGCGAGAGGTGGATCG;CCGGCGGCAGCTGGTGCGGG | -0.26;-0.12;0.86;0.89;0.92;0.92;1.05;1.11;1.11;1.22 |
| PHIP | 0.73087687 | 5.329027453 | 10 | GAACACACACTGACAGCTAT;ACTGACAGCTATAGGGCAGG;CACACTGACAGCTATAGGGC;CGGCTCCACCATTCAAGCAA;CTCCACCATTCAAGCAACGG;CGCCGCCGTTGCTTGAATGG;CTCCGCCGCCGTTGCTTGAA;TGAATGGTGGAGCCGAAGCT;AACACACACTGACAGCTATA;TGTGTTCACGAGCCGAGCTT | 0.19;0.49;0.55;0.66;0.75;0.75;0.87;0.96;0.98;1.1 |
| TP73 | 0.718805836 | 5.090993246 | 10 | GGGCGCGCGAGCCTCCGGGC;GAGGCTCGCGCGCCCGCGAA;AAGGGGACGCAGCGAAACCG;GTTTCGCTGCGTCCCCTTCG;TTTCGCTGCGTCCCCTTCGC;GAAGGGGACGCAGCGAAACC;CGAAGGGGACGCAGCGAAAC;CGGGCGCGCGAGCCTCCGGG;AGGCTCGCGCGCCCGCGAAG;CCGCCCGCGCACCCGCCCGG | -0.56;0.74;0.77;0.78;0.82;0.85;0.87;0.95;0.96;1.0 |
| MCTS1 | 0.718234138 | 4.133202073 | 10 | GAAAGAAATTGAATTTGCAG;GGAAAGAAATTGAATTTGCA;GCCGGCCTTCCTCGTGTGAG;CCGGCAGATCCCCTCACACG;CAGATCCCCTCACACGAGGA;CCTCGTGTGAGGGGATCTGC;ACAGCTCCGACTGTCAGTGC;GGAAGGCCGGCACTGACAGT;GTGCCGGCCTTCCTCGTGTG;TCCCCTCACACGAGGAAGGC | -0.24;-0.23;-0.15;0.98;1.06;1.08;1.08;1.11;1.16;1.33 |
| CSDE1 | 0.666374814 | 4.544453688 | 10 | CGTGCTGCTTATGGCGGCGC;CTTATGGCGGCGCTGGAGAG;GTTGGGGTGAGTACGACCTC;GCTTATGGCGGCGCTGGAGA;TTGGGGTGAGTACGACCTCA;AGAGGGGGCGCTGAGCTGTT;CTACCCCCAGTTGGACCCTG;TGCTTATGGCGGCGCTGGAG;GAGGGGGCGCTGAGCTGTTG;TTATGGCGGCGCTGGAGAGG | -0.14;0.08;0.55;0.7;0.77;0.79;0.79;0.92;1.1;1.1 |

| | | | | | |
|---|---|---|---|---|---|
| AHR | 0.640583635 | 5.025517112 | 10 | CTGCCCCTGTTCTGAGCTGG;CCTGTTCTGAGCTGGCGGTC;TCTGTTCCGAGAGCGTGCCC;CTGTTCTGAGCTGGCGGTCC;TGTTCTGAGCTGGCGGTCCG;CCGGACCGCCAGCTCAGAAC;CGGCTGCCCCTGTTCTGAGC;GGCGGTCCGGGGCACGCTCT;CGGACCGCCAGCTCAGAACA;GGACCGCCAGCTCAGAACAG | 0.3;0.4;0.5;0.61;0.66;0.67;0.69;0.79;0.8;0.98 |
| CHMP5 | 0.636929796 | 3.919781449 | 10 | CTAGTGTTTGGGTTTCTTCG;AGGCTGGGCGGCGGAGCCTT;CAGGCTGGGCGGCGGAGCCT;TCAAGATGAACCGACTCTTC;ACTGACTGCATTGGCACGGT;CTTCGGGAAAGCGAAACCCA;CTCAAGATGAACCGACTCTT;GTTTCGCTTTCCCGAAGAGT;CCTGACTGACTGCATTGGCA;CAGTCAGTCAGGCTGGGCGG | -0.53;0.02;0.04;0.75;0.76;0.89;0.9;1.09;1.18;1.26 |
| NUFIP2 | 0.623269004 | 3.875077845 | 10 | AGCAGCCCGCAGCCTCAGCC;CTTTCAATGGAGGAGAAGCC;CGGCTGGGACTCCCTGGCTG;GTCCCAGCCGCTTTCAATGG;CTCCTCCATTGAAAGCGGCT;GGCTTCTCCTCCATTGAAAG;GGACTCCCTGGCTGAGGCTG;GCAGCCCGCAGCCTCAGCCA;GACTCCCTGGCTGAGGCTGC;GGAGTCCCAGCCGCTTTCAA | -0.2;-0.19;0.4;0.4;0.74;0.8;0.91;0.99;1.11;1.28 |
| CDKN2C | 0.617550374 | 4.696379784 | 10 | TCCGATGCCATCATGCAGCC;CTGGTTAGGAGCAAAGGAAA;CCTTTCCTTTGCTCCTAACC;TGGTTAGGAGCAAAGGAAAG;TGCCATCATGCAGCCTGGTT;TGCAGCCTGGTTAGGAGCAA;CCTGGTTAGGAGCAAAGGAA;TGGCATCGGAGACTGACAGA;ACCAGGCTGCATGATGGCAT;CTCCTAACCAGGCTGCATGA | 0.28;0.36;0.37;0.38;0.45;0.67;0.72;0.81;1.0;1.13 |
| PHF6 | 0.577967211 | 4.52316274 | 10 | TCCTTTCATTCGAGGAGAGA;GAATGAAAGGAAACAACCTC;AGGTTGTTTCCTTTCATTCG;TCCAGCAGTGCCTGAGAGCG;GAGGGTAAAGAGAAAGAAGT;AAAGAGAAAGAAGTCGGTCT;CTCCAGCAGTGCCTGAGAGC;GAGCGGGGCTCTGTCGCCGG;TGAGAGCGGGGCTCTGTCGC;GGCGACAGAGCCCCGCTCTC | -0.13;0.52;0.57;0.58;0.59;0.6;0.66;0.76;0.8;0.81 |
| PAXIP1 | 0.546500933 | 3.86607805 | 10 | CCTCAGGAACTTTGGGCGCC;GAACATCTCCTCAGGAACTT;GAGGTCAAGTATTACGCGGT;GGAGGTCAAGTATTACGCGG;CCAGGCGCCCAAAGTTCCTG;AAG | -0.2;0.25;0.3;0.46;0.52;0.53;0. |

| | | | | | |
|---|---|---|---|---|---|
| | | | | TTCCTGAGGAGATGTTC;CAGGGAGGTCAAGTATTACG;TCCTGAGGAGATGTTCAGGG;AGTTCCTGAGGAGATGTTCA;ACCTCCCTGAACATCTCCTC | 54;0.86;1.03;1.17 |
| GNAI2 | 0.538059944 | 4.478324987 | 10 | TCCCGCAGAGGGCTGGTGGT;GTGGTGGGAGCGGAGTGGGT;GAGTGCTTCCCGCAGAGGGC;TGCTTCCCGCAGAGGGCTGG;AGCCCTCTGCGGGAAGCACT;ACCCGAGTGCTTCCCGCAGA;TCCCACCACCAGCCCTCTGC;CTCCCACCACCAGCCCTCTG;GACCCGAGTGCTTCCCGCAG;GCCCTCTGCGGGAAGCACTC | 0.28;0.31;0.35;0.47;0.47;0.58;0.58;0.65;0.73;0.95 |
| KDM4A | 0.528006394 | 3.512696833 | 10 | CCAAAGCCCCGCCAGGGCTT;GCTGAGCCTAAGCCCTGGCG;GATCGGCCAGTGGCGACAGC;TCAGCTCCTGCTGTCGCCAC;GAGCTGAGCCTAAGCCCTGG;AAAGCAGTAGGATCGGCCAG;TACAGCCCAAAGCCCCGCCA;CAGGAGCTGAGCCTAAGCCC;TCGCCTGCTGGAAAAGCAGT;TGCTGGAAAAGCAGTAGGAT | -0.3;0.09;0.17;0.39;0.57;0.57;0.67;0.73;1.16;1.23 |
| CTNNBL1 | 0.524216678 | 3.42475968 | 10 | AGTGGAGTATTTGCTGGGCC;TGGGCCGGGTACCATGGACG;AAGTGGAGTATTTGCTGGGC;ATTTGCTGGGCCGGGTACCA;CAGGGAAGTGGAGTATTTGC;GCGGTCTGGGCGTGAGTGCA;TCGCCCACGTCCATGGTACC;CAGAAGTTCGCCCACGTCCA;AGGGAAGTGGAGTATTTGCT;GGGCCGGGTACCATGGACGT | -0.18;-0.15;0.05;0.27;0.7;0.75;0.86;0.88;0.91;1.16 |
| INO80E | 0.522163712 | 3.207066087 | 10 | CGGCCCGTTCATGACCGGCC;GCCGGTCATGAACGGGCCGG;TCTGGGCGCCACTCCCGGGC;CGGGCCGGTCATGAACGGGC;GGTAGCGGGAGGGCAGACTC;CTCCCGGGCCGGTCATGAAC;CCACTGCTTGGGGTAGCGGG;ACTCCCGGGCCGGTCATGAA;TCCGCCGGCCCGTTCATGAC;CAGACTCTGGGCGCCACTCC | -0.34;-0.22;-0.16;0.74;0.74;0.76;0.83;0.9;0.94;1.05 |
| INO80B | 0.516725374 | 3.664404896 | 10 | TGGGAGCACCTCTGGGGCTA;AGGGGCCTCCATAGCCCCAG;ACCTCATGAGTAAGCTGTGG;GCGGCGTGGGAGCACCTCTG;TGAGTAAGCTGTGGCGGCGT;GGCGGCGTGGGAGCACCTCT;AGGACCTCATGAGTAAGCTG;GAGCACCTCTGGGGCTATGG;ATGAGTAAGCTGTGGCGGCG;GCCGCCACAGCTTACTCATG | -0.35;0.07;0.28;0.62;0.65;0.65;0.66;0.77;0.85;0.97 |

| | | | | | |
|---|---|---|---|---|---|
| HELZ2 | 0.487558069 | 4.204912615 | 10 | TGAGCGCGGGGGTCCCAGGG;GCTCCAGGAGGGTGAGCGCG;CAGCTCCAGGAGGGTGAGCG;AGCTCCAGGAGGGTGAGCGC;GCAGGGCAGGCAGCTCCAGG;GCTCCTGGGCAGGCTCGGCA;TGAGAGCTCCTGGGCAGGCT;CTCCAGGAGGGTGAGCGCGG;CAGGGCAGGCAGCTCCAGGA;GGGTGAGCGCGGGGGTCCCA | 0.1;0.23;0.37;0.47;0.52;0.56;0.59;0.59;0.72;0.72 |
| SERF2 | 0.467757624 | 3.655469577 | 10 | ACGGGTCATGGCGACGGCAG;AGGGCAAGGAGAAGGGGGCT;AGAAGGGGGCTCGGTGCTCA;AAAGAAAAGGGCAAGGAGAA;GGGACGTAACGGAGGCAGGT;AAGAAAAGGGCAAGGAGAAG;GGCTCGGTGCTCACGGGTCA;GTGCTCACGGGTCATGGCGA;AGAAAAGGGCAAGGAGAAGG;GAAGGGGGCTCGGTGCTCAC | -1.43;-0.04;0.34;0.6;0.72;0.73;0.76;0.85;0.9;1.25 |
| JUNB | 0.464776425 | 3.070606582 | 10 | CCAGCTCCCTGCTGGCTCCG;GAGAGCGGCCAGGCCAGCCT;GAGCTGGGGGAAACGACGCC;CTCTGGCGCGATAGCTTTCC;CAGCAGGGAGCTGGGAGCTG;CTCCCTGCTGGCTCCGAGGC;CTCGGAGCCAGCAGGGAGCT;AGGCCAGCCTCGGAGCCAGC;GGCCAGCCTCGGAGCCAGCA;AGCAGGGAGCTGGGAGCTGG | -0.22;-0.04;-0.02;0.4;0.41;0.43;0.62;0.71;0.74;1.61 |
| ZRSR2 | 0.451225445 | 3.533592854 | 10 | CGGCGCTTACCTTGGTTTCT;TCGGGAAACGTCATCTTCTC;CTCGGGAAACGTCATCTTCT;GGCGCTTACCTTGGTTTCTC;TGACGTTTCCCGAGAAACCA;AAACCAAGGTAAGCGCCGTA;ACCAAGGTAAGCGCCGTACG;TCCCCGTACGGCGCTTACCT;CGCCTGCTCATCTCCCCGTA;AACCAAGGTAAGCGCCGTAC | -0.08;0.1;0.19;0.23;0.28;0.53;0.69;0.79;0.88;0.91 |
| UBA7 | 0.447719575 | 3.331765644 | 10 | CTGTTTGCACTGGCTACAGC;GTCCCTTATTCGCTTGGCCT;TGTTTGCACTGGCTACAGCA;CACTGGCTACAGCAGGGCAC;GAACCAAGGCCAAGCGAATA;GCCAGTGCAAACAGGAACCA;AACCAAGGCCAAGCGAATAA;GCCTTGGTTCCTGTTTGCAC;CTGCTGTAGCCAGTGCAAAC;AGCAGCGTCCCTTATTCGCT | -0.43;-0.06;0.35;0.52;0.58;0.59;0.65;0.68;0.8;0.81 |
| HSH2D | 0.427474271 | 3.793347892 | 10 | GGGAGACCCTTGGATGGCCG;TCCTCGAGGGCTGAGGCTCT;GCGAGGGCCTCGGCCATCCA;CCTCGAGG | 0.03;0.07;0.35;0.4;0.44;0.51;0 |

| | | | | | |
|---|---|---|---|---|---|
| | | | | GCTGAGGCTCTT;CGGCCATCCAAGGGTCTCCC;AGGCTCTTGGGTCACATACC;CGAGGGCCTCGGCCATCCAA;ATACCTGGGAGACCCTTGGA;GGCTCTTGGGTCACATACCT;TCACATACCTGGGAGACCCT | .53;0.6;0.63;0.72 |
| FBXO38 | 0.420223749 | 4.058511469 | 10 | CTCTCAACCACAATAACAGG;CACAATAACAGGCGGAGGGT;TAGGTGAGGCGGCGGCTGAG;AGGTGAGGCGGCGGCTGAGT;CGACCCTCCGCCTGTTATTG;CAACCACAATAACAGGCGGA;TCCCTCTCAACCACAATAAC;GCGGAGGGTCGGCGTAGGTG;AACAGGCGGAGGGTCGGCGT;TCAACCACAATAACAGGCGG | 0.18;0.27;0.35;0.38;0.4;0.41;0.46;0.46;0.6;0.69 |
| UBE2H | 0.416355178 | 3.899435832 | 10 | GCTGCTAGGAGCCGGTCAGA;TAGTGGAGGCTGCTAGGAGC;ACGCTCCCTCCTGCCTGCTG;GAGCCGGTCAGAGGGTGAGT;GGAGCCGGTCAGAGGGTGAG;GCAGGCGAGCGAGCCACAGC;GGCTGCTAGGAGCCGGTCAG;CGCTTATAGTGGAGGCTGCT;AGCCGGTCAGAGGGTGAGTG;CTCCCCACTCACCCTCTGAC | -0.03;0.29;0.34;0.42;0.43;0.49;0.52;0.54;0.56;0.6 |
| WDR26 | 0.41405547 | 3.106938813 | 10 | AGCAGAGCGGAGCGGATCCG;CAACACACCAGGGAGCAGAG;CGGAGCGGATCCGAGGACAG;GCAGCTGCCGCCTCTGTCCT;CACCAGGGAGCAGAGCGGAG;ATCCGCTCCGCTCTGCTCCC;GGGGAAGAATCAACACACCA;GGGGGAAGAATCAACACACC;GACTCCATTAGGCAGCAGCT;AGCGGATCCGAGGACAGAGG | -0.24;-0.14;0.15;0.41;0.47;0.54;0.65;0.73;0.78;0.8 |
| BABAM2 | 0.40744954 | 3.676374989 | 10 | GCGCAGGTCGGTGCGTCTGT;CGCAGGTCGGTGCGTCTGTC;ATCGGTAACCGGCGACTACG;CAGGTCGGTGCGTCTGTCGG;GCAGGTCGGTGCGTCTGTCG;TCGGTAACCGGCGACTACGT;CGTCTGTCGGGGGCGCGCTC;CCTGTACCCCACGTAGTCGC;CCGGCGACTACGTGGGGTAC;CGGTAACCGGCGACTACGTG | -0.05;0.22;0.26;0.39;0.4;0.41;0.42;0.54;0.6;0.88 |
| ACTR5 | 0.387510737 | 3.483929315 | 10 | CGGCACGGGCGTCGCGGAAC;CTCCAGCACTGGGTCCGGTG;CCGGCCTCCAGCACTGGGTC;CACTGGGTCCGGTGCGGCAC;CGCACCGGACCCAGTGCTGG;GCGGCACGGGCGTCGCGGAA;CCGCGACGCCCGTG | -0.02;0.03;0.27;0.31;0.32;0.49;0.52;0.62;0.66;0.68 |

| | | | | | |
|---|---|---|---|---|---|
| | | | | CCGCAC;CCGGTGCGGCACGGGCGTCG;GCACTGGGTCCGGTGCGGCA;TGCCGCACCGGACCCAGTGC | |
| PHC3 | 0.372628501 | 3.674931384 | 10 | ACTGATGTCGTAACTAGGGG;GTGACTGATGTCGTAACTAG;GACTGATGTCGTAACTAGGG;CGTAACTAGGGGGGGCGAGT;GTAACTAGGGGGGCGAGTT;AAGCGGAGTGAGTGACTAGA;GCAGCCCATGTTAGTGATGG;TGACTGATGTCGTAACTAGG;TGGTGACTGATGTCGTAACT;GGTGACTGATGTCGTAACTA | -0.06;0.28;0.33;0.35;0.36;0.4;0.43;0.49;0.51;0.64 |
| PQBP1 | 0.370936905 | 2.301584752 | 10 | CGTGTCGCCGCTACGATACC;GTGTCGCCGCTACGATACCT;GAAGGTCTCATTCGGTGTTT;TCCAGTTACTTTCAGGCTCG;TCCCCGAGCCTGAAAGTAAC;AATGAGACCTTCTCTCAACG;GTGAAGGCCTCGTTGAGAGA;CGTTGAGAGAAGGTCTCATT;AGTCCAGTTACTTTCAGGCT;TGTGGGCCCAGGTATCGTAG | -0.31;-0.27;-0.2;-0.1;0.52;0.66;0.76;0.82;0.87;0.96 |
| UTY | 0.368871889 | 3.305519778 | 10 | TTACAACTTTGTGCTGGTGC;ACACAATTACAACTTTGTGC;ACAAAGTTGTAATTGTGTCA;CAAAGTTGTAATTGTGTCAC;GGGAGGTCGCGGCGGCTGCG;AAGGCAGAAGCTCGCAGTGC;TGGGAGCGGGGAGGTCGCGG;GAGGTCGCGGCGGCTGCGTG;GGAGGTCGCGGCGGCTGCGT;GGCTGCAAGGAAAAAAGCTG | -0.02;0.04;0.14;0.16;0.38;0.49;0.52;0.53;0.64;0.82 |
| SNRNP70 | 0.359433049 | 2.07297596 | 10 | GCCCGCTAAGCGAGCGCACC;GCGACGGAATCAGACGGACG;GGCCTGGTGCGCTCGCTTAG;CGGGTGGCTGAGCAGCGGCC;GCCTGGTGCGCTCGCTTAGC;TGCGCTCGCTTAGCGGGCGA;TAGCGGGCGACGGAATCAGA;AGTCGCTATCGGAGGCCGCG;CCTGCTCCAGTCGCTATCGG;CCGCGCGGGTGGCTGAGCAG | -0.24;-0.22;-0.09;-0.08;0.12;0.13;0.47;0.69;1.0;1.79 |
| KPNB1 | 0.357110471 | 3.14358866 | 10 | CCTCCCTCCAAATGGGCTGC;TCCAAATGGGCTGCTGGCGG;CAAATGGGCTGCTGGCGGCG;CCCTCCAAATGGGCTGCTGG;GGGCTGCTGGCGGCGGGGGA;GCGGCGGGGGAGGGACCCTG;CGGCGGGGGAGGGACCCTGG;CGCGCTCTGAGCTGCCCCCA;GCTCAGAGCGCGCTGGTGCT;GGGGGCAGCTCAGAGCGCGC | -0.15;0.07;0.16;0.21;0.29;0.47;0.51;0.53;0.67;0.81 |

| | | | | | |
|---|---|---|---|---|---|
| STRAP | 0.331617122 | 2.66392643 | 10 | AAGTAGGGGATGGGGAGAAG;GCTGTGAATCGTGGCTGGCC;TGGACGCTGTGAATCGTGGC;GGTGTGGACGCTGTGAATCG;ACGATTCACAGCGTCCACAC;GAGGGAGGAAAGTAGGGGAT;GATGGGGAGAAGCGGAGAAC;AGGGAGGAAAGTAGGGGATG;ATGGGGAGAAGCGGAGAACC;GGAGGGAGGAAAGTAGGGGA | -0.41;-0.07;0.11;0.16;0.36;0.5;0.57;0.58;0.59;0.93 |
| CREBBP | 0.331183119 | 2.939553749 | 10 | GCGGTTGTGGGGCCCGGGAC;CGCGATCTACTCGGCCCCGC;GGGGTGCGCGGGCGGTTGTG;TACTCGGCCCCGCCGGTCCC;CGGGGTGCGCGGGCGGTTGT;GTTGTGGGGCCCGGGACCGG;GTAGATCGCGCTCGAAGCCC;TCGCGCTCGAAGCCCCGGTC;ATCGCGCTCGAAGCCCCGGT;GGCTTCGAGCGCGATCTACT | -0.21;-0.05;0.14;0.33;0.4;0.4;0.4;0.4;0.72;0.8 |
| NUP153 | 0.320933126 | 2.206457963 | 10 | GCGCTGTGCGCACAGGGGAG;ACGACGACGACGACGGCGGC;GGGGCGGGCGCTGTGCGCAC;GGGCGGGCGCTGTGCGCACA;GGCGCTGTGCGCACAGGGGA;CGACGACGACGACGGCGGCG;GACGACGACGACGACGGCGG;AGGGACGACGACGACGACGG;TGCAGGGACGACGACGACGA;GGCGGGCGCTGTGCGCACAG | -0.77;-0.48;-0.1;0.06;0.25;0.64;0.75;0.84;0.97;1.04 |
| ANP32B | 0.306028928 | 3.123885917 | 10 | CTGGGTGGCGGGCGGCGTGC;CCCGGAGCCAAGTTACTCAC;GTGGCGGGCGGCGTGCGGGC;CCCGTGAGTAACTTGGCTCC;AGCCAAGTTACTCACGGGAG;CCCCGGAGCCAAGTTACTCA;CCGTGAGTAACTTGGCTCCG;TCCCGTGAGTAACTTGGCTC;CGTGAGTAACTTGGCTCCGG;CTCCGCTCCCGTGAGTAACT | -0.09;0.07;0.14;0.23;0.25;0.4;0.43;0.46;0.49;0.67 |
| ARNT | 0.305271111 | 2.466940518 | 10 | GGTGGCATCTGCGGCCATGG;CTCTAAAGGGGAGGGGCCGA;GCGGCGACTACTGCCAACCC;TGGGGCGCCTCTAAAGGGGA;TGCCAACCCCGGTGAGGAGA;GGCGGTGGCATCTGCGGCCA;GGACTGGGGCGCCTCTAAAG;GGGGCGCCTCTAAAGGGGAG;GACTACTGCCAACCCCGGTG;GTTGGCAGTAGTCGCCGCCA | -0.33;-0.16;-0.07;0.3;0.31;0.37;0.59;0.6;0.66;0.77 |
| CHEK2 | 0.302448913 | 3.585152345 | 10 | GTCATATGGGGAACTTCTGT;GGAGAGTGTGCGGCTCCAGG;GCTCACGCGGTGAGTCATAT;CGGCTCCAG | 0.13;0.21;0.23;0.26;0.32;0.34; |

| | | | | | |
|---|---|---|---|---|---|
| | | | | GTGGGCTCACG;GGCTCACGCGGTGAGTCATA;GAGAGTGTGCGGCTCCAGGT;TCATATGGGGAACTTCTGTT;CTGAGGCTGCGGAGAGTGTG;TGCGGAGAGTGTGCGGCTCC;ACTCACCGCGTGAGCCCACC | 0.34;0.36;0.41;0.42 |
| NCOA6 | 0.298334173 | 2.747166701 | 10 | TCTGTCCCGCCGCCCGCGCC;CCGGCTCCGTGAGGCCCTGC;TCCGTGAGGCCCTGCCGGGT;CGGCTCCGTGAGGCCCTGCC;GGCCCTGCCGGGTCGGGCTG;CCGTGAGGCCCTGCCGGGTC;CGCCCGCAGCCCGACCCGGC;GCCCGCAGCCCGACCCGGCA;CTGCCGGGTCGGGCTGCGGG;GCGCGGGCGGCGGGACAGAC | -0.11;-0.07;0.06;0.06;0.07;0.46;0.59;0.62;0.65;0.66 |
| FAM214B | 0.282141516 | 3.036539491 | 10 | CGGAACTACAGGTGTGGGGC;ACCCACGGAACTACAGGTGT;GGAACTACAGGTGTGGGGCA;GCCCCACACCTGTAGTTCCG;AACGGACCCACGGAACTAC;TGAGCGTGGAAGAAGGGACG;CCCCACACCTGTAGTTCCGT;GACCCACGGAACTACAGGTG;CCCACGGAACTACAGGTGTG;CCCAAGTGCAACGGGACCCA | 0.07;0.08;0.11;0.12;0.15;0.16;0.32;0.38;0.53;0.9 |
| SUV39H1 | 0.275015952 | 2.349161492 | 10 | AGGCAGAGGCGCGGCCCGC;AAATTTAAAAGGTGAAGCAG;GCCCGAATGTCGTTAGCCGT;AAGCAGTGGAGGCAGAGGCG;CCCGAATGTCGTTAGCCGTG;AAGATGGCGGAAAATTTAAA;AGGTGAAGCAGTGGAGGCAG;AGCAGTGGAGGCAGAGGCGC;TTTAAAAGGTGAAGCAGTGG;TTAGCCGTGGGGAAAGATGG | -0.22;-0.12;-0.11;0.13;0.26;0.28;0.32;0.65;0.75;0.82 |
| CHMP6 | 0.274190731 | 1.81849631 | 10 | ACTTGGTGGGTCCCGGGCCA;TCTGCTTCTTGCGGCCGAAC;CTTGGTGGGTCCCGGGCCAG;GCCAGGGGCGGGCGCCGCCA;TGACGCGGCTCTGCTTCTTG;GCCGCCATGGGTAACCTGTT;CCCATGGCGGCGCCCGCCCC;CCAGGGGCGGGCGCCGCCAT;GCGGCCGAACAGGTTACCCA;GCCGAACAGGTTACCCATGG | -0.45;-0.19;-0.06;0.02;0.12;0.12;0.15;0.54;0.65;1.83 |
| PRR14L | 0.272483631 | 2.838552686 | 10 | GCCAGGTCTACCTGAGCCTA;CAGGTAGACCTGGCGACGAC;GGAGCGCCCCCGGCCTCTGT;AGGTAGACCTGGCGACGACG;TGAGCCTACGGAGCCGACAG;CCGGCCTCTGTCGGCTCCGT;TCAGGTAGACCTGGC | -0.25;0.02;0.17;0.26;0.31;0.37;0.39;0.39;0.5;0.55 |

| | | | | | |
|---|---|---|---|---|---|
| | | | | GACGA;TCTGTCGGCTCCGTAGGCTC;ACGGAGCCGACAGAGGCCGG;TCCGTAGGCTCAGGTAGACC | |
| UBE2L6 | 0.248396701 | 2.536685643 | 10 | CGCATGCTCGCCATCATGTC;CACACTCGGTCCCGACATGA;TCGCATGCTCGCCATCATGT;GAGGGAGGAGGGAATGCGGG;GGCGAGCATGCGAGTGGTGA;CATGATGGCGAGCATGCGAG;ATAGGATGACCCCTTTCTCT;TGGTGAAGGTAACCGCGTAT;GAAAGGGGTCATCCTATACG;CGAGGGGAGCCAAGAGAAAG | -0.29;-0.02;0.1;0.24;0.24;0.25;0.27;0.44;0.61;0.65 |
| SCTR | 0.234709839 | 0.912156748 | 10 | GCGCGCGGCCACCTGGTCCG;GCCCCGGGACTGCTCCTCCT;GGTCCGAGGAGGAGCAGTCC;GGACTGCTCCTCCTCGGACC;CTGCTCCTCCTCGGACCAGG;CCCGACCTGCGGCGGGCCCC;CGCGGCCACCTGGTCCGAGG;GCTTAGCGCGCGCGGCCACC;GGCGGGCGGCTTAGCGCGCG;TCCGAGGAGGAGCAGTCCCG | -0.37;-0.34;-0.27;-0.25;-0.22;-0.04;-0.03;-0.03;0.15;3.74 |
| YPEL5 | 0.231830007 | 1.976787228 | 10 | CGCGGATCTCACCGCCGCTC;CCCGCGGGACTCACCCTGAG;GCGGATCTCACCGCCGCTCA;GCCGCTCAGGGTGAGTCCCG;CCCGAGTCCAGCCCCTCCCG;CTCAGGGTGAGTCCCGCGGG;CTGAGCGGCGGTGAGATCCG;CCGCTCAGGGTGAGTCCCGC;GTGAGATCCGCGGCTGCCAG;GCGGGACTCACCCTGAGCGG | -0.52;-0.19;-0.07;0.03;0.11;0.4;0.41;0.62;0.65;0.88 |
| IKZF5 | 0.217503915 | 2.260481643 | 10 | CGGCATTTAACAAGGCGGTG;GGCGACGGCATTTAACAAGG;TACTCCATGCAGTTGTCAGG;GAAGACGGCGGCGGCGGCGA;TGTGACGGTGACGAAGACGG;GGTGACGAAGACGGCGGCGG;GGCGGCGACGGCATTTAACA;GACGGTGACGAAGACGGCGG;CGGAGACACAACAAAGATGG;GACTGTGACGGTGACGAAGA | -0.31;-0.23;0.12;0.17;0.26;0.33;0.35;0.39;0.52;0.58 |
| KHSRP | 0.208509849 | 2.753385003 | 10 | TGAAGCTGAGGAGGCGGCGG;GCGGCGGCGGCGGCTCAACG;TGGGTTCCCAGAGTGCTCCG;TGGCGCGGAGGCTGAAGCTG;CGGCGGCGGCGGCTCAACGC;GGAGGCTGAAGCTGAGGAGG;GAACAAGGCCTCGCTCCACA;CGCGGAGGCTGAAGCTGAGG;GGCGGCTCAACGCGGGAACA;GGCTGAAGCTGAGGAGGCGG | -1.03;0.13;0.24;0.26;0.28;0.32;0.34;0.44;0.51;0.59 |

| | | | | | |
|---|---|---|---|---|---|
| MAP2K6 | 0.203042219 | 2.39175966 | 10 | GCCTTCCCTAACGTTGCAAC;GCACACCTTGCAAAACAGAC;GCTGTGCTGCAGCTACATGA;CTTCCCTAACGTTGCAACTG;TTTCCCCCAGTTGCAACGTT;TTCCCCCAGTTGCAACGTTA;TGCAGCTACATGATGGGCCA;CCTTCCCTAACGTTGCAACT;TTCCCTAACGTTGCAACTGG;CCCAGTTGCAACGTTAGGGA | -0.28;-0.0;0.08;0.13;0.17;0.18;0.34;0.35;0.5;0.57 |
