## Supplementary material for "Illuminating host-mycobacterial interactions with functional genomic screening to inhibit mycobacterial pathogenesis": Table S8

**Table S8. Oligonucleotides used for THP-1 gene knockout or knockdown cell construction. Related to Figure 1 and Figure 5.**

| | **THP-1 cell pathways** | **Targeting gene** | **sgRNA target sequence** |
|---|---|---|---|
| *M. bovis* BCG infection | Type I IFN signaling pathway | TYK2 | TCAAGCGCAGCCAGTCCCCG |
| | | JAK1 | GGACAGCCGGGACTGGGCGC |
| | | IFNAR1 | GTAACTGGTGGGATCTGCGG |
| | Phagosome function | YPEL5 | CCCGCGGGACTCACCCTGAG |
| | Apoptosis | WDR26 | CAACACACCAGGGAGCAGAG |
| | Eicosanoids metabolism | AHR | TCTGTTCCGAGAGCGTGCCC |
| | Chromatin modification | CREBBP | GTAGATCGCGCTCGAAGCCC |
| | Transcriptional regulation of TP53 | TP73 | AAGGGGACGCAGCGAAACCG |
| | Deubiquitination | UCHL5 | GAGCGAGAGGTGGATCGGGG |
| | Unclear functions | PHIP | TGAATGGTGGAGCCGAAGCT |
| | | TRERF1 | GGAGAGTTTGGAGTTGCTTG |
| Validation of Cas9 function | | AAVS1 | GGGGCCACTAGGGACAGGAT |
| Validation of dCas9 function | | INTS9 | GGCAGGTGGCGGAGATTGCAC |
| | | MCM2 | GGATCGTGGTACTGCTATGG |
| Non-targeting control | | NC1 | GGGCTGAACGCGTATTCGCG |
| | | NC13 | GCTAGTCTGCGTGACGCGTCT |
| | | NC80 | ACATGTGGCTCCGCCCACAG |
| | | NC135 | AGATCTGCTCCATGTCACCA |
